# Gene Expression Changes Occurring at Bolting Time are Associated with Leaf Senescence in Arabidopsis

**DOI:** 10.1101/2020.05.29.109306

**Authors:** Will E Hinckley, Judy A. Brusslan

## Abstract

In plants, the vegetative to reproductive phase transition (termed bolting in Arabidopsis) generally precedes age-dependent leaf senescence (LS). Many studies describe a temporal link between bolting time and LS, as plants that bolt early, senesce early, and plants that bolt late, senesce late. However, the molecular mechanisms underlying this relationship are unknown and are potentially agriculturally important, as they may allow for the development of crops that can overcome early LS caused by stress-related early phase transition. We hypothesized that gene expression changes associated with bolting time were regulating LS. We used a mutant that displays both early bolting and early LS as a model to test this hypothesis. An RNA-seq time series experiment was completed to compare the early bolting mutant to vegetative WT plants of the same age. This allowed us to identify bolting time-associated genes (BAGs) expressed in an older rosette leaf at the time of inflorescence emergence. The BAG list contains many well characterized LS regulators (*ORE1, WRKY45, NAP, WRKY28*), and GO analysis revealed enrichment for LS and LS-related processes. These bolting associated LS regulators likely contribute to the temporal coupling of bolting time to LS.

## Introduction

Leaf senescence (LS) is the sequential death of older leaves, one-by-one, as the plant matures, while whole plant senescence is the simultaneous death of all leaves at the end of the growing season in monocarpic species (Nooden et al., 1997). A visual hallmark of LS is leaf yellowing, caused by chlorophyll degradation (Ougham et al., 2008; Tamary et al., 2019). During these processes, nitrogen (most commonly in the forms of nitrate, asparagine, and glutamine) and other macromolecules are recycled from dying leaves (sources) and relocated to growing tissues (sinks), including the reproductive organs (Havé et al., 2017). A better understanding of the regulation of LS may have important agricultural implications on yield and nutrition content.

Endogenous signaling molecules that control LS have been well characterized. Ethylene, abscisic acid (ABA), jasmonic acid (JA), salicylic acid (SA), and reactive oxygen species (ROS) are known to promote both age dependent and/or dark induced LS (Jing et al., 2005; Khanna-Chopra, 2012; Yuehui et al., 2002; K. Zhang et al., 2013; Zhao et al., 2017). Many genetic and epigenetic regulators of LS have also been identified. (Nicole Ay, Bianka Janack, 2014; Woo et al., 2019). Tri-methylated Lysine 4 on Histone 3 (H3K4me3) is an active gene expression mark associated with increased expression of many LS related genes (Brusslan et al., 2015; P. Liu et al., 2019). Histone Deacetyltransferase 6 (HDA6) regulates both flowering time and LS (Keqiang Wu, Lin Zhang, Changhe Zhou, Chun-Wei Yu, 2008). POWERDRESS interacts with HDA9 to promote aging (X. Chen et al., 2016). The H3K27me3 demethylase REF6 was also shown to promote leaf senescence (X. Wang et al., 2019). H3K9 acetylation via HAC1 (HISTONE ACETYLTRANSFERASE-1) promotes LS, and the transcription factor (TF) ERF022, a potential HAC1 target, positively regulates LS (Hinckley et al., 2019). Other individual TFs have been shown to regulate LS, for example *NAP, WRKY53, WRKY75*, and *ORE1* are positive regulators and JUB1, WRKY54 and WRKY70 negatively regulate LS (Guo et al., 2017; Lei et al., 2020; Miao et al., 2004; Qiu et al., 2015; Zentgraf & Doll, 2019). There are multiple large TF families that are associated with age dependent and dark induced LS (*WRKY, NAC, ERF*) (Bakshi & Oelmüller, 2014; Jiang et al., 2017; H. J. Kim et al., 2016; Koyama et al., 2013; Koyama, 2014; Li et al., 2018). Furthermore, stress, defense, and LS signaling overlap and some TFs are known to bridge stress and LS signaling (*SAG113, NAP, WRKY53*) (Asad et al., 2019; Nir Sade, María del Mar Rubio-Wilhelmi, Kamolchanok Umnajkitikorn, 2018; Woo et al., 2013).

While many regulators have been identified that function at the onset of or during LS, less is known about developmentally early regulators of LS. Kim et al. recently uncovered the NAC troika, consisting of three Arabidopsis NAC TFs that act in young rosette leaves to prevent early LS (H. J. Kim et al., 2018). To our knowledge, this represents the earliest known regulation of LS in Arabidopsis. Still, greater knowledge of developmentally early regulators of LS is needed to better optimize its timing.

Preceding LS is the vegetative to reproductive phase transition, which in Arabidopsis is termed bolting or flowering: the development of the primary inflorescence that produces cauline leaves, inflorescence meristems, and floral meristems. Many different environmental and autonomous cues can induce flowering independently, however all pathways converge on conserved master flowering time regulators: FT, SOC1, and FLC (Mouradov et al., 2002; Song et al., 2018). Stress can also induce flowering (Kiyotoshi Takeno, 2016; Wada & Takeno, 2010). The function of stress signaling in both flowering time and LS may serve as a link that allows reproductive success during biotic or abiotic stress.

A classic study in Arabidopsis showed LS in the fifth and sixth leaves was similar in WT and two late flowering mutants, *co-2* and *fca*, suggesting that leaf age was an important determinant of LS, not the vegetative-reproductive phase transition. Furthermore, LS was unaffected in male sterile (*ms1-1*) mutants, indicating that seed development did not influence LS (Hensel et al., 1993).

Other studies in Arabidopsis, however; show a relationship between the timing of bolting and leaf and/or whole plant senescence. Two ecotypes, Ler and Cvi, showed a negative correlation between bolting and sixth leaf longevity; Ler bolted much later and displayed shorter leaf longevity than Cvi (Luquez VM, Sasal Y, Medrano M, Martín MI, Mujica M, 2006). A similar study using more conventional quantitative methods of LS looked at the relationship between the natural bolting times and LS phenotypes of multiple Arabidopsis ecotypes, and found the opposite result of Luquez et al. Bolting age positively correlated to the chlorophyll content of the five largest leaves (measured at 59 DAS) among eight ecotypes, with later bolting ecotypes having more chlorophyll, implying a delay in LS. In addition, these eight ecotypes revealed a negative correlation between bolting age and percent yellow leaves in the rosette; here later bolting ecotypes had a smaller percentage of senescent leaves (Balazadeh et al., 2008). These findings show that the earlier bolting ecotypes were further progressed into leaf and whole plant senescence than later bolting ecotypes, indicating the timing of LS is related to the bolting time of each individual ecotype. However, these studies are based on the bolting times of each ecotype and do not address the effects of altered bolting time within one ecotype.

A similar positive correlation between bolting age and both leaf and whole plant senescence was seen in a single ecotype model. *KHZ1* and *KHZ2* encode highly similar KH domain-Zn finger proteins. *khz1 khz2* double mutants bolt late and show delayed leaf and whole plant senescence. Overexpression of either KHZ1 or KHZ2 resulted in early bolting, leaf and whole plant senescence (Yan et al., 2017). Here, a positive correlation was seen between the varying bolting times and LS phenotypes in one ecotype. Some genes have been identified that regulate both flowering time and LS in Arabidopsis (*SOC1, WRKY75*), which may partially explain this temporal relationship in the single ecotype model (J. Chen et al., 2017; L. Zhang et al., 2018).

Although there are conflicting data, most recent studies point to a positive correlation between bolting and leaf and whole plant senescence. We hypothesized that there are gene expression changes occuring in the rosette associated with the reproductive transition at the shoot apical meristem that regulate LS. These gene expression changes could be contributing to the positive correlation seen between bolting time and LS. To address this hypothesis, we isolated a mutant that displays both early bolting and early LS. We then completed an RNA-seq time series experiment that compared the early bolting mutants to vegetative WT plants of the same age. Our approach allowed the identification of leaf gene expression changes specifically associated with bolting time. This list of 398 Bolting Associate Genes (BAGs) was enriched for LS and LS-related GO terms, and includes many well characterized LS regulators. Using a LS database (Li, Yang Zhang, Dong Zou, Yi Zhao, Hou-Ling Wang, Yi Zhang, Xinli Xia, Jingchu Luo, Hongwei Guo, 2020), we found that 202 of these BAGs are known to be associated with LS. A list of potential novel early LS regulators was proposed. We then produce a gene regulatory network summarizing BAG interactions generated from publicly available DAP-seq TF binding site data (O’Malley et al., 2016). This study shows that there are leaf gene expression changes associated with bolting that may regulate LS in *Arabidopsis thaliana*. These early regulators of LS likely contribute to the temporal relationship connecting bolting time and LS.

## Results

### *atx* Triple Mutant Phenotype

Previous data show that changing levels of the H3K4me3 mark correlate to changes in gene expression during age-dependent developmental LS (Brusslan et al., 2015). The peak of H3K4me3 marks in Arabidopsis is located about 400 bp downstream of the transcription start site (TSS) and is correlated to high expression of the nearby gene (Z. Zhang et al., 2016). For 389 genes, H3K4me3 marks increased in parallel with gene expression during age-dependent LS, and we named these genes H3K4me3-Senescence Up-Regulated Genes (K4-SURGs). We initially hypothesized that H3K4me3 accumulation might be a mechanism for increasing expression of senescence related genes. To address this hypothesis, we generated H3K4me3 methyltransferase mutants and studied their LS phenotypes.

Class III Arabidopsis trithorax (*ATX)* histone methyltransferases methylate H3K4 (Pontvianne et al., 2010). Different *ATX* enzymes catalyze different methyltransferase activities (mono-, di-, tri-methylation), and some single mutants display more severely altered developmental phenotypes than others (L. Q. Chen et al., 2017; Tamada et al., 2009; Yun et al., 2012). Our single *atx1, atx3*, and *atx4* mutants and double mutant combinations did not display a detectable change in LS, leading to the isolation of a homozygous *atx* triple mutant (*atx1 atx3 atx4*). We isolated two *atx* triple mutants (TM1 and TM2), which contain the same alleles but are derived from different F_2_ populations from distinct individual parents, thus are independent isolates of the same genotype. The (*atx1 atx3 atx4)* triple mutant genotype was confirmed both by PCR and RT-PCR (Supplemental Data File 1).These mutants displayed significantly early bolting (Figure 1A) and significantly early LS, quantified by *NIT2* and *WRKY75* mRNA induction (Figure 1B-C) and by chlorophyll loss (Figure 1D). We also confirmed that early bolting translated to an early flowering phenotype (Supplemental Data File 2, Sheet_5), meaning bolting is an appropriate phenotype marker for the vegetative-reproductive transition. The accumulation of H3K4me3 downstream of the *FLOWERING LOCUS C (FLC)* TSS is associated with high expression of *FLC*, a flowering inhibitor, thereby preventing the vegetative to reproductive transition (bolting)(Pien et al., 2008; Yun et al., 2012). The early bolting phenotype in the *atx* TMs is likely due to a decrease in H3K4me3 accumulation at the *FLC* locus and decreased gene expression of this and related flowering inhibitors.

**Figure 1:**
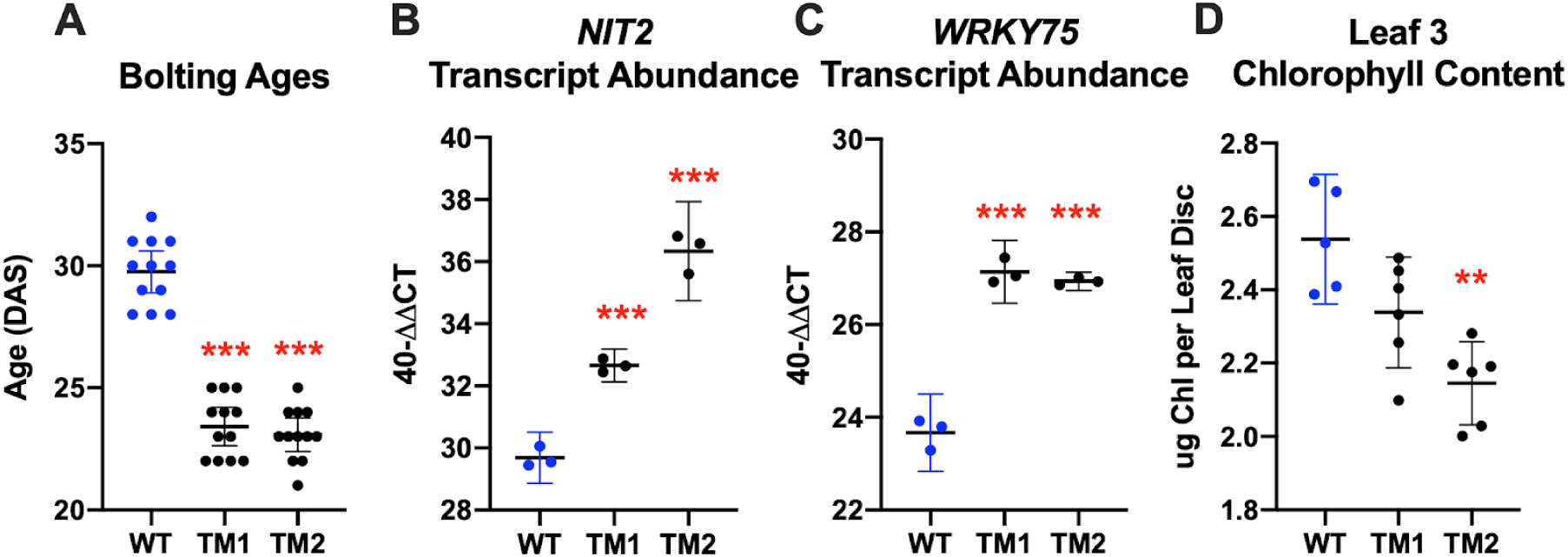
*atx1 atx3 atx4* Triple Mutant (TM) Phenotypes. A: A plant was considered to have bolted when the inflorescence extended 1 centimeter from the base of the rosette. B and C: Real-Time qPCR was used to measure the transcript abundance of two genetic LS markers, *NIT2* and *WRKY75*, in RNA isolated from leaf 6 at day 33. Individual data points shown are the averages of three technical replicates, generated from 6 plants. D: A leaf disc obtained by hole punching leaf 3 harvested at 33 days of age. For statistics, one-way ANOVAs were run. Then, T-tests with Bonferonni corrected p-values were completed to determine significance. For all data, one representative replicate of three is shown. Results were similar in all three replicates, which can be found in Supplemental Data File 2_Sheets 1-4. Error bars display 95% confidence intervals.

*NIT2* encodes a nitrilase and is a robust K4-SURG that serves as an mRNA marker for LS (Brusslan et al., 2012). *WRKY75* is a well characterized positive regulator of LS (T. Kim et al., 2019) that was also identified as a K4-SURG. H3K4me3 is an activating mark, thus if hypomethylation caused by *atx* mutation affected *NIT2* or *WRKY75*, we would expect lower gene expression, but we detected the opposite (Figure 1B-C). This led us to hypothesize that *NIT2* and *WRKY75* induction were not caused by local H3K4me3 changes, rather the coupling of LS to bolting time might be responsible. The genetic mechanism behind this temporal relationship has not been defined, and it became our goal to identify gene expression changes associated with the bolting event that may be regulating LS.

### RNA-seq Time-Series Experimental Design

We completed an RNA-seq time-series experiment that compared the early bolting *atx* TMs to vegetative WT Col-0 plants of the same age over a six-day time course (Figure 2). This single ecotype approach overcomes the genetic variability in different Arabidopsis ecotypes and the confounding factor of ecotypes that are acclimated to different geographical parts of the world being grown in a single environment. If LS-related signaling was being initiated in leaves by bolting, we would expect to see senescence-related differentially expressed genes (DEGs) between the *atx* TMs and WT, and enrichment of senescence-related biological processes within these DEGs.

**Figure 2:**
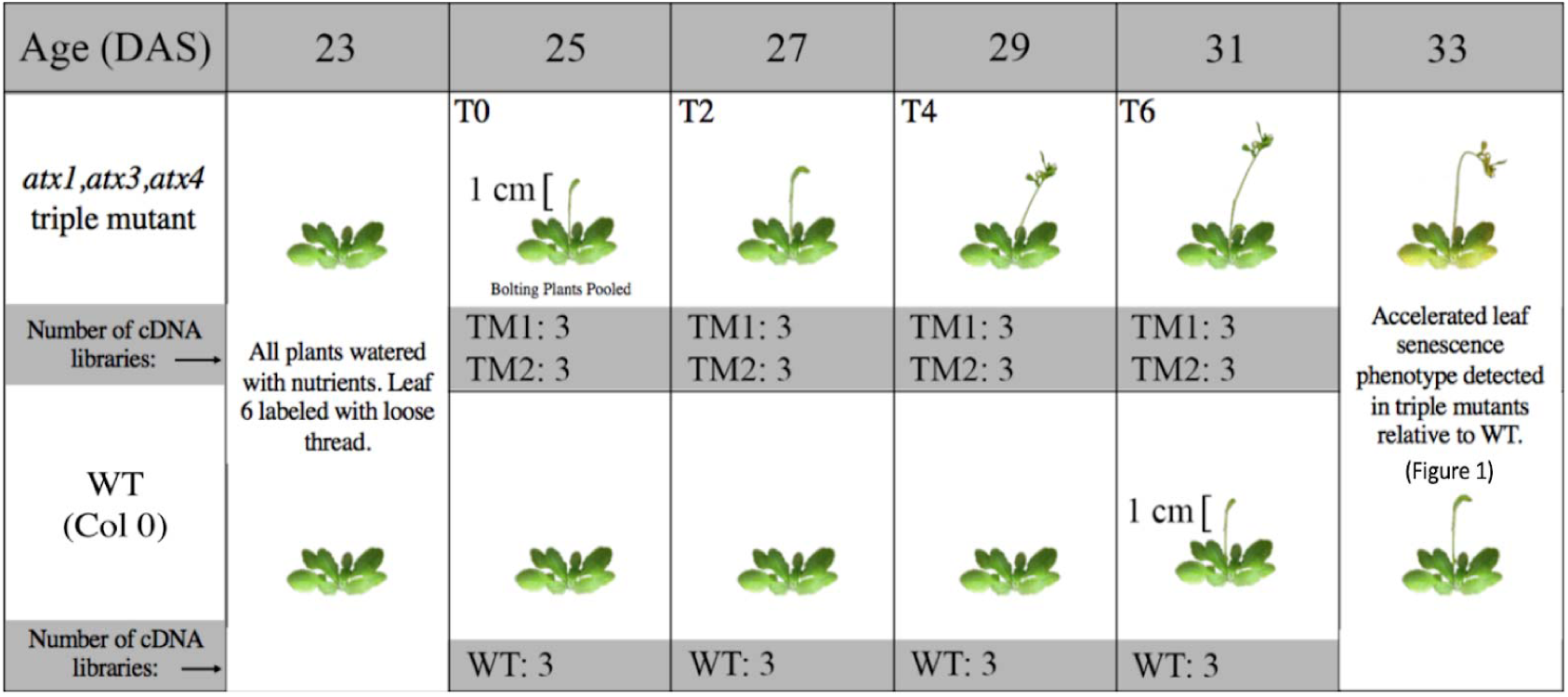
RNA-seq Time Series Experimental Design. WT, *atx* Triple Mutant 1 (TM1), and *atx* Triple Mutant 2 (TM2) were grown in long-day conditions. (n=54 per genotype).□ Bolting age was scored and plants recorded at the peak of TM bolting (days 24 and 25) were grouped. 10 plants were randomly selected from this group four times in two day increments to ensure that plants were developmentally similar. Leaf 6 was harvested from the 10 plants and stored at −80°C. The 10 leaves were homogenized in liquid nitrogen with a mortar and pestle and then separated into three tubes, which were treated as three replicates/libraries. All harvesting was completed between 8:00 and 10:00 AM to prevent interference by diurnal gene expression changes.

Plants that bolted at the peak of *atx* TM bolting (Day 24-25, T0) were grouped. Plants were then randomly selected from this group for leaf 6 harvesting, which began at T0 and continued in two day increments (T2, T4, and T6). This synchronization to bolting ensured the plants were developmentally similar. Vegetative WT leaf 6 control tissue was harvested at the same time points. This design allowed us to differentiate gene expression changes associated with bolting versus those associated with age. cDNA libraries were prepared and subject to high-throughput sequencing (Supplemental Data File 3 reports FPKM values).

As expected, hierarchical transcriptome clustering showed TM1 and TM2 are highly similar at each time point (Figure 3A). All T0 samples clustered together, regardless of genotype, because they are developmentally the most similar. The T2, T4, and T6 transcriptomes from bolting *atx* TMs cluster away from the vegetative WT samples (T0 and T2). WT samples from T4 and T6 cluster with the *atx* TM bolting samples, likely because they are nearing the vegetative-reproductive transition. These clustering patterns were also seen using Principal Component Analysis (PCA, Figure 3B).

**Figure 3:**
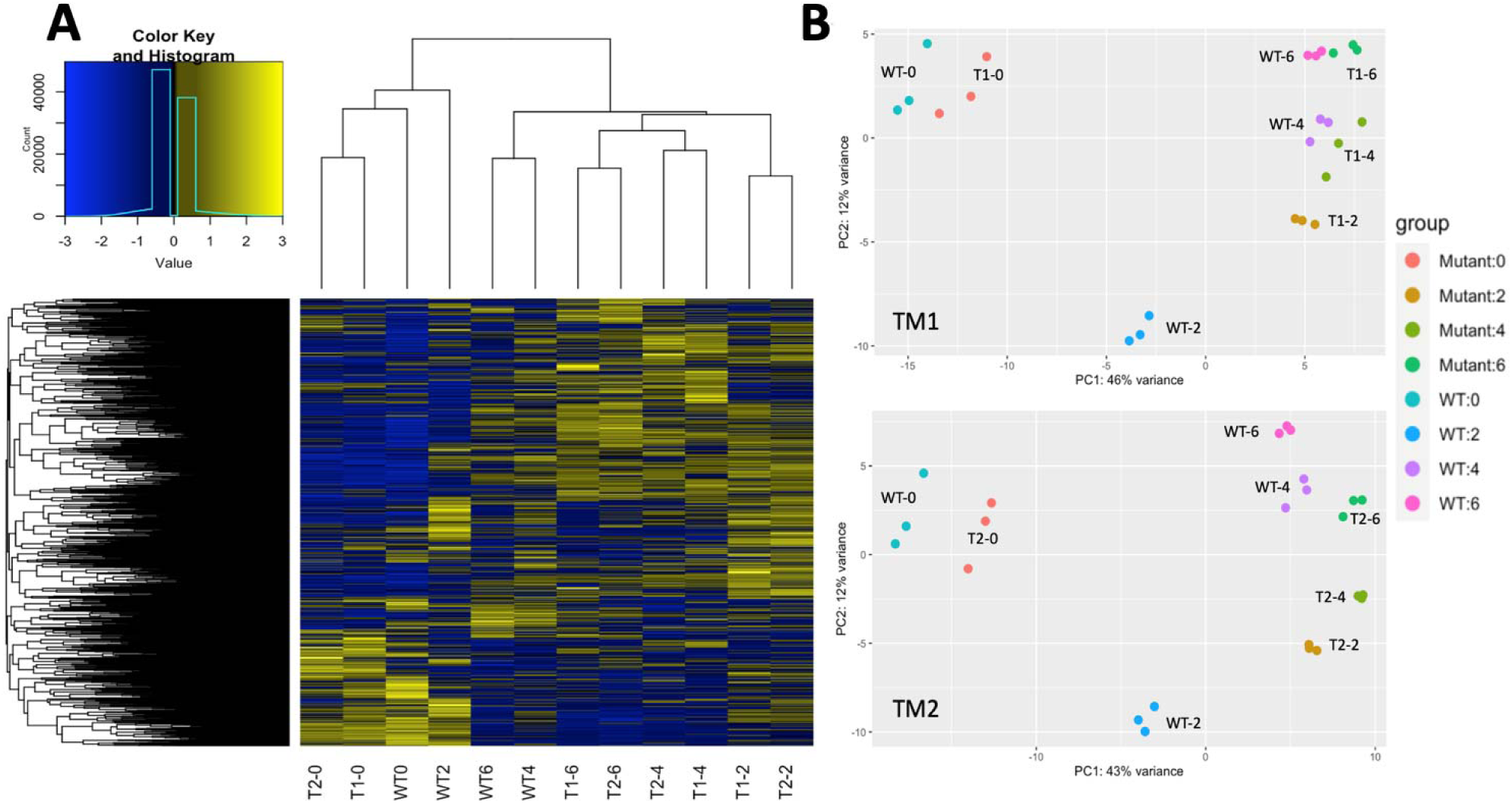
Transcriptome Comparisons. Hierarchical clustering was used to generate a heatmap of transcriptomes, which were generated by averaging expression levels of the three replicates per line at each time point. (T1= *atx* TM1, T2 = *atx* TM2). PCA was completed using the DeSeq2 package for WT vs. TM1 (top) and WT vs. TM2 (bottom) and the resulting charts were manually labeled by genotype and time.

### RNA-seq Data Analysis Strategy

Three methods were used to determine differential expression. A gene needed to be identified by at least two different statistical methods to be considered a DEG. A (Genotype + Time + Time:Genotype) design was used to run an LRT test with (∼Time) as a reduced model in DESeq2, which treated time as a continuous variable and removed genes that show the same expression change over time in all samples (Love et al., 2014). edgeR was used to compare each mutant to WT separately at each time point, meaning time was treated as a factor (Mark D. Robinson, Davis J. McCarthy, 2009). Lastly, a T-test based method was written in R (All code used for differential expression analysis can be found in Supplemental Data File 4). The use of multiple methods increased stringency to prevent spurious detection of differential gene expression. Our list of 750 DEGs (Supplemental Data File 5, Sheet 3) contained many bolting/flowering time regulators (*SOC1, FT, MED18, FLC, FPA, CIB1, CIB5, SEP1, SEP3, MAF4, and MAF5*), providing confidence that our harvesting method and differential expression analysis isolated bolting related gene expression changes.

We then sought to validate these results in WT Col-0 plants undergoing the normal (not early) vegetative-reproductive phase transition in other datasets (Supplemental File 5, Sheet 4). While not synchronized to bolting, Breeze *et al*. completed a similar developmental time series experiment in Arabidopsis (Breeze et al., 2011). Leaf 7 was harvested from WT Col-0 in two day increments. We selected the time point at which plants began to flower in their time series (Day 21) and included the three subsequent time points. DEGs from these four time points were overlapped with our DEG list. Gaudin *et al*. identified genes downstream of FT and SOC1 by monitoring transcriptome changes in WT Col-0 Arabidopsis plants as they transitioned from short day (SD) to long day (LD) photoperiods (Gaudin et al., 2019). This photoperiodic transition induces flowering, and while their transcriptomes were generated from pre-bolting plants, the genetic changes associated with the SD-LD transition are related to bolting.

Compared to our experiment, plants in these two experiments were solely WT genotypes, were not developmentally synchronized, were grown in different chambers, and DEG lists were generated using different statistical methods. However, all datasets covered the vegetative-reproductive transition. It is important to note that *FT* and *SOC1* were both found to be differentially expressed in all three experiments, indicating that canonical flowering time-related gene expression was occurring in each dataset.

### Bolting Associated Genes (BAGs)

In order to be considered a bolting-associated gene (BAG), a gene had to be identified by at least two out of the three statistical methods used in our analysis, and it had to be validated in at least one of the WT time series experiments. Default parameters on Genesect from VirtualPlant 1.3 were used to determine that all gene lists displayed significant overlap (p<.001) (Katari et al., 2010). Genes identified in our experiment that did not overlap with the WT experiments may be false positives or may be specifically associated with early flowering or changes in H3K4me3 accumulation. This approach stringently identified 398 genes that are differentially expressed at bolting time (Supplemental Data File 5, Sheet 5).

A script was written in R to visualize the expression profiles of all 398 BAGs. (Supplemental Data File 6). Select gene expression profiles are shown along with the corresponding profiles from the WT time-series experiments (Figure 5). *WRKY45* increased over time in all samples except for the vegetative WT control in our experiment. *DTX50* also showed strong induction in each experiment, though it decreased back to basal levels after four days, indicating a more transient expression. *PSK4* increases at T0 and then maintains expression levels higher than WT at all subsequent time points, which corresponds to the clear induction of *PSK4* in both WT time series. *SAG20* did not show as clear of a trend of expression over time, but it was consistently higher than vegetative WT levels. *SAG20* increases expression in both WT time series. *ANT* appears to decrease expression in all datasets undergoing the vegetative reproductive transition. WT follows the same trend, with a delayed decrease in *ANT* expression compared to the *atx* TMs, likely as WT was nearing bolting time.

**Figure 4:**
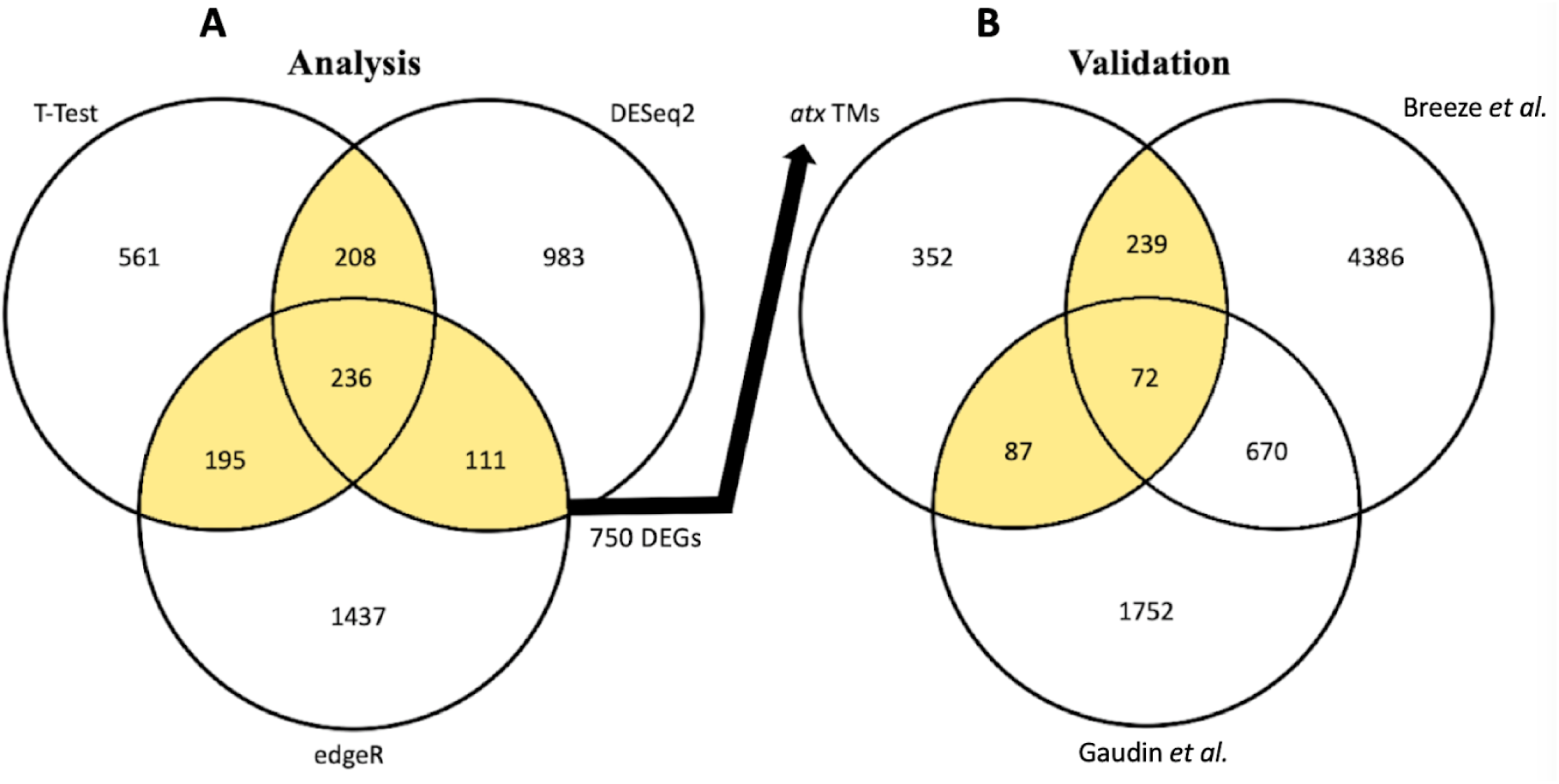
Differential Expression Analysis and Validation. A: DEGs lists from our T-test method (1200), DESeq2 (1536), and edgeR (1978) were overlapped. DEGs identified by at least two of the three statistical methods (750) were considered for validation. B: In order to be considered bolting associated, a gene needed to be differentially expressed in our *atx* triple mutant analysis and at least one of the publicly available WT time-series experiments. Highlighted portions in the Venn diagram mark those selected and used for validation from our analysis, or which genes were considered to be BAGs after validation. Supplemental Data File 5 contains gene lists.

**Figure 5:**
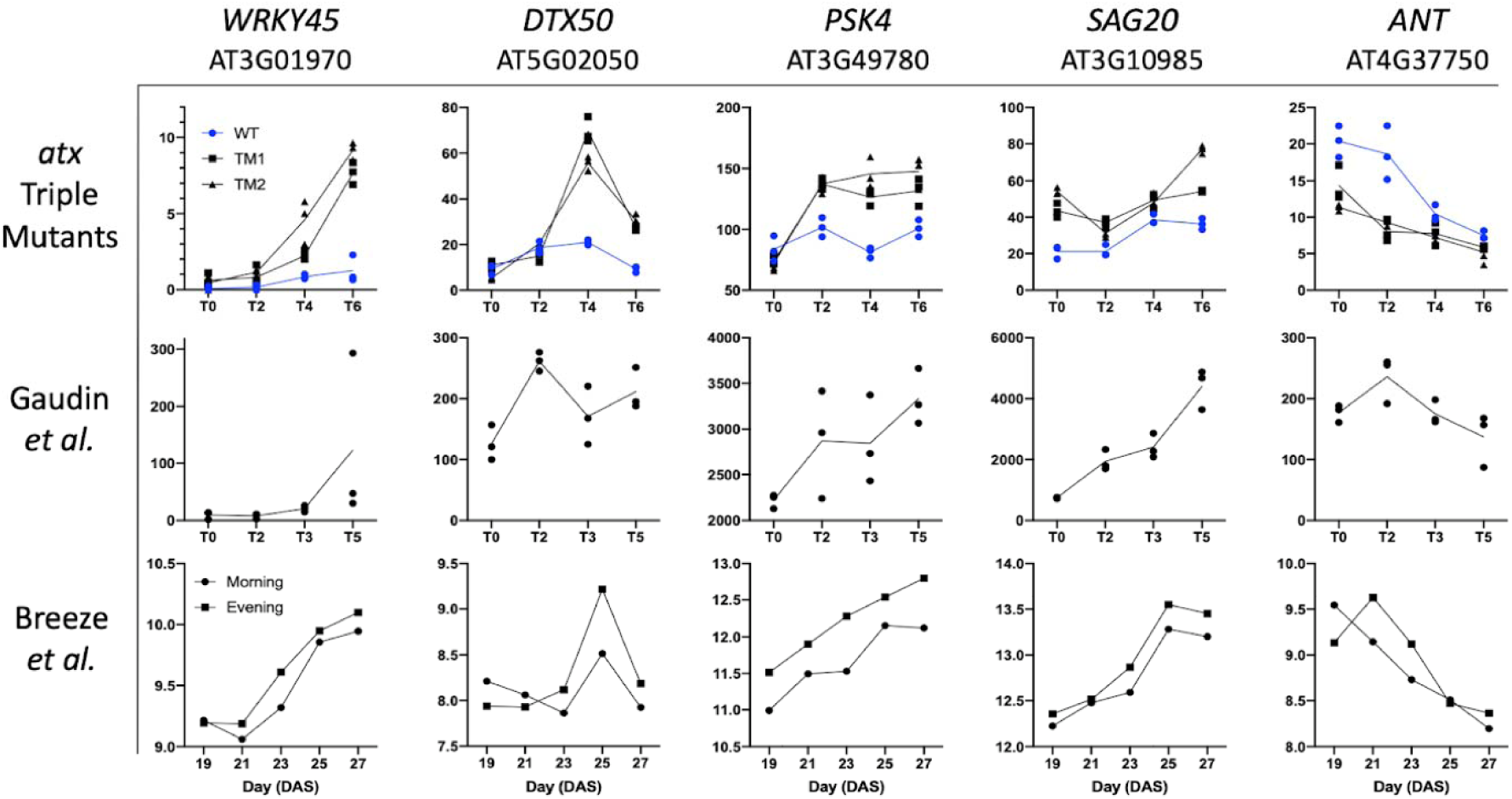
Examples of BAG Expression Across Experiments. Raw data from each experiment were used to generate these graphs in GraphPad PRISM. Data from the *atx* TM and Gaudin *et al*. experiments represent FPKM from RNA-seq, while Breeze *et al*. is Lowess normalized averaged (4 reps) signals in log space from a microarray experiment. Bolting occurs at T0 in the *atx* TM experiment and at Day 21 in the Breeze *et al*. experiment. T0 in the Gaudin *et al*. experiment represents the time of photoperiodic transition from short day to long day conditions. Samples from plants undergoing the vegetative-reprodutive transition are shown in black while vegetative control plants are shown in blue.

The 398 BAGs are enriched for both LS and LS-related biological processes (Figure 6A). The Panther Gene Ontology analysis allows extraction of the input DEGs associated with each enriched term (Huaiyu Mi, Anushya Muruganujan, 2012; Thomas et al., 2003)(Supplemental Data File 7). We intersected these input BAGs associated with individual enriched GO terms with the times that they were differentially expressed in our time series experiment (in TM2 from the edgeR analysis). We found that on average, among the genes associated with enriched GO terms, there was a significantly higher proportion of BAGs differentially expressed at T2 compared to all other time points (Figure 6B). This indicates that the BAGs associated with enriched GO terms change expression more frequently at T2, two days after bolting. These genes associated with enriched GO terms were changing at the time point most closely following the emergence of the bolt, supporting the hypothesis that the initial elongation of the bolt is the stimulus of LS-related signaling. This is further supported by the heatmap showing that the most dramatic differences in BAG expression between the *atx* TMs and WT are seen between T0 and T2 (Figure 6C). Furthermore, Short Time-series Expression Miner (STEM) clustering analysis was completed to determine if TM1 and TM2 displayed similar BAG gene expression patterns. STEM identified mostly the same significant gene expression clusters in both mutants (Figure 6D) (Ernst & Bar-Joseph, 2006).

**Figure 6:**
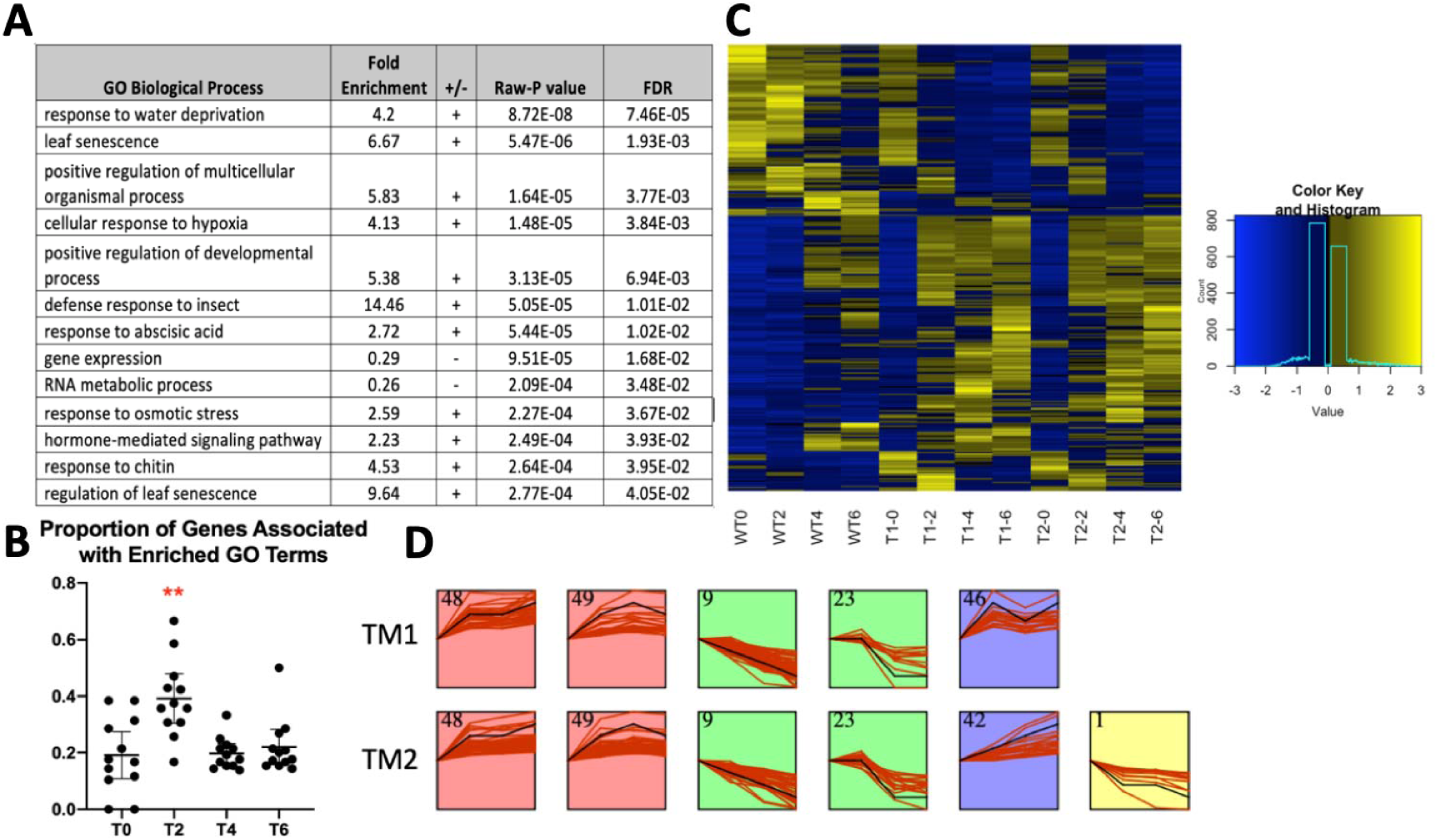
Bolting Associated Genes (BAGs) A: Enriched GO terms from a Panther Gene Ontology analysis are shown. B. An ANOVA run on the time-intersected enriched BAGs generated a significant P-value (1.3E-05). Pairwise T-tests were then completed using a Bonferonni corrected p-value (0.05/6 = 0.0083), which indicated that T2 was significantly different than all other time points [p= 0.0003 (T0), 0.0002 (T4), 0.0008 (T6)]. No other significant results were detected. Error bars show the 95 percent confidence intervals. C: A heatmap showing BAG expression levels in all 12 transcriptomes. D. Short Time Series Expression Miner (STEM) was used to find significant clusters of gene expression changes.

### Potential Novel Early Regulators of LS

We then wanted to identify genes within the BAG list that either regulate or are associated with LS. Zhang *et al*. generated a database of genes known to be associated with LS (LSD 3.0)(Zhonghai Li., 2019). Using VirtualPlant, we found a significant overlap between BAGs and LSD 3.0 (p<.001). 202 of the 398 BAGs (50.7%) were shared (Supplemental Data File 5, Sheet 6). While some of these known LS-associated BAGs likely contribute to the coupling of LS to flowering time, we also wanted to find potential novel LS regulators. Kim et al completed a dark-induced detached leaf senescence (DILS) RNA-seq time series experiment with WT Col-0 plants that we used to find genes that change expression during DILS (GSE99754) (Jeongsik Kim, Su Jin Park, Il Hwan Lee, Hyosub Chu, Christopher A Penfold, Jin Hee Kim, Vicky Buchanan-Wollaston, Hong Gil Nam, Hye Ryun Woo, 2018). We compared gene expression from T0 (before dark treatment) and T3 (3 days into dark treatment) using DESeq2 with default parameters and standard cutoffs (P<0.05, FDR<0.05, and a log2 fold change>2) (Supplemental Data File 5, Sheet 7). We intersected the DILS DEGs with the BAG and LSD gene lists, and a significant overlap was found (p< 0.001). 91 BAGs were shared in the DILS experiment, but were not in the LSD (Figure 7A, Supplemental Data File 5, Sheet 8). 68 (74.7%) of these 91 genes were downregulated during *atx* TM bolting and DILS, while 15 (16.5%) were upregulated during *atx* TM bolting and DILS. Seven of these 91 genes were upregulated in bolting *atx* TMs, but downregulated during DILS and one gene was downregulated during *atx* TM bolting, but upregulated during DILS. A subset of BAGs that change expression during DILS, but are not present in the LSD, are signaling molecules that may represent novel early regulators of LS (Figure 7B).

**Figure 7:**
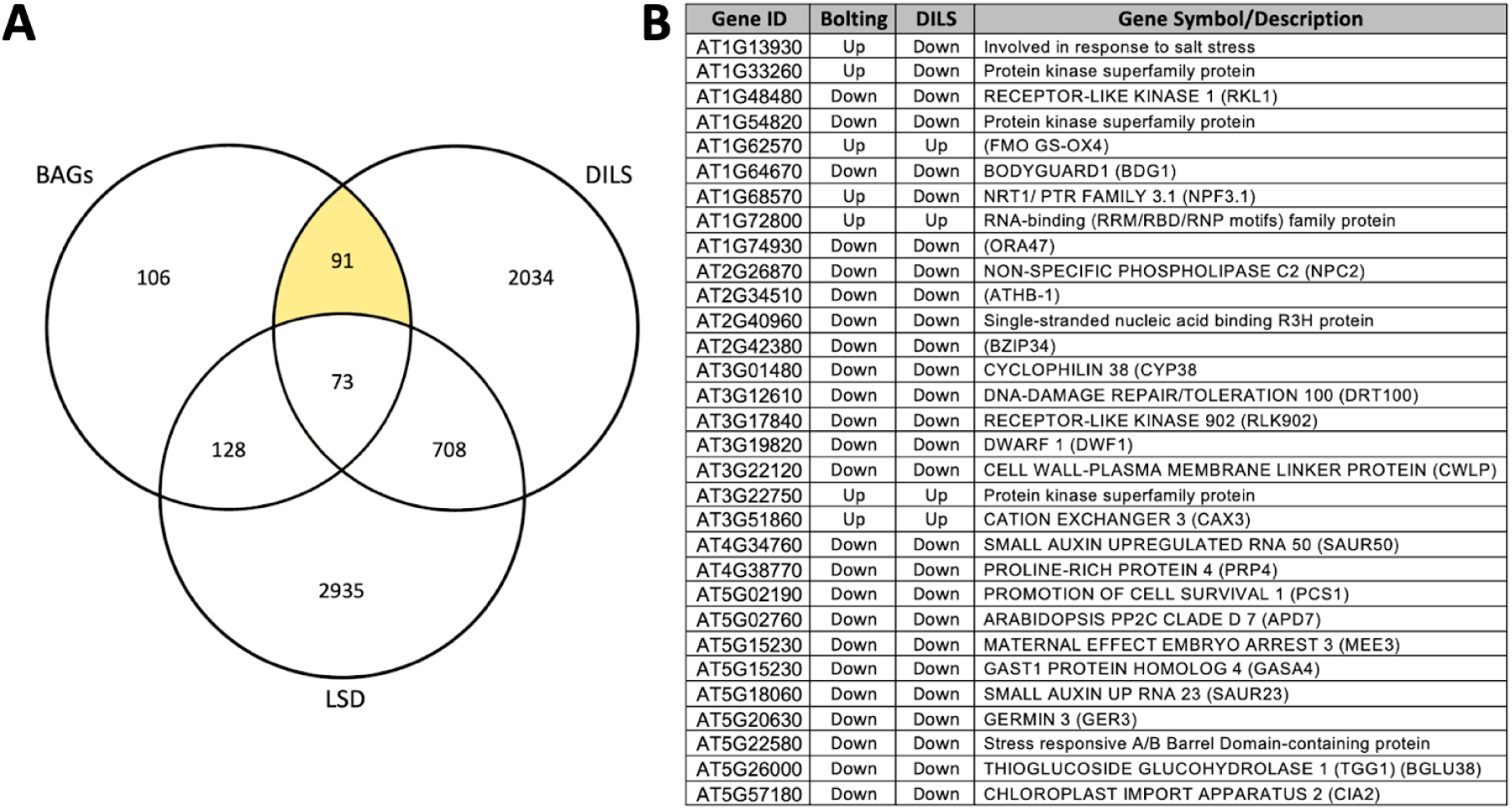
Potential Novel LS Regulators. The Venn Diagram shows overlap between the 398 BAGs, the 2906 DEGs from the DILS dataset, and the 3844 SAGs in the Leaf Senescence Database (LSD). The 91 BAGs that were differentially expressed in the DILS dataset but not in the LSD are highlighted. Select signaling genes from this list of 91 genes are shown in the table along with their direction of gene expression at bolting and during DILS. Gene Symbols/Descriptions were obtained using the bulk data retrieval tool in TAIR.

### Bolting Associated Gene Regulatory Network (GRN)

We sought to identify genetic interactions between BAGs, of which many are transcription factors (TFs). Recently, the development of DNA-Affinity-Purification sequencing (DAP-seq) has allowed for high throughput screening for TF binding sites (Bartlett et al., 2017). We wrote a script that uses DAP-seq data from GEO to generate lists of bound targets for the bolting associated TFs for which data were available. (GSE60141) (Supplemental Data Files 8 and 9)(O’Malley et al., 2016). We then intersected these data with the time and direction of differential expression from our experiment, and annotation in the LSD (Supplemental Data File 10). These data were uploaded to Cytoscape for visualization (Shannon et al., 2003). To reduce network density, nodes are only shown at the first time they were differentially expressed. A side effect of the display is that some interactions appear reversed in time. An example of this is with ANAC046 regulating *ERF054. ANAC046* is differentially expressed at T2, but *ERF054* is differentially expressed at every time point. Since ERF054 is only shown at T0, the first time it is differentially expressed, it appears that ANAC046 is conferring temporally backwards regulation.

All nodes in the network are BAGs. The network shows that there are bolting time associated TFs that can bind and might explain the differential expression of BAGs. There appears to be a shift from upregulation (red) to downregulation (blue) from T0 to T6. While most of the TFs appear to act independently, some interactions between TFs were identified in the DAP-seq data. *ERF054* is bound by ORE1, ANAC046, and GBF3 while *ANAC046* is bound by HB34. Select signaling molecules were included in the subnetwork focusing on ERF054 (Figure 9). All TFs in the subnetworks confer differential expression at multiple time points and have both shared and independent targets.

**Figure 8:**
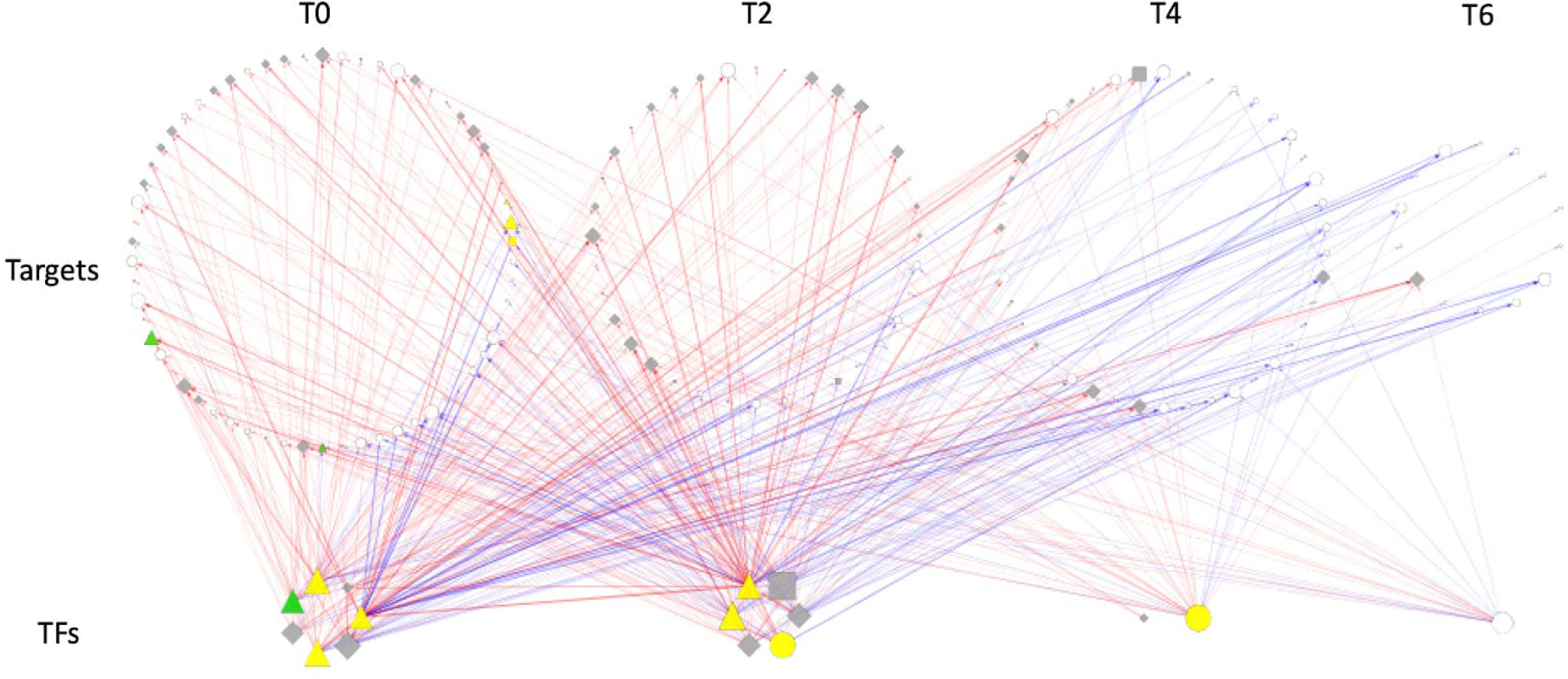
Bolting Time-Associated Gene Regulatory Network. The network was manually constructed as described in the Methods, and uploaded to Cytoscape for visualization. White nodes were not present in the LSD. Grey nodes were present, but had an unclear function listed in the LSD. Yellow nodes promote LS while green nodes prevent LS. Upregulation is shown by red edges and downregulation is shown with blue edges. Node shape refers to the type of evidence used to determine LS association in LSD 3.0; diamonds represent genomic (microarray) data, triangles represent mutant studies, and squares represent molecular data.

**Figure 9:**
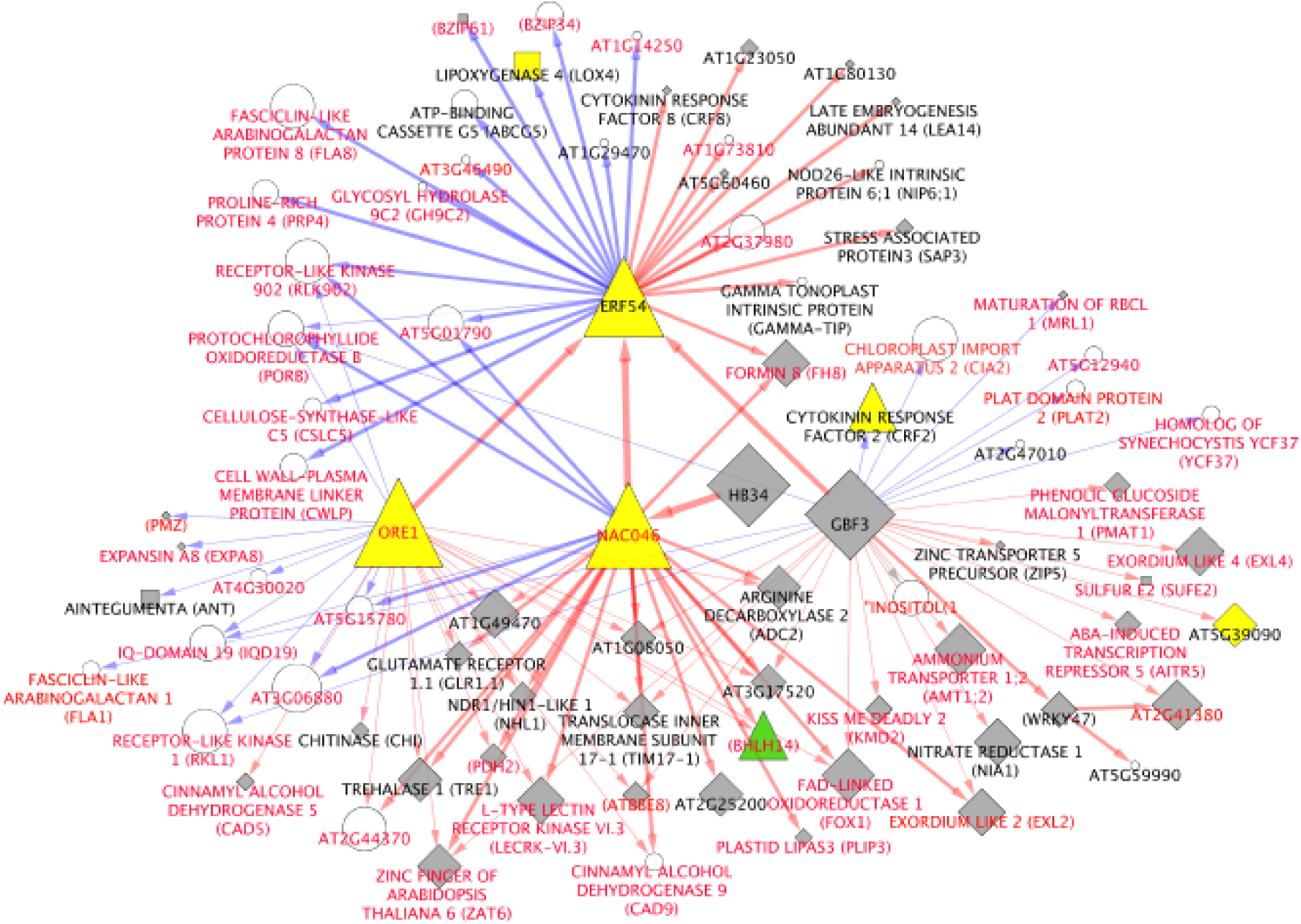
*ERF054* Subnetwork. White nodes were not present in the LSD. Grey nodes were present, but had an unclear function. Yellow nodes promote LS while green nodes prevent LS. Upregulation is shown by red edges and downregulation is shown with blue edges. Node shape refers to the type of evidence used to determine LS association; diamonds represent genomic data, triangles represent mutant studies, and squares represent molecular data. Red labels refer to genes that are differentially expressed during DILS and black labels refer to genes not differentially expressed during DILS. Potential novel LS regulators are shown as white nodes with red labels.

DAP-seq binding data show multiple senescence associated TFs (ORE1, ANAC046, HB34, and GBF3) converging towards *ERF054*. This may explain the gene expression profile of *ERF054*, both in our experiment and Breeze et al. (Supplemental Data File 6, At4g28140). Note that time resolution of data are not shown in the subnetworks. Only *ORE1* binds to *ERF054* at T0, explaining the significantly increased expression of *ERF054* at T0 in the *atx* TMs relative to WT. However, even greater induction of *ERF054* gene expression is seen in the *atx* TMs at T2 relative to WT, when other TFs (*GBF3, ANAC046*, and *HB34*, which itself binds *ANAC046*) are differentially expressed and can also bind to *ERF054*. This network also shows that many targets of these TFs are differentially expressed in DILS but not included in the LSD (white nodes with red labels). These data further support these genes being novel LS regulators, as they are directly bound by well characterized LS-promoting TFs.

Other subnetworks were created by selecting known TFs known to confer LS-related signaling: WRKY45 (Figure 10A) and HB34 (Figure 10B). WRKY45 is a positive regulator of LS (L. Chen et al., 2017) that increases expression at the time of bolting (Figure 5). *WRKY45* expression is induced during DILS, and it is present in the LSD. The subnetwork shows WRKY45 binds to many LS-associated genes (grey nodes), and many genes that are differentially expressed during DILS (nodes labeled in red). For example, WRKY45 binds to *AHK4*, which is downregulated at bolting and is a negative regulator or LS (Riefler et al., 2006). This interaction supports WRKY45 being a positive regulator of LS. WRKY45 also binds to genes that are differentially expressed during DILS, but are not present in the LSD (white nodes with red labels). The same is seen for HB34. HB34 binds and may activate the expression of *ANAC046*, a positive regulator of LS (Oda-Yamamizo et al., 2016). It also binds to known SAGs and potential novel regulators of LS. Interestingly, most of the potentially novel LS regulators are downregulated in the subnetworks (Figures 9 and 10: *CIA2, DWF1, RKL1, CMCU, BZIP34, RLK902*).

**Figure 10:**
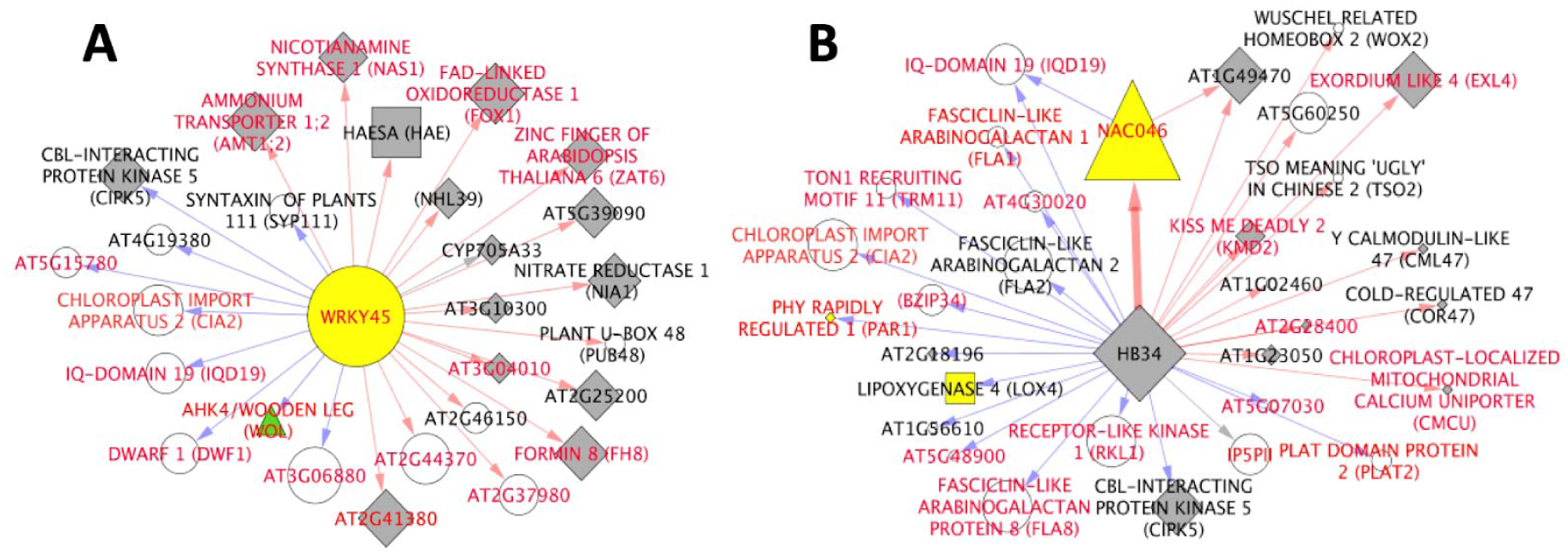
Select Subnetworks. The network was manually constructed in R (description in methods), then uploaded to Cytoscape for visualization. White nodes were not present in the LSD. Grey nodes were present, but had an unclear function. Yellow nodes promote LS while green nodes prevent LS. Upregulation is shown by red edges and downregulation is shown with blue edges. Node shape refers to the type of evidence used to determine LS association; diamonds represent genomic data, triangles represent mutant studies, and squares represent molecular data.

## Discussion

The goal of our study was to better understand the molecular connection between the vegetative-reproductive transition and LS. We hypothesized that there are LS-related gene expression changes associated with the bolting event. Using RNA-seq, we generated a list of 398 bolting associated genes (BAGs). 202 of these BAGs were present in the Leaf Senescence Database (LSD), some of which are likely responsible for temporally connecting LS to bolting time. We also identified 91 BAGs that are differentially expressed during dark-induced leaf senescence (DILS) but are not present in the LSD. Further study of these 91 genes may reveal that some are novel early regulators of LS.

### *atx* Triple Mutant Phenotypes

While the two *atx* triple mutants (TM1 and TM2) contain the same *atx1 atx3* and *atx4* alleles, we feel justified in treating them as two independent early flowering mutants: TM2 appears to have a stronger early LS phenotype, it consistently produced more DEGs than TM1, and clustered further away from WT than TM1 did in PCA analysis (Figure 3B). We think that by overlapping the results between TM1 and TM2, we reduced the probability of false discoveries. Furthermore, TM2 consistently displayed a slightly more severe early LS phenotype than TM1 (Figure 1D, Supplemental Data File 2). There are likely background mutations unique to TM1 or TM2 that contribute to this difference. Regardless, all BAGs were significantly differentially expressed in both TM1 and TM2. Early flowering has been reported in other *atx* mutants (Berr et al., 2015; Yun et al., 2012) which is consistent with our findings. Chen et al isolated an (*atx3, atx4, atx5)* triple mutant but did not report a change in flowering time or LS (L. Q. Chen et al., 2017). The five *ATX* enzymes are classified into two clades (*ATX1* and *ATX2*), and (*ATX3, ATX4*, and *ATX5)*, thus their genetic divergence may explain the difference in phenotypes between the (*atx1 atx3 atx4*) and the (*axt3 atx4 atx5*) triple mutants.

### RNA-seq Time Series Experiment

Synchronizing tissue harvesting to bolting differentiated bolting-associated and age-associated changes in gene expression. Multiple statistical approaches were used for the identification of 750 initial DEGs since there is not one perfect statistical approach for our time-series analysis. Treating time as a continuous variable as we did in DESeq2 helps to identify time-resolved gene expression changes. However, our edgeR based method treated time as a factor, which may have allowed greater detection of transient changes in gene expression. It is common to see high overlap between edgeR and DESeq2, however that typically occurs when the same underlying statistical design is used. Here, we did not expect strong overlap as the programs were run with different designs. Even with the varying designs, Genesect found significant overlap between these gene lists (p< 0.001).

Lack of reproducibility for large dataset analysis is a chronic issue (Joël Simoneau, Simon Dumontier, Ryan Gosselin, 2019; Porra et al., 1989). While some programs are commonly used for differential expression analysis (edgeR, DESeq2, limma), we felt justified to include a T-test method, as it increased transparency and showed that a bulk of the DEGs identified by the two commonly used programs were supported by simple T-tests. Our stringent overlapping method should reduce the risk of false positives.

It was also important to validate our results in WT plants because the *atx* TMs may have unknown epigenetic effects. It would have been ideal to repeat the RNA-seq time-series experiment, but to center it around bolting WT plants, and use early and late flowering controls, however our use of publicly available data was more practical and cost effective. While this is one limitation to our study, two published experimental designs were similar to ours. These two WT datasets used similarly aged leaves, similar time resolution, and were centered around the vegetative-reproductive transition, which was made apparent by the identification of *FT* and *SOC1* as DEGs in both. As expected, our dataset had stronger overlap with Breeze *et al*. than Gaudin *et al*., as plants in the Breeze *et al*. dataset were bolting. Gaudin *et al*. was specifically looking at the genetic changes associated with *FT* and *SOC1* expression, and their plants were not yet bolting. Both datasets had multiple replicates for high confidence output. Ultimately, changes in gene expression will need to be validated with real-time qPCR of separate biological replicates.

### Bolting Associated Genes (BAGs)

The 398 BAGs were enriched for LS and LS-related biological process GO terms. This supports the hypothesis that bolting stimulates LS-related signaling. The following known LS-associated genes were identified in our *atx* triple mutant analysis and then further validated in **<u>both</u>** WT time series experiments. *SOC1* is a classic example of a gene that positively regulates both flowering time and LS (J. Chen et al., 2017). *ANAC032* positively regulates flowering time and LS and is responsive to oxidative stress (Mahmood et al., 2016). *WRKY45* positively regulates LS, and WRKY45 overexpression lines displayed early bolting along with early LS, suggesting WRKY45 may play a role in regulating both flowering time and LS (L. Chen et al., 2017). *SAG20* and *SEN1* are both LS marker (Fernández-Calvino et al., 2016; Schenk et al., 2005; Weaver et al., 1998). *DEAR1* promotes cell death and SA synthesis (Tsutsui et al., 2009). Mutation of *ELS1*, a close homolog of the BAG *DTX50* results in both early flowering and early LS (Z. Wang et al., 2016). *DTX50* is a transiently expressed ABA transporter (H. Zhang et al., 2014). The induction of *DTX50* at bolting time in all experiments indicates ABA flux is associated with bolting, which could be promoting LS. *ADP7 (SSPP)* is a senescence suppressed protein phosphatase (Xiao et al., 2015). Some genes identified in all three experiments were also found to be K4-SURGs (*WRKY45*, mate efflux AT1G33110, peroxidase superfamily protein At2g371300, *BGLU11, CAX3, CTPS4, PSK4*, and *TBL38* (Brusslan et al., 2015).

Other LS related BAGs were shared between the *atx* TMs and Breeze *et al*. dataset. *ORE1* (*ANAC092*) is a well studied promoter of LS (Qiu et al., 2015). *WRKY28* is responsible for activating SA biosynthesis genes and promotes LS (Tian et al., 2020; van Verk et al., 2011). *SAG13* is a ROS responsive BAG that regulates DILS (Dhar et al., 2020). When knocked out in higher order mutants to reduce redundancy with other CRF TFs, *crf2* mutants showed delayed LS, suggesting the BAG *CRF2* works with other CRF TFs to promote LS (Raines et al., 2016). *BOI* attenuates cell death and regulates flowering time (Nguyen et al., 2015). BHLH112 plays roles in both flowering time and stress response (H. C. Chen et al., 2017; Y. Liu et al., 2015). The following genes were shared between the atx TMs and the Gaudin et al dataset. *WRKY46* is responsible for activating SA biosynthesis genes and promotes LS (van Verk et al., 2011). *WRKY48* is a stress and pathogen induced regulator of plant defense. (Deng-Hui et al., 2008). The BAG *SAUR41* acts redundantly with *SAUR49* to promote LS by regulating *SSPP*, another BAG with a known role in LS (Wen et al., 2020; Xiao et al., 2015).

Other BAGs associated with flowering time, seed development, and seed nutrient content were also identified. *UMAMI28* is responsible for transporting amino acids to the developing seeds (Müller et al., 2015). *SUS3* regulates sugar metabolism in developing seeds (Angeles-Núñez & Tiessen, 2010). The BAG *SEP1* works redundantly with *SEP2* and *SEP3* to regulate flower and ovule development (Pelaz et al., 2000). *CURVY1* regulates both flowering time and seed development (Gachomo et al., 2014). *FUL (AGL8)* and *BFT* regulate flowering time (Marian Bemer, Hilda van Mourik, Jose M Muiño, Cristina Ferrándiz, Kerstin Kaufmann, 2017; Ryu et al., 2011; Vicente Balanzà, Irene Martínez-Fernández, 2014). These genes change expression in the 6th rosette leaf, suggesting they may contribute to processes other than flower and seed development and nutrition.

The abundance of enriched genes peaking at T2 supports our hypothesis that bolting stimulates LS signaling. Significantly more genes associated with enriched processes were differentially expressed two days after the bolting event. The heatmap (Figure 6C) also highlights the most differential expression in TMs relative to WT between T0 and T2. Furthermore, half of the BAG list was found to be in the LSD, meaning half of the BAGs are known to be associated with LS. These findings further support that the bolting event stimulates LS related signaling in mature leaf 6, and that our harvesting method captured the signaling stimulated by bolting.

### Potential Novel LS Regulators

Ninety-one BAGs were not present in the LSD, but were differentially regulated during DILS (Figure 7B). A majority of the potential novel LS regulators (91%) change expression in the same direction during both bolting and DILS, but seven (7.7%) are upregulated at bolting and downregulated during DILS. These seven genes might prevent LS prior to and during bolting, and then must be downregulated for normal or dark-induced LS to proceed. Only one gene was downregulated in bolting *atx* TMs but upregulated during DILS. Some of these 91 genes regulate processes that are known to be related to LS. For example, *ORA47* regulates both ABA and JA synthesis (H. Chen et al., 2016). The BAG BZIP34 forms heterodimers with BZIP61, which is a BAG in the LSD that is also differentially expressed during DILS (Gibalová et al., 2009)(LSD database annotation). BZIP61 and BZIP34, which share 71% amino acid identity, may work together to regulate LS downstream at bolting time. While many of these 91 genes have known functions related to LS, a phenotype screen of verified mutants is needed to support their function.

### GRNs

The main GRN (Figure 8) shows that BAG TFs can regulate the expression of other BAGs at every time point in our experiment. This format was obtained by separating TFs and targets. Nodes were then separated by time. Some targets were removed from the subnetworks to reduce density, however no genes that were differentially expressed during DILS (red labels) or were potential novel LS regulators (red labels on white nodes) were removed. Subnetworks (Figures 9 and 10) show how individual TFs have the potential to regulate many well known regulators of development and LS. CML47 inhibits SA accumulation and defense, thus the upregulation of CML47 at bolting may serve as a mechanism to prevent inappropriately early LS (Lu et al., 2018). NAP is a positive regulator of LS, thus its decrease in expression at bolting may also be working to prevent early LS (Lei et al., 2020). *HAI1 (SAG113)* is ABA responsive and is downstream of NAP (Wong et al., 2019; K. Zhang & Gan, 2012). KMD2 inhibits CK signaling, which delays LS, thus the induction of KMD2 at the time of bolting might promote LS (H. J. Kim et al., 2013). *NIA1* and *NIA2* are responsible for nitrate reduction in plants and were both significantly upregulated after bolting time (Olas & Wahl, 2019). N metabolism and transport are a major component of LS. Both *NIA1* and *NIA2* are bound by many well characterized LS regulators that change expression at bolting time (Figures 9 and 10).

### Conclusion

We have identified 398 BAGs expressed in mature leaf 6 that change expression at the time of bolting. 202 of these BAGs are known to be associated with LS (Zhonghai Li et al., 2020). Many of these BAGs are also differentially expressed during DILS. Furthermore, we identify a set of genes that change expression during bolting and DILS, but are not in the LSD, meaning some may be novel early regulators of LS. Together, this study identifies LS-related gene expression changes that occur in a specific mature rosette leaf two days after the vegetative to reproductive transition at the shoot apical meristem.

## Abbreviations

ABA: abscisic acid
BAGs: Bolting Associated Genes
CK: Cytokinin
DEG: Differentially Expressed Genes
DILS: Dark Induced Leaf Senescence
GEO: Gene Expression Omnibus
GO: Gene Ontology
GRN: Gene Regulatory Network
JA: Jasmonic acid
LS: Leaf Senescence
LSD: Leaf Senescence Database
PCA: Principal Component Analysis
PCR: Polymerase Chain Reaction
RT-PCR: Reverse Transcriptase Polymerase Chain Reaction
qPCR: Quantitative Polymerase Chain Reaction
SA: Salicylic Acid
T0: Bolting Time
T2: 2 Days After Bolting
T4: 4 Days After Bolting
T6: 6 Days After Bolting
TMs: *atx* Triple Mutants
WT: Wildtype

## Materials/Methods

### Plant growth conditions

*Arabidopsis thaliana* plants were grown in Sunshine® Mix #1 Fafard®-1P RSi soil (Sungro Horticulture), which was treated with Gnatrol WDG (Valent Professional Products) (0.3 g/500 ml H_2_O) to inhibit the growth of fungus gnat larvae. Plants were sub-irrigated with Gro-Power 4-8-2 (Gro-Power, Inc., 8 ml per gallon), and grown in Percival AR66L2X growth chambers under a 20:4 light:dark diurnal cycle (Long Day) with a light intensity of 28 μmoles photons m^-2^ s^-1^. The low light intensity prevents light stress in older leaves, which was evident as anthocyanin accumulation at higher light intensities. To compensate for the reduced light intensity, the day length was extended. The petiole of the sixth leaf to emerge was marked with a thread on individual plants.

### Genotype Analysis

Genomic DNA was isolated from two-three leaves using Plant DNAzol Reagent (Thermo Fisher) following manufacturer’s instructions. Pellets were dried at room temperature for at least two hours, and resuspended in 30 μl TE (10 mM Tris, pH 8.0, 1 mM EDTA) overnight at 4°C. One microliter of genomic DNA was used as a template in PCR reactions with primers listed in supplemental file 1. All standard PCR reactions were performed with a 57°C annealing temperature using *Taq* polymerase with Standard *Taq* Buffer (New England Biolabs).

### Chlorophyll Analysis

One hole-punch was removed from each marked or detached leaf and incubated in 800 µl N,N-dimethyl formamide (DMF) overnight in the dark. 200 µl of sample was transferred to a quartz microplate (Molecular Devices) and absorbance at 664 and 647 nm was measured with a BioTek Synergy H1 plate reader. Absorbance readings were used to determine chlorophyll concentration (Porra et al., 1989). For each genotype/condition, *n* = 6 single hole punches from 6 individual plants.

### Gene Expression

Total RNA was isolated from leaf 6 using Trizol reagent. 1,000 ng of extracted RNA was used as a template for cDNA synthesis using MMLV-reverse transcriptase (New England Biolabs) and random hexamers to prime cDNA synthesis. The cDNA was diluted 16-fold and used as a template for real-time qPCR using either ABsolute QPCR Mix, SYBR Green, ROX (Thermo Scientific) or qPCRBIO SyGreen Blue Mix Hi-Rox (PCR Biosystems), in Step One Plus or Quant Studio 6 Flex (Thermo Fisher) qPCR machines. All real-time qPCR reactions were run with a 61°C annealing temperature, and normalized to *ACT2*.

### RNA-seq library construction

Ten plants, per line at each time point were selected for harvesting from the developmentally synchronized group of bolting plants. Leaf 6 was harvested from all 10 plants and immediately flash-frozen together. A mortar and pestle were used to grind tissue in liquid nitrogen. Homogenized tissue was separated evenly into three tubes to be treated as three replicates. The Breath-Adaptive Directional sequencing (BrAD-seq) (Townsley et al., 2015) protocol was completed to generate cDNA libraries. A 1/50th dilution of the final library was used with *ACT2* primers in qPCR to check for library consistency. The Illumina-ready cDNA libraries that were then sequenced at the UCI Genome High-Throughput Facility (GHTF).

### RNA-seq data analysis

Raw data were aligned to the TAIR10 genome and counted using Kallisto. Data were then exported to R and rounded to the nearest integer for differential expression analyses. DESeq2 was completed using a 2 factor (Genotype + Time + Genotype:Time) model that treated time as a continuous variable, rather than a category (Supplemental Data File 4). An 0.05 p-value and adjusted P-Value (FDR) cutoffs was used to determine significance. PCA was completed with DESeq2 (Love et al., 2014). Comparisons were made at each time point with edgeR with a p-value cutoff of 0.05 and a 1.5 fold change cutoff for significance (Supplemental Data File 4) (Mark D. Robinson, Davis J. McCarthy, 2009). A simple T-test based approach was written in R that used a 0.05 p-value and 2-fold change cutoff (Supplemental Data File 4) and used to compare WT and *atx* TMs at each time point. For all three approaches, WT was compared with TM1 and TM2 separately. The overlap between the TM1 vs WT and TM2 vs WT DEG lists was determined for each statistical method separately using R. The intersection of these DEG lists between the three methods was then identified using http://www.interactivenn.net/. Heatmaps were generated using heatmap.2.

### GRN Construction

A script was written in Xcode to be run on a Macbook to analyze DAP-seq data in the NARROWPEAK format (DAPSEQ.sh). Chromosome sequences were provided in the same working directory and BLASTn was run in the terminal. The list of BAGs was overlapped with the total list of transcription factors in *Arabidopsis thaliana*. http://planttfdb.cbi.pku.edu.cn/index.php?sp=Ath. These bolting associated TFs were then overlapped with the mmc2 file provided with the Ecker *et al*. DAP-seq data set to determine which TFs were present. (GEO Dataset GSE60141). The col samples were used for each of the 21 TFs for which binding data were available. R was used to find the intersection between the lists of TF binding targets and the BAGs. The resulting data were uploaded to cytoscape for visualization.

### Accession Numbers

RNA-seq data from this study was made publicly available on GEO as GSE134177.

### Supplemental Data Files

Supplemental Data File 1. T-DNA insertions disrupt gene expression in all three *atx* mutant alleles.

Supplemental Data File 2. *atx* TM phenotype data: All three replicates of raw triple mutant phenotype data are shown along with appropriate statistics and the primers used in this study.

Supplemental Data File 3. Raw FPKM data from the TM RNA-seq Experiment

Supplemental Data File 4. Differential Expression Analysis Code: Annotated code intended for R (Version 1.2.5033) is included for all three DE analyses.

Supplemental Data File 5. Gene Lists

Supplemental Data File 6. BAG expression profiles

Supplemental Data File 7. Reformat.sh: This script was written for a Mac terminal and it takes JSON output files from the Panther Gene Ontology tool and reformats it to make it able to be intersected with time data and used as input in the dcast command in R.

Supplemental Data File 8. DAP-seq.sh: This script takes DAP-seq data in the Narrowpeak format and generates gene names associated with the input genomic locations. It is specific for Arabidopsis and the chromosome sequencences must be present in the same working directory as the program. One must also be able to run BlastN in the terminal. This can be modified to work for ChIP-seq data.

Supplemental Data File 9. DAP-seq binding data for BAG TFs. These data were obtained using the DAP-seq program and include gene accession numbers and associated primary gene symbols generated using the bulk data retrieval tool in TAIR.

Supplemental Data File 10. GRN input file with annotations

## Notes

**Funding Statement:** This work was supported by the NIH SCORE program 1SC3GM113810 to JAB and the NIH RISE program to WEH 2R25GM071638-09A1

https://www.ncbi.nlm.nih.gov/geo/query/acc.cgi?acc=GSE134177

### Competing Interest Statement

The authors have declared no competing interest.

