## Supplementals for "Gene Expression Changes Occurring at Bolting Time are Associated with Leaf Senescence in Arabidopsis": Supplemental_Data_File_4.docx

***DESeq2: LRT,***

if (!requireNamespace("BiocManager", quietly = TRUE))

install.packages("BiocManager")

##Just TM1, repeat with Tm2,then overlap the DEG lists

library(DESeq2)

#subset just the numerical data (remove the AT numbers)

mydata <- as.data.frame(X200322_T1Counts[c(2:25)])

#add the AT numbers back in as row names

row.names(mydata) <- X200322_T1Counts$target_id

#Round the data

mydata<-round(mydata)

#Define experimental design

T1dds <- DESeqDataSetFromMatrix(countData = mydata,

colData = X200322_T1ES,

design= ~ ~ Genotype + Time+Genotype:Time)

#remove genes with consistently low expression

nrow(T1dds)

keep <- rowSums(counts(T1dds)) > 1

T1dds <- T1dds[keep,]

nrow(T1dds)

#Run the DE analysis

T1dds <- DESeq(T1dds, test="LRT", reduced = ~ Time)

#Return Results, degs different between genotypes

resT1 <- results(T1dds, contrast=c("Genotype","WT","Mutant"))

mcols(resT1, use.names = TRUE)

summary(resT1)

#Subset genes with p<.05 and FDR<.05

T1Sig <- subset(resT1,abs(padj)< 0.05)

write.csv(T1Sig, file = "200325T1DeSEQ2.csv")

##This is identical to the previous chunk of code, just repeated for the second mutant

library(DESeq2)

mydata1 <- as.data.frame(X200322_T2Counts[c(2:25)]) #subset just the numerical data (remove the AT numbers)

row.names(mydata1) <- X200322_T2Counts$target_id #add the AT numbers back in as row names

mydata1<-round(mydata1)

T2dds <- DESeqDataSetFromMatrix(countData = mydata1,

colData = X200322_T2ES,

design= ~ ~ Genotype + Time+Genotype:Time)

nrow(T2dds)

keep <- rowSums(counts(T2dds)) > 1

T2dds <- T2dds[keep,]

nrow(T2dds)

T2dds <- DESeq(T2dds, test="LRT", reduced = ~ Time)

resT2 <- results(T2dds, contrast=c("Genotype","WT","Mutant"))

mcols(resT2, use.names = TRUE)

summary(resT2)

T2Sig <- subset(resT2,abs(padj)< 0.05)

write.csv(T2Sig, file = "200325T1DeSEQ2.csv")

T1AT<-as.data.frame(rownames(T1Sig))

T2AT<-as.data.frame(rownames(T2Sig))

#overlap the DEG lists from T1 and T2

DESeq2DEGs<- as.data.frame(intersect(T1AT$`rownames(T1Sig)`,T2AT$`rownames(T2Sig)`))

write.csv(DESeq2DEGs, file = "200325DESeq2DEGList.csv")

***#edgeR model:***

library(edgeR)

library(limma)

library(RColorBrewer)

library(HTSFilter)

#TM1

#subset just the numerical data (remove the AT numbers)

mydata <- as.data.frame(X200322_T2Counts[c(2:25)])

#add the AT numbers back in as row

row.names(mydata) <- X200322_T2Counts$target_id

#define the groups for design (combo contains genotype and time in one string (WT0,WT2, etc)

group<-X200322_T2ES$Combo

#make matrix

y <- DGEList(counts=(mydata), group=group)

#remove genes with low expression

keep <- filterByExpr(y)

y <- y[keep, , keep.lib.sizes=FALSE]

#calculate the library sizes for normalization

y <- calcNormFactors(y)

#Read to make sure it worked and check the groups look right

y$samples

#Assign the groups to the design matrix, to group which samples will be tested together

design <- model.matrix(~0+group)

#Define what comparisons to make, here I assign WT at time point 0 to mutant at time point 0

my.contrasts <- makeContrasts(

T0 = groupWT0-groupMut0,

T2 = groupWT2-groupMut2,

T4 = groupWT4-groupMut4,

T6 = groupWT6-groupMut6,

levels=design)

#estimate dispersion, I think this finds variance of mean relative to library size per gene?

y <- estimateDisp(y, design)

#combine the design and count matrix data.

fit <- glmQLFit(y, design)

#Run the tests

T1T0 <- glmQLFTest(fit, contrast=my.contrasts[,"T0"])

T1T2 <- glmQLFTest(fit, contrast=my.contrasts[,"T2"])

T1T4 <- glmQLFTest(fit, contrast=my.contrasts[,"T4"])

T1T6 <- glmQLFTest(fit, contrast=my.contrasts[,"T6"])

#isolate transcripts with p<.05 and a logFC greater than 1.5

T1T0Sig<-subset(T1T0$table, PValue<.05)

T1T2Sig<-subset(T1T2$table, PValue<.05)

T1T4Sig<-subset(T1T4$table, PValue<.05)

T1T6Sig<-subset(T1T6$table, PValue<.05)

T1T0SigLog<-subset(T1T0Sig, abs(logFC)>1.5)

T1T2SigLog<-subset(T1T2Sig, abs(logFC)>1.5)

T1T4SigLog<-subset(T1T4Sig, abs(logFC)>1.5)

T1T6SigLog<-subset(T1T6Sig, abs(logFC)>1.5)

#get transcript names

T1T0SigAT<-as.data.frame(rownames(T1T0SigLog))

T1T2SigAT<-as.data.frame(rownames(T1T2SigLog))

T1T4SigAT<-as.data.frame(rownames(T1T4SigLog))

T1T6SigAT<-as.data.frame(rownames(T1T6SigLog))

write.csv(T1T0SigAT, file = "edgeRT2-0Sig.csv")

write.csv(T1T2SigAT, file = "edgeRT2-2Sig.csv")

write.csv(T1T4SigAT, file = "edgeRT2-4Sig.csv")

write.csv(T1T6SigAT, file = "edgeRT2-6Sig.csv")

I repeated this identical code with TM2, then reported the intersection between TM1 vs. WT and TM2 vs. WT.

T-Test based approach:

library (vegan)

library (Hmisc)

library (ggplot2)

library(reshape)

library(reshape2)

library(splitstackshape)

library(plyr)

library(gdata)

ncol(FPKMafterfilter1) #how many columns (should be 36, but 37 with labels)

mydata <- as.data.frame(FPKMafterfilter1[c(2:37)]) #subset just the numerical data (remove the AT numbers)

row.names(mydata) <- FPKMafterfilter1$X1 #add the AT numbers back in as row names

#everything below separates out samples by replicates

W0 <- as.data.frame(mydata[c(1:3)])

W2 <- as.data.frame(mydata[c(4:6)])

W4 <- as.data.frame(mydata[c(7:9)])

W6 <- as.data.frame(mydata[c(10:12)])

T10 <- as.data.frame(mydata[c(13:15)])

T12 <- as.data.frame(mydata[c(16:18)])

T14 <- as.data.frame(mydata[c(19:21)])

T16 <- as.data.frame(mydata[c(22:24)])

T20 <- as.data.frame(mydata[c(25:27)])

T22 <- as.data.frame(mydata[c(28:30)])

T24 <- as.data.frame(mydata[c(31:33)])

T26 <- as.data.frame(mydata[c(34:36)])

#get the average of three replicates in each file

W0mean <- as.data.frame(rowMeans(W0[,1:3]))

W2mean <- as.data.frame(rowMeans(W2[,1:3]))

W4mean <- as.data.frame(rowMeans(W4[,1:3]))

W6mean <- as.data.frame(rowMeans(W6[,1:3]))

T10mean <- as.data.frame(rowMeans(T10[,1:3]))

T12mean <- as.data.frame(rowMeans(T12[,1:3]))

T14mean <- as.data.frame(rowMeans(T14[,1:3]))

T16mean <- as.data.frame(rowMeans(T16[,1:3]))

T20mean <- as.data.frame(rowMeans(T20[,1:3]))

T22mean <- as.data.frame(rowMeans(T22[,1:3]))

T24mean <- as.data.frame(rowMeans(T24[,1:3]))

T26mean <- as.data.frame(rowMeans(T26[,1:3]))

#calculate fold change for TM over WT.

T10Fold1<- as.data.frame((T10mean/W0mean))

T20Fold1<- as.data.frame((T20mean/W0mean))

T12Fold1<- as.data.frame((T12mean/W2mean))

T22Fold1<- as.data.frame((T22mean/W2mean))

T14Fold1<- as.data.frame((T14mean/W4mean))

T24Fold1<- as.data.frame((T24mean/W4mean))

T16Fold1<- as.data.frame((T16mean/W6mean))

T26Fold1<- as.data.frame((T26mean/W6mean))

#rename the columns to say "Fold"

names(T10Fold1)[1] <- "Fold"

names(T20Fold1)[1] <- "Fold"

names(T12Fold1)[1] <- "Fold"

names(T22Fold1)[1] <- "Fold"

names(T14Fold1)[1] <- "Fold"

names(T24Fold1)[1] <- "Fold"

names(T16Fold1)[1] <- "Fold"

names(T26Fold1)[1] <- "Fold"

#Creating files for sig data to be put into

test2 <- NULL

test4<- NULL

#Overall ttest: sta is needed for directionality, so the data can be subset to up or downregulated for fold change calcs

#T0, WT vs TM1 and TM2

for (j in seq(nrow(W0)))

{

ID_REF<-rownames(W0[j,]) #to extract the probe id

sta1<-t.test(W0[j,], T10[j,])$statistic #to run the t.test and extract the $statistic (i.e., the difference in the mean vale)

sig1<-t.test(W0[j,], T10[j,])$p.value #to (re)run the t.test and extract the significance value (i.e., p-value)

test1<-cbind(ID_REF,sta1,sig1)	# this juxtaposing sta and sig

test2<-rbind(test2,test1) # to same sta and sig

ID_REF<-rownames(W0[j,]) #to extract the probe id

sta2<-t.test(W0[j,], T20[j,])$statistic #to run the t.test and extract the $statistic (i.e., the difference in the mean vale)

sig2<-t.test(W0[j,], T20[j,])$p.value #to (re)run the t.test and extract the significance value (i.e., p-value)

test3<-cbind(ID_REF,sta2,sig2)	# this juxtaposing sta and sig

test4<-rbind(test4,test3) # to same sta and sig

}

#Add Fold Change Data into it

head(test2)

test2<- as.data.frame(test2)

test2$Fold<- T10Fold1

#make a numeric dataset

test2$sta1<-as.numeric(as.character(test2$sta1))

test2$sig1<-as.numeric(as.character(test2$sig1))

#isoalte the significant results with pvalue of .05 and a 2 fold change

sig_wt_T1_0 <- subset(test2,abs(sig1)<.05)

sig2foldT101 <- subset(sig_wt_T1_0,abs(Fold)>2)

sig2foldT102 <- subset(sig_wt_T1_0,abs(Fold)<0.5)

T1_0_Sig <- combine(sig2foldT101,sig2foldT102)

#remove excess files so they dont stack up

rm(sig_wt_T1_0)

rm(sig2foldT101)

rm(sig2foldT102)

rm(sta1)

rm(sig1)

rm(test1)

rm(test2)

#repeat for TM2

test4<- as.data.frame(test4)

test4$Fold<- T20Fold1

#make a numeric dataset

test4$sta2<-as.numeric(as.character(test4$sta2))

test4$sig2<-as.numeric(as.character(test4$sig2))

#isoalte the significant results with a bonferonni correction., .05/19437=.00000257

sig_wt_T2_0 <-subset(test4,abs(sig2)<.05)

sig2foldT201 <-subset(sig_wt_T2_0,abs(Fold)>2)

sig2foldT202 <- subset(sig_wt_T2_0,abs(Fold)<0.5)

T2_0_Sig <- combine(sig2foldT201,sig2foldT202)

rm(sig_wt_T2_0)

rm(sig2foldT201)

rm(sig2foldT202)

rm(sta2)

rm(sig2)

rm(test3)

rm(test4)

#start for the next time point

test6 <- NULL

test8<- NULL

#T2, WT vs TM1 and TM2

for (j in seq(nrow(W2)))

{

ID_REF<-rownames(W2[j,]) #to extract the probe id

sta3<-t.test(W2[j,], T12[j,])$statistic #to run the t.test and extract the $statistic (i.e., the difference in the mean vale)

sig3<-t.test(W2[j,], T12[j,])$p.value #to (re)run the t.test and extract the significance value (i.e., p-value)

test5<-cbind(ID_REF,sta3,sig3)	# this juxtaposing sta and sig

test6<-rbind(test6,test5) # to same sta and sig

ID_REF<-rownames(W2[j,]) #to extract the probe id

sta4<-t.test(W2[j,], T22[j,])$statistic #to run the t.test and extract the $statistic (i.e., the difference in the mean vale)

sig4<-t.test(W2[j,], T22[j,])$p.value #to (re)run the t.test and extract the significance value (i.e., p-value)

test7<-cbind(ID_REF,sta4,sig4)	# this juxtaposing sta and sig

test8<-rbind(test8,test7) # to same sta and sig

}

test6<- as.data.frame(test6)

test6$Fold<- T12Fold1

#make a numeric dataset

test6$sta3<-as.numeric(as.character(test6$sta3))

test6$sig3<-as.numeric(as.character(test6$sig3))

sig_wt_T1_2 <-subset(test6,abs(sig3)<.05)

sig2foldT121 <-subset(sig_wt_T1_2,abs(Fold)>2)

sig2foldT122 <- subset(sig_wt_T1_2,abs(Fold)<0.5)

T1_2_Sig <- combine(sig2foldT121,sig2foldT122)

test8<- as.data.frame(test8)

test8$Fold<- T22Fold1

#make a numeric dataset

test8$sta4<-as.numeric(as.character(test8$sta4))

test8$sig4<-as.numeric(as.character(test8$sig4))

sig_wt_T2_2 <-subset(test8,abs(sig4)<.05)

sig2foldT221 <-subset(sig_wt_T2_2,abs(Fold)>2)

sig2foldT222 <- subset(sig_wt_T2_2,abs(Fold)<0.5)

T2_2_Sig <- combine(sig2foldT221,sig2foldT222)

rm(sig_wt_T2_2)

rm(sig2foldT121)

rm(sig2foldT122)

rm(sig2foldT221)

rm(sig2foldT222)

rm(sta3)

rm(sig3)

rm(test5)

rm(test6)

rm(test7)

rm(test8)

#repeat for the third time point

test10= NULL

test12= NULL

#WT vs TM1 and Tm2 at T4

for (j in seq(nrow(W4)))

{

ID_REF<-rownames(W4[j,]) #to extract the probe id

sta5<-t.test(W4[j,], T14[j,])$statistic #to run the t.test and extract the $statistic (i.e., the difference in the mean vale)

sig5<-t.test(W4[j,], T14[j,])$p.value #to (re)run the t.test and extract the significance value (i.e., p-value)

test9<-cbind(ID_REF,sta5,sig5)	# this juxtaposing sta and sig

test10<-rbind(test10,test9) # to same sta and sig

ID_REF<-rownames(W4[j,]) #to extract the probe id

sta6<-t.test(W4[j,], T24[j,])$statistic #to run the t.test and extract the $statistic (i.e., the difference in the mean vale)

sig6<-t.test(W4[j,], T24[j,])$p.value #to (re)run the t.test and extract the significance value (i.e., p-value)

test11<-cbind(ID_REF,sta6,sig6)	# this juxtaposing sta and sig

test12<-rbind(test12,test11) # to same sta and sig

}

test10<- as.data.frame(test10)

test10$Fold<- T14Fold1

#make a numeric dataset

test10$sta5<-as.numeric(as.character(test10$sta5))

test10$sig5<-as.numeric(as.character(test10$sig5))

sig_wt_T1_4 <-subset(test10,abs(sig5)<.05)

sig2foldT141 <-subset(sig_wt_T1_4,abs(Fold)>2)

sig2foldT142 <- subset(sig_wt_T1_4,abs(Fold)<0.5)

T1_4_Sig <- combine(sig2foldT141,sig2foldT142)

test12<- as.data.frame(test12)

test12$Fold<- T24Fold1

#make a numeric dataset

test12$sta6<-as.numeric(as.character(test12$sta6))

test12$sig6<-as.numeric(as.character(test12$sig6))

sig_wt_T2_4 <-subset(test12,abs(sig6)<.05)

sig2foldT241<-subset(sig_wt_T2_4,abs(Fold)>2)

sig2foldT242 <- subset(sig_wt_T2_4,abs(Fold)<0.5)

T2_4_Sig <- combine(sig2foldT241,sig2foldT242)

rm(sta4)

rm(sig4)

rm(sta5)

rm(sig5)

rm(sta6)

rm(sig6)

rm(sig_wt_T1_4)

rm(sig_wt_T2_4)

rm(sig2foldT141)

rm(sig2foldT142)

rm(sig2foldT241)

rm(sig2foldT242)

rm(sta3)

rm(sig3)

rm(test9)

rm(test10)

rm(test11)

rm(test12)

test14= NULL

test16= NULL

#WT vs TM1 and Tm2 at T6

for (j in seq(nrow(W6)))

{

ID_REF<-rownames(W6[j,]) #to extract the probe id

sta7<-t.test(W6[j,], T16[j,])$statistic #to run the t.test and extract the $statistic (i.e., the difference in the mean vale)

sig7<-t.test(W6[j,], T16[j,])$p.value #to (re)run the t.test and extract the significance value (i.e., p-value)

test13<-cbind(ID_REF,sta7,sig7)	# this juxtaposing sta and sig

test14<-rbind(test14,test13) # to same sta and sig

ID_REF<-rownames(W6[j,]) #to extract the probe id

sta8<-t.test(W6[j,], T26[j,])$statistic #to run the t.test and extract the $statistic (i.e., the difference in the mean vale)

sig8<-t.test(W6[j,], T26[j,])$p.value #to (re)run the t.test and extract the significance value (i.e., p-value)

test15<-cbind(ID_REF,sta8,sig8)	# this juxtaposing sta and sig

test16<-rbind(test16,test15) # to same sta and sig

}

test14<- as.data.frame(test14)

test14$Fold<- T16Fold1

#make a numeric dataset

test14$sta7<-as.numeric(as.character(test14$sta7))

test14$sig7<-as.numeric(as.character(test14$sig7))

sig_wt_T1_6 <-subset(test14,abs(sig7)<.05)

sig2foldT161 <-subset(sig_wt_T1_6,abs(Fold)>2)

sig2foldT162 <- subset(sig_wt_T1_6,abs(Fold)<0.5)

T1_6_Sig <- combine(sig2foldT161,sig2foldT162)

test16<- as.data.frame(test16)

test16$Fold<- T26Fold1

#make a numeric dataset

test16$sta8<-as.numeric(as.character(test16$sta8))

test16$sig8<-as.numeric(as.character(test16$sig8))

sig_wt_T2_6 <-subset(test16,abs(sig8)<.05)

sig2foldT261 <-subset(sig_wt_T2_6,abs(Fold)>2)

sig2foldT262 <- subset(sig_wt_T2_6,abs(Fold)<0.5)

T2_6_Sig <- combine(sig2foldT261,sig2foldT262)

rm(sta7)

rm(sig7)

rm(sta8)

rm(sig8)

rm(sig_wt_T1_6)

rm(sig_wt_T2_6)

rm(sig_wt_T1_2)

rm(sig2foldT161)

rm(sig2foldT162)

rm(sig2foldT261)

rm(sig2foldT262)

rm(test13)

rm(test14)

rm(test15)

rm(test16)

#Overlap the Datasets (Tm1 and Tm2)

T0DEGs <- merge(T1_0_Sig, T2_0_Sig, by = "ID_REF")

T2DEGs <- merge(T1_2_Sig, T2_2_Sig, by = "ID_REF")

T4DEGs <- merge(T1_4_Sig, T2_4_Sig, by = "ID_REF")

T6DEGs <- merge(T1_6_Sig, T2_6_Sig, by = "ID_REF")

#Put all DEGs AT numbers onto single files

T0AT <-as.data.frame(T0DEGs$ID_REF)

T2AT <-as.data.frame(T2DEGs$ID_REF)

T4AT <-as.data.frame(T4DEGs$ID_REF)

T6AT <-as.data.frame(T6DEGs$ID_REF)

#Remove all of the extra files!

rm(T10Fold1)

rm(T20Fold1)

rm(T12Fold1)

rm(T22Fold1)

rm(T14Fold1)

rm(T16Fold1)

rm(T24Fold1)

rm(T26Fold1)

rm(W0)

rm(W2)

rm(W4)

rm(W6)

rm(T10)

rm(T12)

rm(T14)

rm(T16)

rm(T20)

rm(T22)

rm(T24)

rm(T26)

rm(W0mean)

rm(W2mean)

rm(W4mean)

rm(W6mean)

rm(T10mean)

rm(T12mean)

rm(T14mean)

rm(T16mean)

rm(T20mean)

rm(T22mean)

rm(T24mean)

rm(T26mean)

rm(T1_0_Sig)

rm(T2_0_Sig)

rm(T1_2_Sig)

rm(T2_2_Sig)

rm(T1_4_Sig)

rm(T2_4_Sig)

rm(T1_6_Sig)

rm(T2_6_Sig)

rm(mydata)

RESULTS:
