## Supplementals for "Gene Expression Changes Occurring at Bolting Time are Associated with Leaf Senescence in Arabidopsis": Supplemental_Data_File_6.pdf

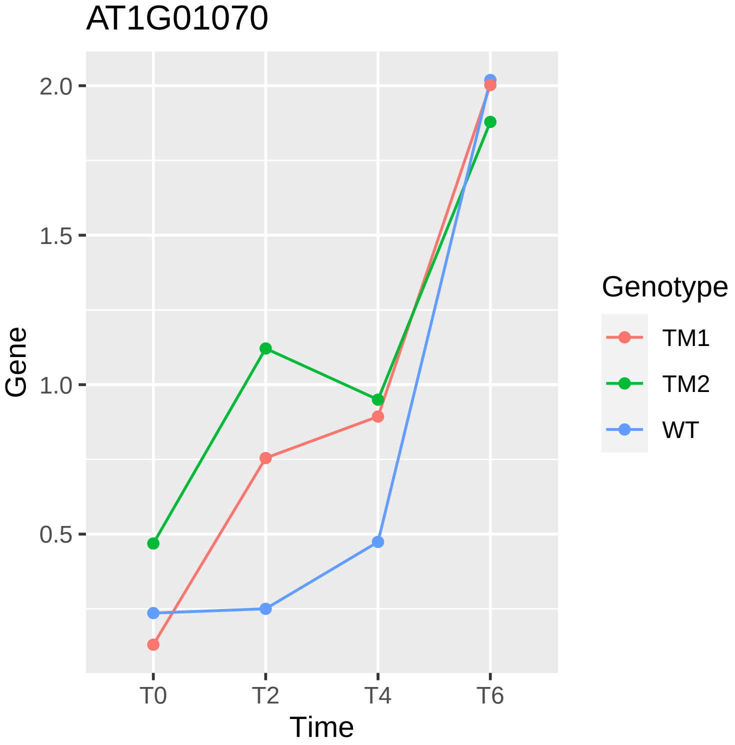
AT1G01070
2.0
1.5
1.0
0.5
Gene
T0
T2
T4
T6
Time
Genotype
TM1
TM2
WT

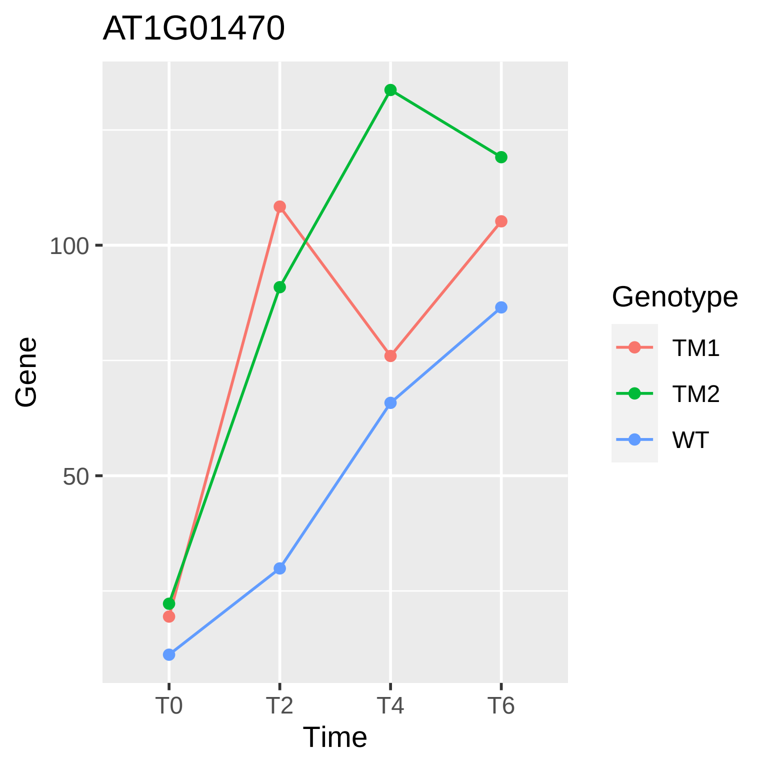
AT1G01470
Gene
100
50
T0
T2
T4
T6
Time
Genotype
TM1
TM2
WT

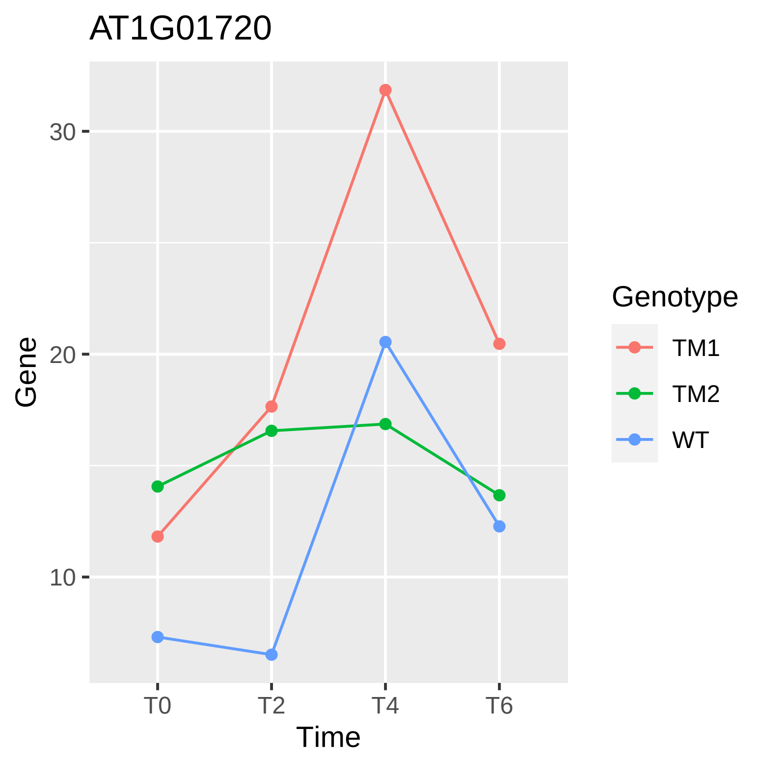
AT1G01720
30
20
10
Gene
T0
T2
T4
T6
Time
Genotype
TM1
TM2
WT

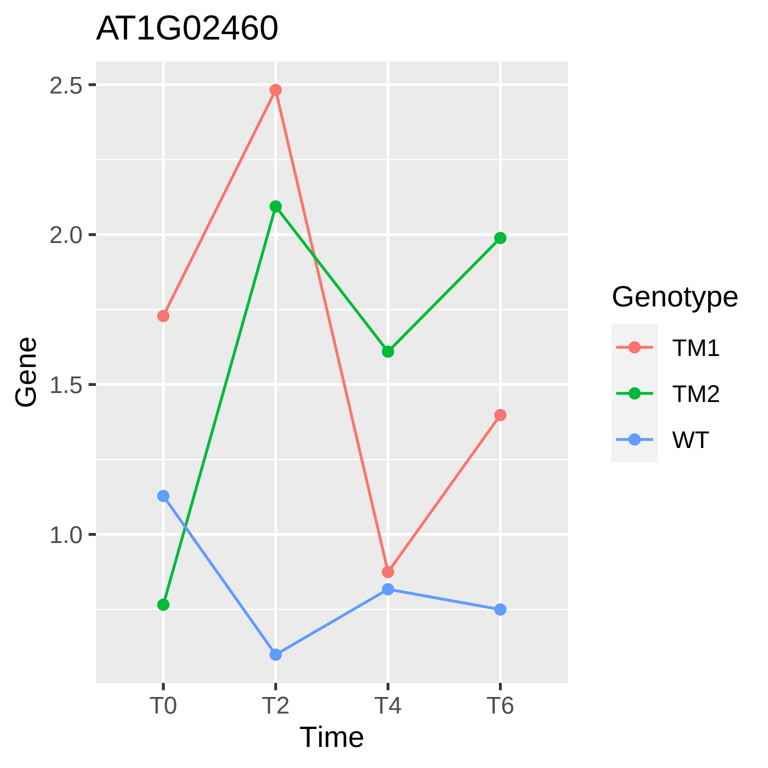
AT1G02460
Gene
2.5
2.0
1.5
1.0
T0
T2
T4
T6
Time
Genotype
TM1
TM2
WT

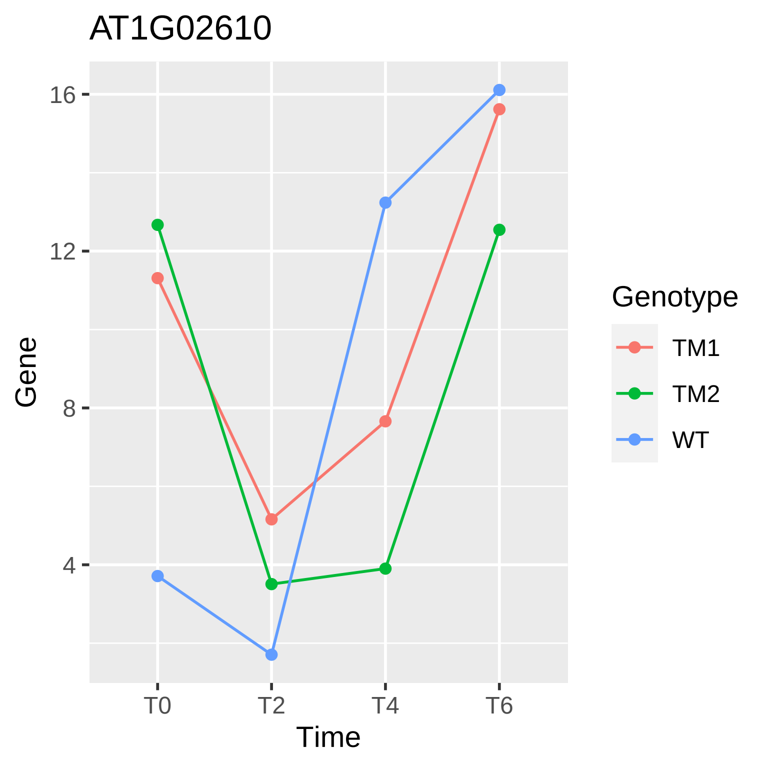
AT1G02610
Gene
16
12
8
4
T0
T2
T4
T6
Time
Genotype
TM1
TM2
WT

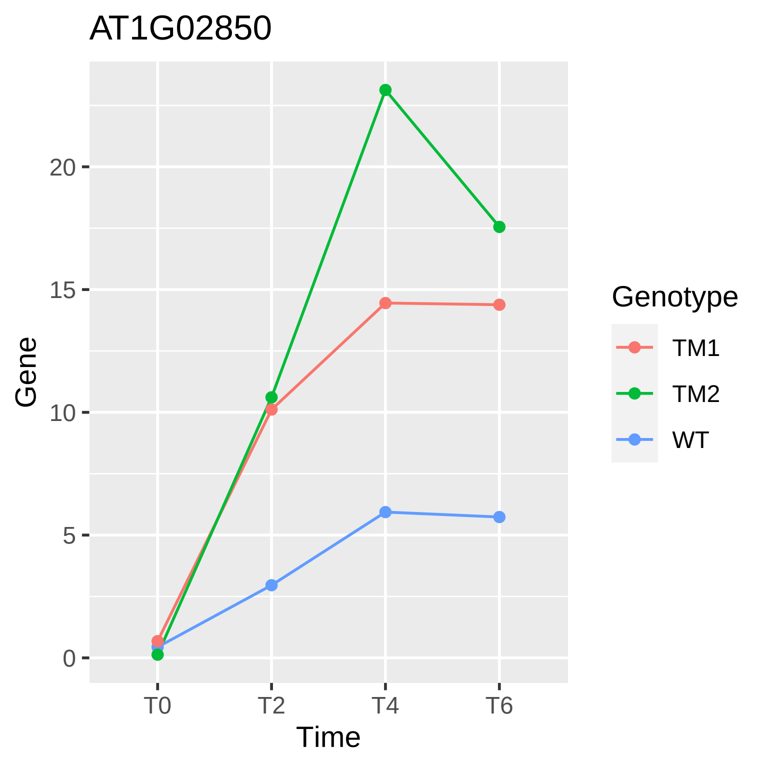
AT1G02850
Gene
20
15
10
5
0
T0
T2
T4
T6
Time
Genotype
TM1
TM2
WT

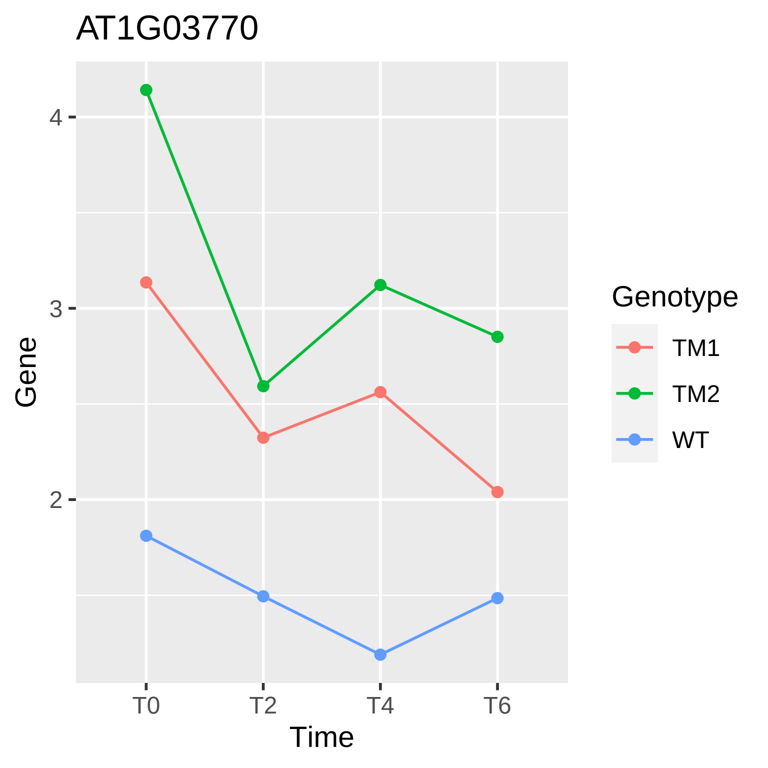
AT1G03770
Gene
4
3
2
T0
T2
T4
T6
Time
Genotype
TM1
TM2
WT

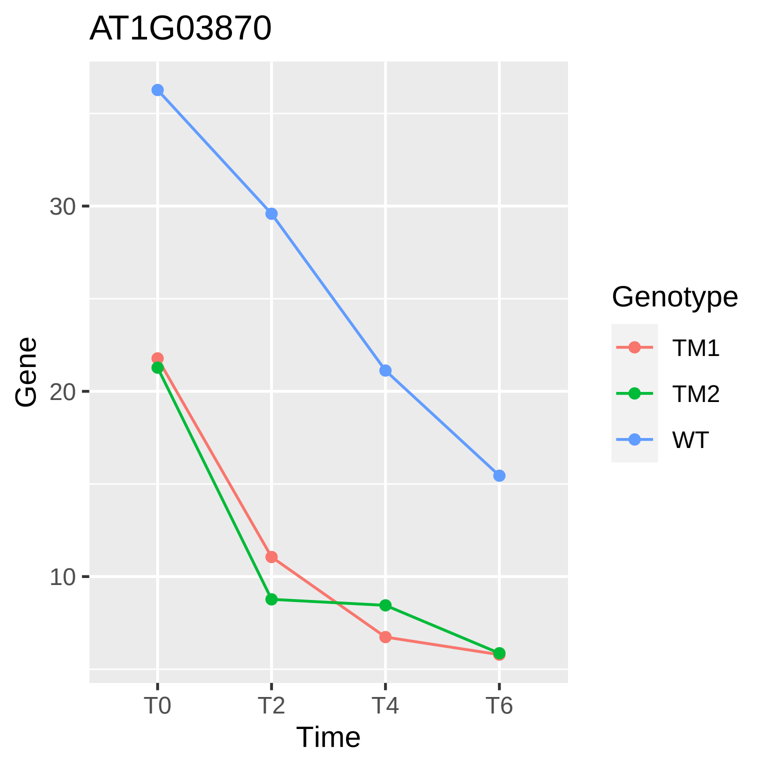
AT1G03870
Gene
30
20
10
T0
T2
T4
T6
Time
Genotype
TM1
TM2
WT

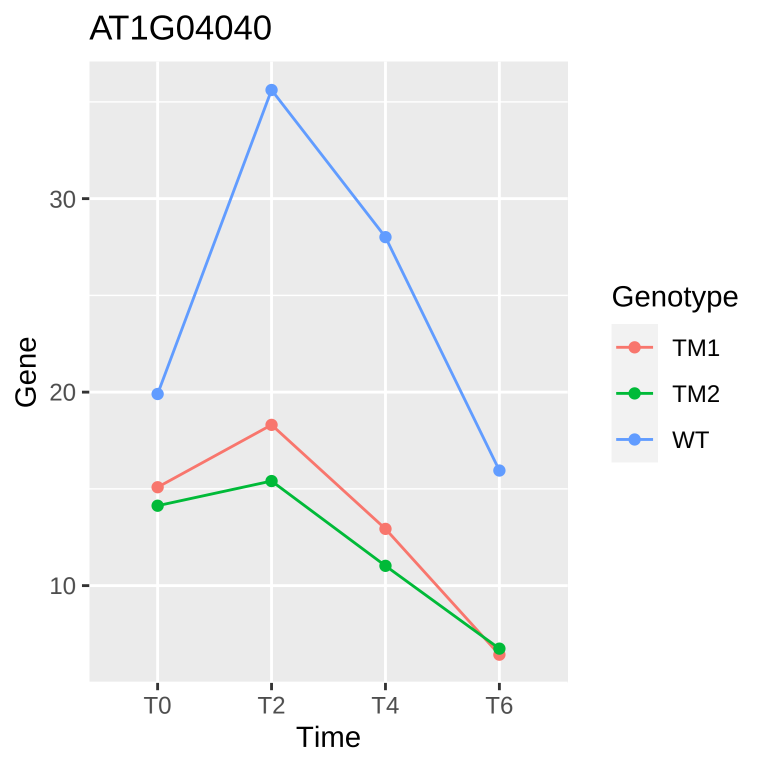
AT1G04040
Gene
30
20
10
T0
T2
T4
T6
Time
Genotype
TM1
TM2
WT

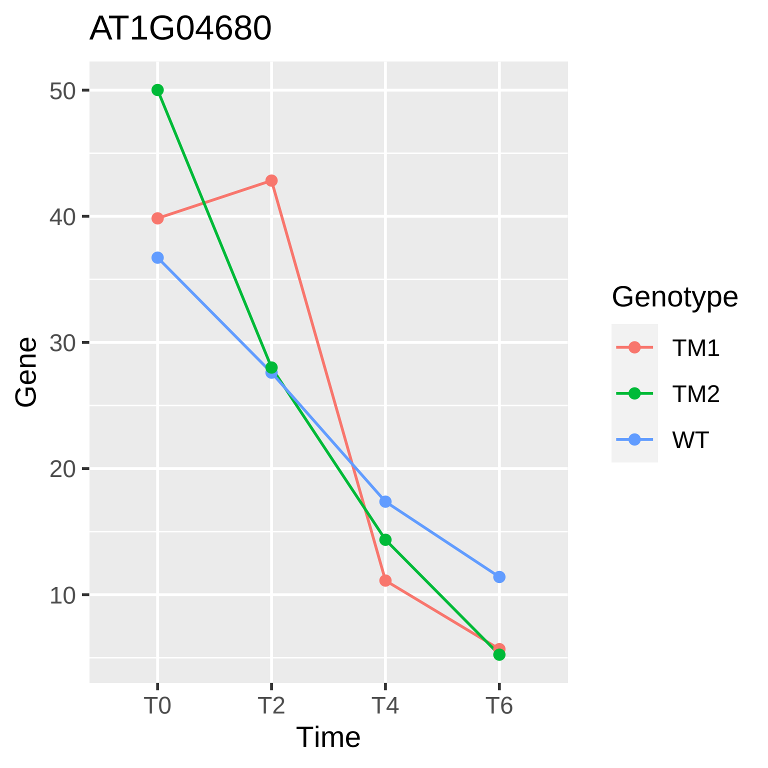
AT1G04680
Gene
50
40
30
20
10
T0
T2
T4
T6
Time
Genotype
TM1
TM2
WT

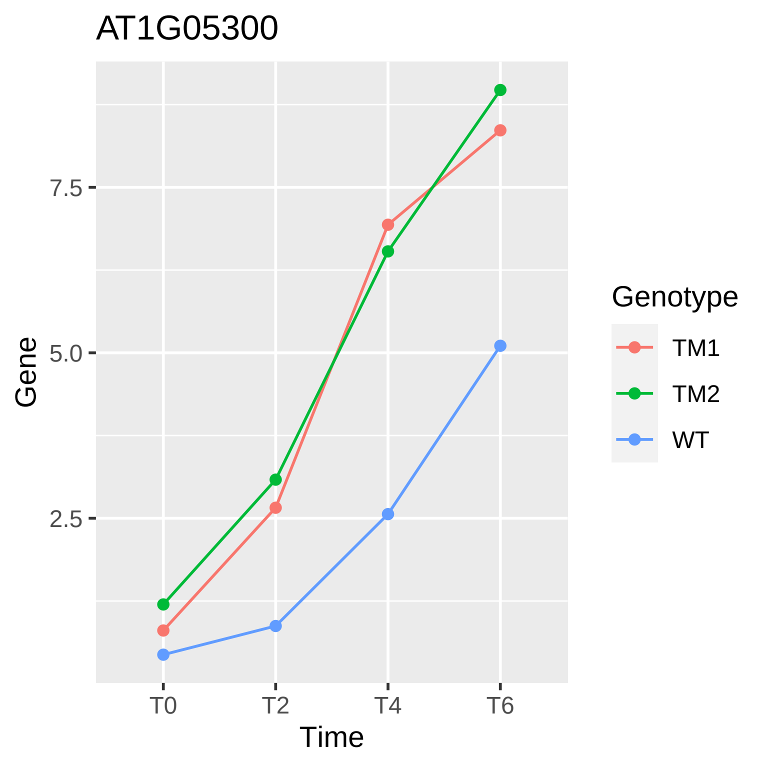

AT1G05300
Gene
7.5
5.0
2.5
T0
T2
T4
T6
Time
Genotype
TM1
TM2
WT

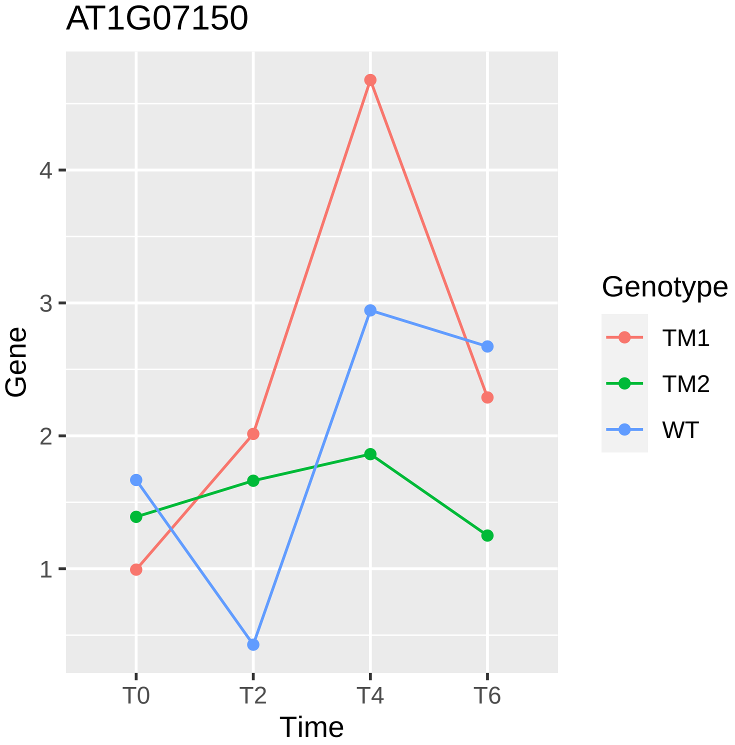
AT1G07150
Gene
4
3
2
1
T0
T2
T4
T6
Time
Genotype
TM1
TM2
WT

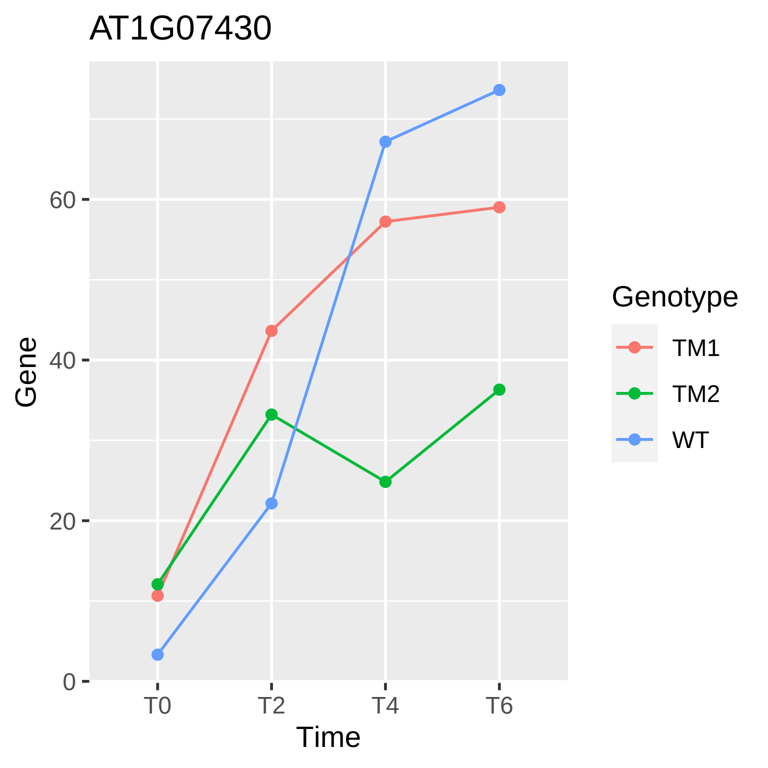
AT1G07430
Gene
60
40
20
0
T0
T2
T4
T6
Time
Genotype
TM1
TM2
WT

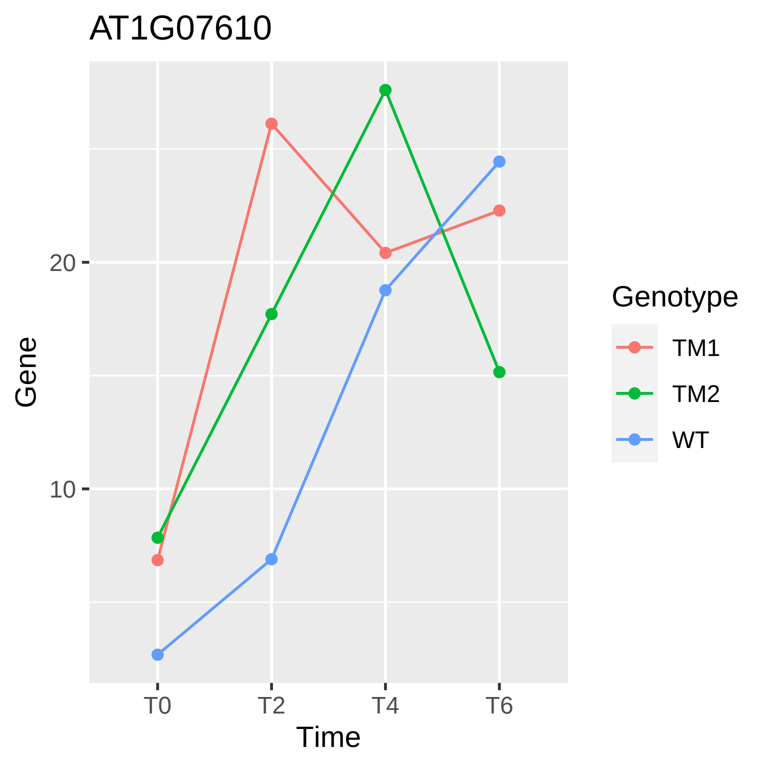
AT1G07610
20
10
Gene
T0
T2
T4
T6
Time
Genotype
TM1
TM2
WT

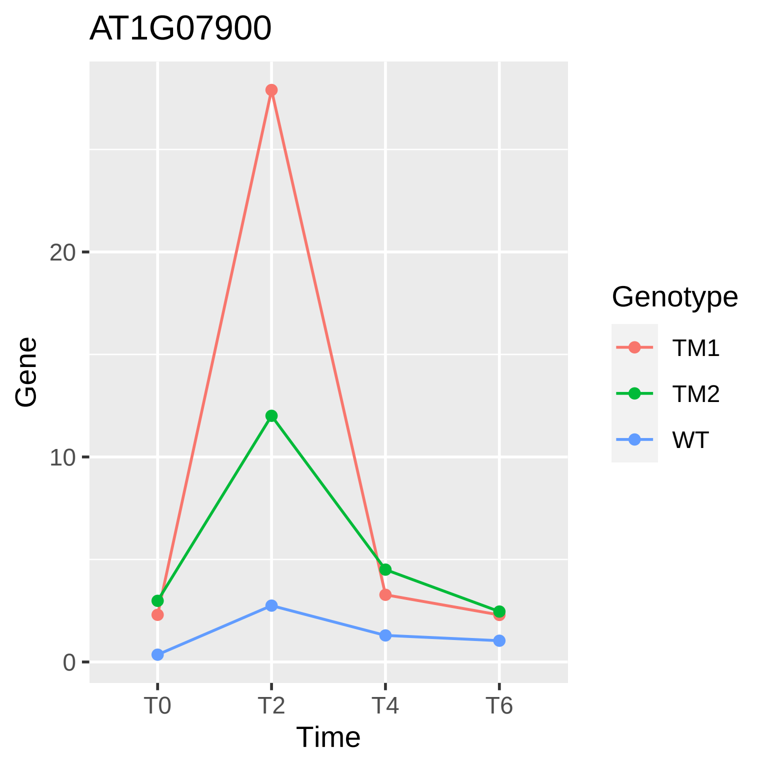

AT1G07900
Gene
20
10
0
T0
T2
T4
T6
Time
Genotype
TM1
TM2
WT

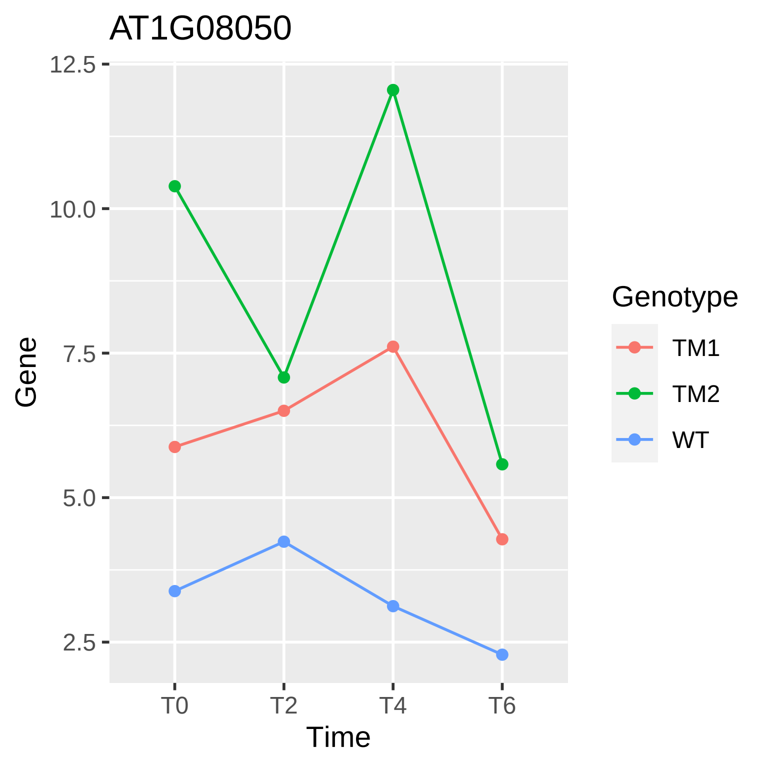

AT1G08050
12.5
10.0
7.5
5.0
2.5
Gene
T0
T2
T4
T6
Time
Genotype
TM1
TM2
WT

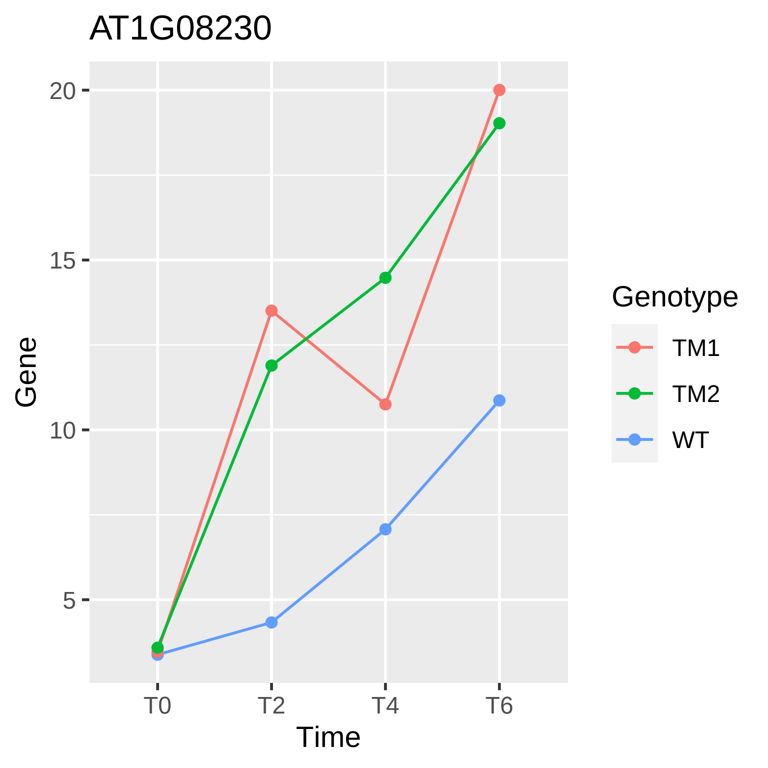
AT1G08230
Gene
20
15
10
5
T0
T2
T4
T6
Time
Genotype
TM1
TM2
WT

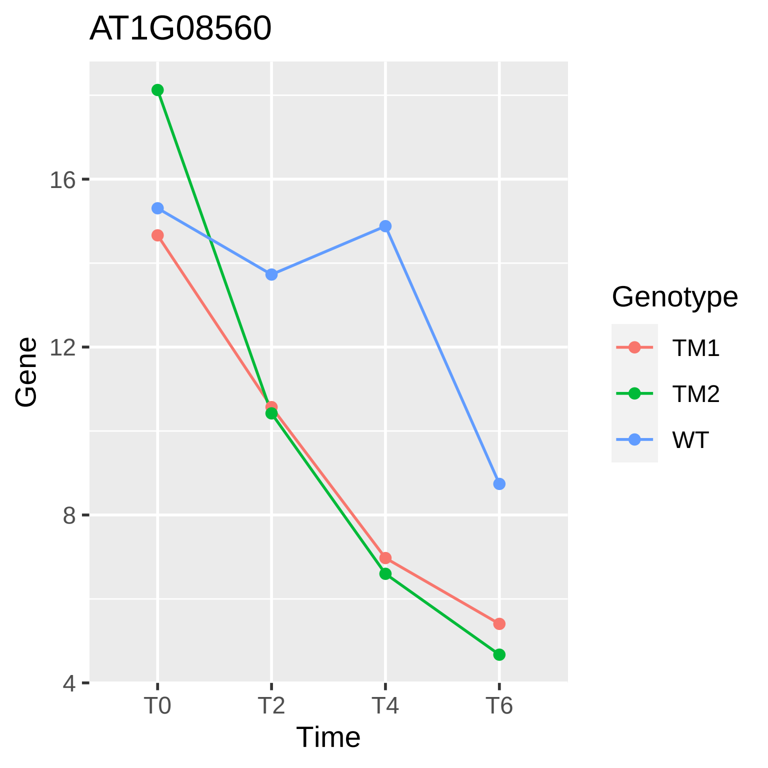

AT1G08560
Gene
16
12
8
4
T0
T2
T4
T6
Time
Genotype
TM1
TM2
WT

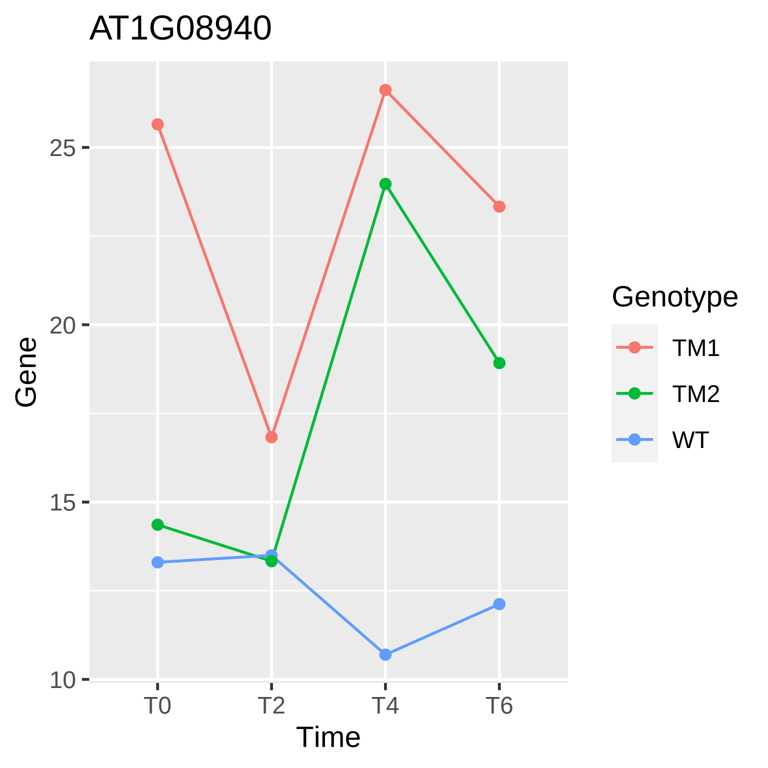

AT1G08940
Gene
25
20
15
10
T0
T2
T4
T6
Time
Genotype
TM1
TM2
WT

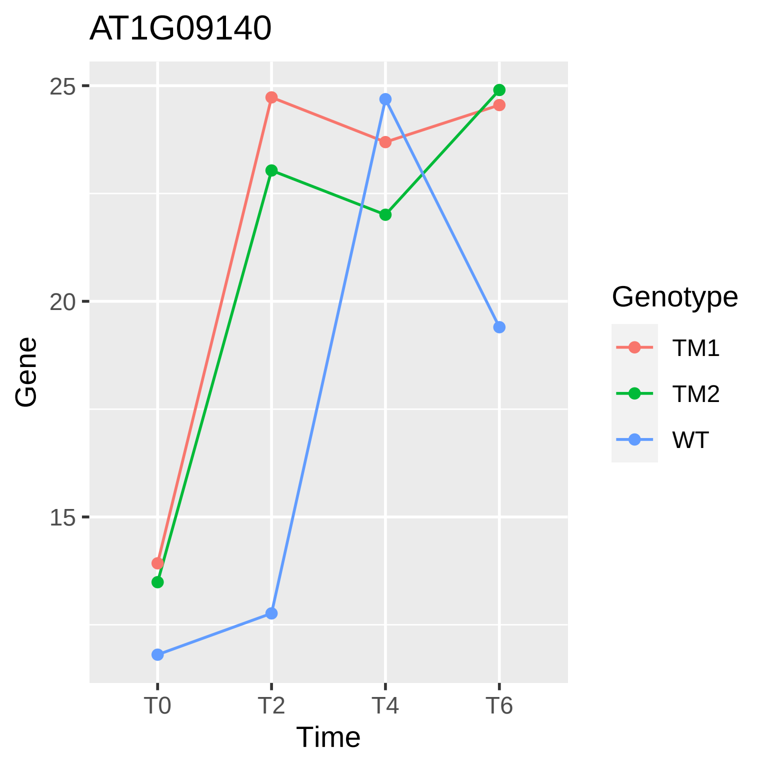

AT1G09140
25
20
15
Gene
T0
T2
T4
T6
Time
Genotype
TM1
TM2
WT

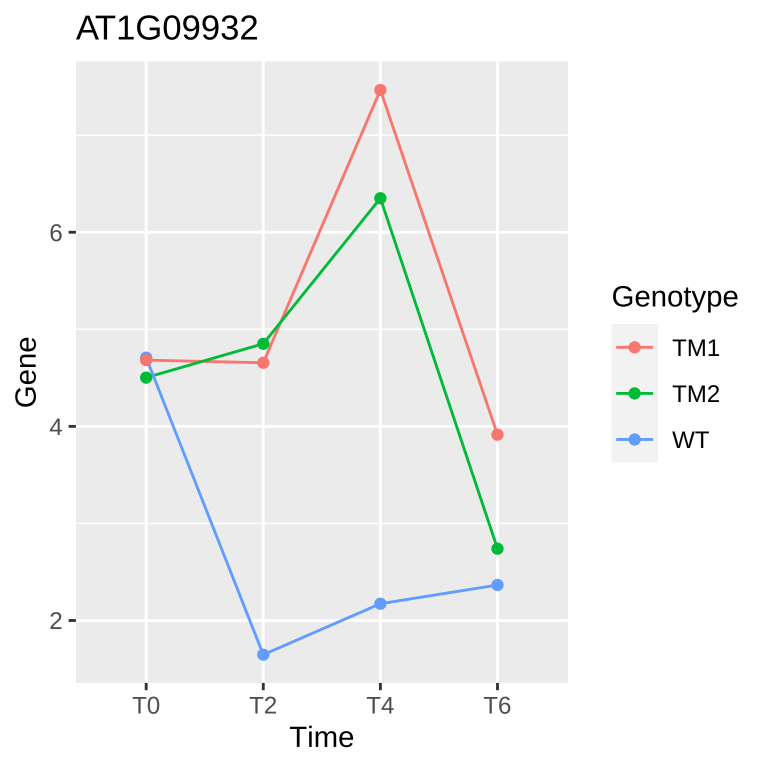
AT1G09932
Gene
6
4
2
T0
T2
T4
T6
Time
Genotype
TM1
TM2
WT

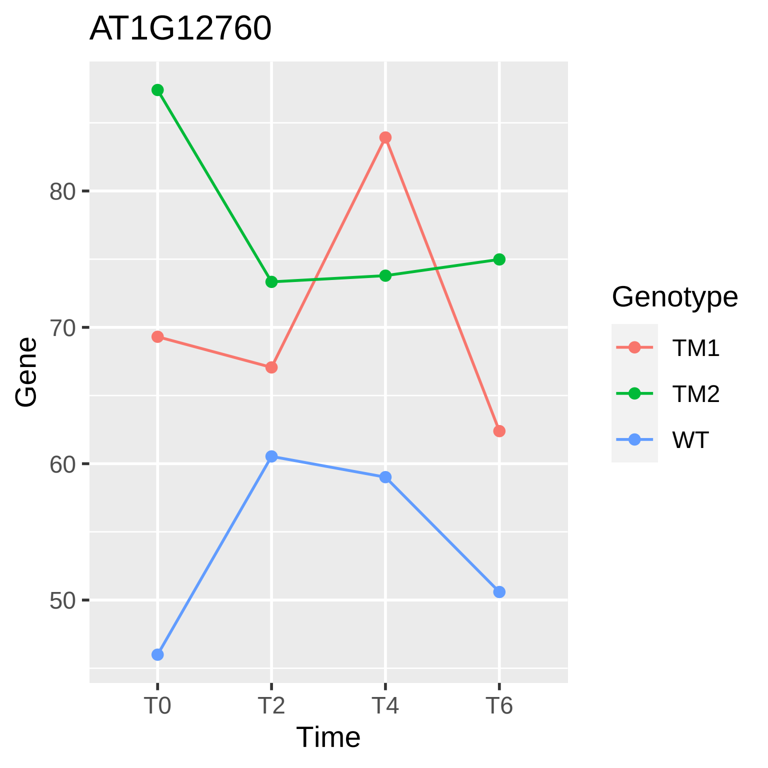

AT1G12760
80
70
60
50
Gene
T0
T2
T4
T6
Time
Genotype
TM1
TM2
WT

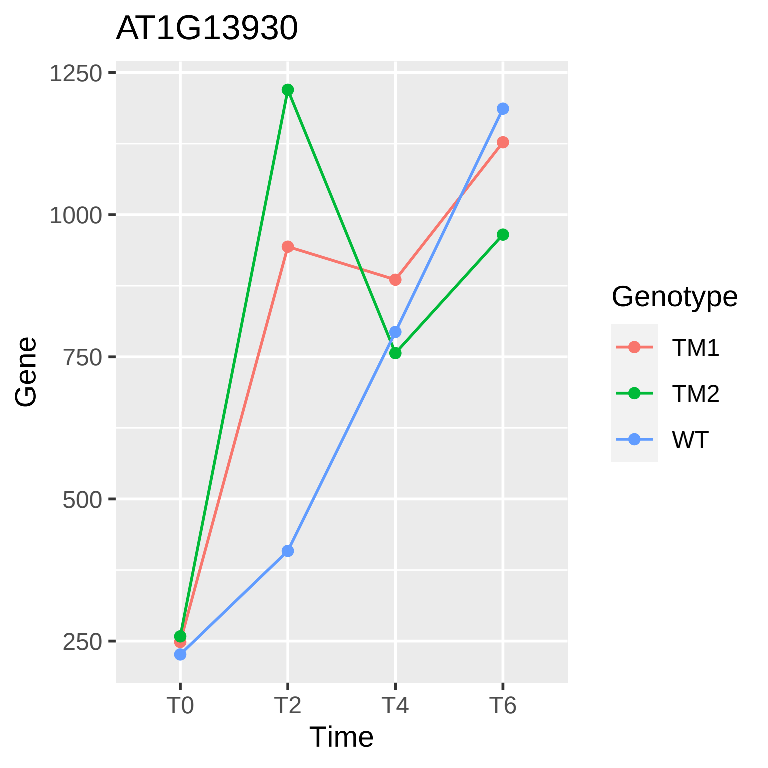
AT1G13930
Gene
1250
1000
750
500
250
T0
T2
T4
T6
Time
Genotype
TM1
TM2
WT

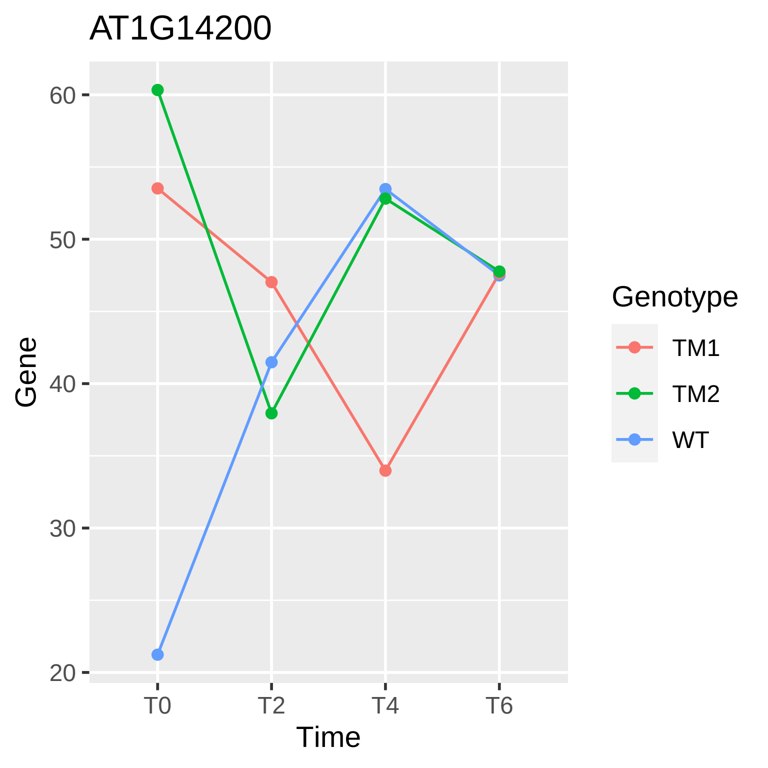
AT1G14200
60
50
40
30
20
Gene
T0
T2
T4
T6
Time
Genotype
TM1
TM2
WT

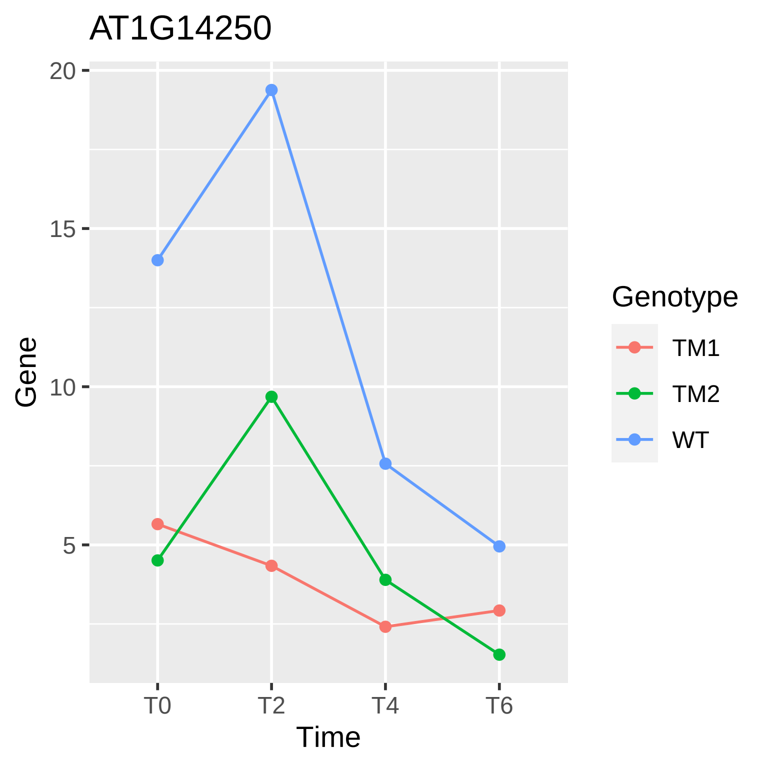
AT1G14250
20
15
10
5
Gene
T0
T2
T4
T6
Time
Genotype
TM1
TM2
WT

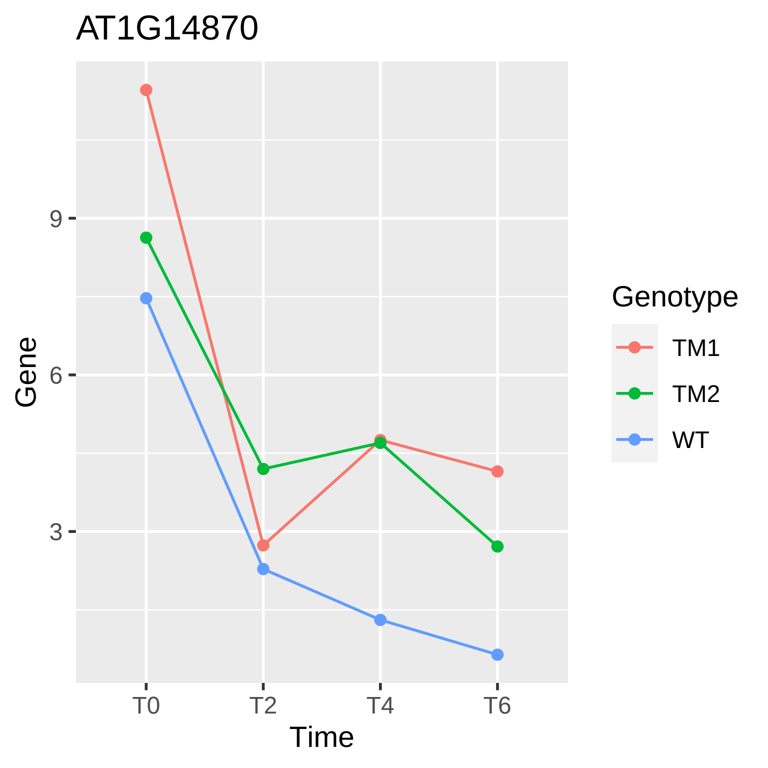

AT1G14870
Gene
9
6
3
T0
T2
T4
T6
Time
Genotype
TM1
TM2
WT

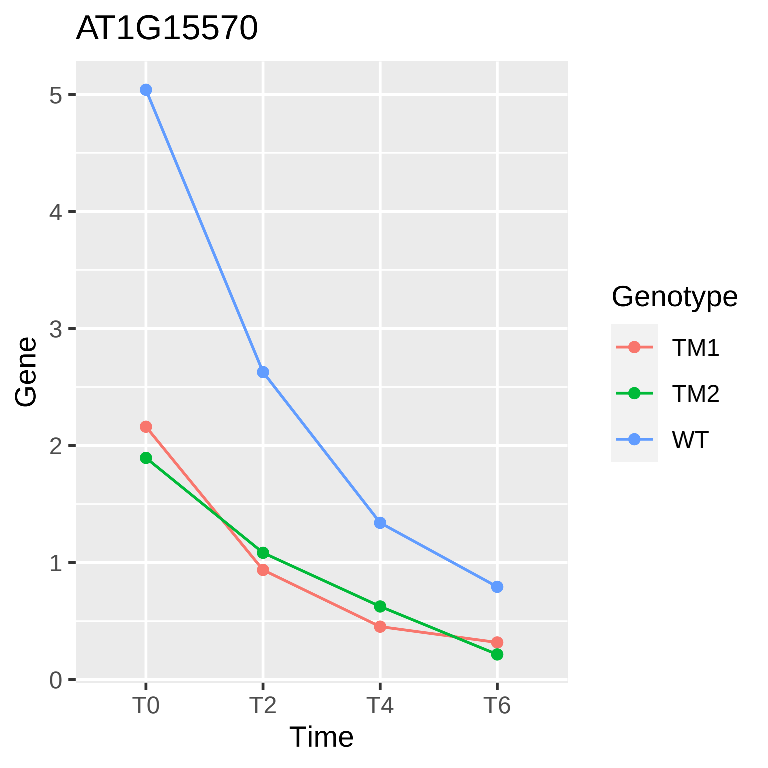
AT1G15570
Gene
5
4
3
2
1
0
T0
T2
T4
T6
Time
Genotype
TM1
TM2
WT

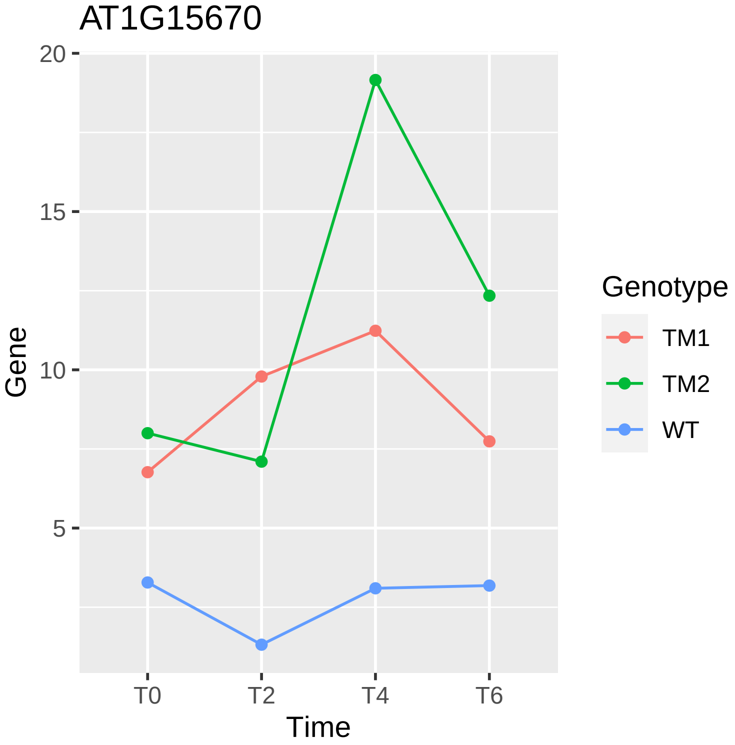
AT1G15670
20
15
10
5
Gene
T0
T2
T4
T6
Time
Genotype
TM1
TM2
WT

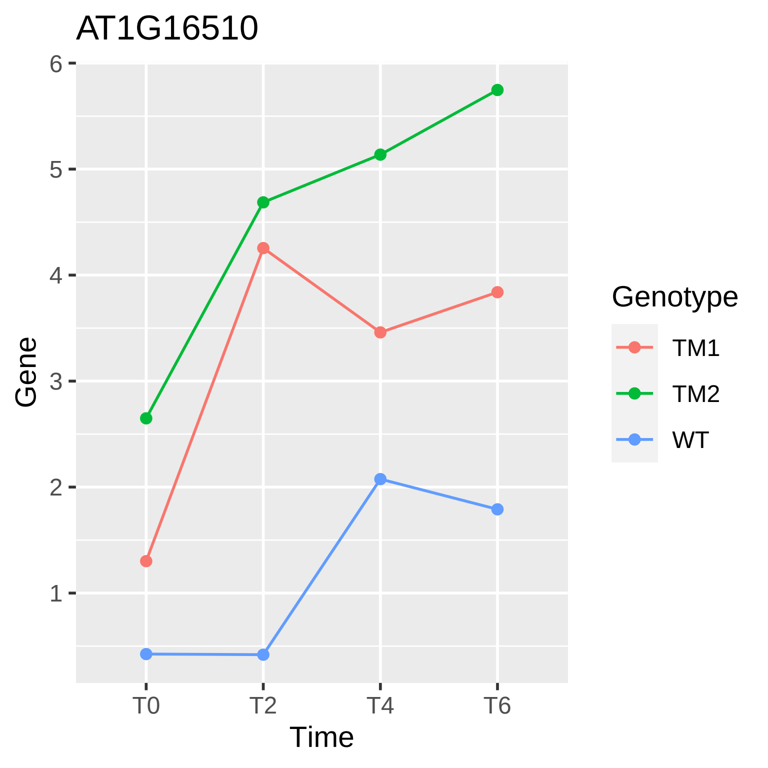
AT1G16510
Gene
6
5
4
3
2
1
T0
T2
T4
T6
Time
Genotype
TM1
TM2
WT

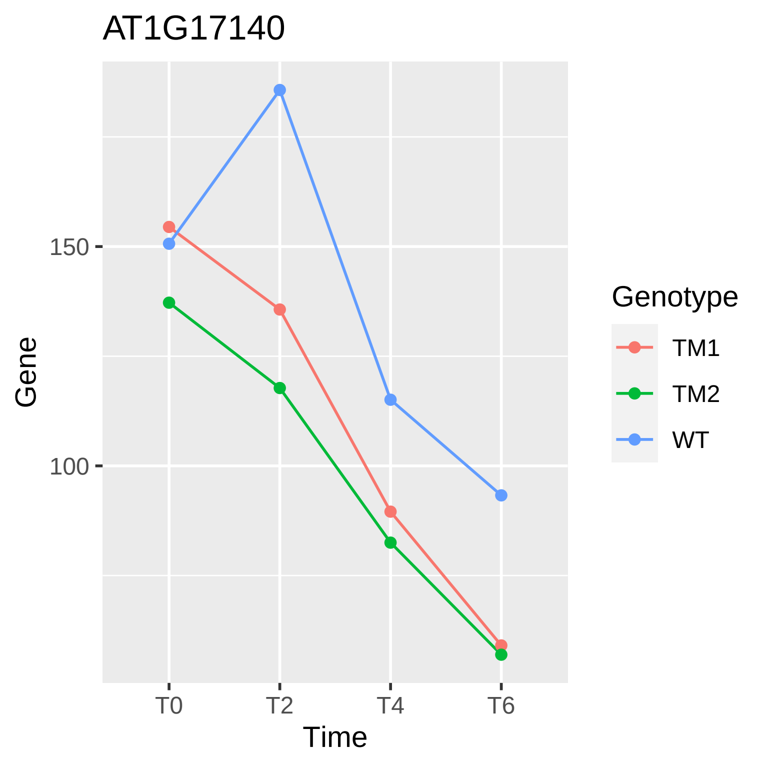
AT1G17140
Gene
150
100
T0
T2
T4
T6
Time
Genotype
TM1
TM2
WT

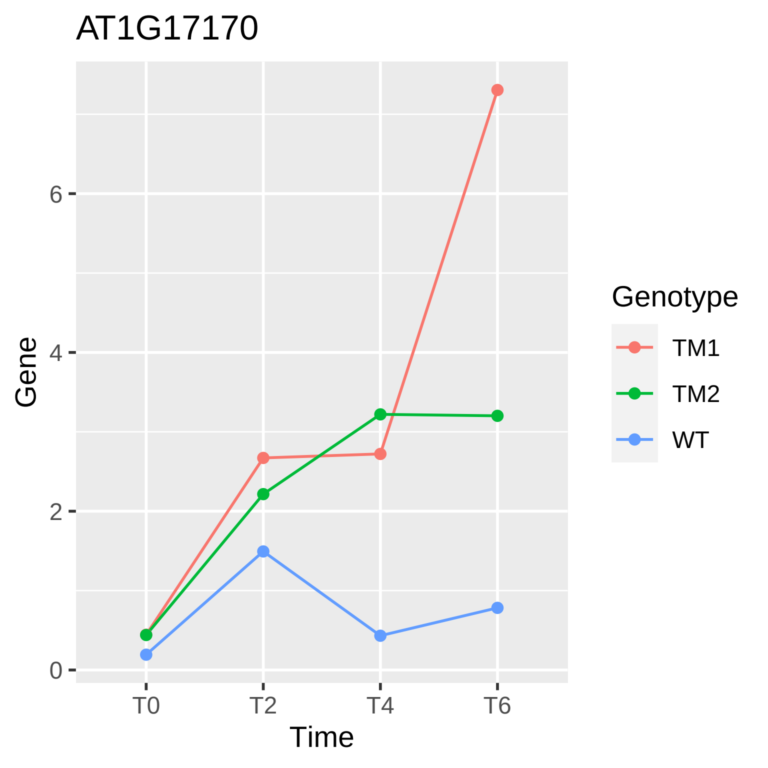
AT1G17170
Gene
6
4
2
0
T0
T2
T4
T6
Time
Genotype
TM1
TM2
WT

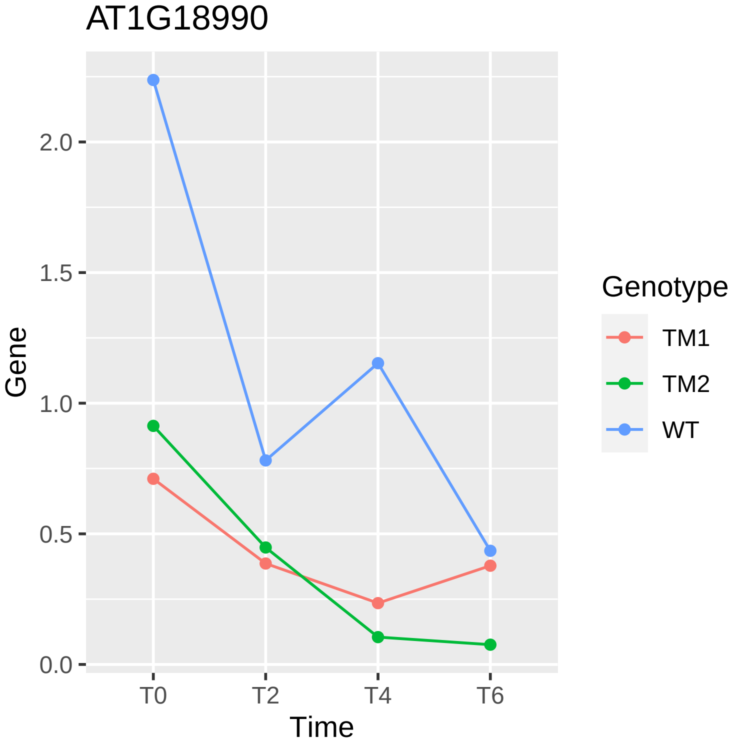
AT1G18990
Gene
2.0
1.5
1.0
0.5
0.0
T0
T2
T4
T6
Time
Genotype
TM1
TM2
WT

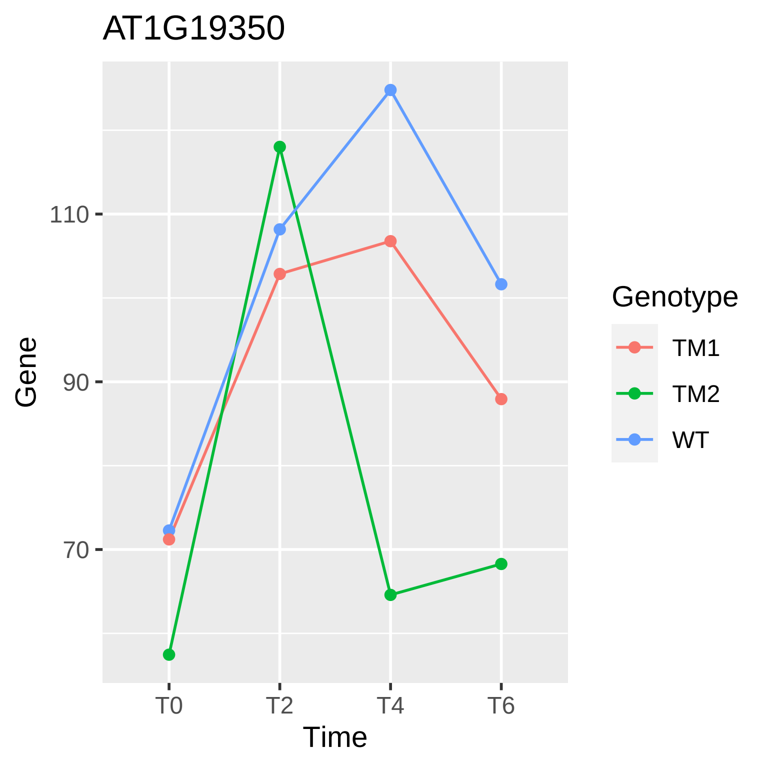

AT1G19350
Gene
110
90
70
T0
T2
T4
T6
Time
Genotype
TM1
TM2
WT

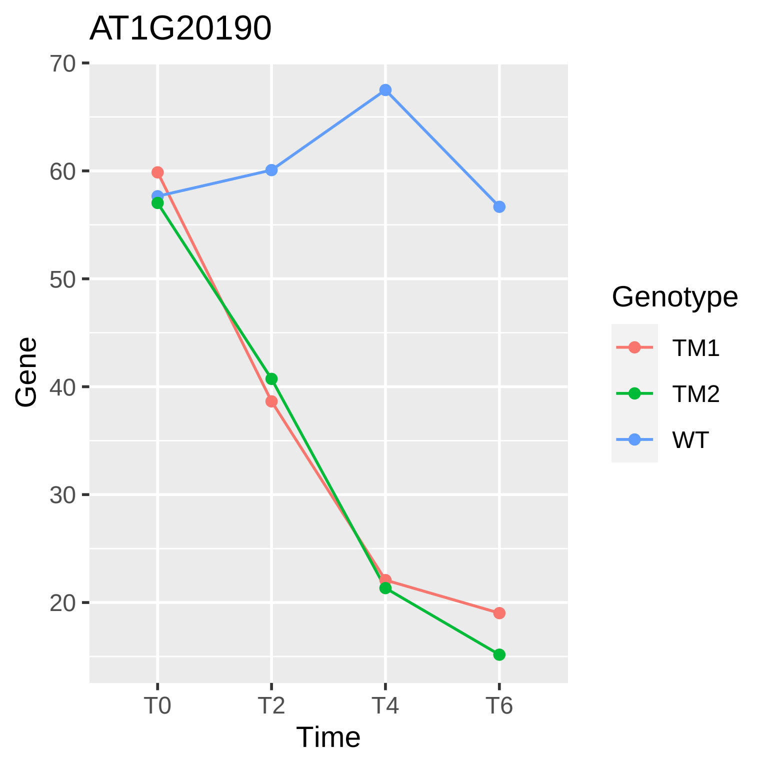
AT1G20190
70
60
50
40
30
20
Gene
T0
T2
T4
T6
Time
Genotype
TM1
TM2
WT

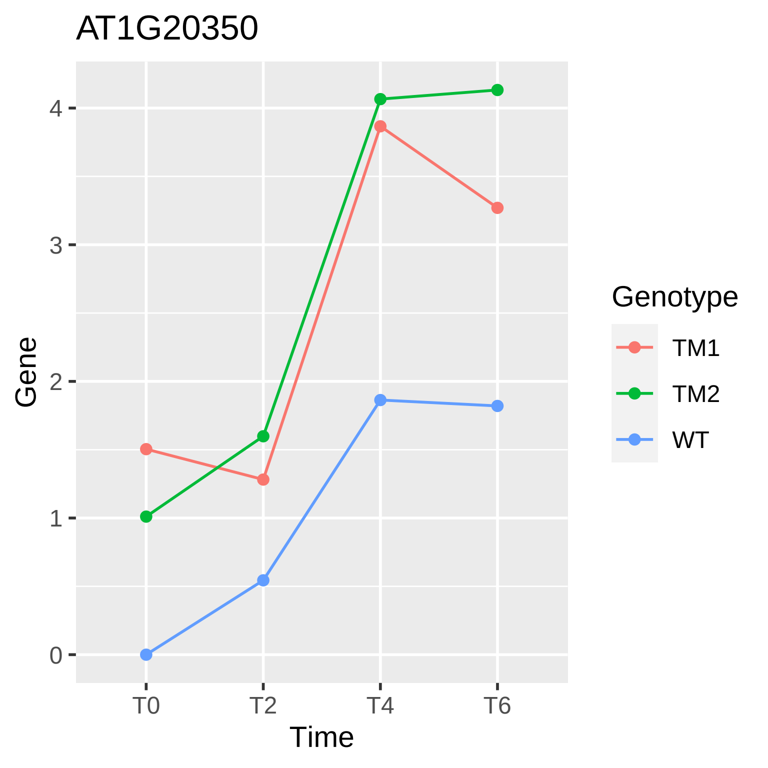
AT1G20350
Gene
4
3
2
1
0
T0
T2
T4
T6
Time
Genotype
TM1
TM2
WT

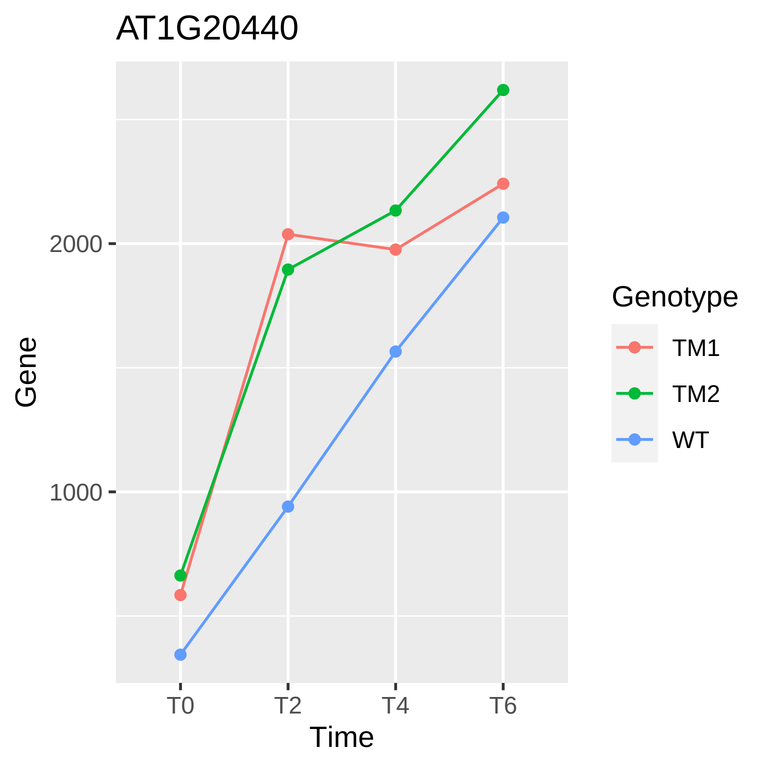
AT1G20440
2000
1000
Gene
T0
T2
T4
T6
Time
Genotype
TM1
TM2
WT

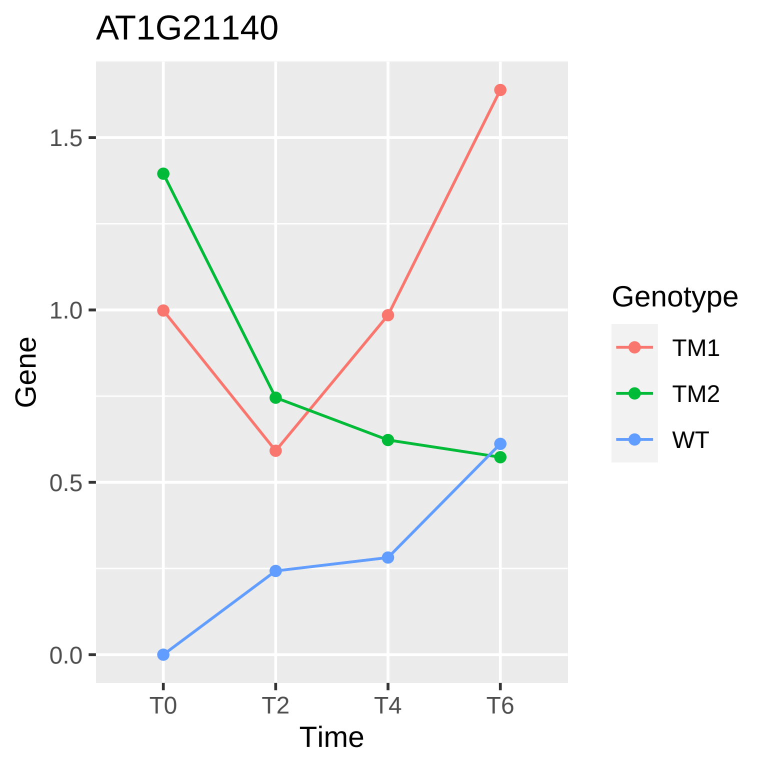
AT1G21140
1.5
1.0
0.5
0.0
Gene
T0
T2
T4
T6
Time
Genotype
TM1
TM2
WT

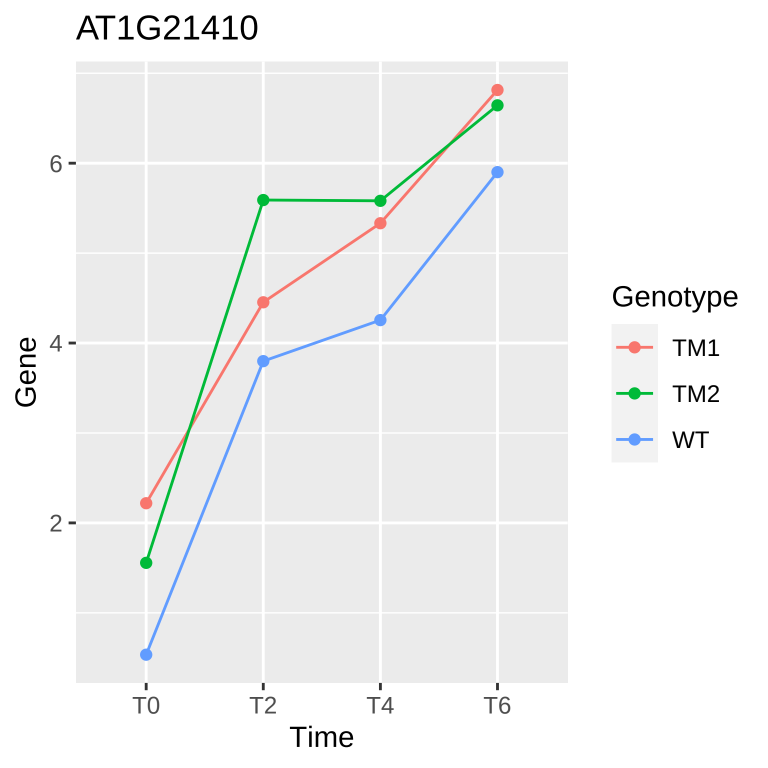
AT1G21410
Gene
6
4
2
T0
T2
T4
T6
Time
Genotype
TM1
TM2
WT

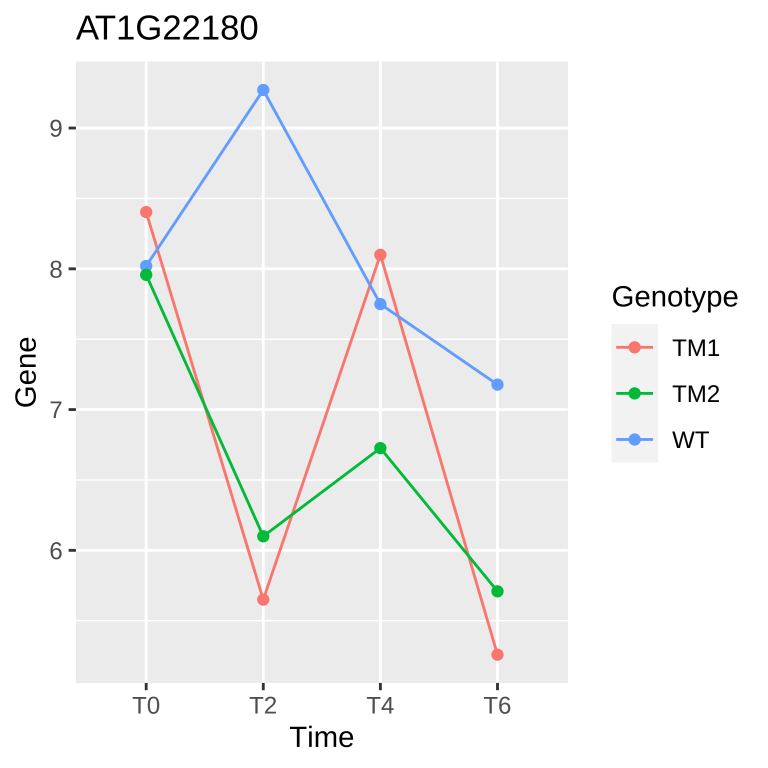
AT1G22180
Gene
9
8
7
6
T0
T2
T4
T6
Time
Genotype
TM1
TM2
WT

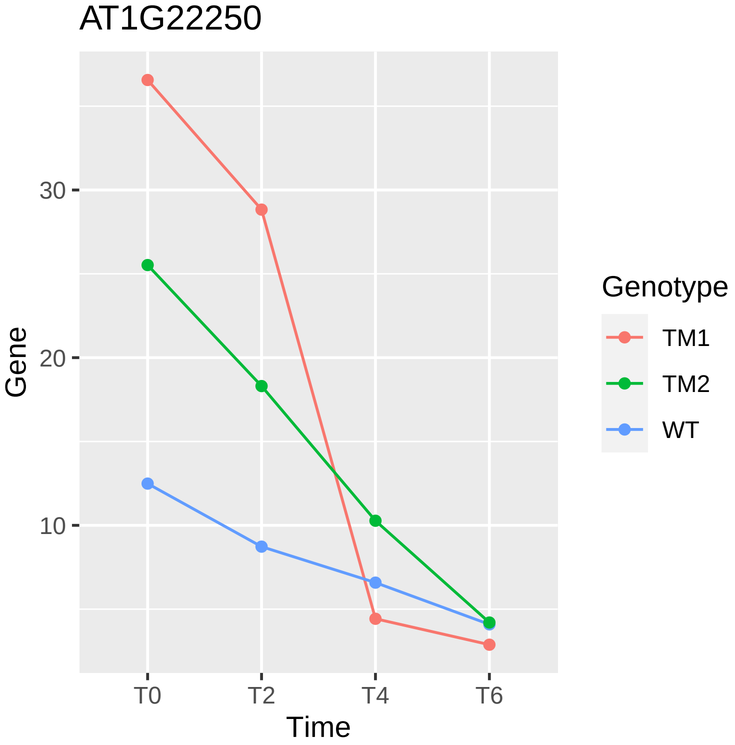
AT1G22250
30
20
10
Gene
T0
T2
T4
T6
Time
Genotype
TM1
TM2
WT

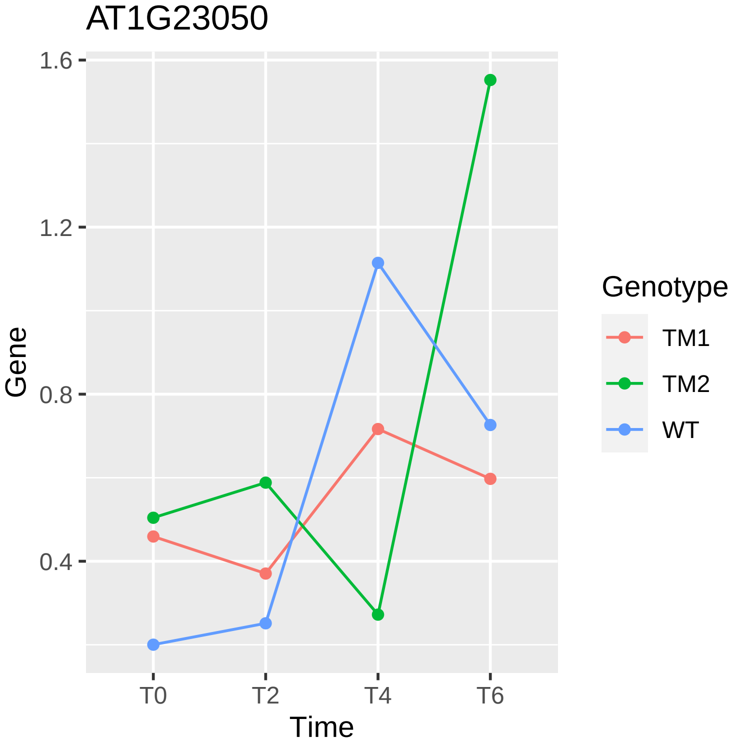

AT1G23050
1.6
1.2
0.8
0.4
Gene
T0
T2
T4
T6
Time
Genotype
TM1
TM2
WT

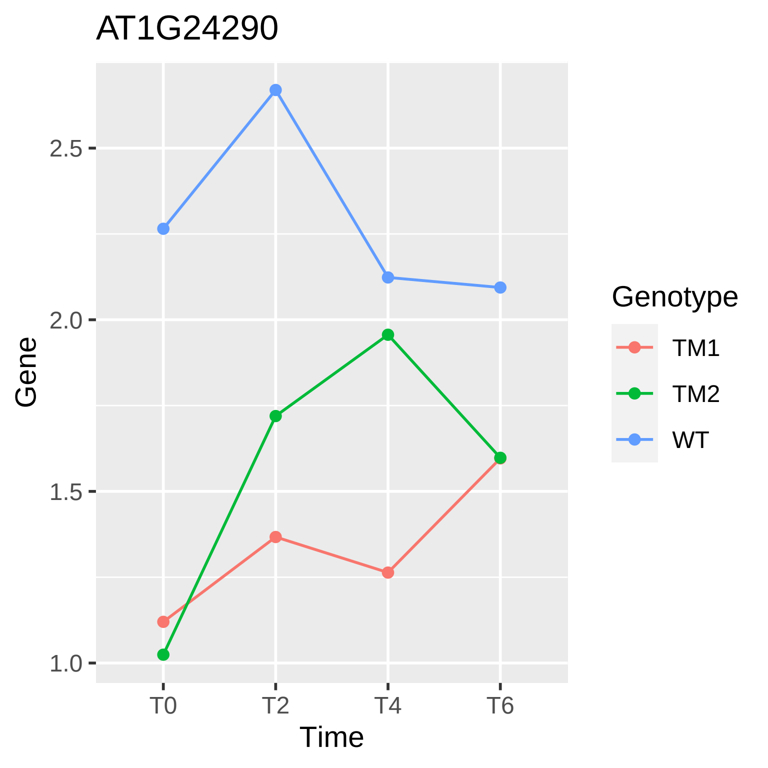
AT1G24290
Gene
2.5
2.0
1.5
1.0
T0
T2
T4
T6
Time
Genotype
TM1
TM2
WT

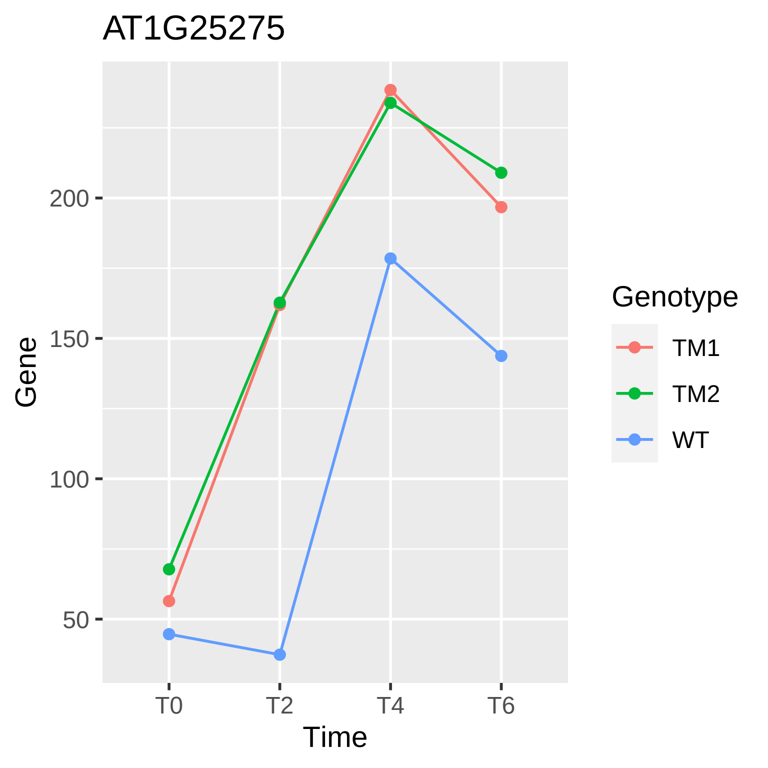
AT1G25275
Gene
200
150
100
50
T0
T2
T4
T6
Time
Genotype
TM1
TM2
WT

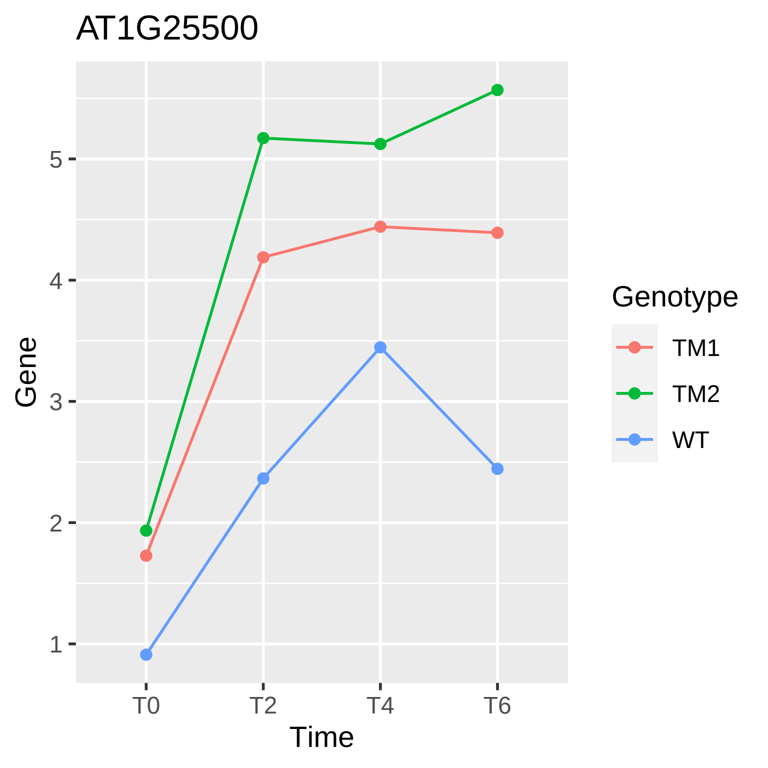
AT1G25500
Gene
1
2
3
4
5
T0
T2
T4
T6
Time
Genotype
TM1
TM2
WT

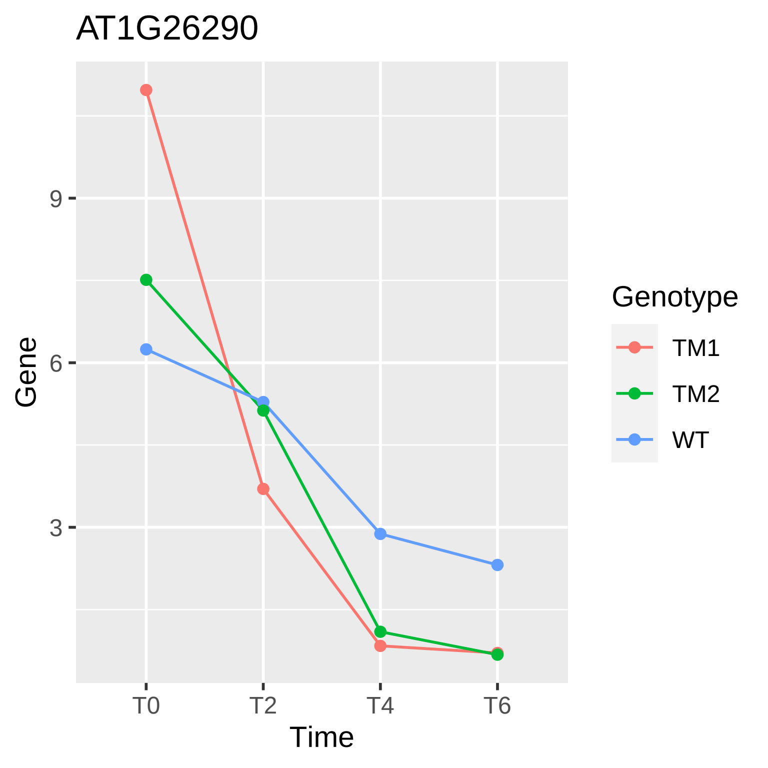

AT1G26290
Gene
9
6
3
T0
T2
T4
T6
Time
Genotype
TM1
TM2
WT

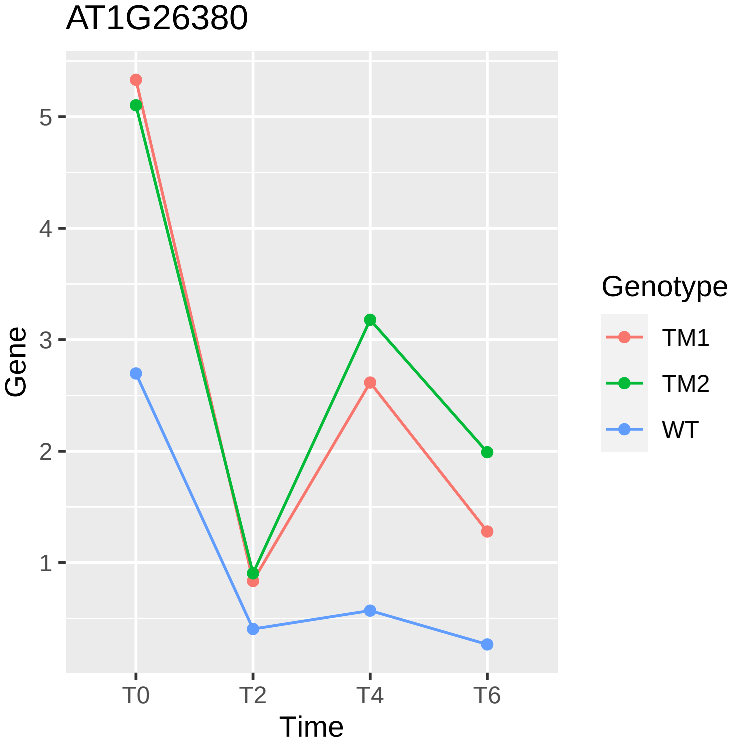

AT1G26380
Gene
Time
T0
T2
T4
T6
1
2
3
4
5
Genotype
TM1
TM2
WT

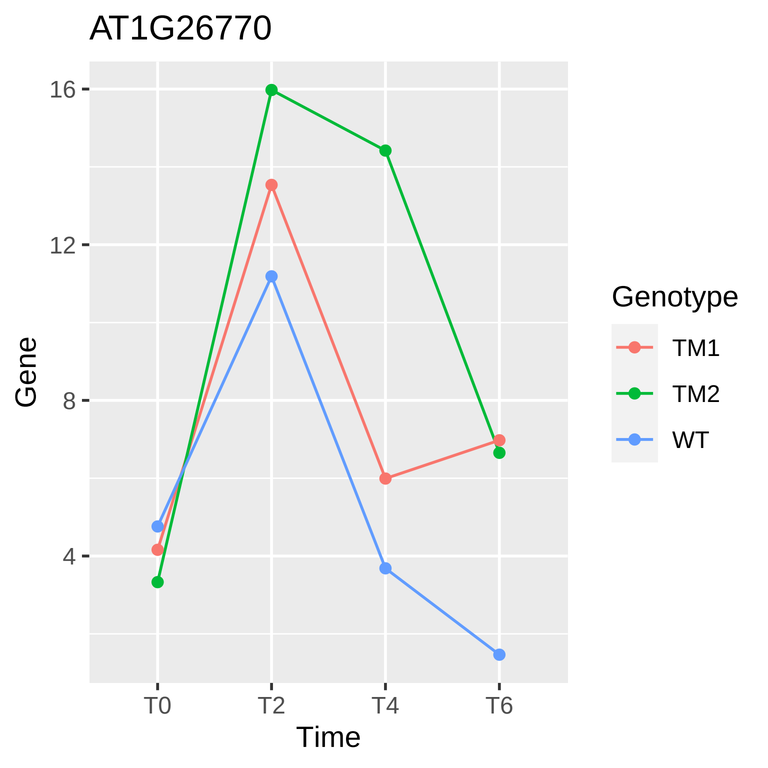

AT1G26770
16
12
8
4
Gene
T0
T2
T4
T6
Time
Genotype
TM1
TM2
WT

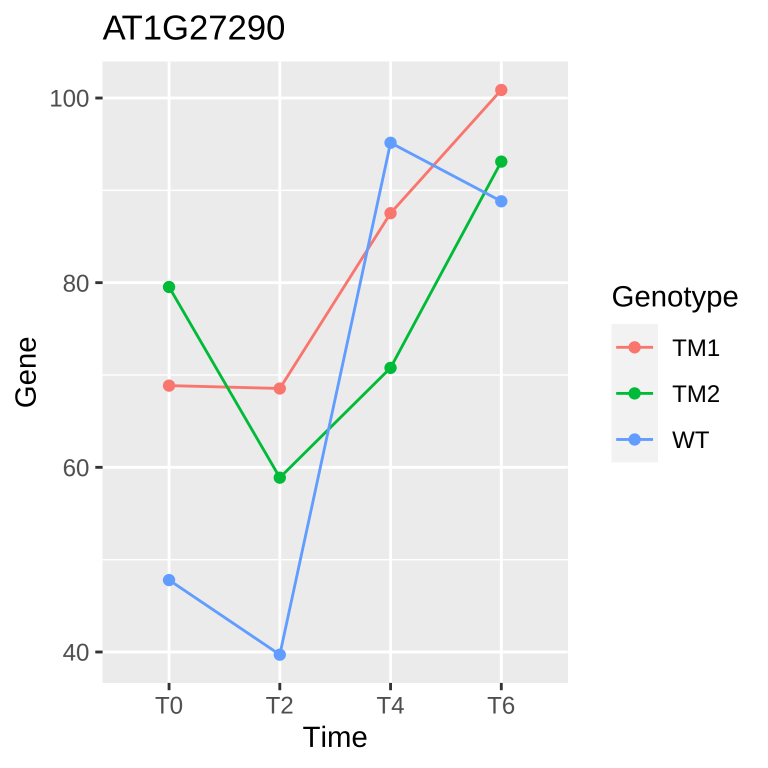
AT1G27290
Gene
100
80
60
40
T0
T2
T4
T6
Time
Genotype
TM1
TM2
WT

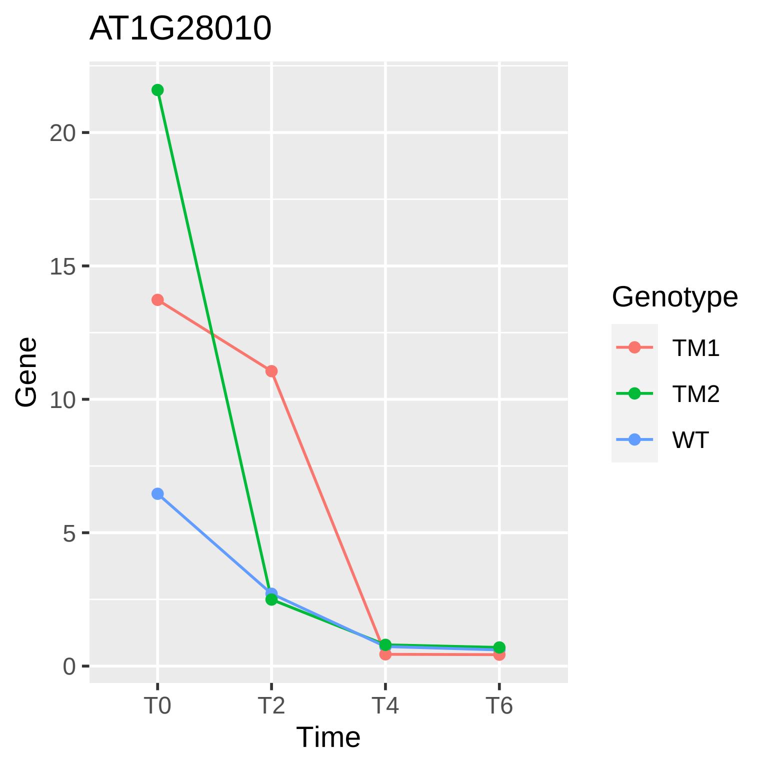

AT1G28010
Gene
20
15
10
5
0
T0
T2
T4
T6
Time
Genotype
TM1
TM2
WT

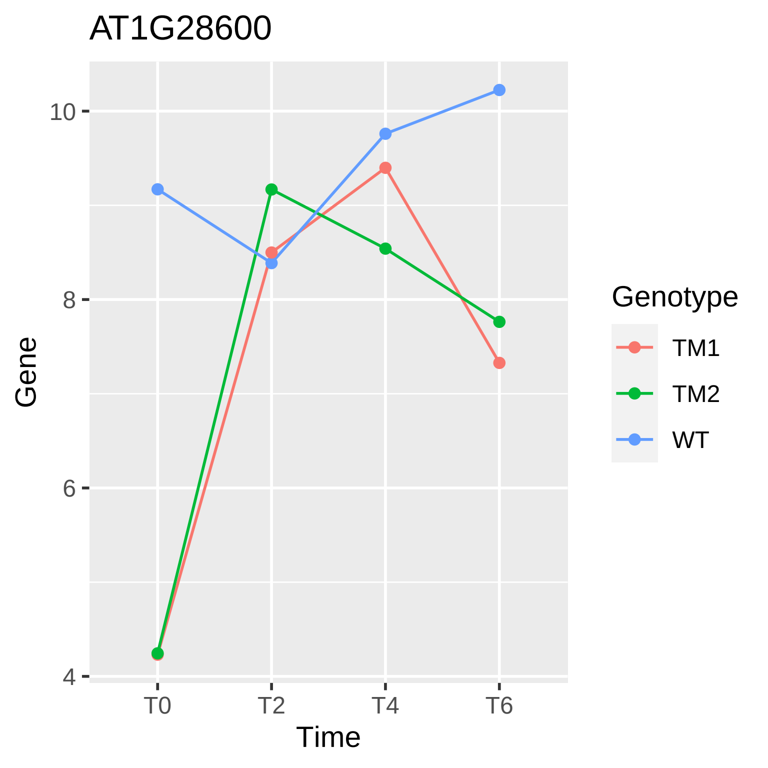

AT1G28600
Gene
Time
10
8
6
4
T0
T2
T4
T6
Genotype
TM1
TM2
WT

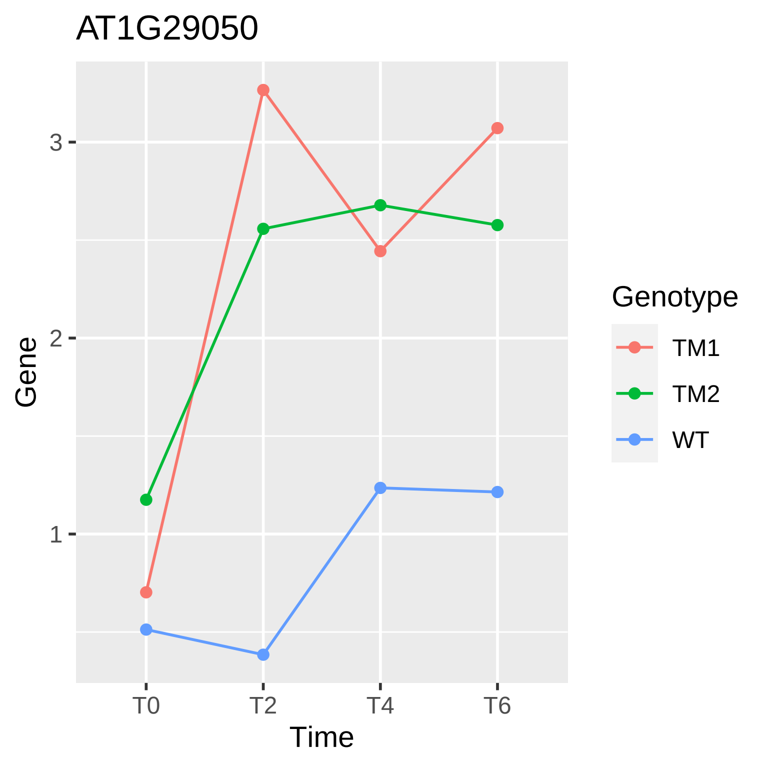
AT1G29050
Gene
3
2
1
T0
T2
T4
T6
Time
Genotype
TM1
TM2
WT

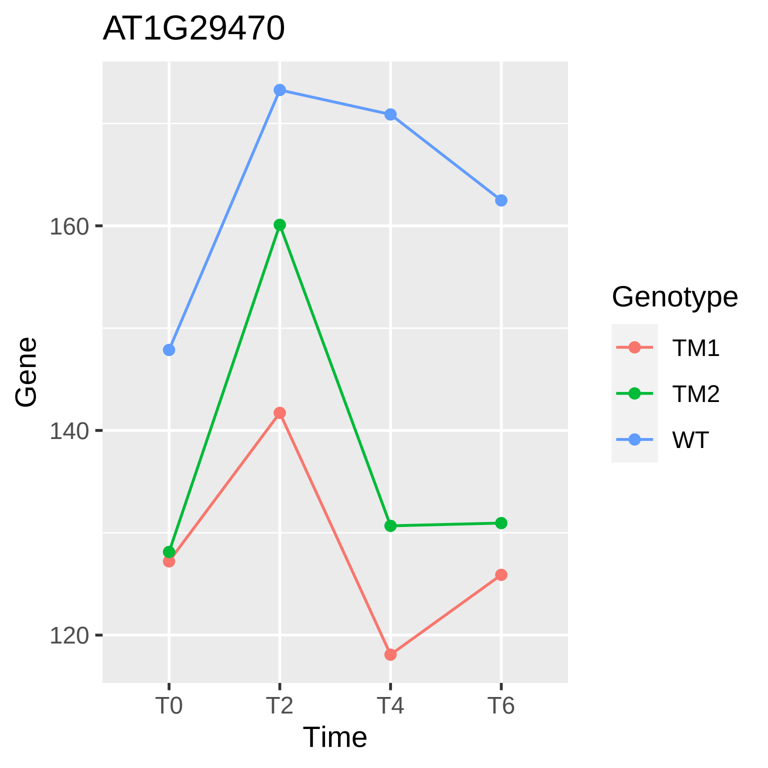

AT1G29470
Gene
160
140
120
T0
T2
T4
T6
Time
Genotype
TM1
TM2
WT

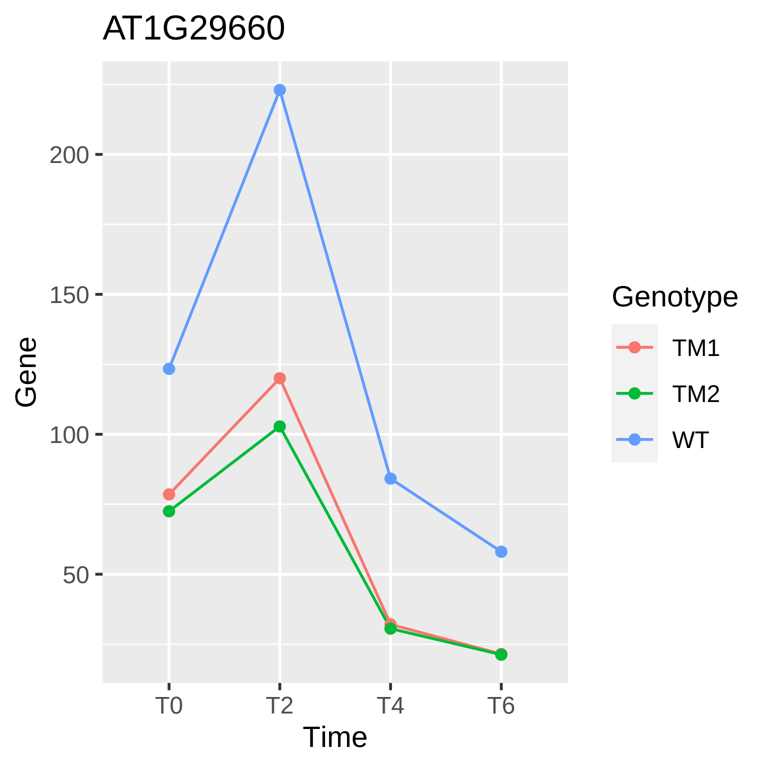
AT1G29660
Gene
200
150
100
50
T0
T2
T4
T6
Time
Genotype
TM1
TM2
WT

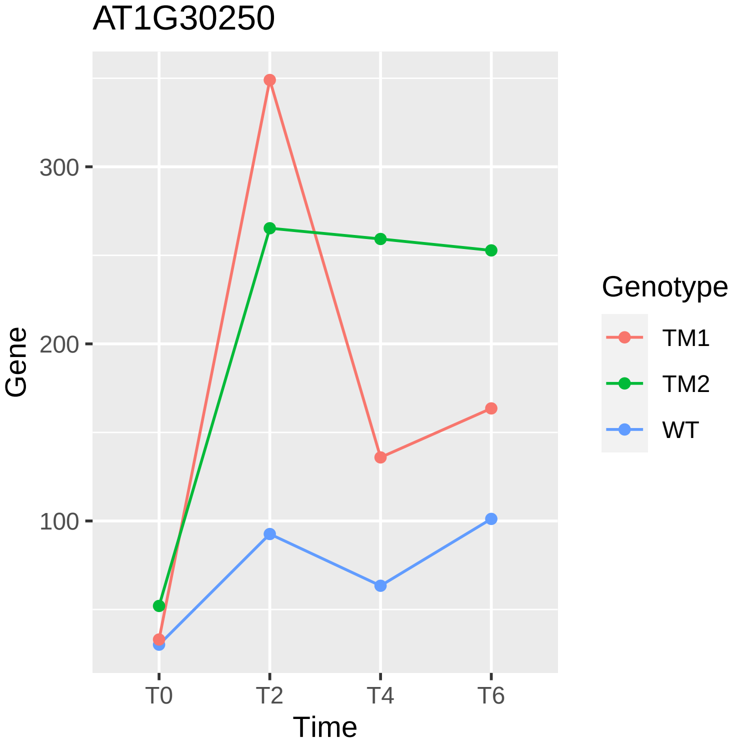
AT1G30250
Gene
300
200
100
T0
T2
T4
T6
Time
Genotype
TM1
TM2
WT

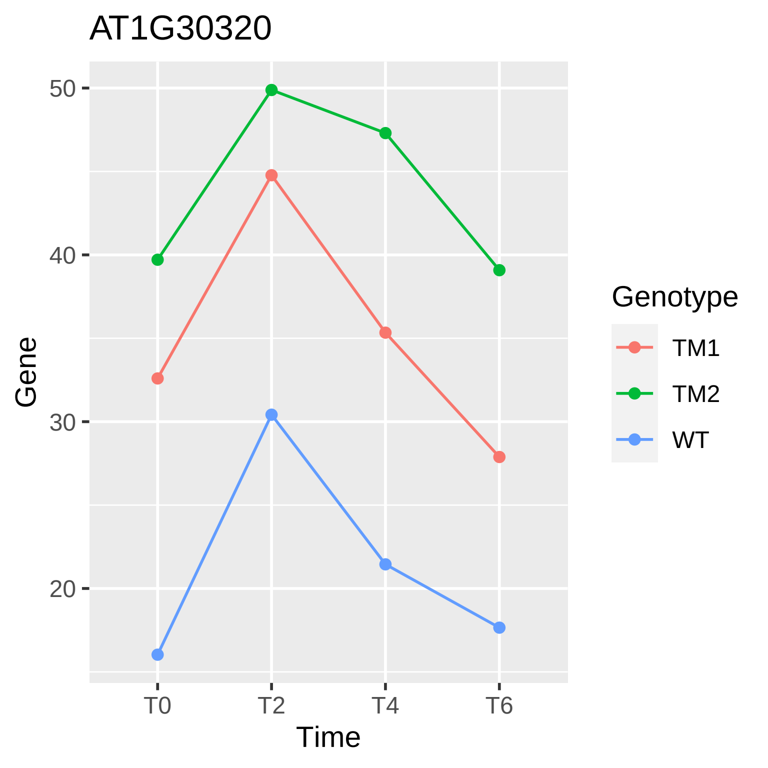

AT1G30320
50
40
30
20
Gene
T0
T2
T4
T6
Time
Genotype
TM1
TM2
WT

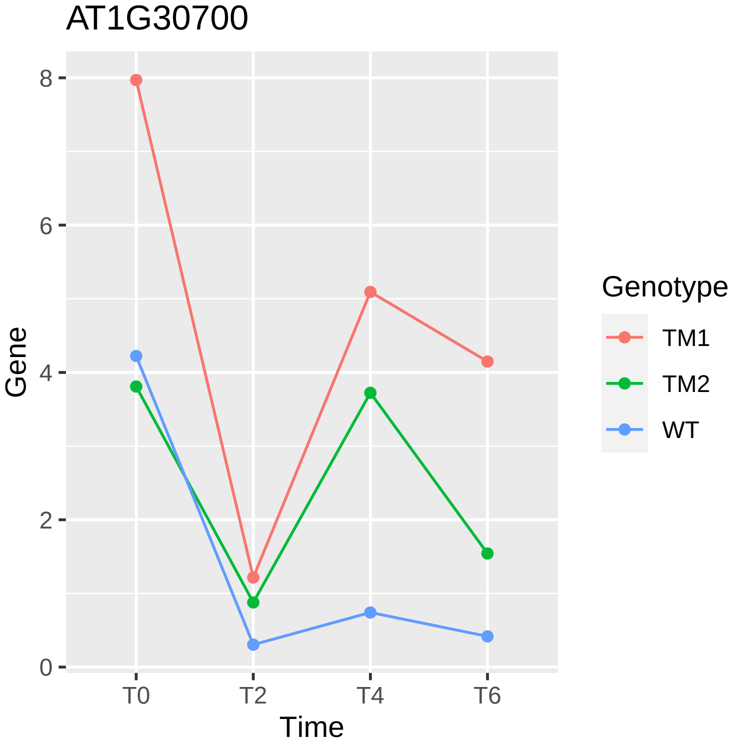

AT1G30700
Gene
8
6
4
2
0
T0
T2
T4
T6
Time
Genotype
TM1
TM2
WT

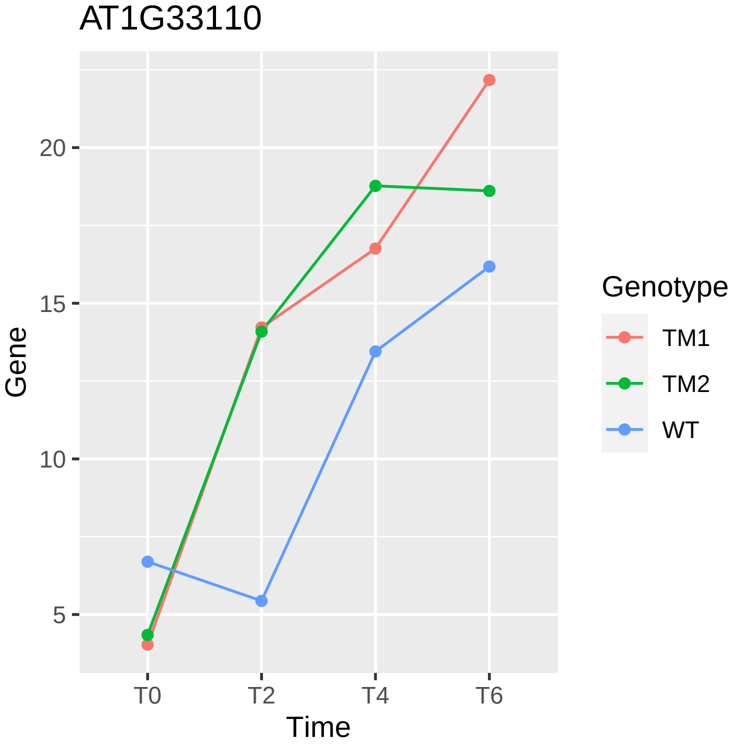
AT1G33110
Gene
20
15
10
5
T0
T2
T4
T6
Time
Genotype
TM1
TM2
WT

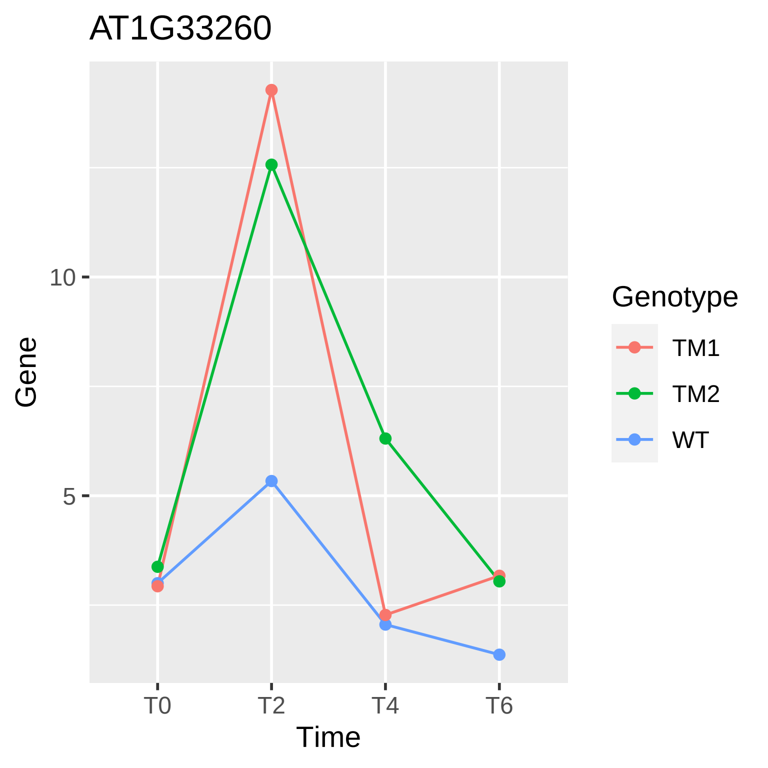

AT1G33260
10
5
Gene
T0
T2
T4
T6
Time
Genotype
TM1
TM2
WT

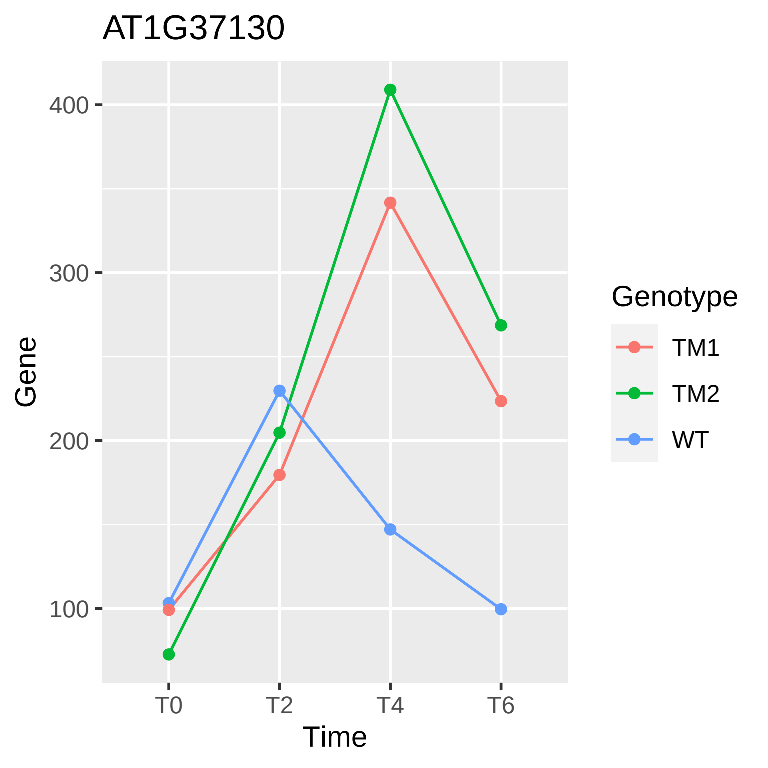
AT1G37130
400
300
200
100
Gene
T0
T2
T4
T6
Time
Genotype
TM1
TM2
WT

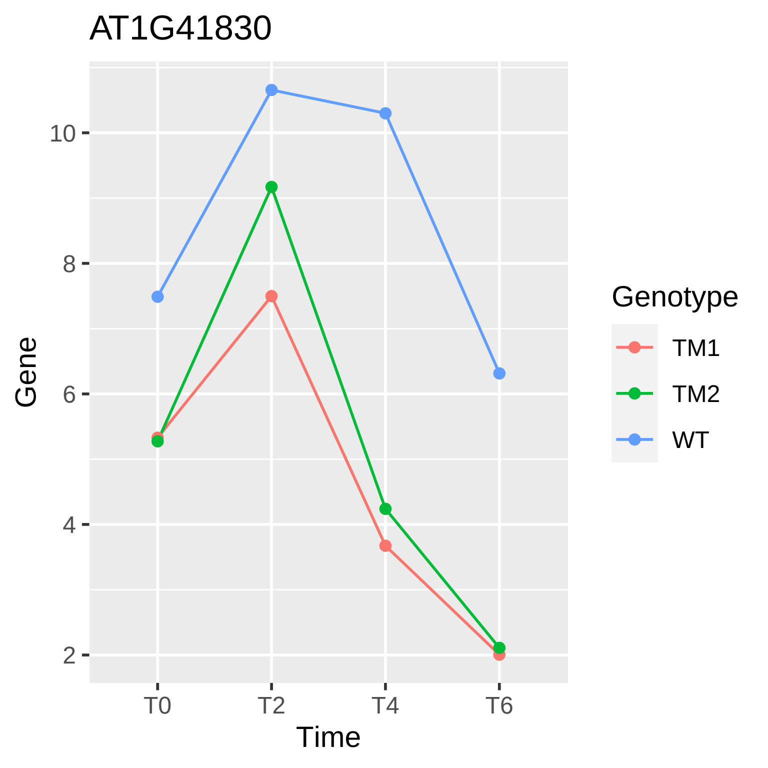

AT1G41830
Gene
10
8
6
4
2
T0
T2
T4
T6
Time
Genotype
TM1
TM2
WT

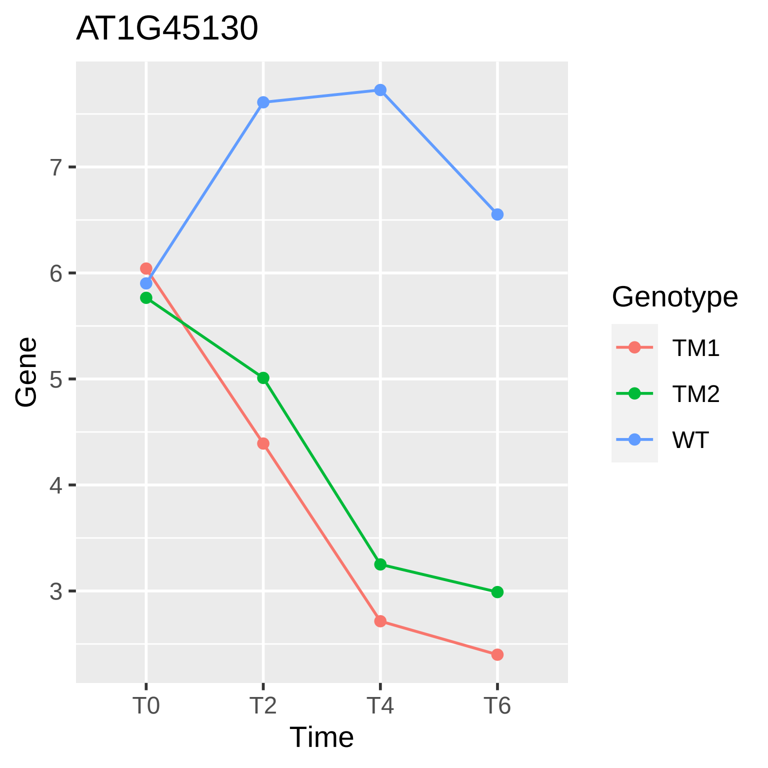
AT1G45130
Gene
7
6
5
4
3
T0
T2
T4
T6
Time
Genotype
TM1
TM2
WT

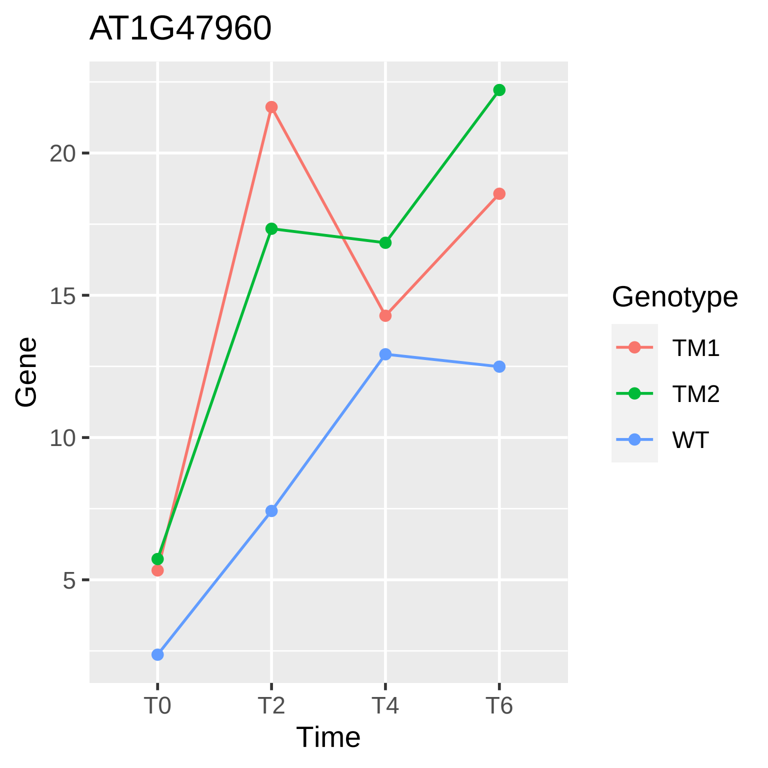

AT1G47960
Gene
20
15
10
5
T0
T2
T4
T6
Time
Genotype
TM1
TM2
WT

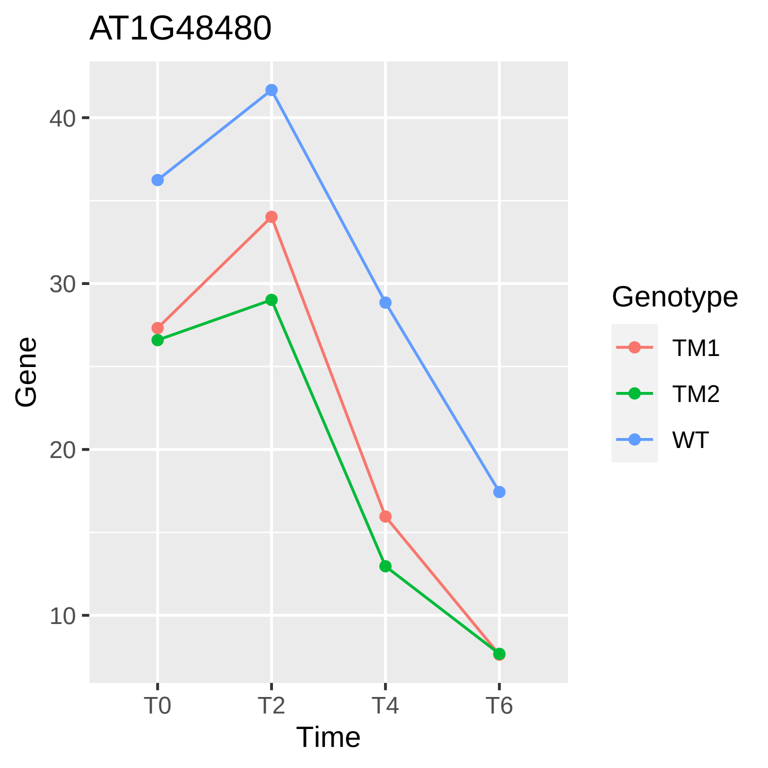
AT1G48480
Gene
Time
T0
T2
T4
T6
10
20
30
40
Genotype
TM1
TM2
WT

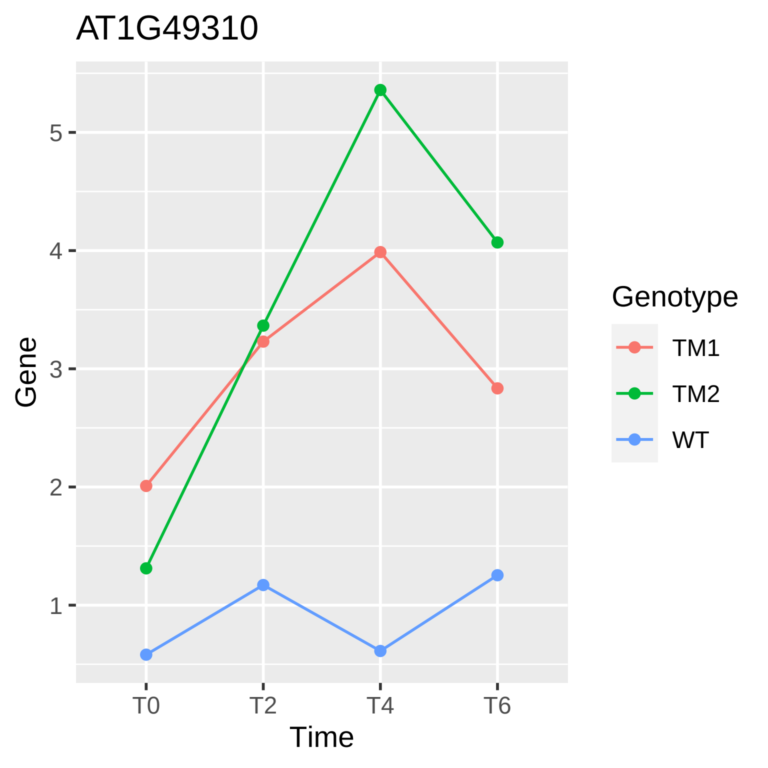

AT1G49310
Gene
Time
T0
T2
T4
T6
1
2
3
4
5
Genotype
TM1
TM2
WT

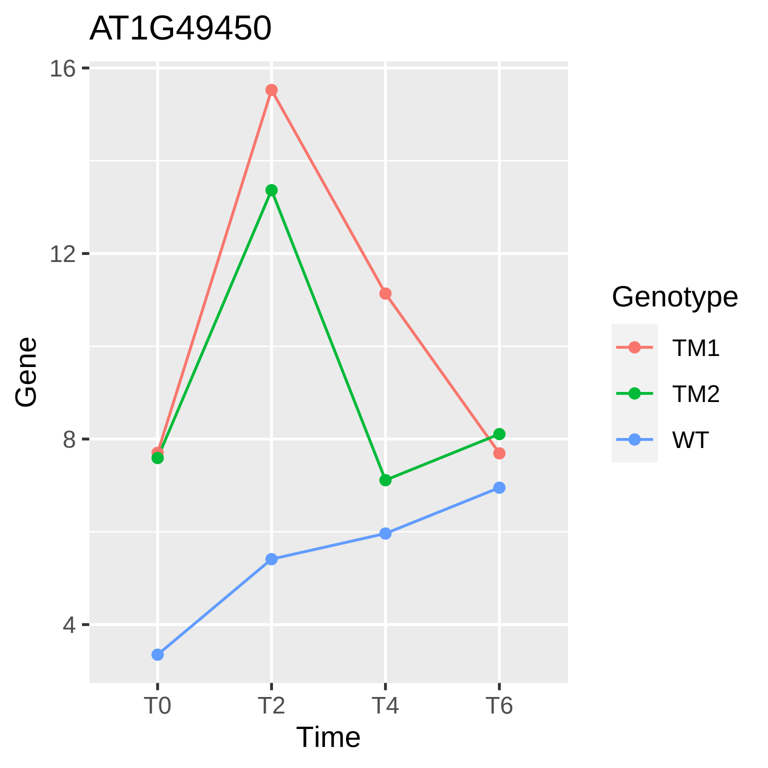
AT1G49450
16
12
8
4
Gene
T0
T2
T4
T6
Time
Genotype
TM1
TM2
WT

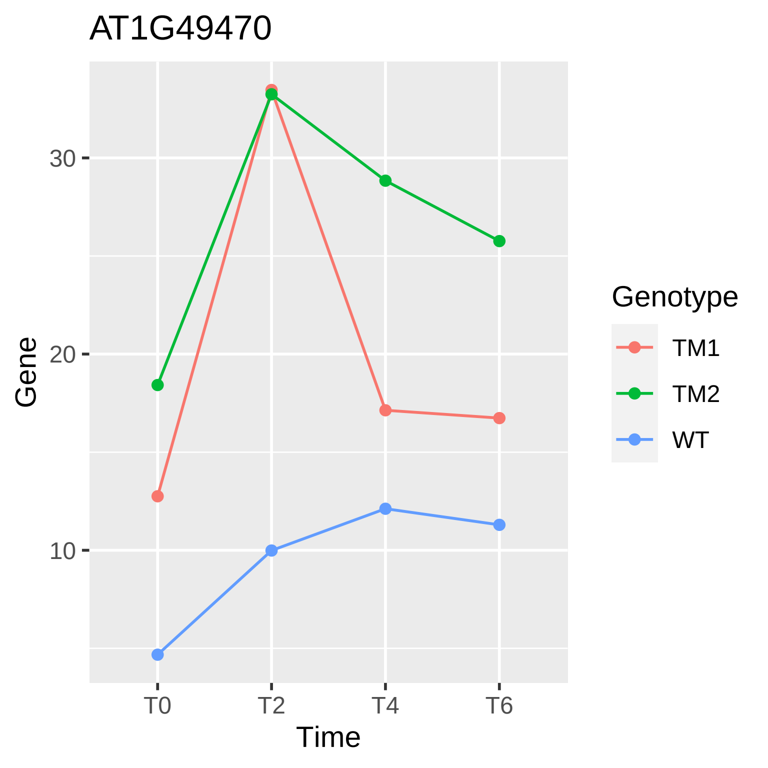
AT1G49470
Gene
30
20
10
T0
T2
T4
T6
Time
Genotype
TM1
TM2
WT

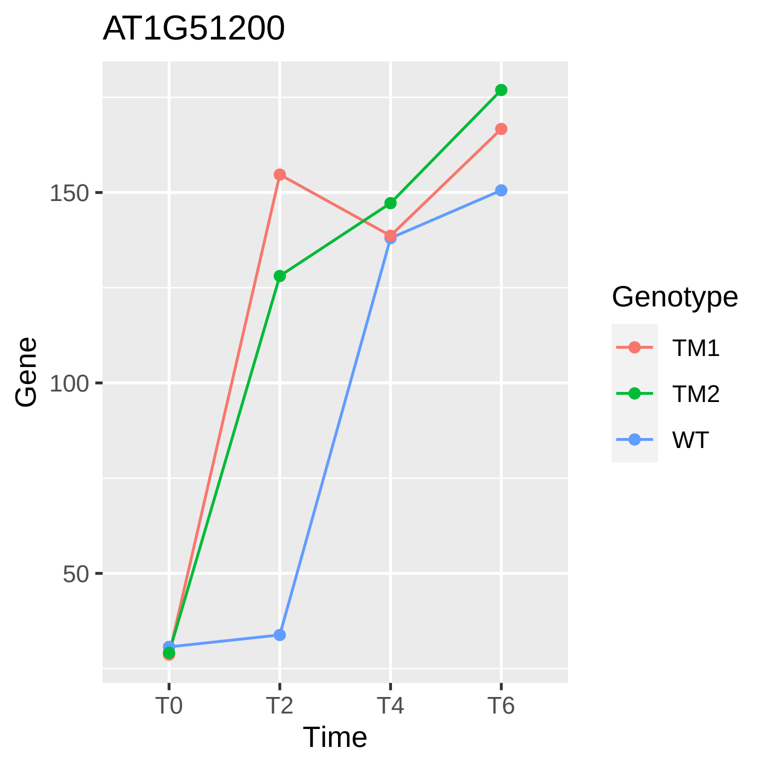

AT1G51200
Gene
150
100
50
T0
T2
T4
T6
Time
Genotype
TM1
TM2
WT

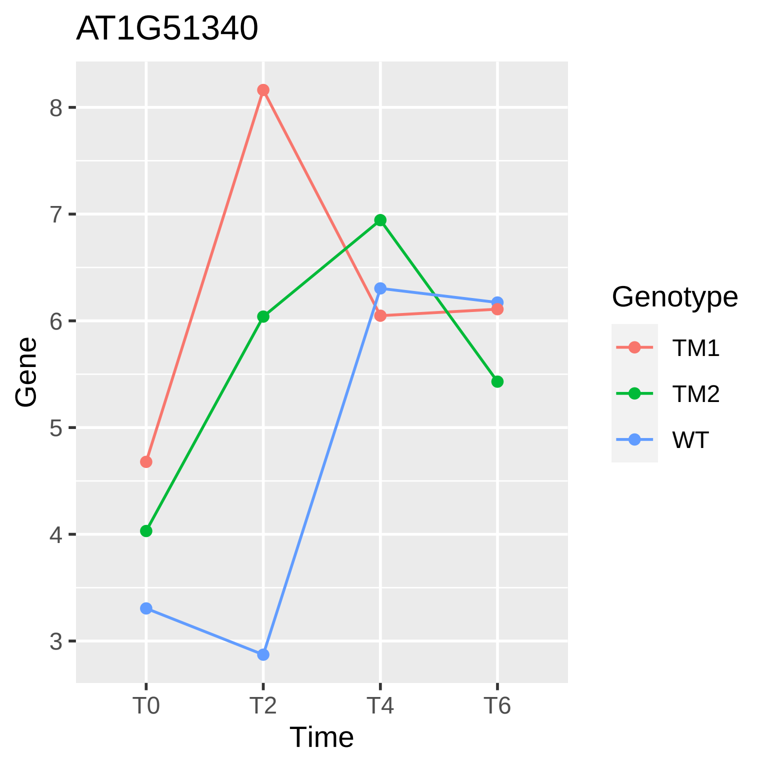
AT1G51340
Gene
8
7
6
5
4
3
T0
T2
T4
T6
Time
Genotype
TM1
TM2
WT

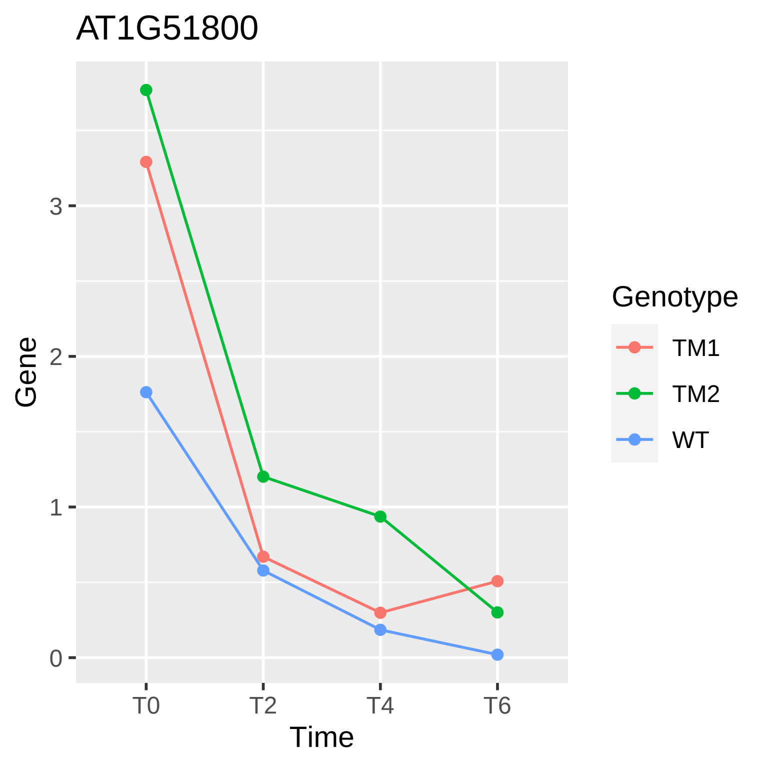

AT1G51800
Gene
0
1
2
3
T0
T2
T4
T6
Time
Genotype
TM1
TM2
WT

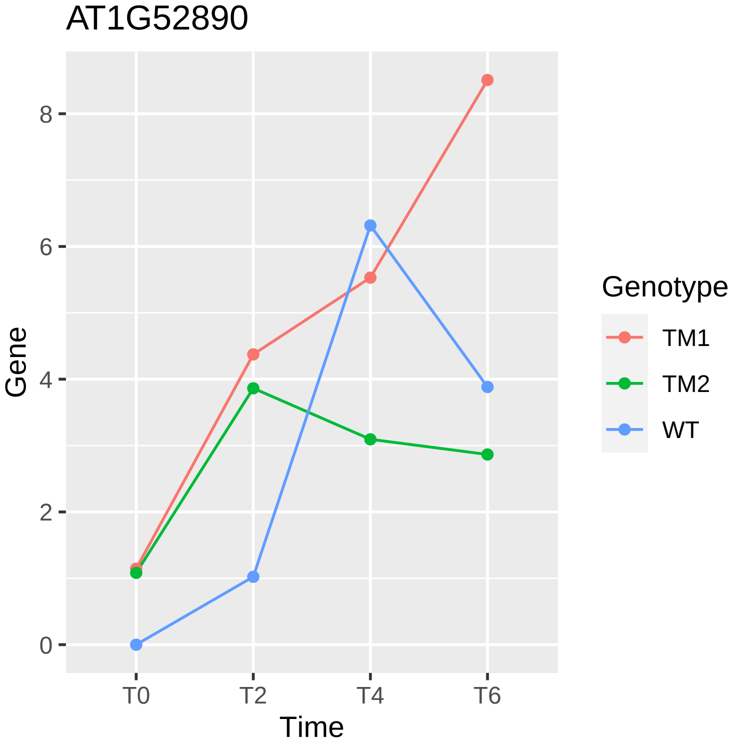

AT1G52890
Gene
Time
T0
T2
T4
T6
0
2
4
6
8
Genotype
TM1
TM2
WT

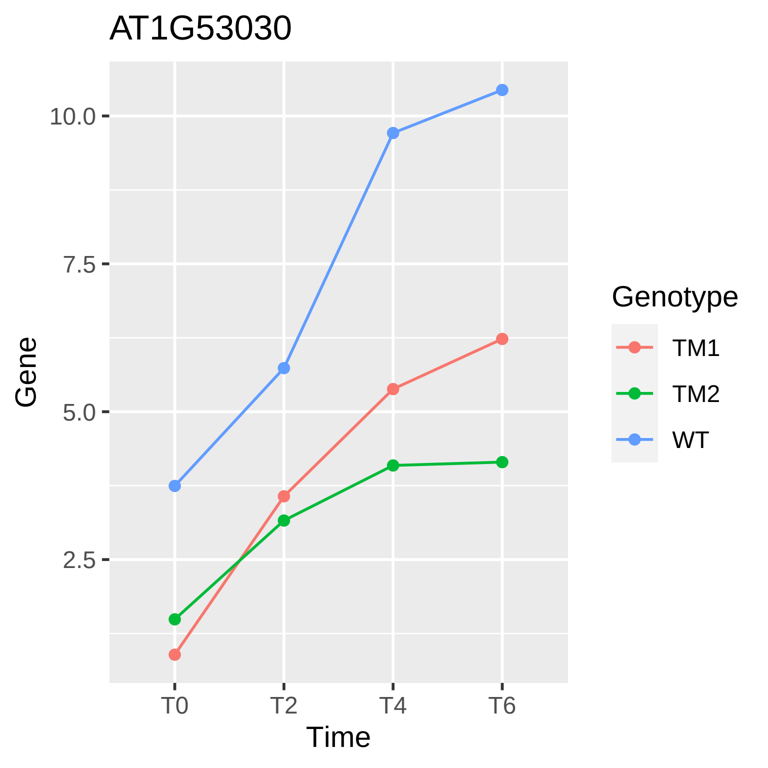
AT1G53030
10.0
7.5
5.0
2.5
Gene
T0
T2
T4
T6
Time
Genotype
TM1
TM2
WT

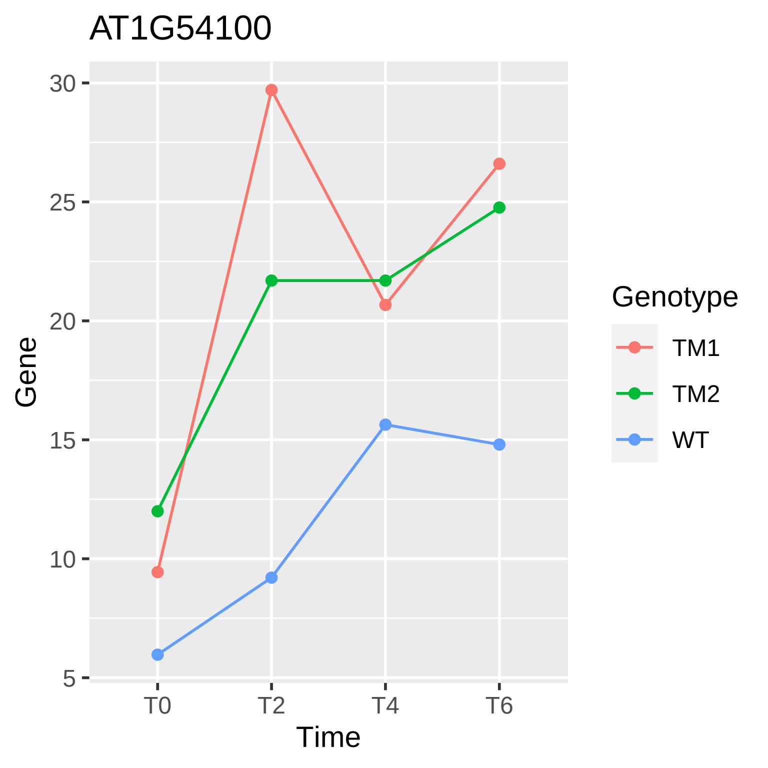

AT1G54100
Gene
30
25
20
15
10
5
T0
T2
T4
T6
Time
Genotype
TM1
TM2
WT

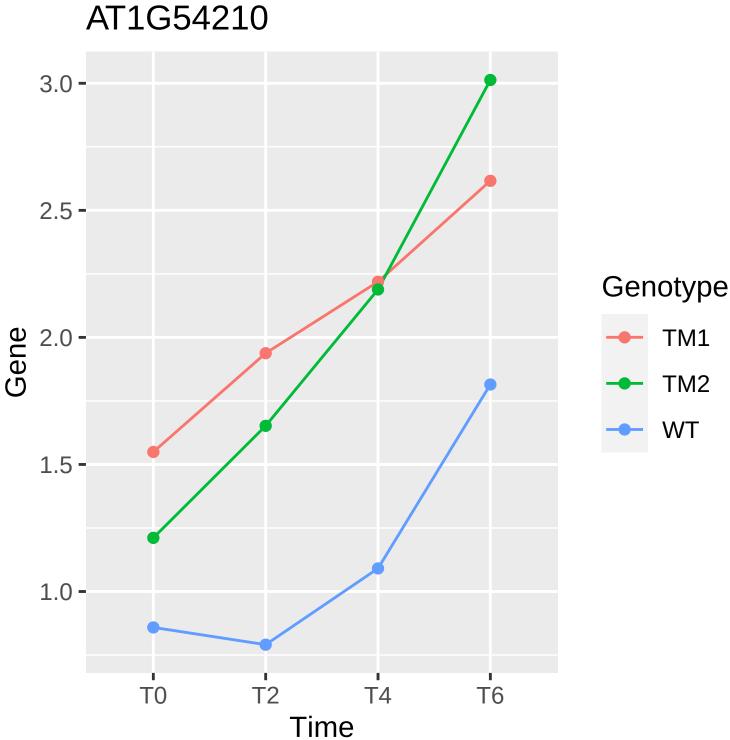
AT1G54210
3.0
2.5
2.0
1.5
1.0
Gene
T0
T2
T4
T6
Time
Genotype
TM1
TM2
WT

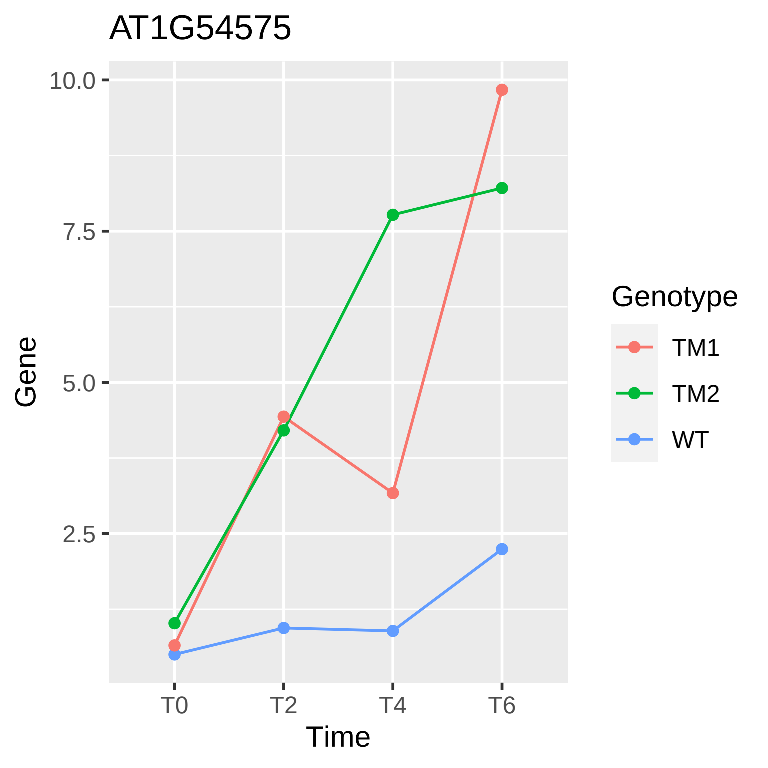

AT1G54575
10.0
7.5
5.0
2.5
Gene
T0
T2
T4
T6
Time
Genotype
TM1
TM2
WT

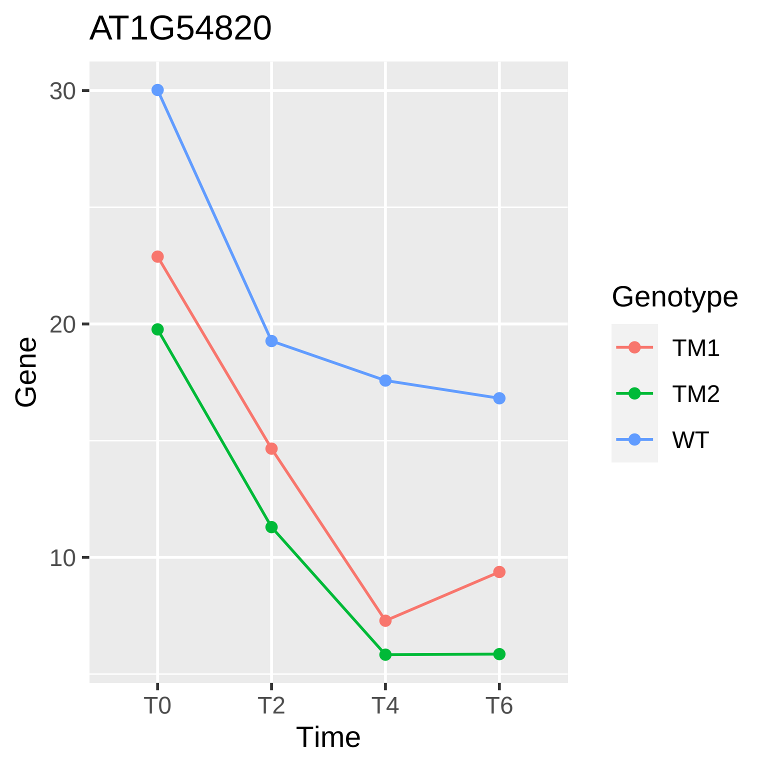

AT1G54820
30
20
10
Gene
T0
T2
T4
T6
Time
Genotype
TM1
TM2
WT

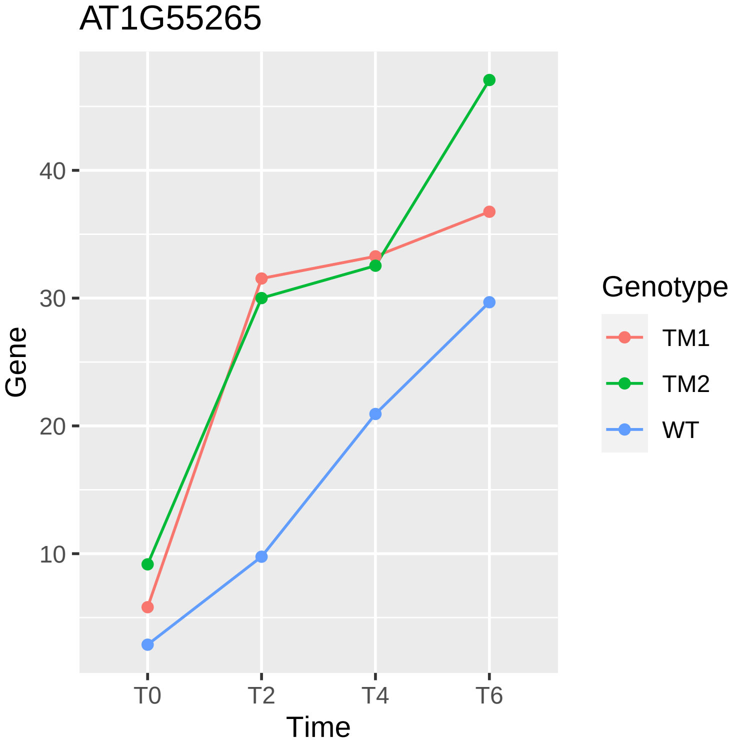
AT1G55265
40
30
20
10
Gene
T0
T2
T4
T6
Time
Genotype
TM1
TM2
WT

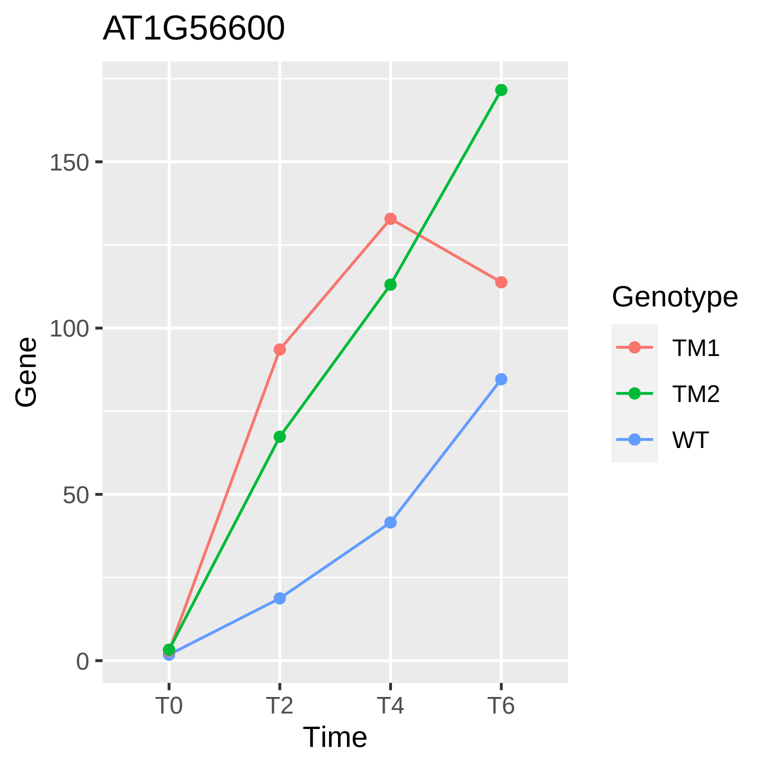

AT1G56600
Gene
150
100
50
0
T0
T2
T4
T6
Time
Genotype
TM1
TM2
WT

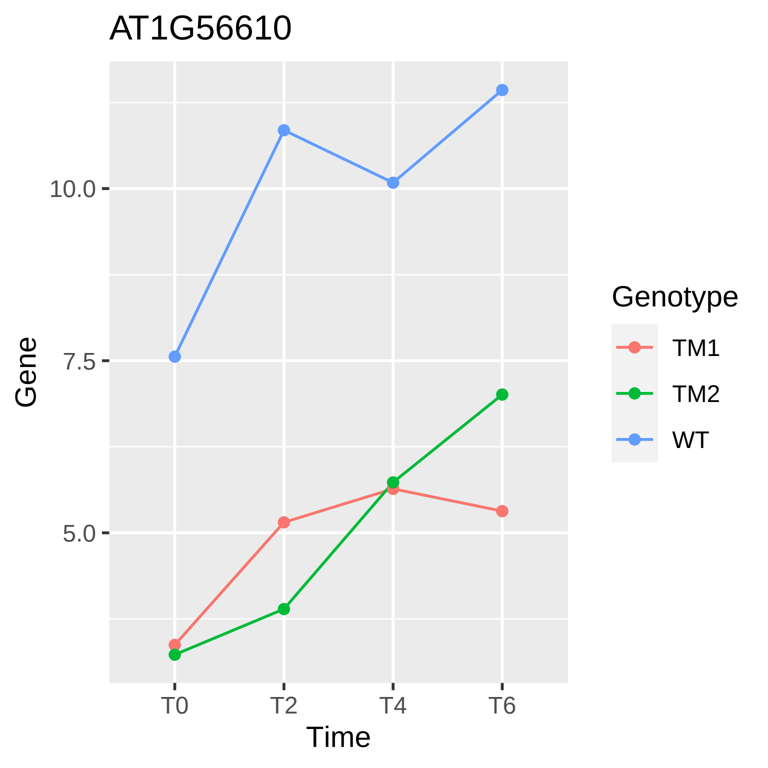
AT1G56610
Gene
10.0
7.5
5.0
T0
T2
T4
T6
Time
Genotype
TM1
TM2
WT

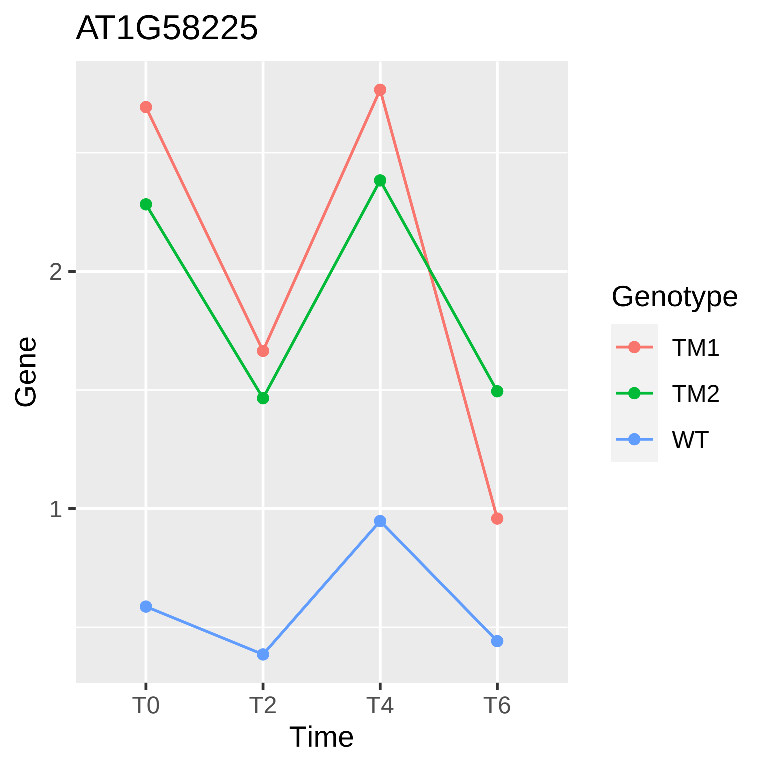
AT1G58225
Gene
2
1
T0
T2
T4
T6
Time
Genotype
TM1
TM2
WT

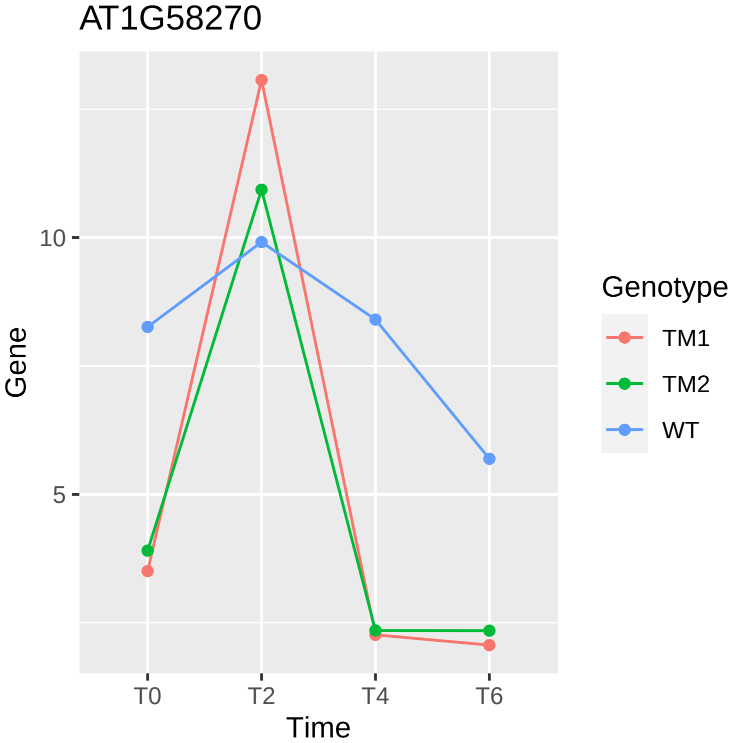

AT1G58270
Gene
10
5
T0
T2
T4
T6
Time
Genotype
TM1
TM2
WT

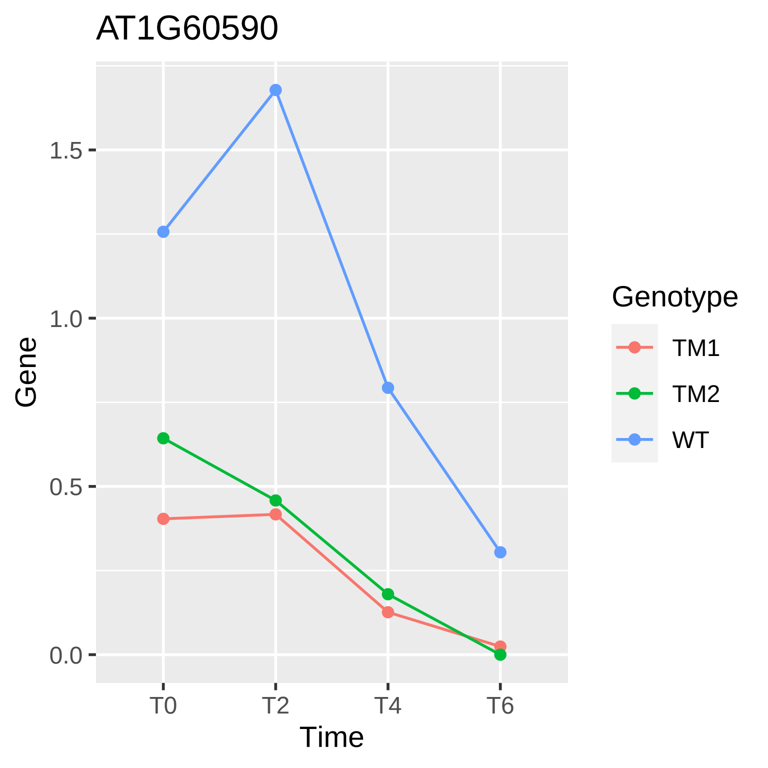
AT1G60590
Gene
1.5
1.0
0.5
0.0
T0
T2
T4
T6
Time
Genotype
TM1
TM2
WT

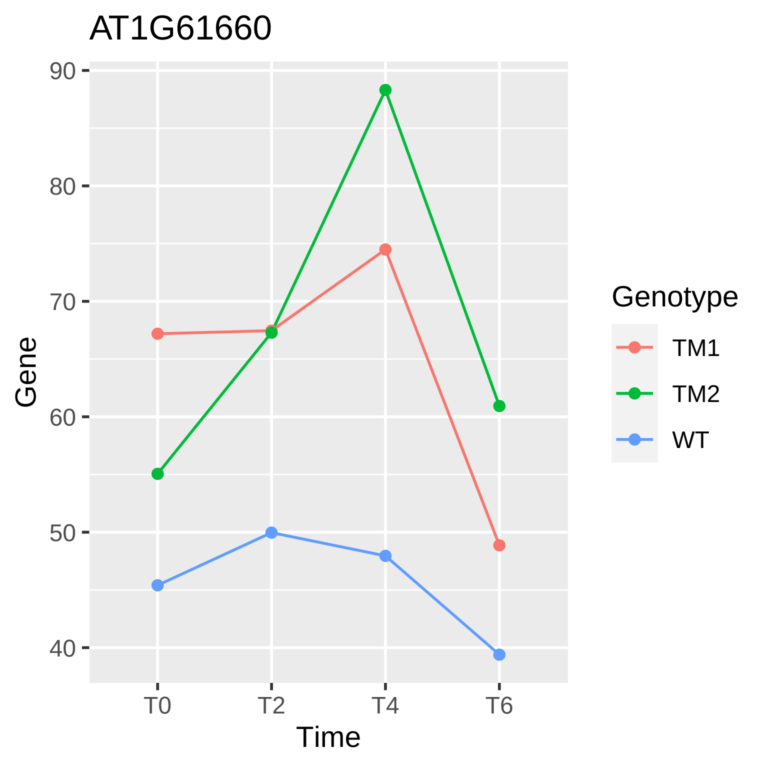

AT1G61660
90
80
70
60
50
40
Gene
T0
T2
T4
T6
Time
Genotype
TM1
TM2
WT

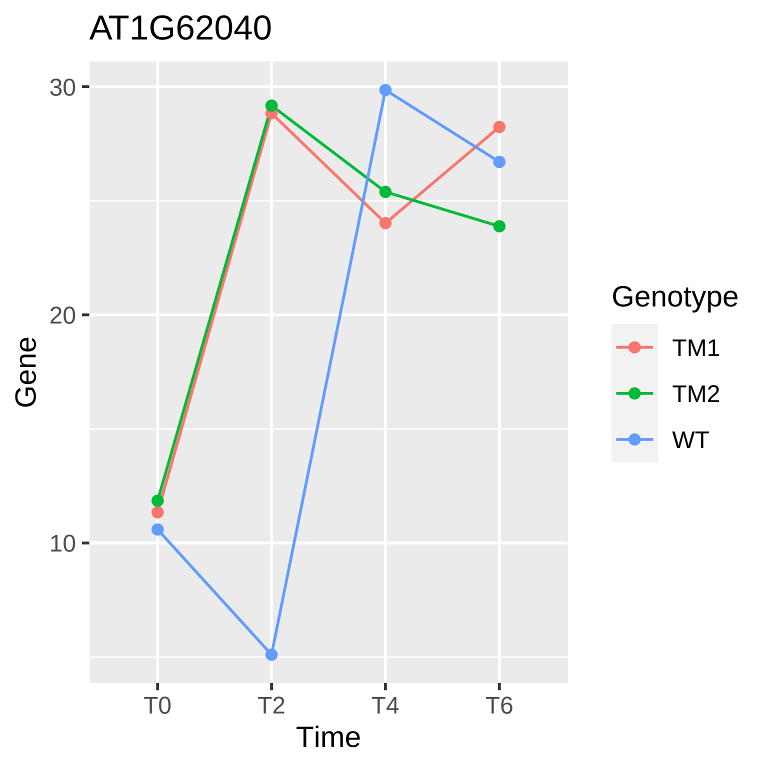

AT1G62040
30
20
10
Gene
T0
T2
T4
T6
Time
Genotype
TM1
TM2
WT

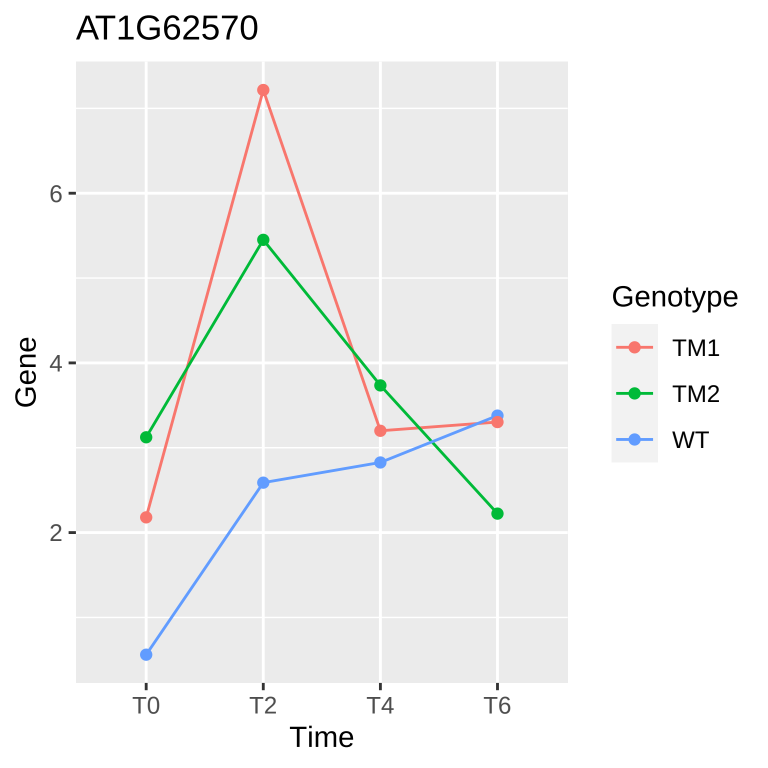

AT1G62570
Gene
6
4
2
T0
T2
T4
T6
Time
Genotype
TM1
TM2
WT

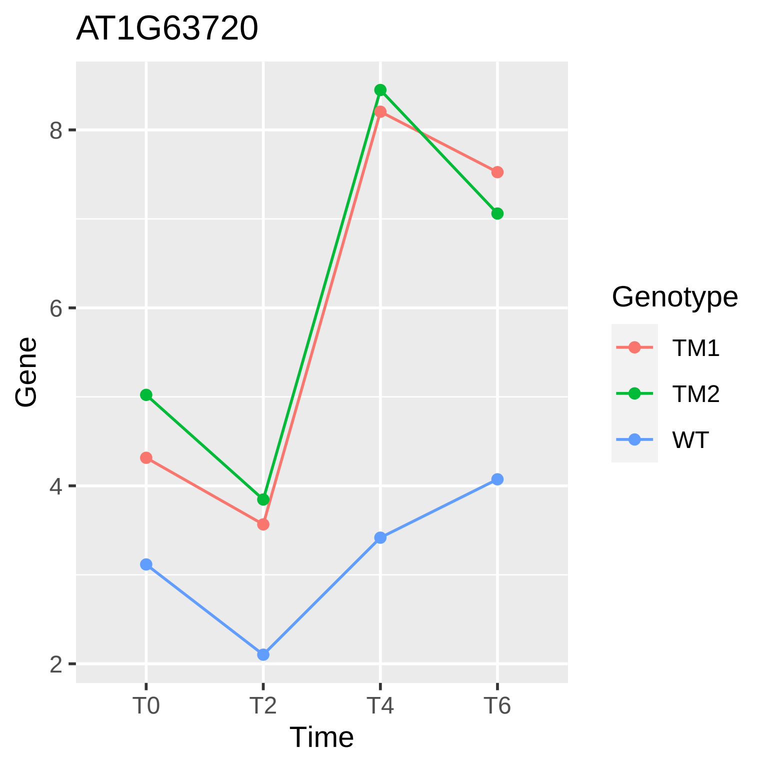

AT1G63720
Gene
8
6
4
2
T0
T2
T4
T6
Time
Genotype
TM1
TM2
WT

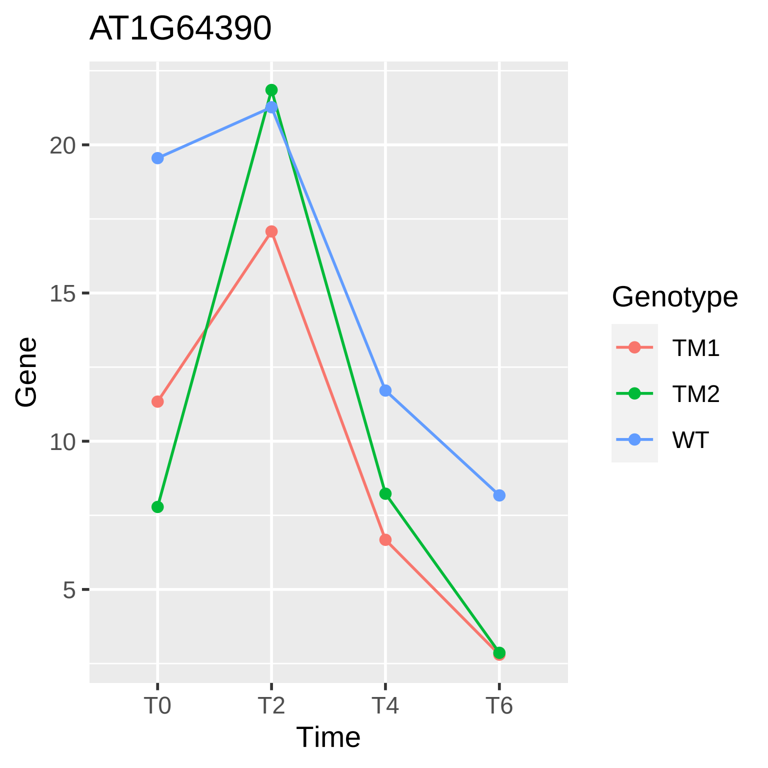

AT1G64390
Gene
20
15
10
5
T0
T2
T4
T6
Time
Genotype
TM1
TM2
WT

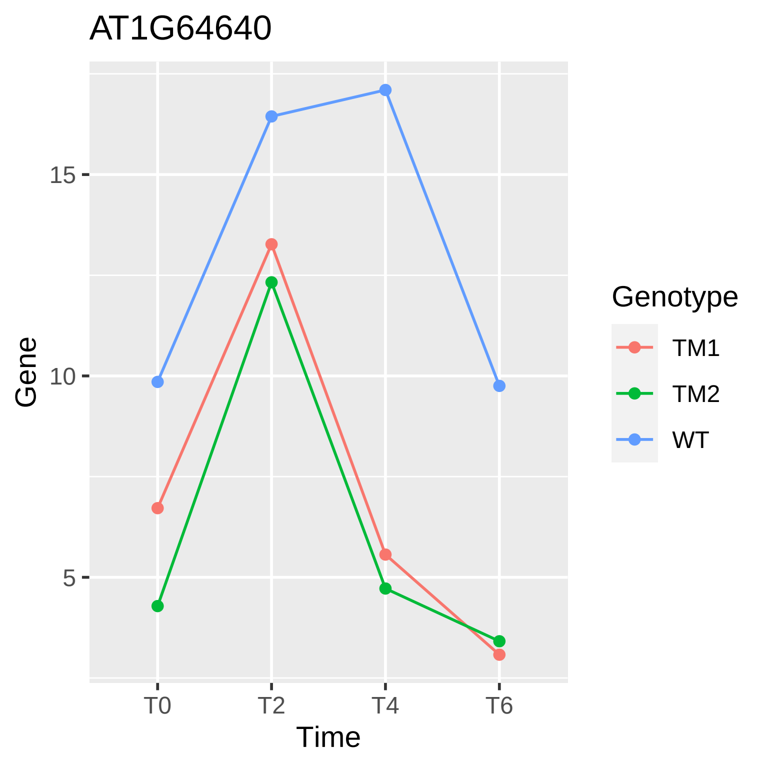
AT1G64640
Gene
15
10
5
T0
T2
T4
T6
Time
Genotype
TM1
TM2
WT

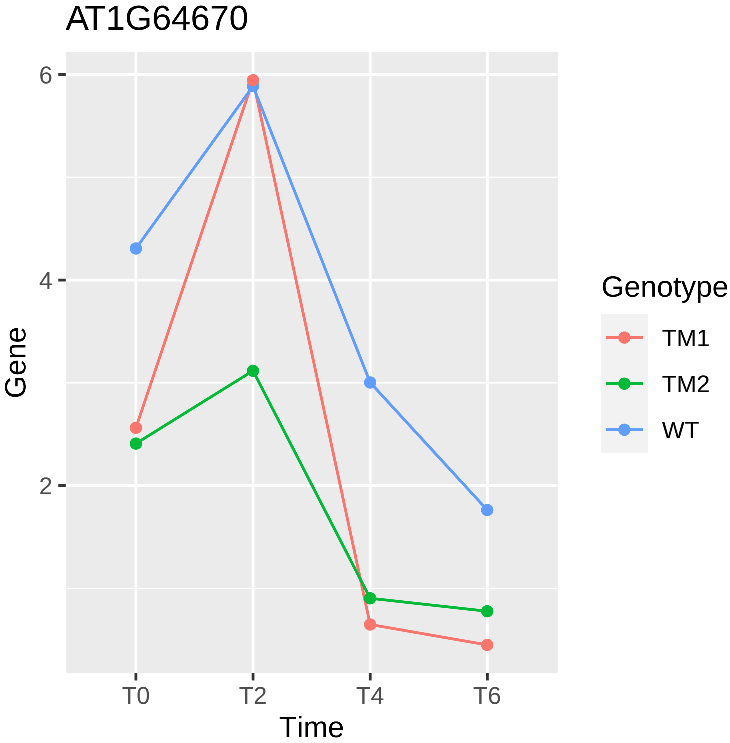

AT1G64670
6
4
2
Gene
T0
T2
T4
T6
Time
Genotype
TM1
TM2
WT

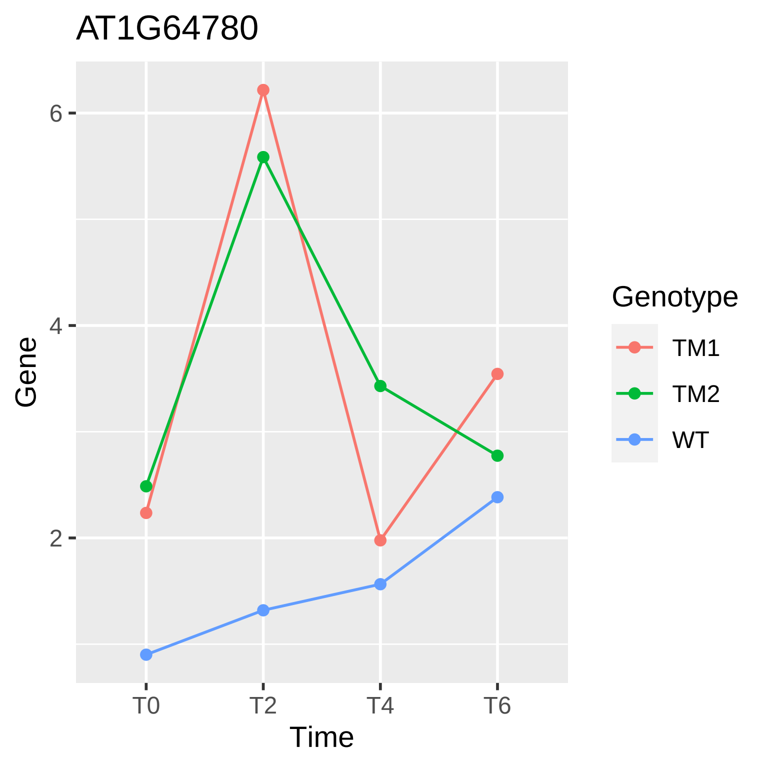

AT1G64780
Gene
6
4
2
T0
T2
T4
T6
Time
Genotype
TM1
TM2
WT

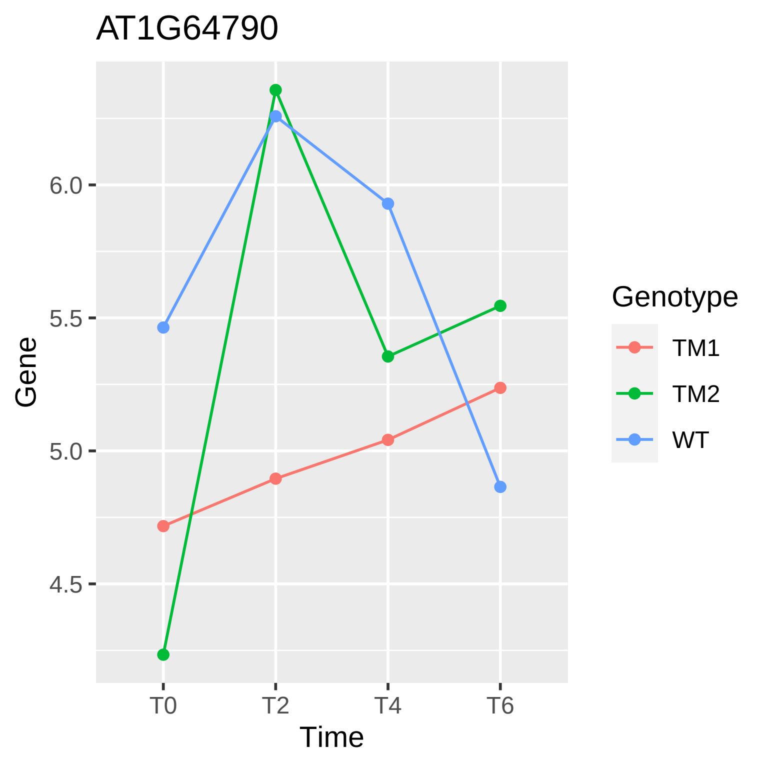

AT1G64790
Gene
6.0
5.5
5.0
4.5
T0
T2
T4
T6
Time
Genotype
TM1
TM2
WT

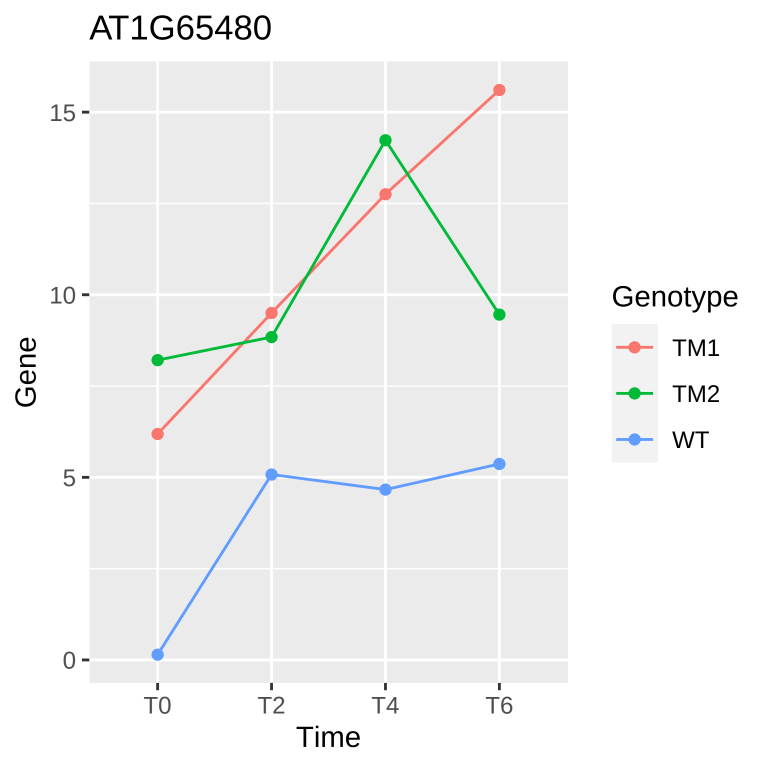
AT1G65480
Gene
15
10
5
0
T0
T2
T4
T6
Time
Genotype
TM1
TM2
WT

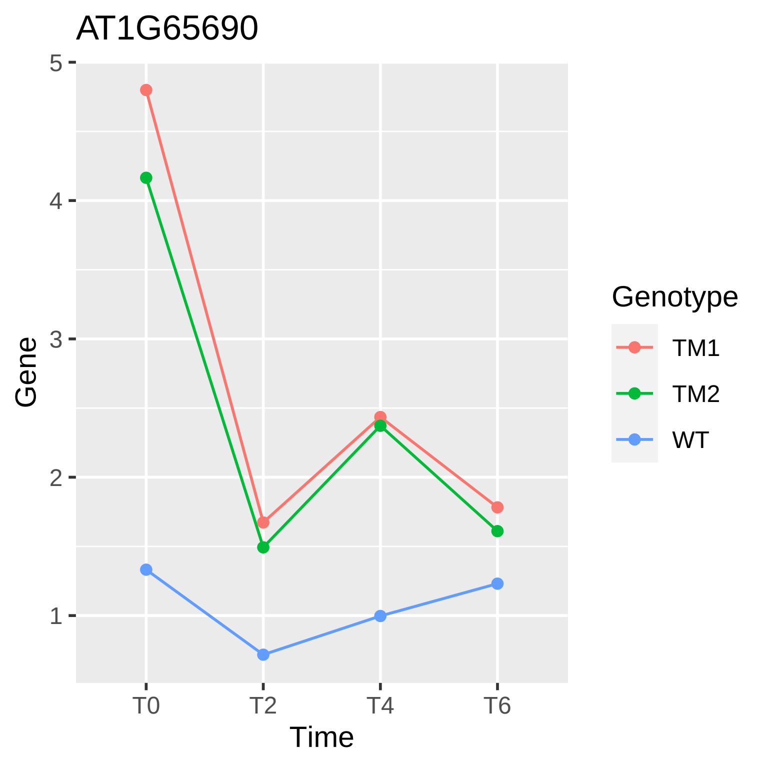

AT1G65690
Gene
5
4
3
2
1
T0
T2
T4
T6
Time
Genotype
TM1
TM2
WT

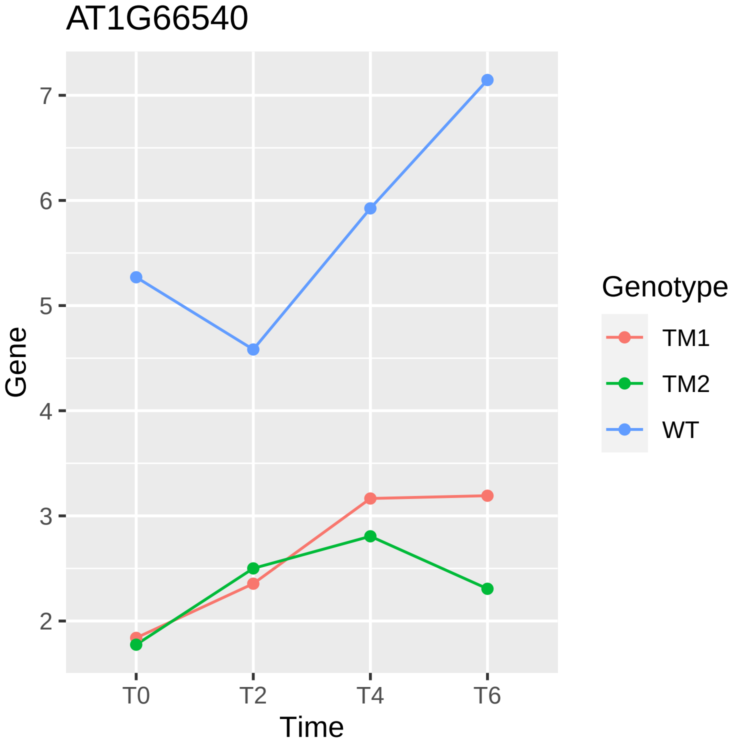
AT1G66540
7
6
5
4
3
2
Gene
T0
T2
T4
T6
Time
Genotype
TM1
TM2
WT

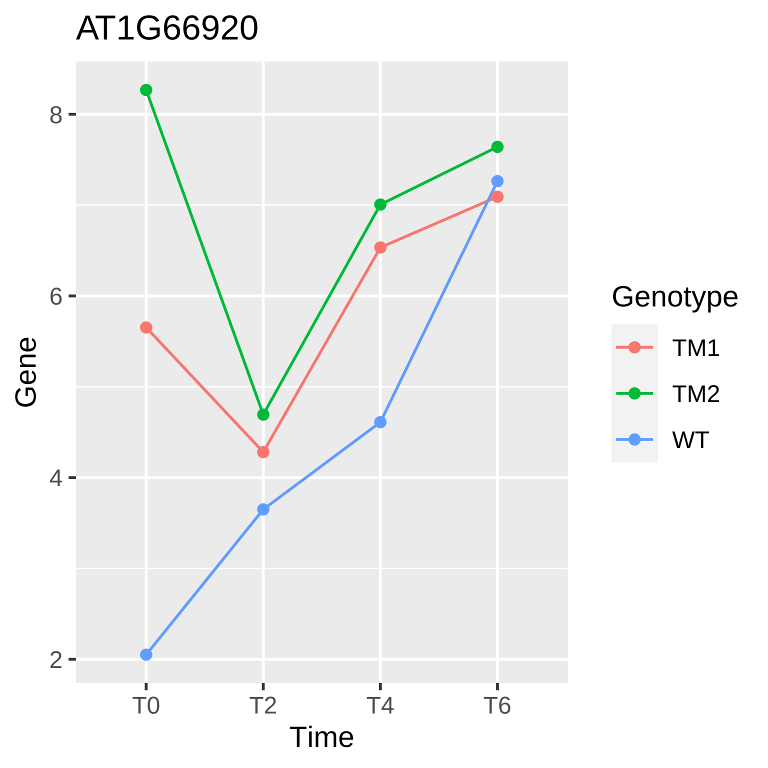

AT1G66920
Gene
Time
T0
T2
T4
T6
2
4
6
8
Genotype
TM1
TM2
WT

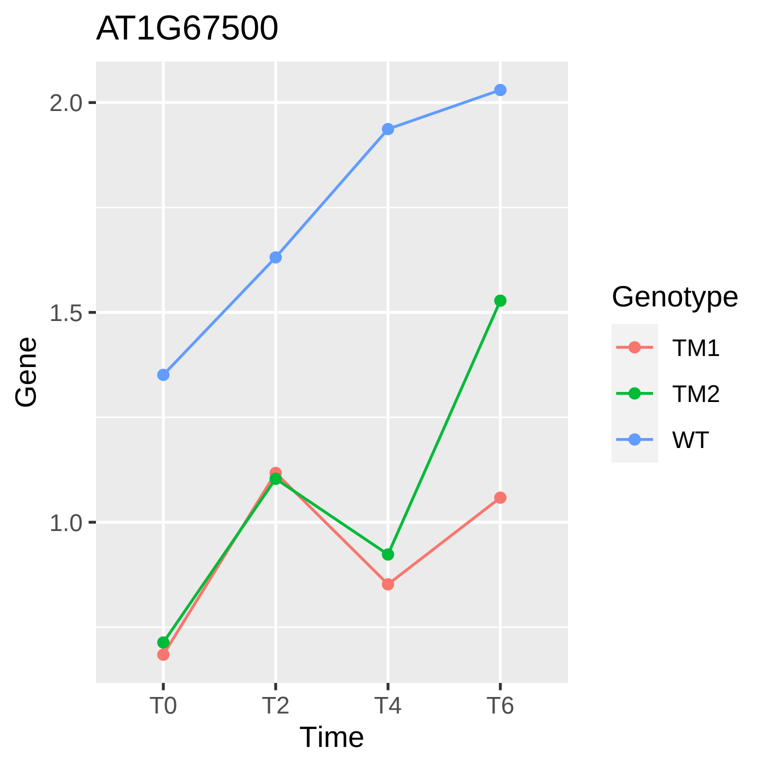
AT1G67500
Gene
2.0
1.5
1.0
T0
T2
T4
T6
Time
Genotype
TM1
TM2
WT

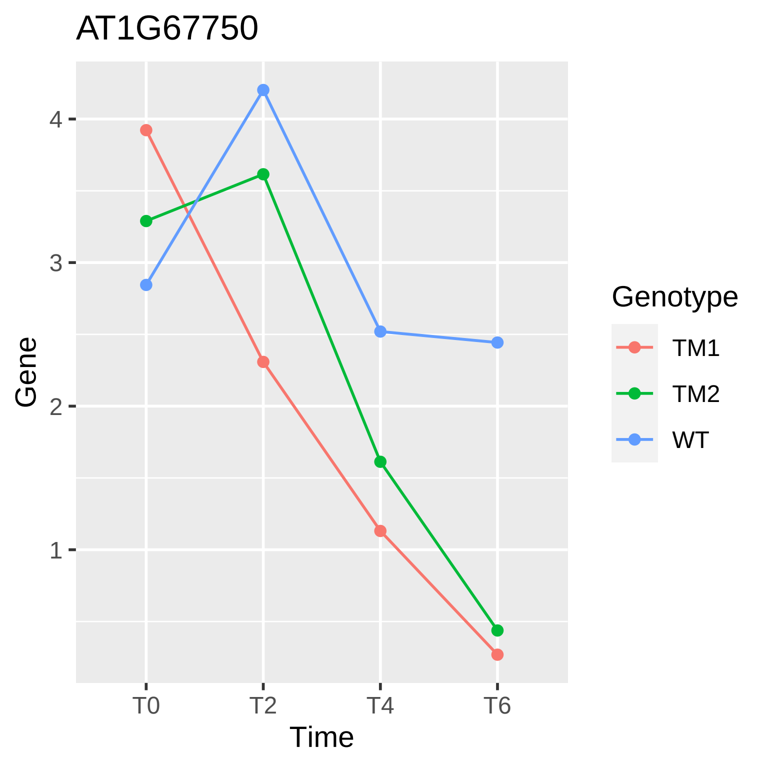
AT1G67750
Gene
4
3
2
1
T0
T2
T4
T6
Time
Genotype
TM1
TM2
WT

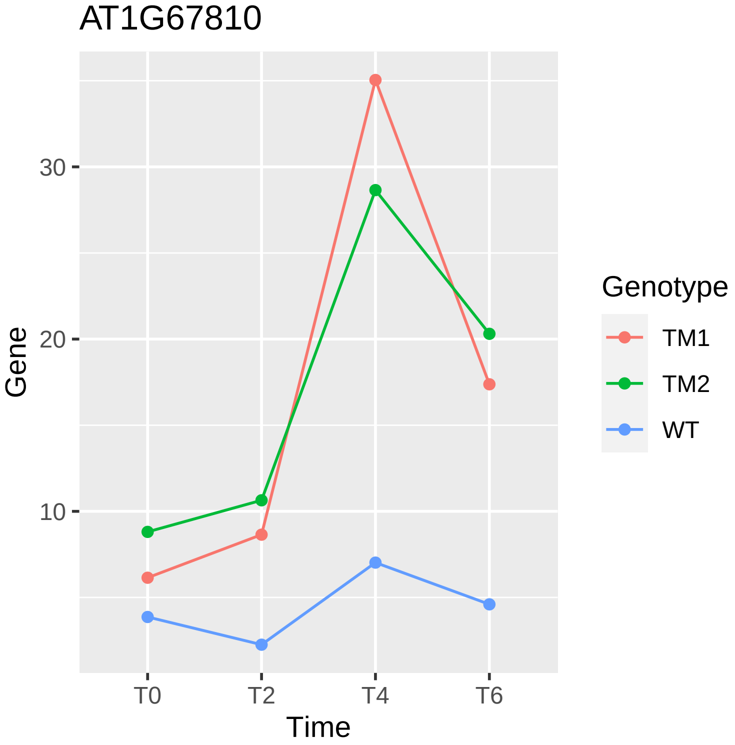
AT1G67810
Gene
30
20
10
T0
T2
T4
T6
Time
Genotype
TM1
TM2
WT

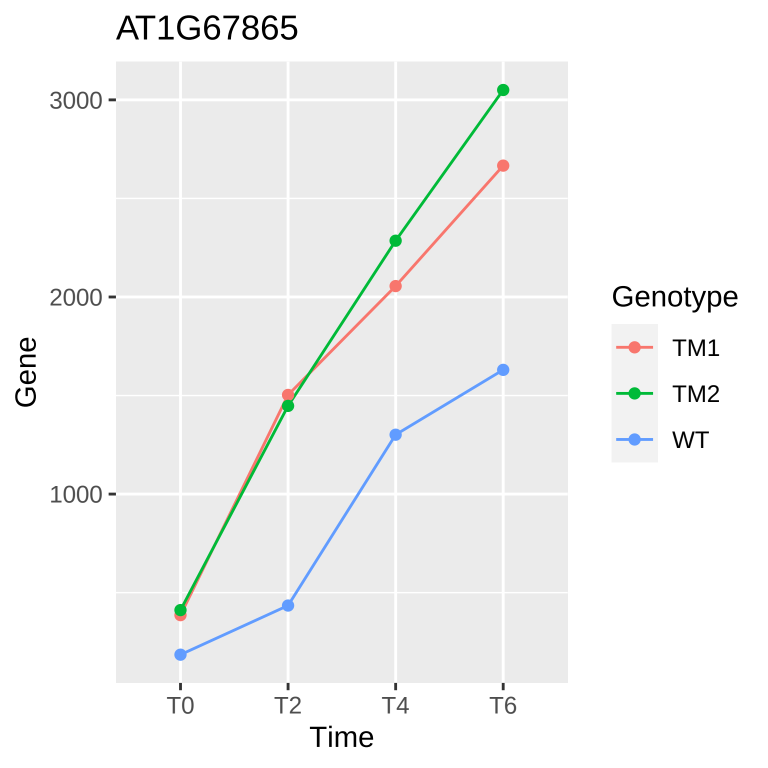

AT1G67865
3000
2000
1000
Gene
T0
T2
T4
T6
Time
Genotype
TM1
TM2
WT

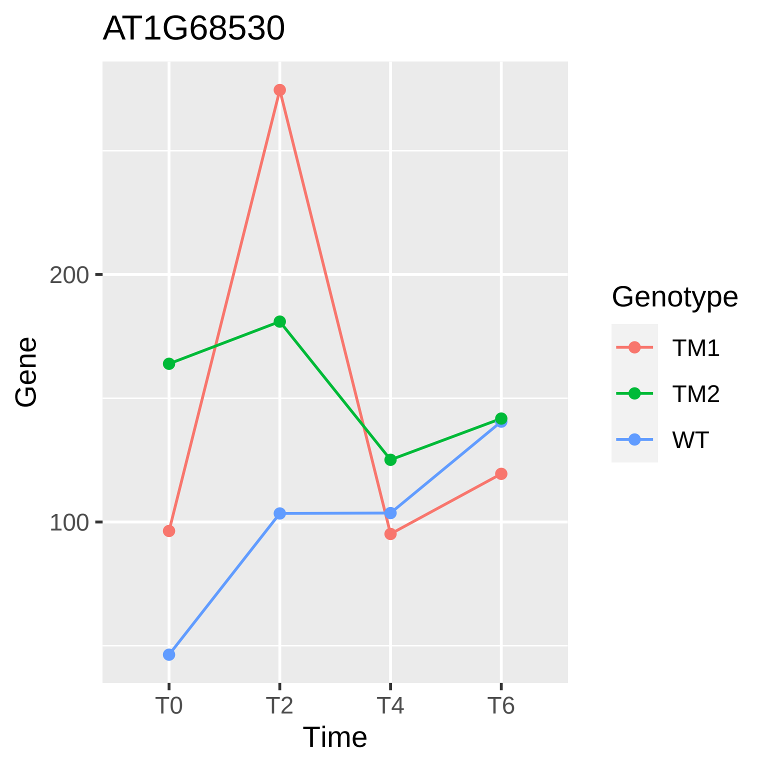
AT1G68530
200
100
Gene
T0
T2
T4
T6
Time
Genotype
TM1
TM2
WT

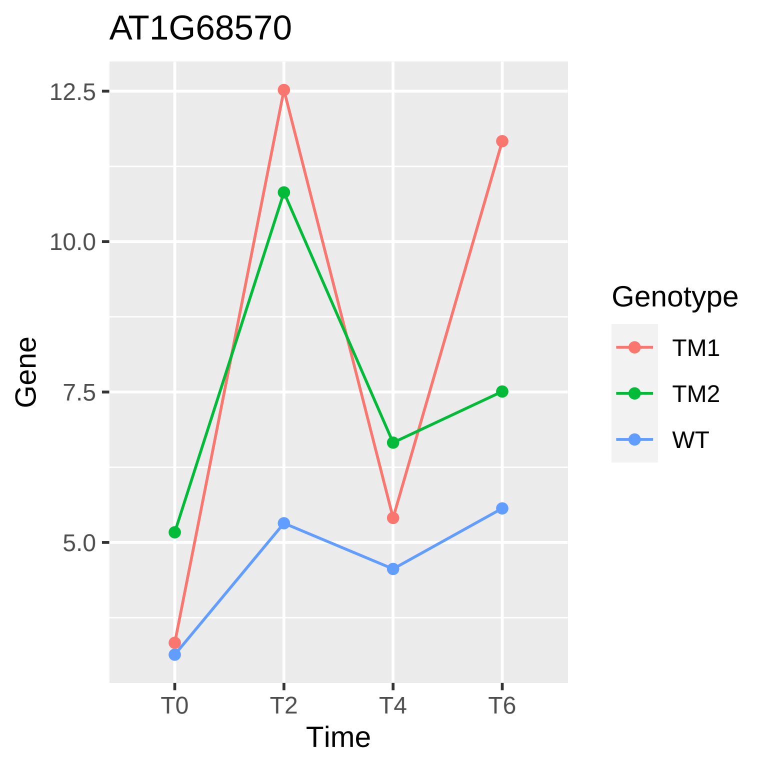

AT1G68570
12.5
10.0
7.5
5.0
Gene
T0
T2
T4
T6
Time
Genotype
TM1
TM2
WT

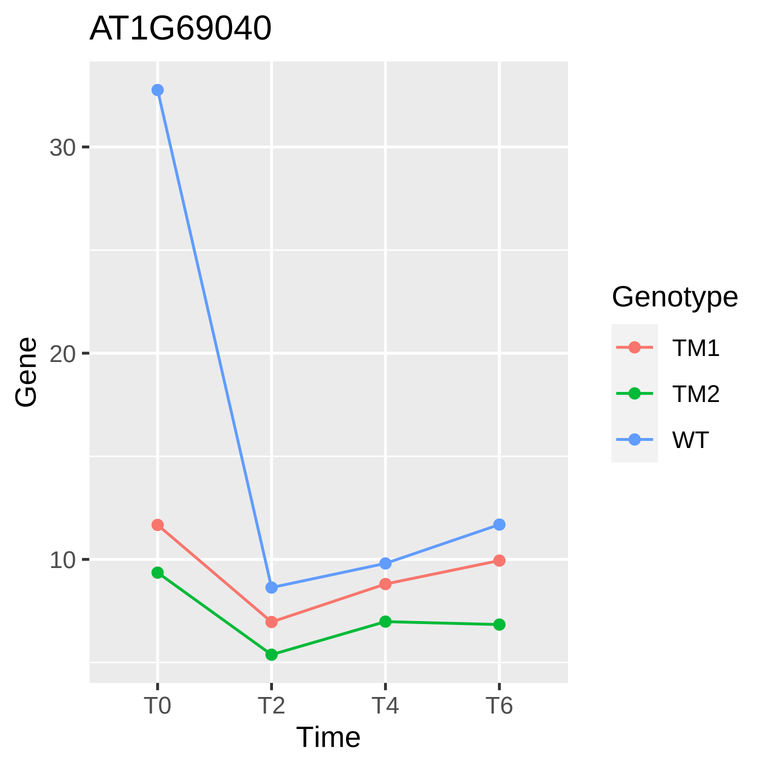

AT1G69040
30
20
10
Gene
T0
T2
T4
T6
Time
Genotype
TM1
TM2
WT

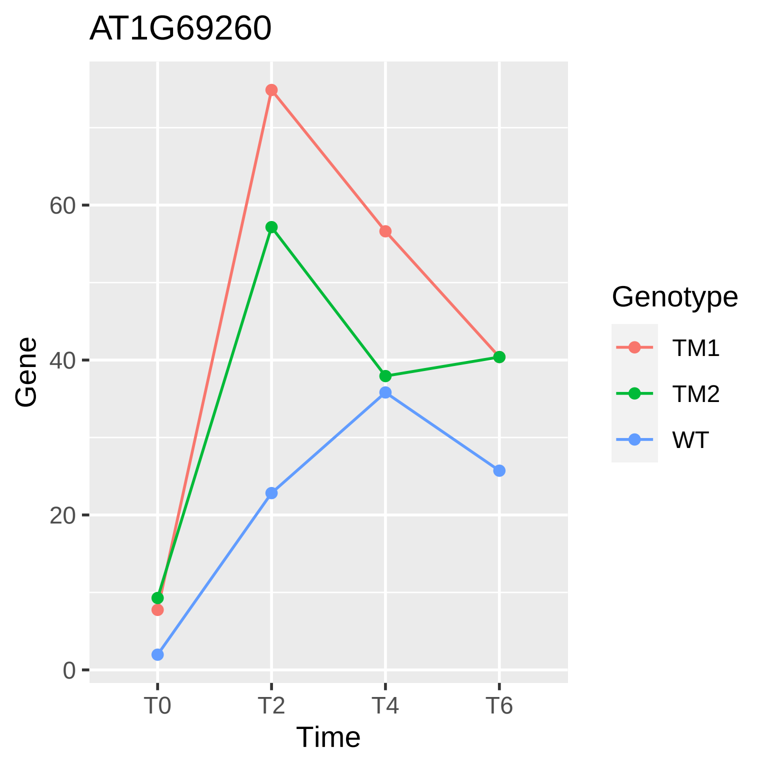

AT1G69260
Gene
60
40
20
0
T0
T2
T4
T6
Time
Genotype
TM1
TM2
WT

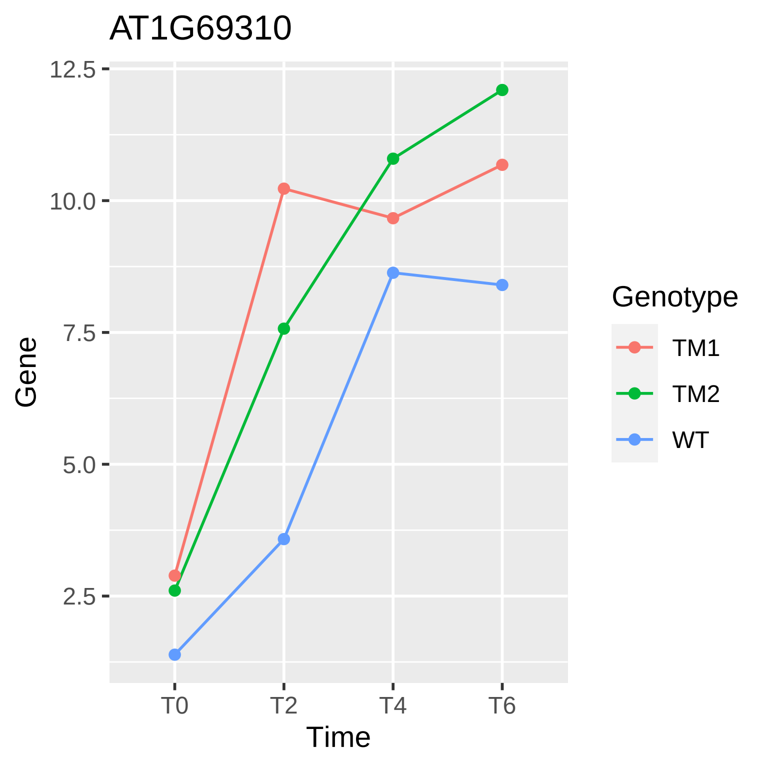
AT1G69310
12.5
10.0
7.5
5.0
2.5
Gene
T0
T2
T4
T6
Time
Genotype
TM1
TM2
WT

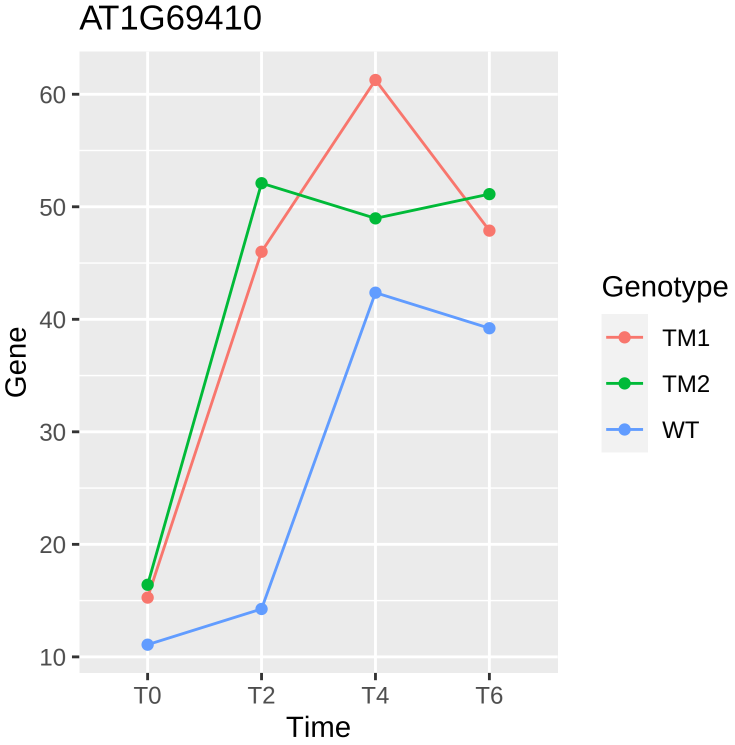

AT1G69410
Gene
60
50
40
30
20
10
T0
T2
T4
T6
Time
Genotype
TM1
TM2
WT

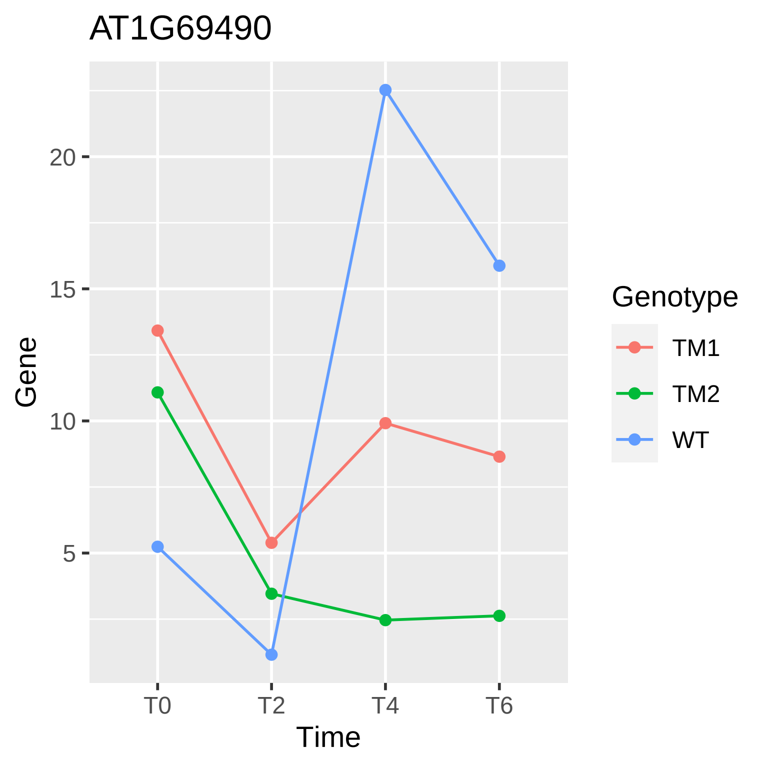

AT1G69490
Gene
20
15
10
5
T0
T2
T4
T6
Time
Genotype
TM1
TM2
WT

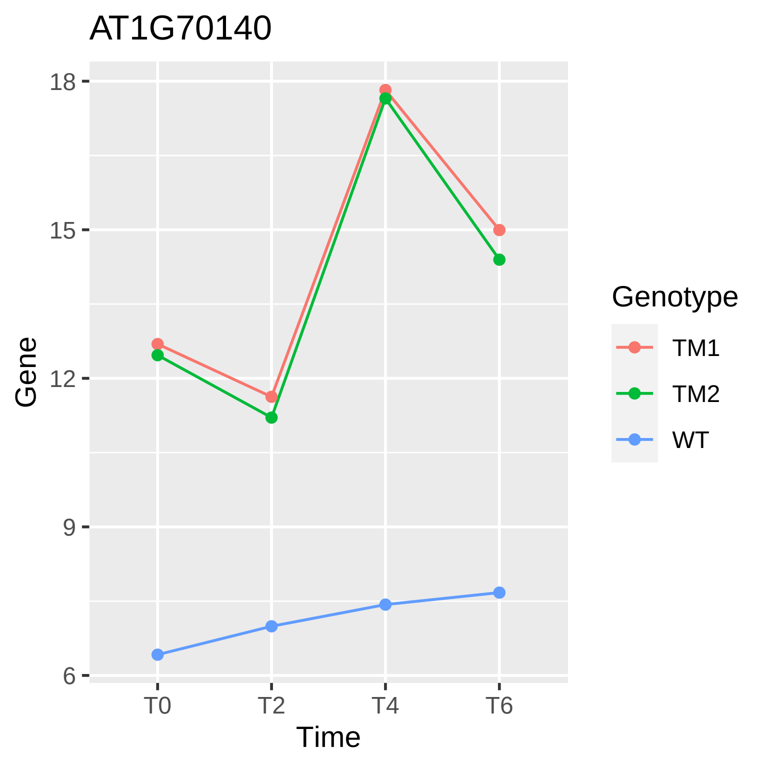
AT1G70140
Gene
18
15
12
9
6
T0
T2
T4
T6
Time
Genotype
TM1
TM2
WT

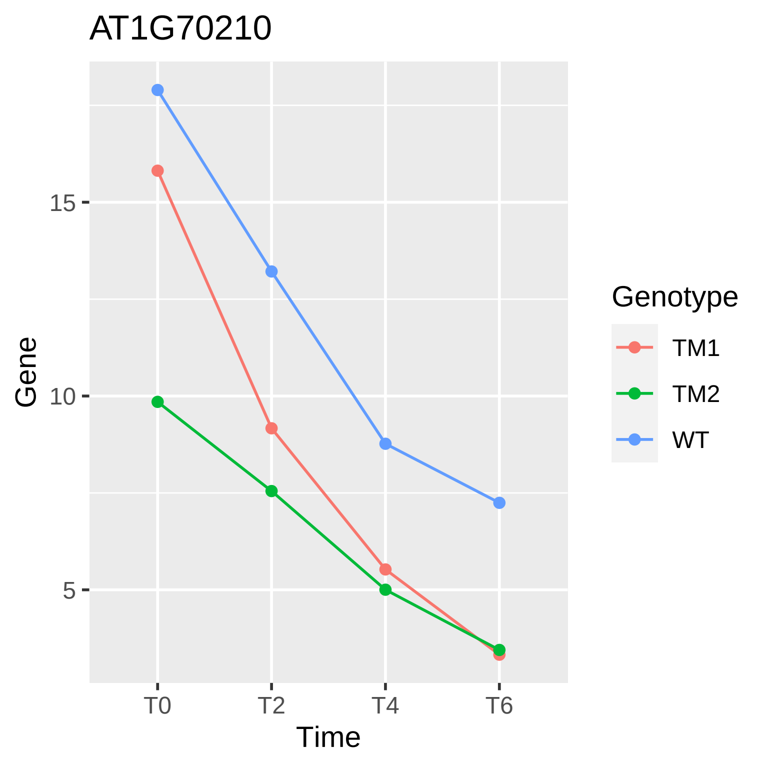

AT1G70210
Gene
15
10
5
T0
T2
T4
T6
Time
Genotype
TM1
TM2
WT

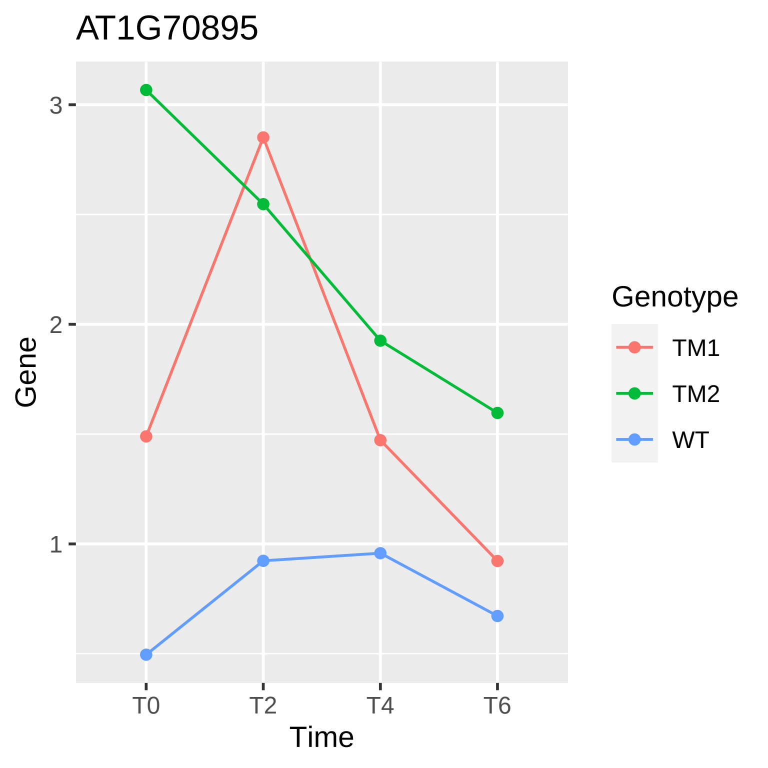
AT1G70895
Gene
3
2
1
T0
T2
T4
T6
Time
Genotype
TM1
TM2
WT

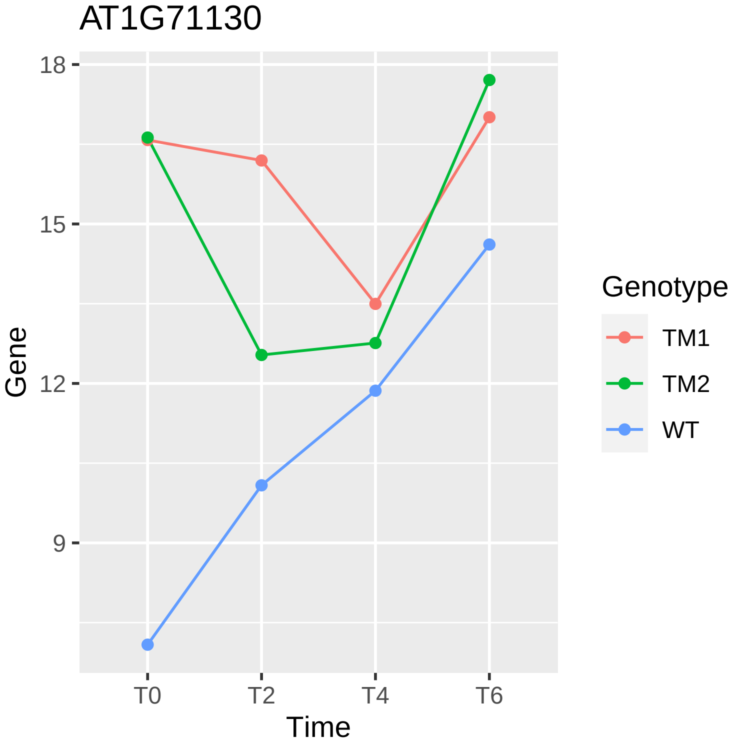

AT1G71130
18
15
12
9
Gene
T0
T2
T4
T6
Time
Genotype
TM1
TM2
WT

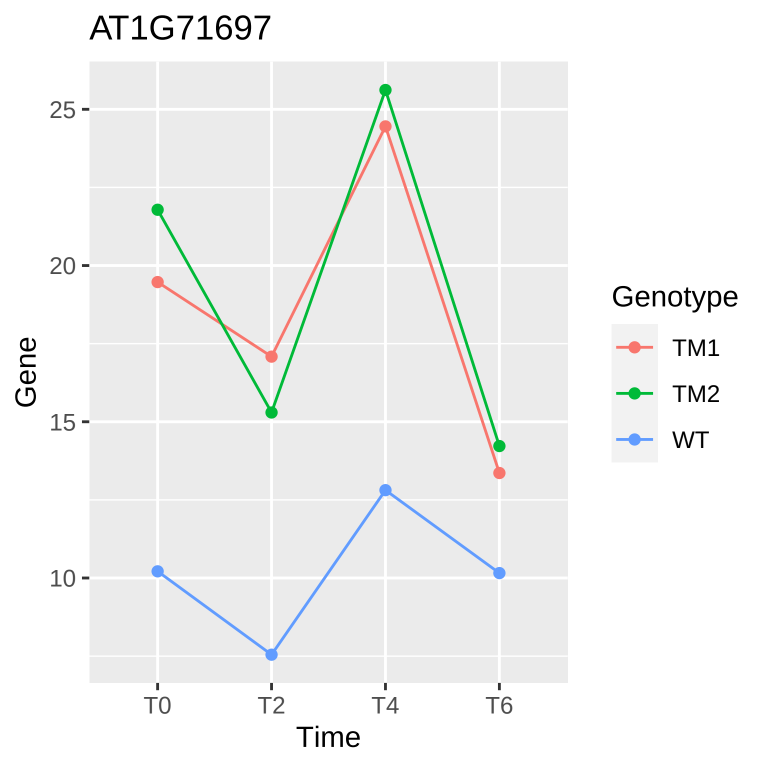
AT1G71697
25
20
15
10
Gene
T0
T2
T4
T6
Time
Genotype
TM1
TM2
WT

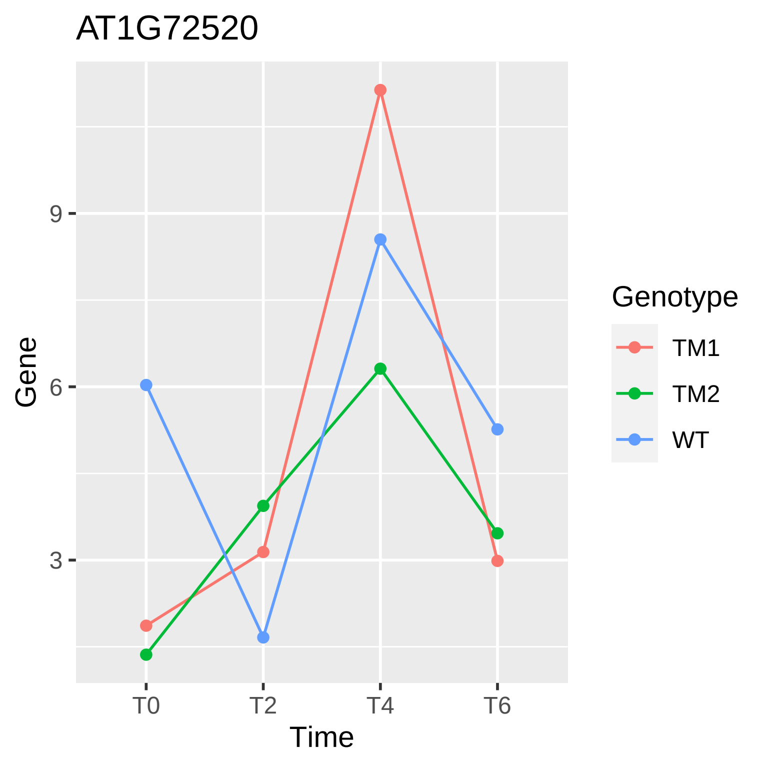
AT1G72520
Gene
9
6
3
T0
T2
T4
T6
Time
Genotype
TM1
TM2
WT

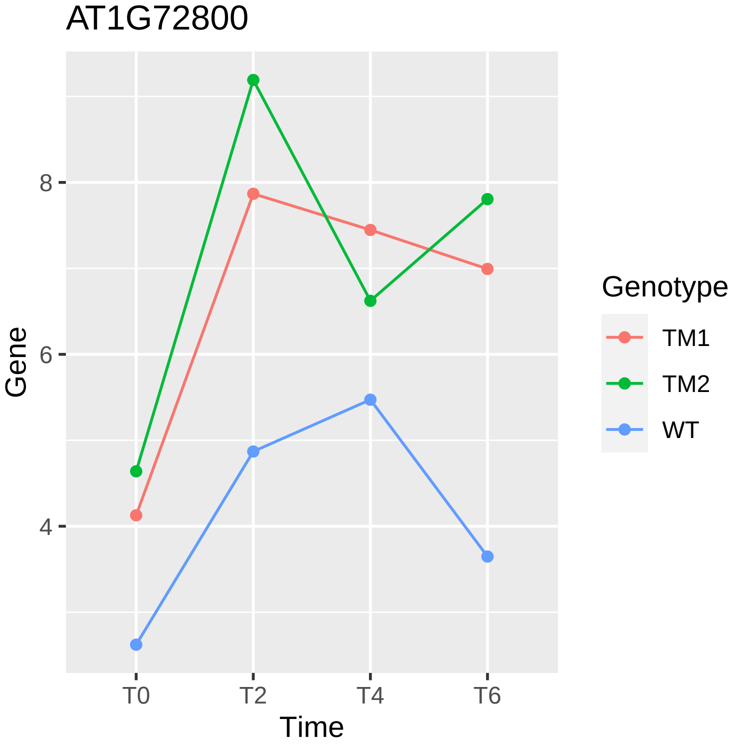

AT1G72800
Gene
8
6
4
T0
T2
T4
T6
Time
Genotype
TM1
TM2
WT

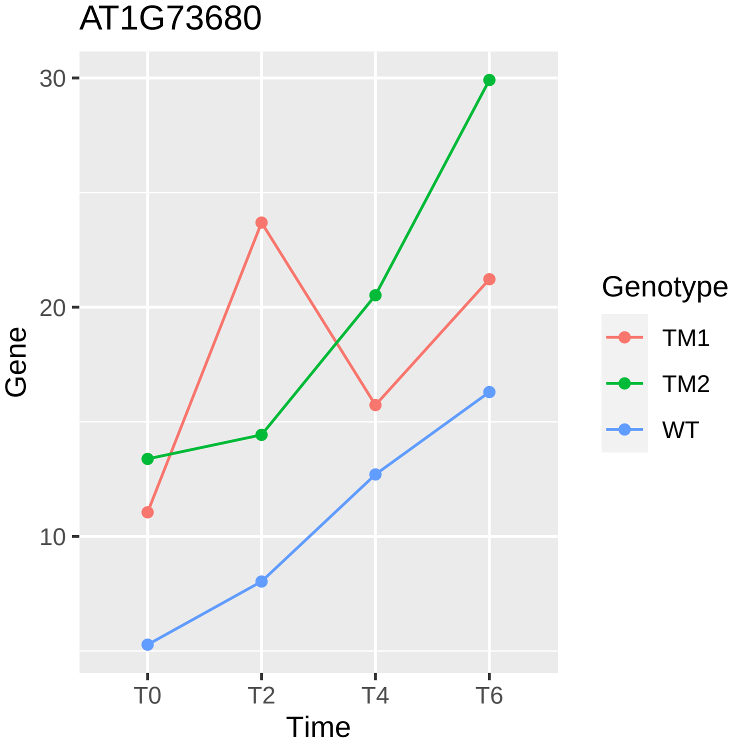

AT1G73680
30
20
10
Gene
T0
T2
T4
T6
Time
Genotype
TM1
TM2
WT

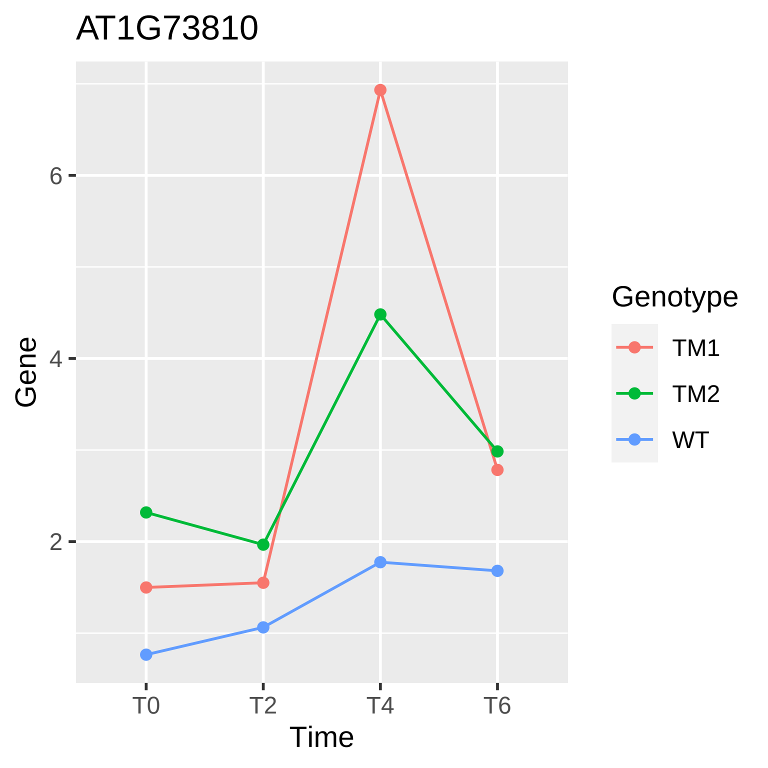

AT1G73810
Gene
6
4
2
T0
T2
T4
T6
Time
Genotype
TM1
TM2
WT

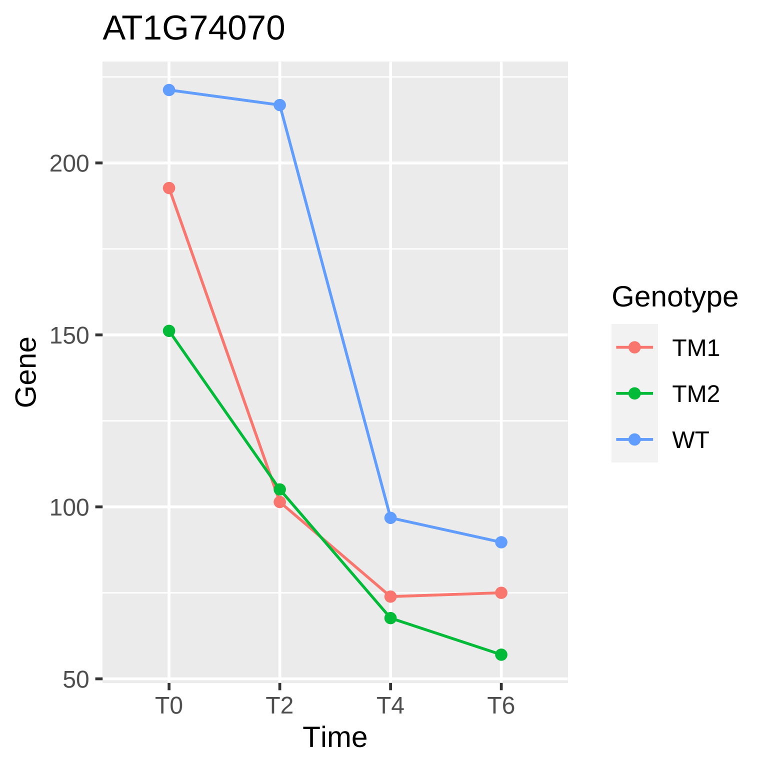

AT1G74070
Gene
200
150
100
50
T0
T2
T4
T6
Time
Genotype
TM1
TM2
WT

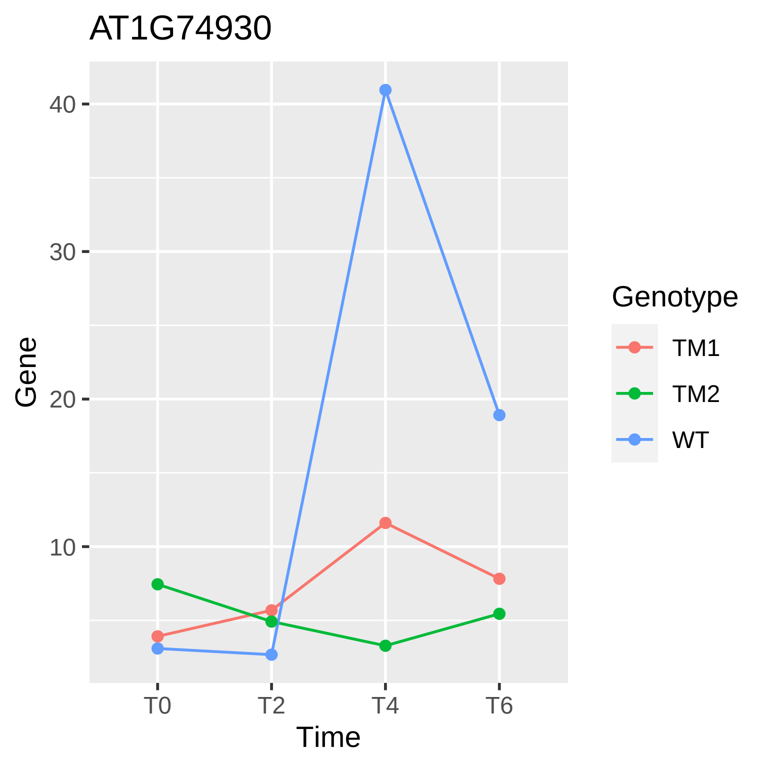
AT1G74930
40
30
20
10
Gene
T0
T2
T4
T6
Time
Genotype
TM1
TM2
WT

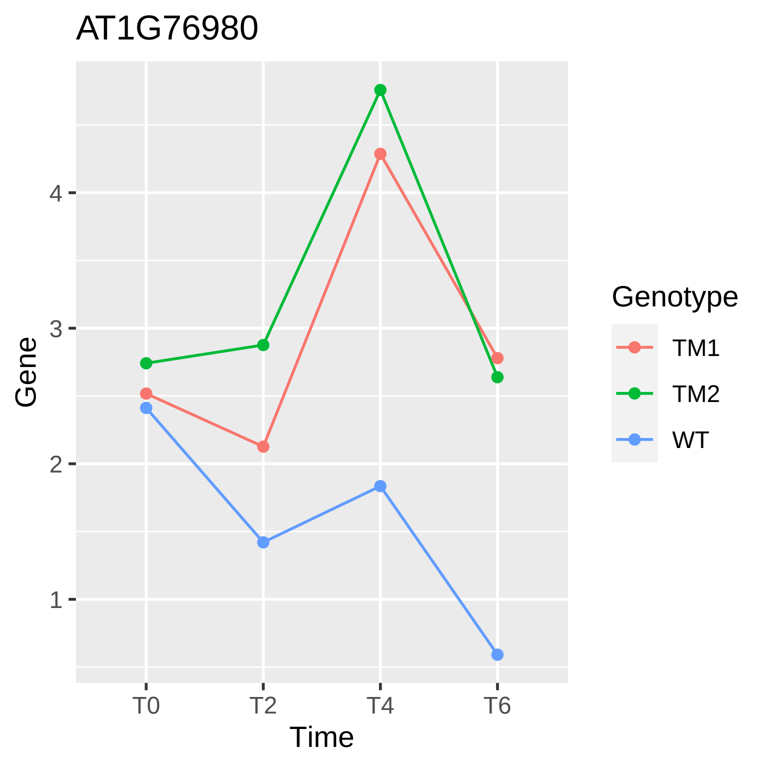
AT1G76980
Gene
4
3
2
1
T0
T2
T4
T6
Time
Genotype
TM1
TM2
WT

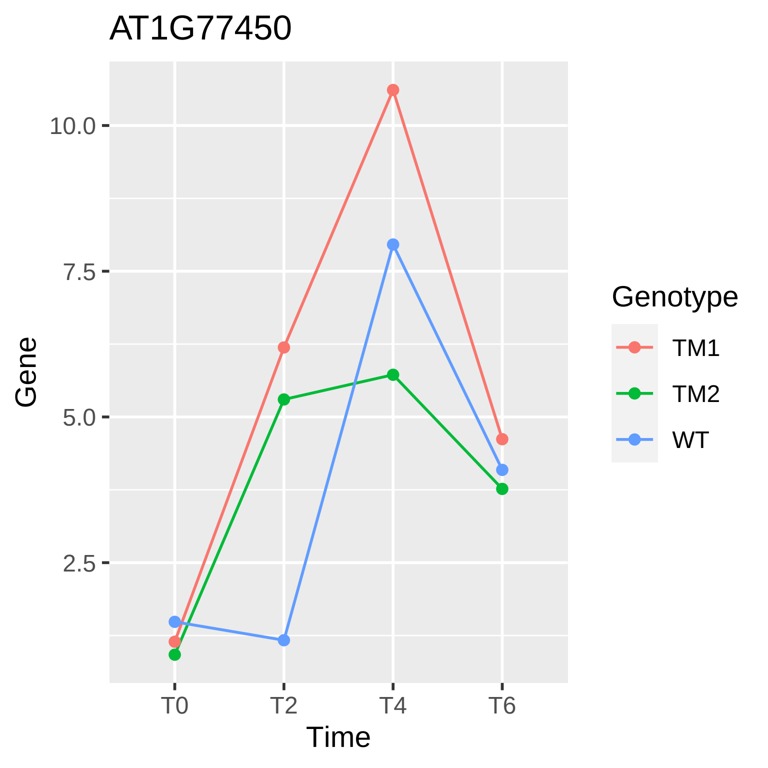
AT1G77450
10.0
7.5
5.0
2.5
Gene
T0
T2
T4
T6
Time
Genotype
TM1
TM2
WT

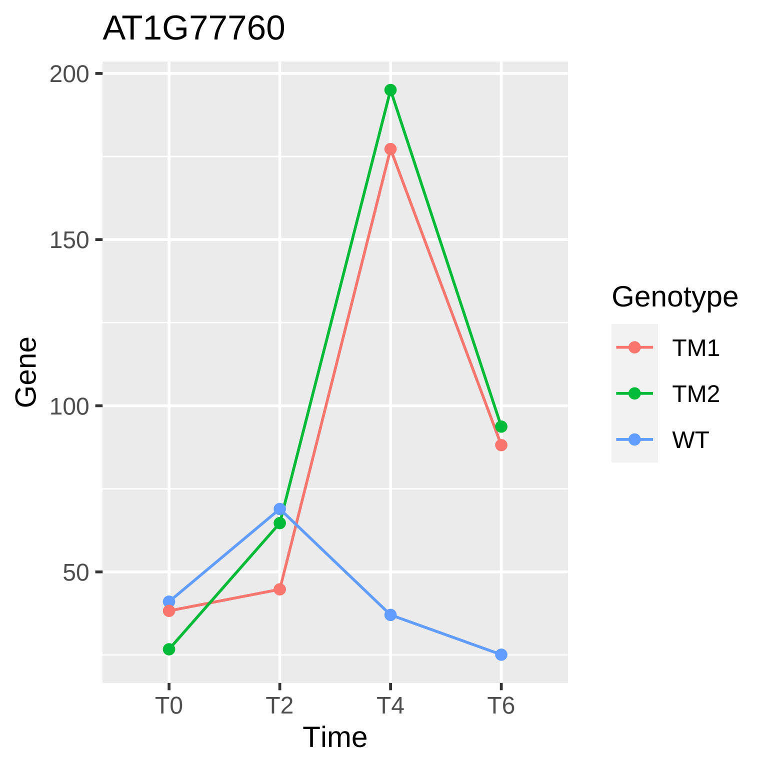

AT1G77760
200
150
100
50
Gene
T0
T2
T4
T6
Time
Genotype
TM1
TM2
WT

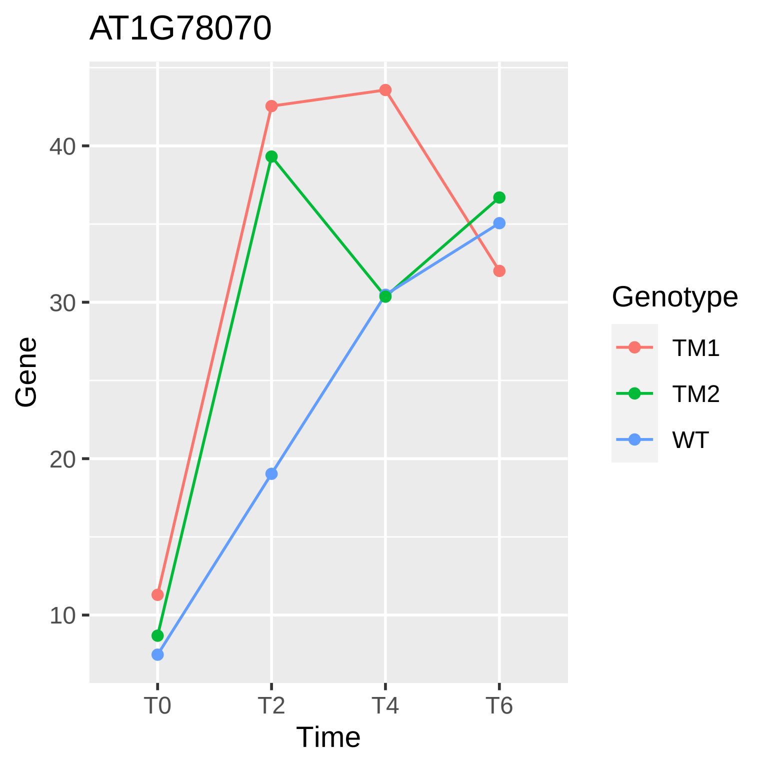

AT1G78070
40
30
20
10
Gene
T0
T2
T4
T6
Time
Genotype
TM1
TM2
WT

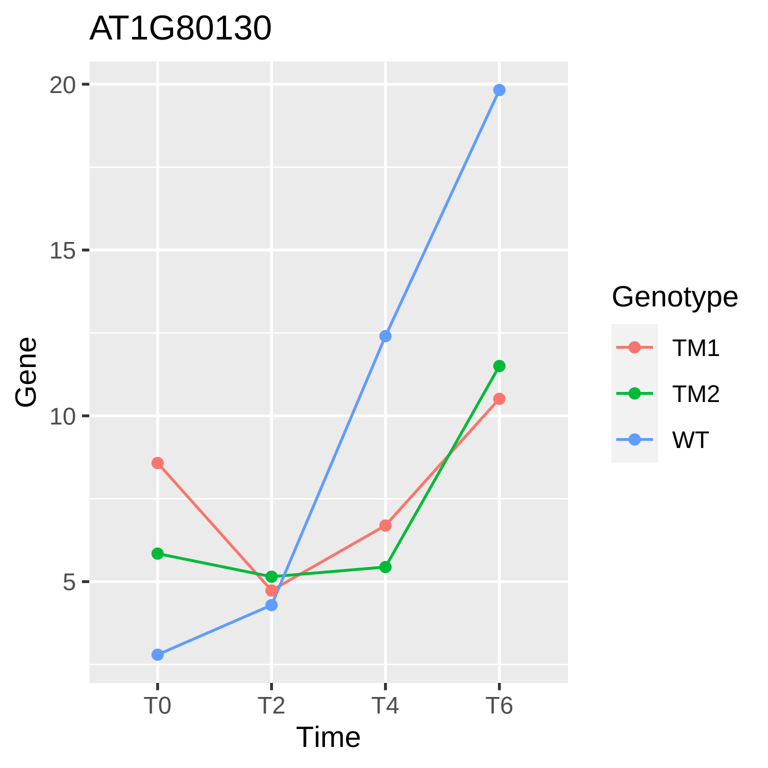
AT1G80130
Gene
20
15
10
5
T0
T2
T4
T6
Time
Genotype
TM1
TM2
WT

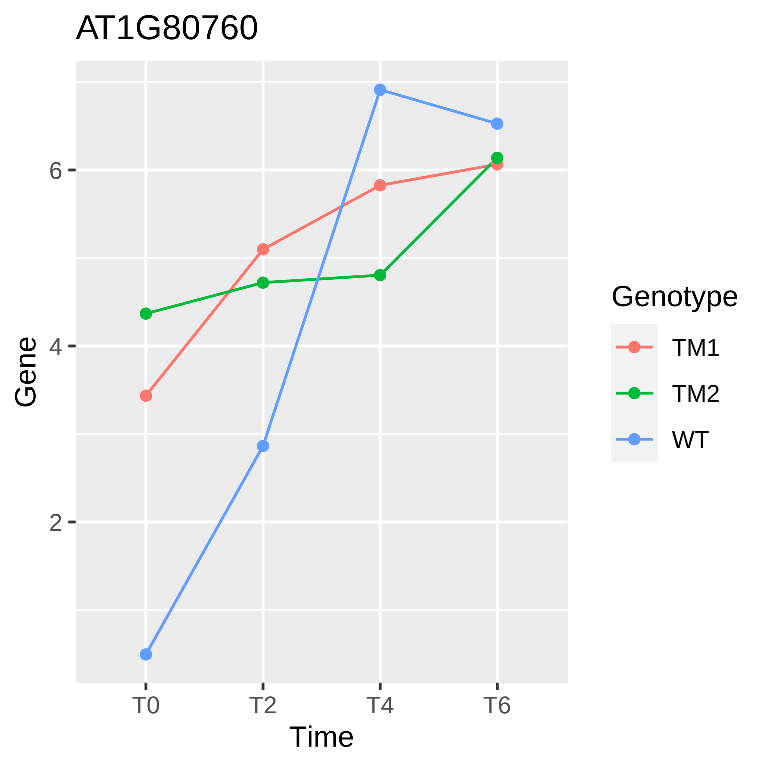
AT1G80760
Gene
6
4
2
T0
T2
T4
T6
Time
Genotype
TM1
TM2
WT

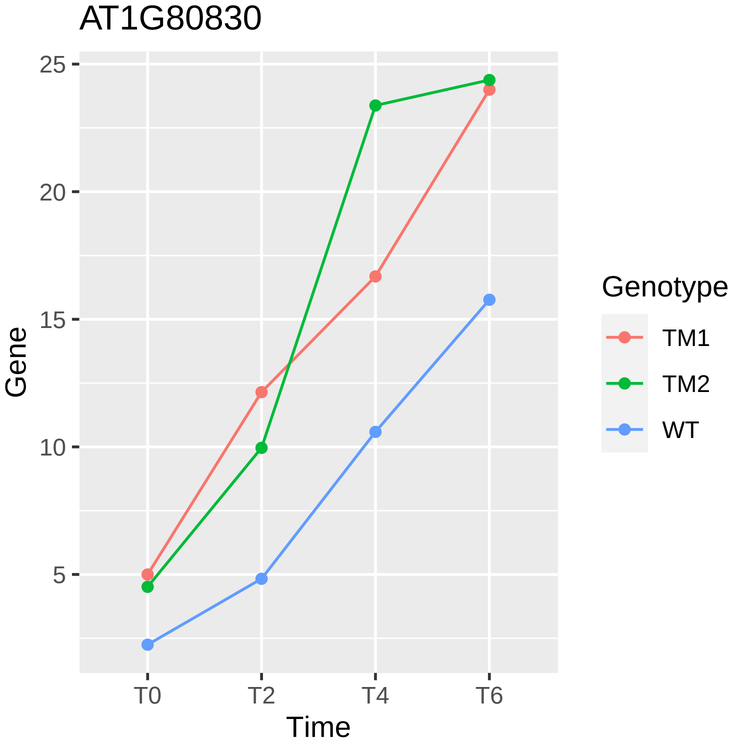
AT1G80830
25
20
15
10
5
Gene
T0
T2
T4
T6
Time
Genotype
TM1
TM2
WT

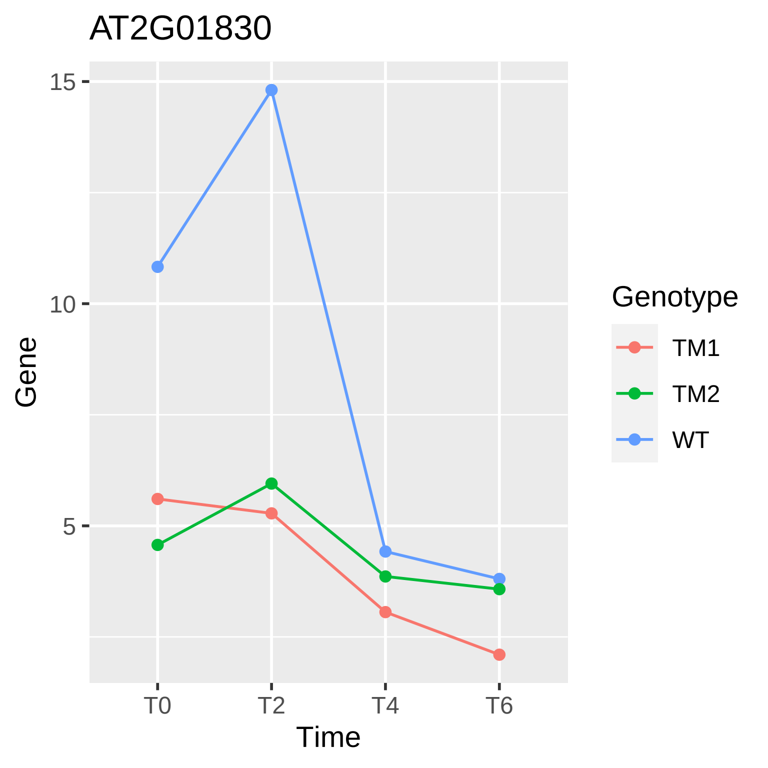

AT2G01830
15
10
5
Gene
T0
T2
T4
T6
Time
Genotype
TM1
TM2
WT

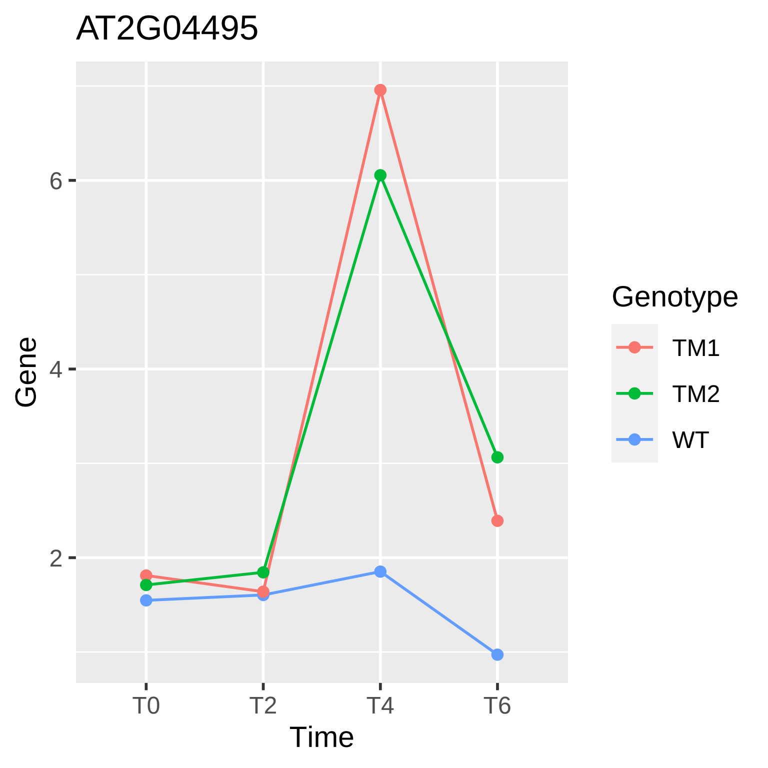
AT2G04495
Gene
6
4
2
T0
T2
T4
T6
Time
Genotype
TM1
TM2
WT

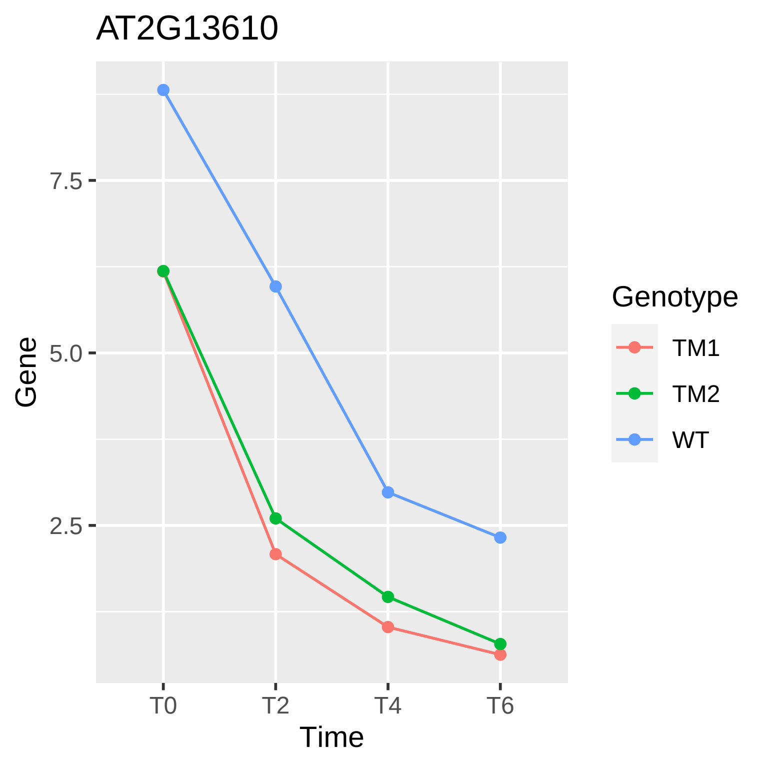

AT2G13610
Gene
7.5
5.0
2.5
T0
T2
T4
T6
Time
Genotype
TM1
TM2
WT

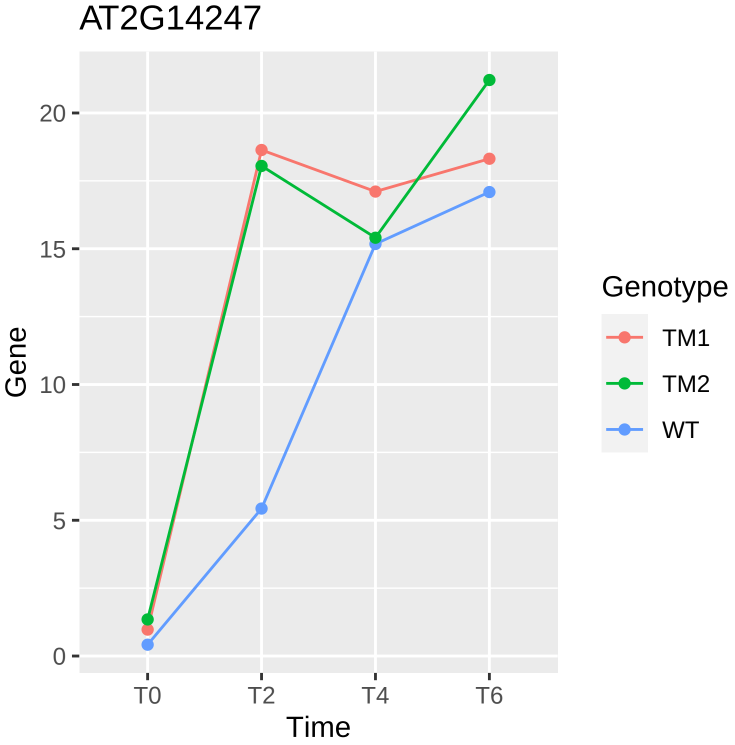

AT2G14247
Gene
0
5
10
15
20
T0
T2
T4
T6
Time
Genotype
TM1
TM2
WT

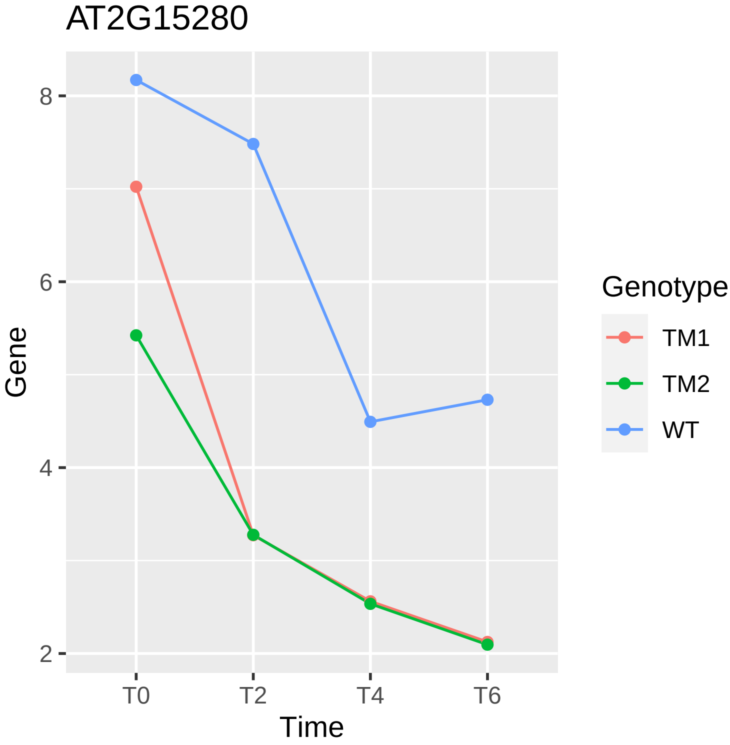
AT2G15280
Gene
8
6
4
2
T0
T2
T4
T6
Time
Genotype
TM1
TM2
WT

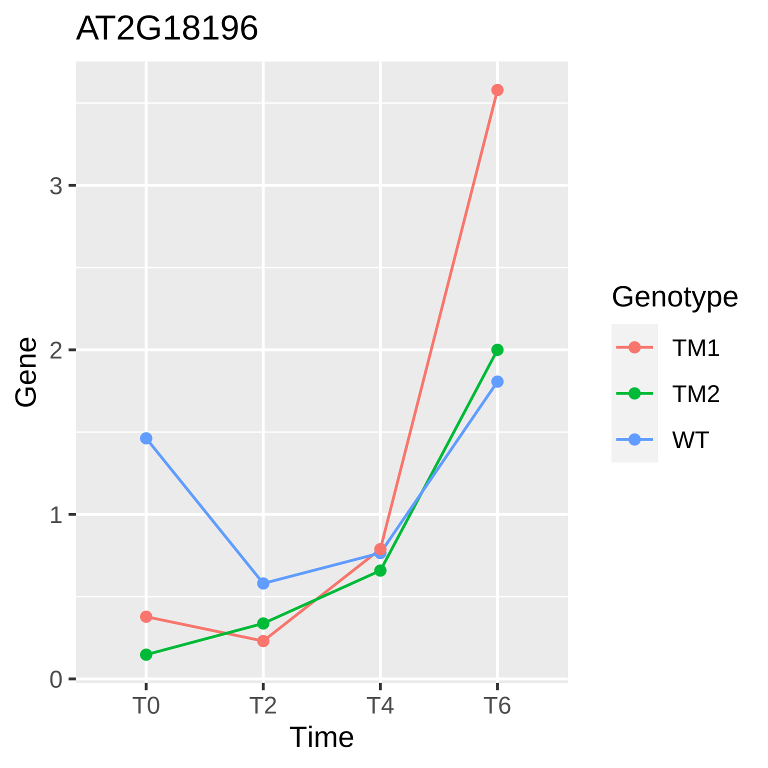
AT2G18196
Gene
Time
T0
T2
T4
T6
0
1
2
3
Genotype
TM1
TM2
WT

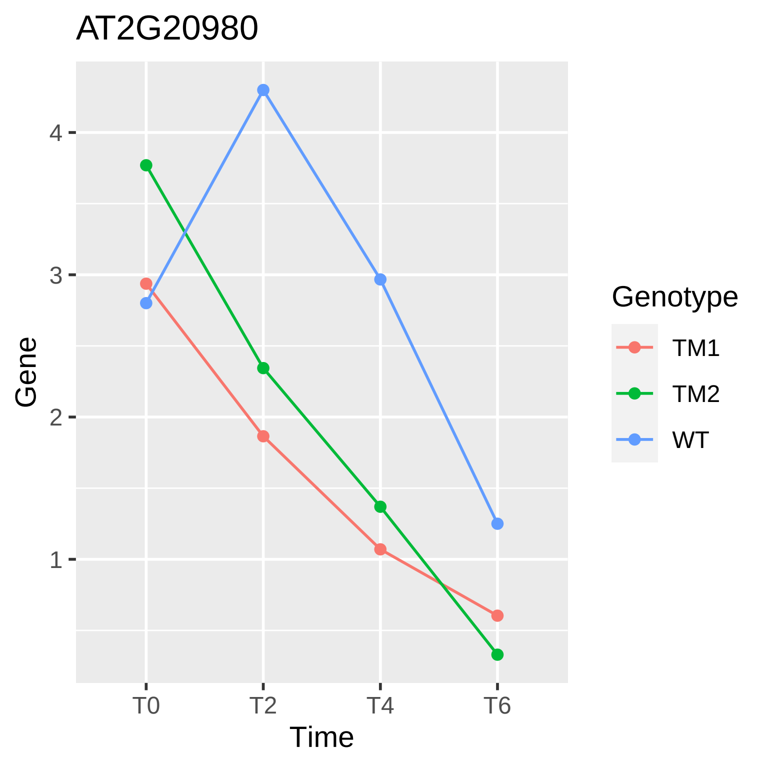
AT2G20980
Gene
4
3
2
1
T0
T2
T4
T6
Time
Genotype
TM1
TM2
WT

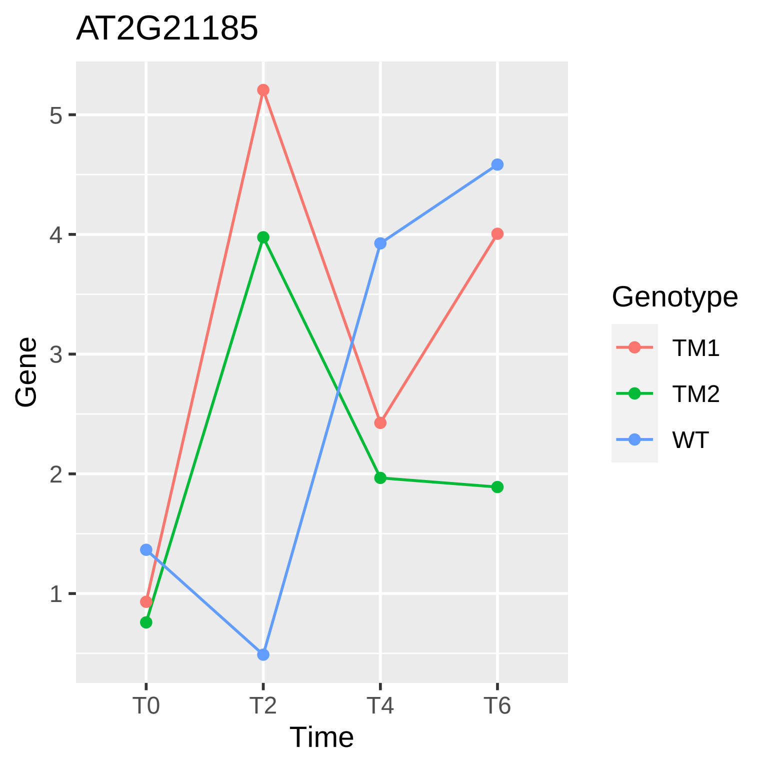

AT2G21185
Gene
5
4
3
2
1
T0
T2
T4
T6
Time
Genotype
TM1
TM2
WT

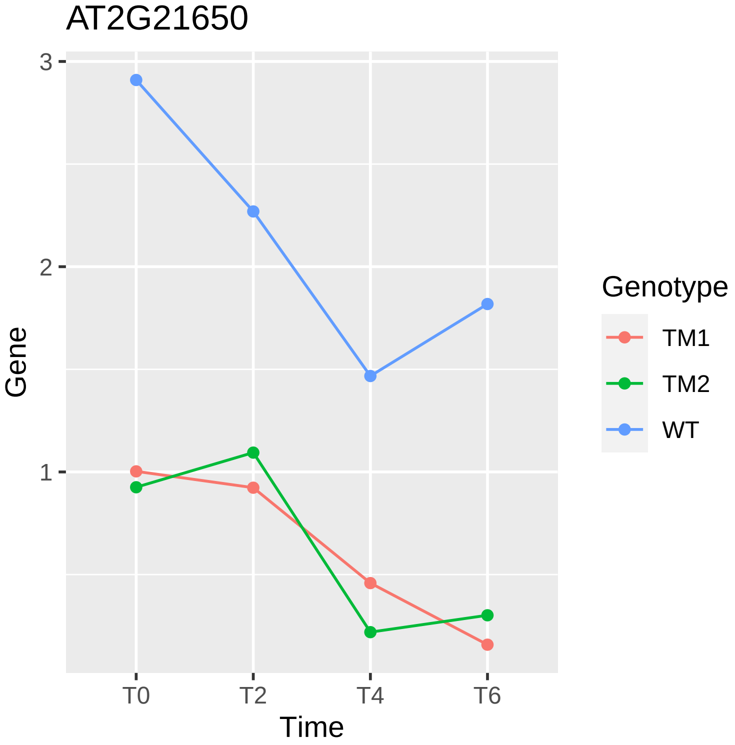

AT2G21650
Gene
3
2
1
T0
T2
T4
T6
Time
Genotype
TM1
TM2
WT

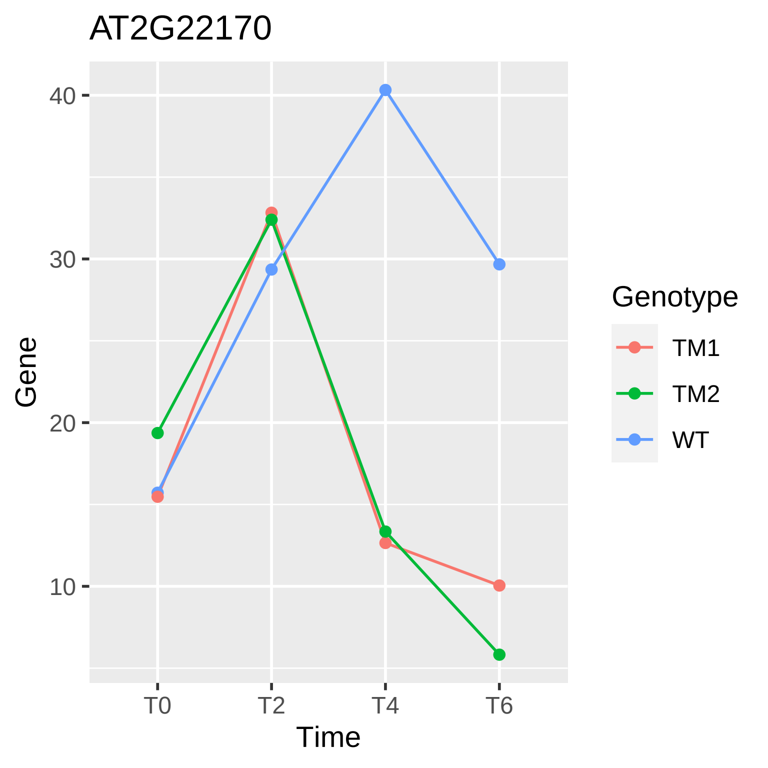

AT2G22170
40
30
20
10
Gene
T0
T2
T4
T6
Time
Genotype
TM1
TM2
WT

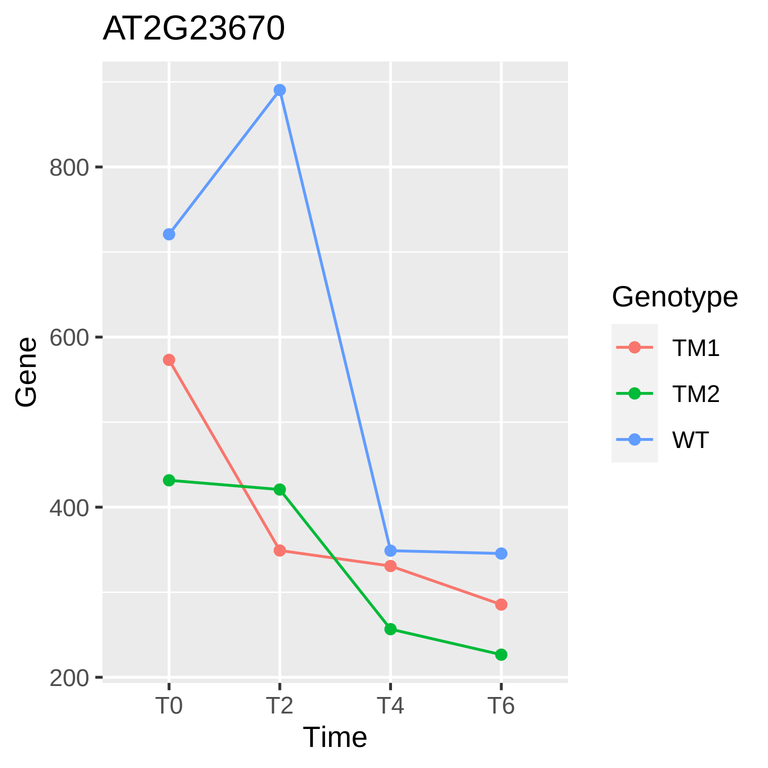
AT2G23670
800
600
400
200
Gene
T0
T2
T4
T6
Time
Genotype
TM1
TM2
WT

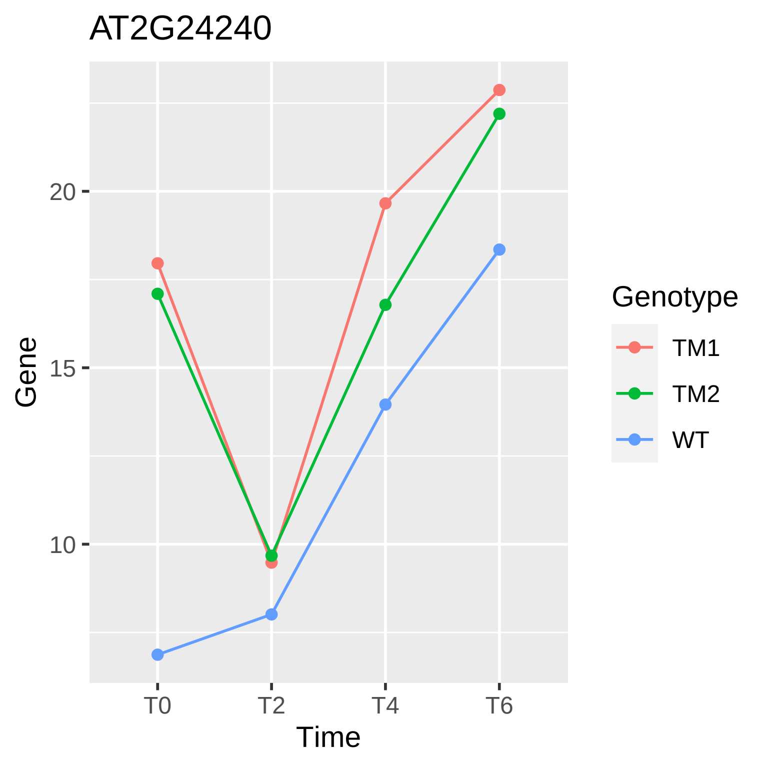
AT2G24240
Gene
20
15
10
T0
T2
T4
T6
Time
Genotype
TM1
TM2
WT

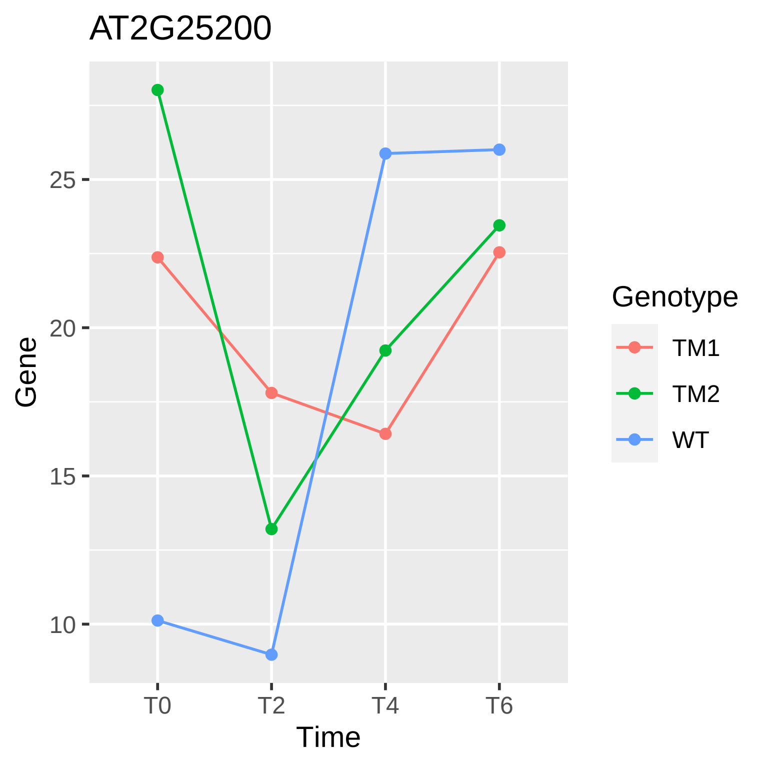

AT2G25200
Gene
25
20
15
10
T0
T2
T4
T6
Time
Genotype
TM1
TM2
WT

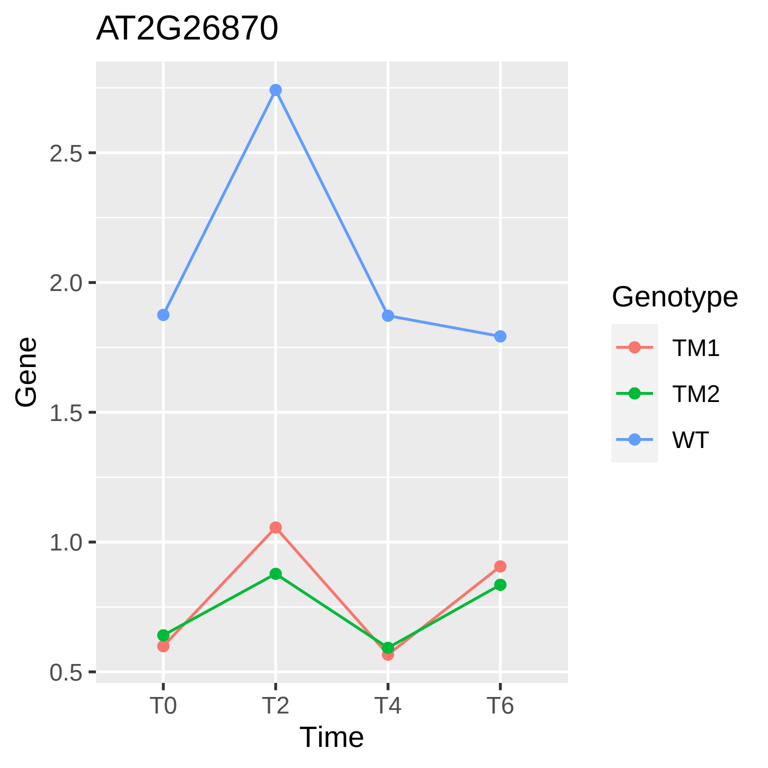

AT2G26870
Gene
2.5
2.0
1.5
1.0
0.5
T0
T2
T4
T6
Time
Genotype
TM1
TM2
WT

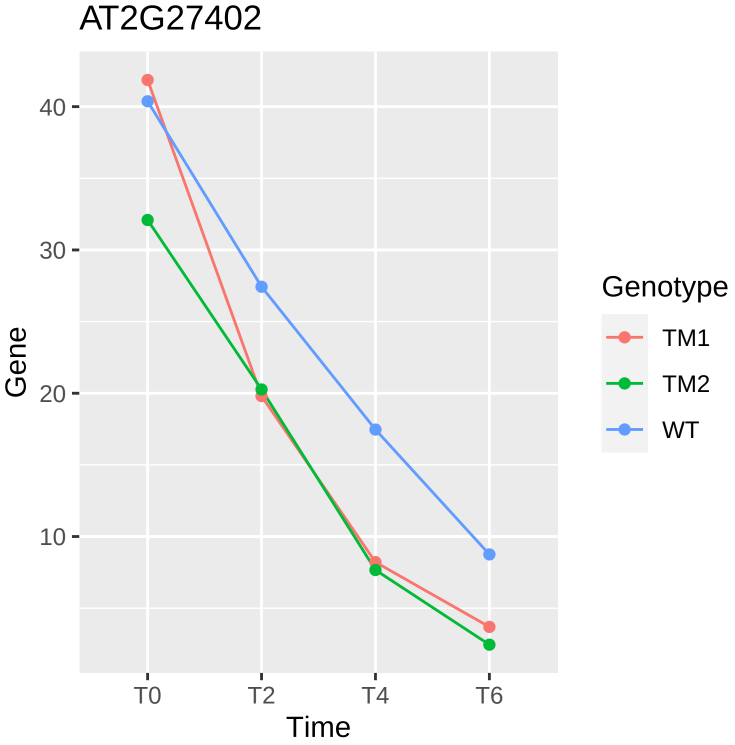

AT2G27402
Gene
40
30
20
10
T0
T2
T4
T6
Time
Genotype
TM1
TM2
WT

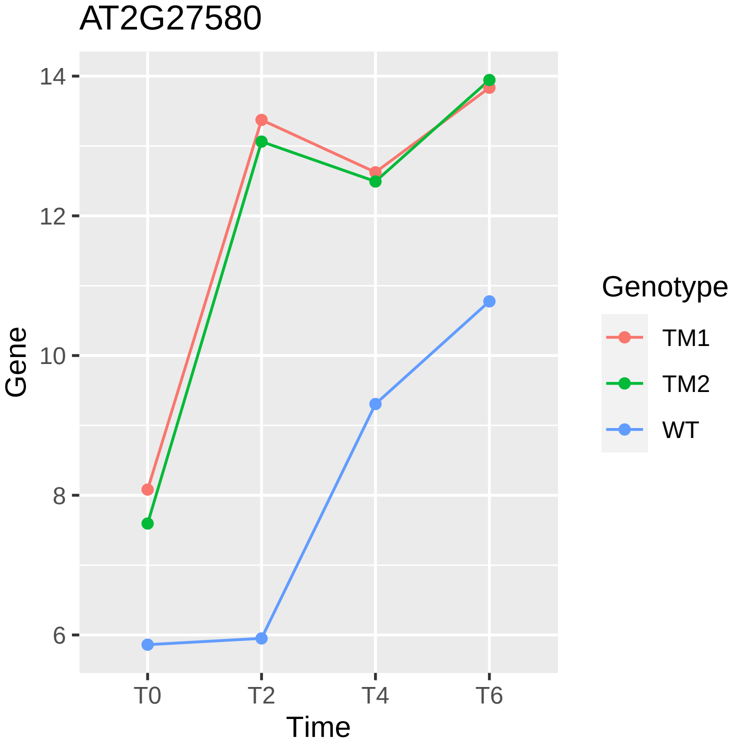
AT2G27580
Gene
14
12
10
8
6
T0
T2
T4
T6
Time
Genotype
TM1
TM2
WT

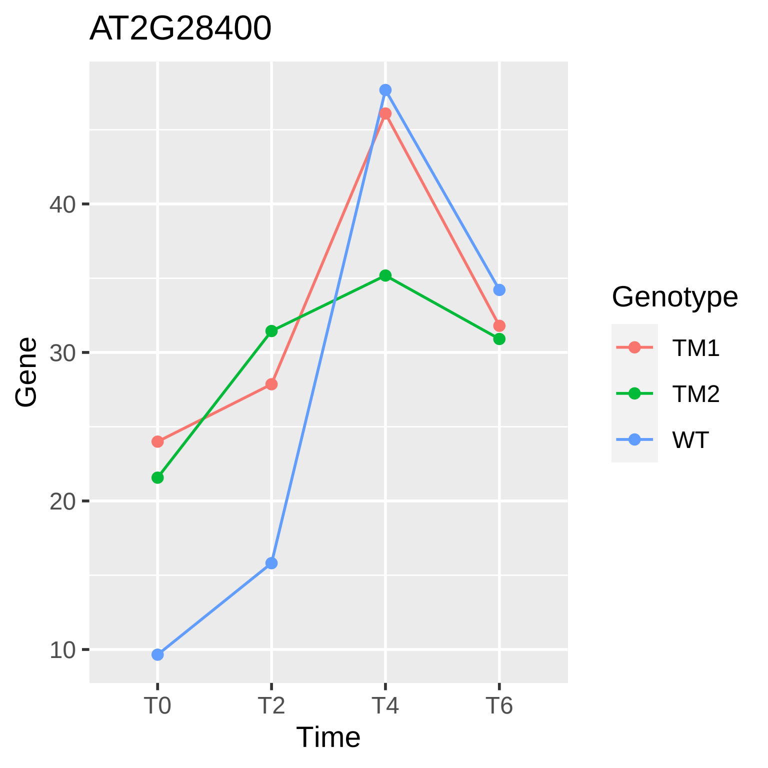

AT2G28400
Gene
40
30
20
10
T0
T2
T4
T6
Time
Genotype
TM1
TM2
WT

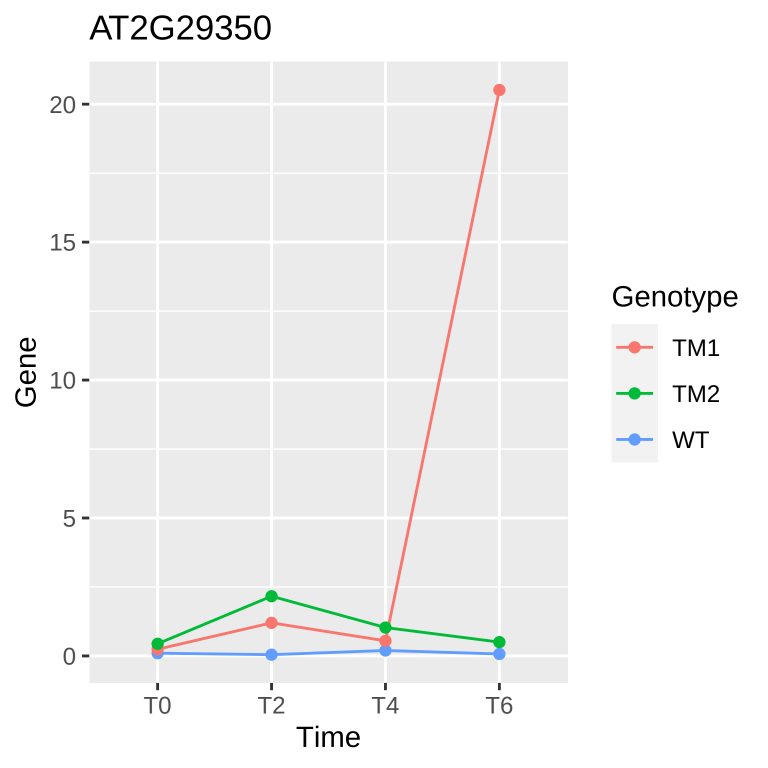

AT2G29350
Gene
20
15
10
5
0
T0
T2
T4
T6
Time
Genotype
TM1
TM2
WT

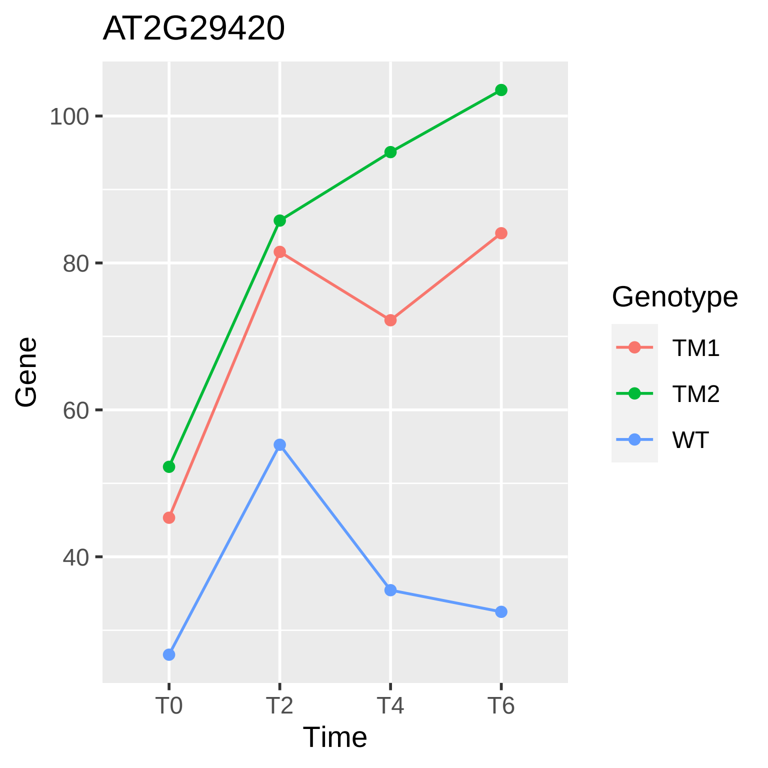
AT2G29420
Gene
100
80
60
40
T0
T2
T4
T6
Time
Genotype
TM1
TM2
WT

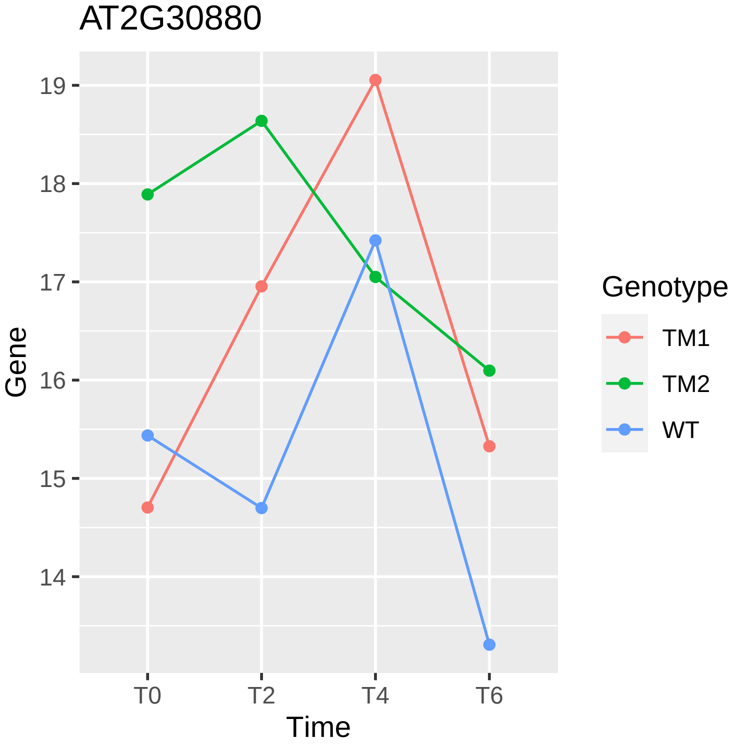

AT2G30880
Gene
19
18
17
16
15
14
T0
T2
T4
T6
Time
Genotype
TM1
TM2
WT

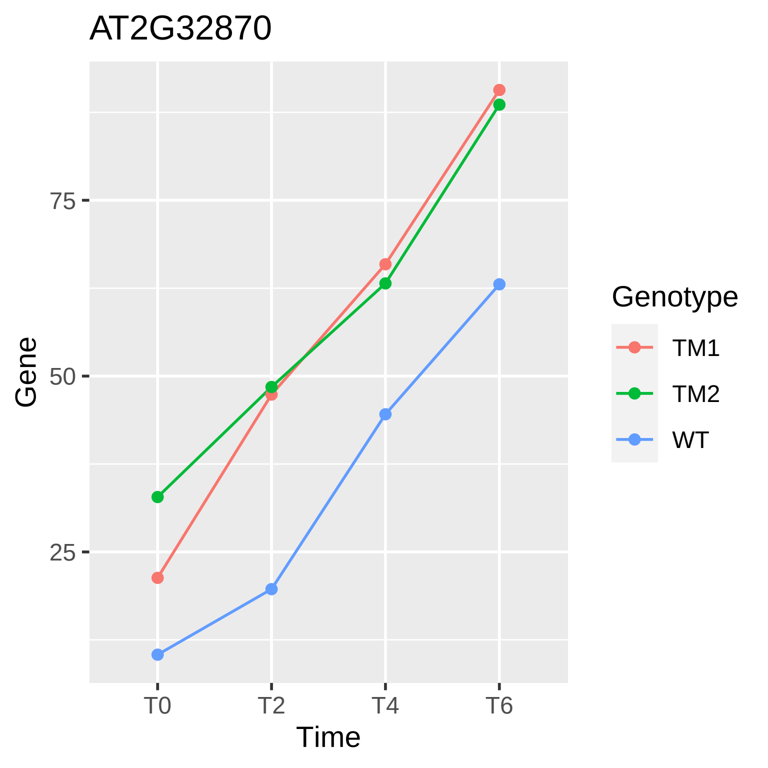
AT2G32870
Gene
75
50
25
T0
T2
T4
T6
Time
Genotype
TM1
TM2
WT

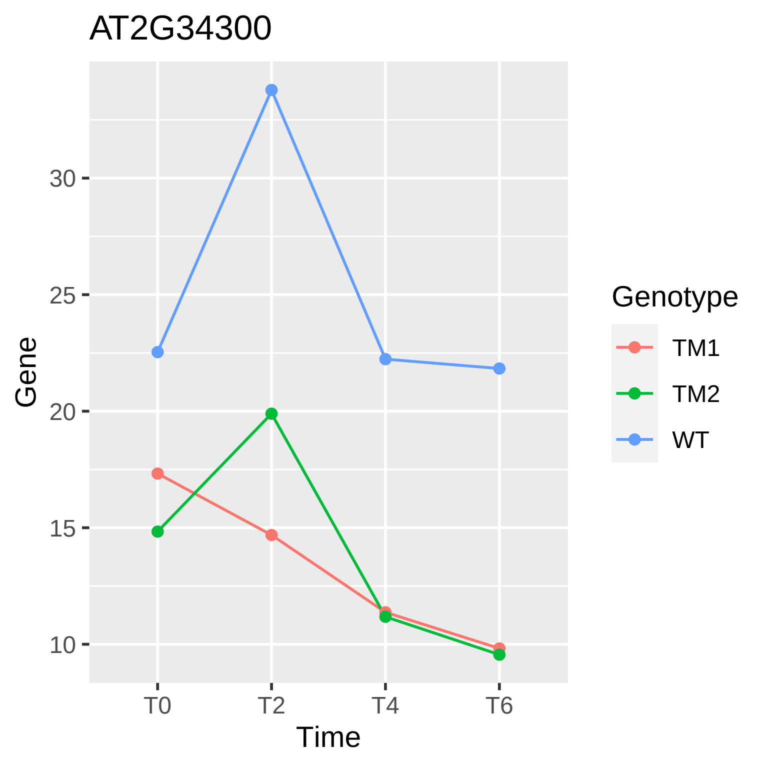
AT2G34300
Gene
30
25
20
15
10
T0
T2
T4
T6
Time
Genotype
TM1
TM2
WT

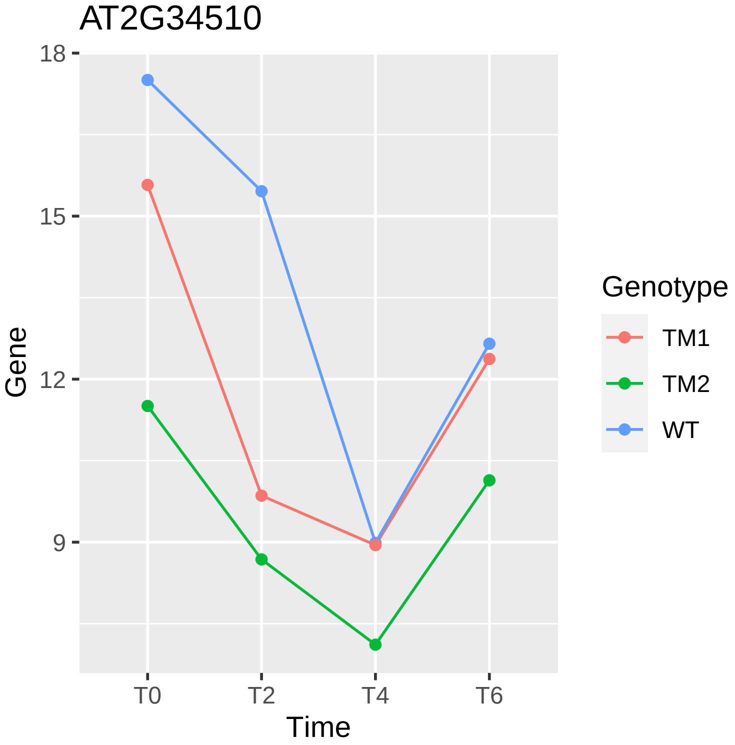

AT2G34510
18
15
12
9
Gene
T0
T2
T4
T6
Time
Genotype
TM1
TM2
WT

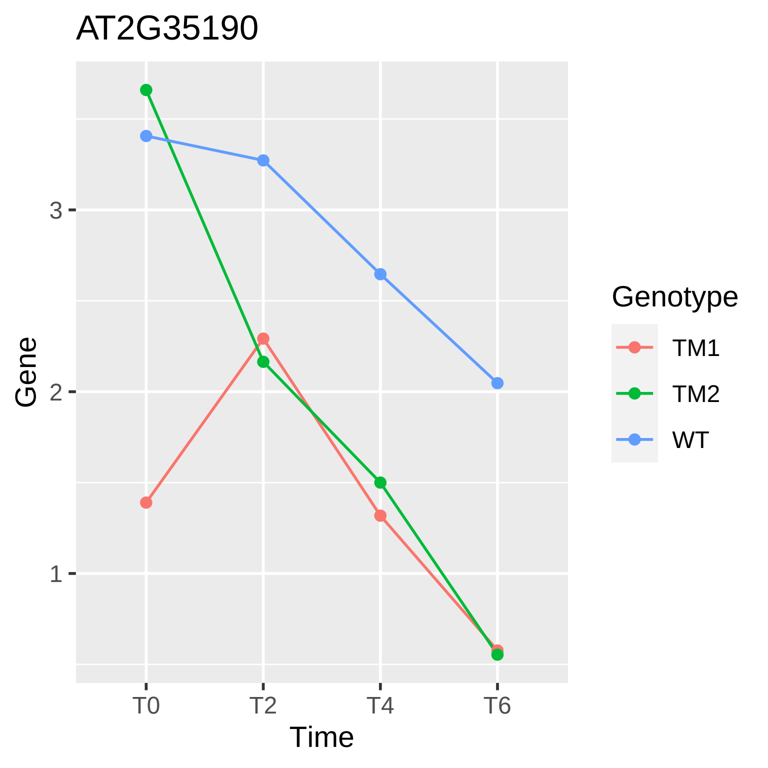
AT2G35190
Gene
Time
3
2
1
T0
T2
T4
T6
Genotype
TM1
TM2
WT

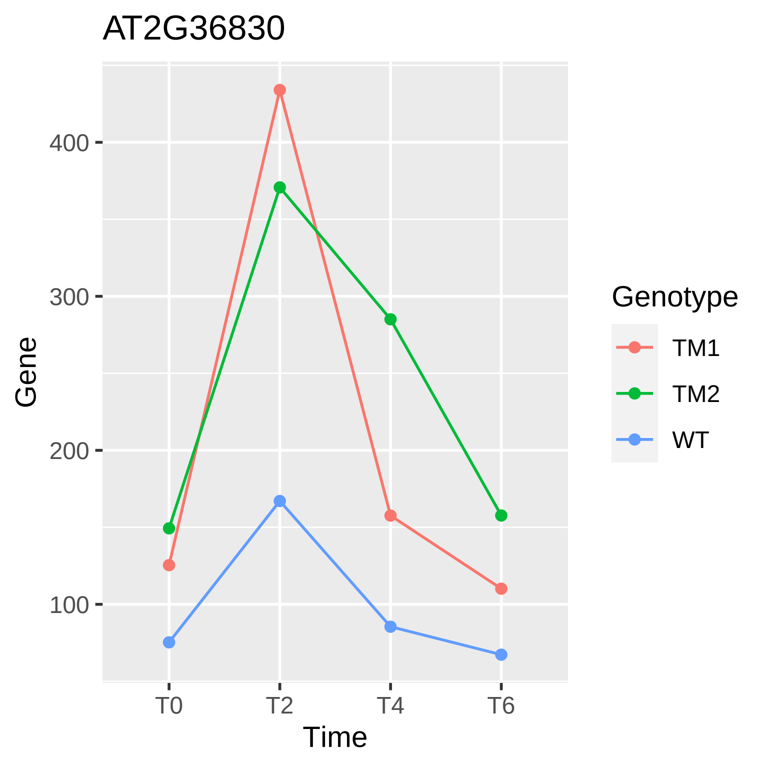
AT2G36830
Gene
400
300
200
100
T0
T2
T4
T6
Time
Genotype
TM1
TM2
WT

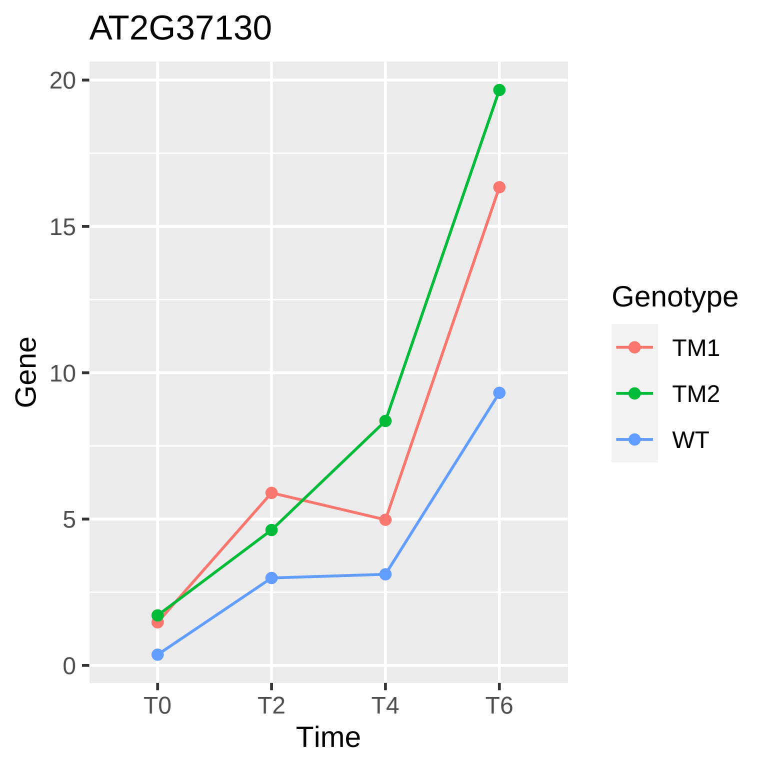

AT2G37130
Gene
0
5
10
15
20
T0
T2
T4
T6
Time
Genotype
TM1
TM2
WT

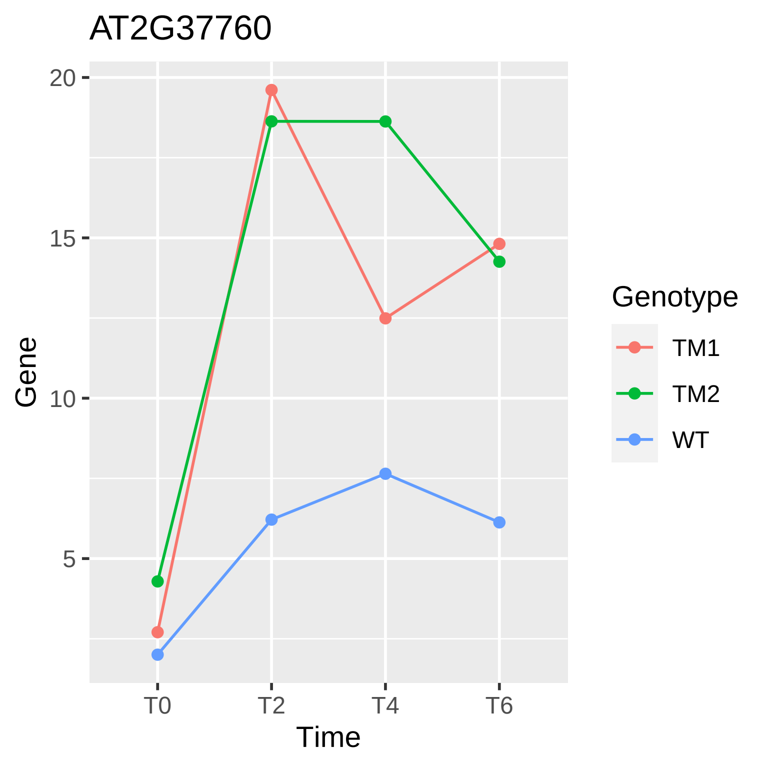

AT2G37760
20
15
10
5
Gene
T0
T2
T4
T6
Time
Genotype
TM1
TM2
WT

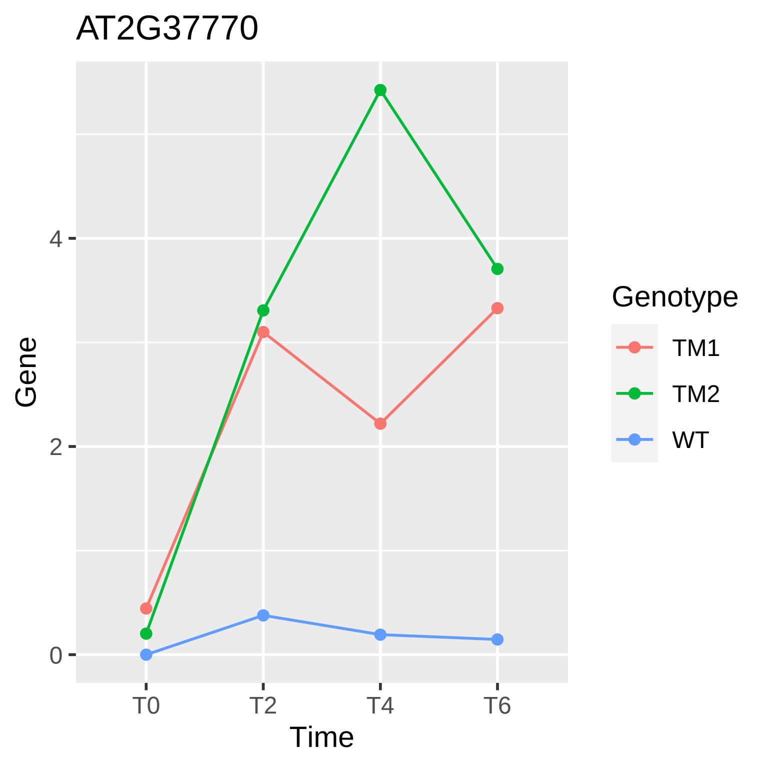
AT2G37770
Gene
4
2
0
T0
T2
T4
T6
Time
Genotype
TM1
TM2
WT

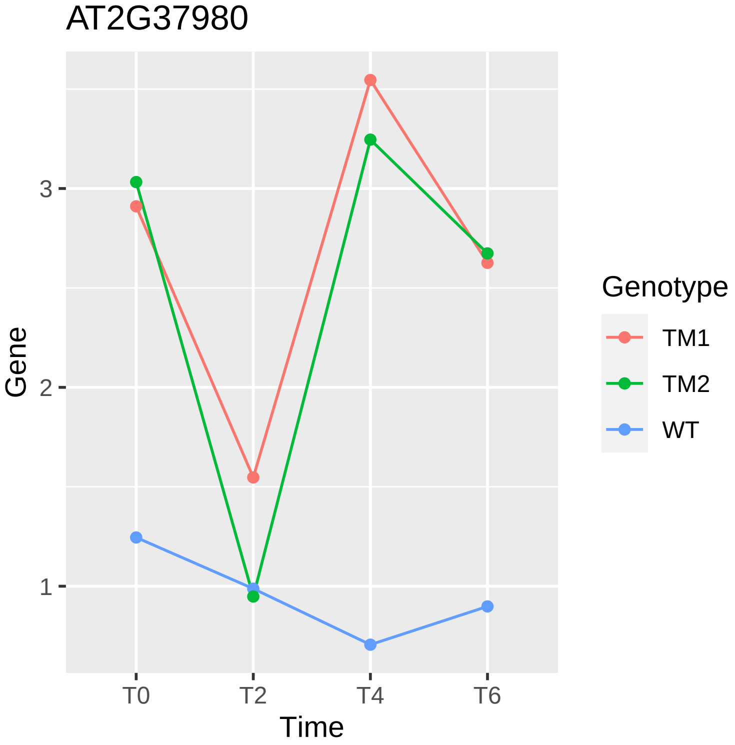
AT2G37980
Gene
3
2
1
T0
T2
T4
T6
Time
Genotype
TM1
TM2
WT

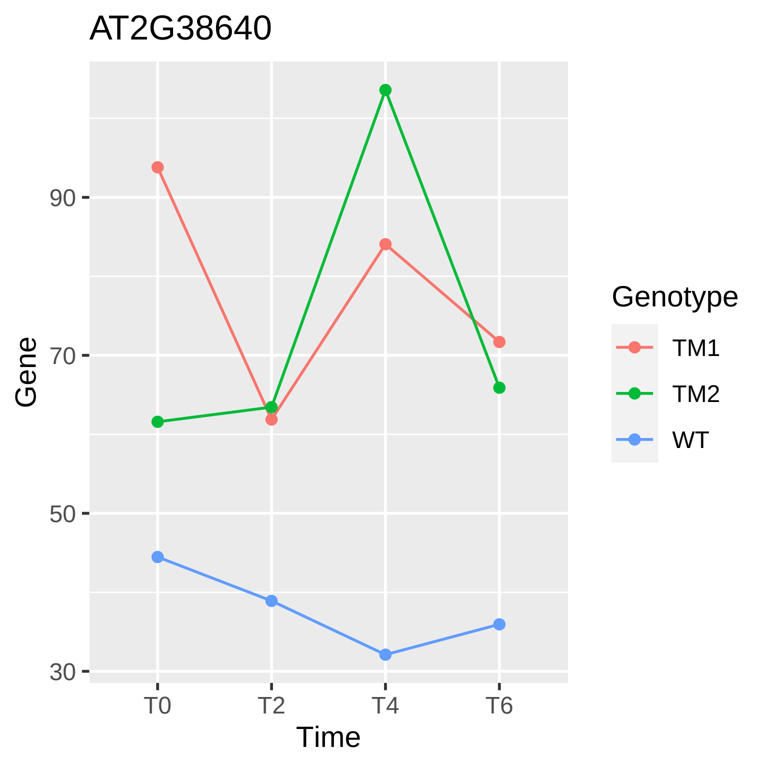

AT2G38640
Gene
90
70
50
30
T0
T2
T4
T6
Time
Genotype
TM1
TM2
WT

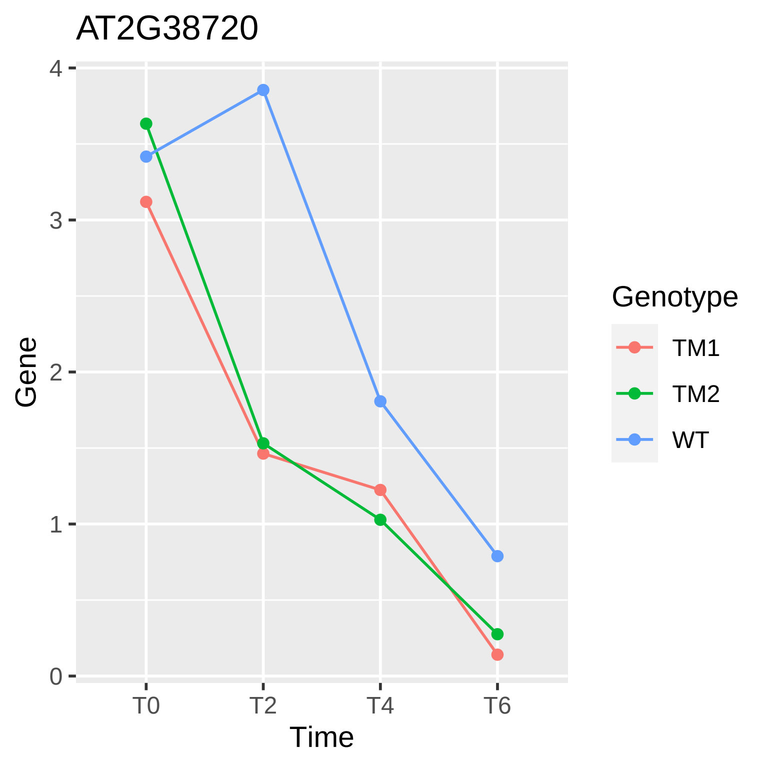
AT2G38720
Gene
0
1
2
3
4
T0
T2
T4
T6
Time
Genotype
TM1
TM2
WT

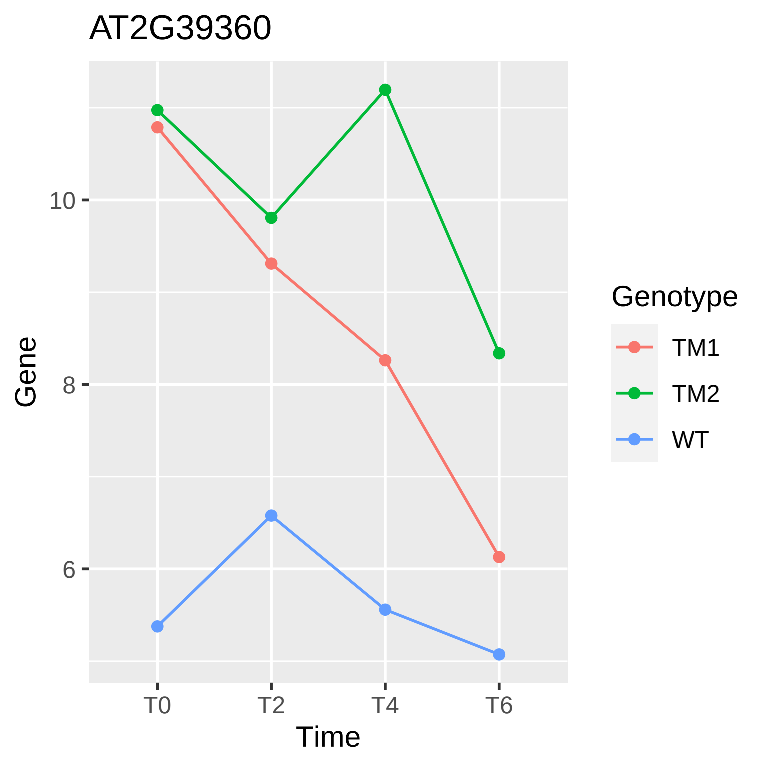
AT2G39360
Gene
10
8
6
T0
T2
T4
T6
Time
Genotype
TM1
TM2
WT

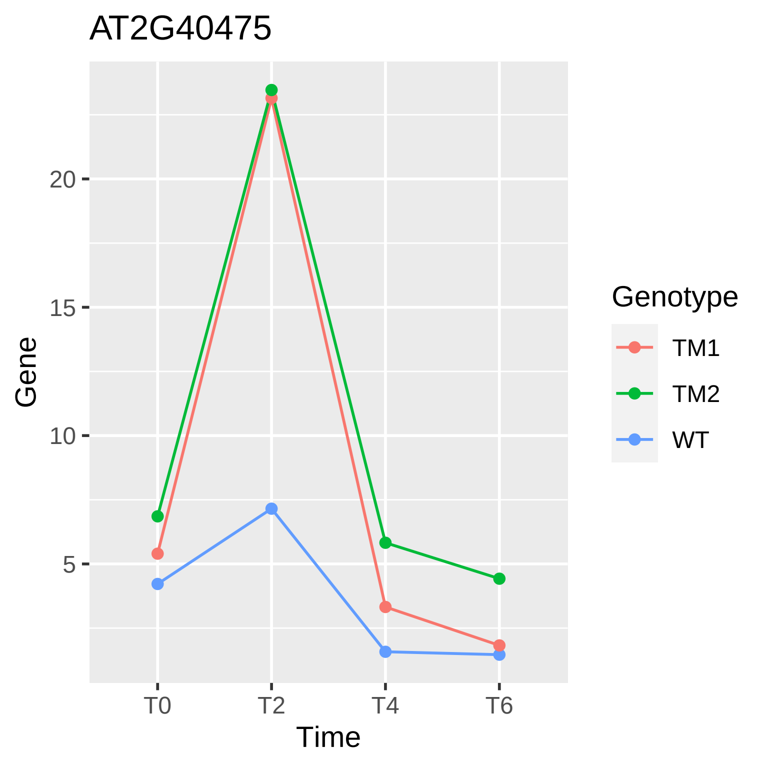

AT2G40475
Gene
20
15
10
5
T0
T2
T4
T6
Time
Genotype
TM1
TM2
WT

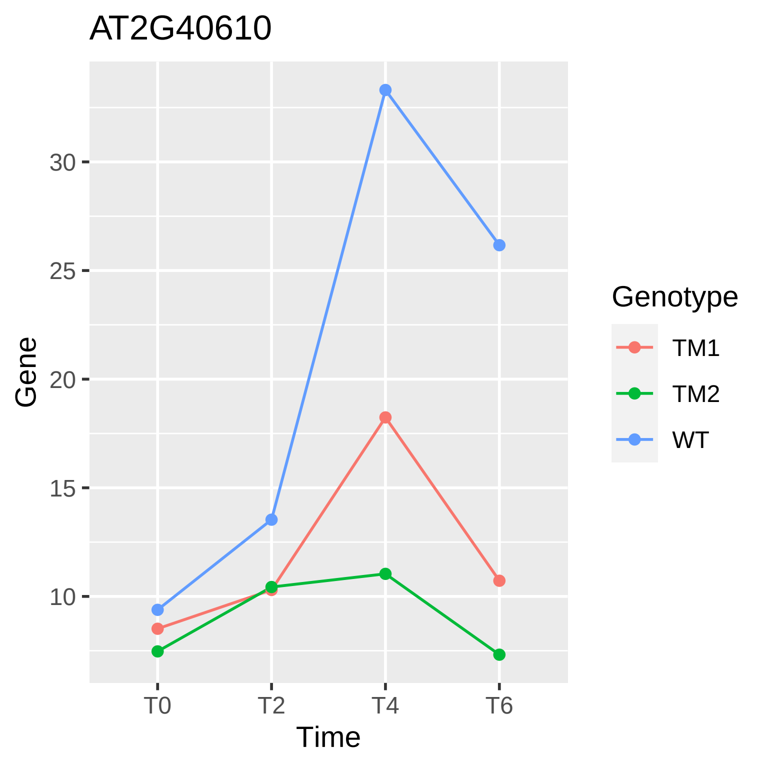

AT2G40610
Gene
30
25
20
15
10
T0
T2
T4
T6
Time
Genotype
TM1
TM2
WT

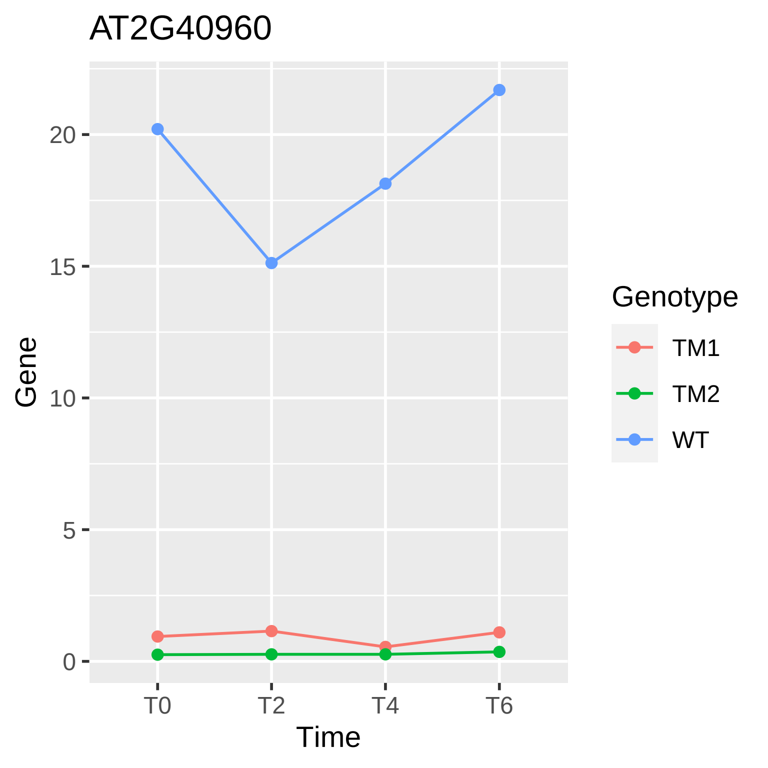

AT2G40960
Gene
20
15
10
5
0
T0
T2
T4
T6
Time
Genotype
TM1
TM2
WT

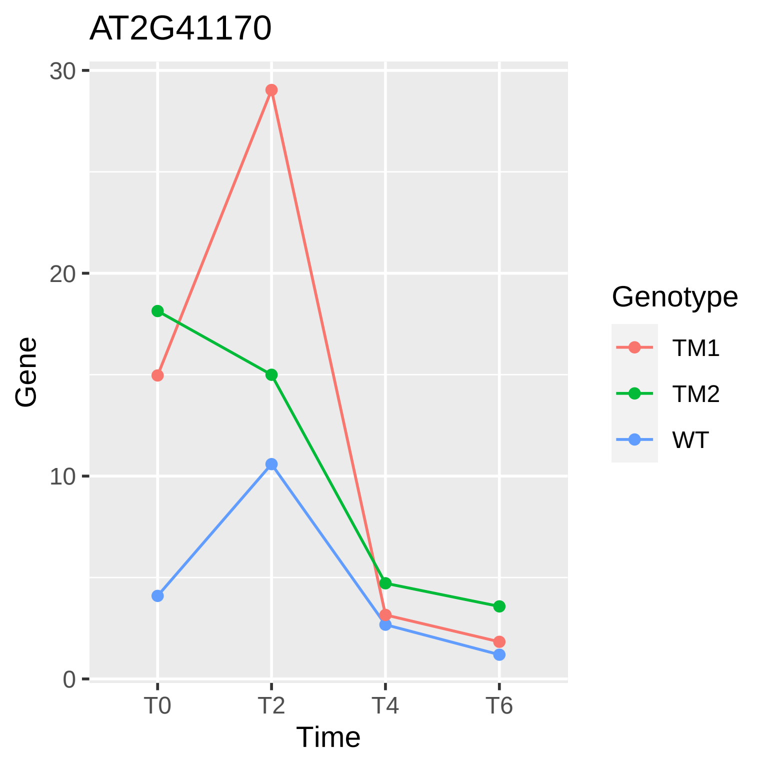

AT2G41170
30
20
10
0
Gene
T0
T2
T4
T6
Time
Genotype
TM1
TM2
WT

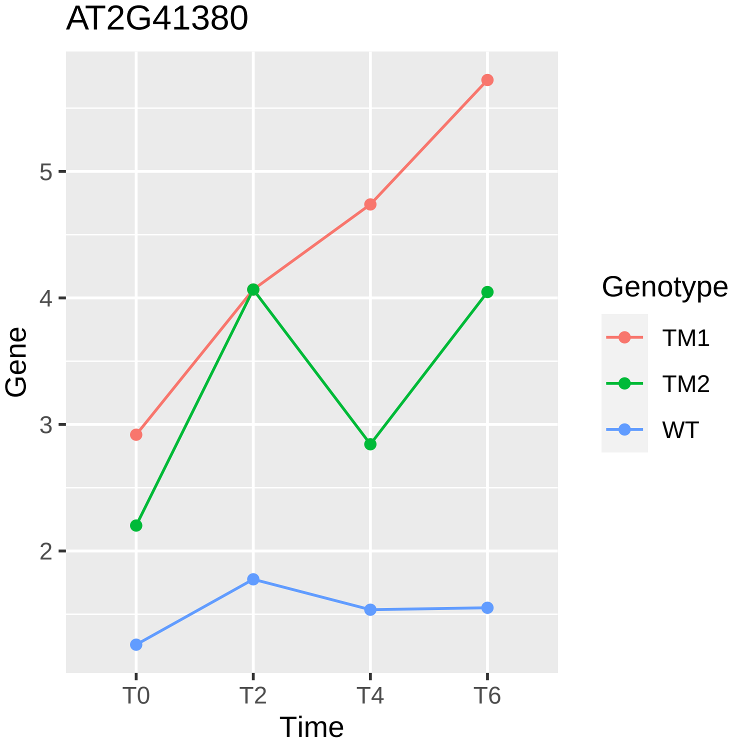
AT2G41380
Gene
5
4
3
2
T0
T2
T4
T6
Time
Genotype
TM1
TM2
WT

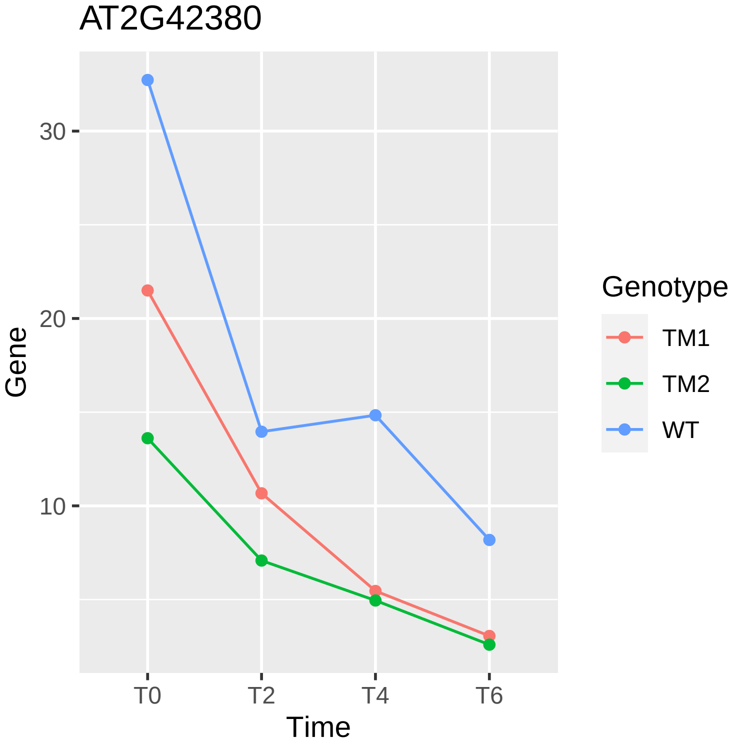

AT2G42380
Gene
30
20
10
T0
T2
T4
T6
Time
Genotype
TM1
TM2
WT

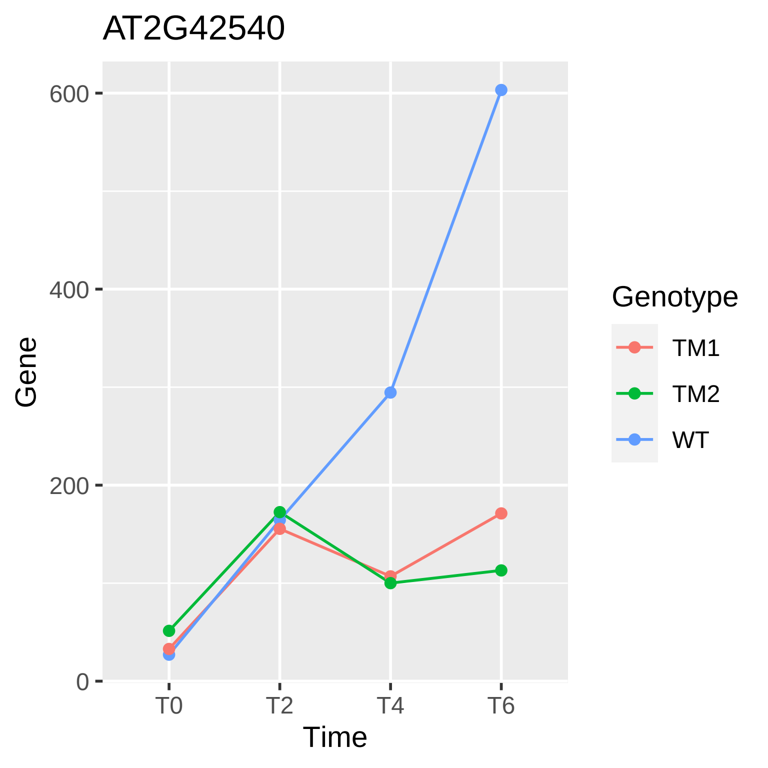
AT2G42540
600
400
200
0
Gene
T0
T2
T4
T6
Time
Genotype
TM1
TM2
WT

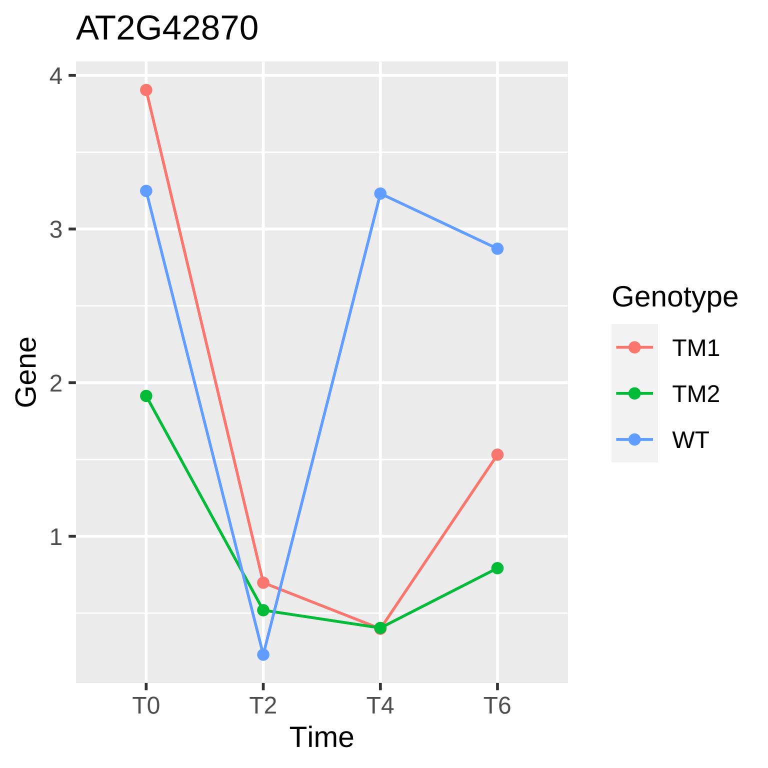
AT2G42870
Gene
4
3
2
1
T0
T2
T4
T6
Time
Genotype
TM1
TM2
WT

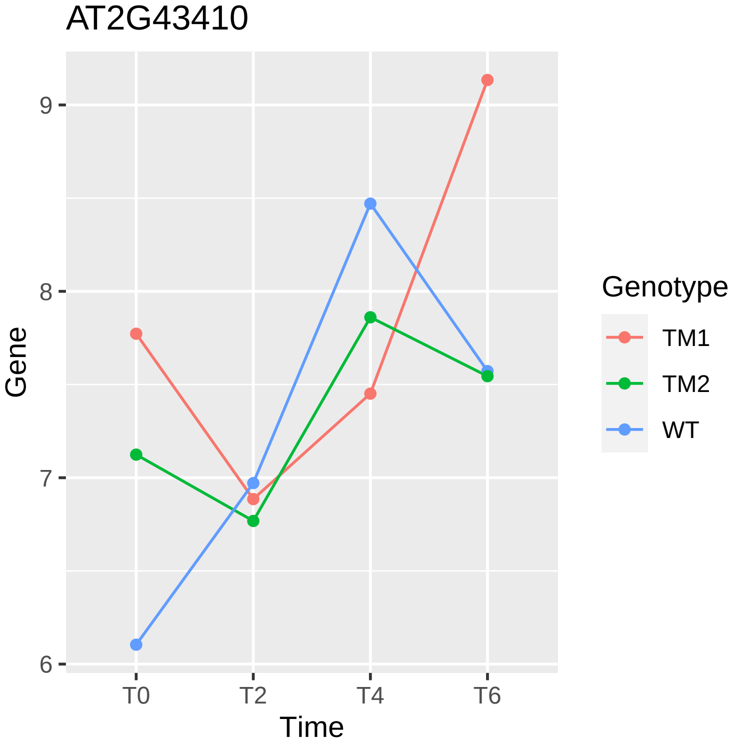

AT2G43410
Gene
9
8
7
6
T0
T2
T4
T6
Time
Genotype
TM1
TM2
WT

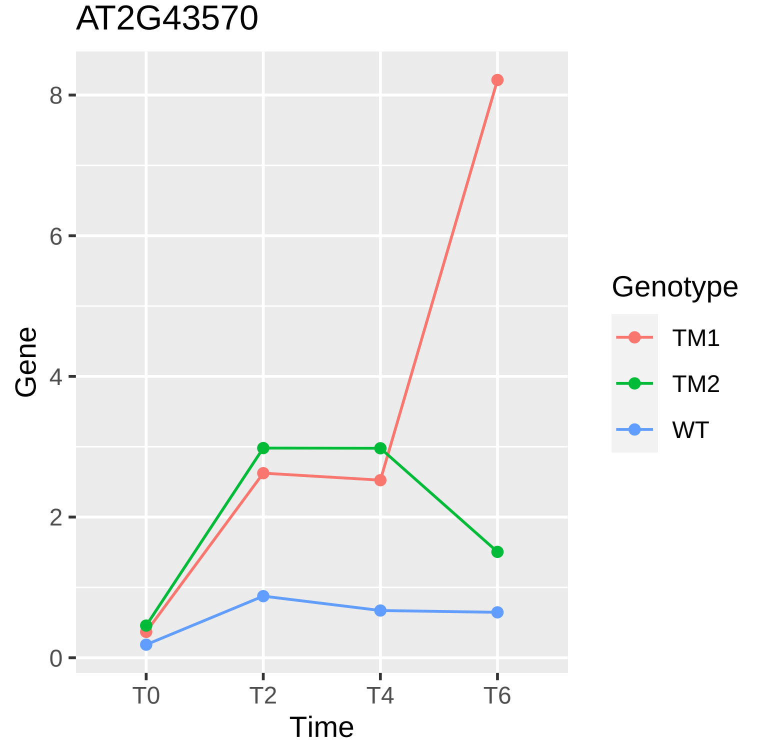
AT2G43570
Gene
Time
0
2
4
6
8
T0
T2
T4
T6
Genotype
TM1
TM2
WT

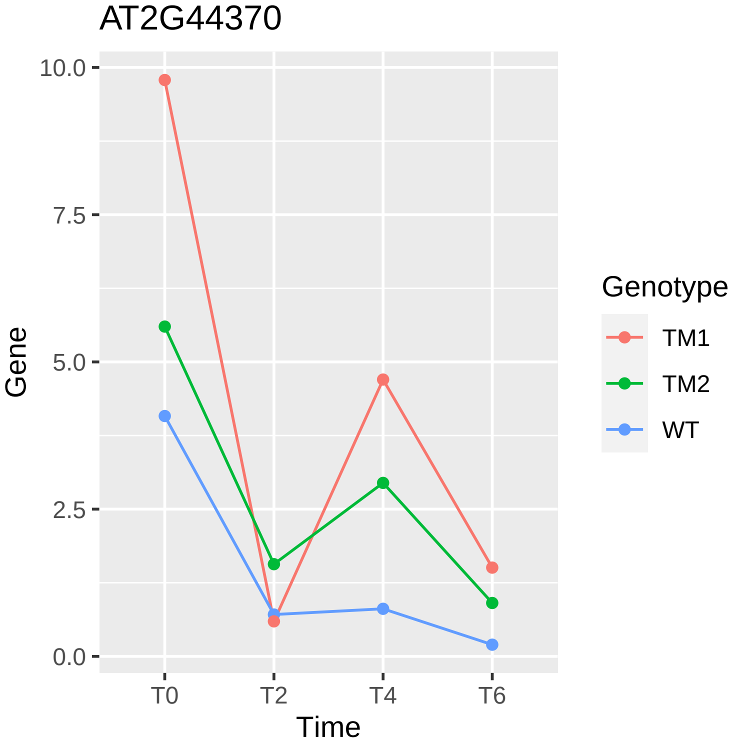
AT2G44370
Gene
10.0
7.5
5.0
2.5
0.0
T0
T2
T4
T6
Time
Genotype
TM1
TM2
WT

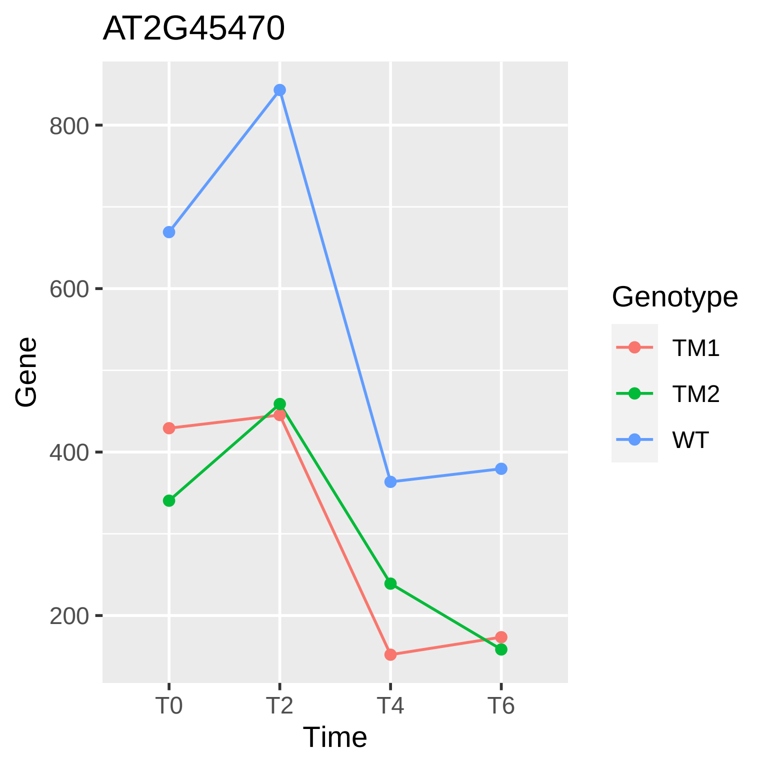

AT2G45470
800
600
400
200
Gene
T0
T2
T4
T6
Time
Genotype
TM1
TM2
WT

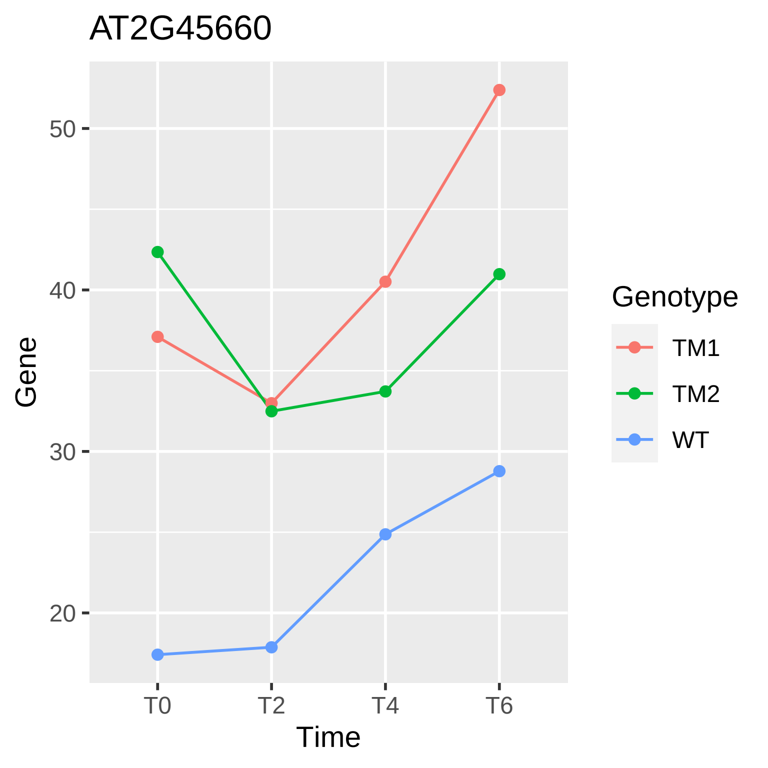

AT2G45660
Gene
50
40
30
20
T0
T2
T4
T6
Time
Genotype
TM1
TM2
WT

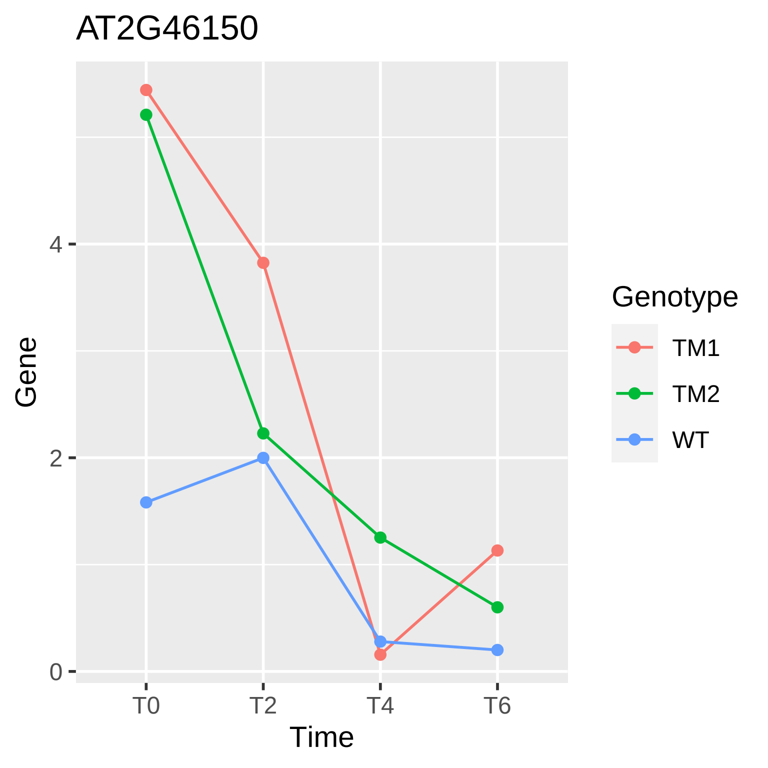

AT2G46150
Gene
0
2
4
T0
T2
T4
T6
Time
Genotype
TM1
TM2
WT

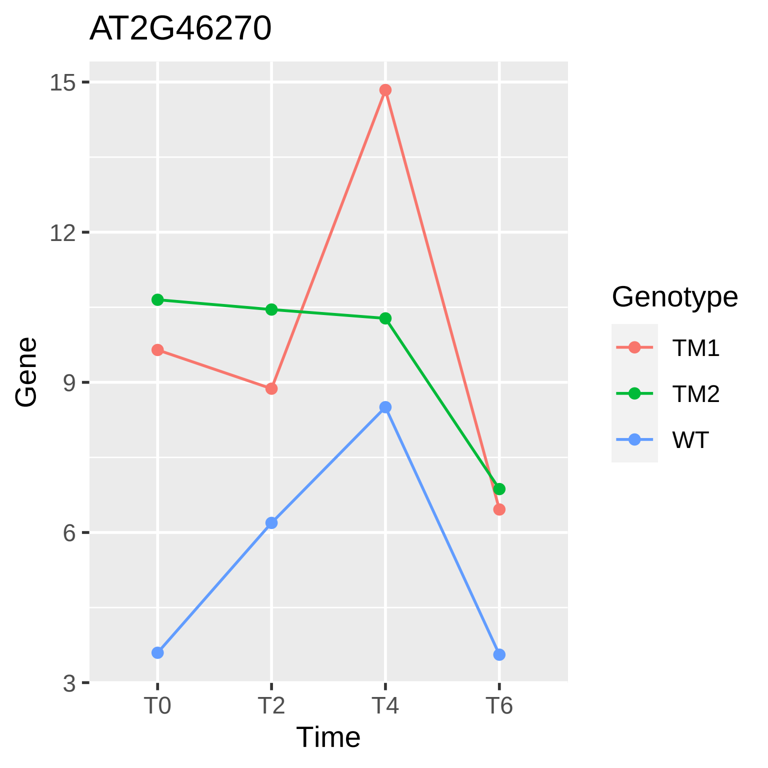

AT2G46270
15
12
9
6
3
Gene
T0
T2
T4
T6
Time
Genotype
TM1
TM2
WT

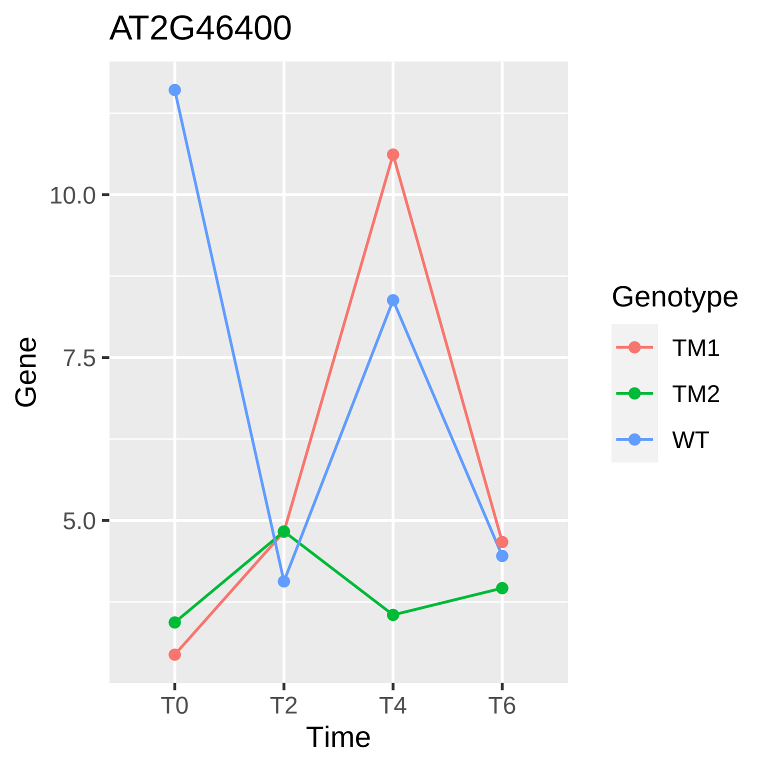

AT2G46400
Gene
10.0
7.5
5.0
T0
T2
T4
T6
Time
Genotype
TM1
TM2
WT

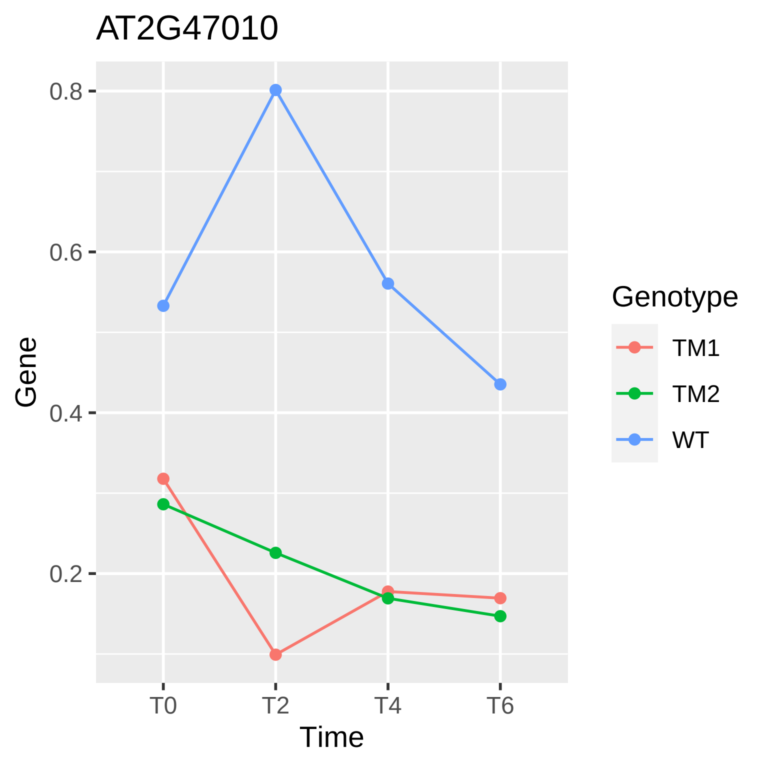
AT2G47010
0.8
0.6
0.4
0.2
Gene
T0
T2
T4
T6
Time
Genotype
TM1
TM2
WT

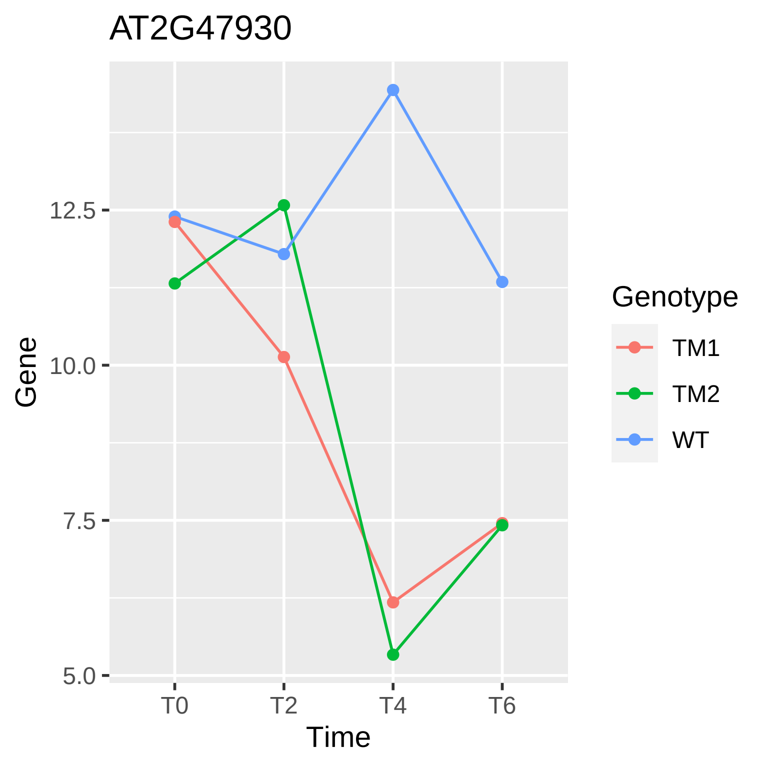
AT2G47930
Gene
12.5
10.0
7.5
5.0
T0
T2
T4
T6
Time
Genotype
TM1
TM2
WT

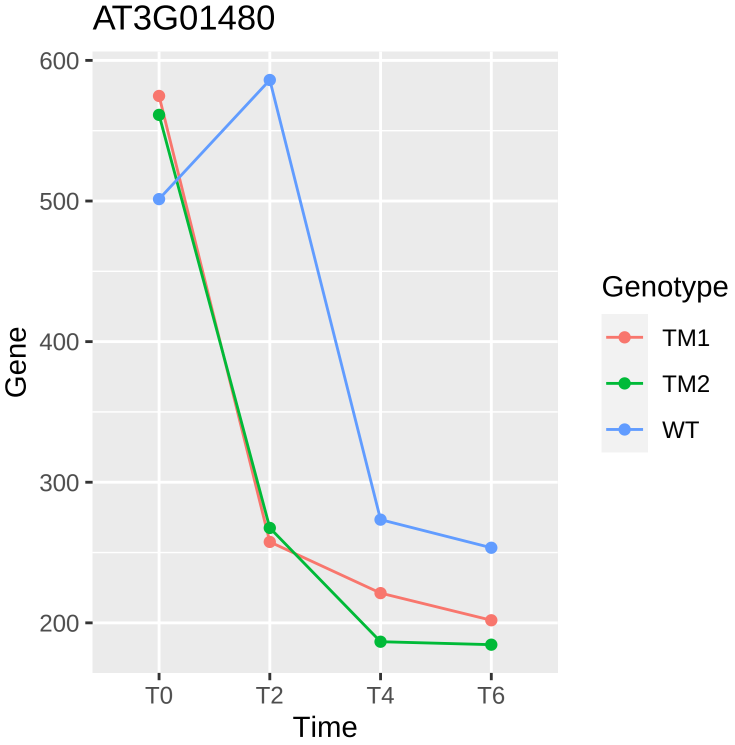
AT3G01480
600
500
400
300
200
Gene
T0
T2
T4
T6
Time
Genotype
TM1
TM2
WT

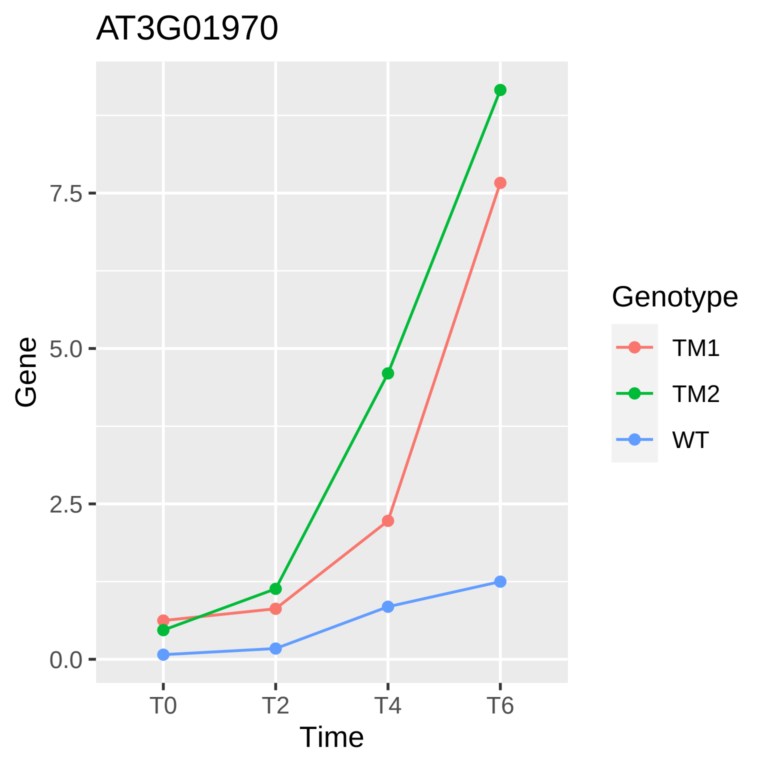
AT3G01970
Gene
7.5
5.0
2.5
0.0
T0
T2
T4
T6
Time
Genotype
TM1
TM2
WT

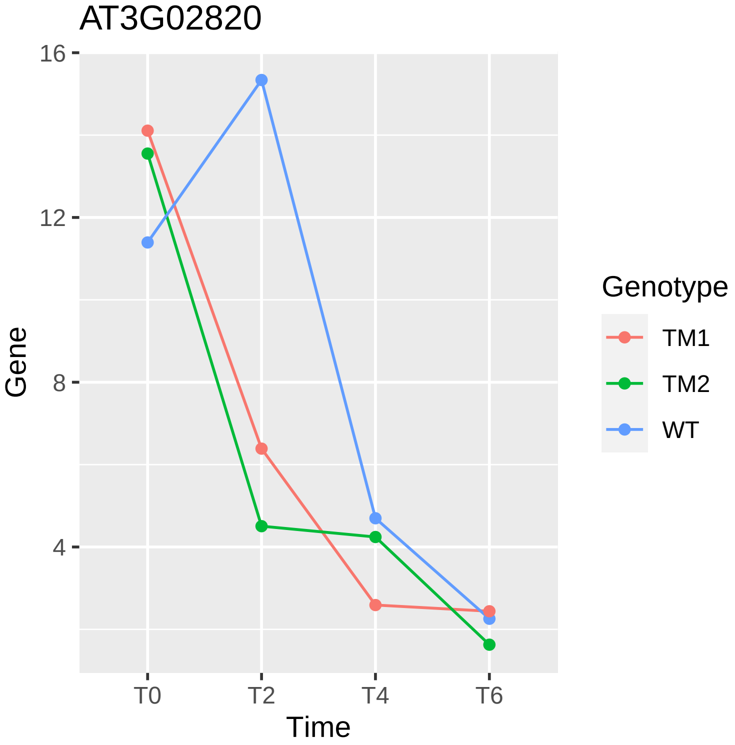
AT3G02820
16
12
8
4
Gene
T0
T2
T4
T6
Time
Genotype
TM1
TM2
WT

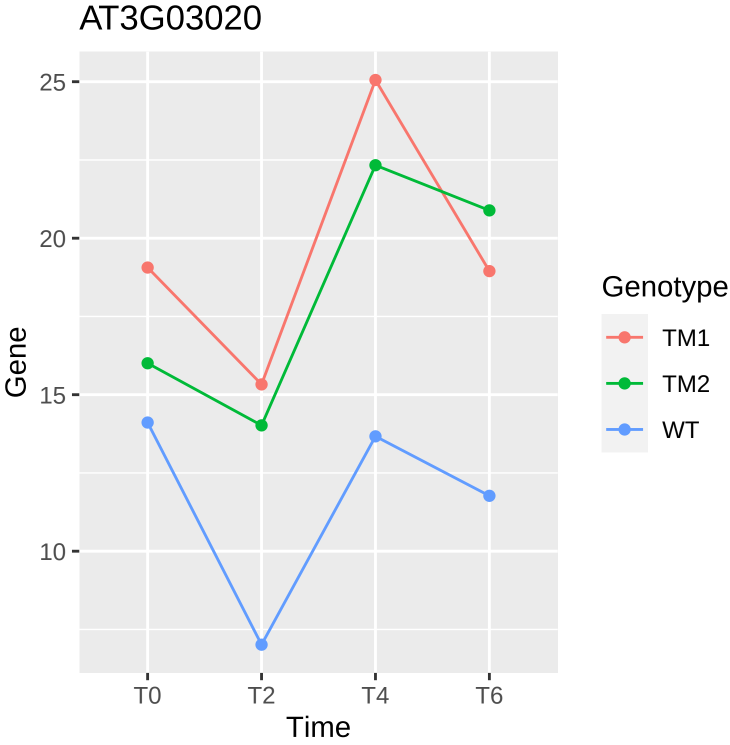
AT3G03020
25
20
15
10
Gene
T0
T2
T4
T6
Time
Genotype
TM1
TM2
WT

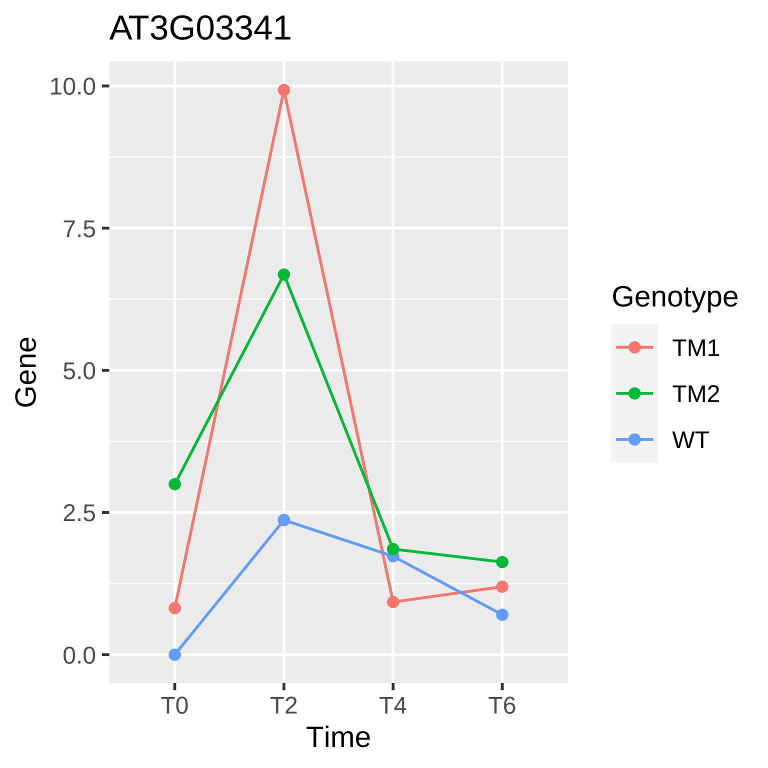
AT3G03341
10.0
7.5
5.0
2.5
0.0
Gene
T0
T2
T4
T6
Time
Genotype
TM1
TM2
WT

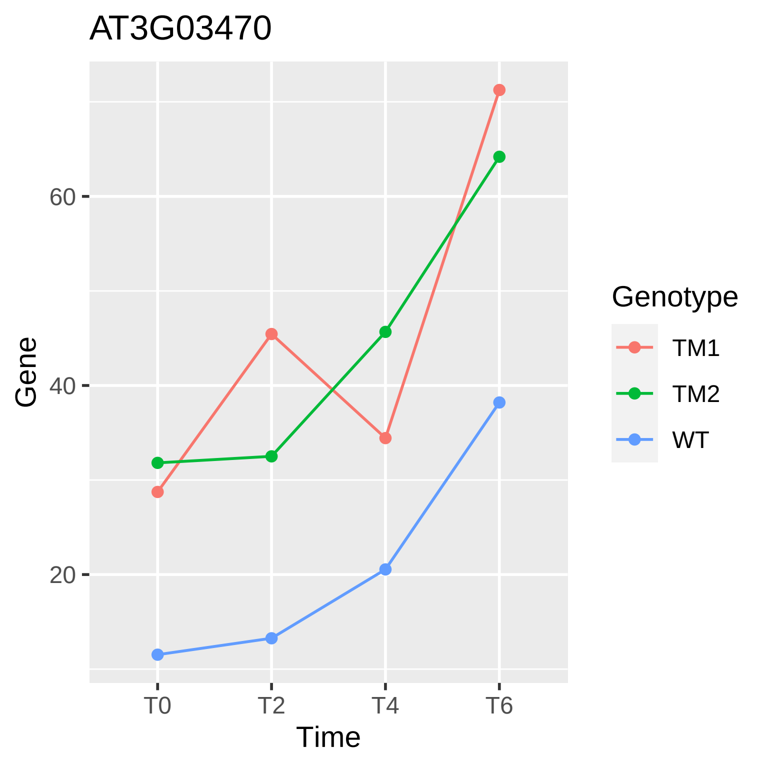
AT3G03470
Gene
60
40
20
T0
T2
T4
T6
Time
Genotype
TM1
TM2
WT

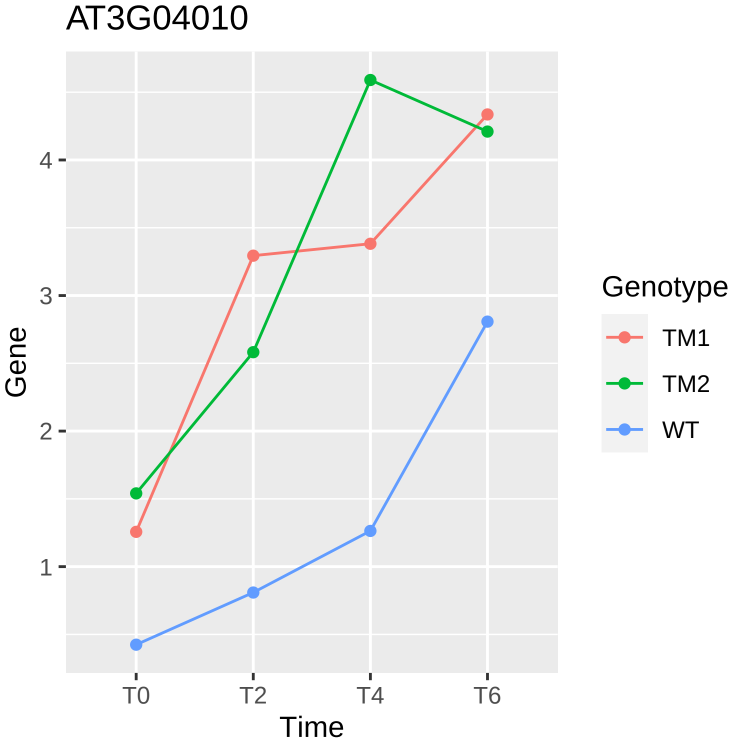

AT3G04010
Gene
4
3
2
1
T0
T2
T4
T6
Time
Genotype
TM1
TM2
WT

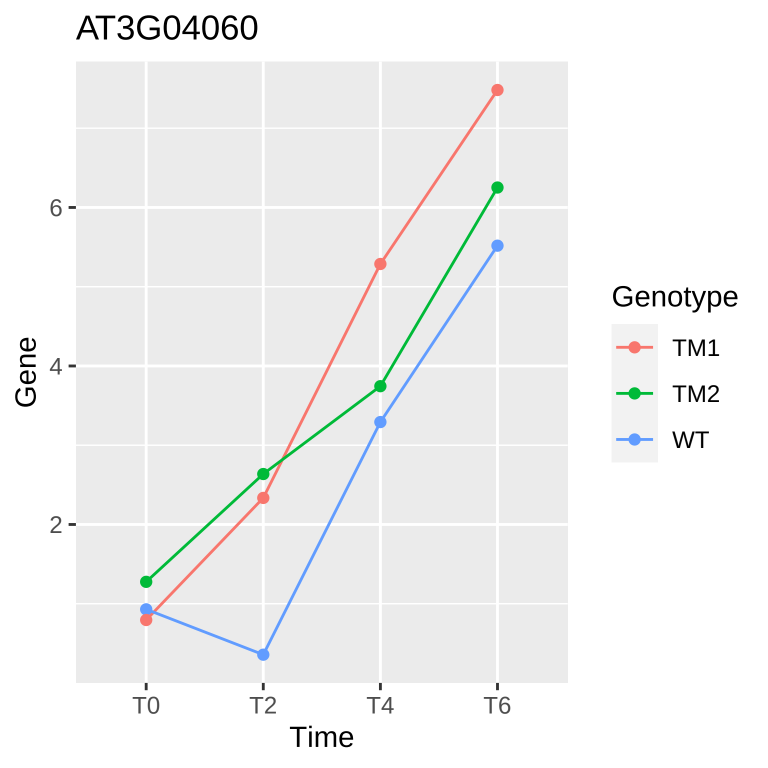
AT3G04060
Gene
2
4
6
T0
T2
T4
T6
Time
Genotype
TM1
TM2
WT

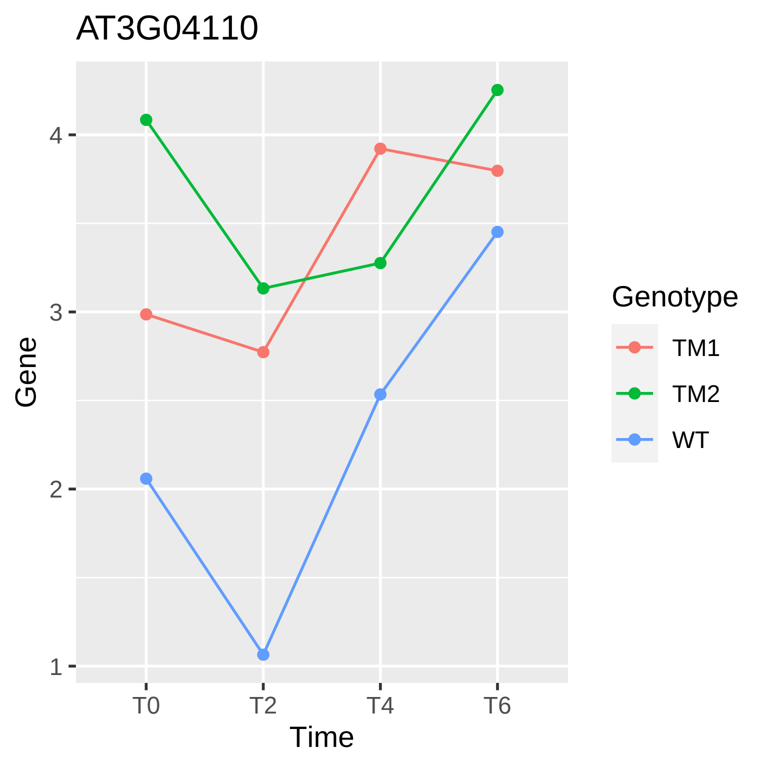
AT3G04110
Gene
1
2
3
4
T0
T2
T4
T6
Time
Genotype
TM1
TM2
WT

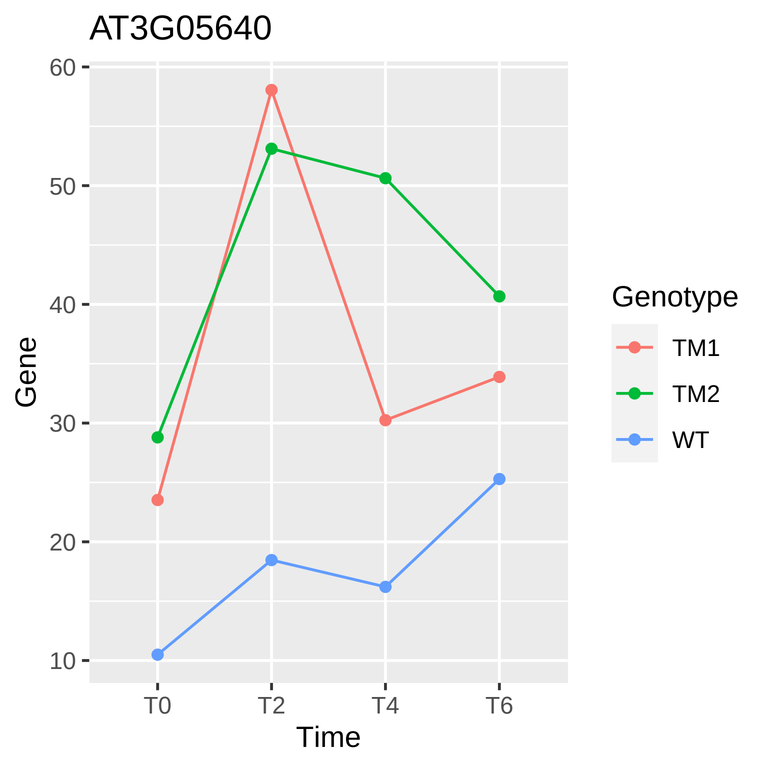
AT3G05640
Gene
60
50
40
30
20
10
T0
T2
T4
T6
Time
Genotype
TM1
TM2
WT

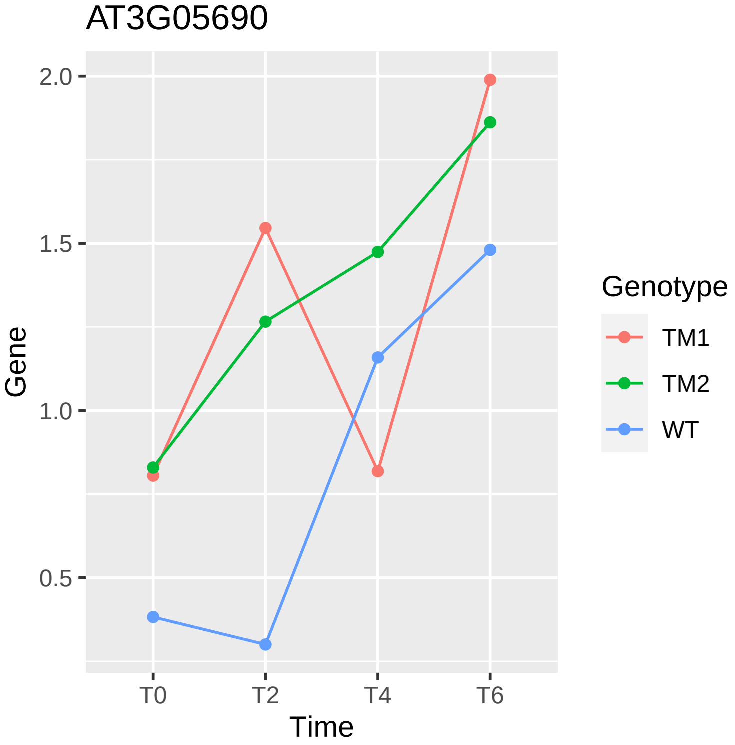
AT3G05690
Gene
2.0
1.5
1.0
0.5
T0
T2
T4
T6
Time
Genotype
TM1
TM2
WT

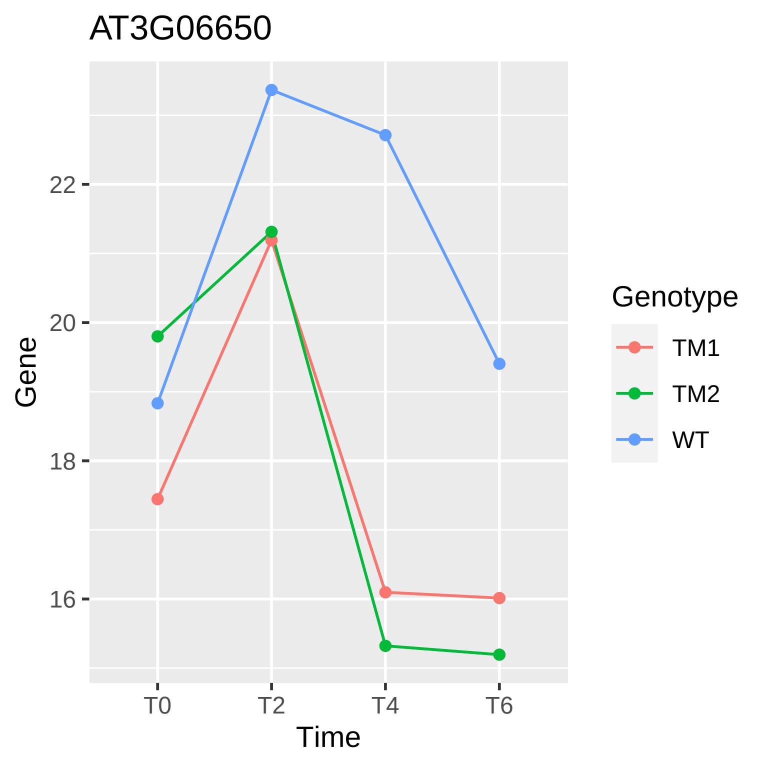
AT3G06650
Gene
22
20
18
16
T0
T2
T4
T6
Time
Genotype
TM1
TM2
WT

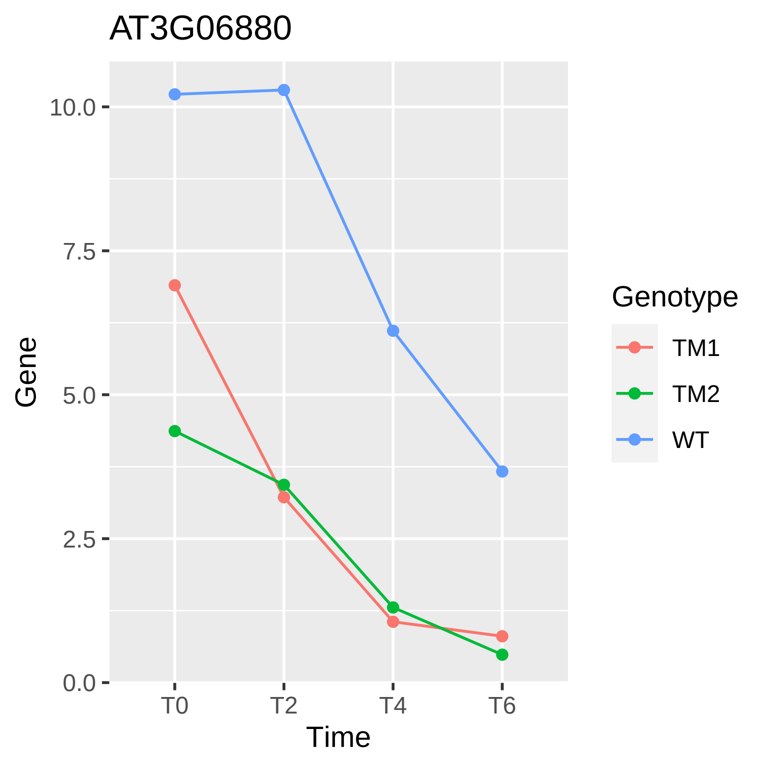

AT3G06880
Gene
10.0
7.5
5.0
2.5
0.0
T0
T2
T4
T6
Time
Genotype
TM1
TM2
WT

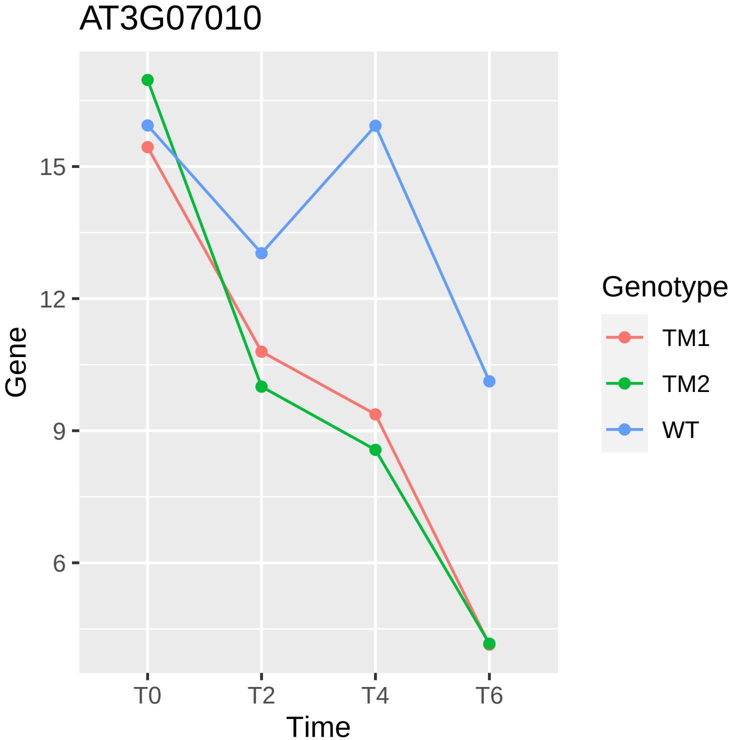
AT3G07010
Gene
15
12
9
6
T0
T2
T4
T6
Time
Genotype
TM1
TM2
WT

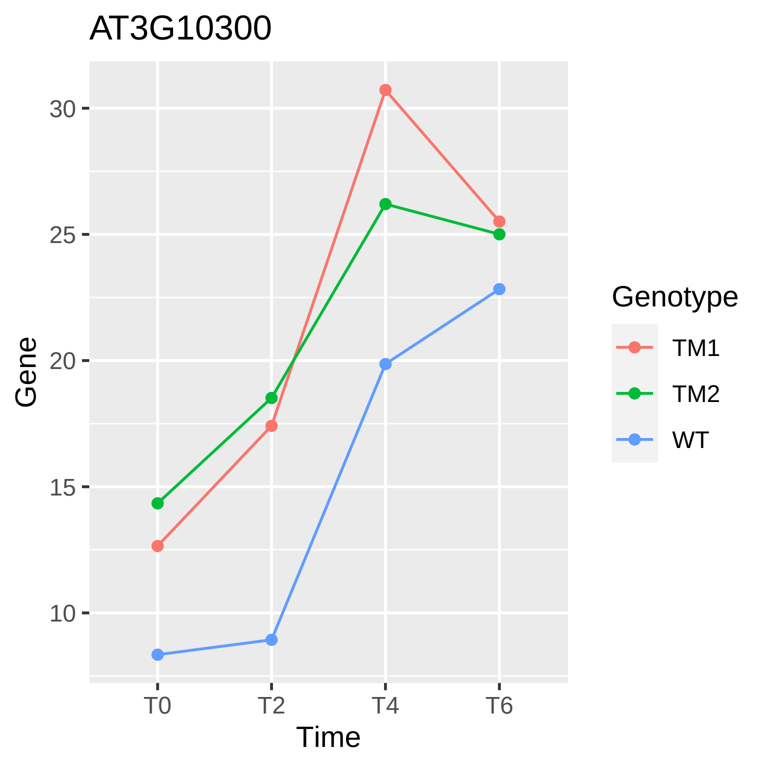

AT3G10300
Gene
Time
30
25
20
15
10
T0
T2
T4
T6
Genotype
TM1
TM2
WT

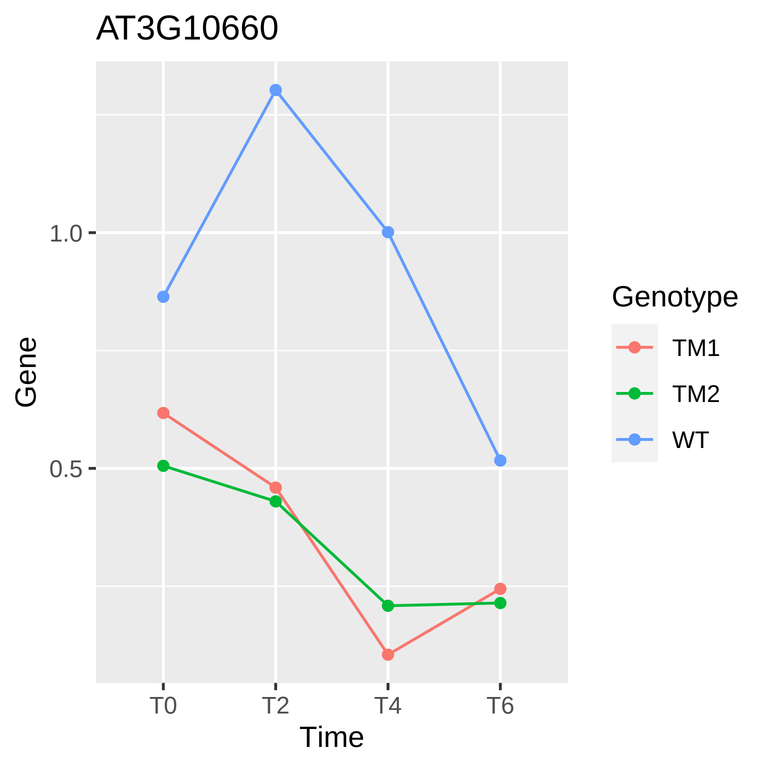

AT3G10660
1.0
0.5
Gene
T0
T2
T4
T6
Time
Genotype
TM1
TM2
WT

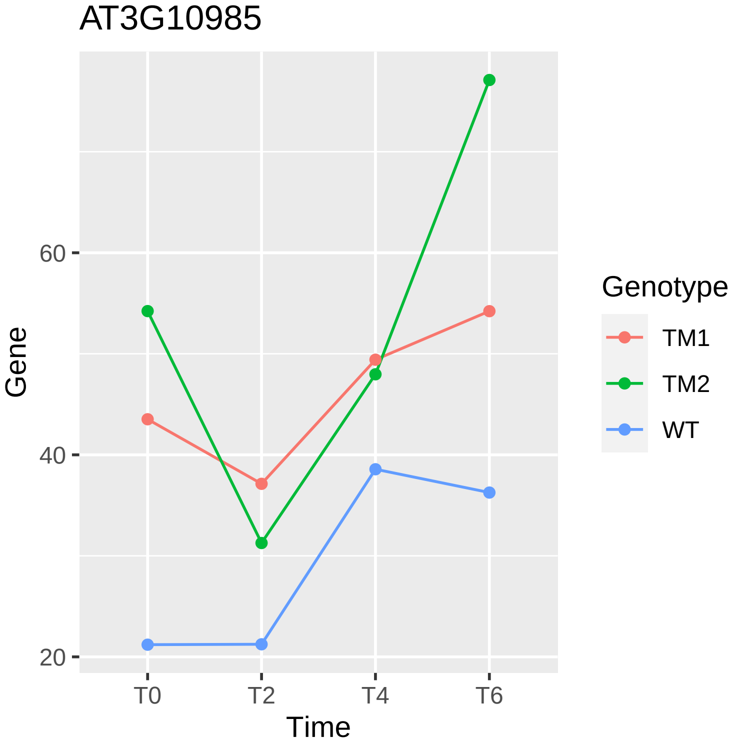
AT3G10985
Gene
60
40
20
T0
T2
T4
T6
Time
Genotype
TM1
TM2
WT

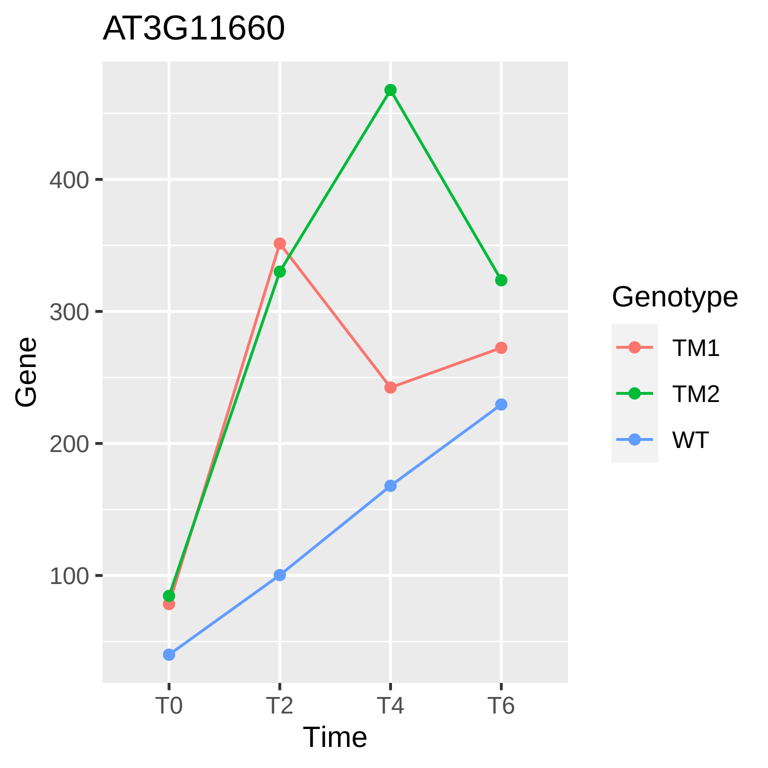

AT3G11660
Gene
400
300
200
100
T0
T2
T4
T6
Time
Genotype
TM1
TM2
WT

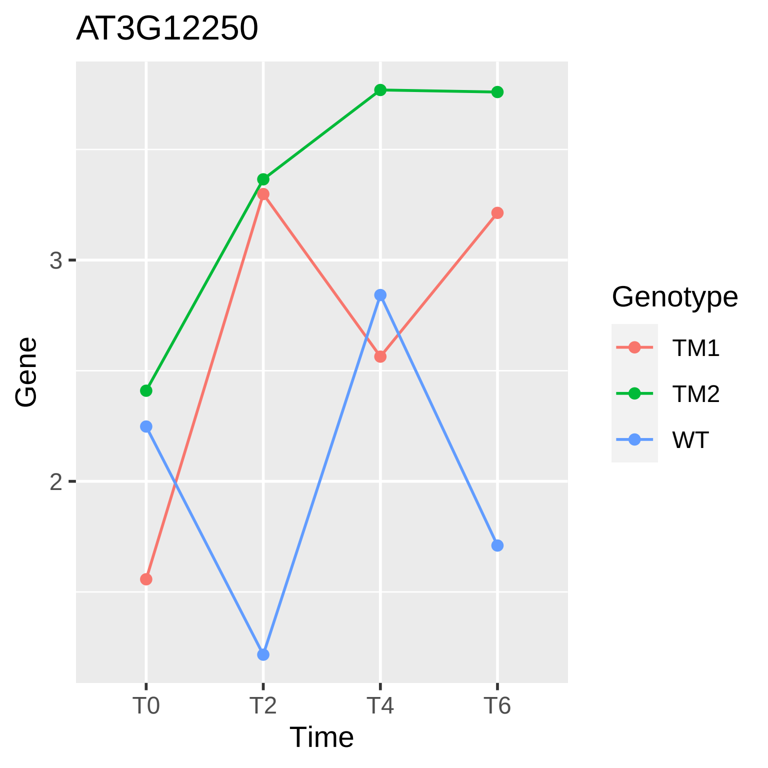

AT3G12250
Gene
3
2
T0
T2
T4
T6
Time
Genotype
TM1
TM2
WT

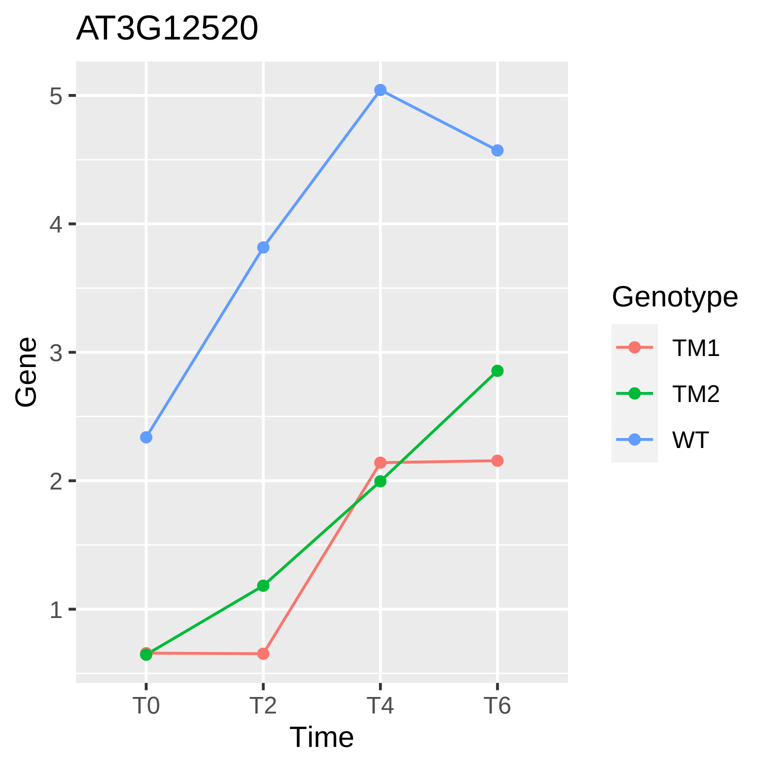

AT3G12520
Gene
Time
T0
T2
T4
T6
1
2
3
4
5
Genotype
TM1
TM2
WT

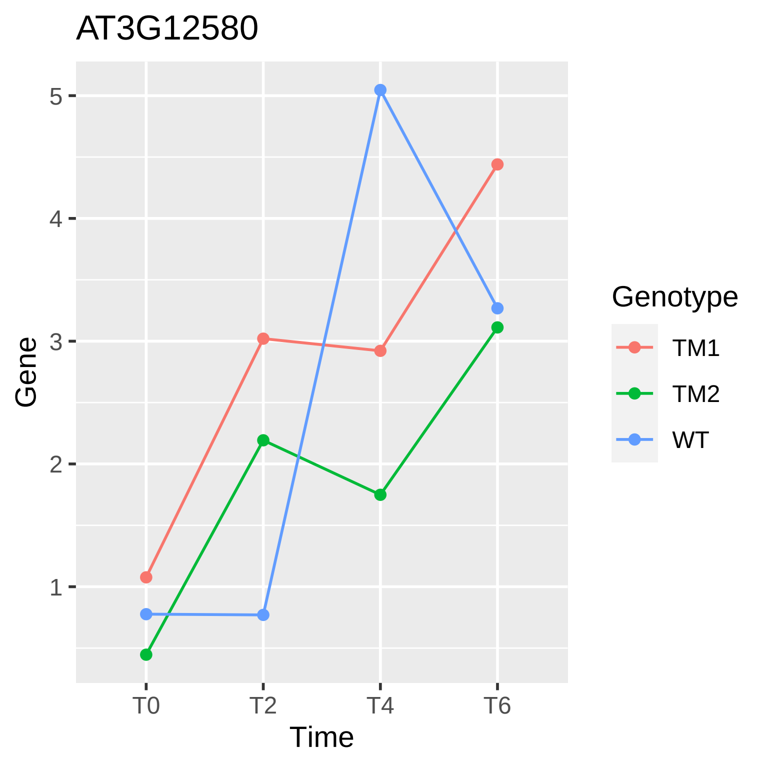
AT3G12580
Gene
5
4
3
2
1
T0
T2
T4
T6
Time
Genotype
TM1
TM2
WT

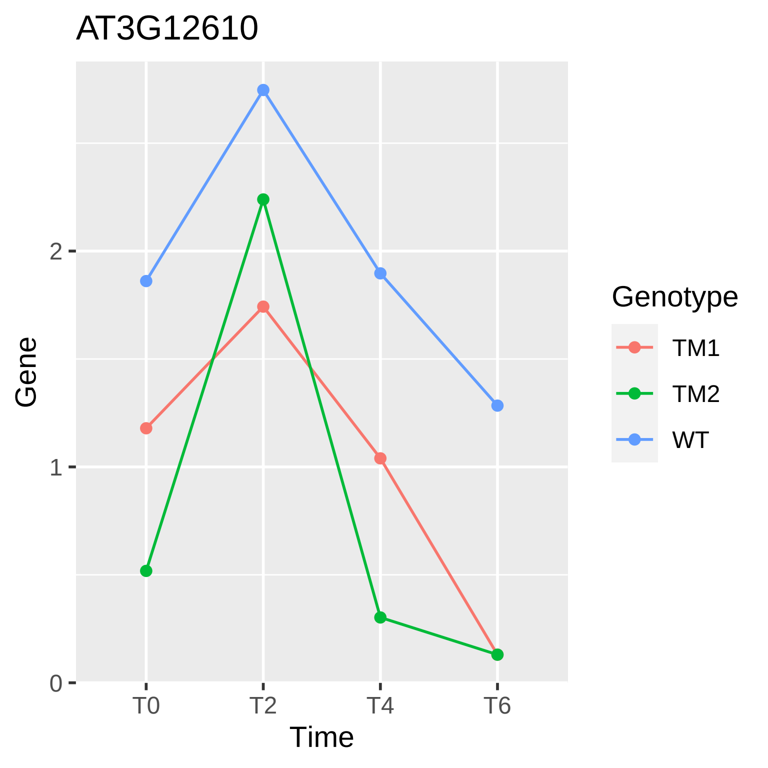

AT3G12610
Gene
2
1
0
T0
T2
T4
T6
Time
Genotype
TM1
TM2
WT

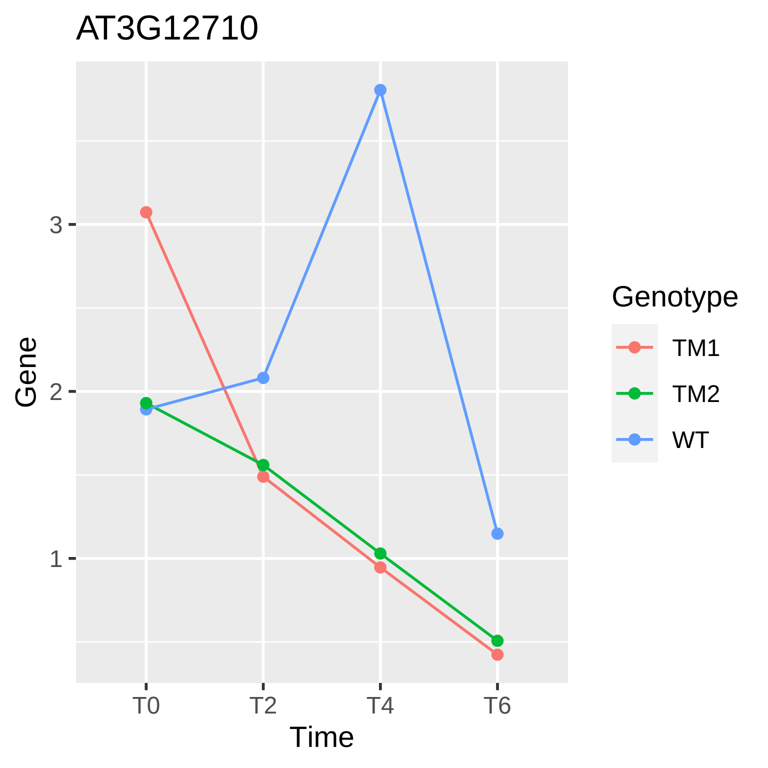

AT3G12710
Gene
3
2
1
T0
T2
T4
T6
Time
Genotype
TM1
TM2
WT

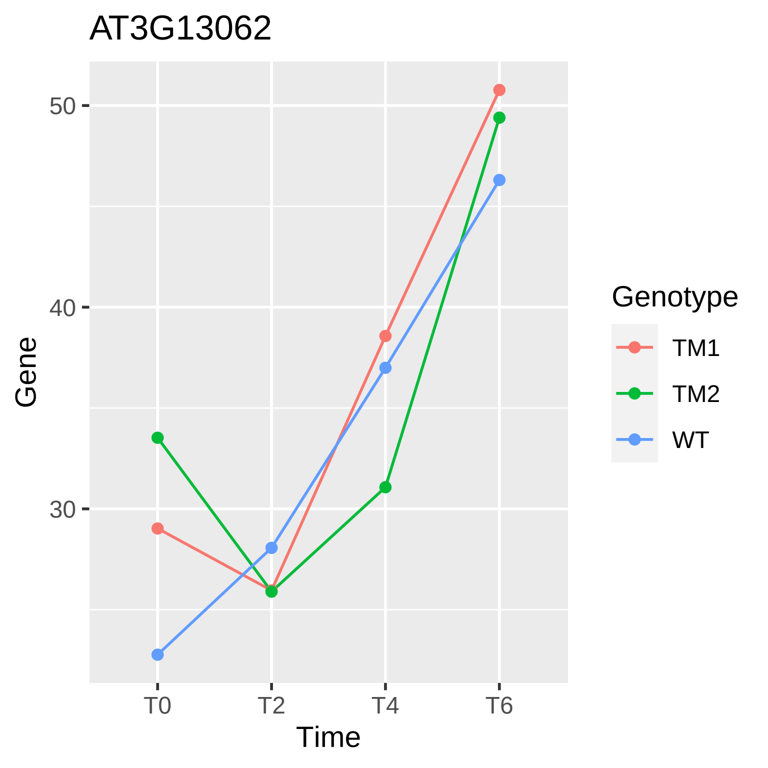

AT3G13062
50
40
30
Gene
T0
T2
T4
T6
Time
Genotype
TM1
TM2
WT

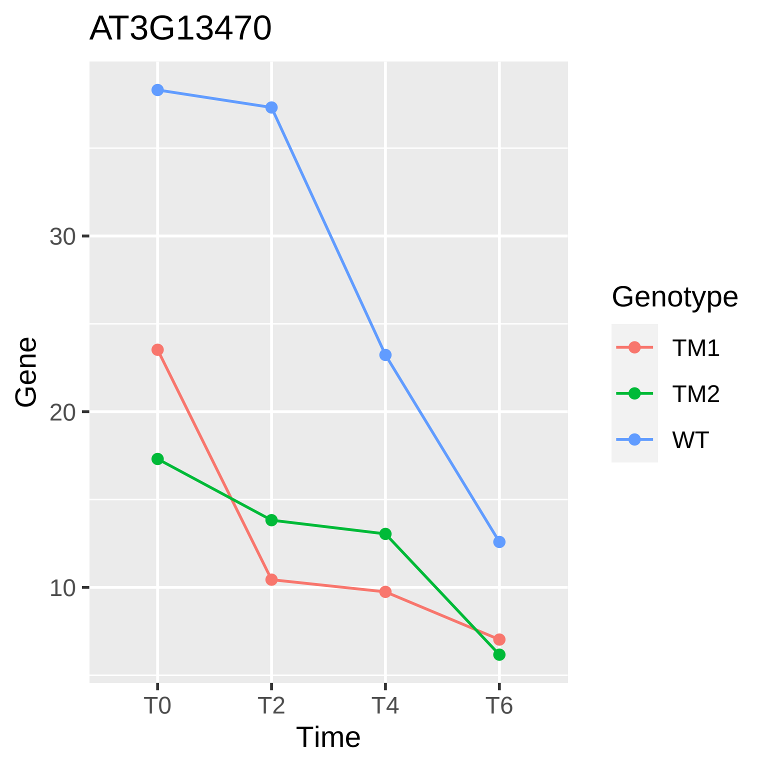
AT3G13470
30
20
10
Gene
T0
T2
T4
T6
Time
Genotype
TM1
TM2
WT

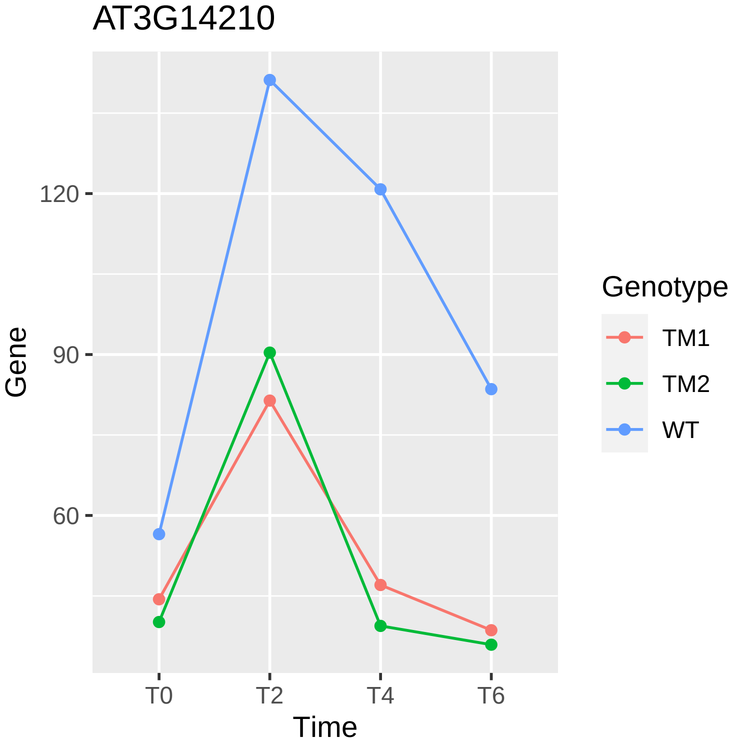
AT3G14210
Gene
120
90
60
T0
T2
T4
T6
Time
Genotype
TM1
TM2
WT

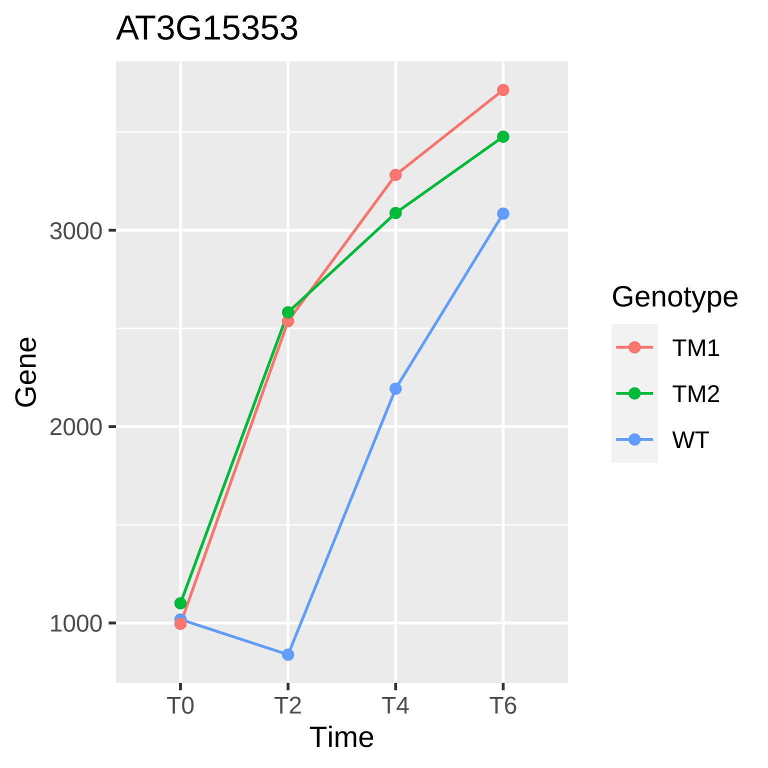
AT3G15353
3000
2000
1000
Gene
T0
T2
T4
T6
Time
Genotype
TM1
TM2
WT

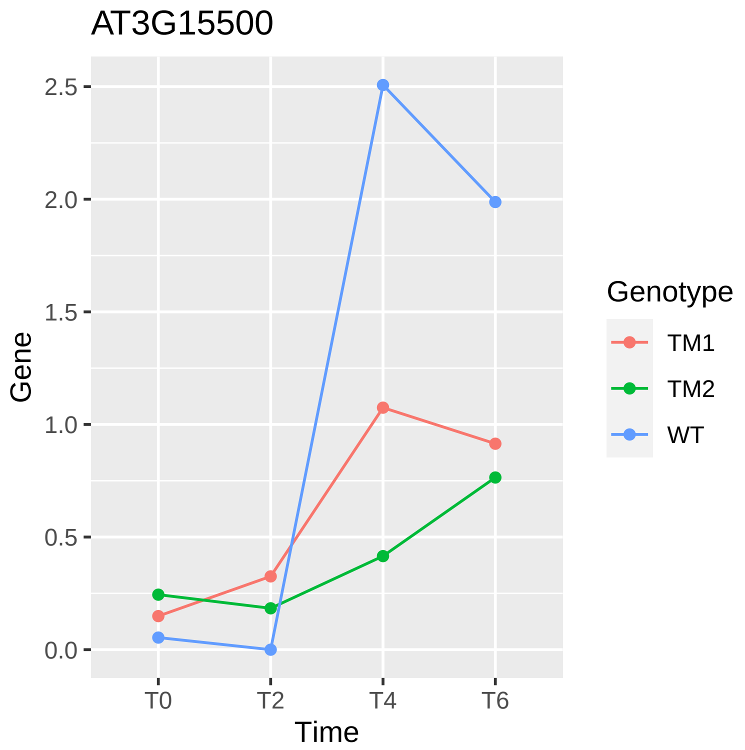

AT3G15500
Gene
2.5
2.0
1.5
1.0
0.5
0.0
T0
T2
T4
T6
Time
Genotype
TM1
TM2
WT

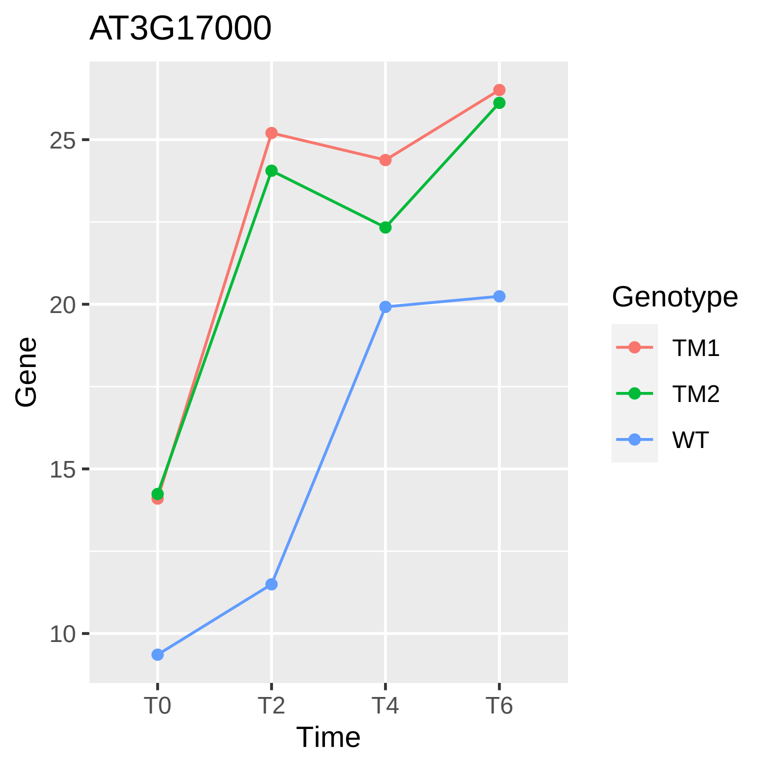

AT3G17000
25
20
15
10
Gene
T0
T2
T4
T6
Time
Genotype
TM1
TM2
WT

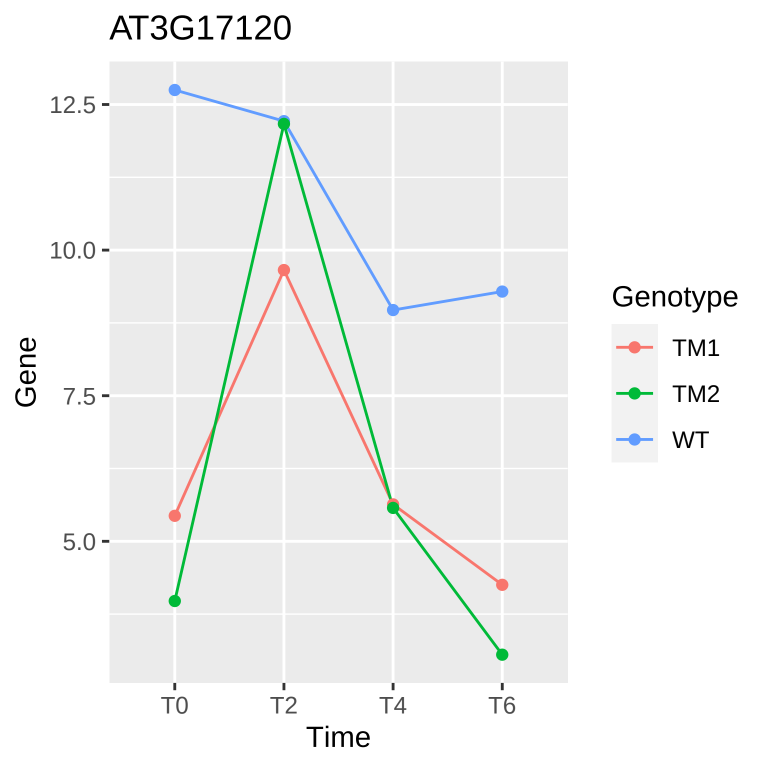
AT3G17120
12.5
10.0
7.5
5.0
Gene
T0
T2
T4
T6
Time
Genotype
TM1
TM2
WT

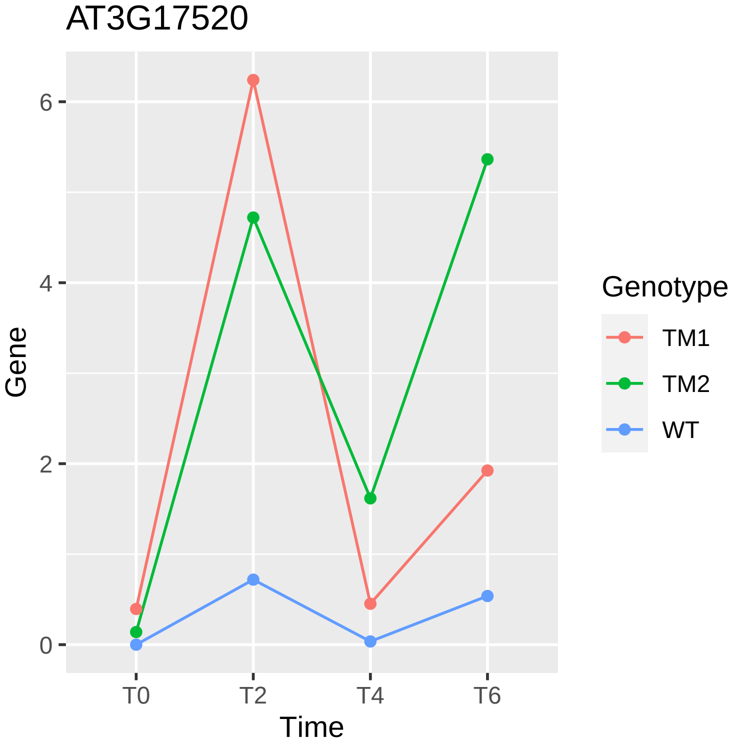

AT3G17520
Gene
Time
0
2
4
6
T0
T2
T4
T6
Genotype
TM1
TM2
WT

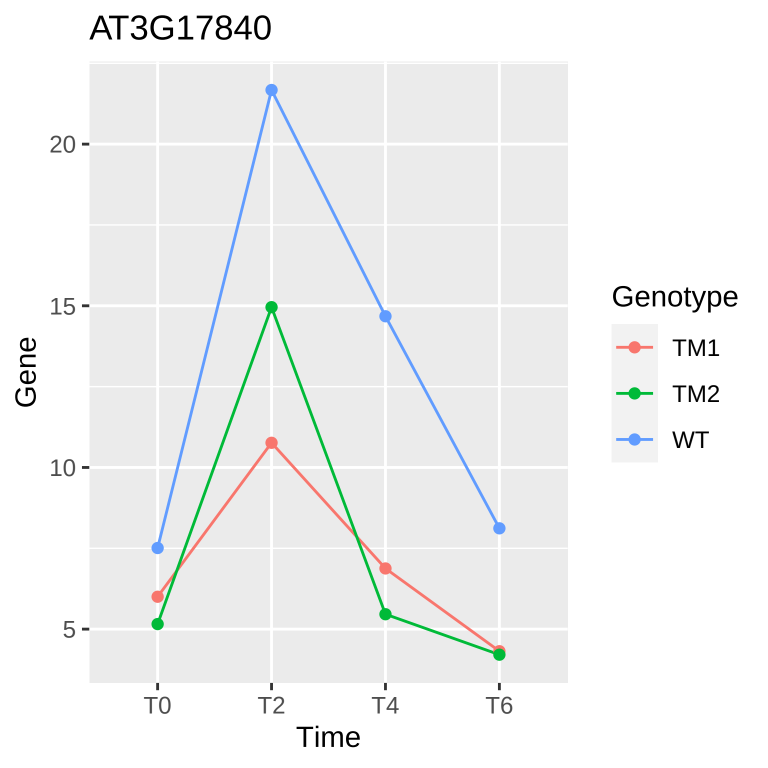

AT3G17840
Gene
20
15
10
5
T0
T2
T4
T6
Time
Genotype
TM1
TM2
WT

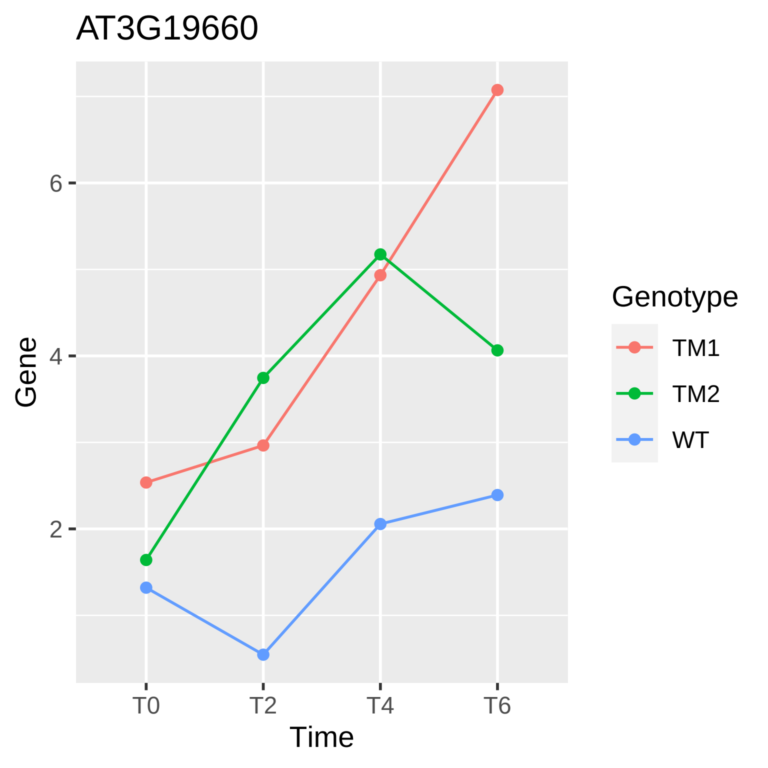
AT3G19660
Gene
6
4
2
T0
T2
T4
T6
Time
Genotype
TM1
TM2
WT

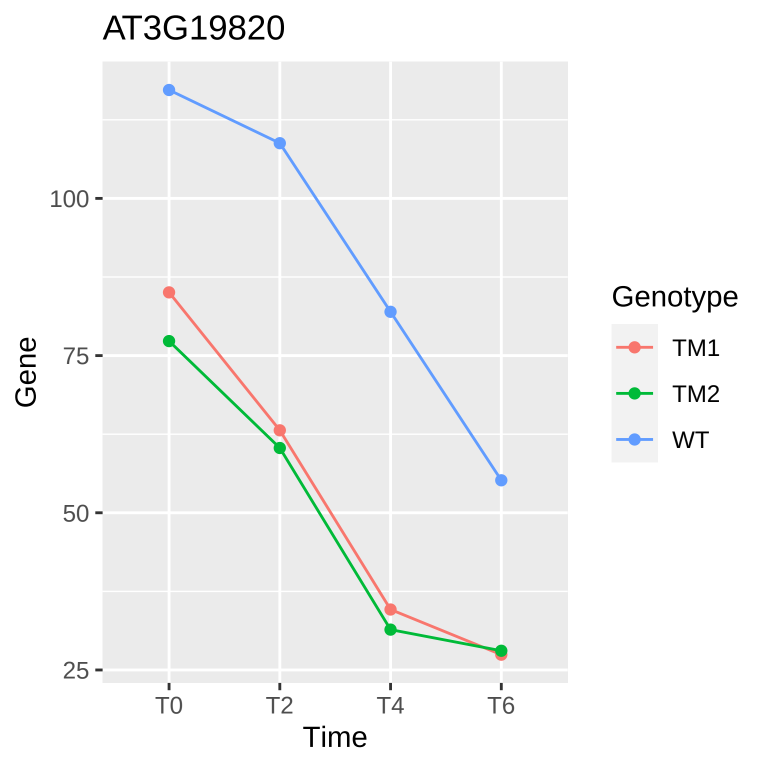

AT3G19820
Gene
100
75
50
25
T0
T2
T4
T6
Time
Genotype
TM1
TM2
WT

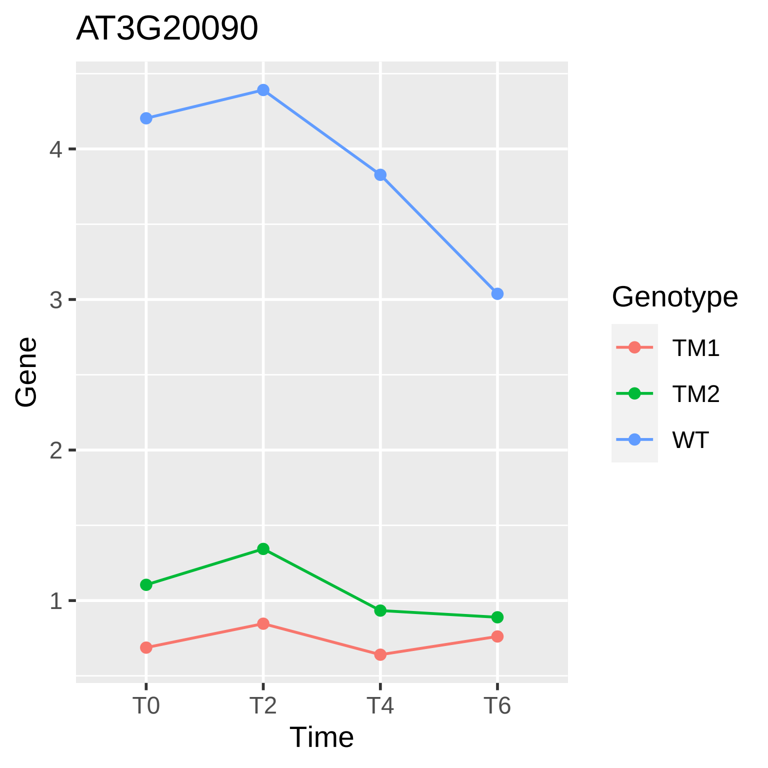
AT3G20090
Gene
4
3
2
1
T0
T2
T4
T6
Time
Genotype
TM1
TM2
WT

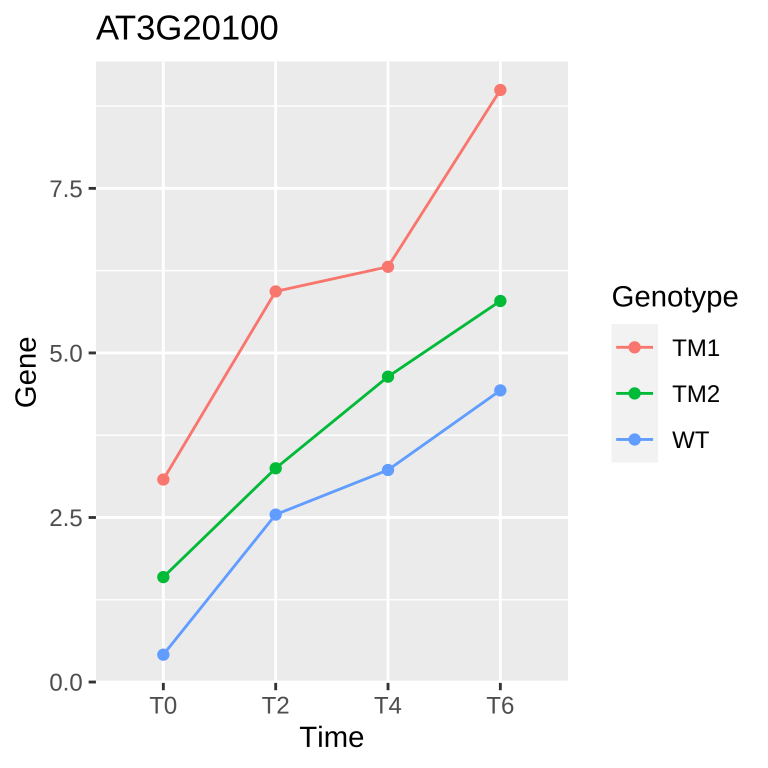

AT3G20100
Gene
7.5
5.0
2.5
0.0
T0
T2
T4
T6
Time
Genotype
TM1
TM2
WT

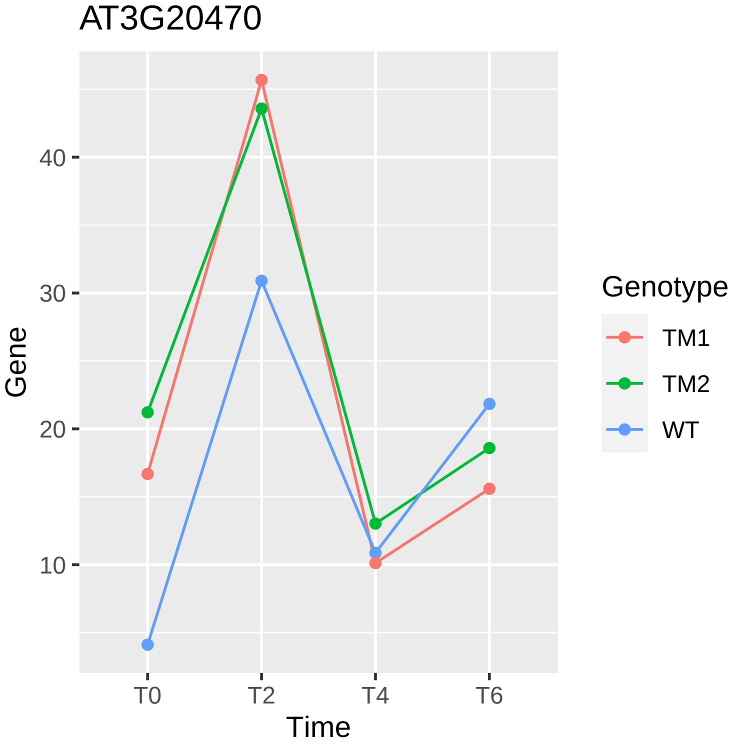
AT3G20470
Gene
40
30
20
10
T0
T2
T4
T6
Time
Genotype
TM1
TM2
WT

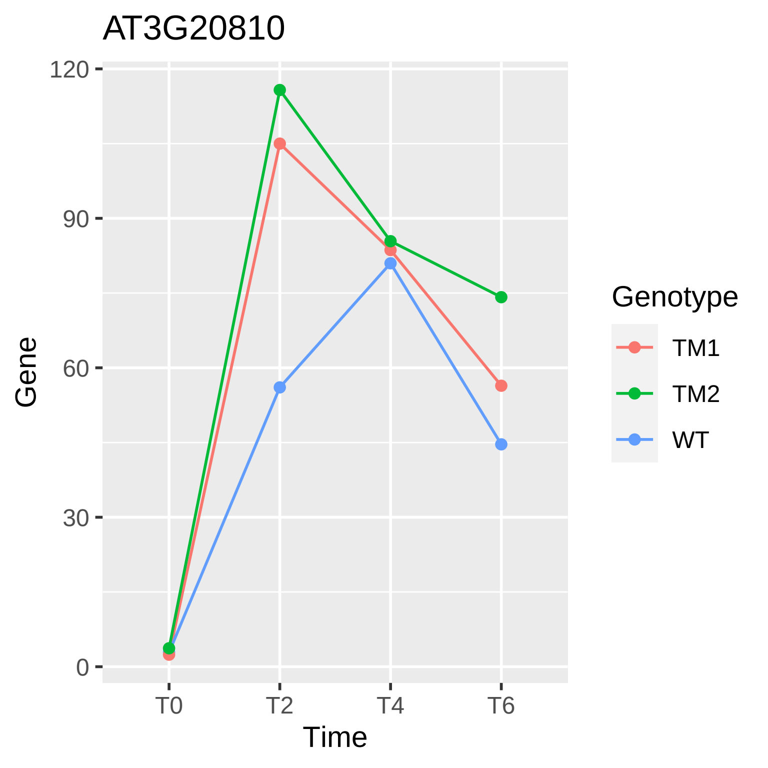
AT3G20810
Gene
120
90
60
30
0
T0
T2
T4
T6
Time
Genotype
TM1
TM2
WT

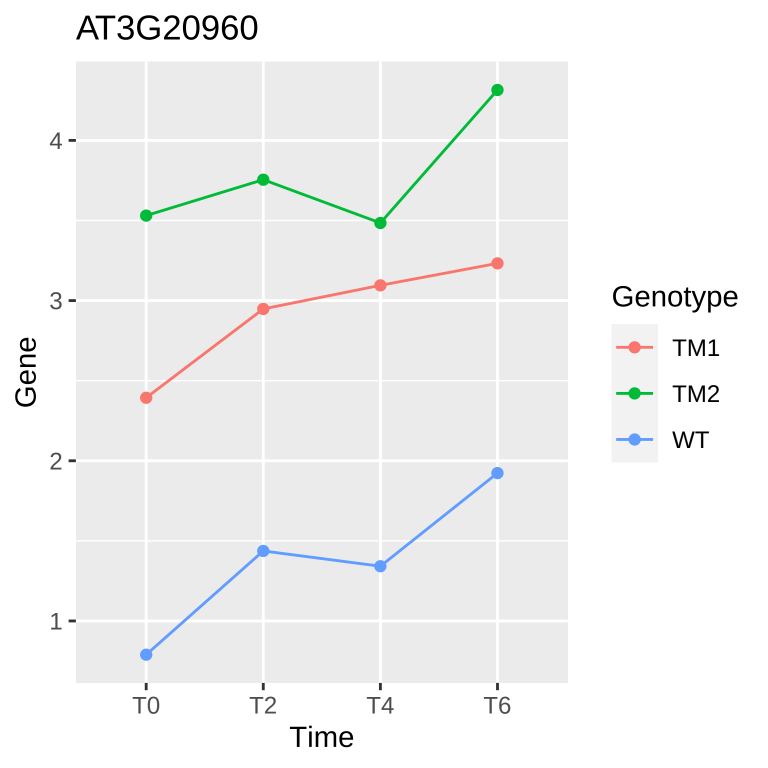
AT3G20960
Gene
4
3
2
1
T0
T2
T4
T6
Time
Genotype
TM1
TM2
WT

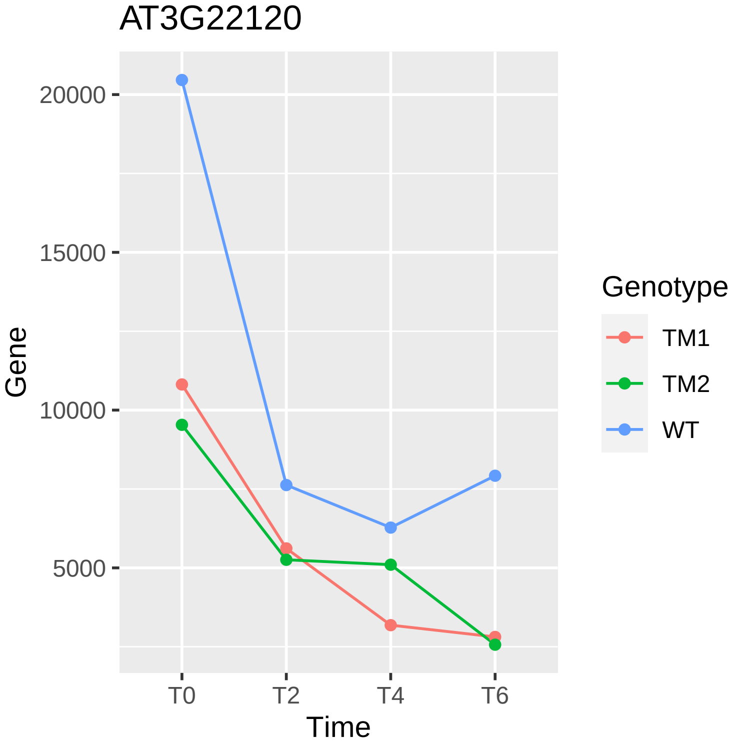
AT3G22120
20000
15000
10000
5000
Gene
T0
T2
T4
T6
Time
Genotype
TM1
TM2
WT

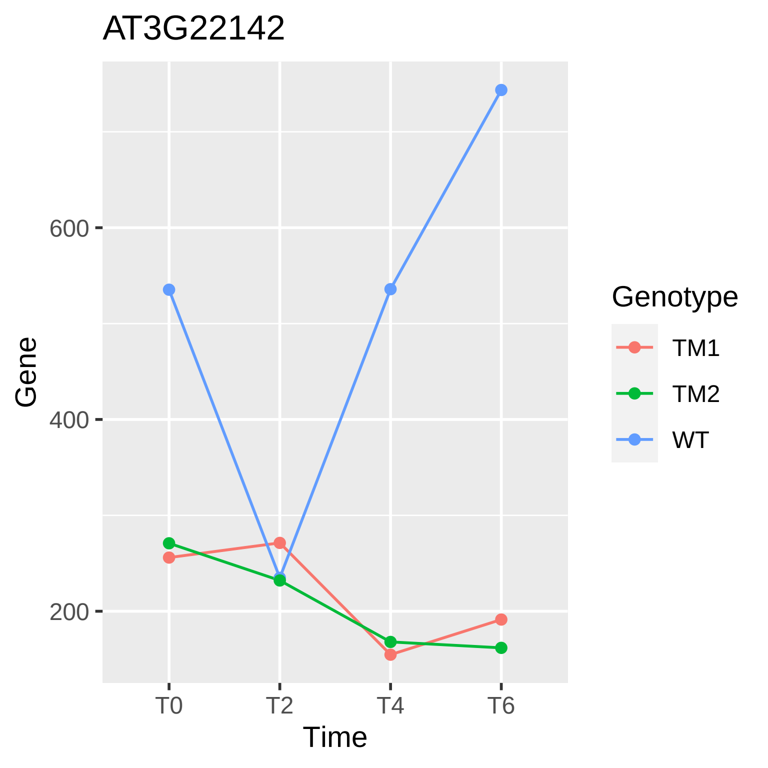
AT3G22142
Gene
600
400
200
T0
T2
T4
T6
Time
Genotype
TM1
TM2
WT

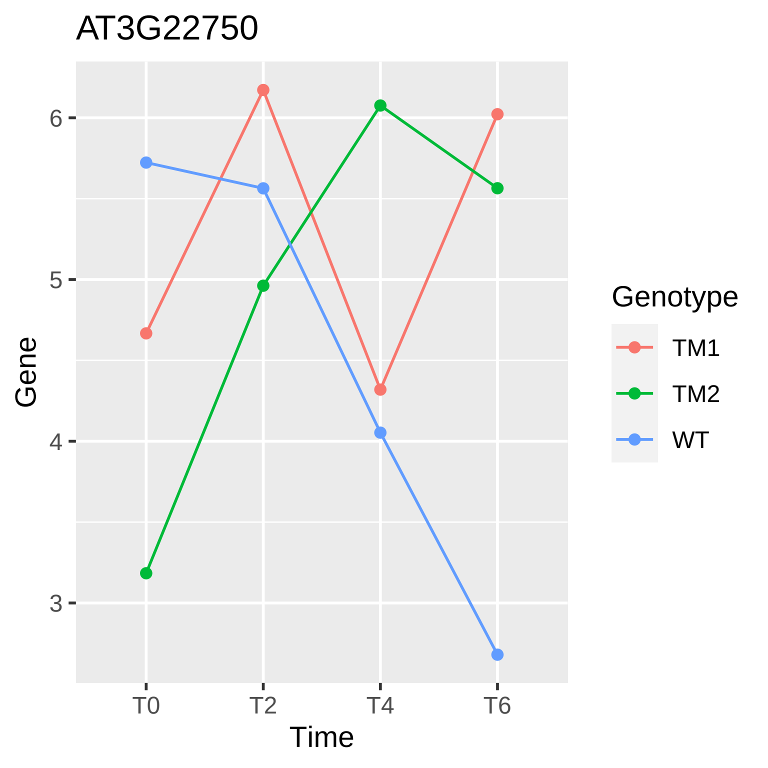
AT3G22750
Gene
6
5
4
3
T0
T2
T4
T6
Time
Genotype
TM1
TM2
WT

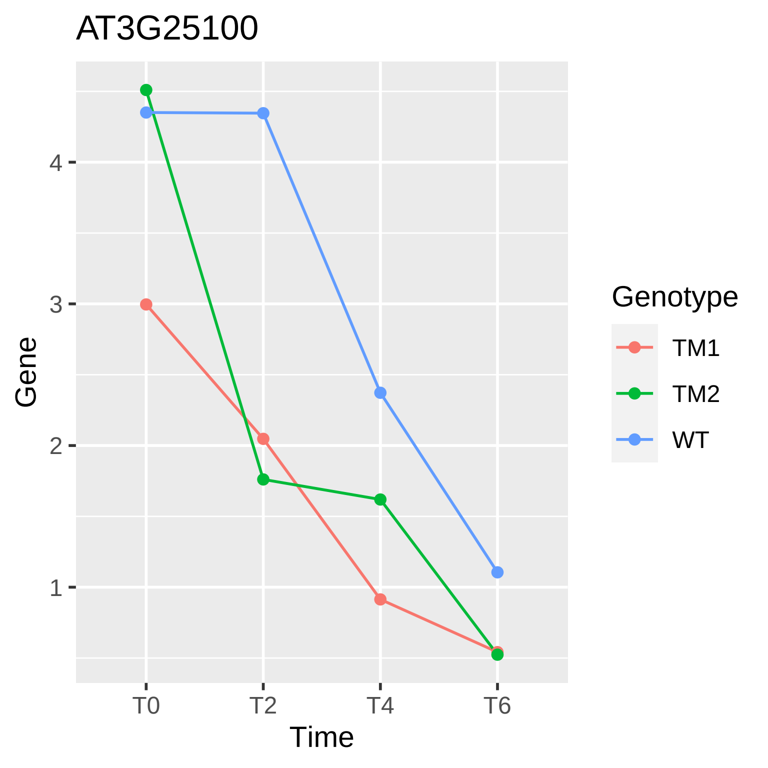
AT3G25100
Gene
4
3
2
1
T0
T2
T4
T6
Time
Genotype
TM1
TM2
WT

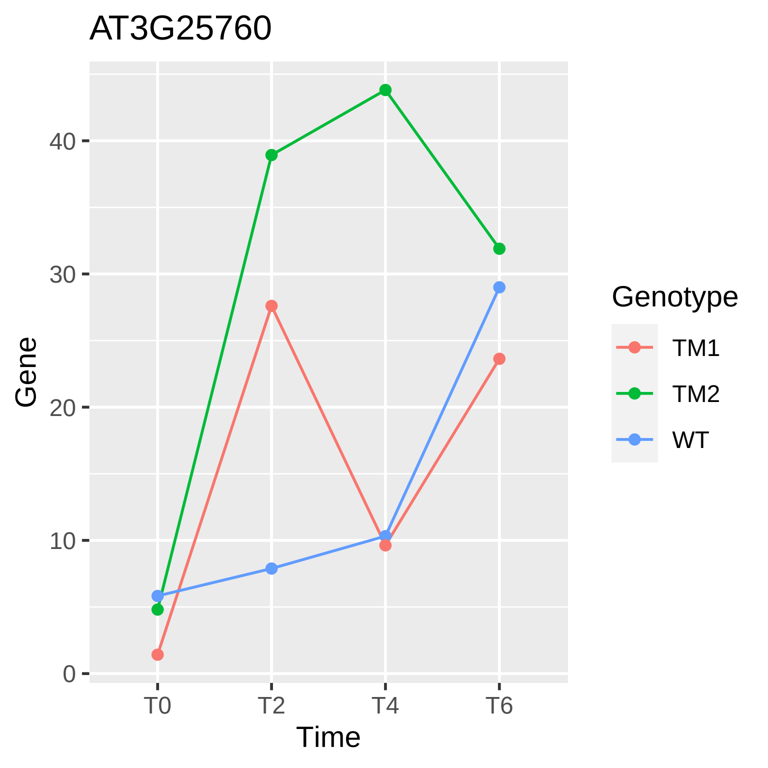
AT3G25760
Gene
Time
0
10
20
30
40
T0
T2
T4
T6
Genotype
TM1
TM2
WT

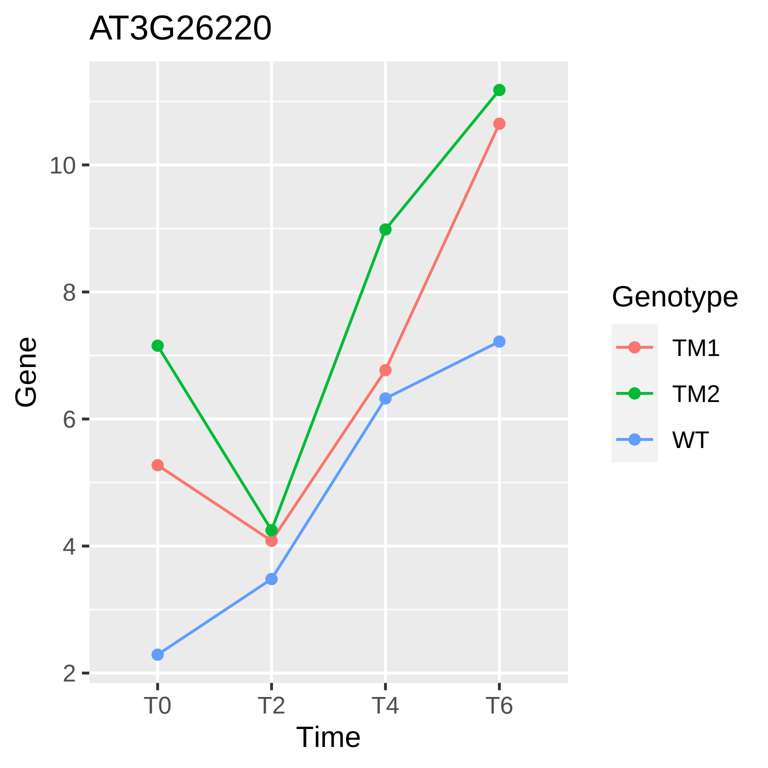

AT3G26220
Gene
10
8
6
4
2
T0
T2
T4
T6
Time
Genotype
TM1
TM2
WT

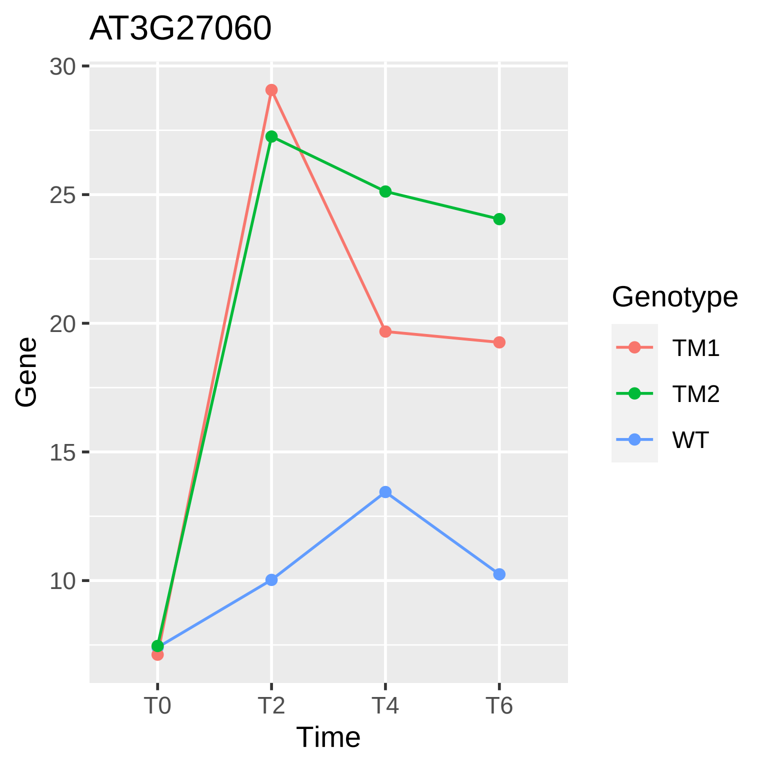

AT3G27060
30
25
20
15
10
Gene
T0
T2
T4
T6
Time
Genotype
TM1
TM2
WT

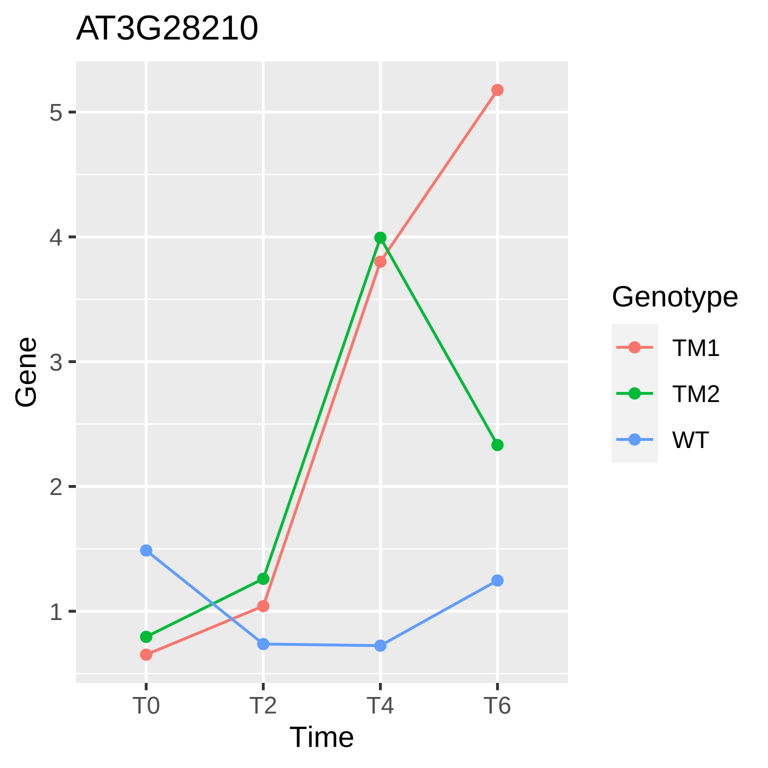
AT3G28210
Gene
5
4
3
2
1
T0
T2
T4
T6
Time
Genotype
TM1
TM2
WT

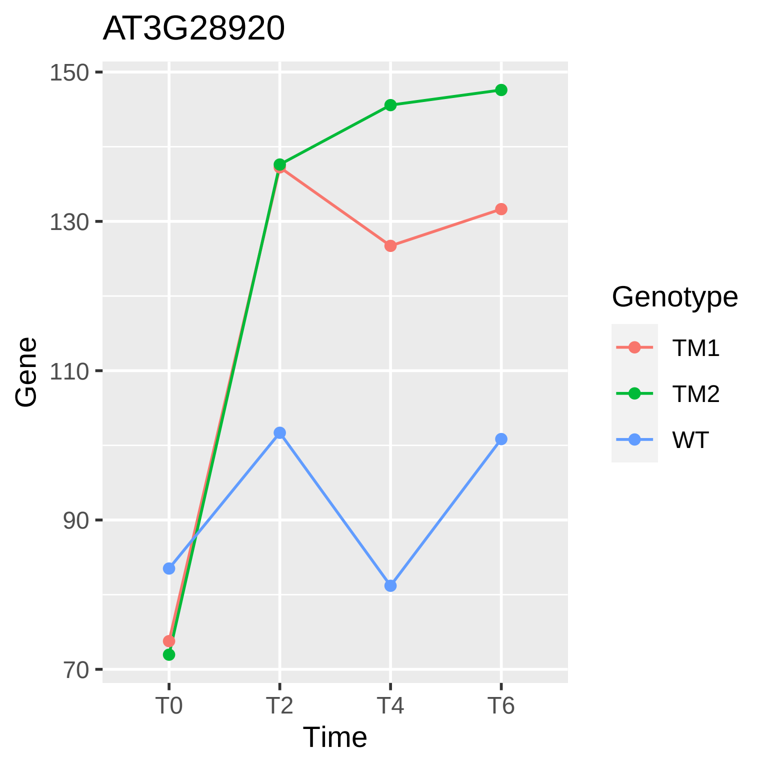

AT3G28920
Gene
150
130
110
90
70
T0
T2
T4
T6
Time
Genotype
TM1
TM2
WT

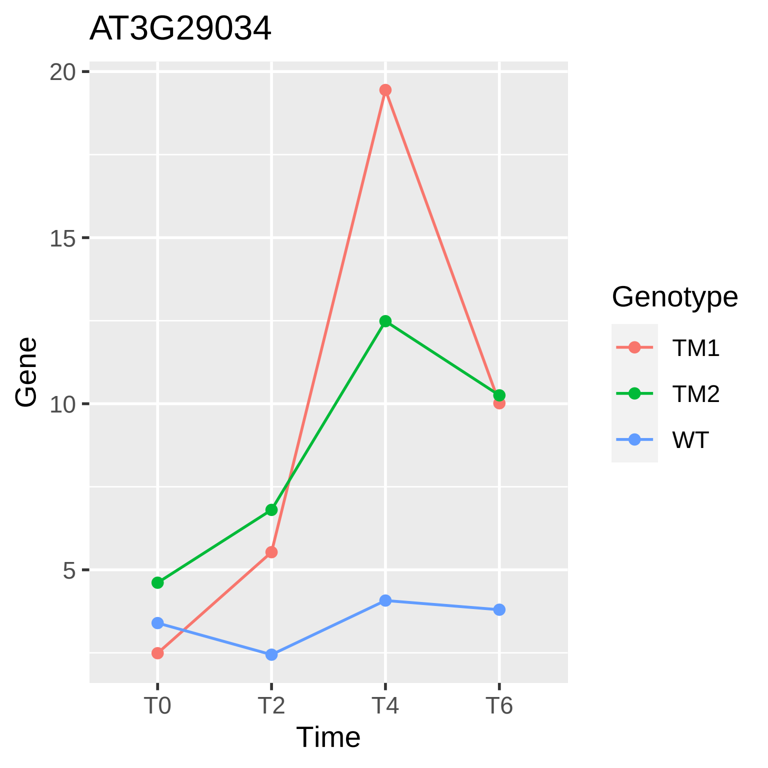
AT3G29034
Gene
20
15
10
5
T0
T2
T4
T6
Time
Genotype
TM1
TM2
WT

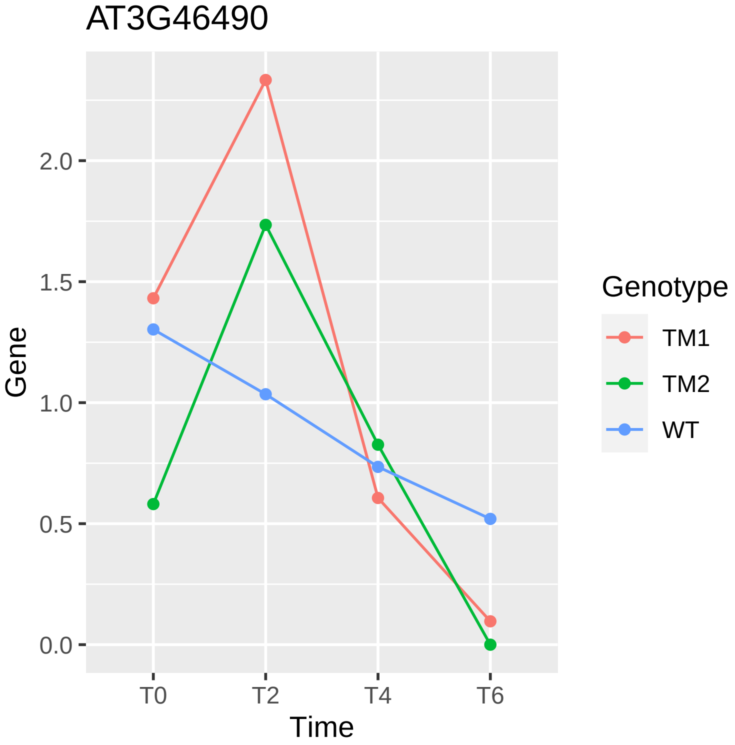
AT3G46490
Gene
2.0
1.5
1.0
0.5
0.0
T0
T2
T4
T6
Time
Genotype
TM1
TM2
WT

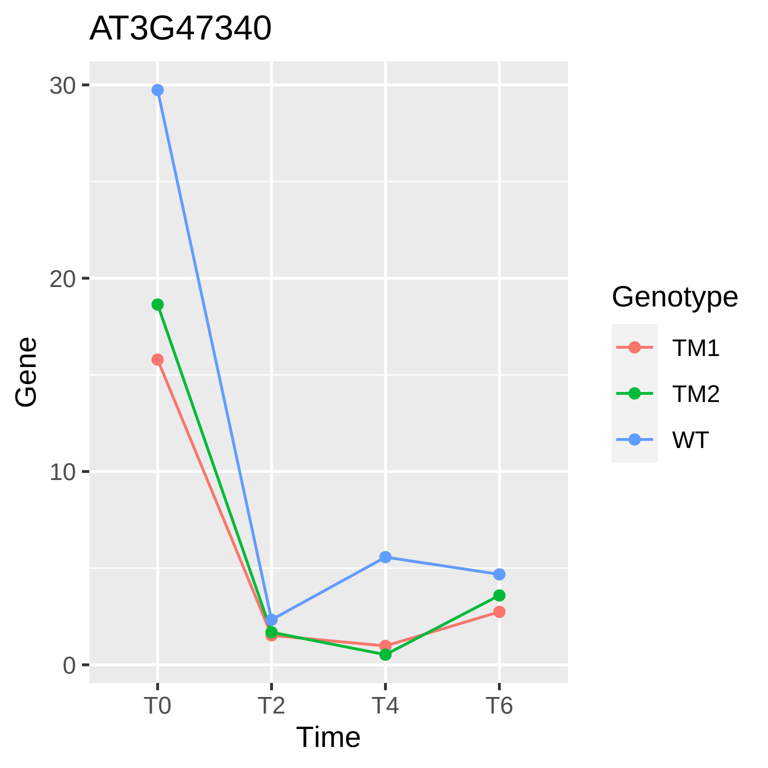

AT3G47340
30
20
10
0
Gene
T0
T2
T4
T6
Time
Genotype
TM1
TM2
WT

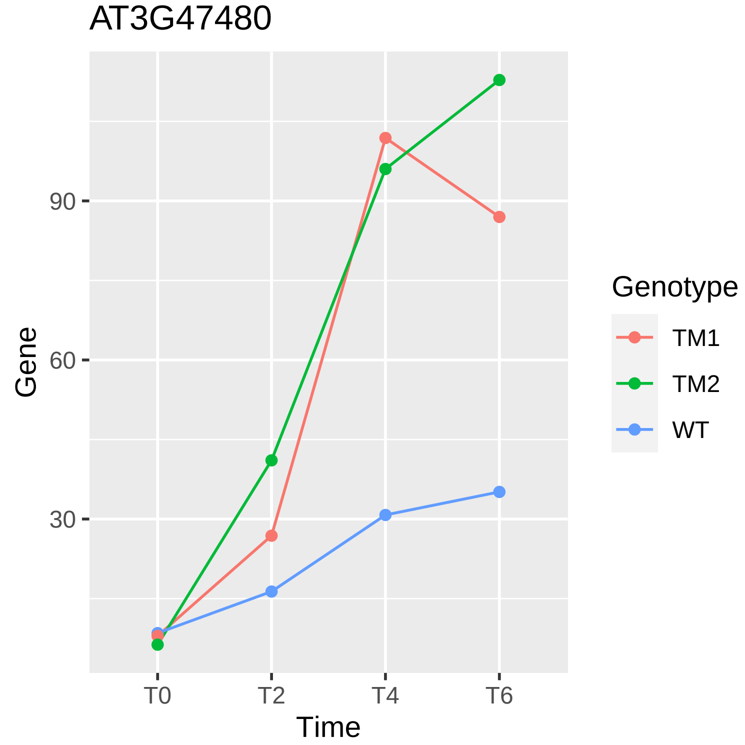
AT3G47480
Gene
90
60
30
T0
T2
T4
T6
Time
Genotype
TM1
TM2
WT

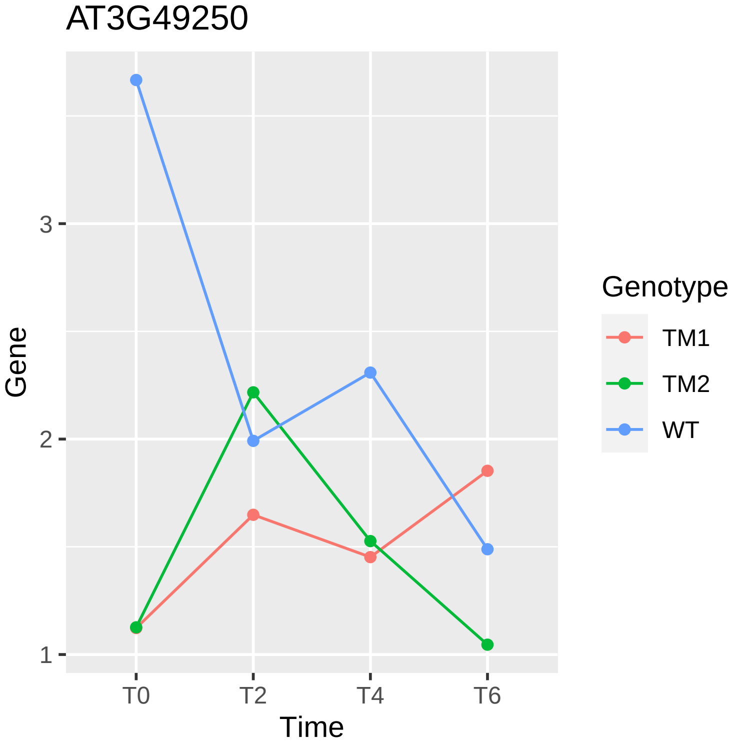
AT3G49250
Gene
3
2
1
T0
T2
T4
T6
Time
Genotype
TM1
TM2
WT

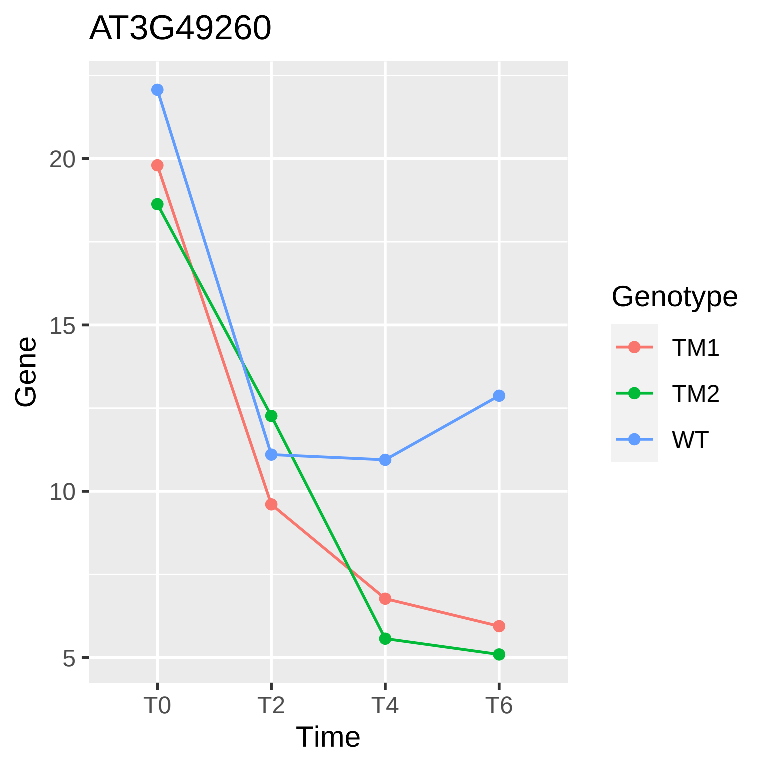
AT3G49260
Gene
20
15
10
5
T0
T2
T4
T6
Time
Genotype
TM1
TM2
WT

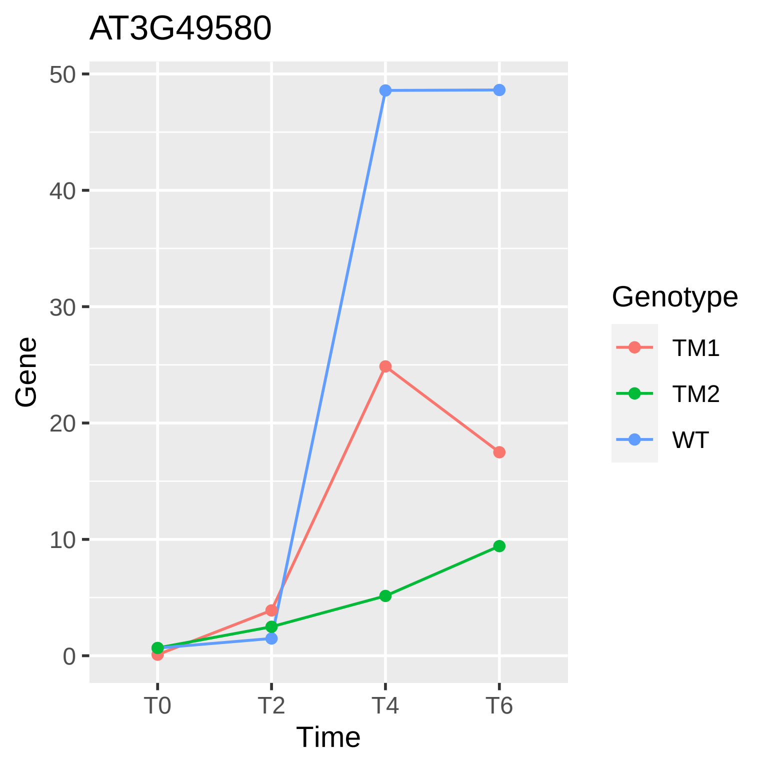

AT3G49580
Gene
0
10
20
30
40
50
T0
T2
T4
T6
Time
Genotype
TM1
TM2
WT

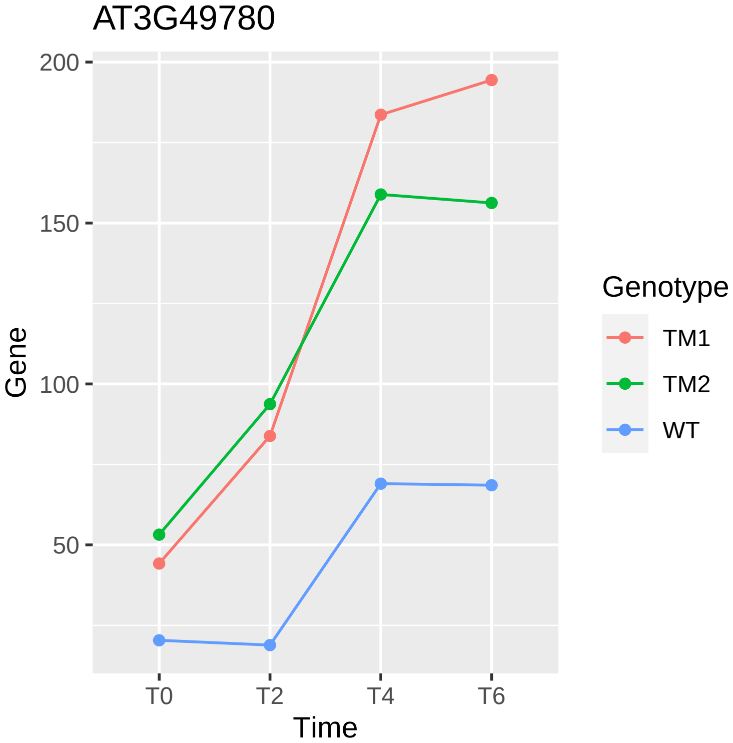
AT3G49780
200
150
100
50
Gene
T0
T2
T4
T6
Time
Genotype
TM1
TM2
WT

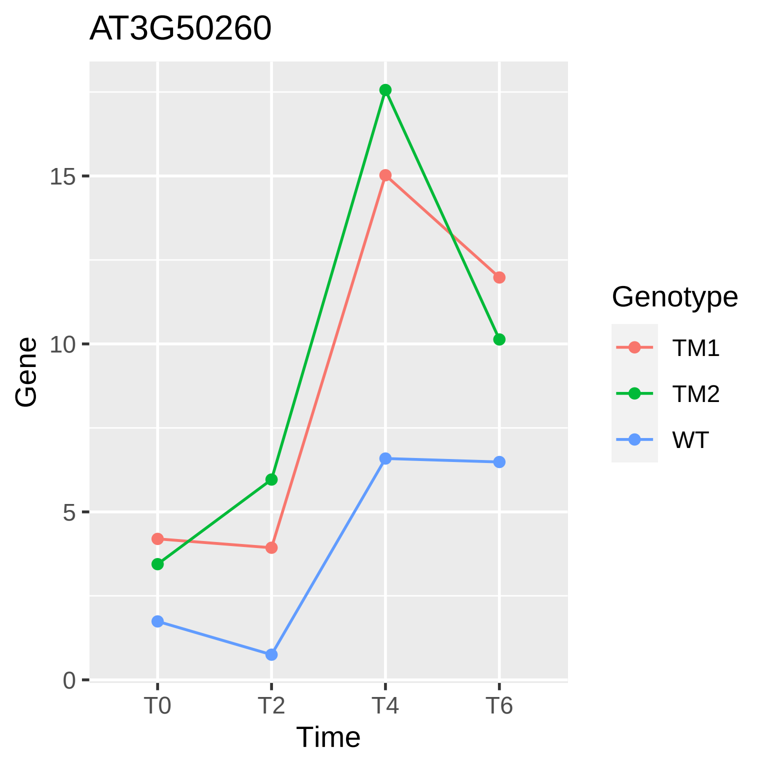
AT3G50260
Gene
15
10
5
0
T0
T2
T4
T6
Time
Genotype
TM1
TM2
WT

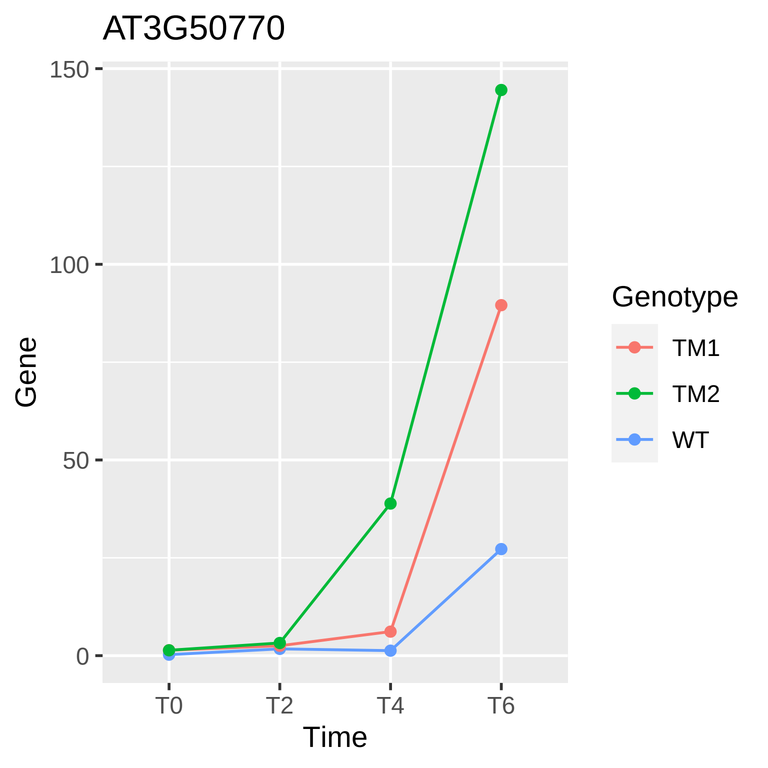
AT3G50770
150
100
50
0
Gene
T0
T2
T4
T6
Time
Genotype
TM1
TM2
WT

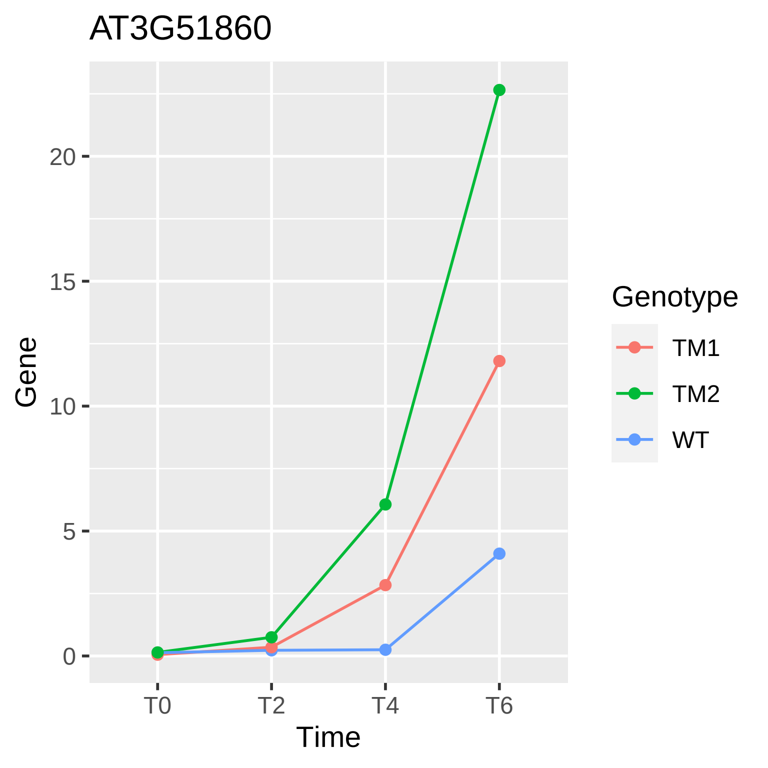

AT3G51860
Gene
20
15
10
5
0
T0
T2
T4
T6
Time
Genotype
TM1
TM2
WT

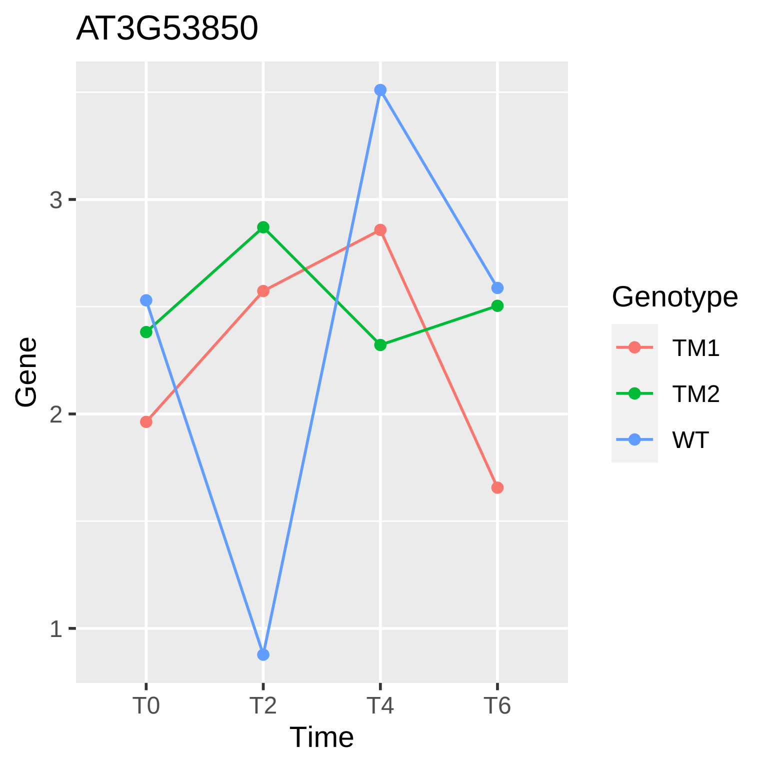

AT3G53850
Gene
Time
3
2
1
T0
T2
T4
T6
Genotype
TM1
TM2
WT

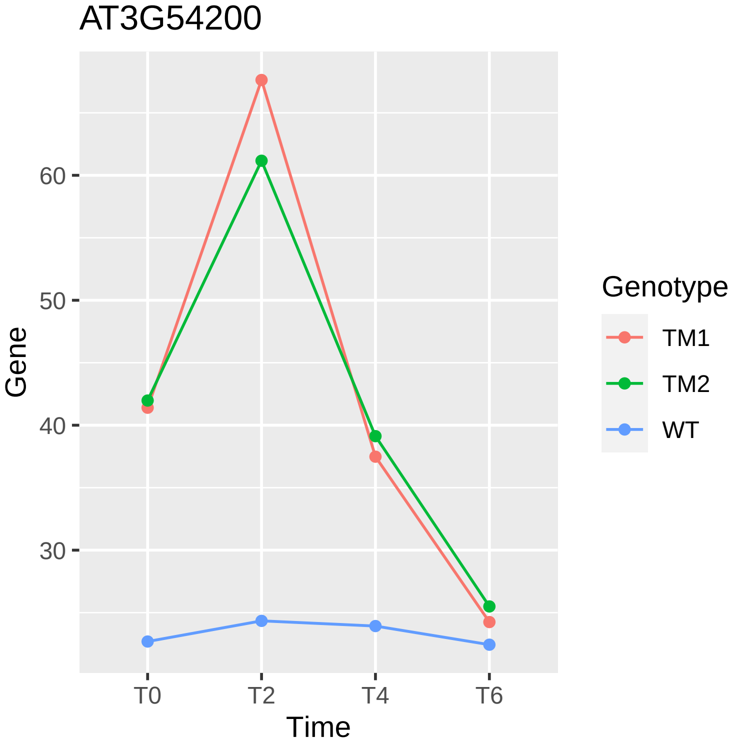

AT3G54200
Gene
60
50
40
30
T0
T2
T4
T6
Time
Genotype
TM1
TM2
WT

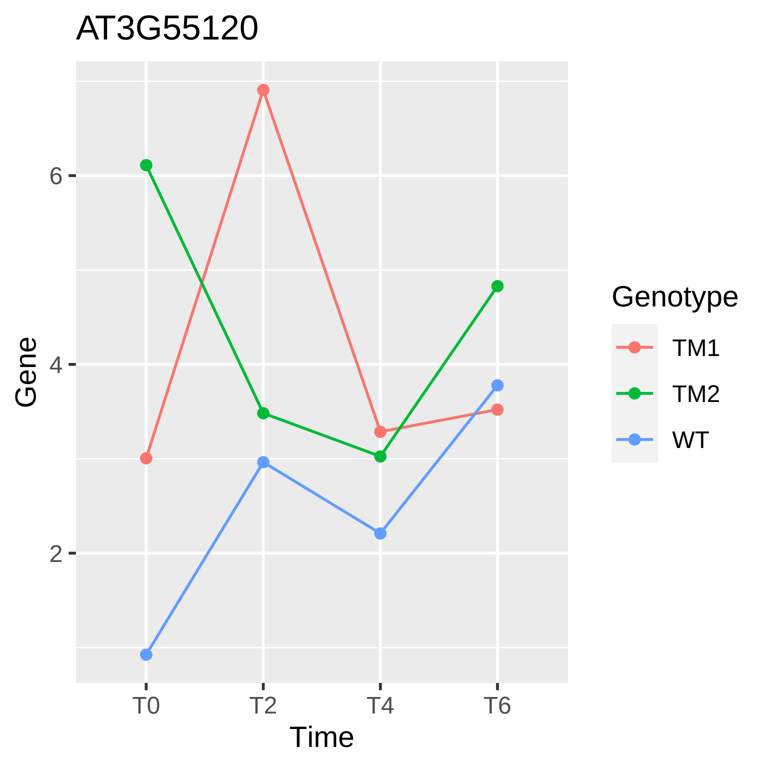

AT3G55120
Gene
6
4
2
T0
T2
T4
T6
Time
Genotype
TM1
TM2
WT

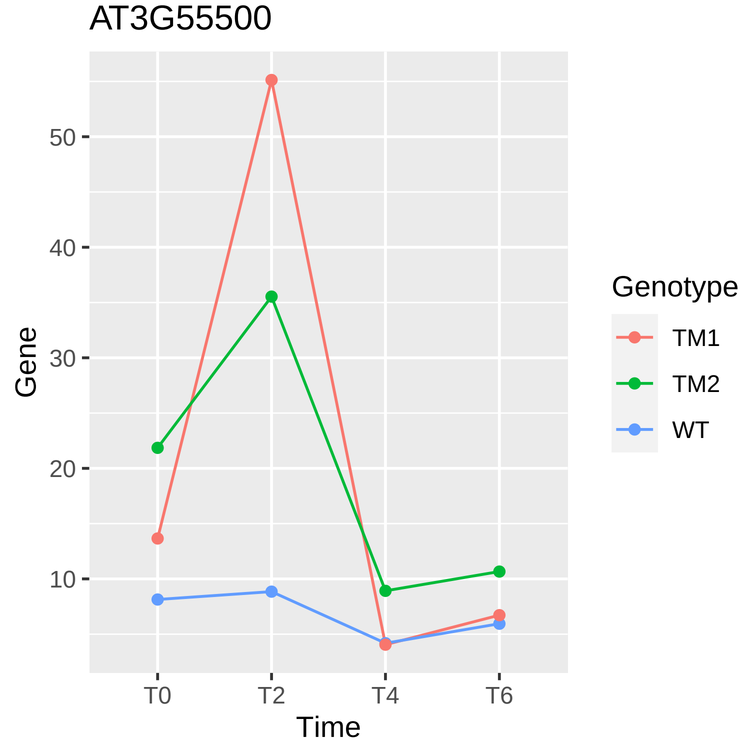
AT3G55500
Gene
50
40
30
20
10
T0
T2
T4
T6
Time
Genotype
TM1
TM2
WT

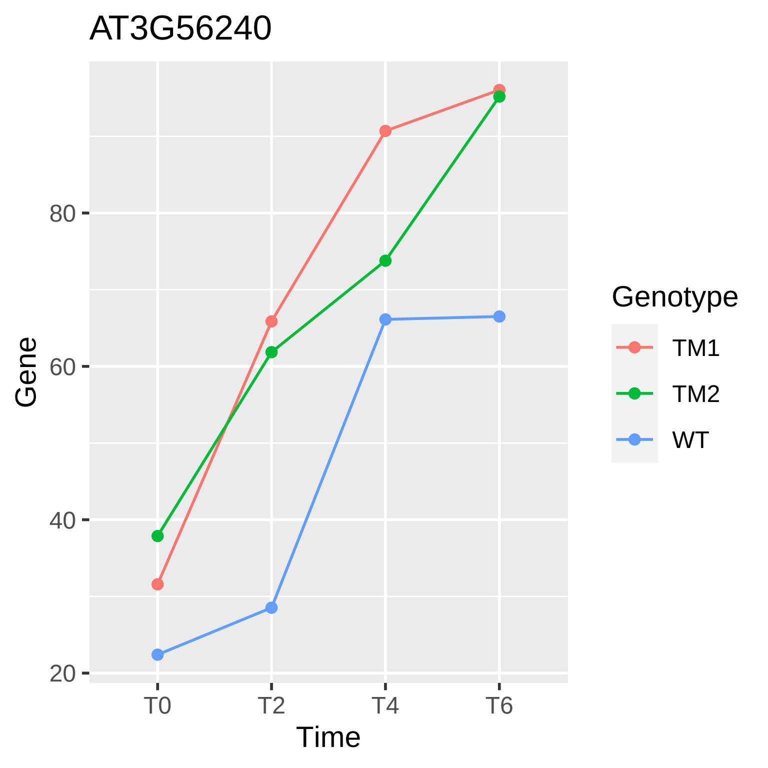

AT3G56240
Gene
80
60
40
20
T0
T2
T4
T6
Time
Genotype
TM1
TM2
WT

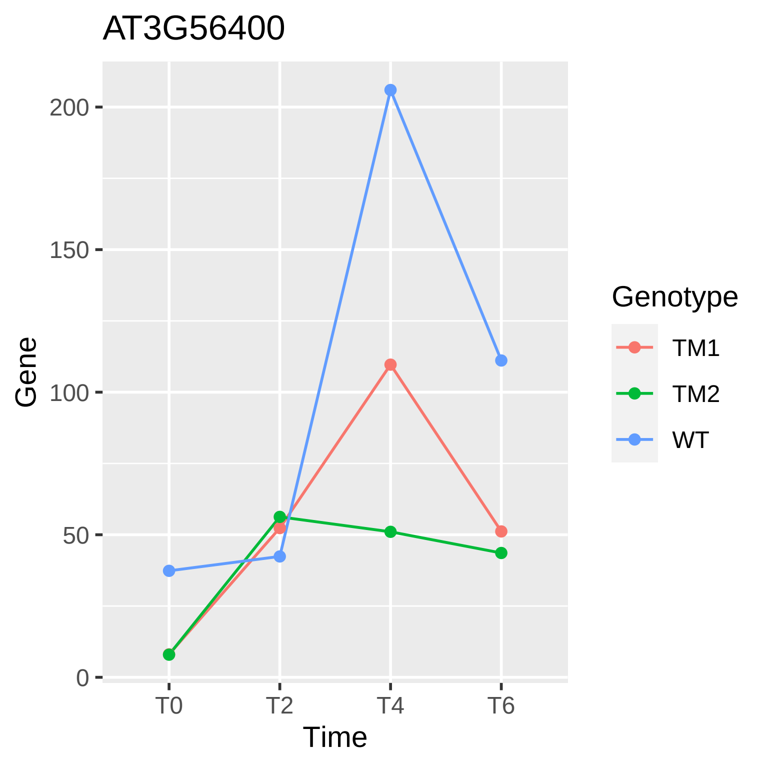
AT3G56400
Gene
200
150
100
50
0
T0
T2
T4
T6
Time
Genotype
TM1
TM2
WT

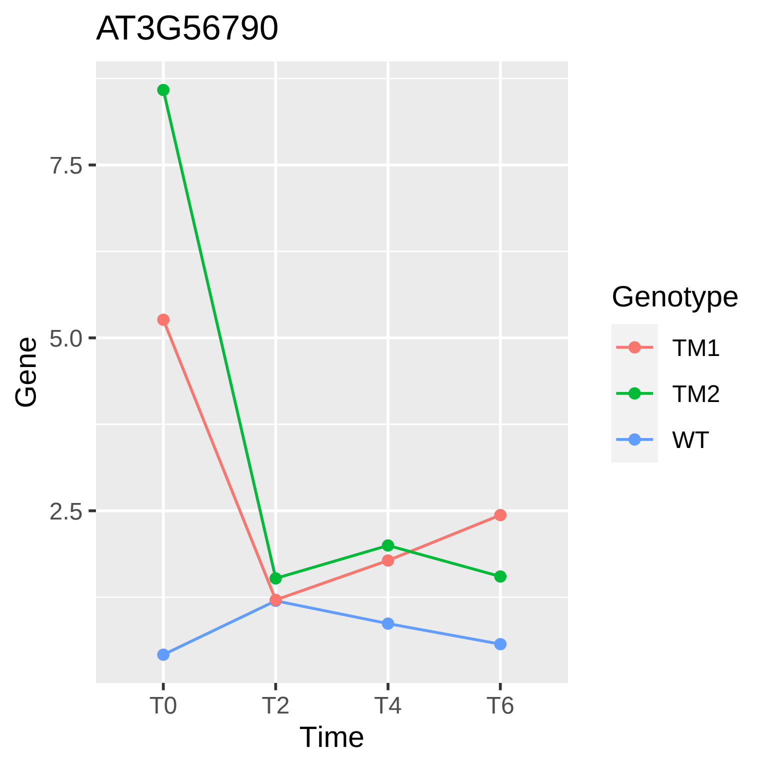

AT3G56790
7.5
5.0
2.5
Gene
T0
T2
T4
T6
Time
Genotype
TM1
TM2
WT

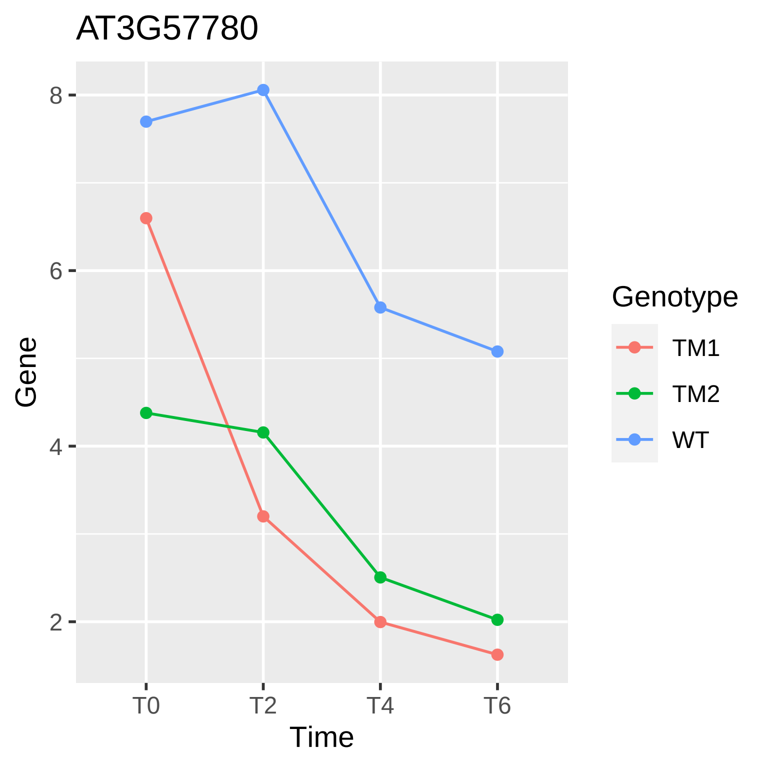

AT3G57780
Gene
8
6
4
2
T0
T2
T4
T6
Time
Genotype
TM1
TM2
WT

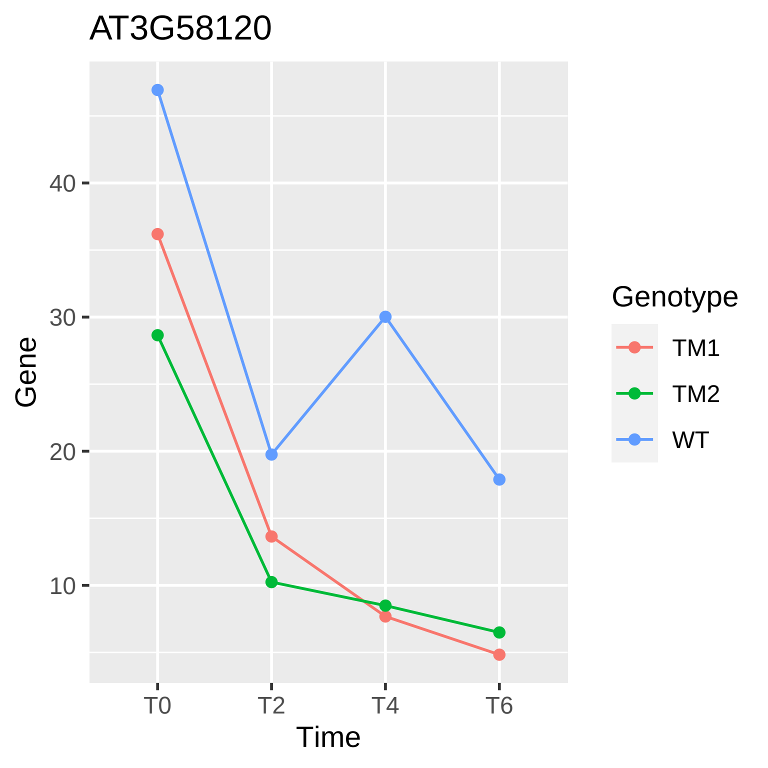

AT3G58120
Gene
40
30
20
10
T0
T2
T4
T6
Time
Genotype
TM1
TM2
WT

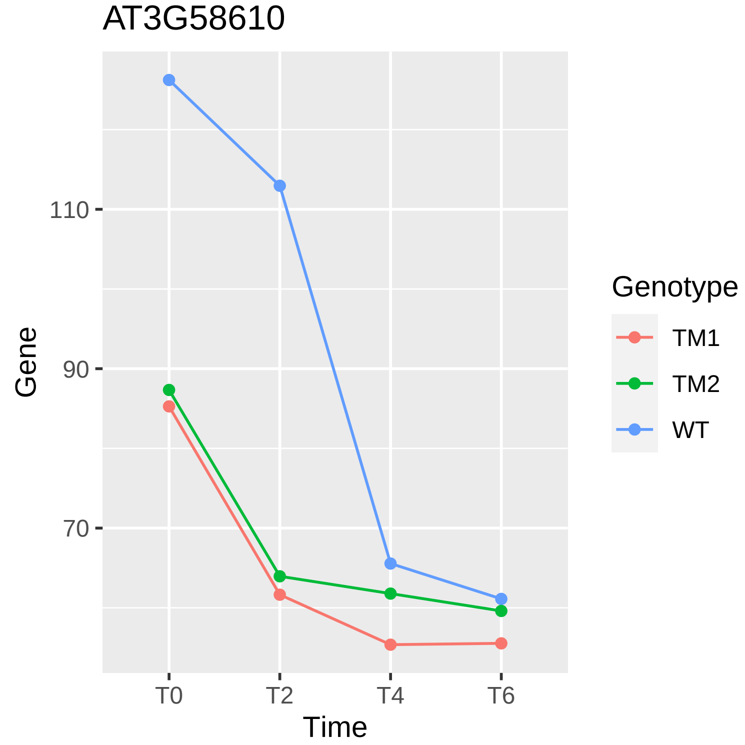
AT3G58610
110
90
70
Gene
T0
T2
T4
T6
Time
Genotype
TM1
TM2
WT

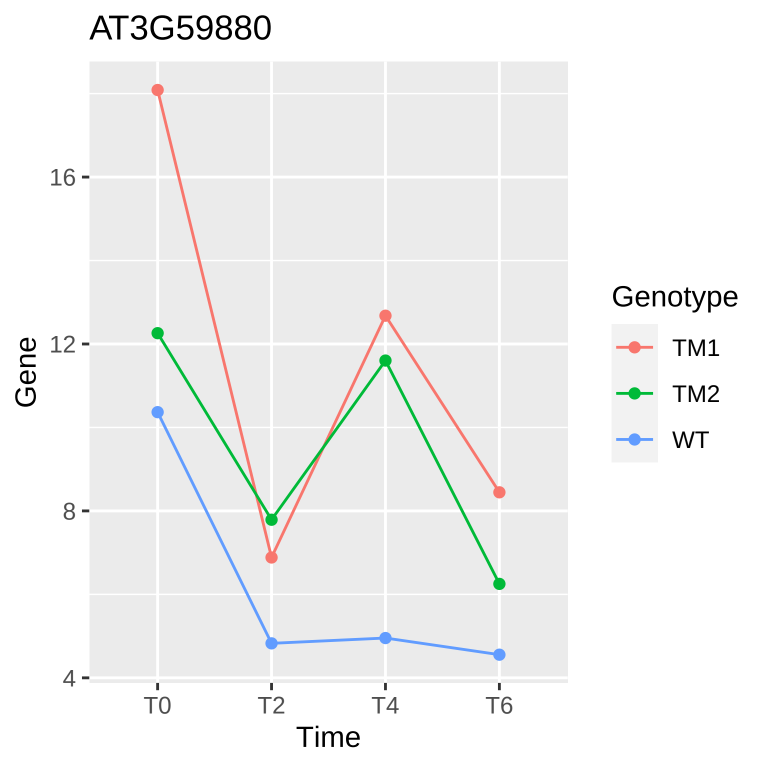

AT3G59880
Gene
16
12
8
4
T0
T2
T4
T6
Time
Genotype
TM1
TM2
WT

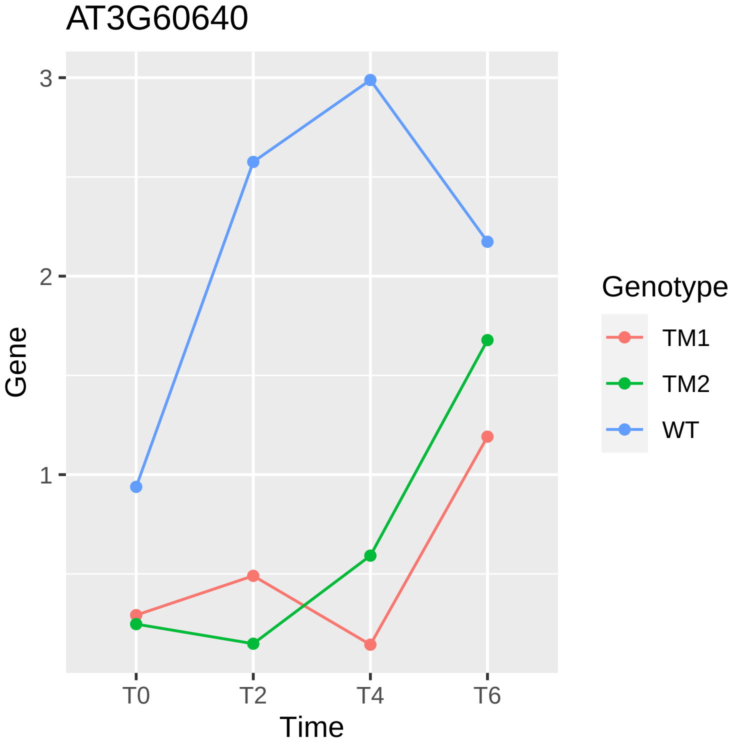
AT3G60640
3
2
1
Gene
T0
T2
T4
T6
Time
Genotype
TM1
TM2
WT

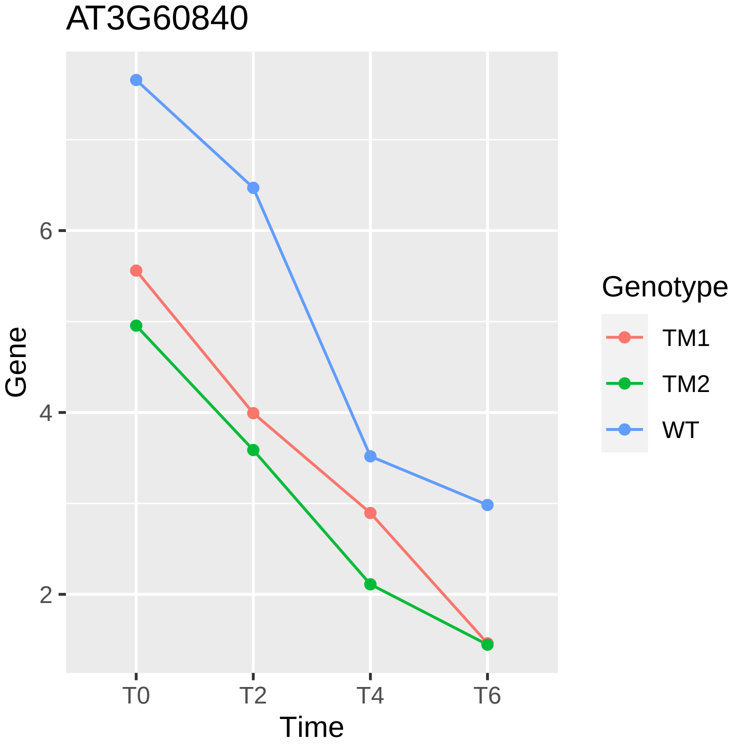

AT3G60840
Gene
6
4
2
T0
T2
T4
T6
Time
Genotype
TM1
TM2
WT

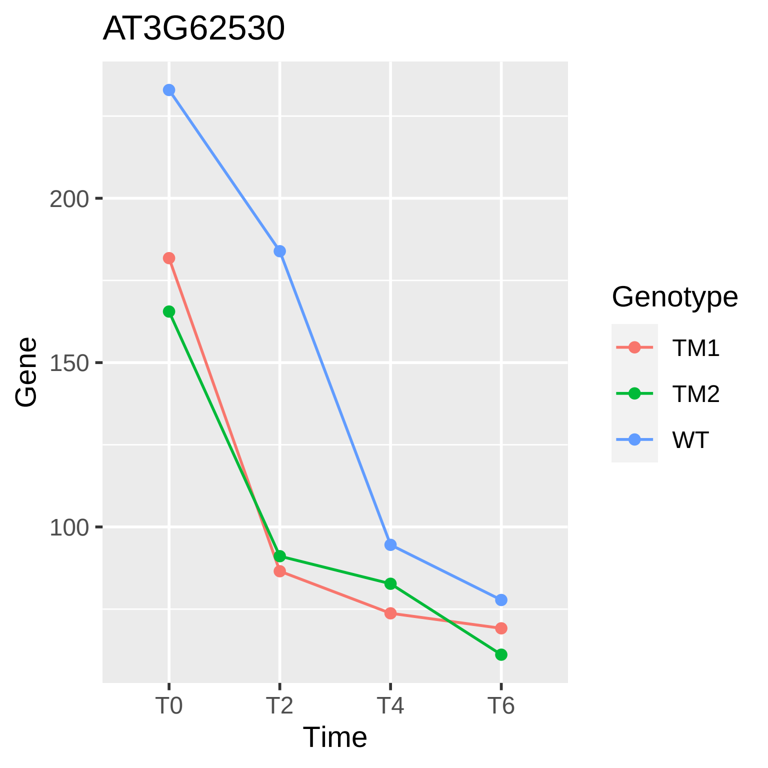
AT3G62530
200
150
100
Gene
T0
T2
T4
T6
Time
Genotype
TM1
TM2
WT

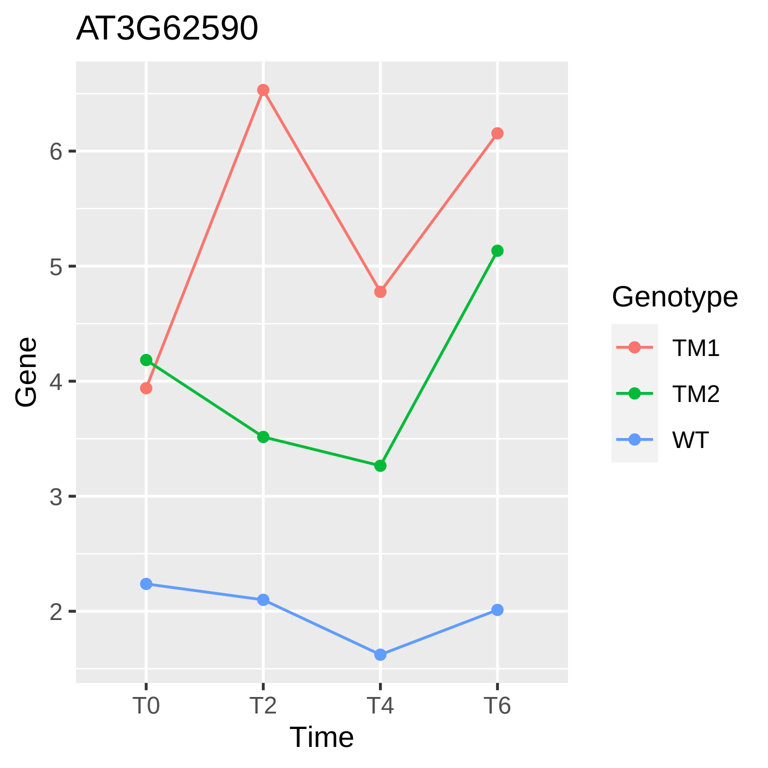
AT3G62590
Gene
6
5
4
3
2
T0
T2
T4
T6
Time
Genotype
TM1
TM2
WT

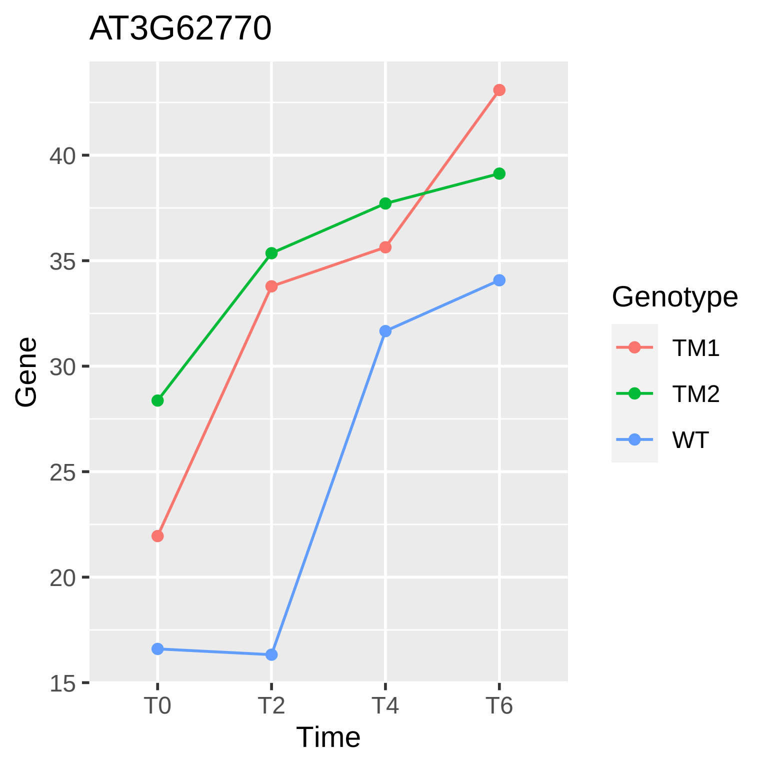

AT3G62770
Gene
40
35
30
25
20
15
T0
T2
T4
T6
Time
Genotype
TM1
TM2
WT

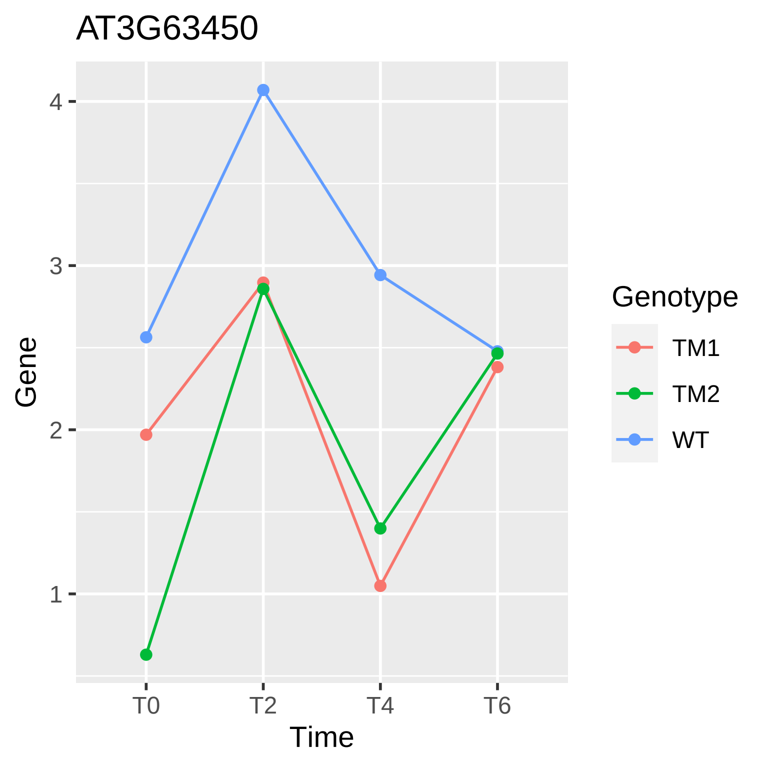

AT3G63450
Gene
4
3
2
1
T0
T2
T4
T6
Time
Genotype
TM1
TM2
WT

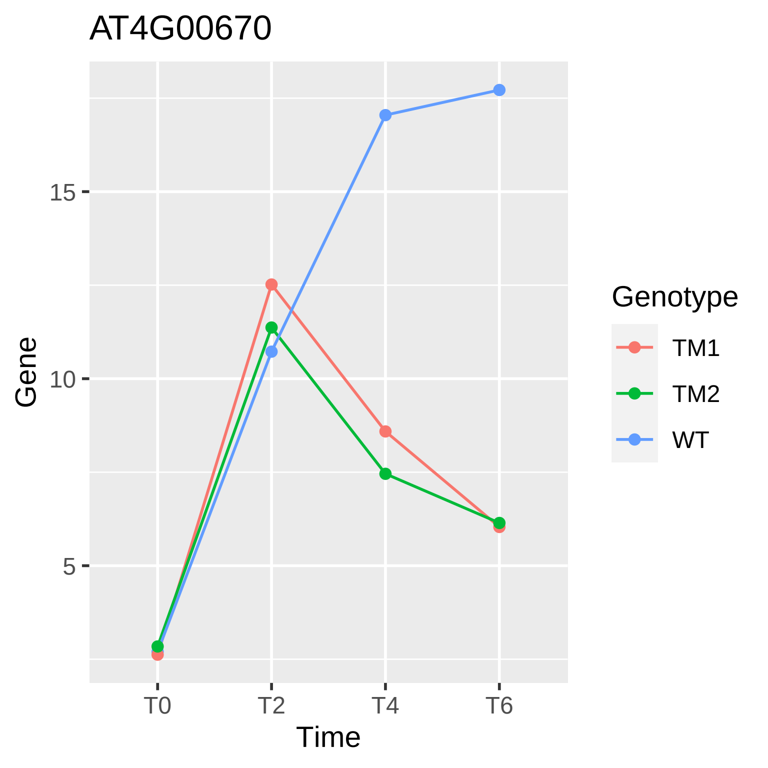
AT4G00670
Gene
15
10
5
T0
T2
T4
T6
Time
Genotype
TM1
TM2
WT

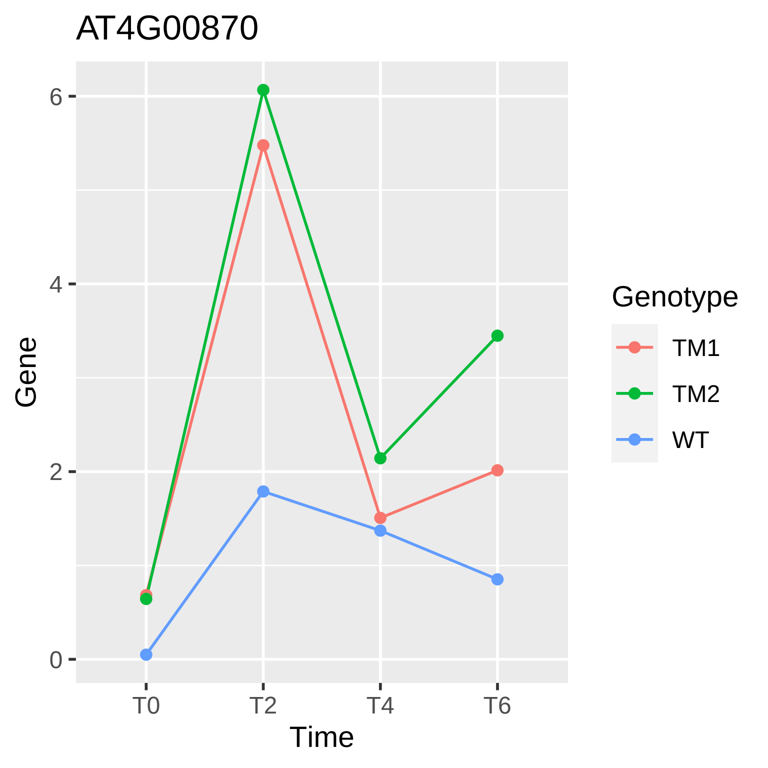

AT4G00870
Gene
6
4
2
0
T0
T2
T4
T6
Time
Genotype
TM1
TM2
WT

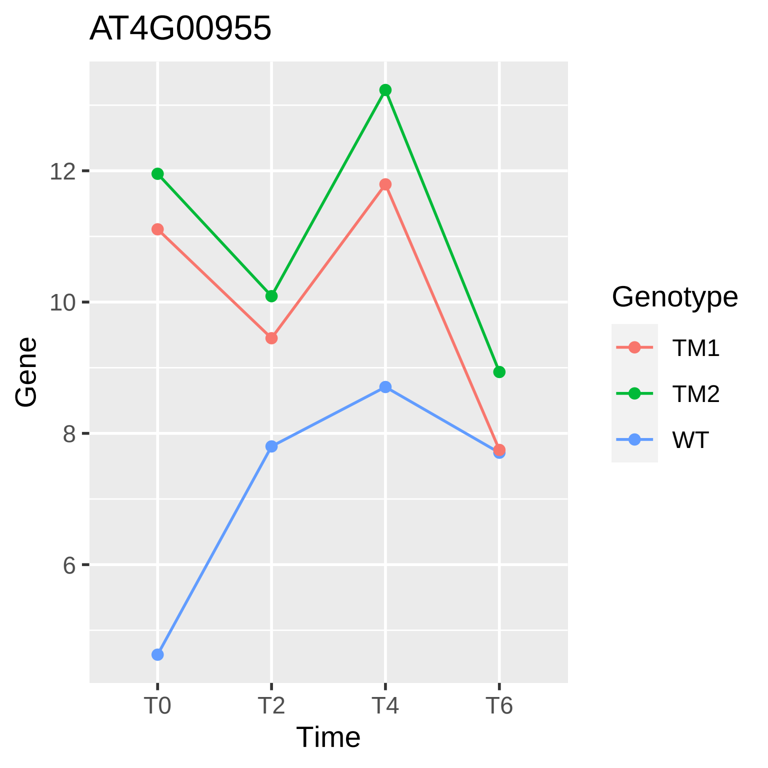
AT4G00955
Gene
Time
T0
T2
T4
T6
6
8
10
12
Genotype
TM1
TM2
WT

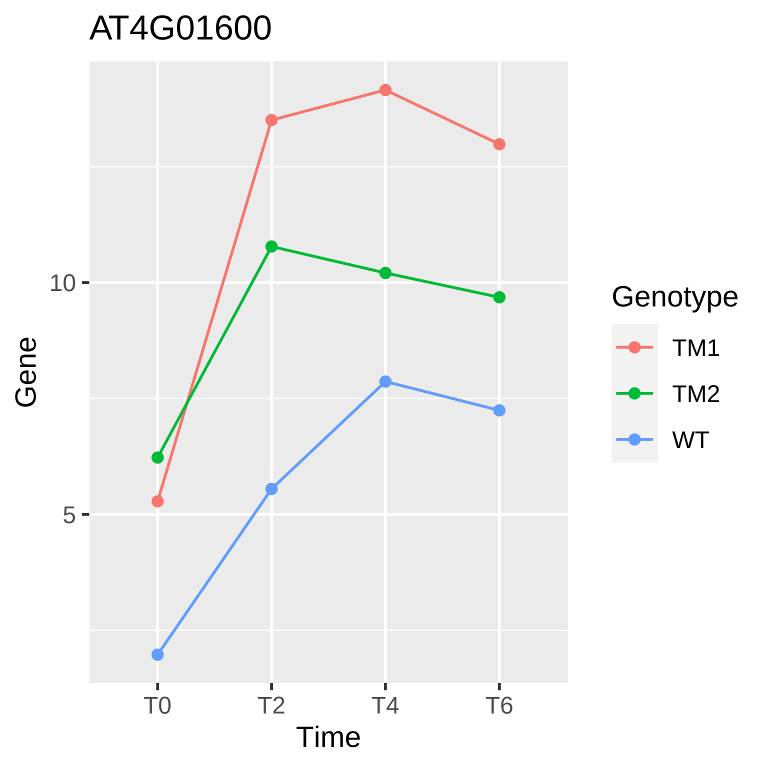
AT4G01600
Gene
10
5
T0
T2
T4
T6
Time
Genotype
TM1
TM2
WT

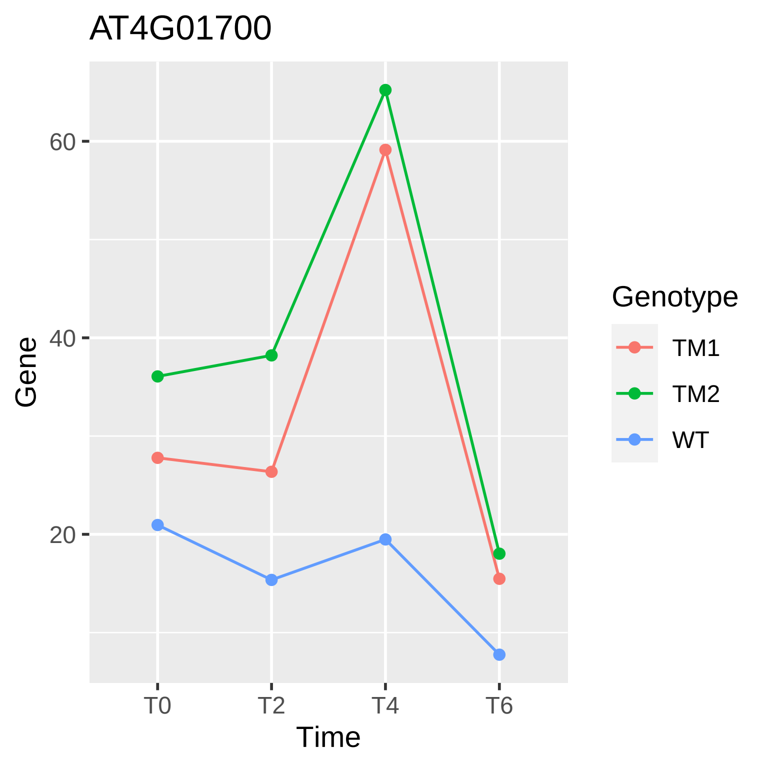

AT4G01700
60
40
20
Gene
T0
T2
T4
T6
Time
Genotype
TM1
TM2
WT

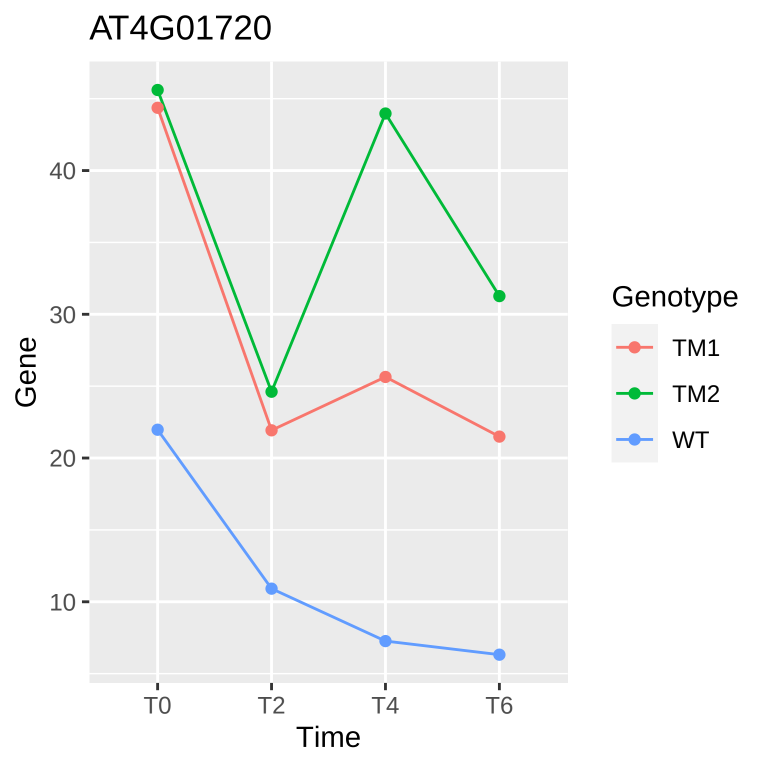

AT4G01720
Gene
40
30
20
10
T0
T2
T4
T6
Time
Genotype
TM1
TM2
WT

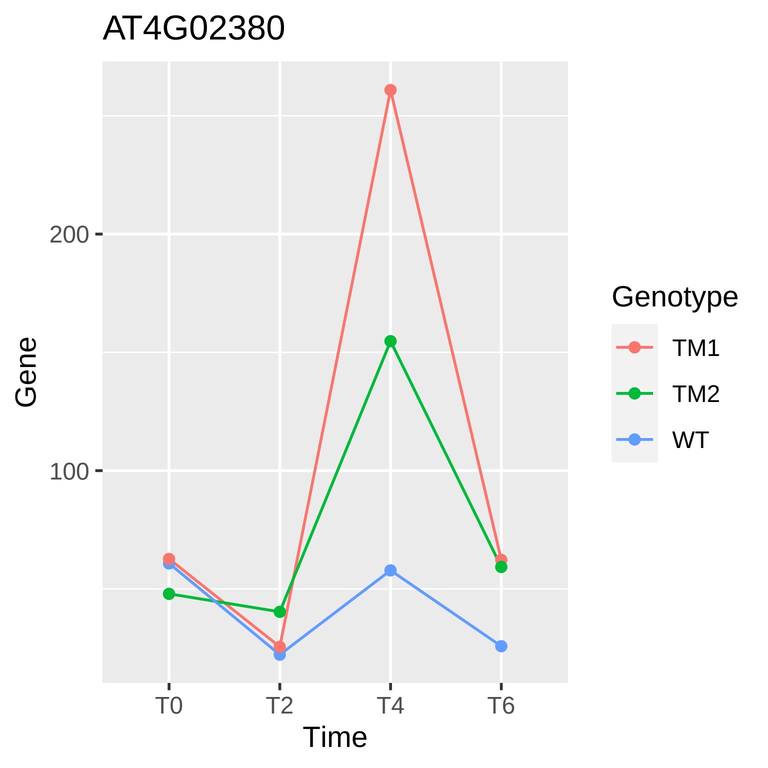
AT4G02380
Gene
200
100
T0
T2
T4
T6
Time
Genotype
TM1
TM2
WT

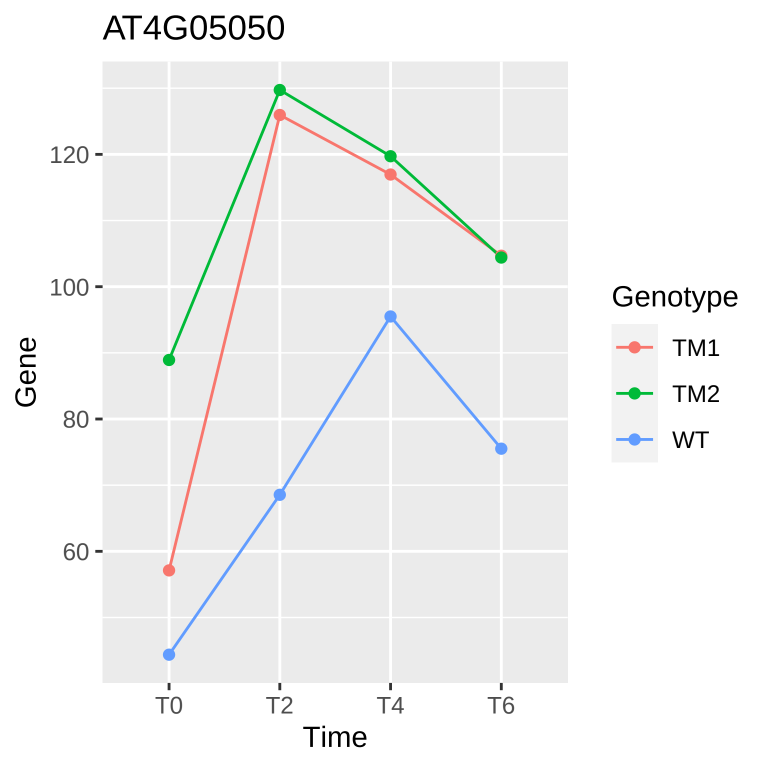
AT4G05050
Gene
120
100
80
60
T0
T2
T4
T6
Time
Genotype
TM1
TM2
WT

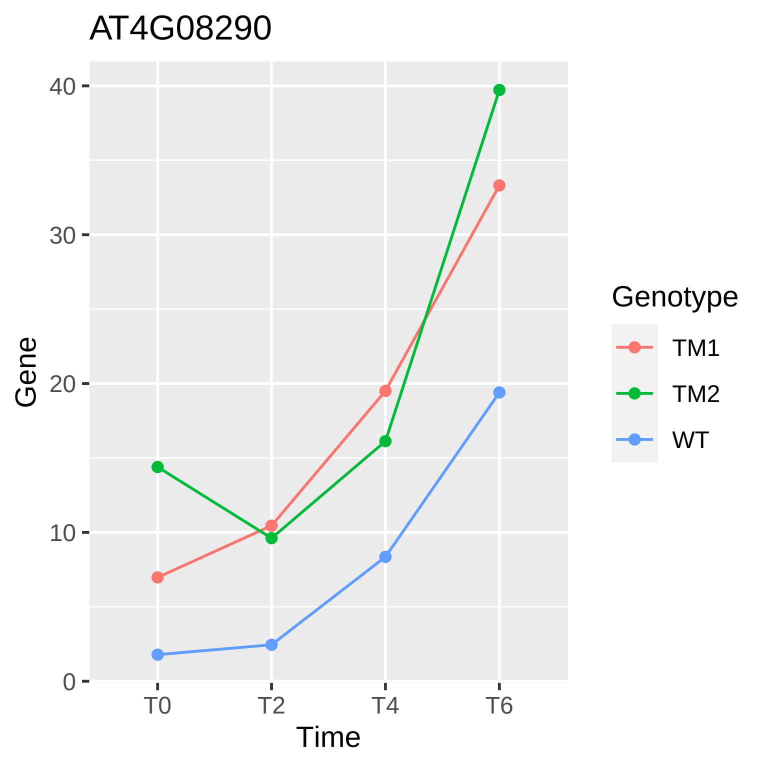
AT4G08290
Gene
40
30
20
10
0
T0
T2
T4
T6
Time
Genotype
TM1
TM2
WT

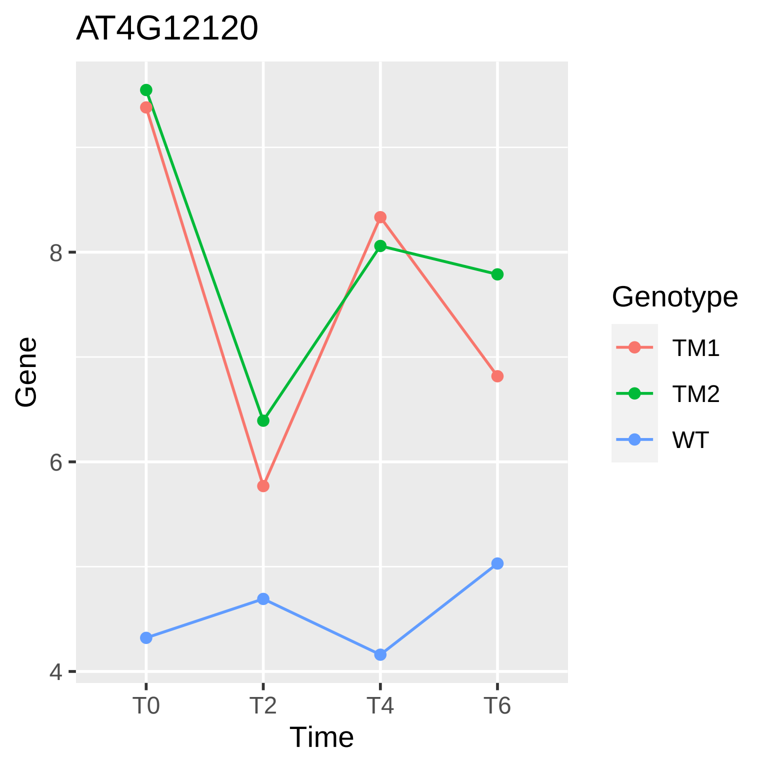
AT4G12120
Gene
8
6
4
T0
T2
T4
T6
Time
Genotype
TM1
TM2
WT

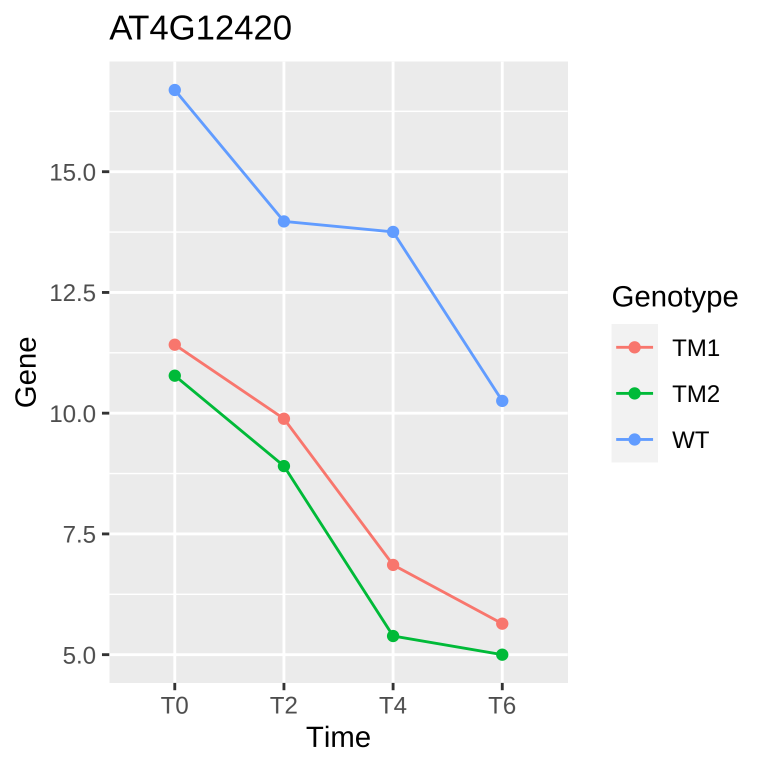

AT4G12420
Gene
15.0
12.5
10.0
7.5
5.0
T0
T2
T4
T6
Time
Genotype
TM1
TM2
WT

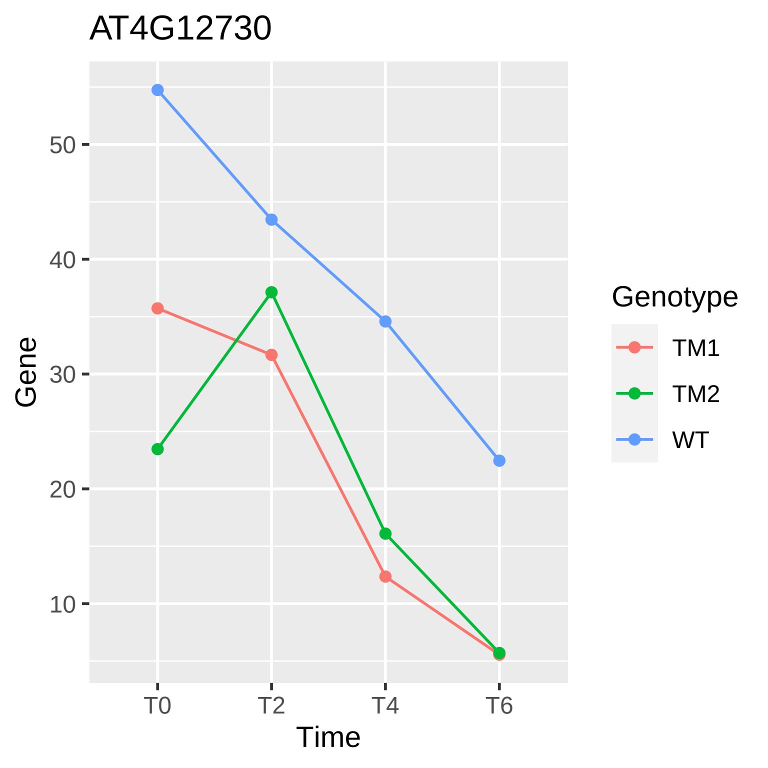
AT4G12730
Gene
50
40
30
20
10
T0
T2
T4
T6
Time
Genotype
TM1
TM2
WT

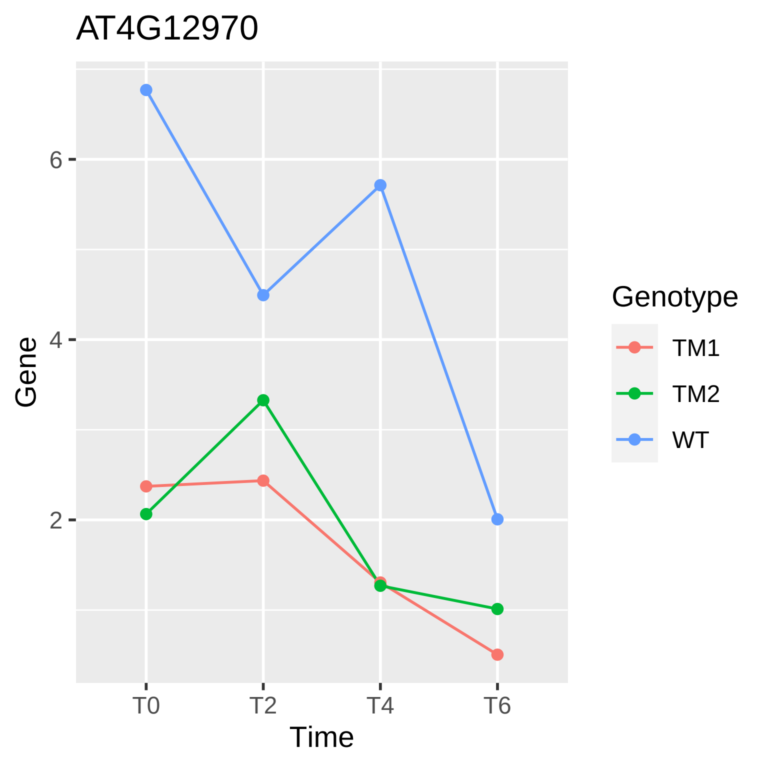

AT4G12970
Gene
6
4
2
T0
T2
T4
T6
Time
Genotype
TM1
TM2
WT

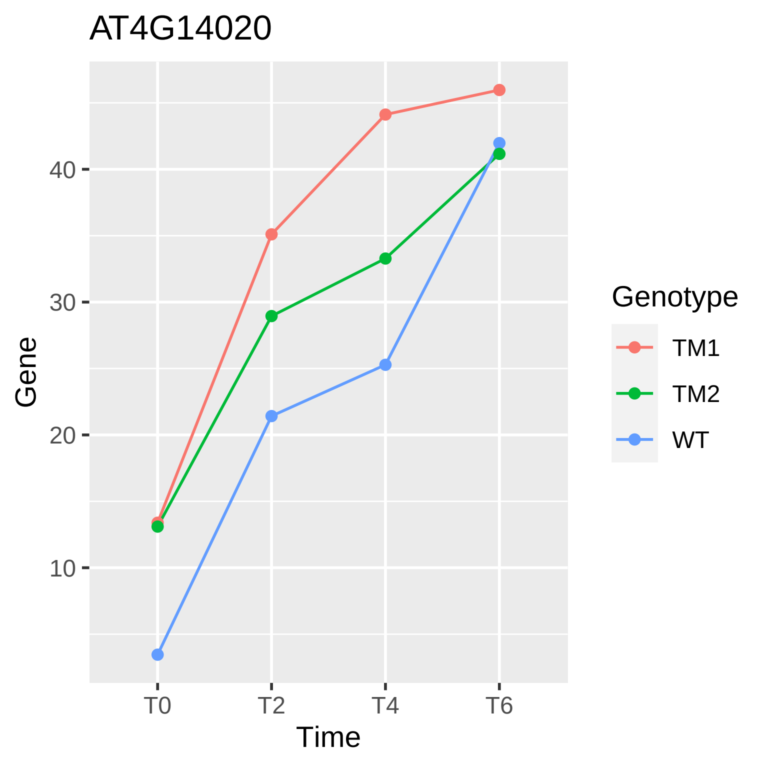
AT4G14020
Gene
10
20
30
40
T0
T2
T4
T6
Time
Genotype
TM1
TM2
WT

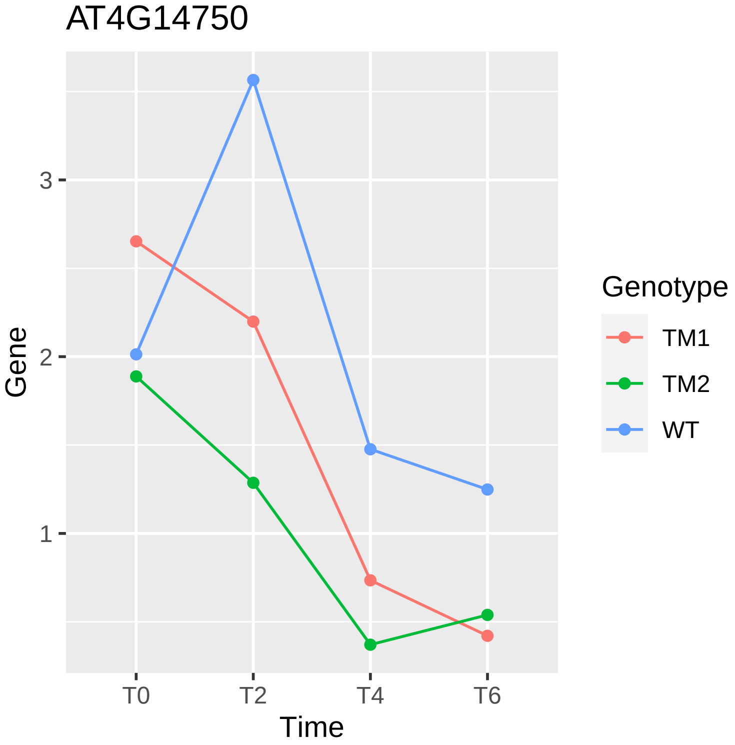
AT4G14750
Gene
3
2
1
T0
T2
T4
T6
Time
Genotype
TM1
TM2
WT

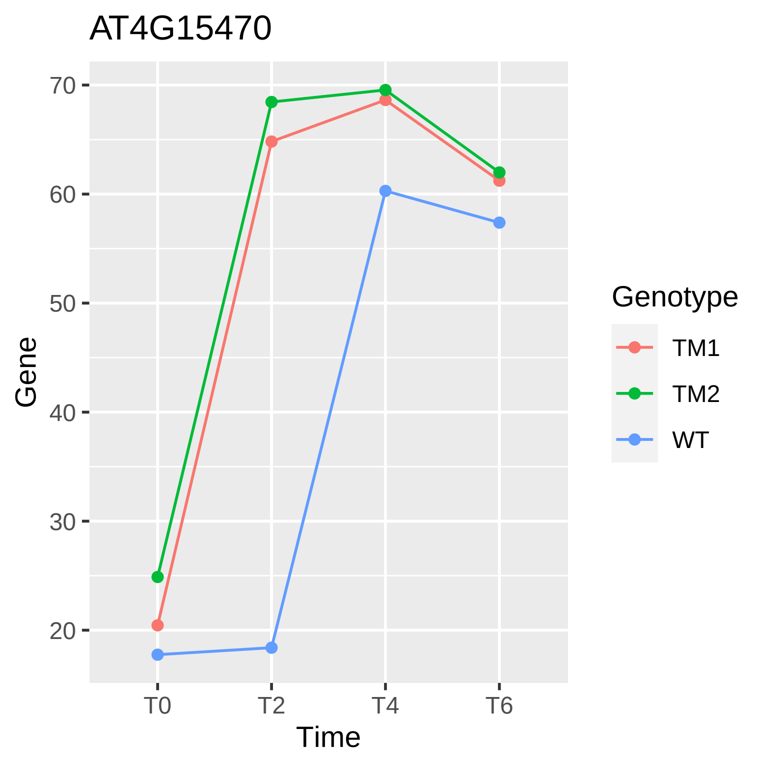

AT4G15470
70
60
50
40
30
20
Gene
T0
T2
T4
T6
Time
Genotype
TM1
TM2
WT

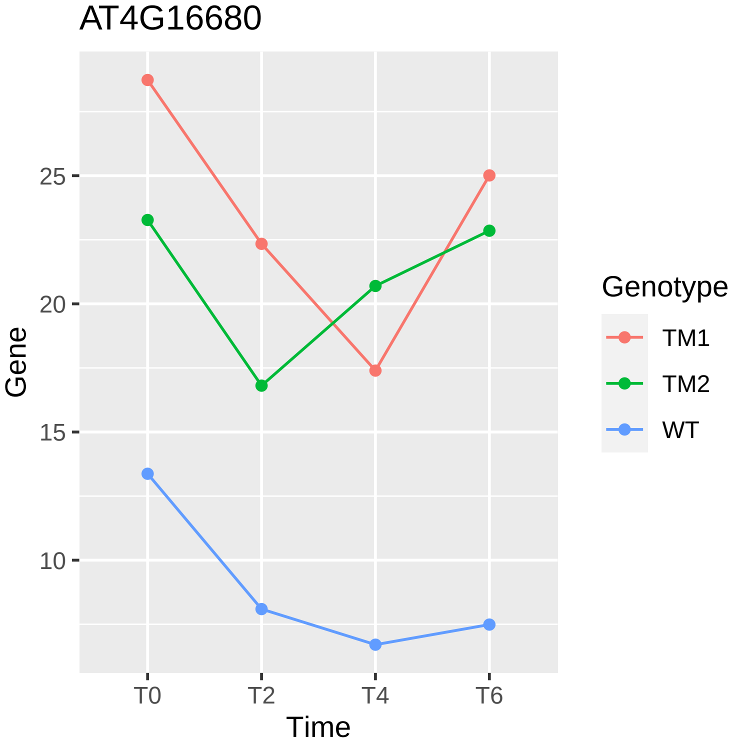
AT4G16680
Gene
25
20
15
10
T0
T2
T4
T6
Time
Genotype
TM1
TM2
WT

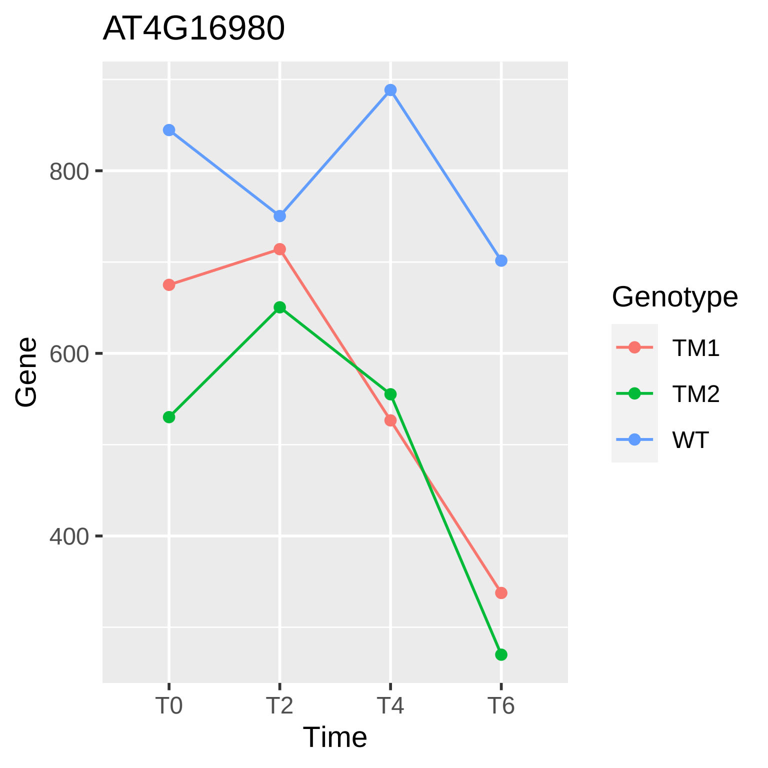

AT4G16980
Gene
800
600
400
T0
T2
T4
T6
Time
Genotype
TM1
TM2
WT

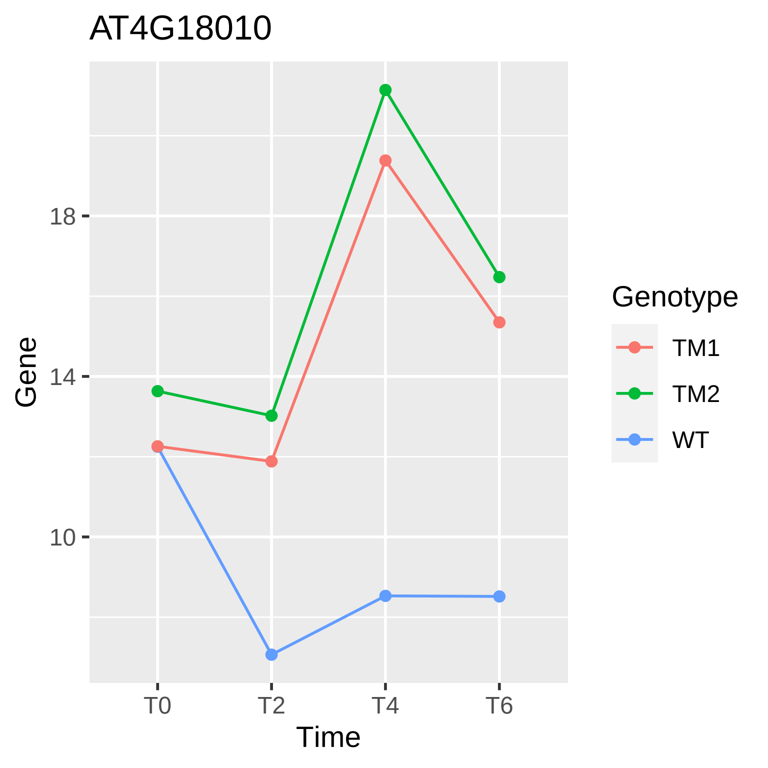

AT4G18010
18
14
10
Gene
T0
T2
T4
T6
Time
Genotype
TM1
TM2
WT

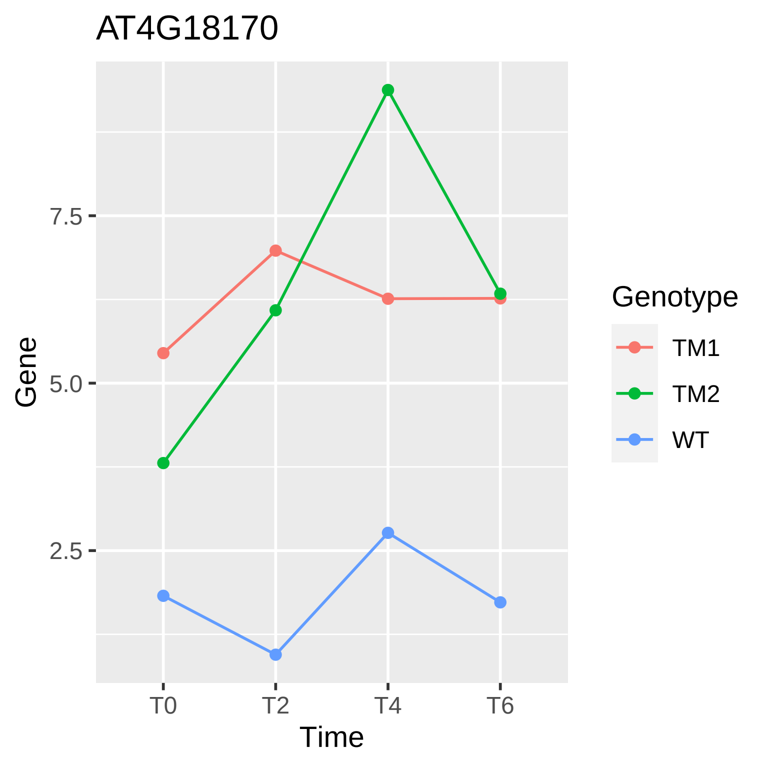

AT4G18170
Gene
7.5
5.0
2.5
T0
T2
T4
T6
Time
Genotype
TM1
TM2
WT

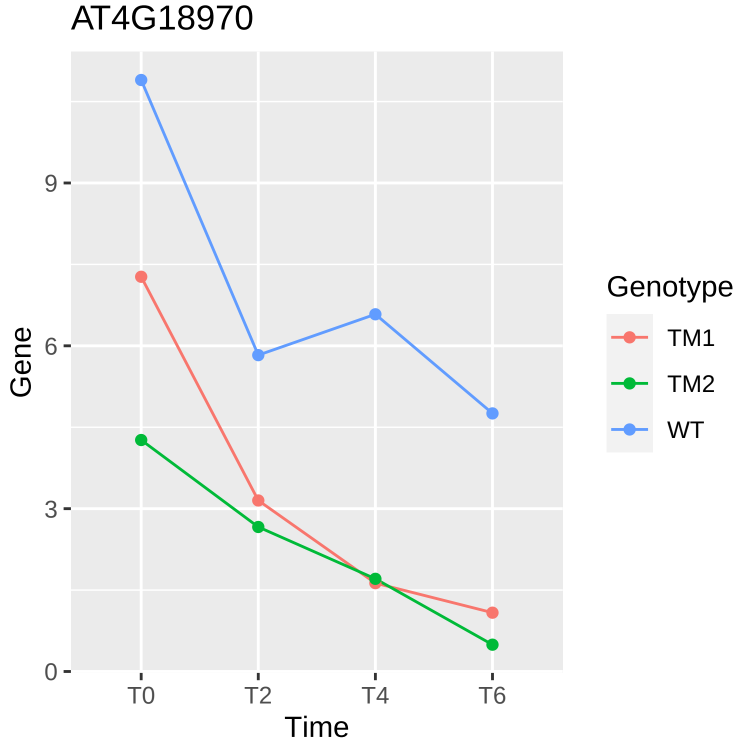
AT4G18970
Gene
9
6
3
0
T0
T2
T4
T6
Time
Genotype
TM1
TM2
WT

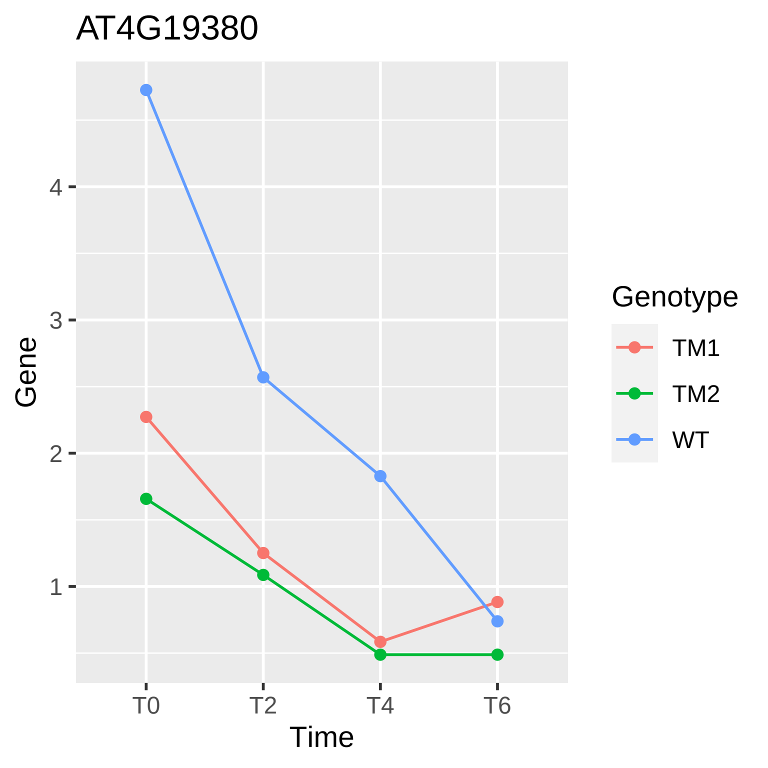

AT4G19380
Gene
4
3
2
1
T0
T2
T4
T6
Time
Genotype
TM1
TM2
WT

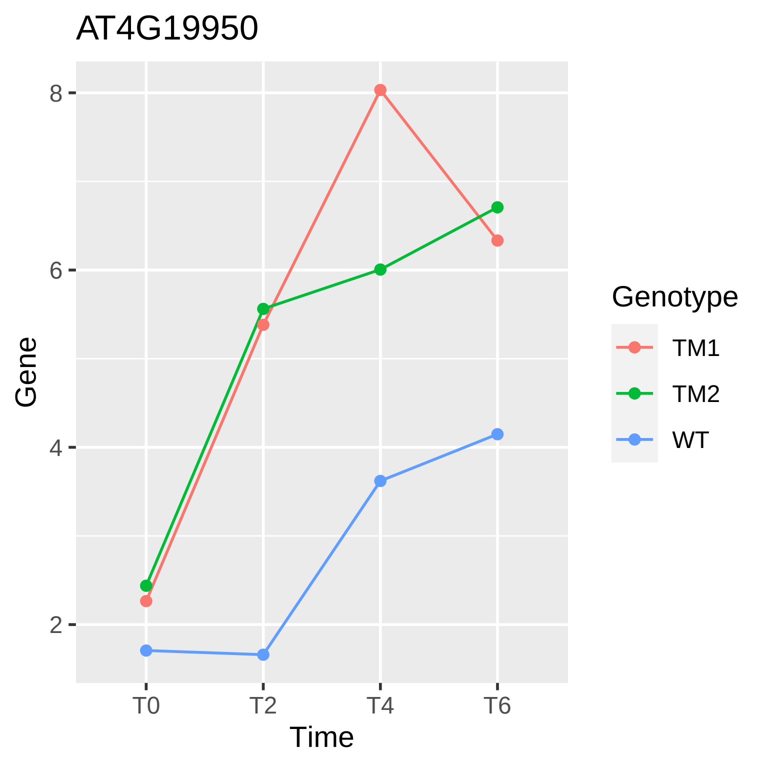
AT4G19950
8
6
4
2
Gene
T0
T2
T4
T6
Time
Genotype
TM1
TM2
WT

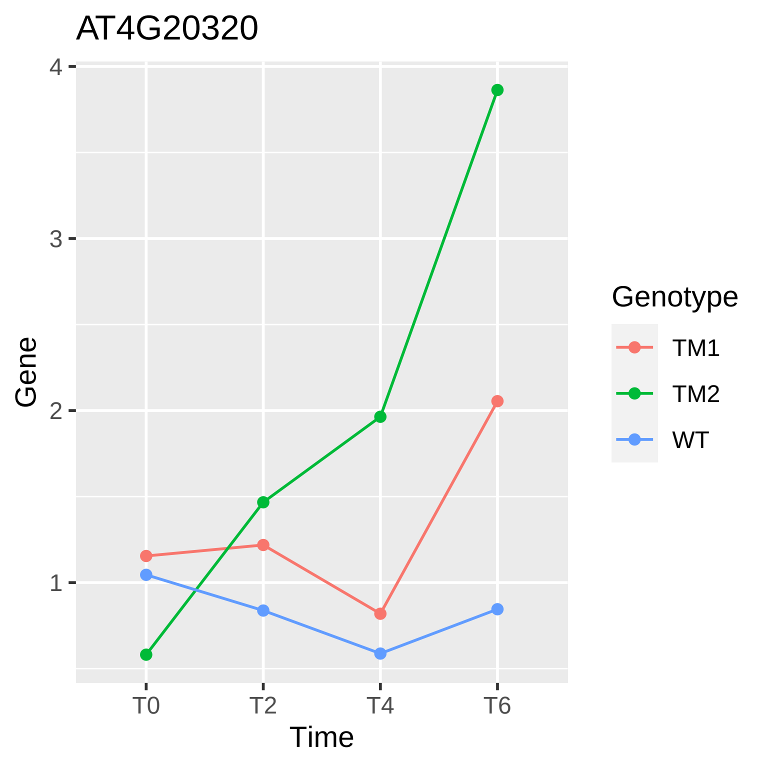
AT4G20320
Gene
4
3
2
1
T0
T2
T4
T6
Time
Genotype
TM1
TM2
WT

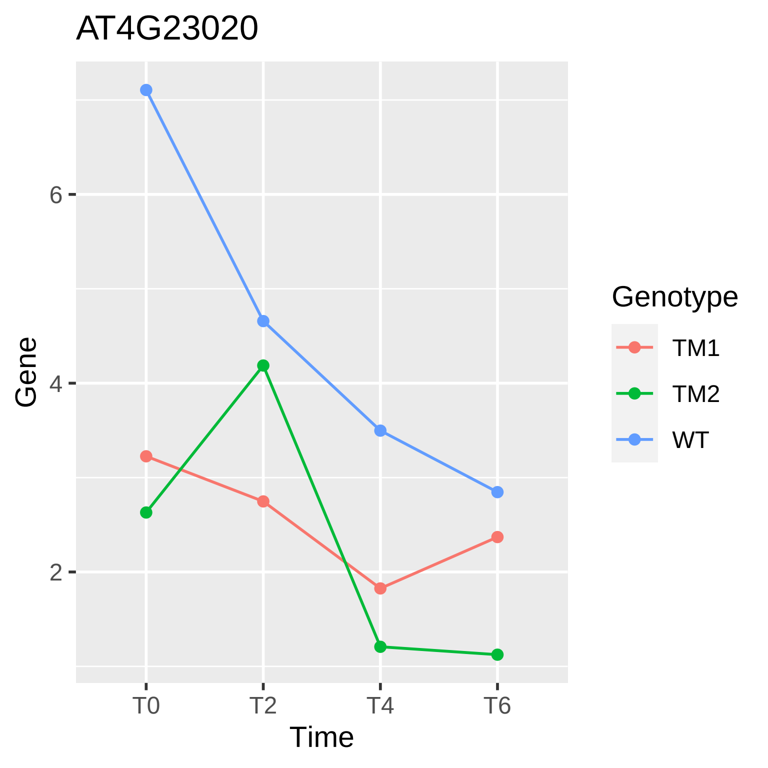

AT4G23020
Gene
6
4
2
T0
T2
T4
T6
Time
Genotype
TM1
TM2
WT

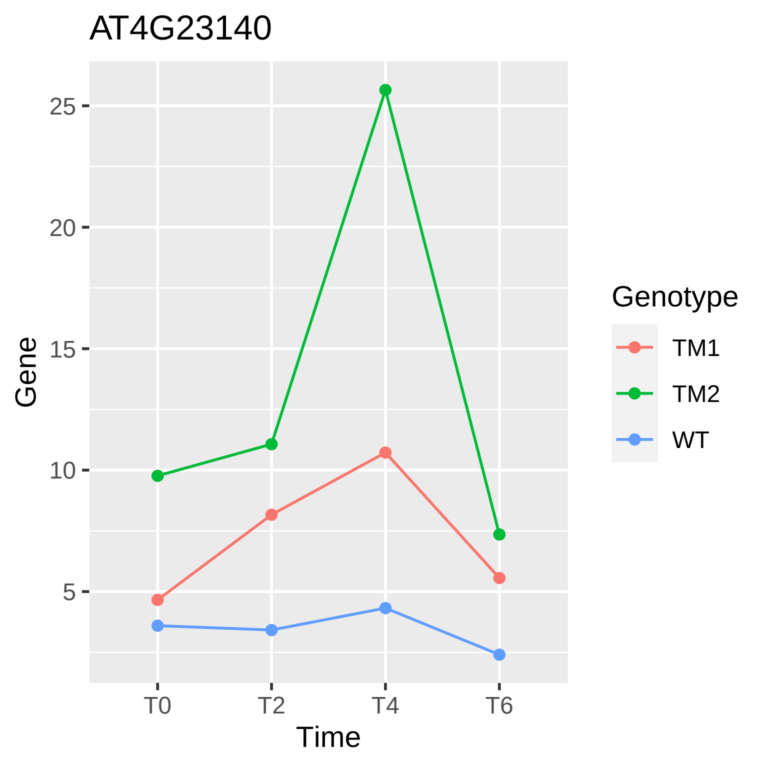

AT4G23140
Gene
25
20
15
10
5
T0
T2
T4
T6
Time
Genotype
TM1
TM2
WT

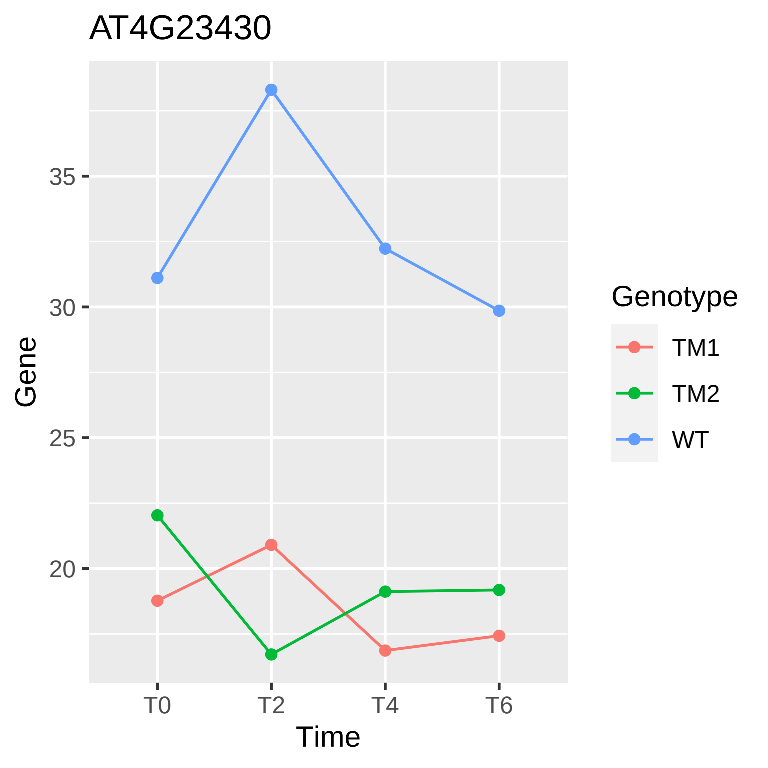
AT4G23430
35
30
25
20
Gene
T0
T2
T4
T6
Time
Genotype
TM1
TM2
WT

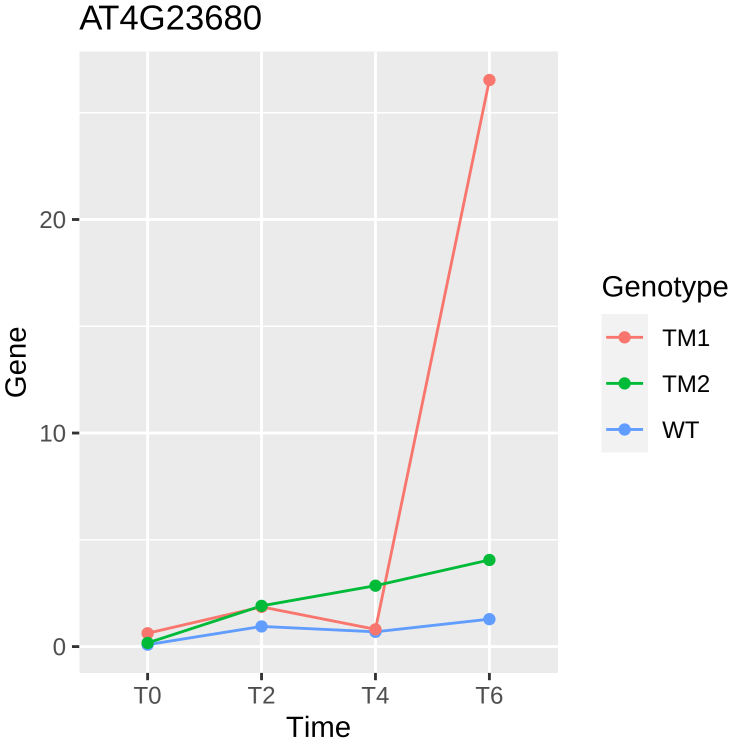

AT4G23680
Gene
20
10
0
T0
T2
T4
T6
Time
Genotype
TM1
TM2
WT

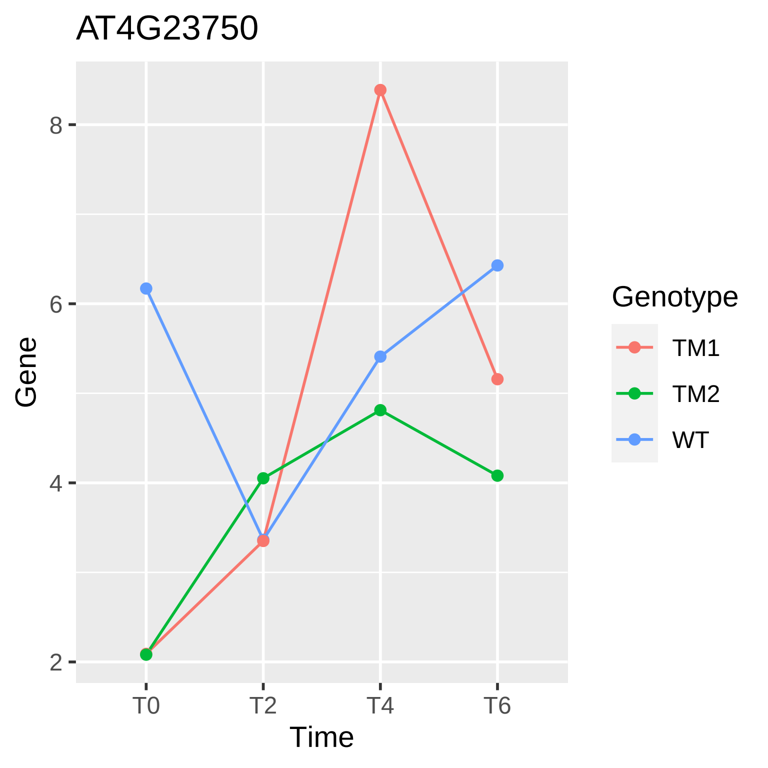

AT4G23750
Gene
8
6
4
2
T0
T2
T4
T6
Time
Genotype
TM1
TM2
WT

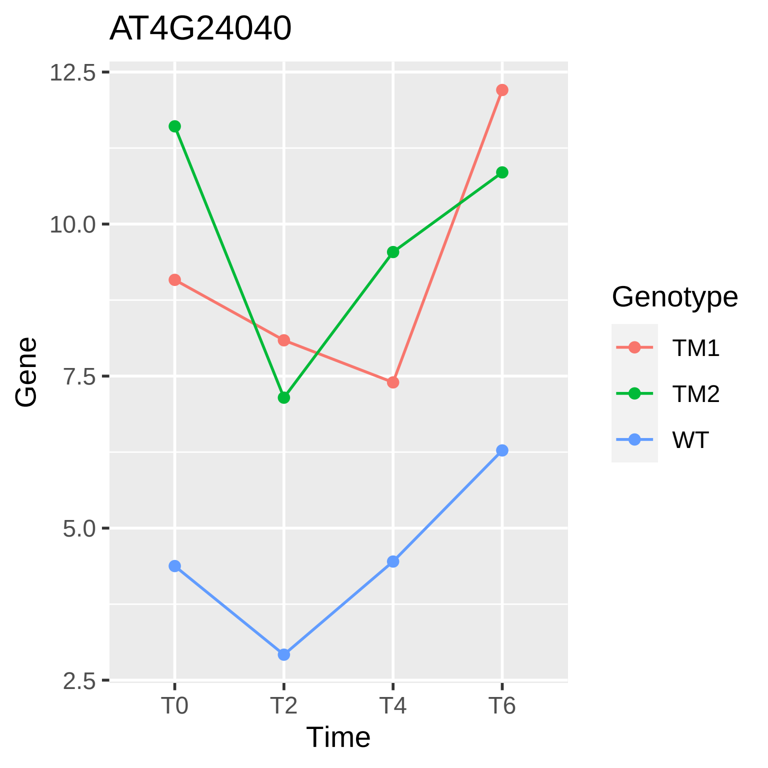
AT4G24040
12.5
10.0
7.5
5.0
2.5
Gene
T0
T2
T4
T6
Time
Genotype
TM1
TM2
WT

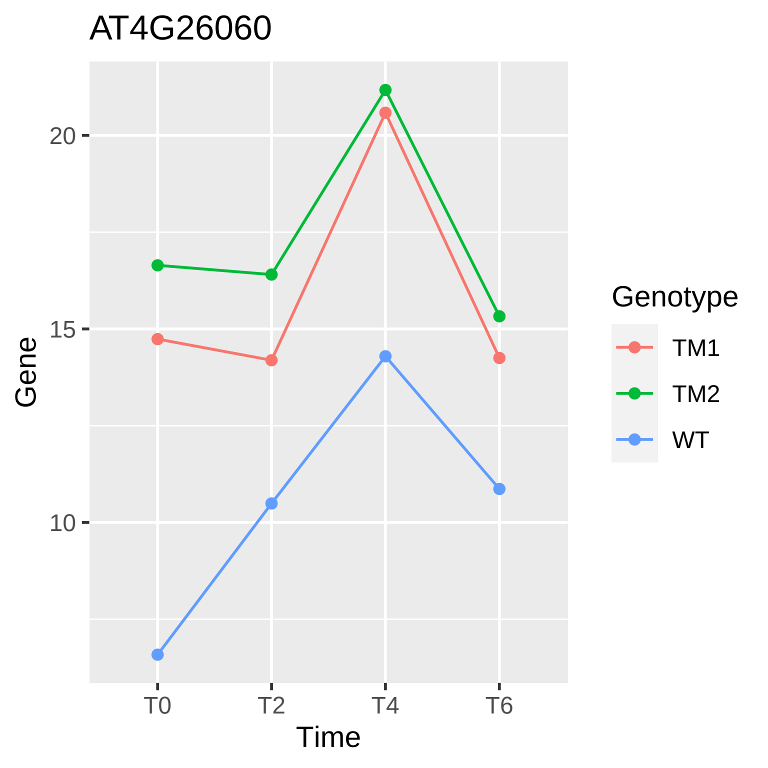
AT4G26060
Gene
20
15
10
T0
T2
T4
T6
Time
Genotype
TM1
TM2
WT

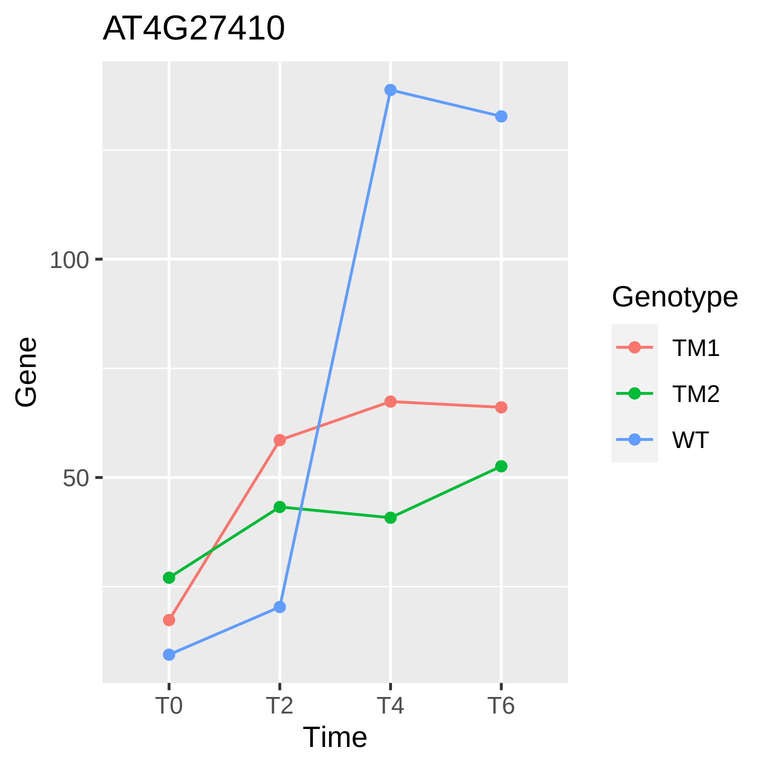
AT4G27410
Gene
100
50
T0
T2
T4
T6
Time
Genotype
TM1
TM2
WT

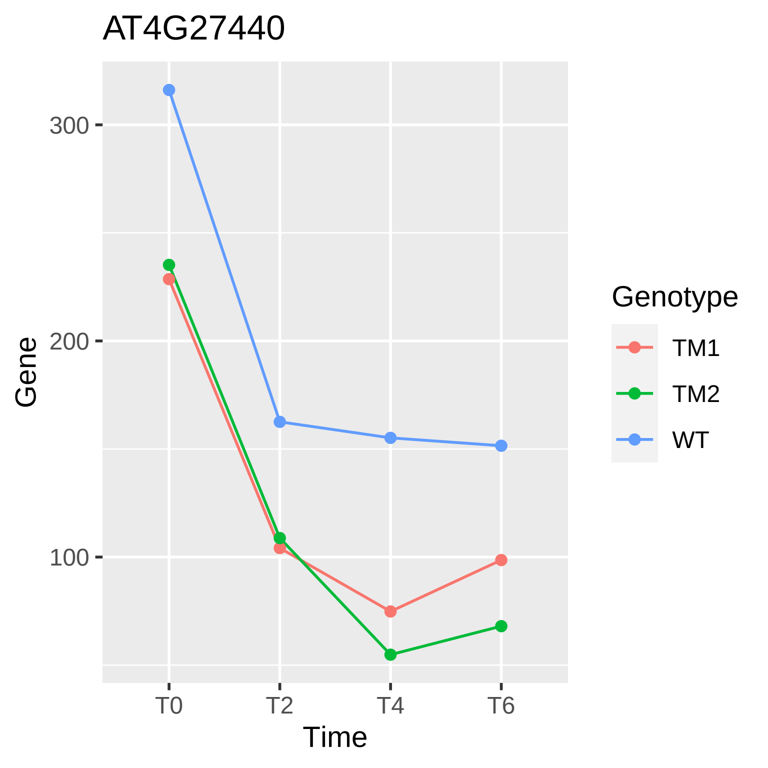
AT4G27440
300
200
100
Gene
T0
T2
T4
T6
Time
Genotype
TM1
TM2
WT

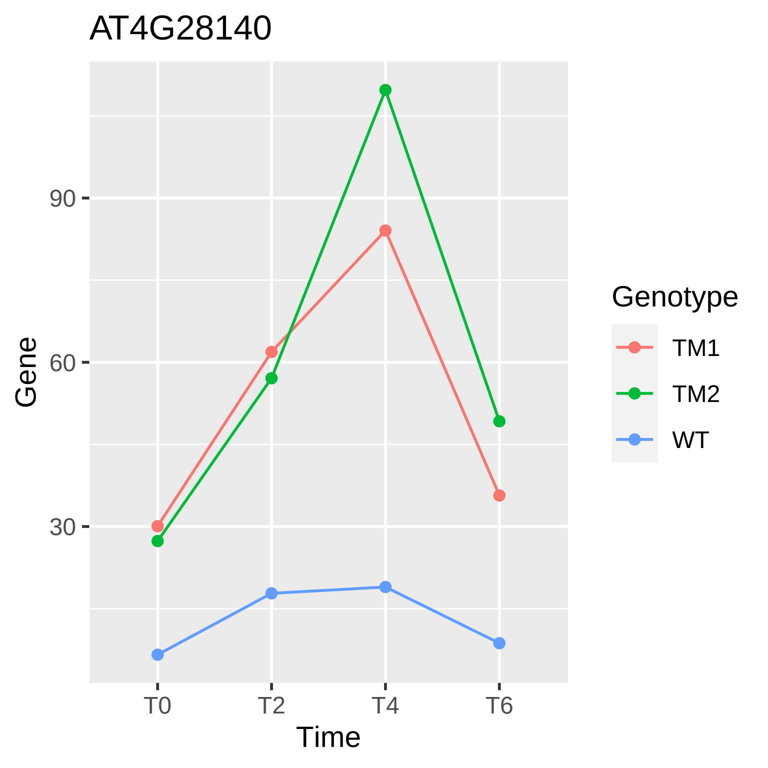
AT4G28140
Gene
90
60
30
T0
T2
T4
T6
Time
Genotype
TM1
TM2
WT

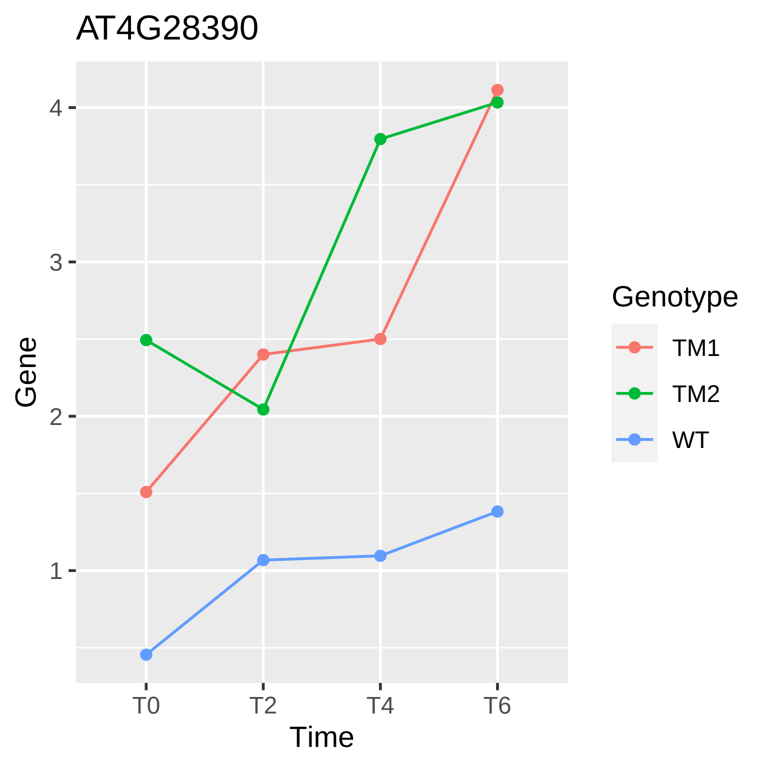
AT4G28390
Gene
4
3
2
1
T0
T2
T4
T6
Time
Genotype
TM1
TM2
WT

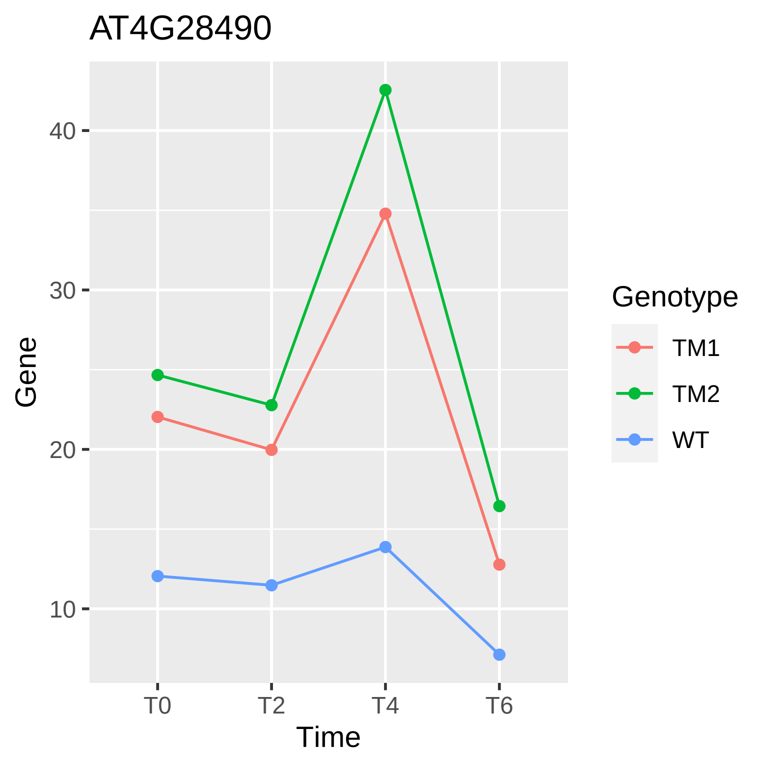
AT4G28490
Gene
40
30
20
10
T0
T2
T4
T6
Time
Genotype
TM1
TM2
WT

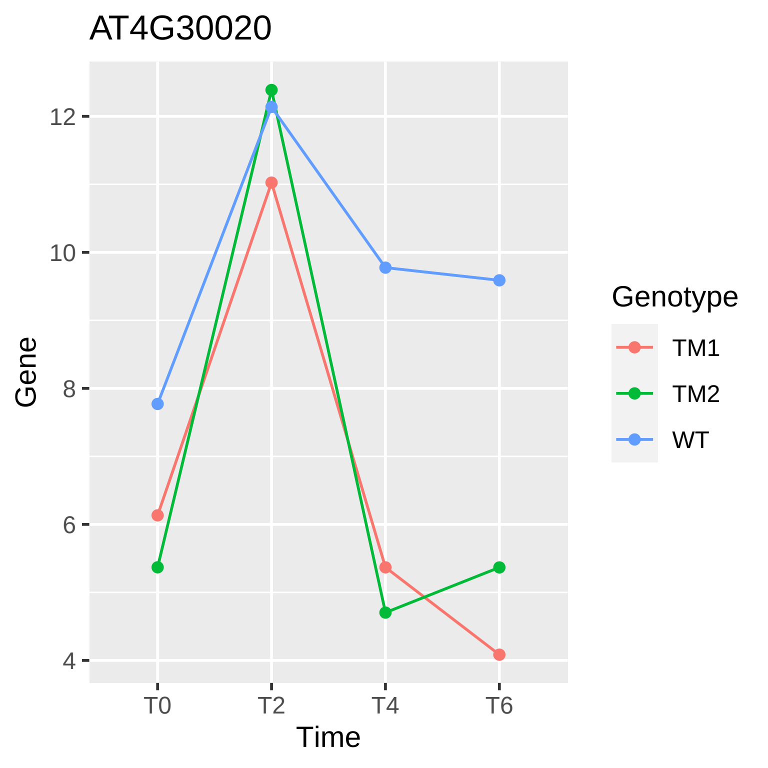

AT4G30020
Gene
Time
12
10
8
6
4
T0
T2
T4
T6
Genotype
TM1
TM2
WT

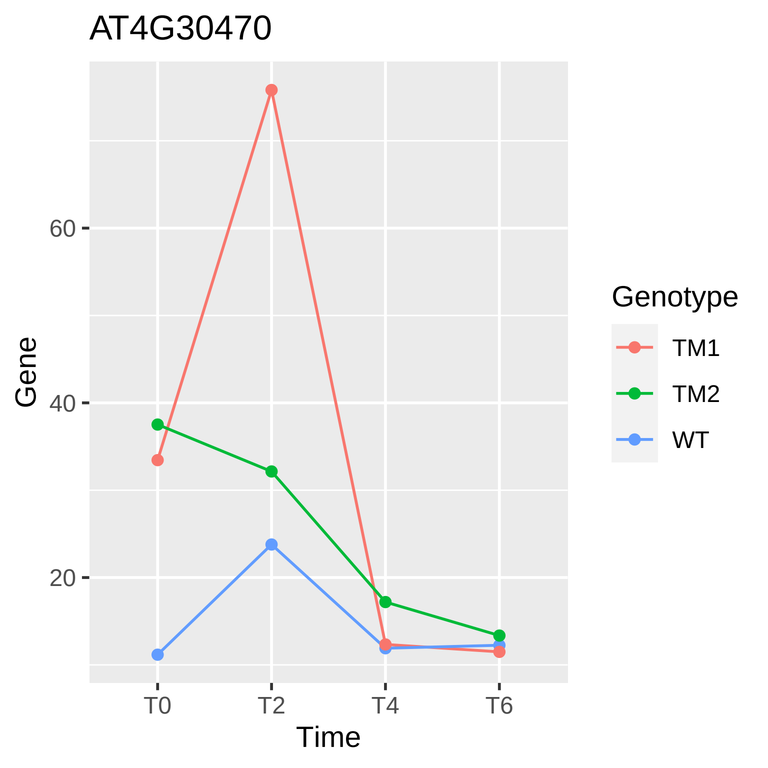
AT4G30470
Gene
60
40
20
T0
T2
T4
T6
Time
Genotype
TM1
TM2
WT

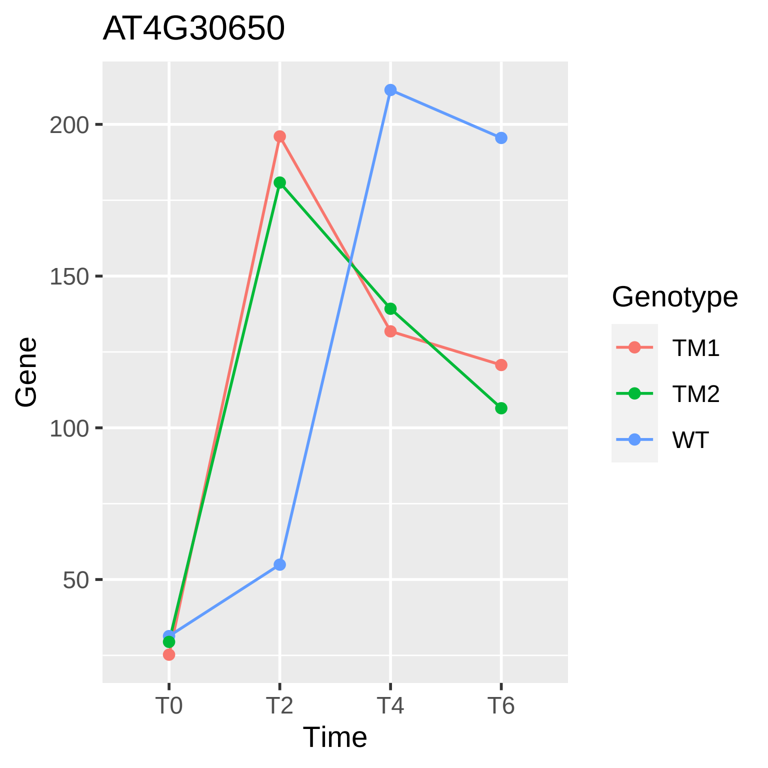

AT4G30650
Gene
200
150
100
50
T0
T2
T4
T6
Time
Genotype
TM1
TM2
WT

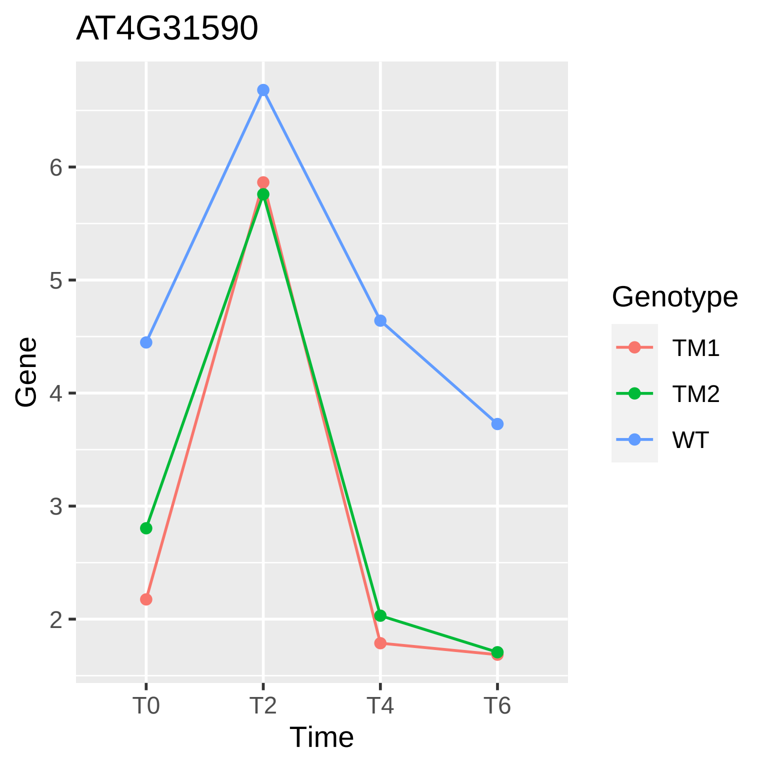
AT4G31590
Gene
6
5
4
3
2
T0
T2
T4
T6
Time
Genotype
TM1
TM2
WT

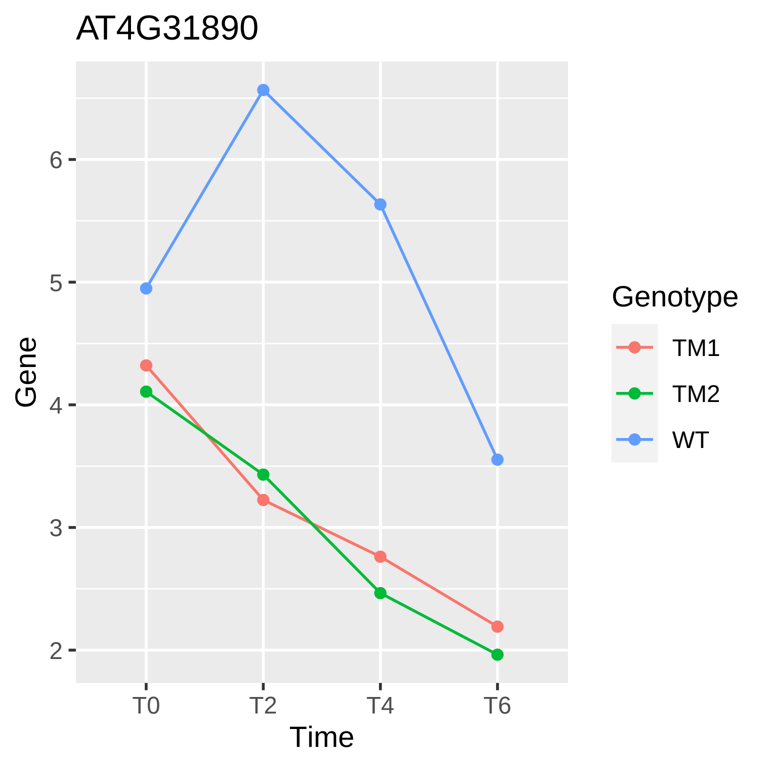

AT4G31890
Gene
Time
T0
T2
T4
T6
2
3
4
5
6
Genotype
TM1
TM2
WT

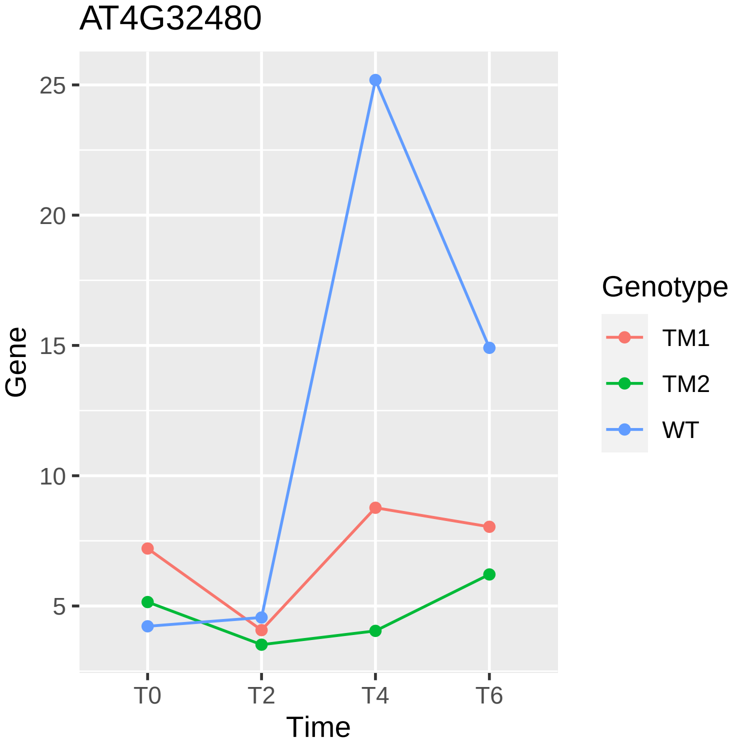
AT4G32480
Gene
25
20
15
10
5
T0
T2
T4
T6
Time
Genotype
TM1
TM2
WT

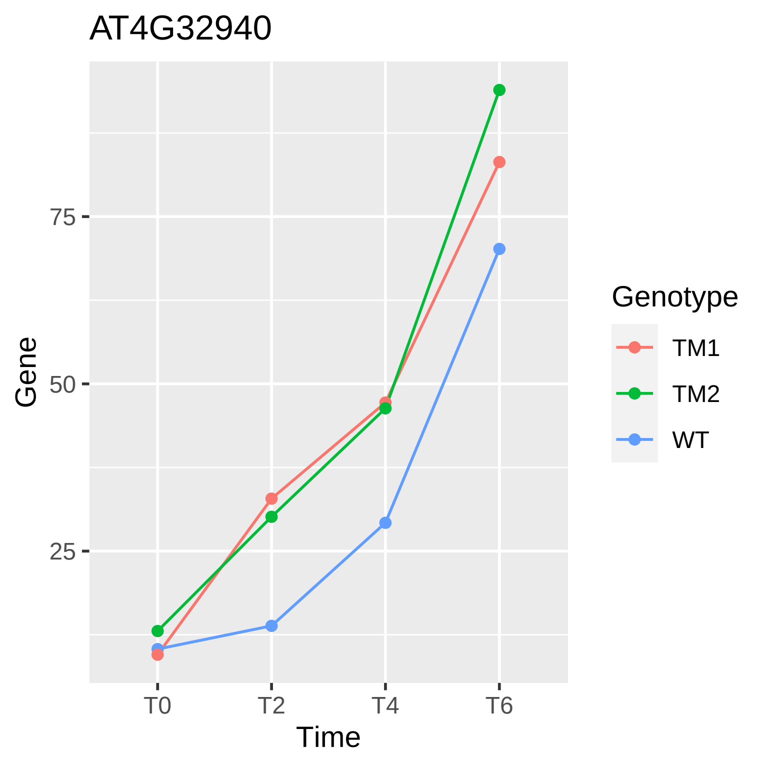
AT4G32940
Gene
75
50
25
T0
T2
T4
T6
Time
Genotype
TM1
TM2
WT

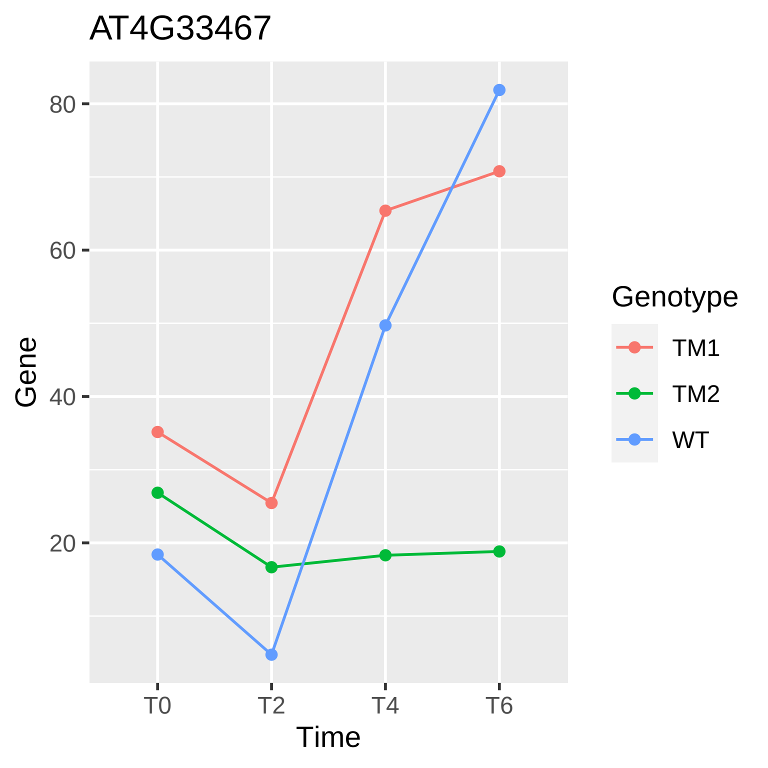

AT4G33467
80
60
40
20
Gene
T0
T2
T4
T6
Time
Genotype
TM1
TM2
WT

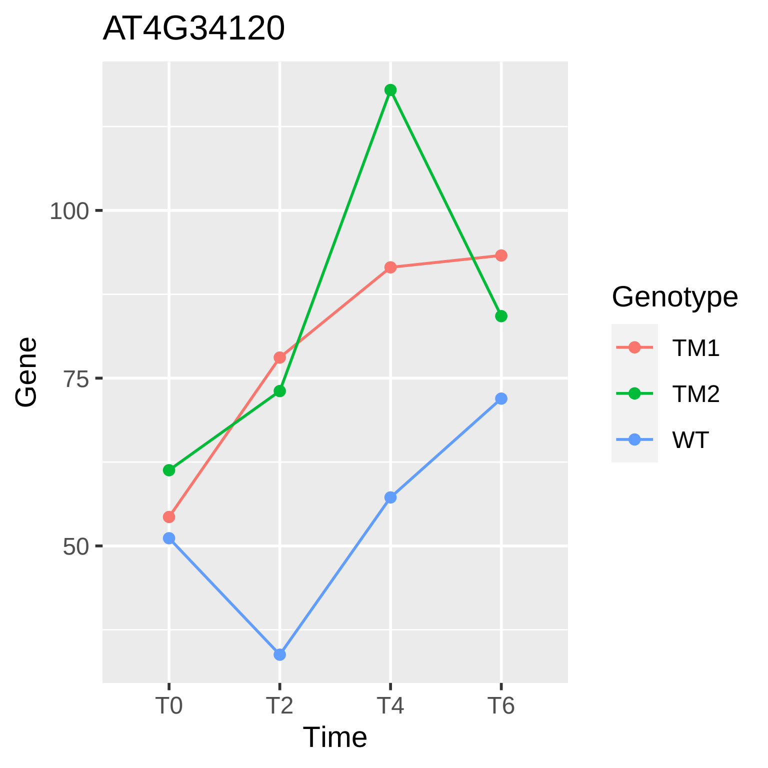
AT4G34120
Gene
100
75
50
T0
T2
T4
T6
Time
Genotype
TM1
TM2
WT

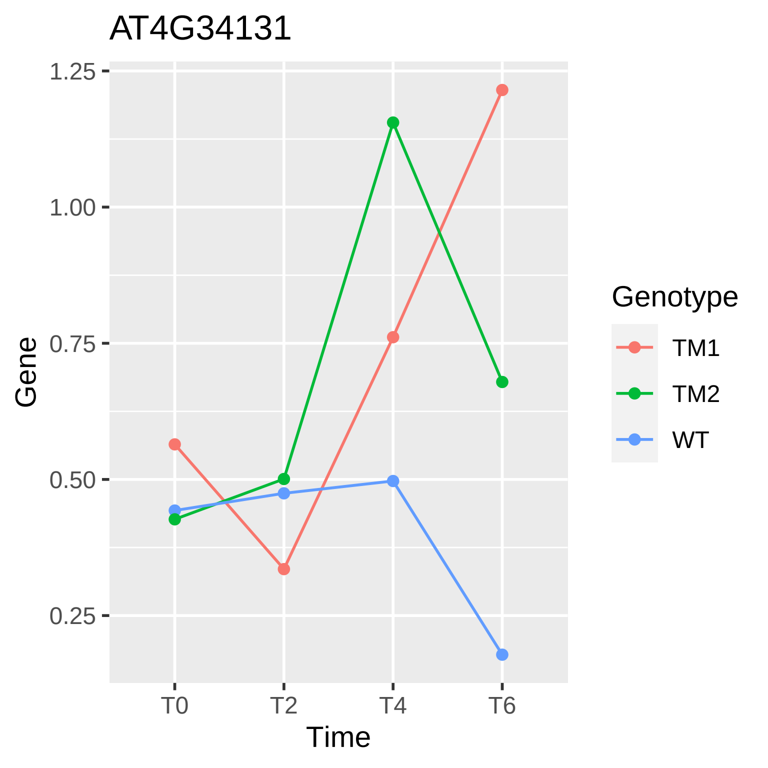
AT4G34131
Gene
1.25
1.00
0.75
0.50
0.25
T0
T2
T4
T6
Time
Genotype
TM1
TM2
WT

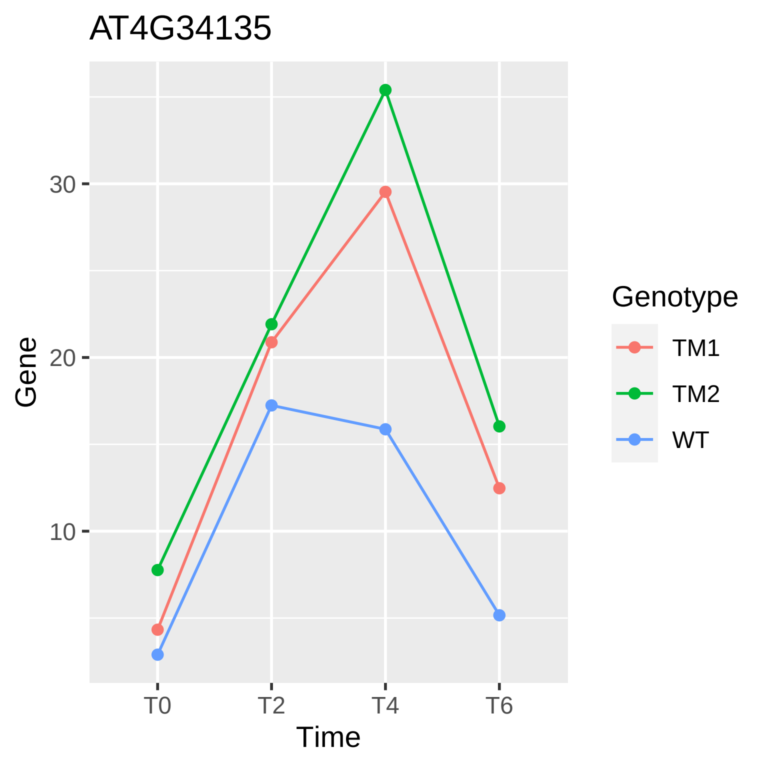
AT4G34135
Gene
30
20
10
T0
T2
T4
T6
Time
Genotype
TM1
TM2
WT

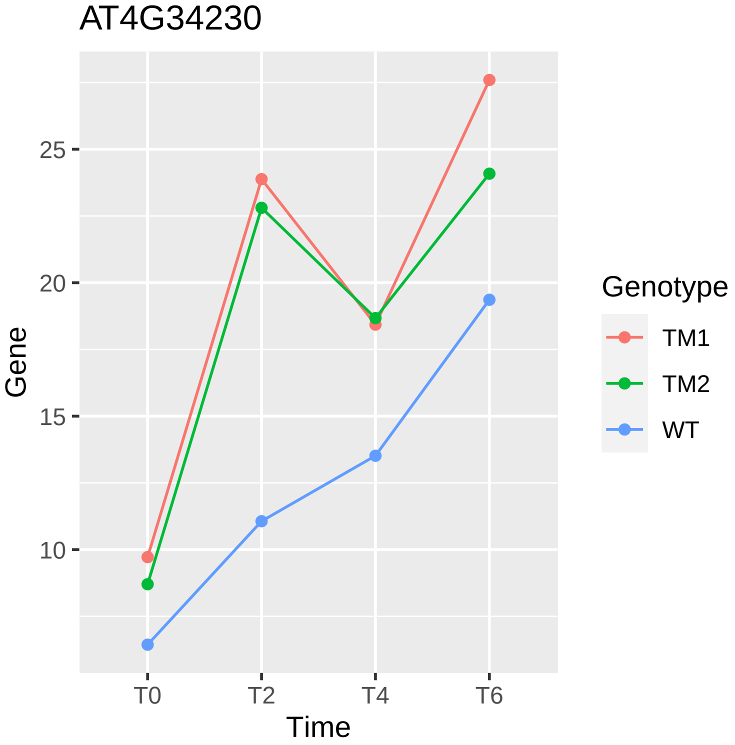

AT4G34230
Gene
25
20
15
10
T0
T2
T4
T6
Time
Genotype
TM1
TM2
WT

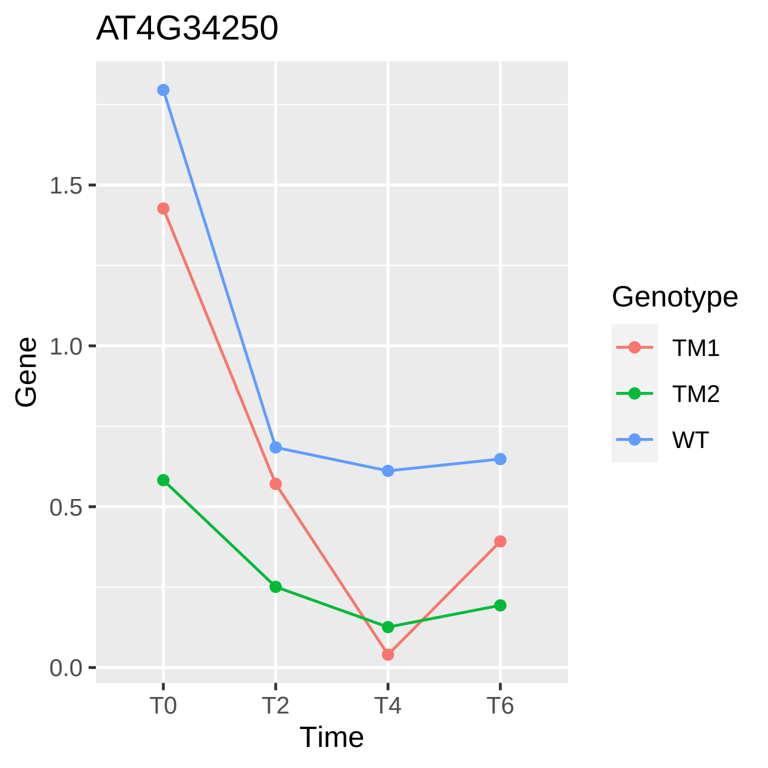

AT4G34250
Gene
0.0
0.5
1.0
1.5
T0
T2
T4
T6
Time
Genotype
TM1
TM2
WT

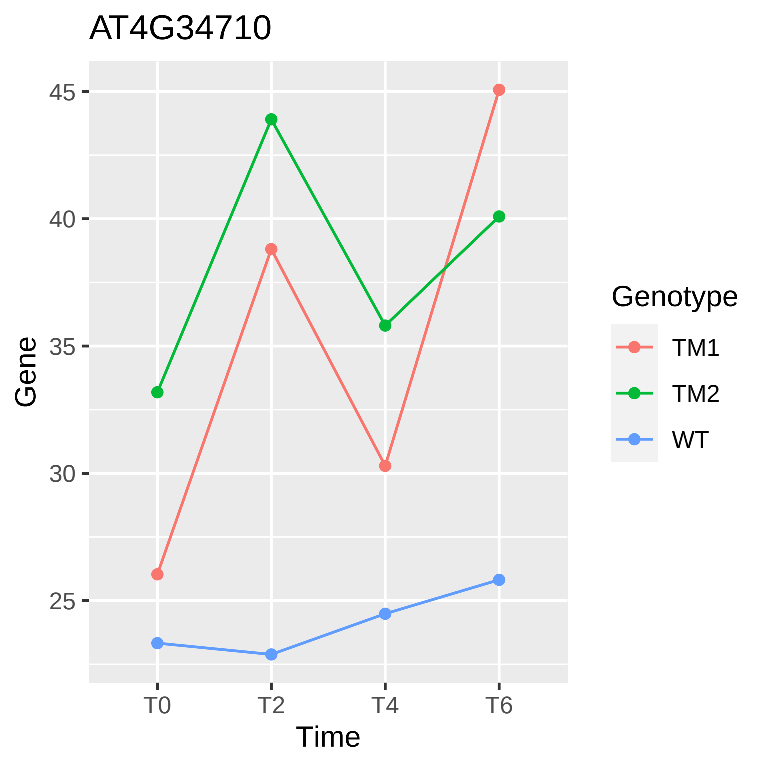
AT4G34710
45
40
35
30
25
Gene
T0
T2
T4
T6
Time
Genotype
TM1
TM2
WT

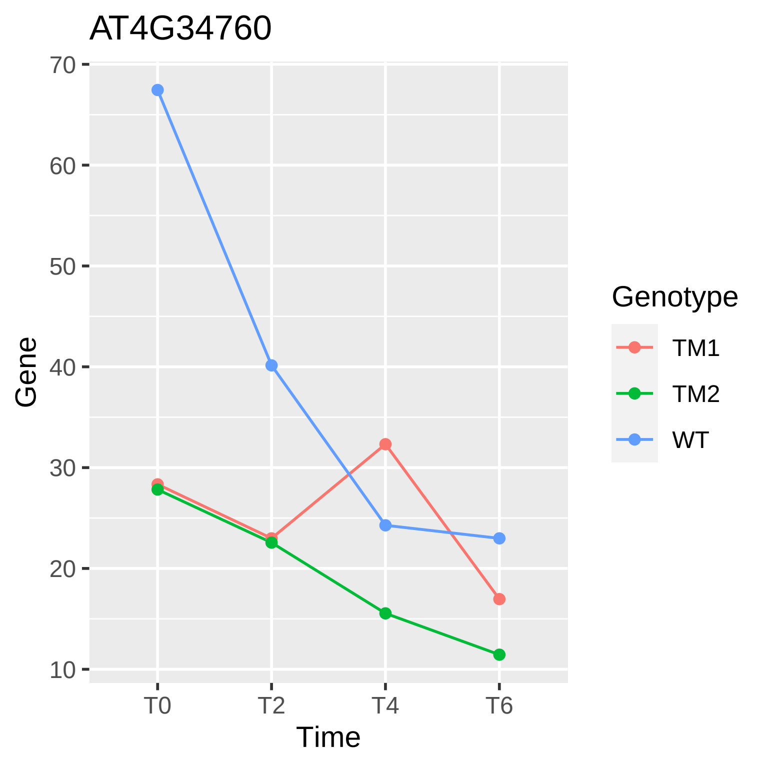
AT4G34760
70
60
50
40
30
20
10
Gene
T0
T2
T4
T6
Time
Genotype
TM1
TM2
WT

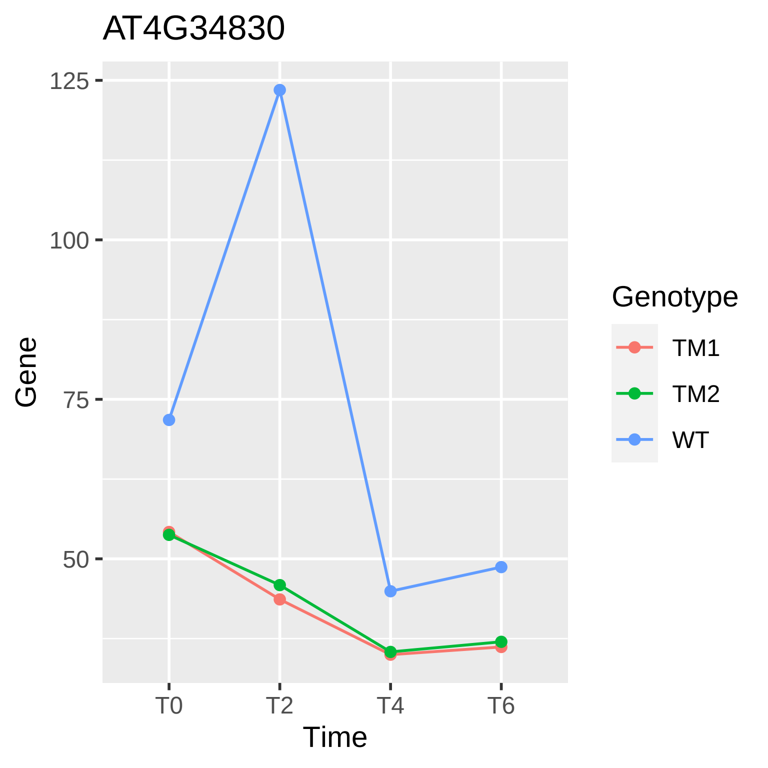
AT4G34830
125
100
75
50
Gene
T0
T2
T4
T6
Time
Genotype
TM1
TM2
WT

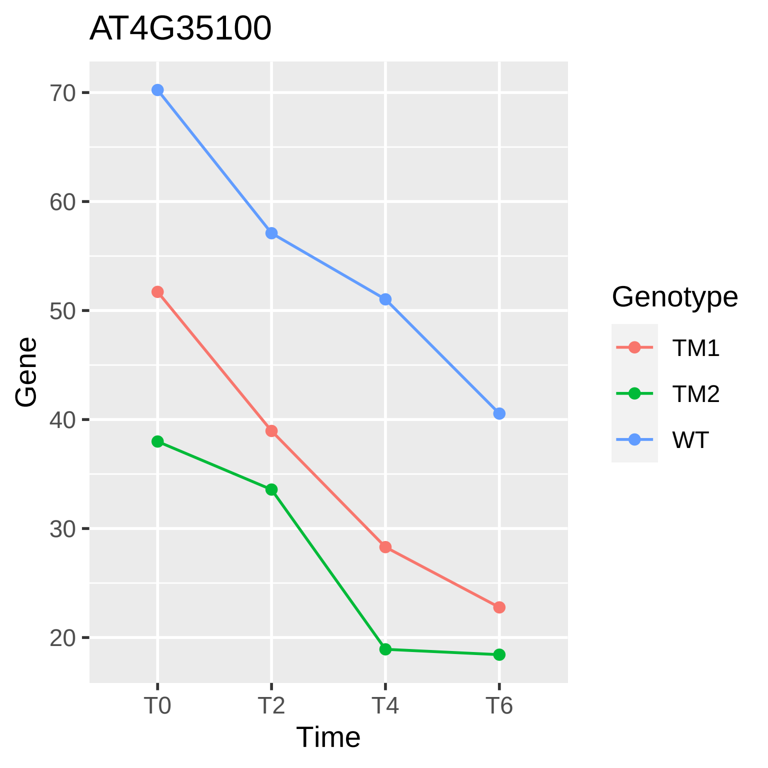

AT4G35100
70
60
50
40
30
20
Gene
T0
T2
T4
T6
Time
Genotype
TM1
TM2
WT

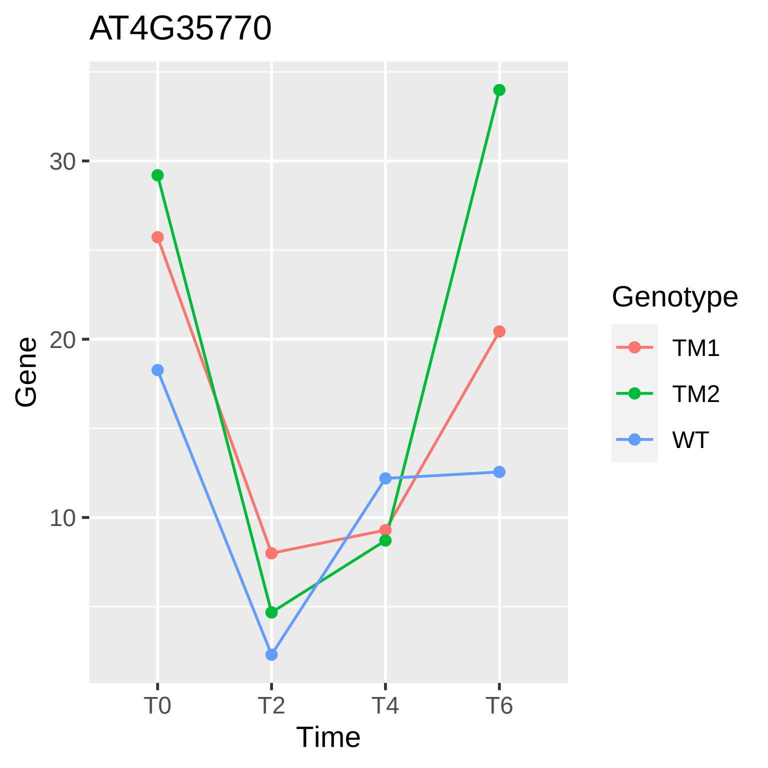
AT4G35770
Gene
30
20
10
T0
T2
T4
T6
Time
Genotype
TM1
TM2
WT

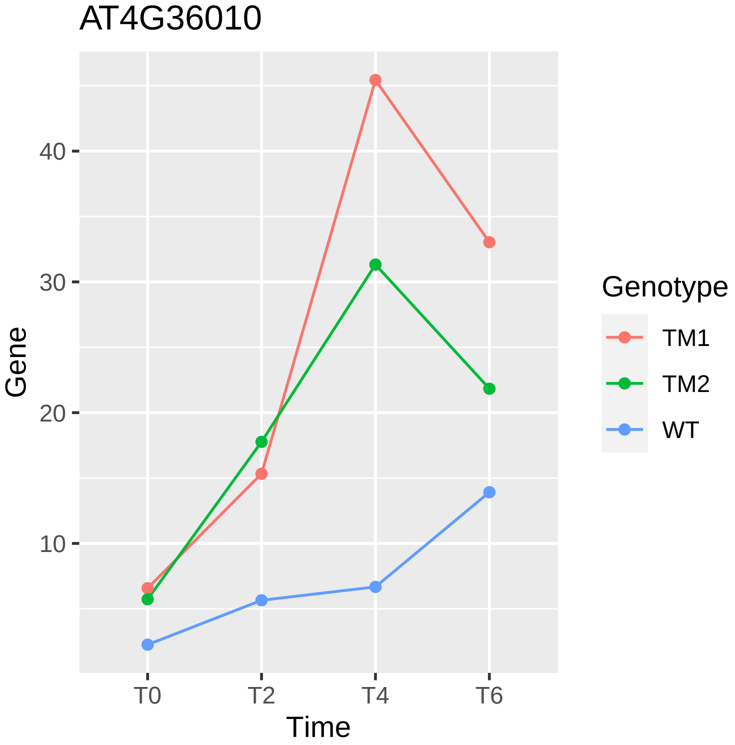
AT4G36010
Gene
40
30
20
10
T0
T2
T4
T6
Time
Genotype
TM1
TM2
WT

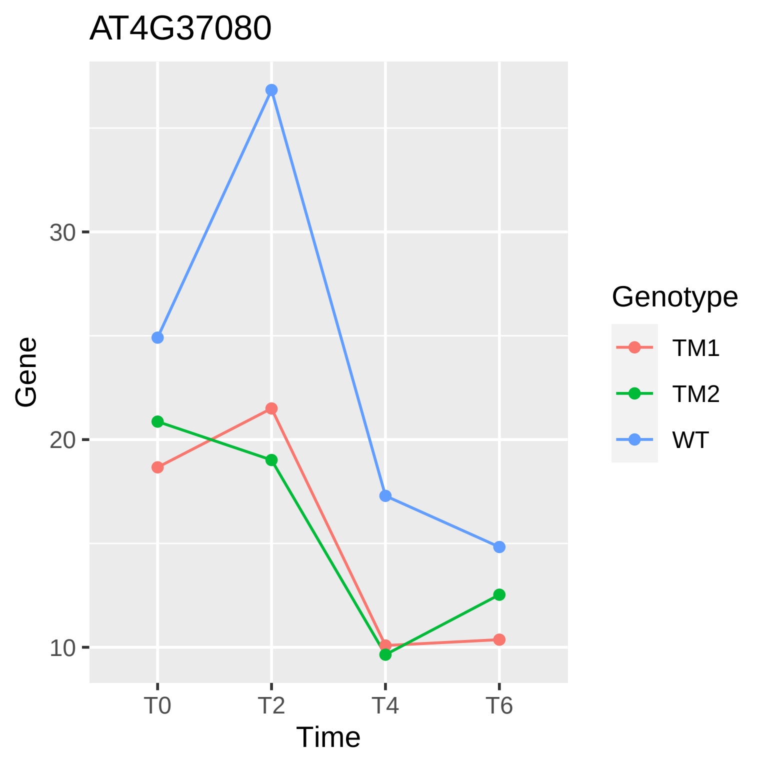
AT4G37080
Gene
30
20
10
T0
T2
T4
T6
Time
Genotype
TM1
TM2
WT

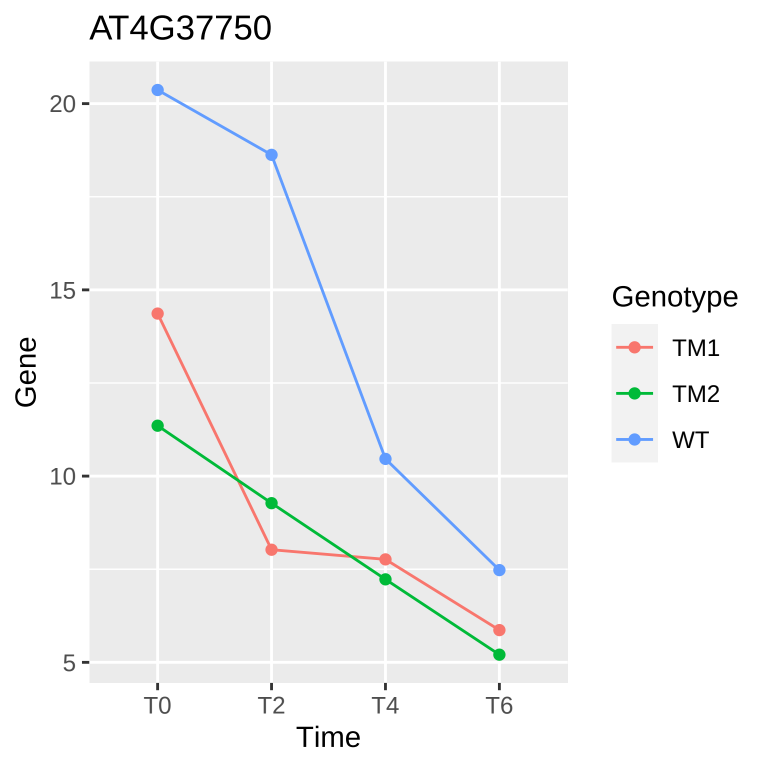

AT4G37750
20
15
10
5
Gene
T0
T2
T4
T6
Time
Genotype
TM1
TM2
WT

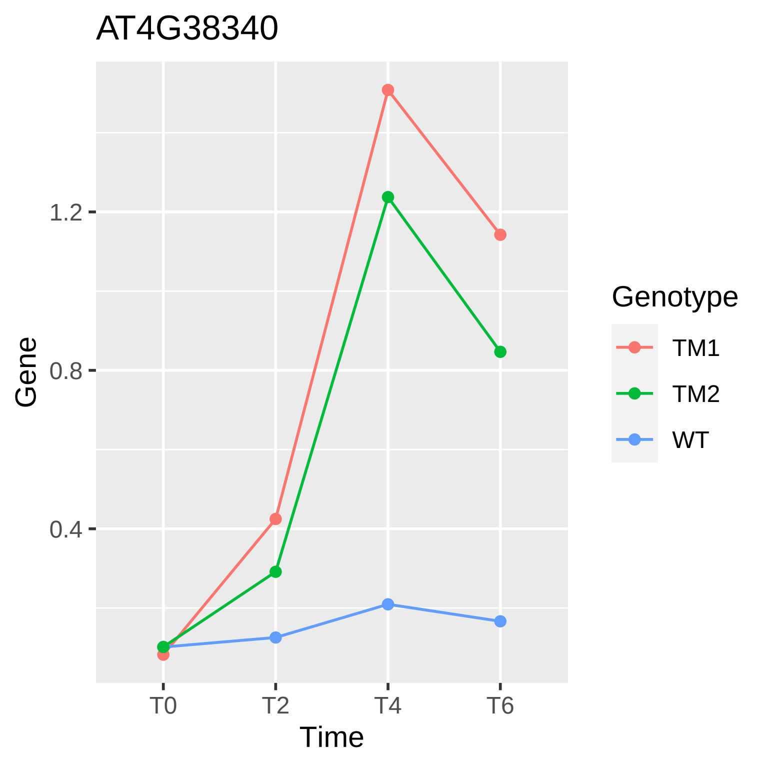
AT4G38340
Gene
1.2
0.8
0.4
T0
T2
T4
T6
Time
Genotype
TM1
TM2
WT

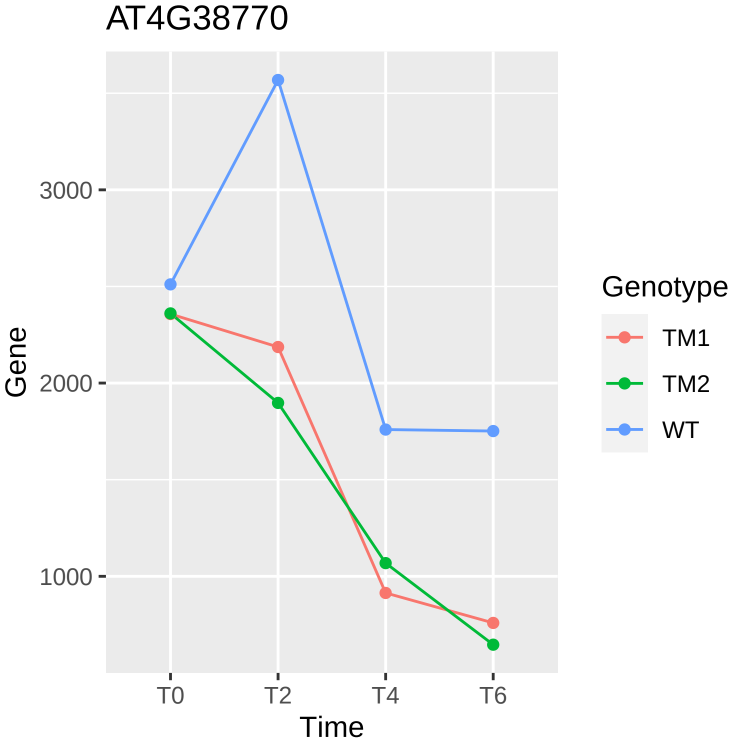

AT4G38770
Gene
3000
2000
1000
T0
T2
T4
T6
Time
Genotype
TM1
TM2
WT

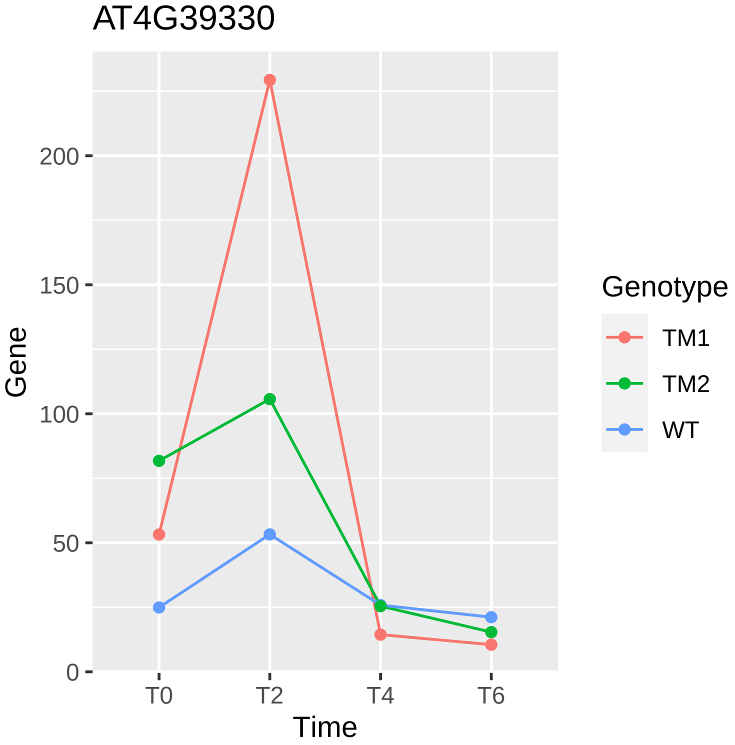
AT4G39330
Gene
200
150
100
50
0
T0
T2
T4
T6
Time
Genotype
TM1
TM2
WT

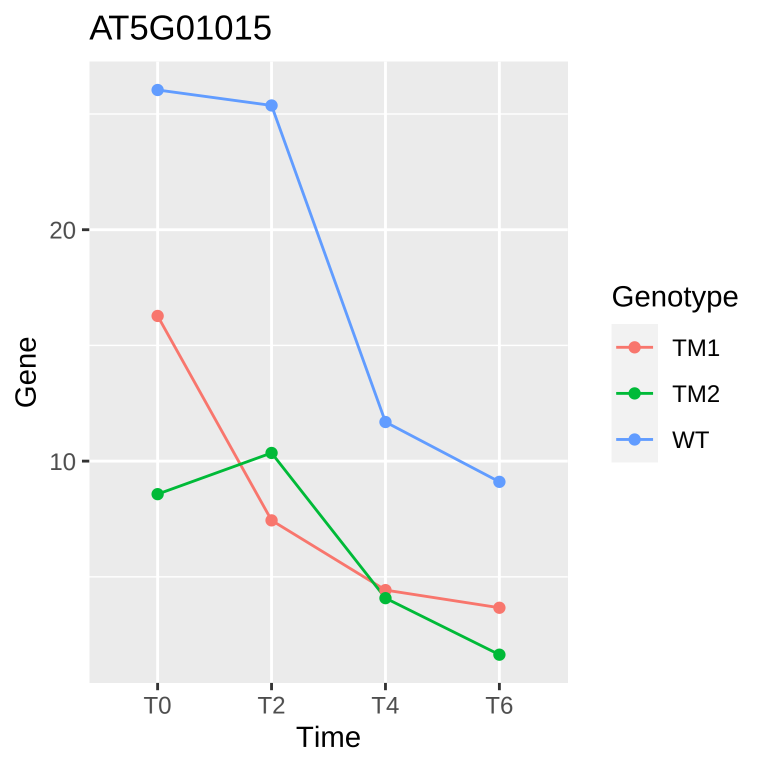

AT5G01015
Gene
20
10
T0
T2
T4
T6
Time
Genotype
TM1
TM2
WT

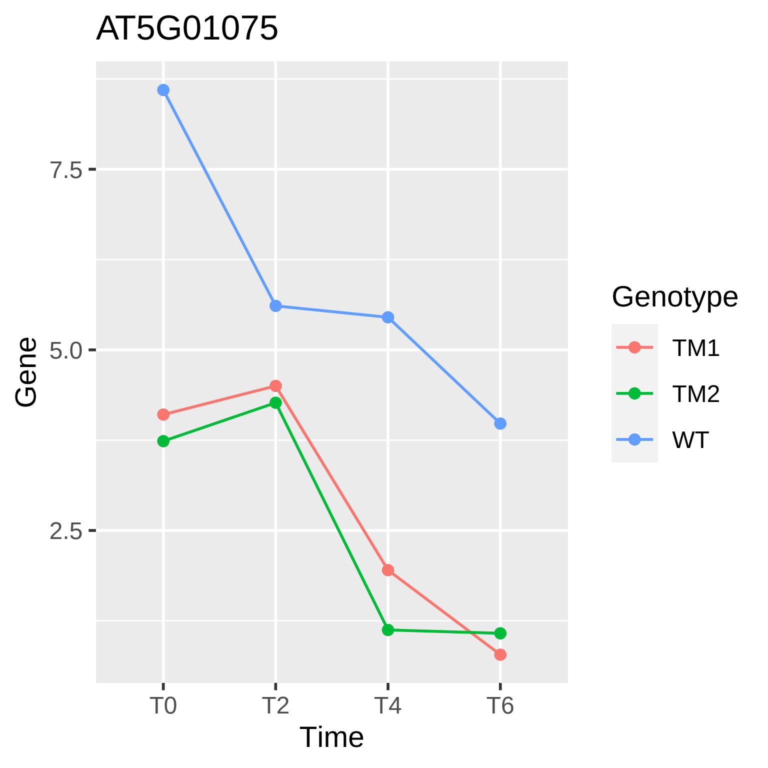

AT5G01075
Gene
7.5
5.0
2.5
T0
T2
T4
T6
Time
Genotype
TM1
TM2
WT

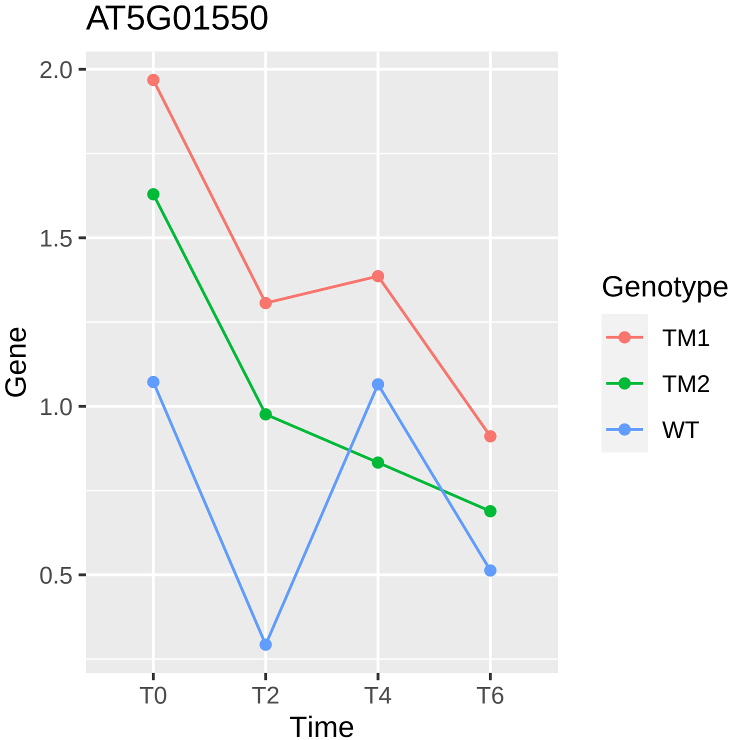
AT5G01550
2.0
1.5
1.0
0.5
Gene
T0
T2
T4
T6
Time
Genotype
TM1
TM2
WT

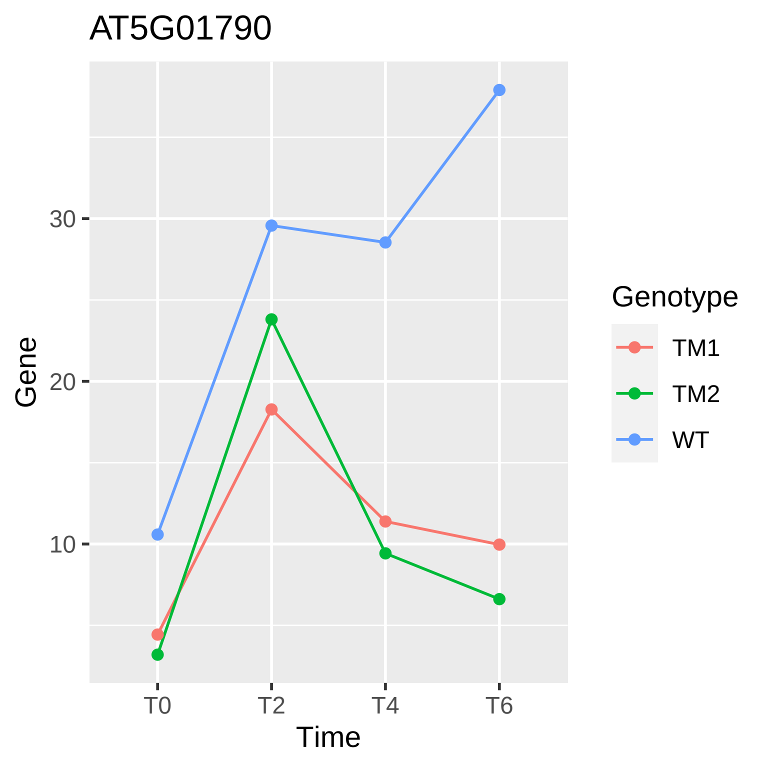

AT5G01790
Gene
30
20
10
T0
T2
T4
T6
Time
Genotype
TM1
TM2
WT

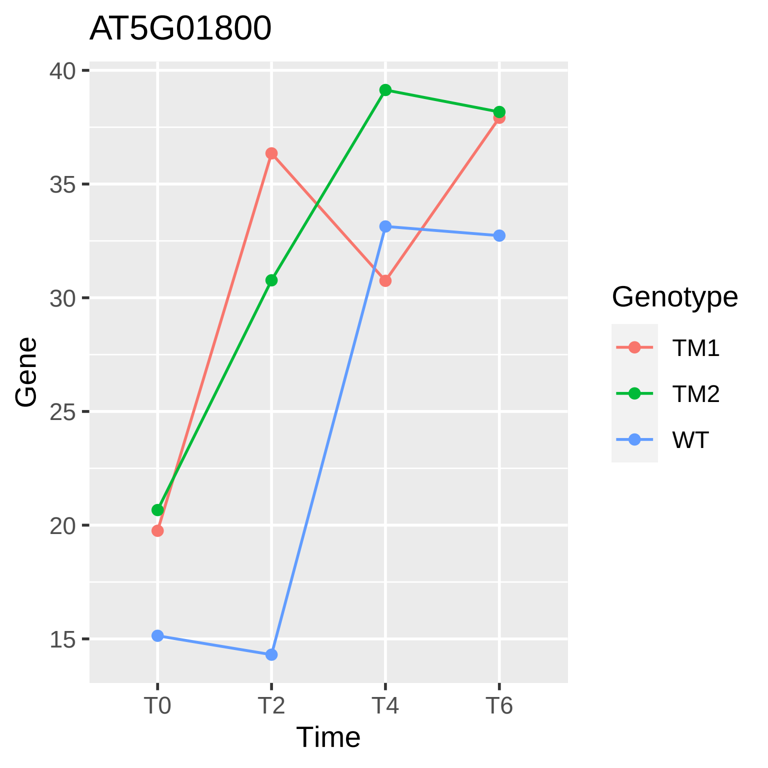
AT5G01800
Gene
40
35
30
25
20
15
T0
T2
T4
T6
Time
Genotype
TM1
TM2
WT

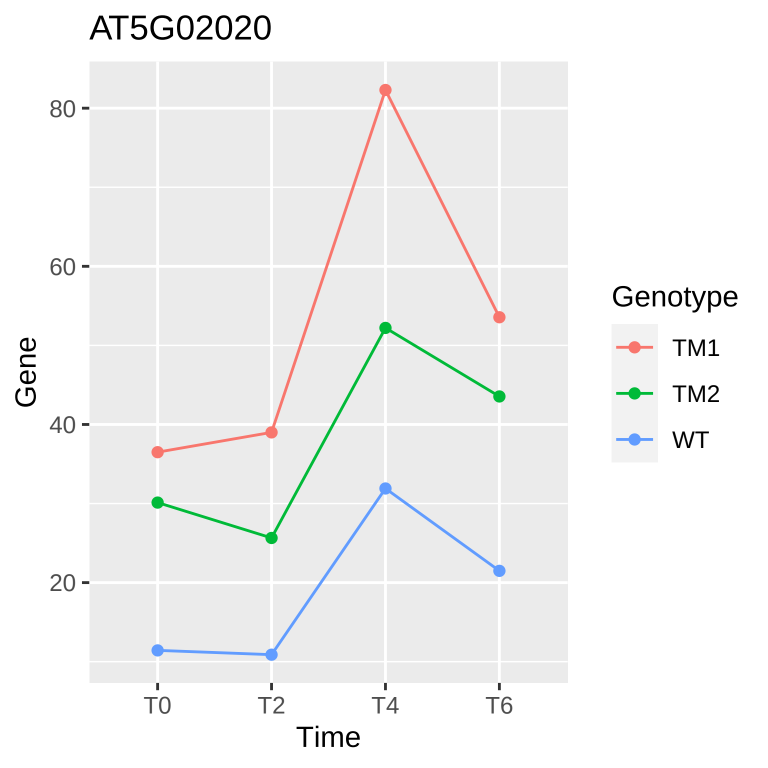

AT5G02020
80
60
40
20
Gene
T0
T2
T4
T6
Time
Genotype
TM1
TM2
WT

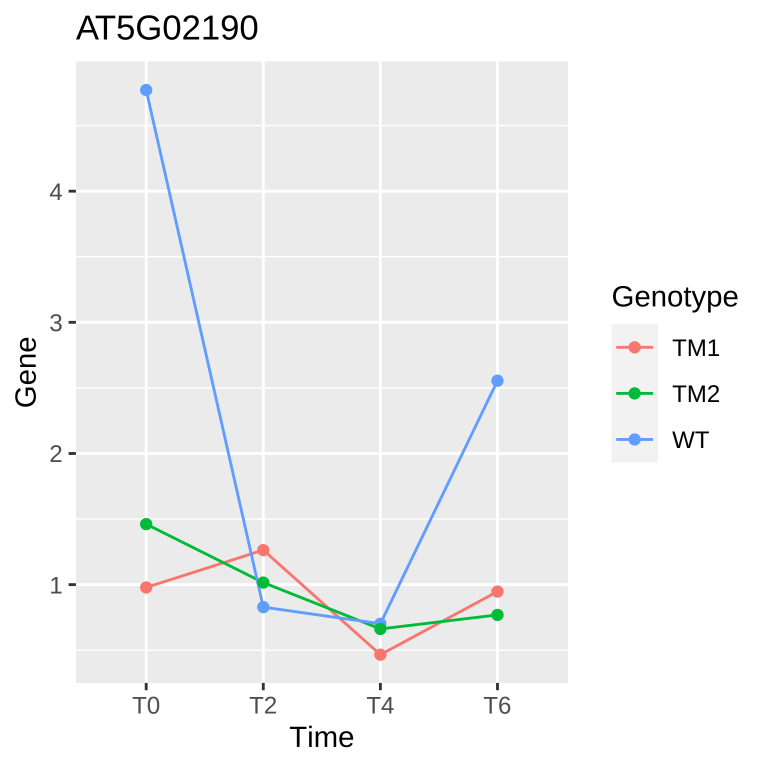

AT5G02190
Gene
1
2
3
4
T0
T2
T4
T6
Time
Genotype
TM1
TM2
WT

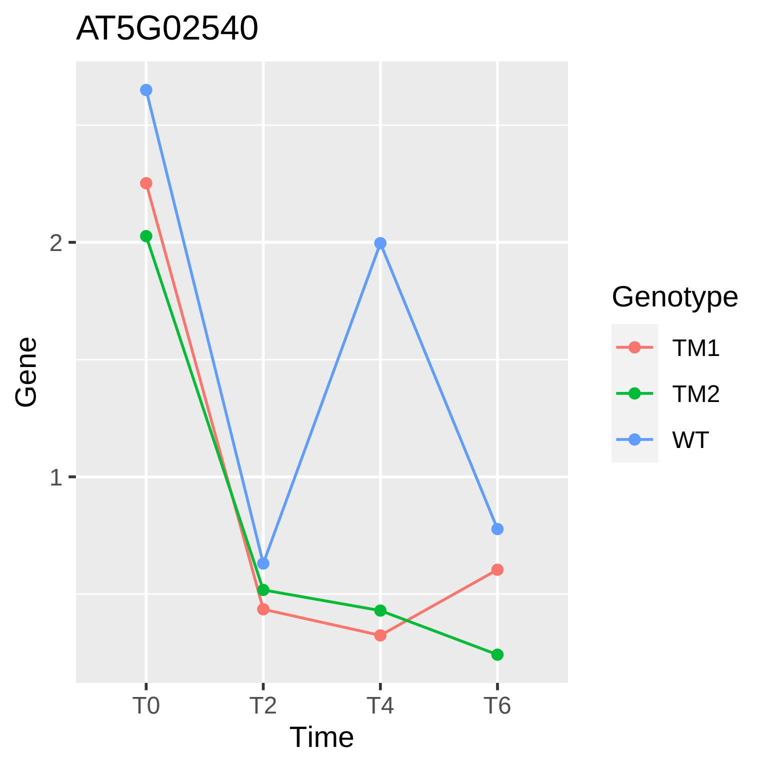

AT5G02540
Gene
2
1
T0
T2
T4
T6
Time
Genotype
TM1
TM2
WT

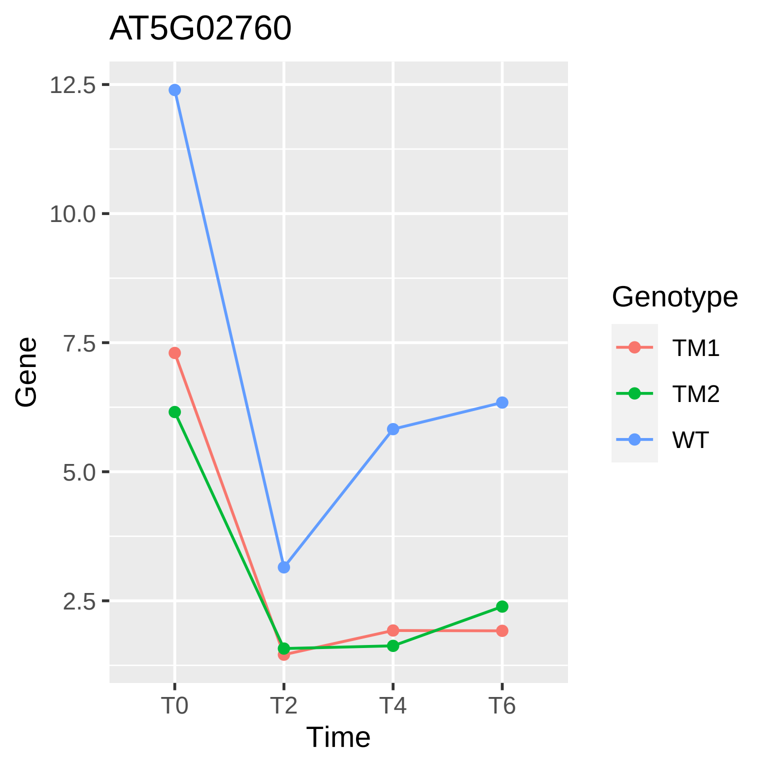

AT5G02760
12.5
10.0
7.5
5.0
2.5
Gene
T0
T2
T4
T6
Time
Genotype
TM1
TM2
WT

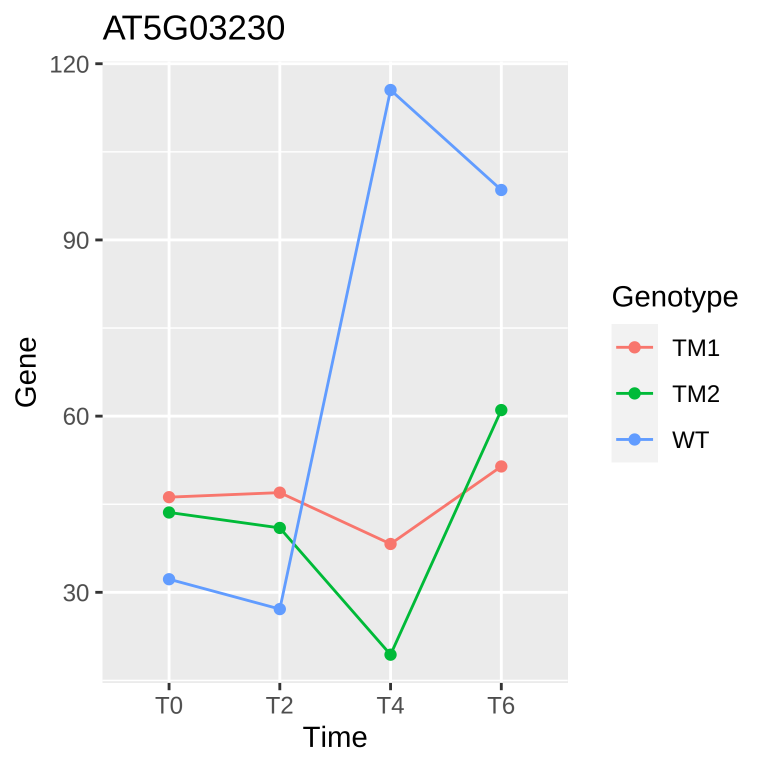

AT5G03230
120
90
60
30
Gene
T0
T2
T4
T6
Time
Genotype
TM1
TM2
WT

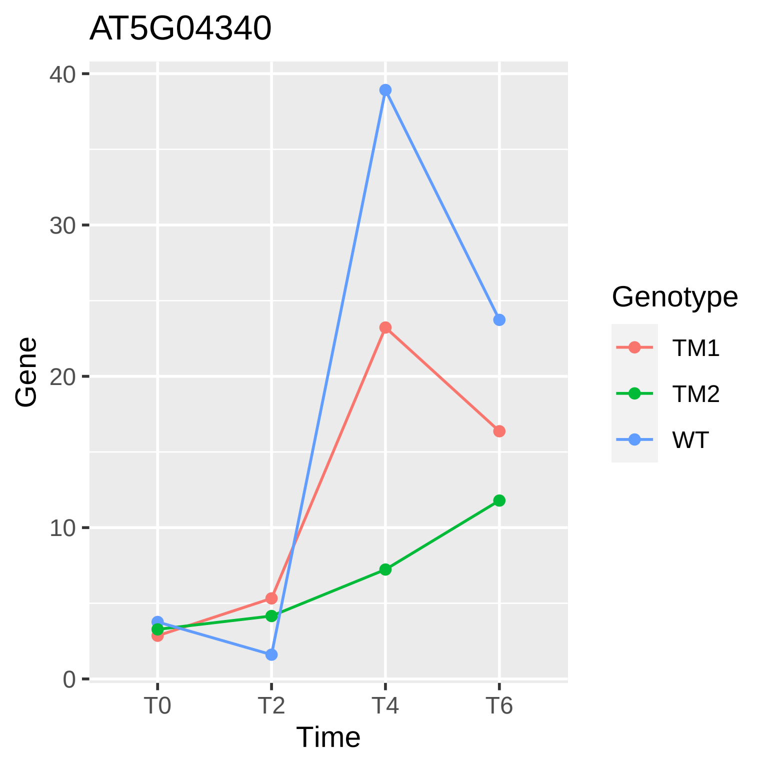

AT5G04340
Gene
0
10
20
30
40
T0
T2
T4
T6
Time
Genotype
TM1
TM2
WT

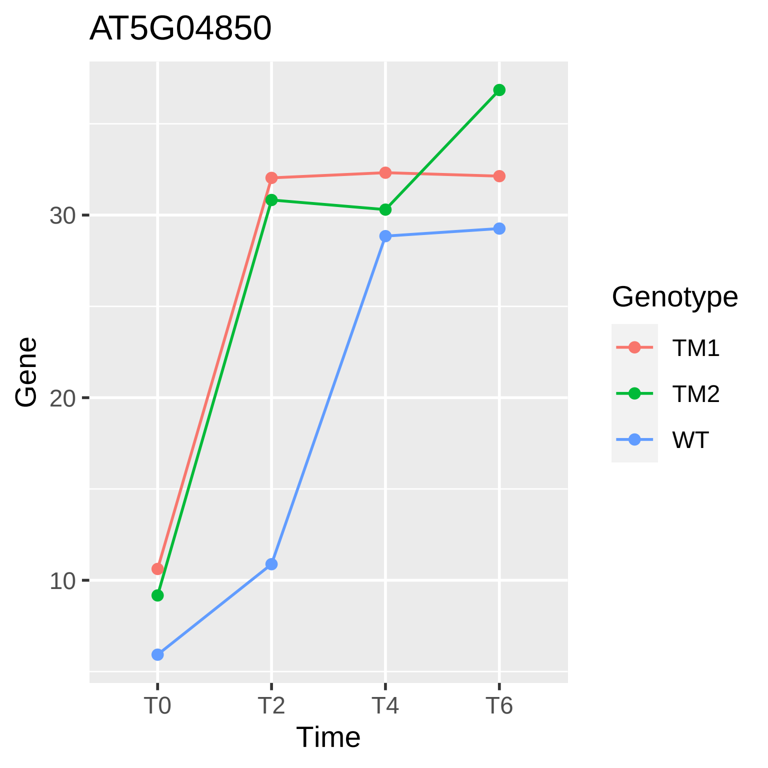
AT5G04850
Gene
30
20
10
T0
T2
T4
T6
Time
Genotype
TM1
TM2
WT

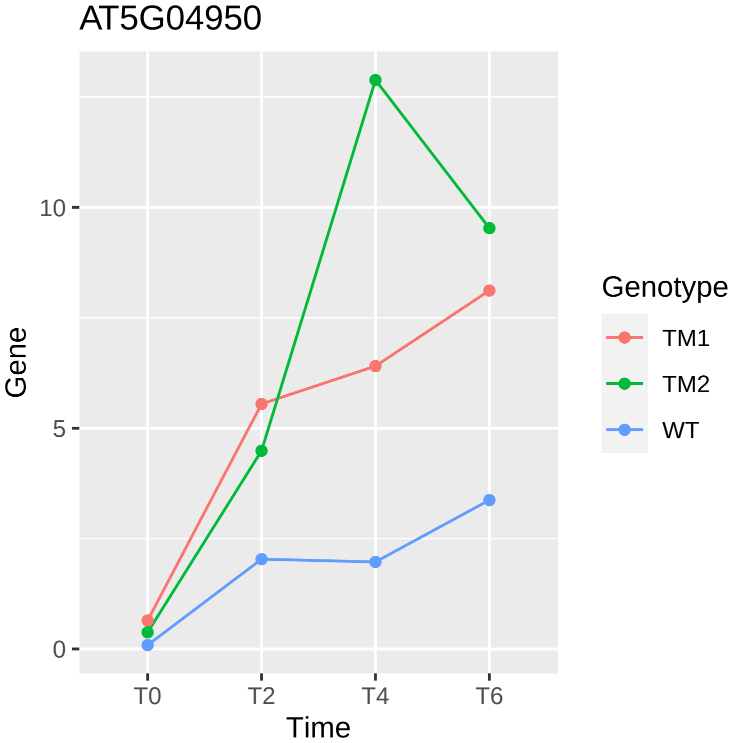

AT5G04950
10
5
0
Gene
T0
T2
T4
T6
Time
Genotype
TM1
TM2
WT

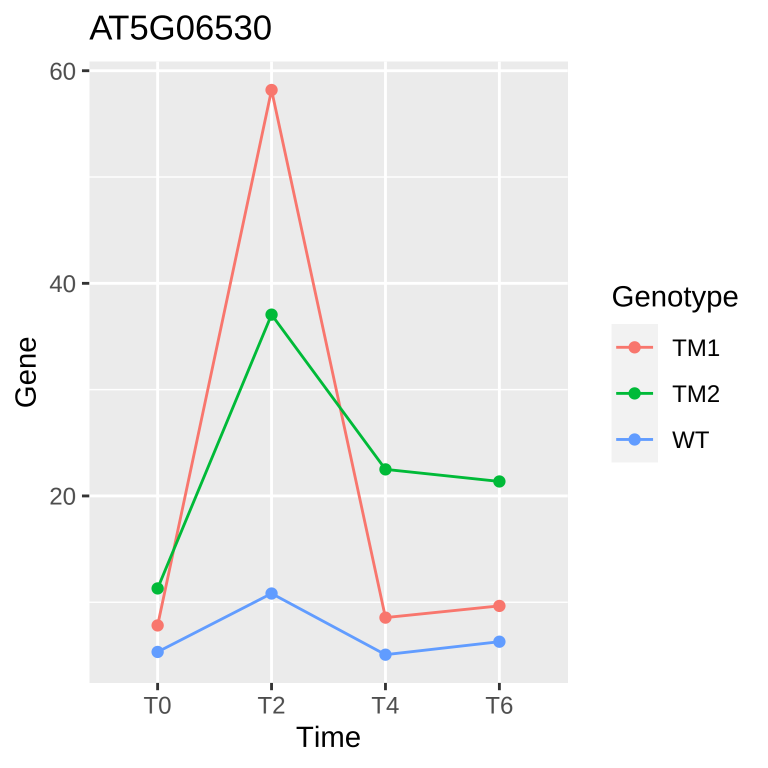
AT5G06530
60
40
20
Gene
T0
T2
T4
T6
Time
Genotype
TM1
TM2
WT

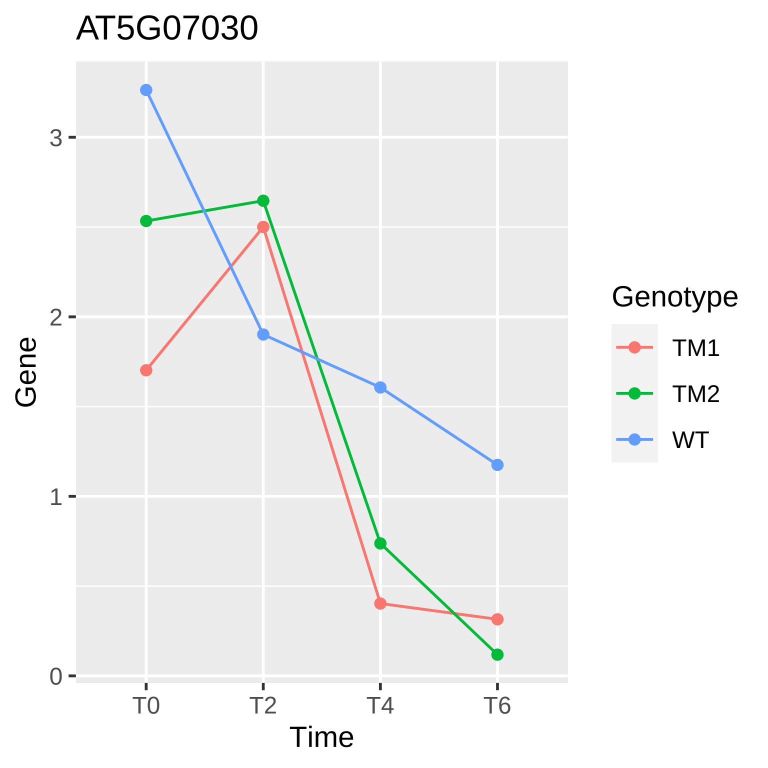
AT5G07030
3
2
1
0
Gene
T0
T2
T4
T6
Time
Genotype
TM1
TM2
WT

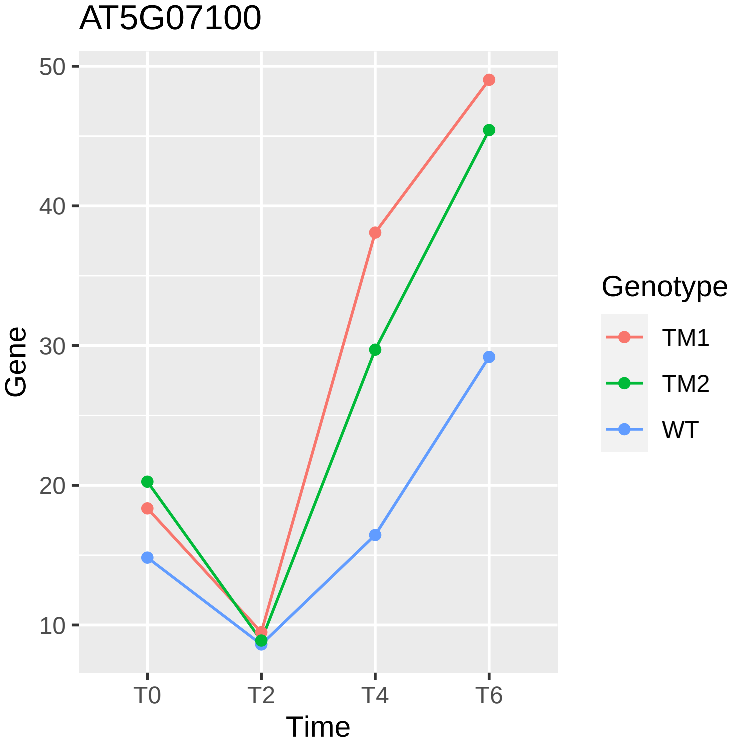
AT5G07100
Gene
50
40
30
20
10
T0
T2
T4
T6
Time
Genotype
TM1
TM2
WT

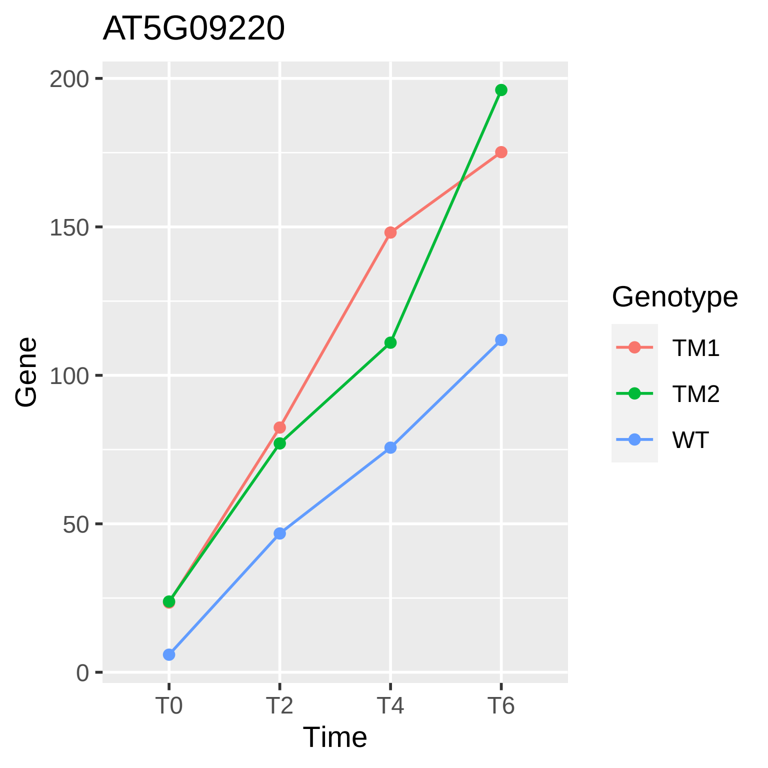

AT5G09220
200
150
100
50
0
Gene
T0
T2
T4
T6
Time
Genotype
TM1
TM2
WT

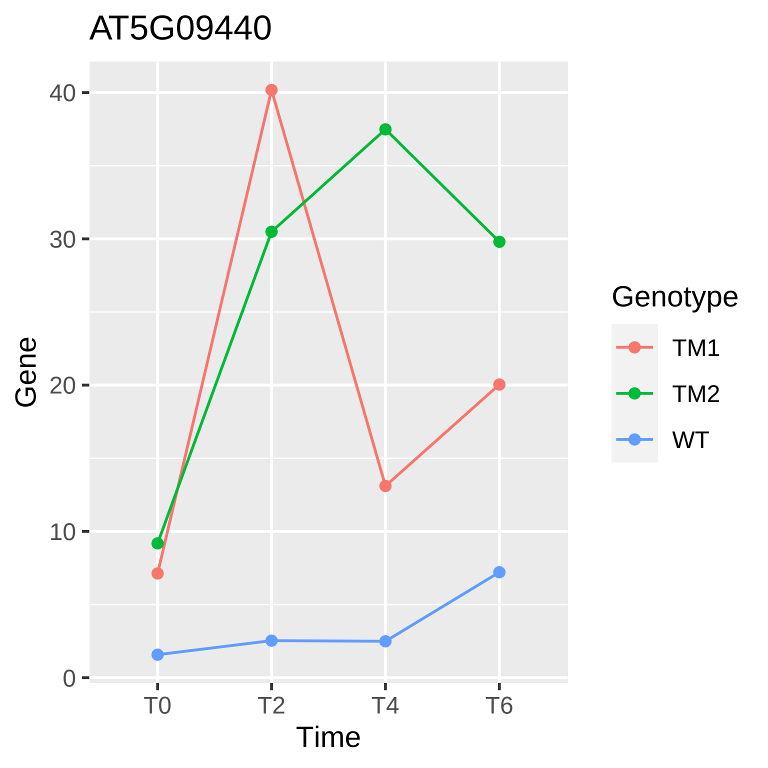
AT5G09440
Gene
40
30
20
10
0
T0
T2
T4
T6
Time
Genotype
TM1
TM2
WT

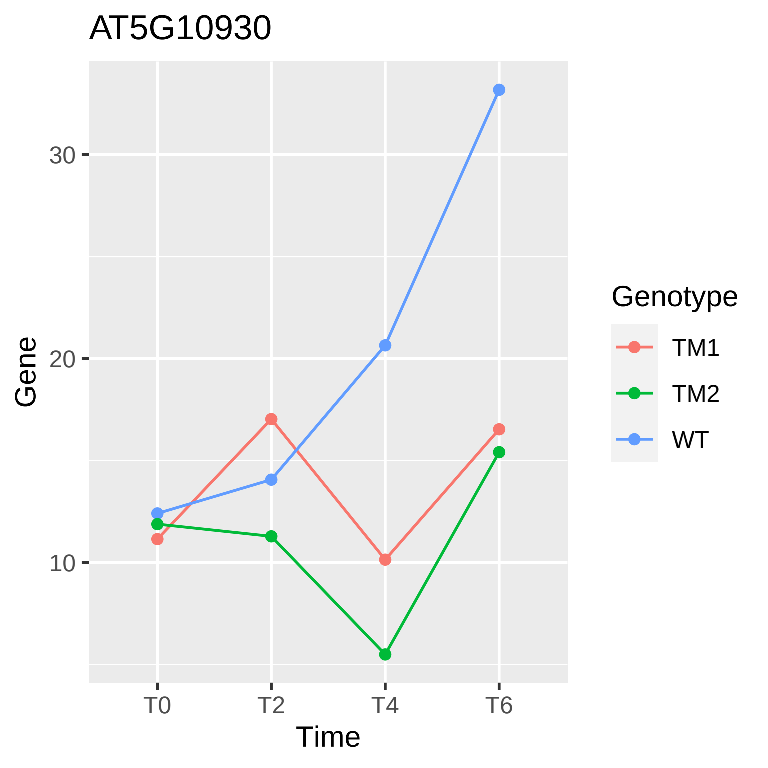

AT5G10930
Gene
30
20
10
T0
T2
T4
T6
Time
Genotype
TM1
TM2
WT

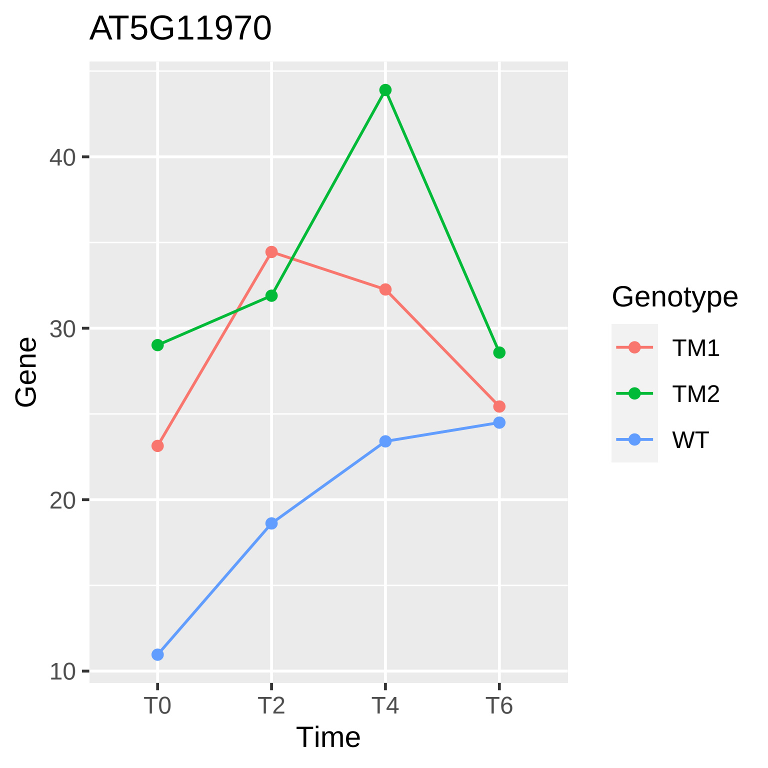

AT5G11970
Gene
40
30
20
10
T0
T2
T4
T6
Time
Genotype
TM1
TM2
WT

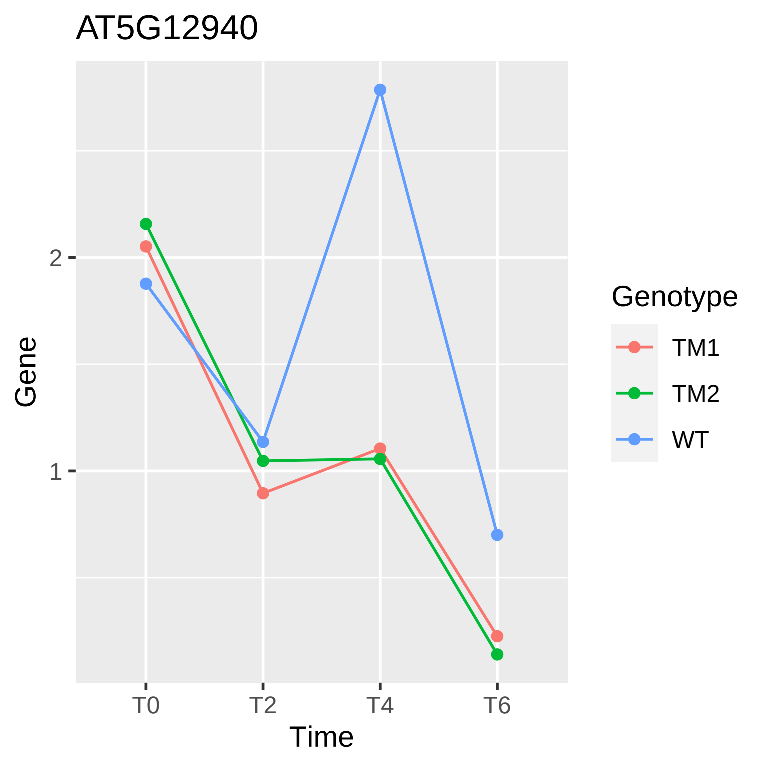

AT5G12940
Gene
Time
T0
T2
T4
T6
1
2
Genotype
TM1
TM2
WT

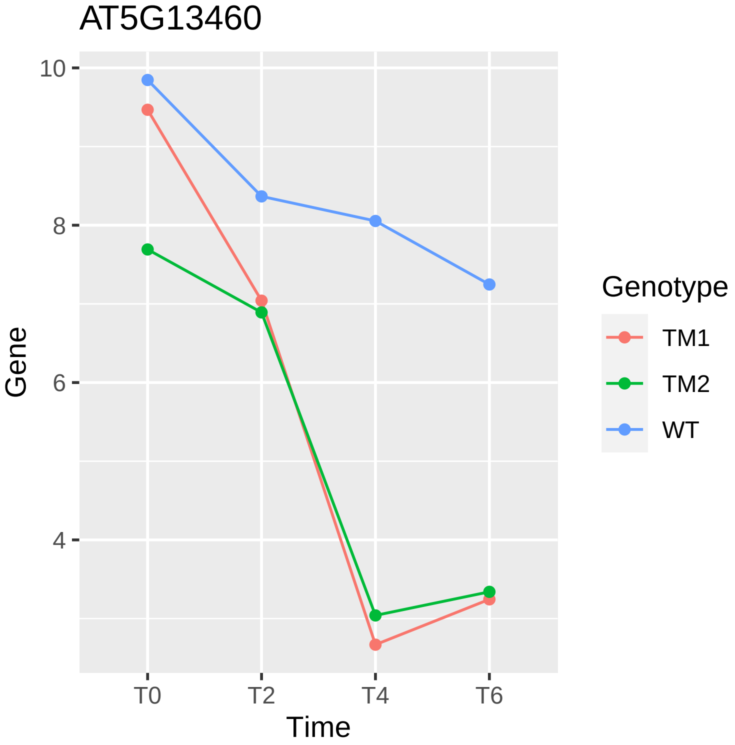
AT5G13460
10
8
6
4
Gene
T0
T2
T4
T6
Time
Genotype
TM1
TM2
WT

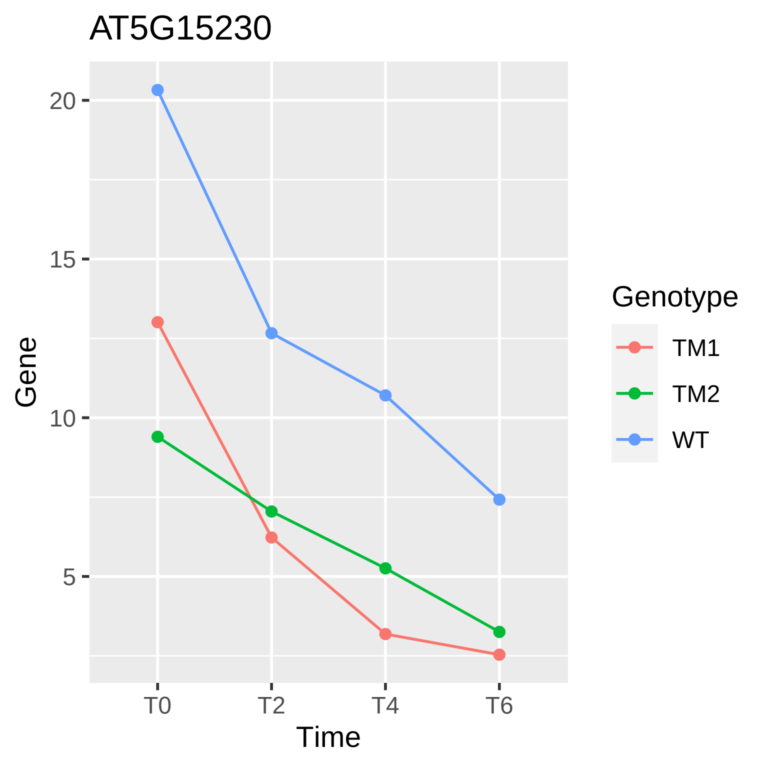

AT5G15230
20
15
10
5
Gene
T0
T2
T4
T6
Time
Genotype
TM1
TM2
WT

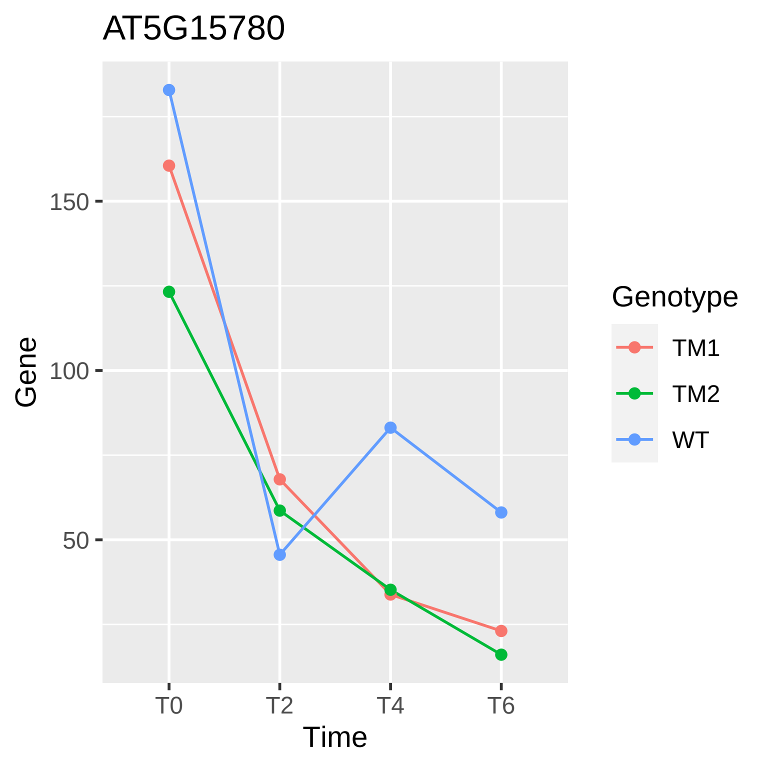
AT5G15780
Gene
150
100
50
T0
T2
T4
T6
Time
Genotype
TM1
TM2
WT

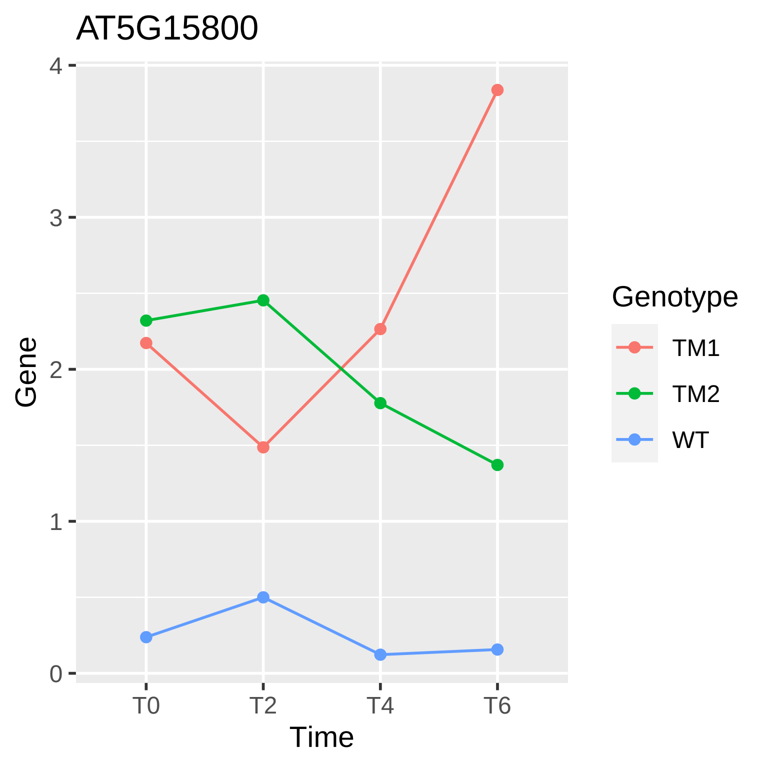

AT5G15800
Gene
4
3
2
1
0
T0
T2
T4
T6
Time
Genotype
TM1
TM2
WT

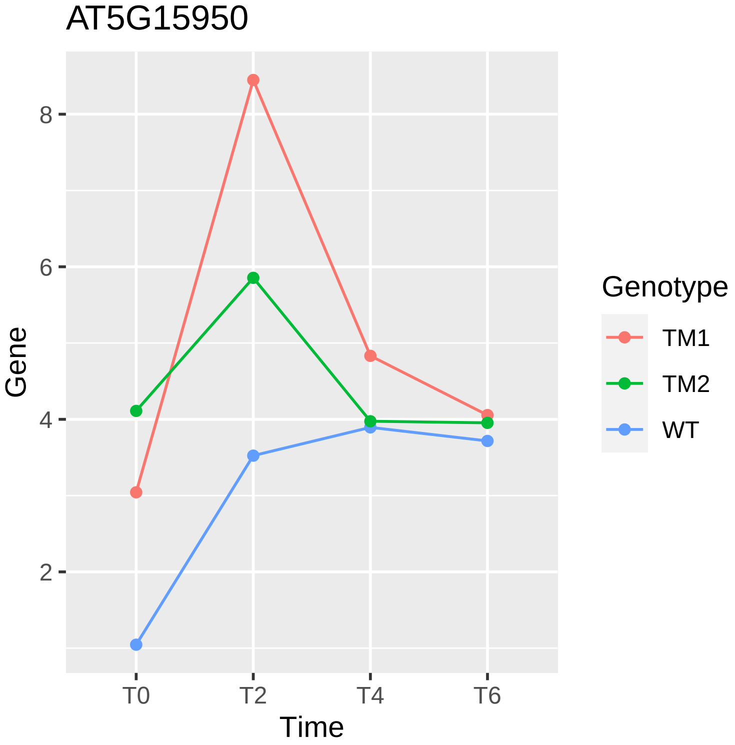
AT5G15950
Gene
8
6
4
2
T0
T2
T4
T6
Time
Genotype
TM1
TM2
WT

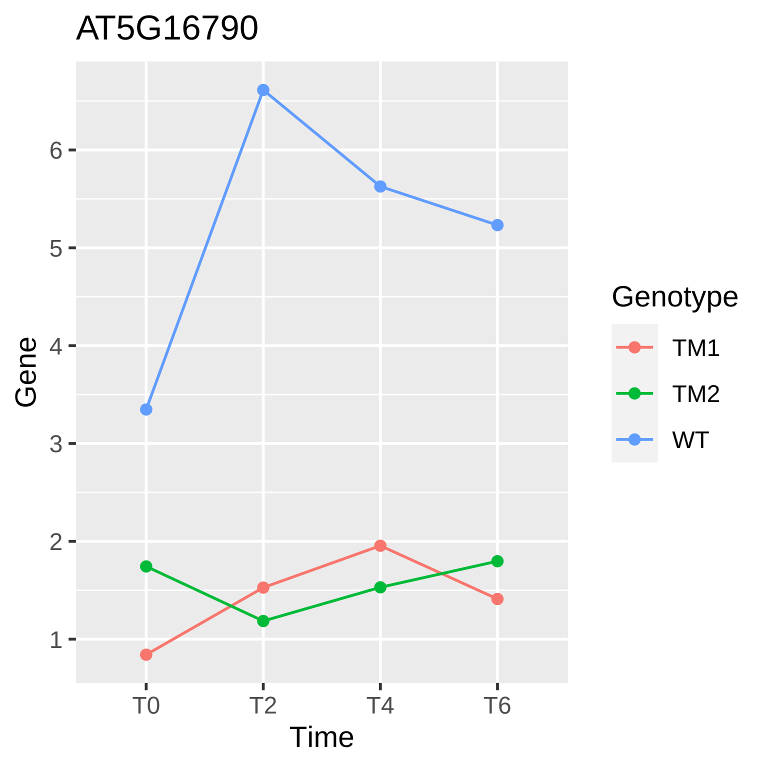
AT5G16790
Gene
6
5
4
3
2
1
T0
T2
T4
T6
Time
Genotype
TM1
TM2
WT

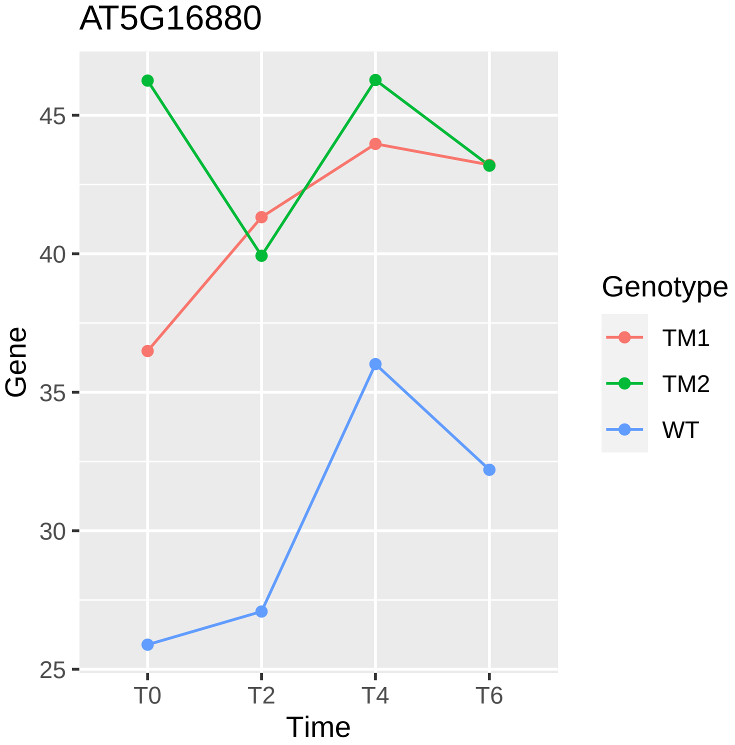

AT5G16880
Gene
45
40
35
30
25
T0
T2
T4
T6
Time
Genotype
TM1
TM2
WT

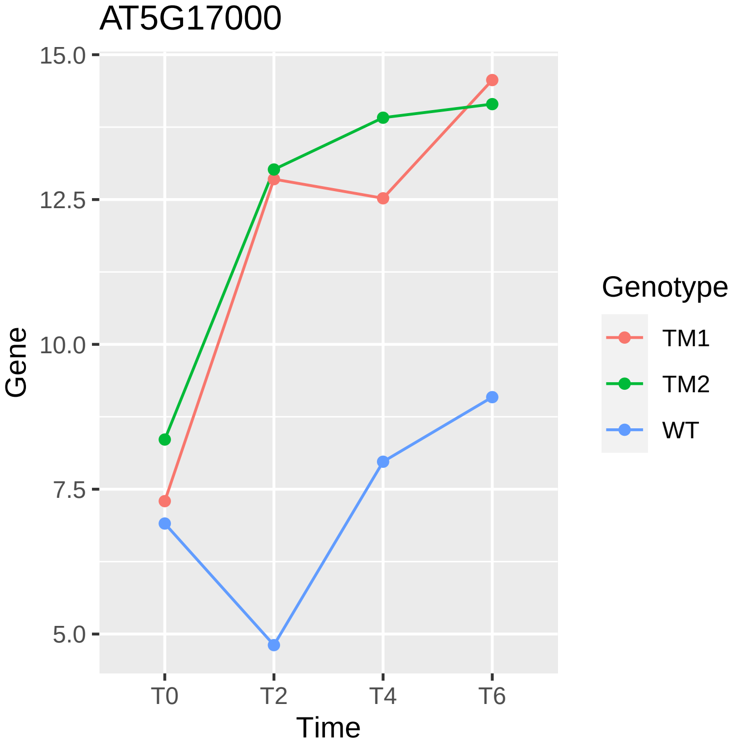
AT5G17000
15.0
12.5
10.0
7.5
5.0
Gene
T0
T2
T4
T6
Time
Genotype
TM1
TM2
WT

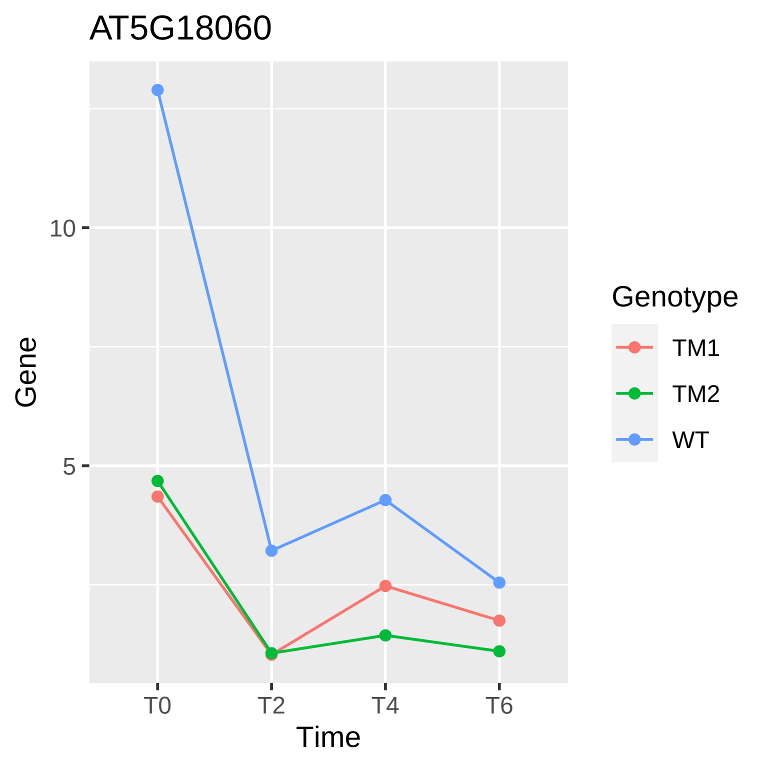

AT5G18060
Gene
10
5
T0
T2
T4
T6
Time
Genotype
TM1
TM2
WT

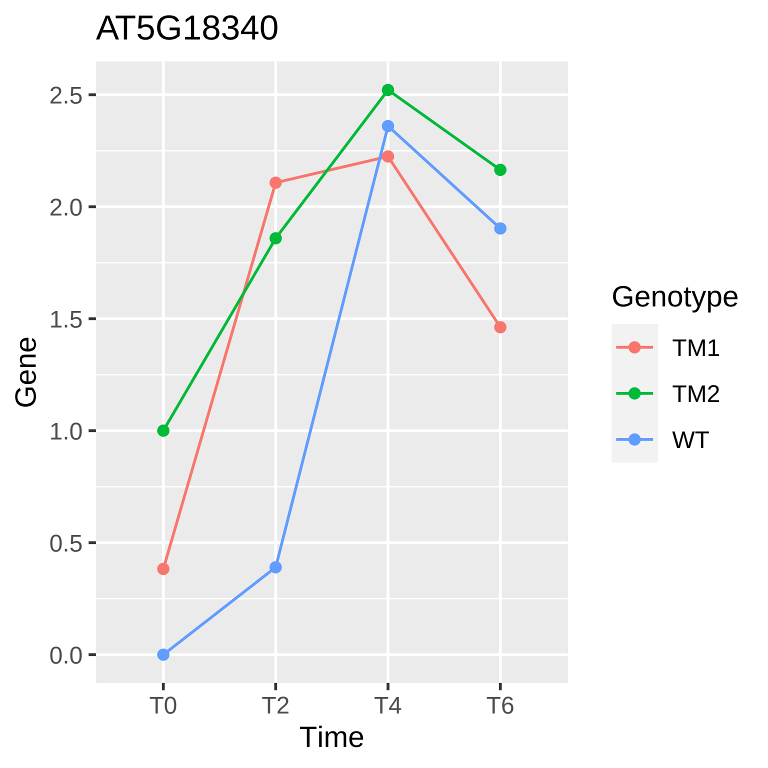
AT5G18340
2.5
2.0
1.5
1.0
0.5
0.0
Gene
T0
T2
T4
T6
Time
Genotype
TM1
TM2
WT

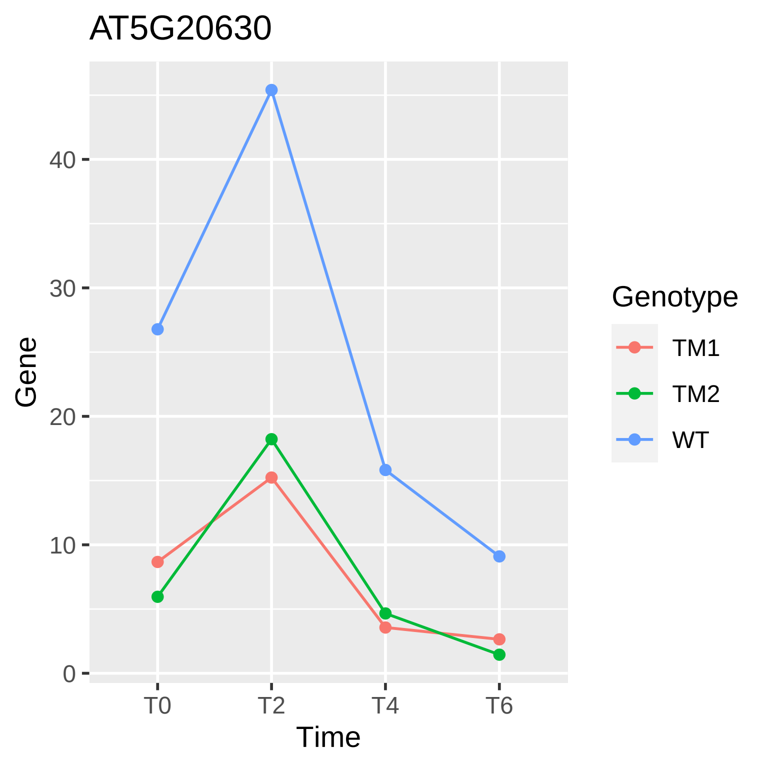

AT5G20630
Gene
Time
0
10
20
30
40
T0
T2
T4
T6
Genotype
TM1
TM2
WT

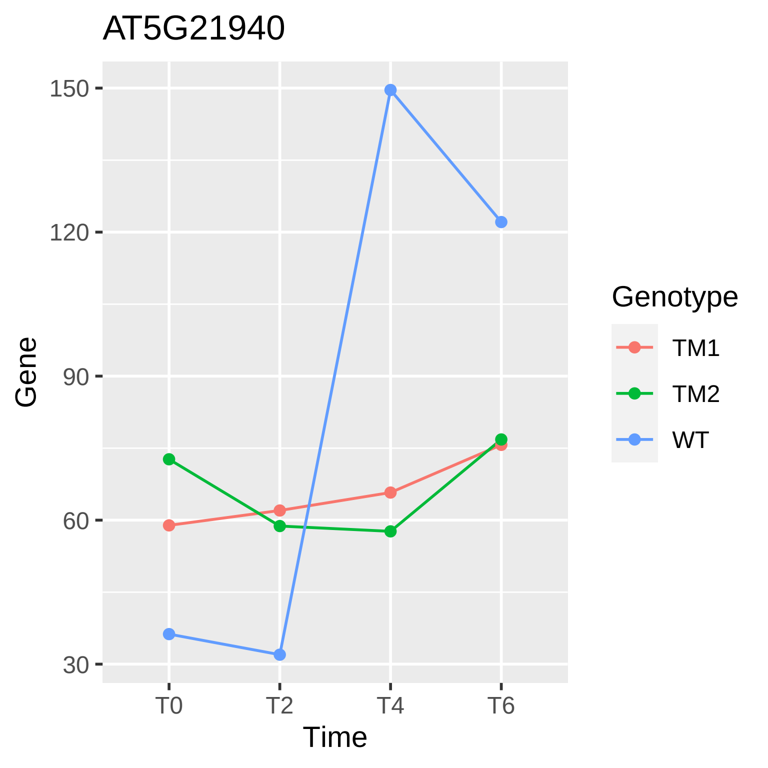

AT5G21940
150
120
90
60
30
Gene
T0
T2
T4
T6
Time
Genotype
TM1
TM2
WT

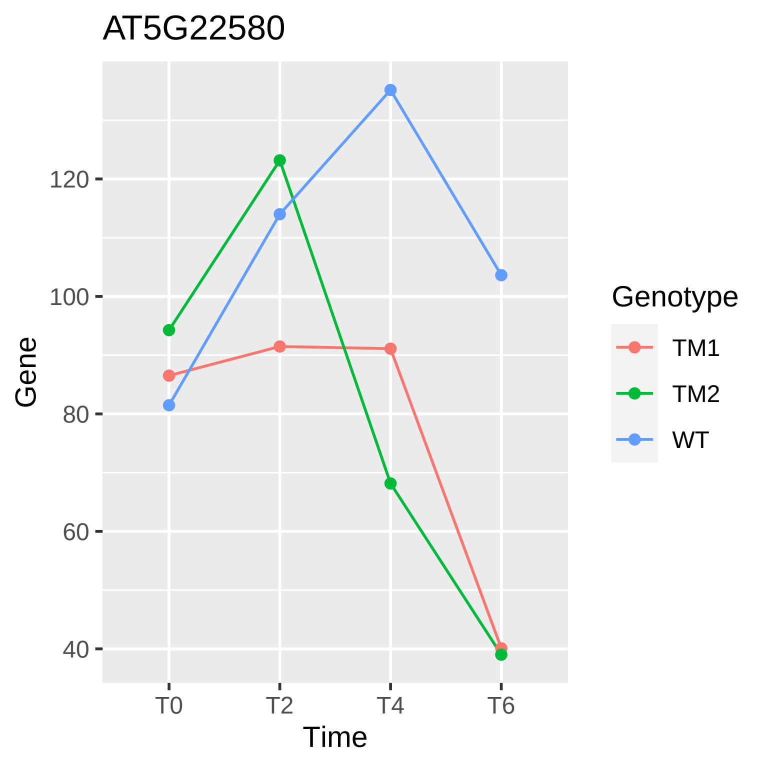

AT5G22580
Gene
120
100
80
60
40
T0
T2
T4
T6
Time
Genotype
TM1
TM2
WT

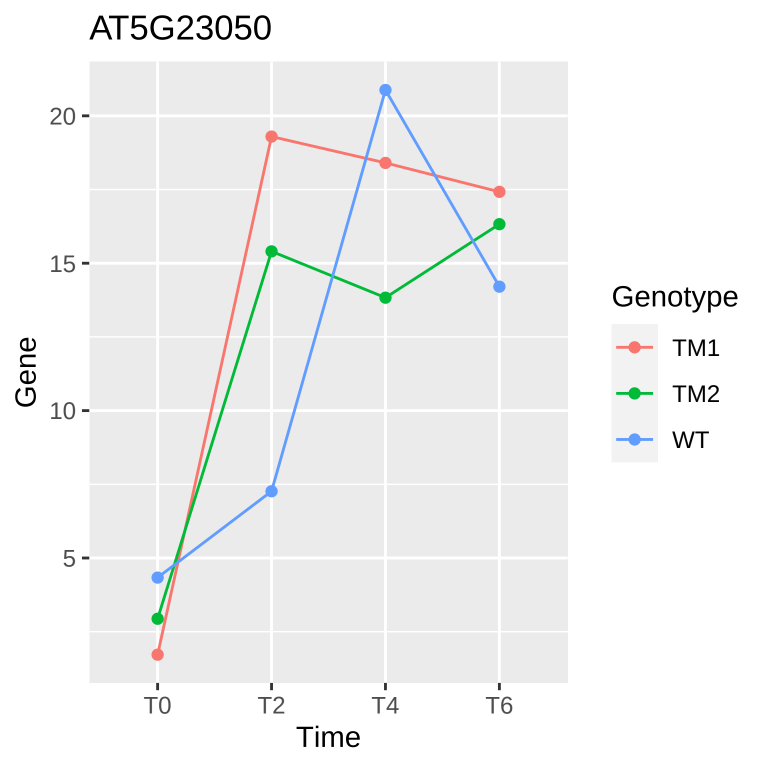

AT5G23050
Gene
20
15
10
5
T0
T2
T4
T6
Time
Genotype
TM1
TM2
WT

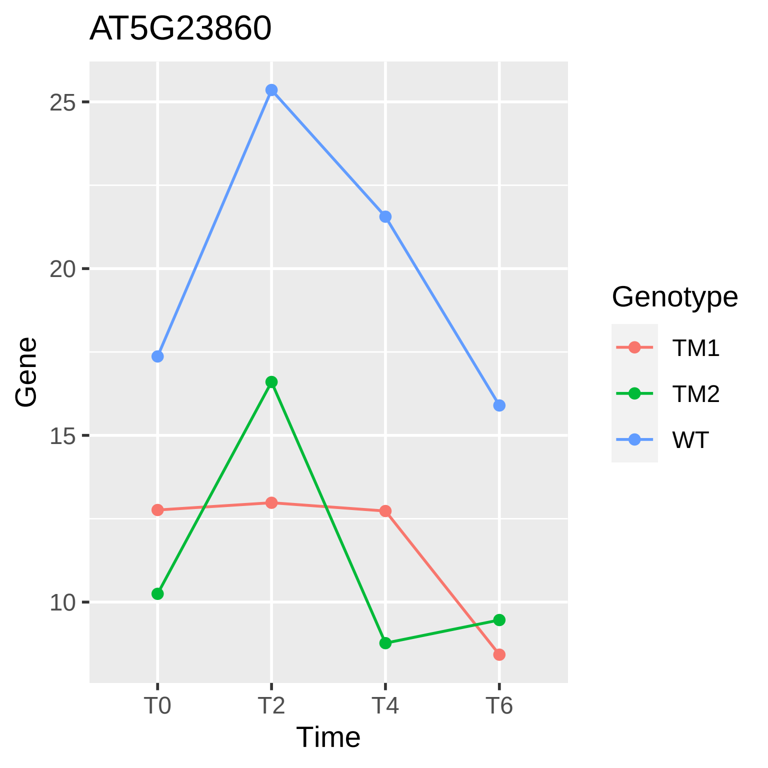

AT5G23860
25
20
15
10
Gene
T0
T2
T4
T6
Time
Genotype
TM1
TM2
WT

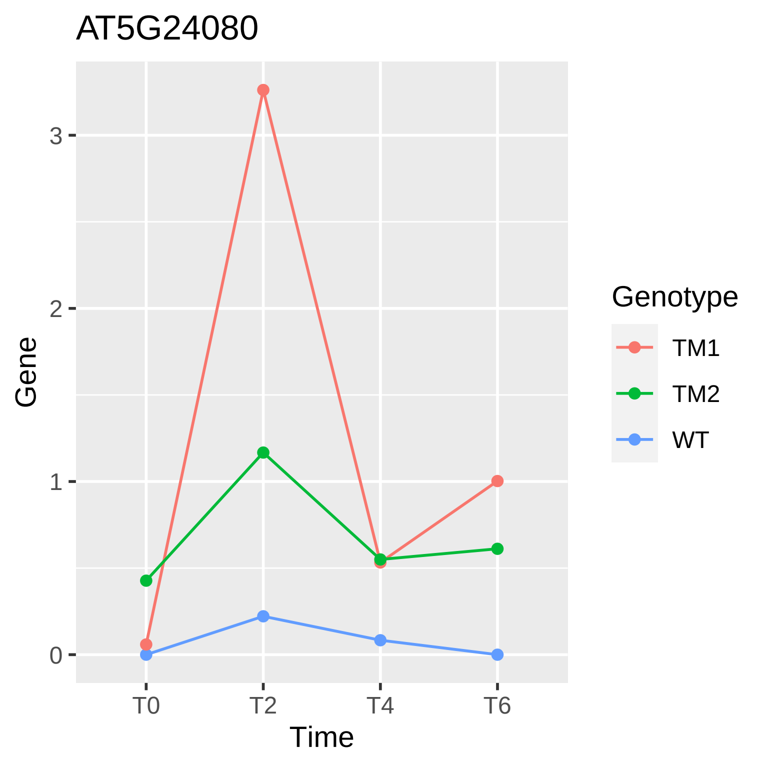
AT5G24080
Gene
3
2
1
0
T0
T2
T4
T6
Time
Genotype
TM1
TM2
WT

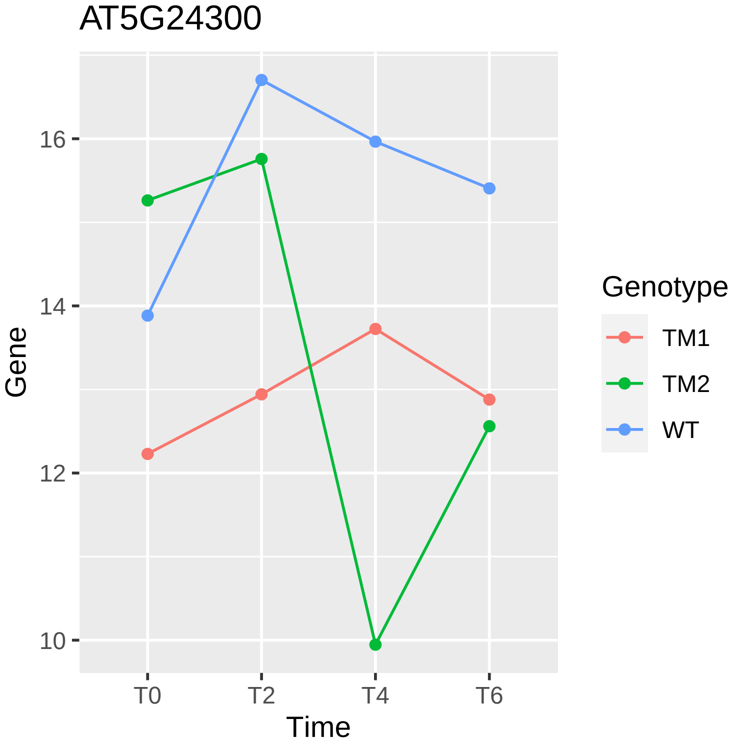
AT5G24300
Gene
16
14
12
10
T0
T2
T4
T6
Time
Genotype
TM1
TM2
WT

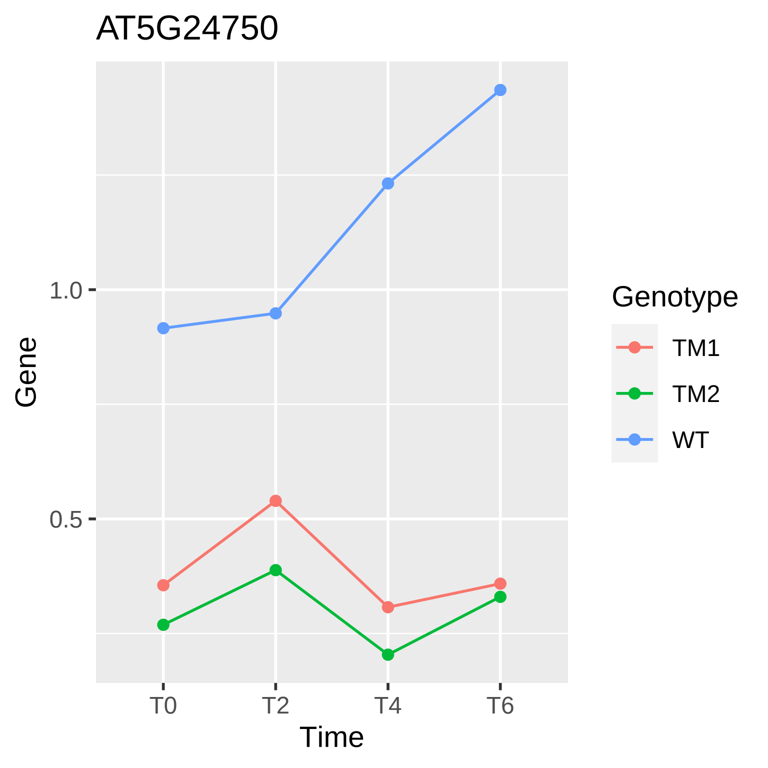

AT5G24750
Gene
1.0
0.5
T0
T2
T4
T6
Time
Genotype
TM1
TM2
WT

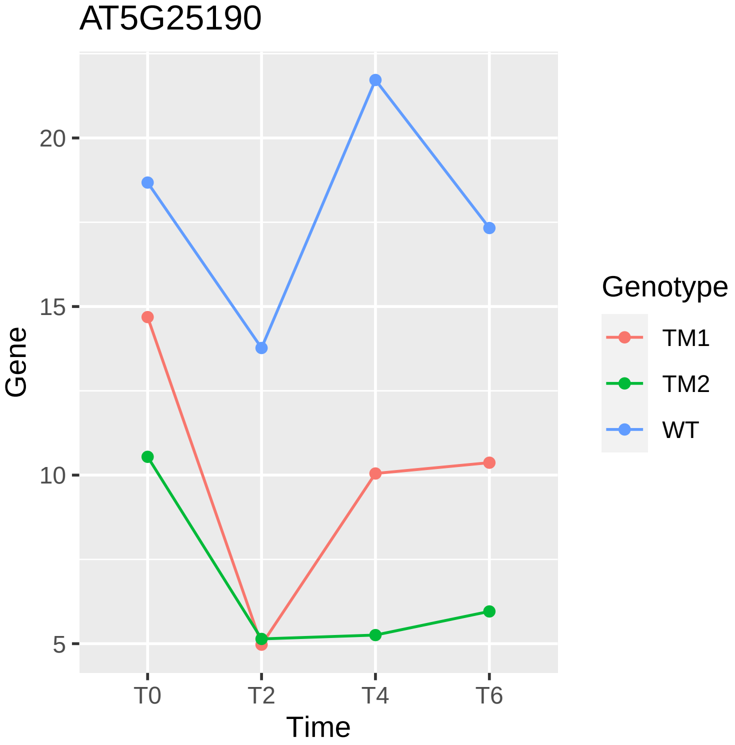

AT5G25190
Gene
Time
20
15
10
5
T0
T2
T4
T6
Genotype
TM1
TM2
WT

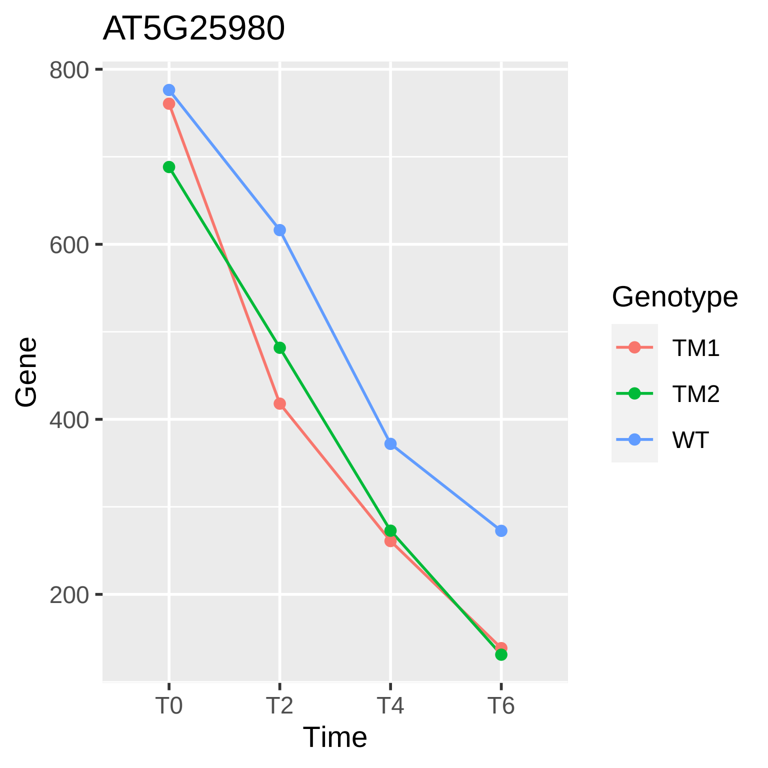

AT5G25980
800
600
400
200
Gene
T0
T2
T4
T6
Time
Genotype
TM1
TM2
WT

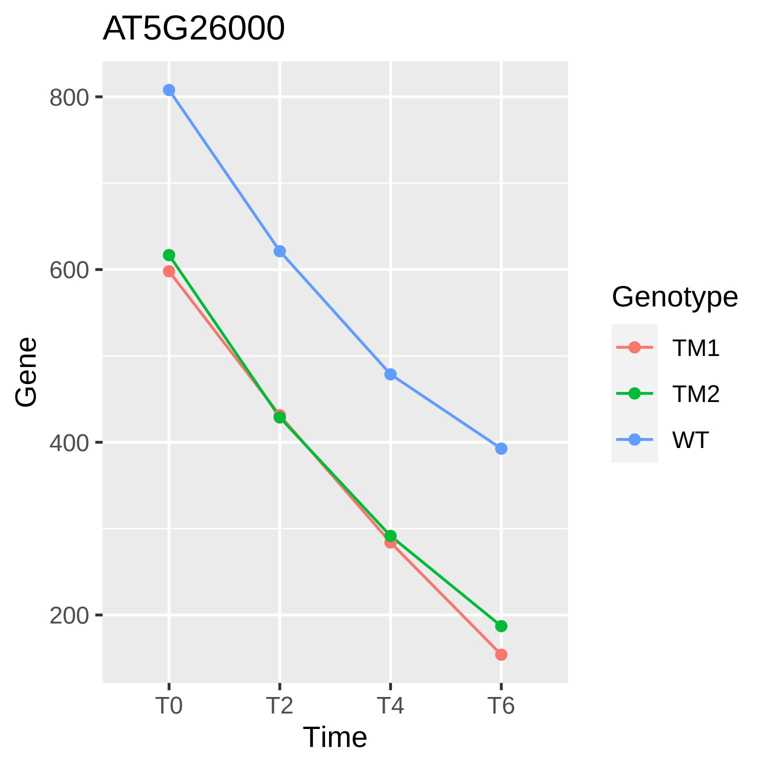
AT5G26000
800
600
400
200
Gene
T0
T2
T4
T6
Time
Genotype
TM1
TM2
WT

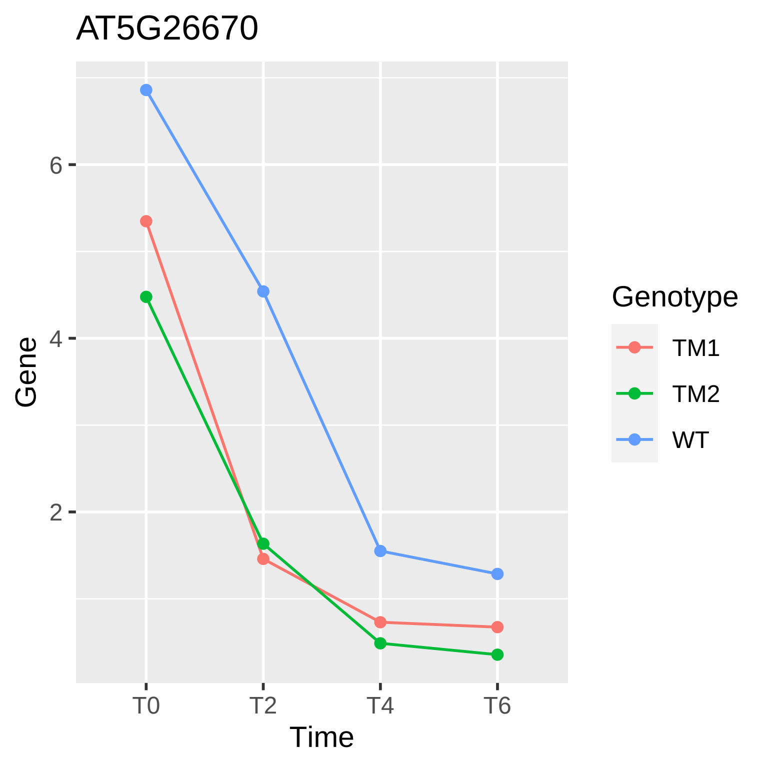
AT5G26670
Gene
6
4
2
T0
T2
T4
T6
Time
Genotype
TM1
TM2
WT

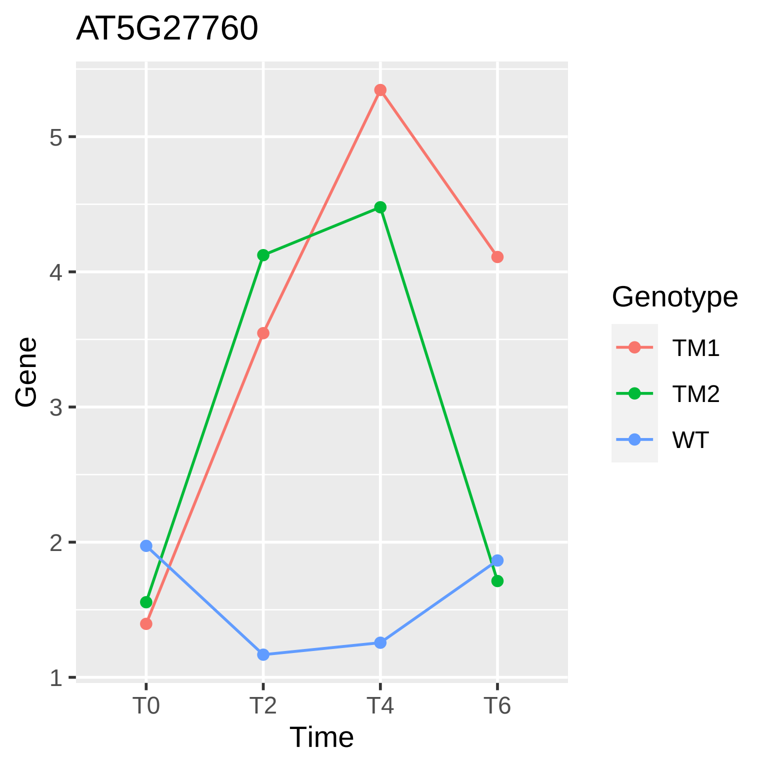
AT5G27760
Gene
5
4
3
2
1
T0
T2
T4
T6
Time
Genotype
TM1
TM2
WT

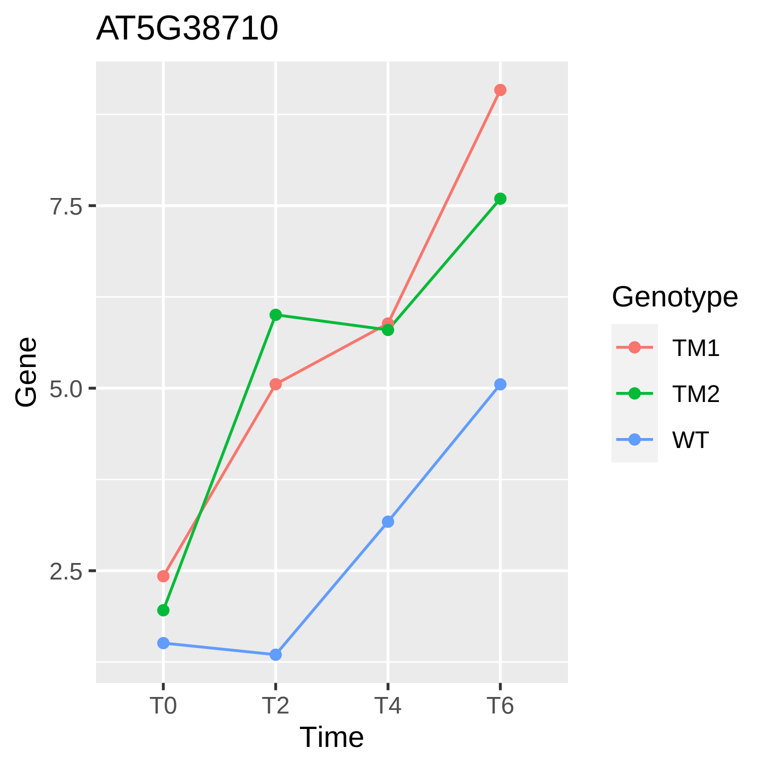

AT5G38710
Gene
7.5
5.0
2.5
T0
T2
T4
T6
Time
Genotype
TM1
TM2
WT

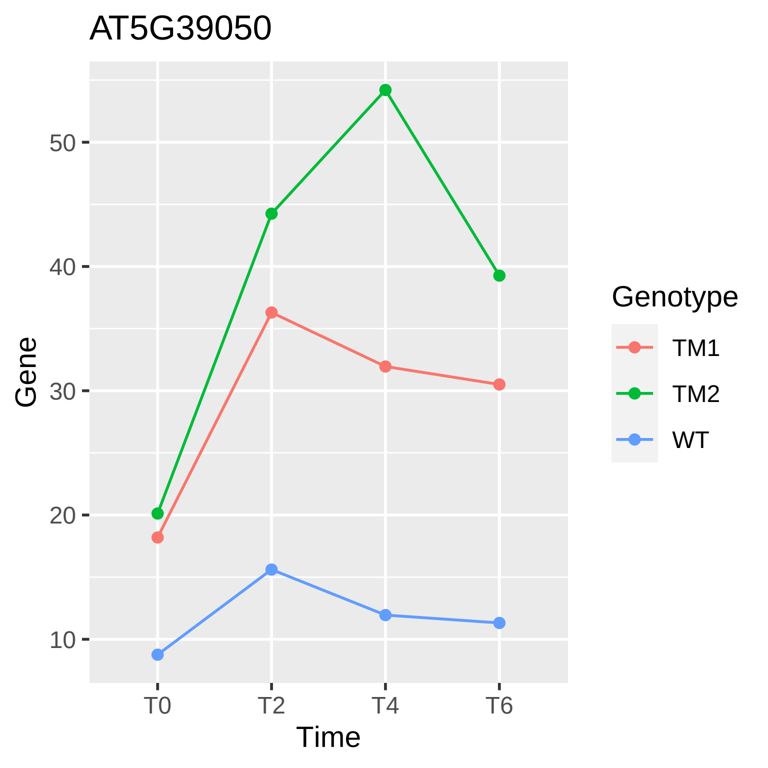

AT5G39050
Gene
50
40
30
20
10
T0
T2
T4
T6
Time
Genotype
TM1
TM2
WT

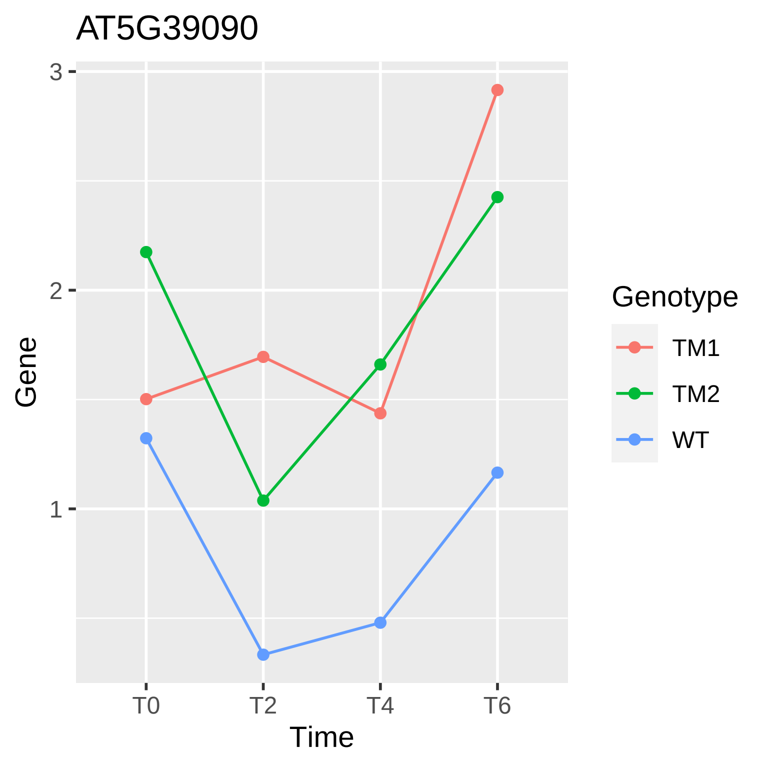

AT5G39090
Gene
3
2
1
T0
T2
T4
T6
Time
Genotype
TM1
TM2
WT

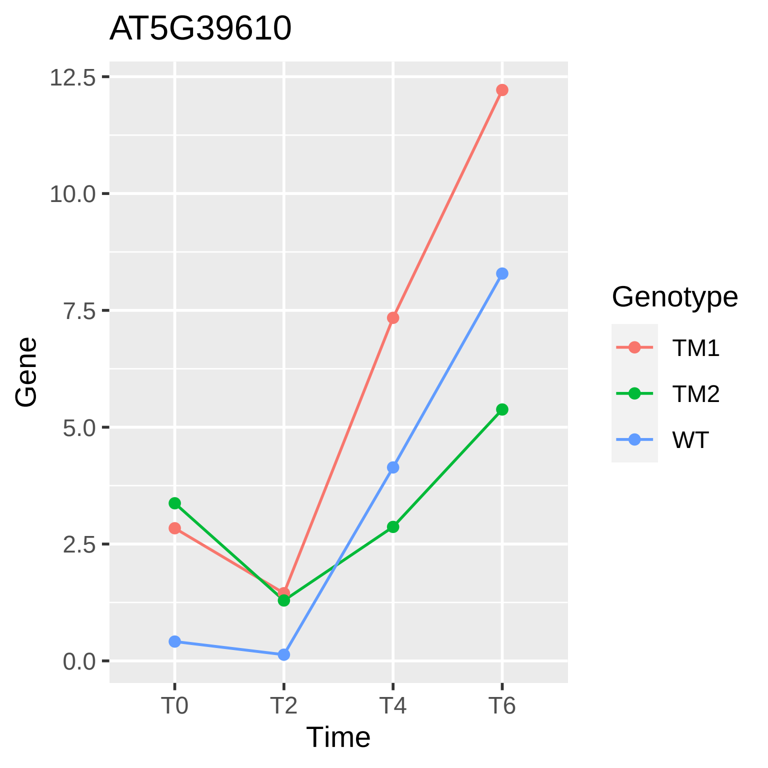
AT5G39610
12.5
10.0
7.5
5.0
2.5
0.0
Gene
T0
T2
T4
T6
Time
Genotype
TM1
TM2
WT

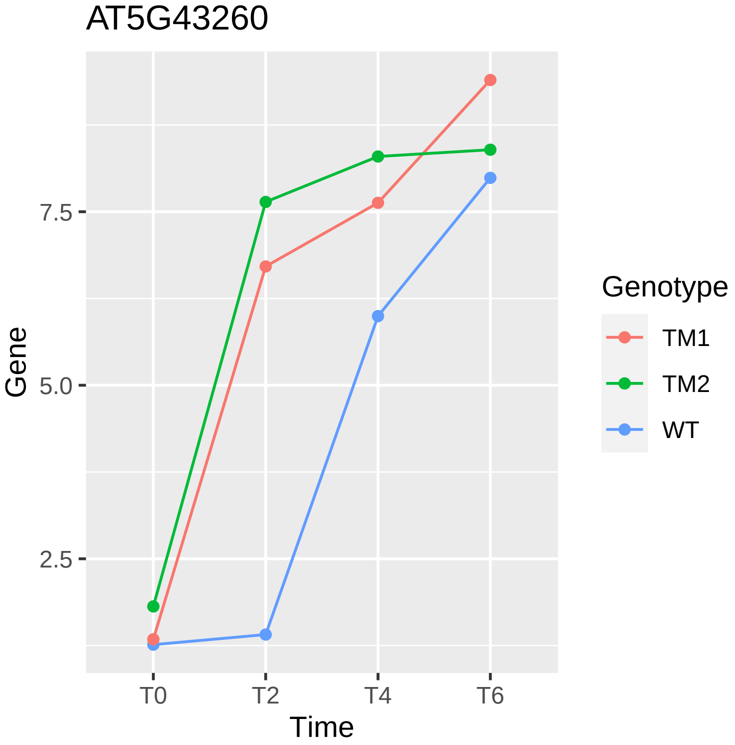
AT5G43260
Gene
7.5
5.0
2.5
T0
T2
T4
T6
Time
Genotype
TM1
TM2
WT

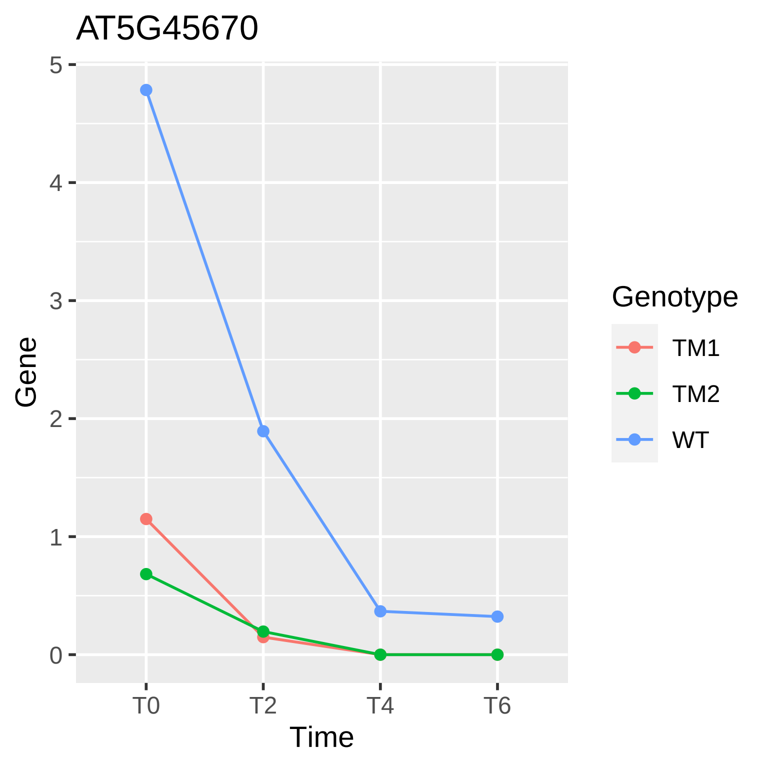
AT5G45670
Gene
0
1
2
3
4
5
T0
T2
T4
T6
Time
Genotype
TM1
TM2
WT

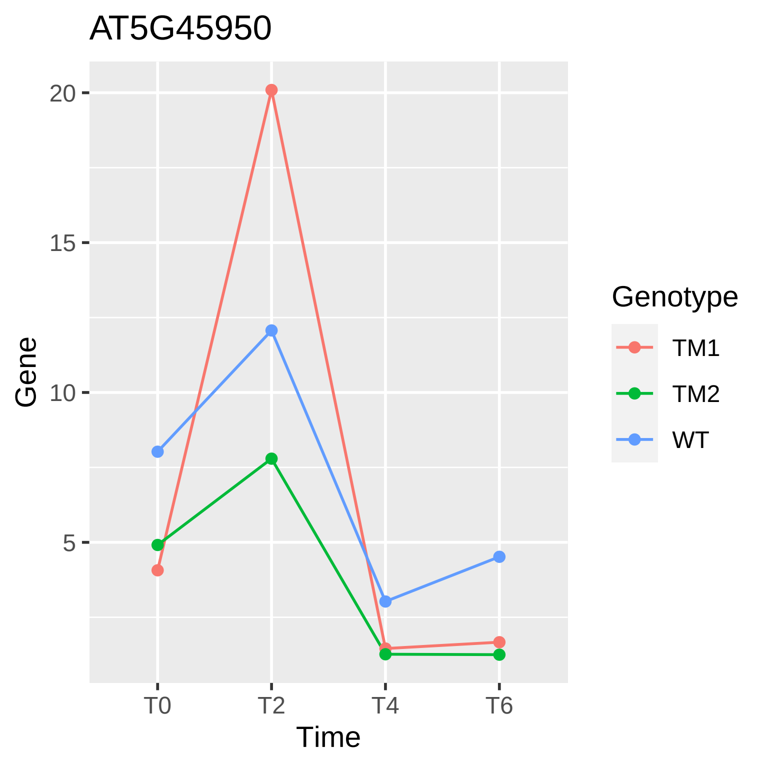
AT5G45950
20
15
10
5
Gene
T0
T2
T4
T6
Time
Genotype
TM1
TM2
WT

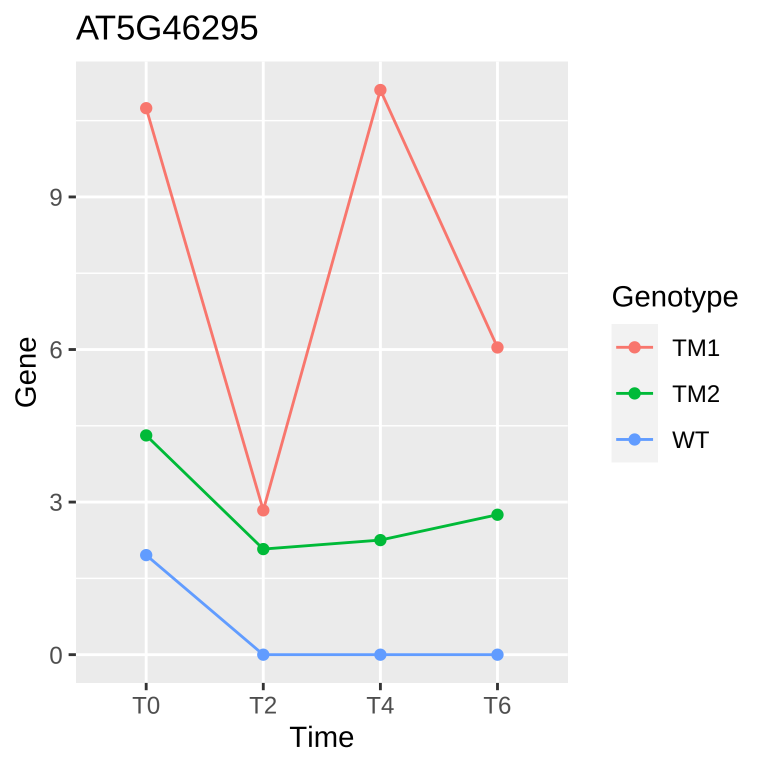

AT5G46295
Gene
9
6
3
0
T0
T2
T4
T6
Time
Genotype
TM1
TM2
WT

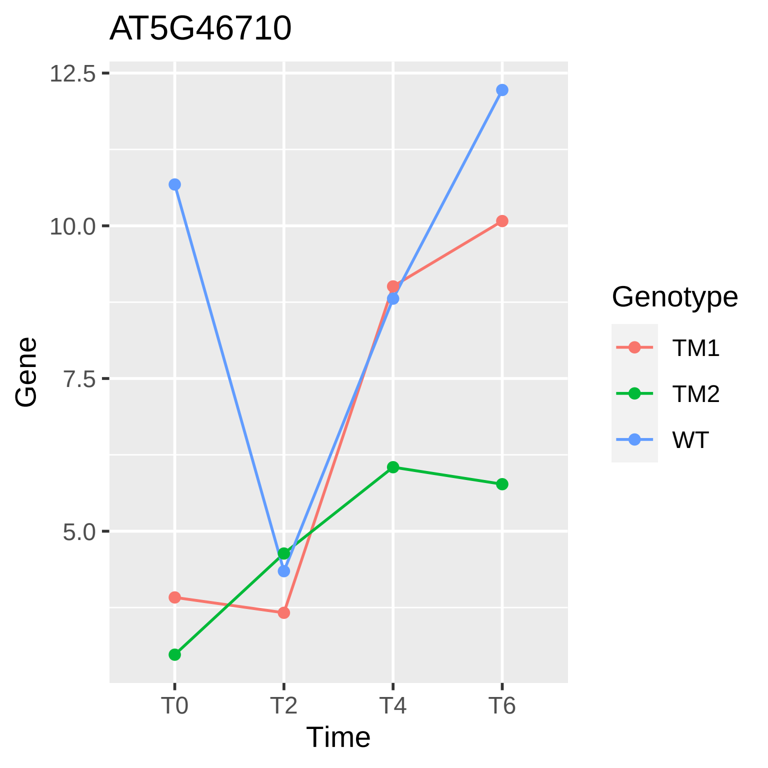
AT5G46710
12.5
10.0
7.5
5.0
Gene
T0
T2
T4
T6
Time
Genotype
TM1
TM2
WT

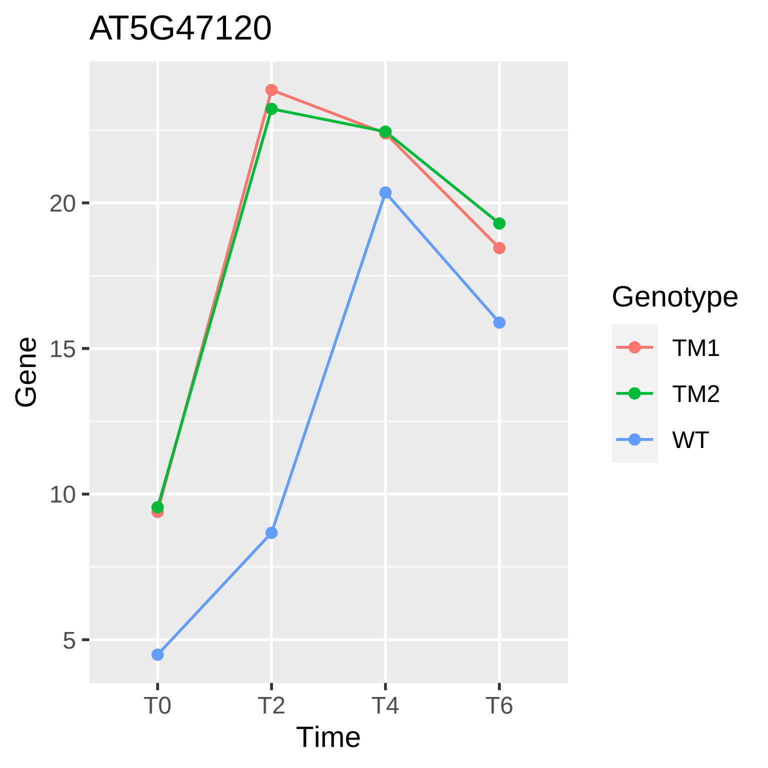
AT5G47120
Gene
20
15
10
5
T0
T2
T4
T6
Time
Genotype
TM1
TM2
WT

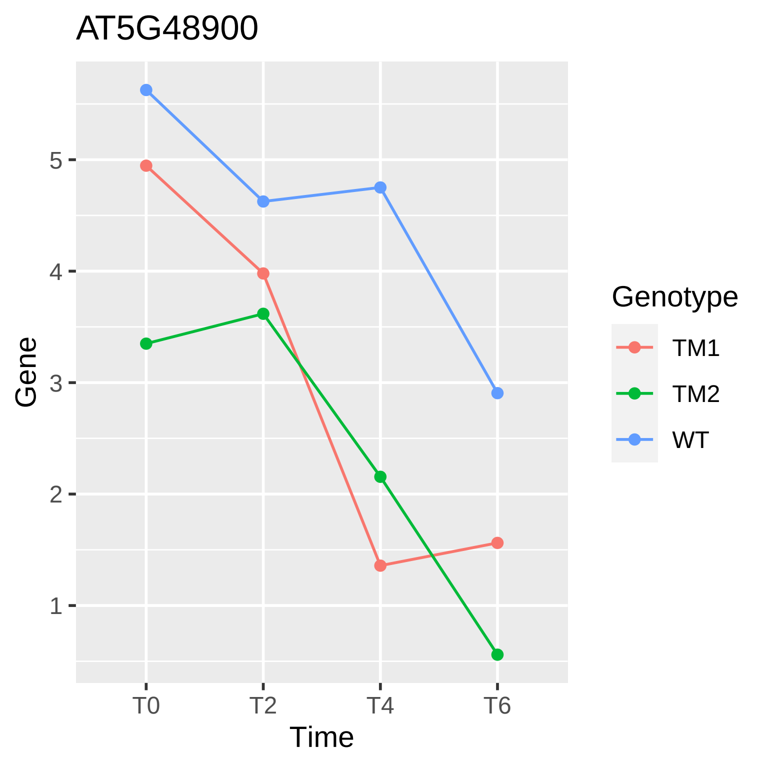
AT5G48900
Gene
5
4
3
2
1
T0
T2
T4
T6
Time
Genotype
TM1
TM2
WT

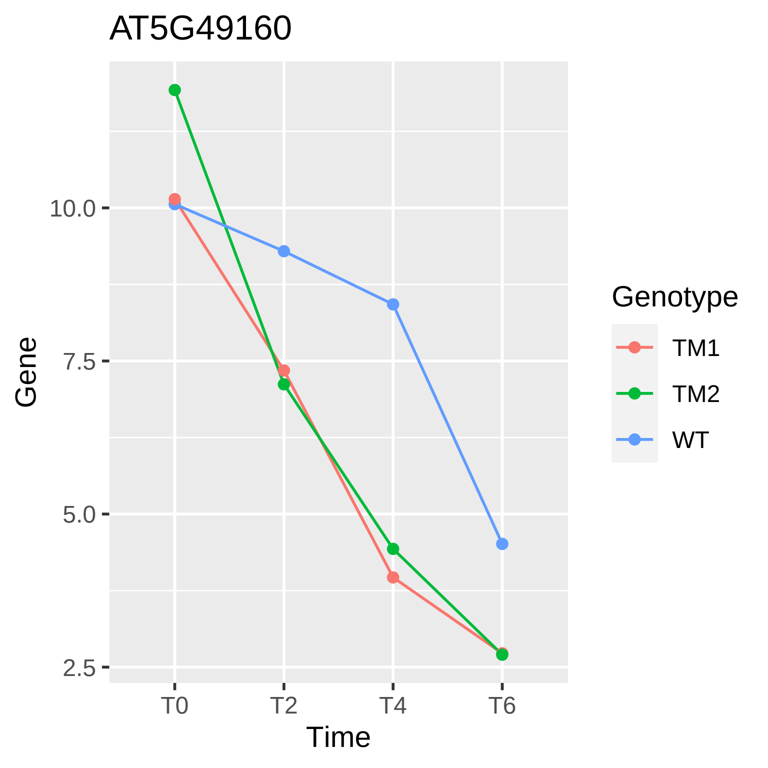
AT5G49160
Gene
10.0
7.5
5.0
2.5
T0
T2
T4
T6
Time
Genotype
TM1
TM2
WT

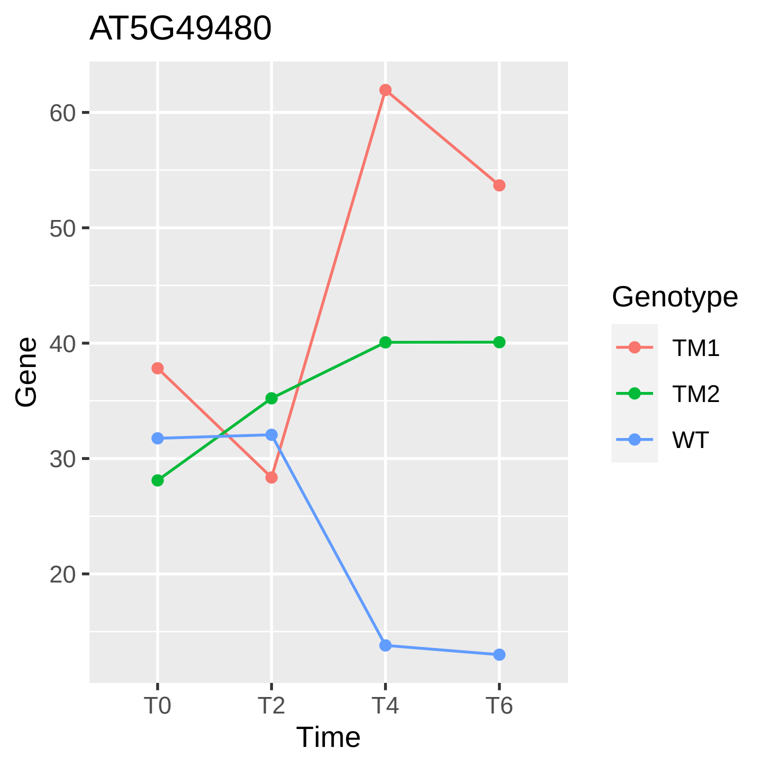
AT5G49480
Gene
60
50
40
30
20
T0
T2
T4
T6
Time
Genotype
TM1
TM2
WT

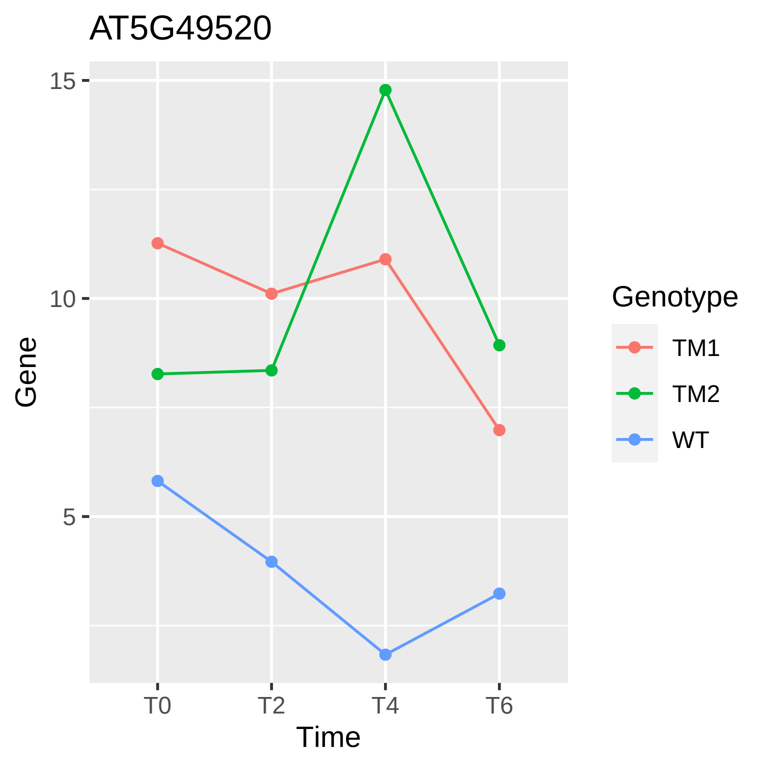
AT5G49520
15
10
5
Gene
T0
T2
T4
T6
Time
Genotype
TM1
TM2
WT

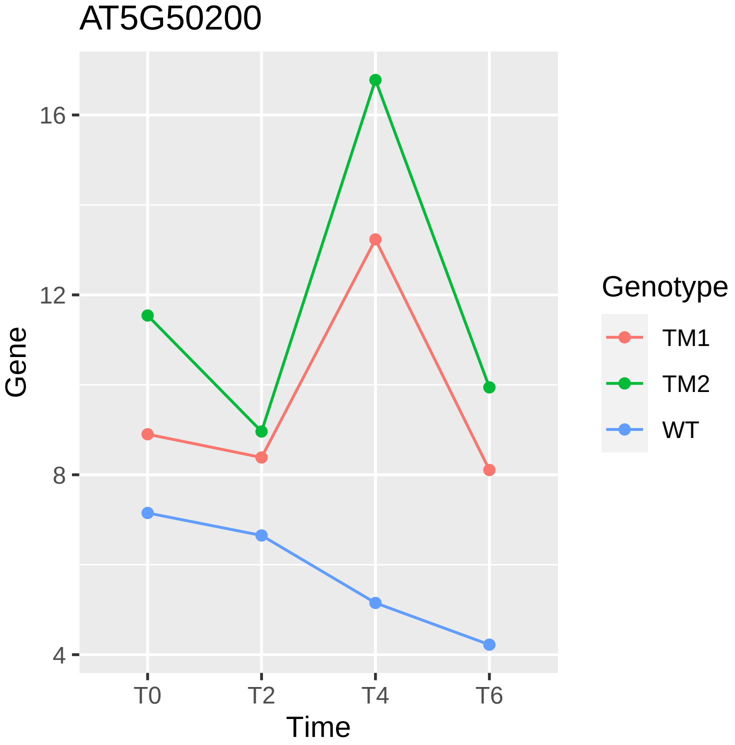
AT5G50200
Gene
16
12
8
4
T0
T2
T4
T6
Time
Genotype
TM1
TM2
WT

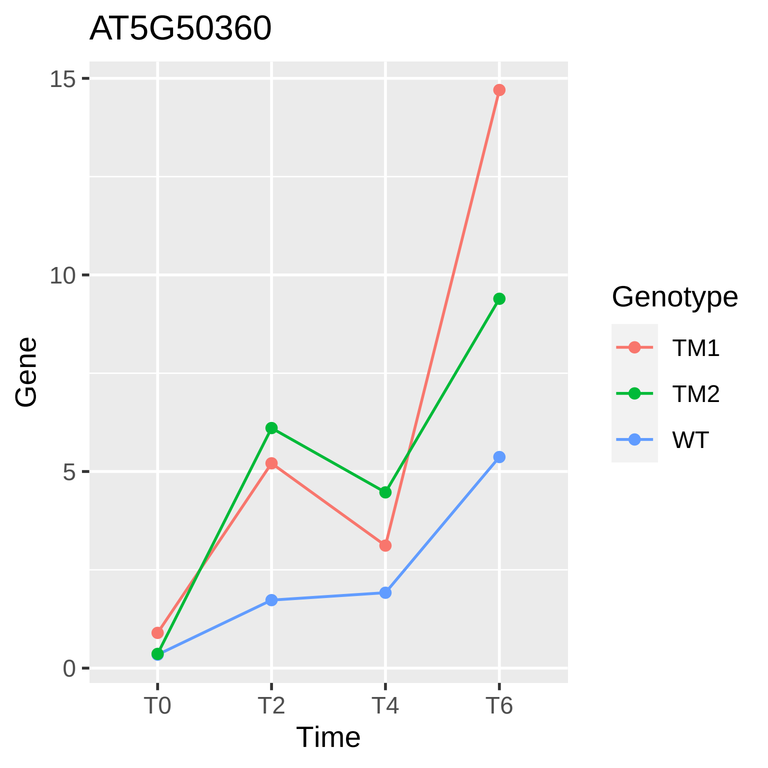

AT5G50360
15
10
5
0
Gene
T0
T2
T4
T6
Time
Genotype
TM1
TM2
WT

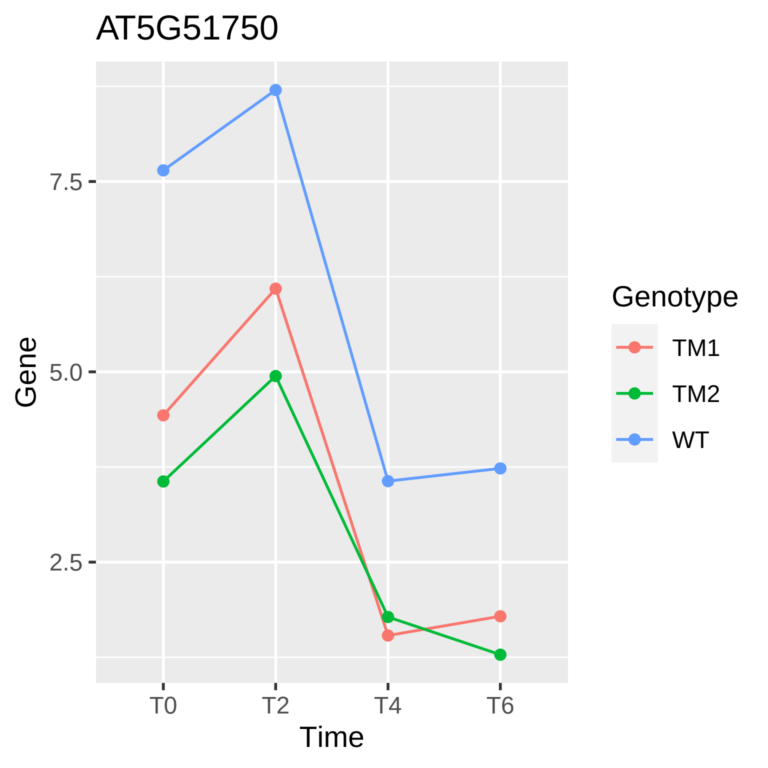
AT5G51750
Gene
7.5
5.0
2.5
T0
T2
T4
T6
Time
Genotype
TM1
TM2
WT

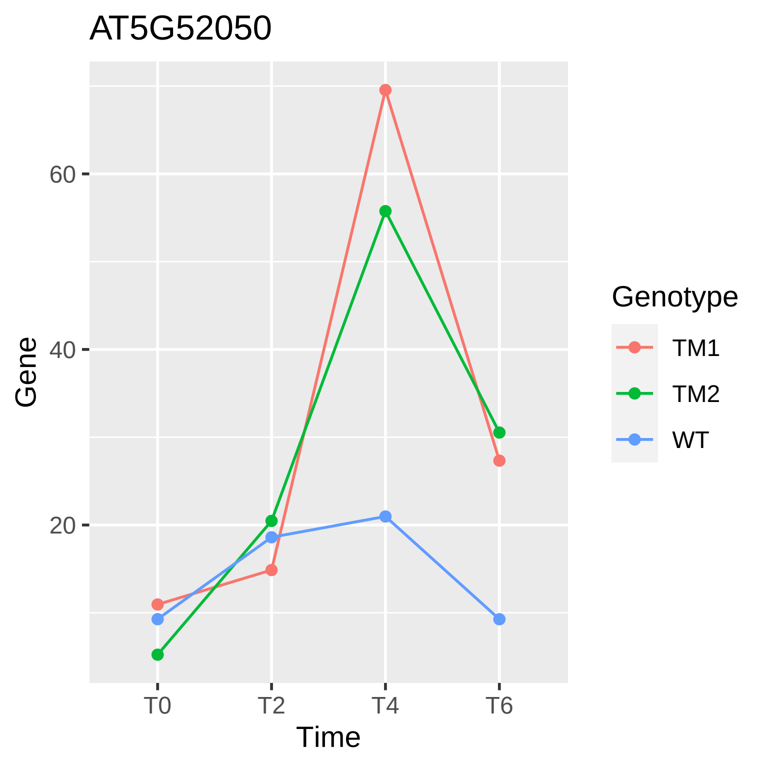

AT5G52050
Gene
60
40
20
T0
T2
T4
T6
Time
Genotype
TM1
TM2
WT

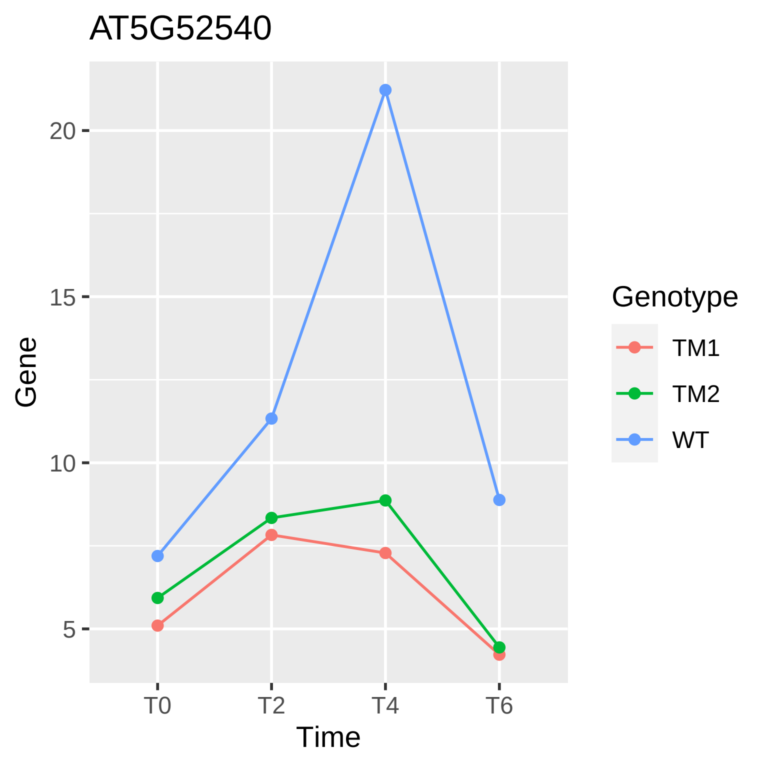

AT5G52540
Gene
Time
T0
T2
T4
T6
5
10
15
20
Genotype
TM1
TM2
WT

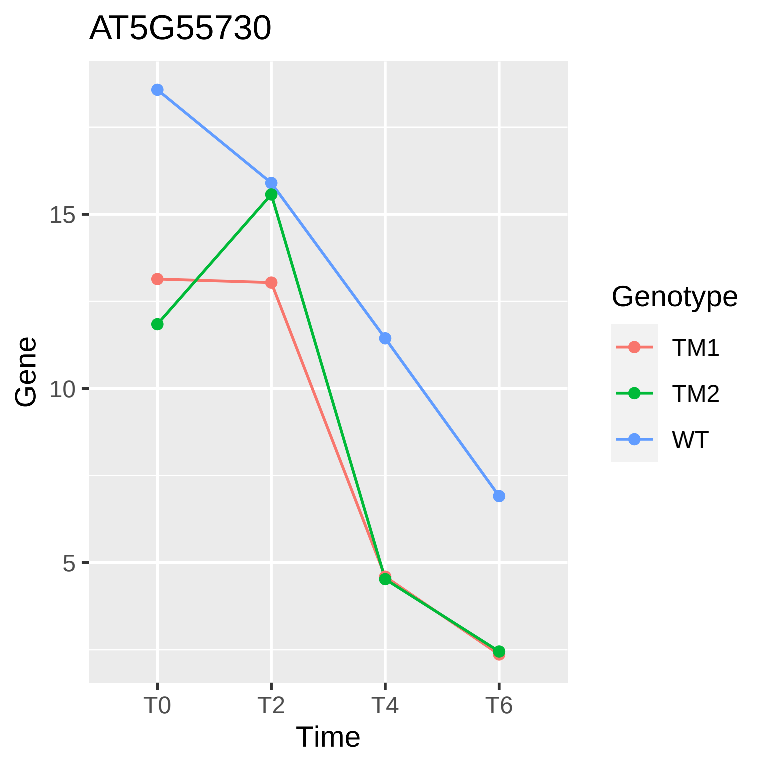
AT5G55730
Gene
15
10
5
T0
T2
T4
T6
Time
Genotype
TM1
TM2
WT

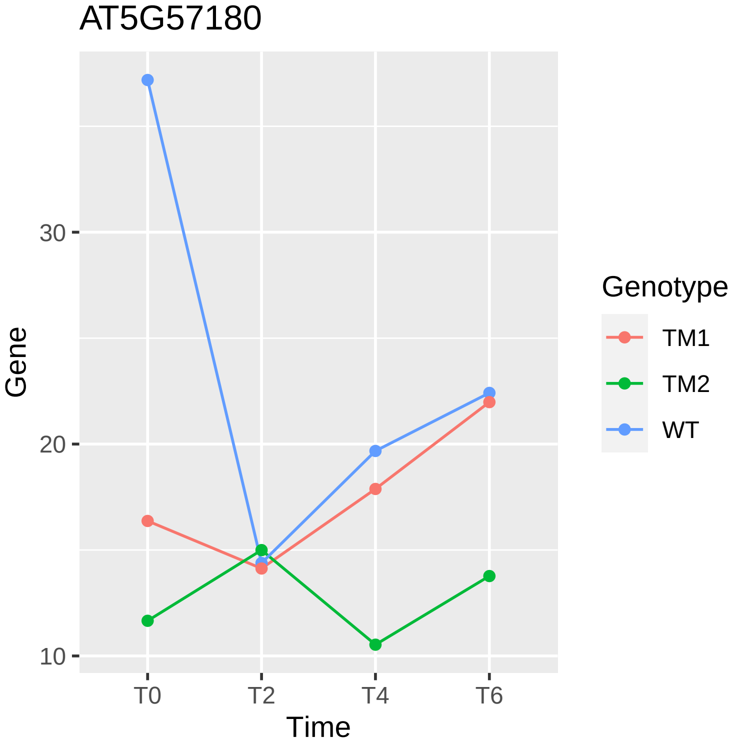
AT5G57180
Gene
30
20
10
T0
T2
T4
T6
Time
Genotype
TM1
TM2
WT

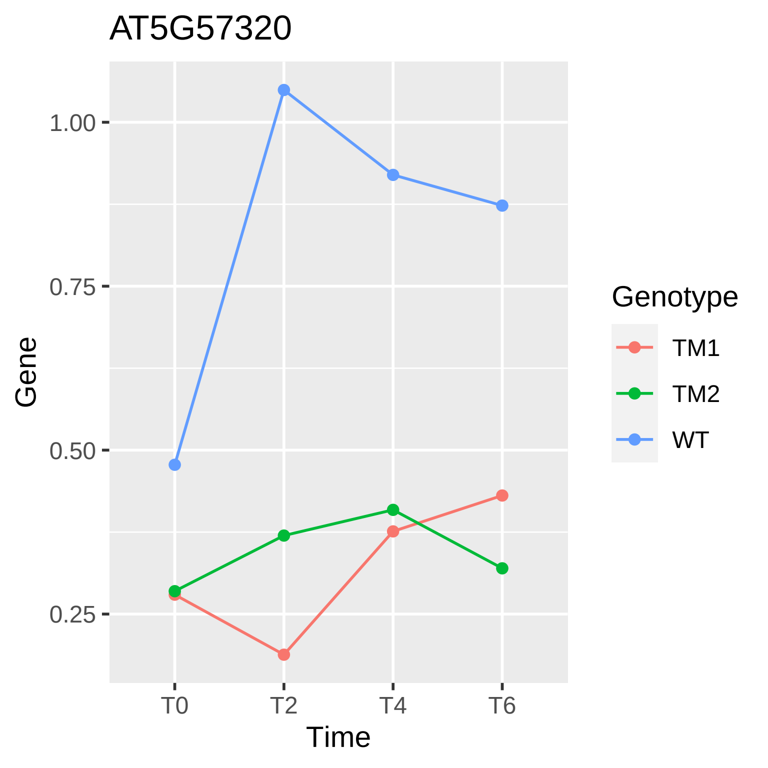
AT5G57320
Gene
1.00
0.75
0.50
0.25
T0
T2
T4
T6
Time
Genotype
TM1
TM2
WT

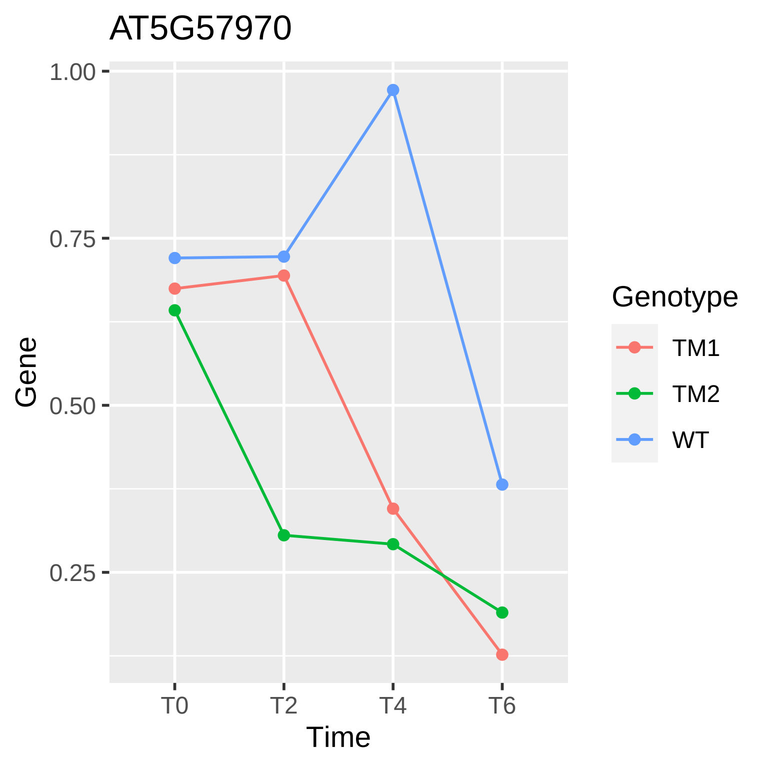

AT5G57970
1.00
0.75
0.50
0.25
Gene
T0
T2
T4
T6
Time
Genotype
TM1
TM2
WT

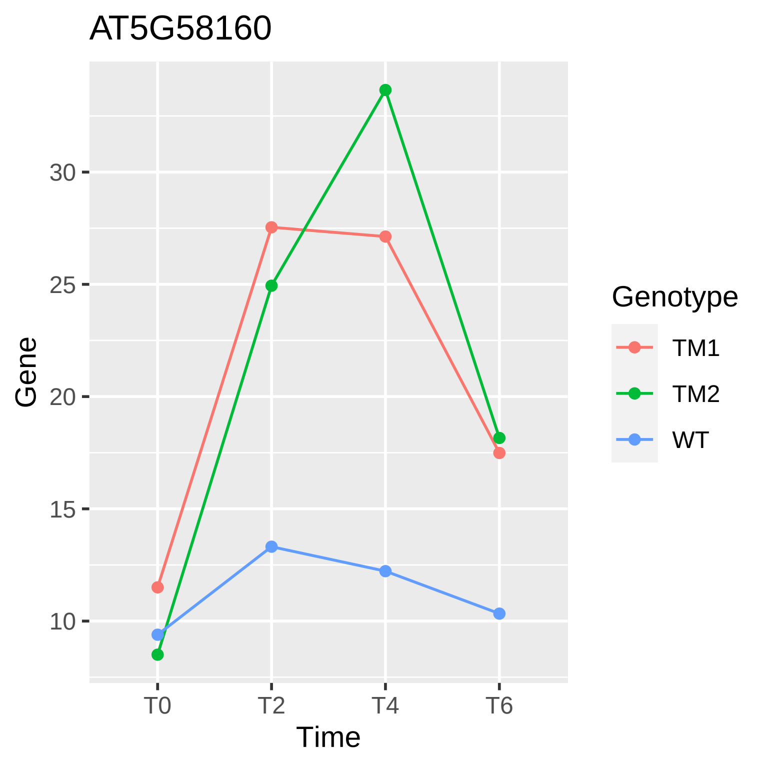

AT5G58160
Gene
30
25
20
15
10
T0
T2
T4
T6
Time
Genotype
TM1
TM2
WT

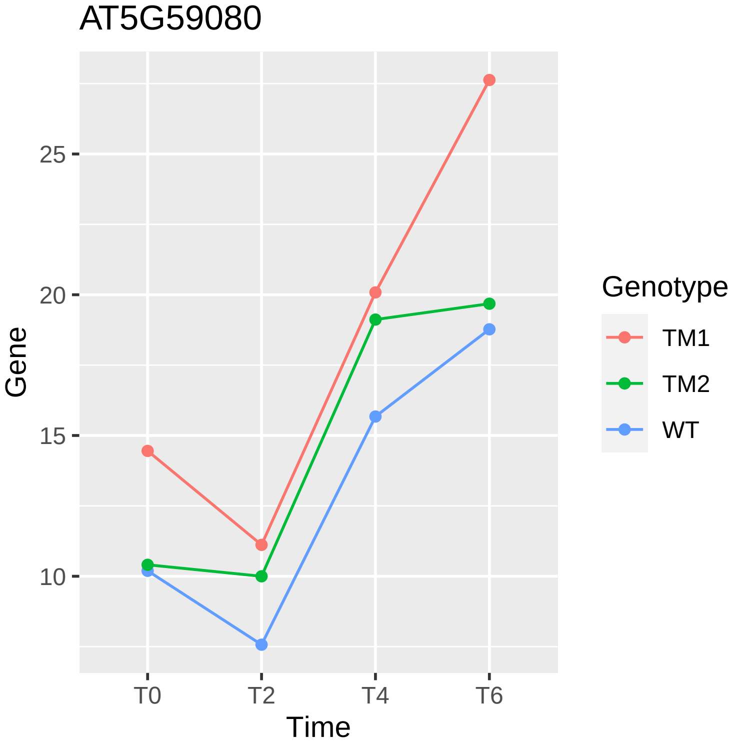

AT5G59080
Gene
25
20
15
10
T0
T2
T4
T6
Time
Genotype
TM1
TM2
WT

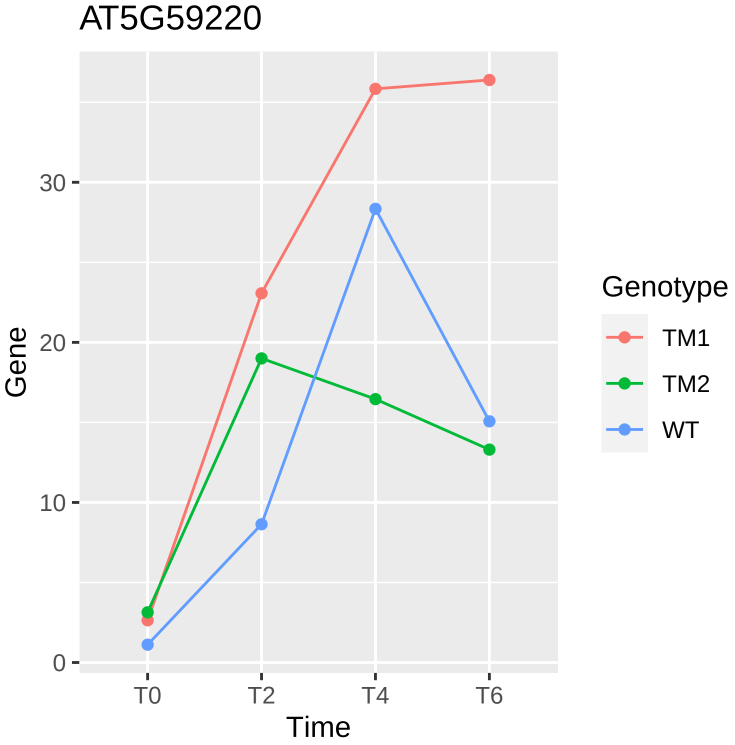

AT5G59220
Gene
0
10
20
30
T0
T2
T4
T6
Time
Genotype
TM1
TM2
WT

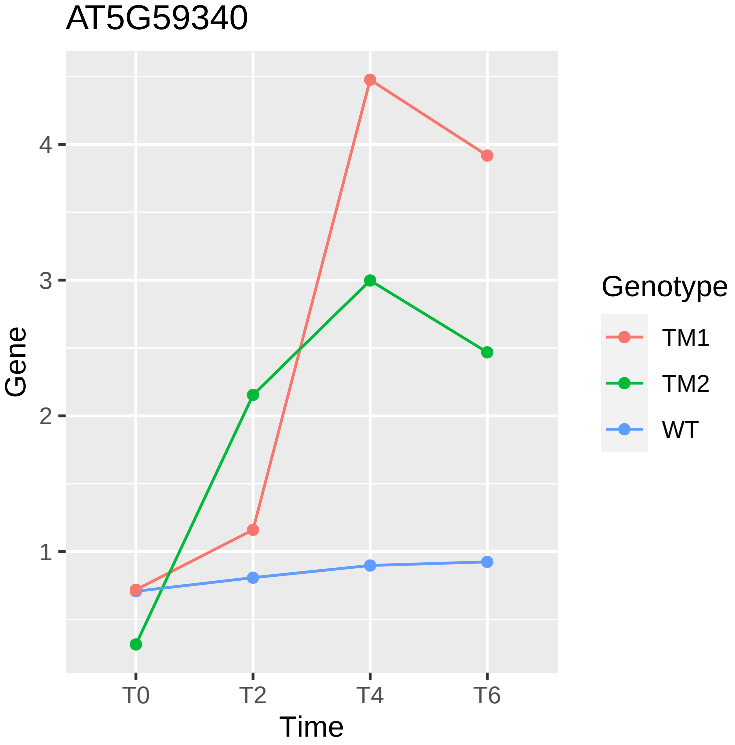
AT5G59340
Gene
1
2
3
4
T0
T2
T4
T6
Time
Genotype
TM1
TM2
WT

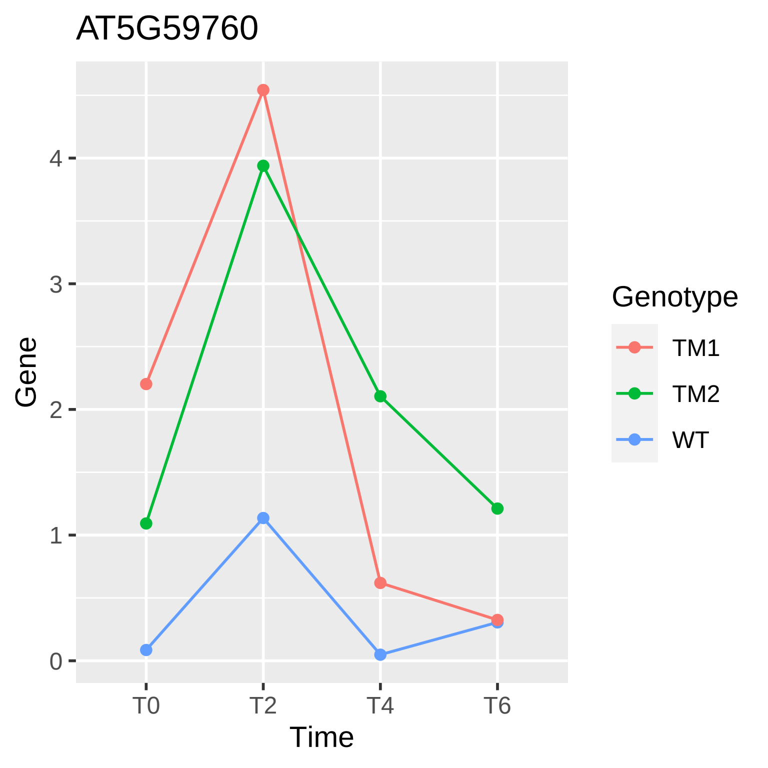
AT5G59760
Gene
0
1
2
3
4
T0
T2
T4
T6
Time
Genotype
TM1
TM2
WT

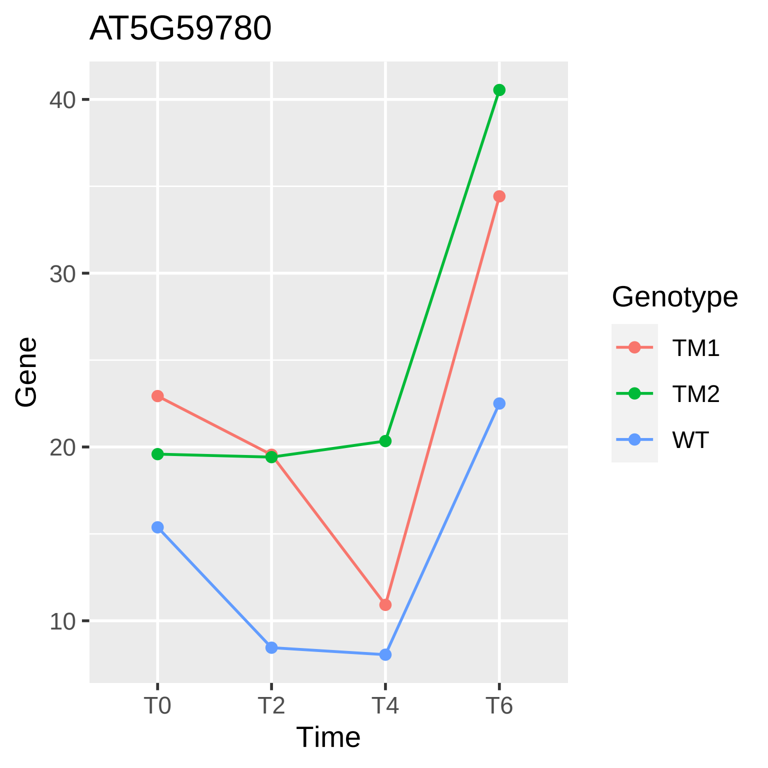

AT5G59780
40
30
20
10
Gene
T0
T2
T4
T6
Time
Genotype
TM1
TM2
WT

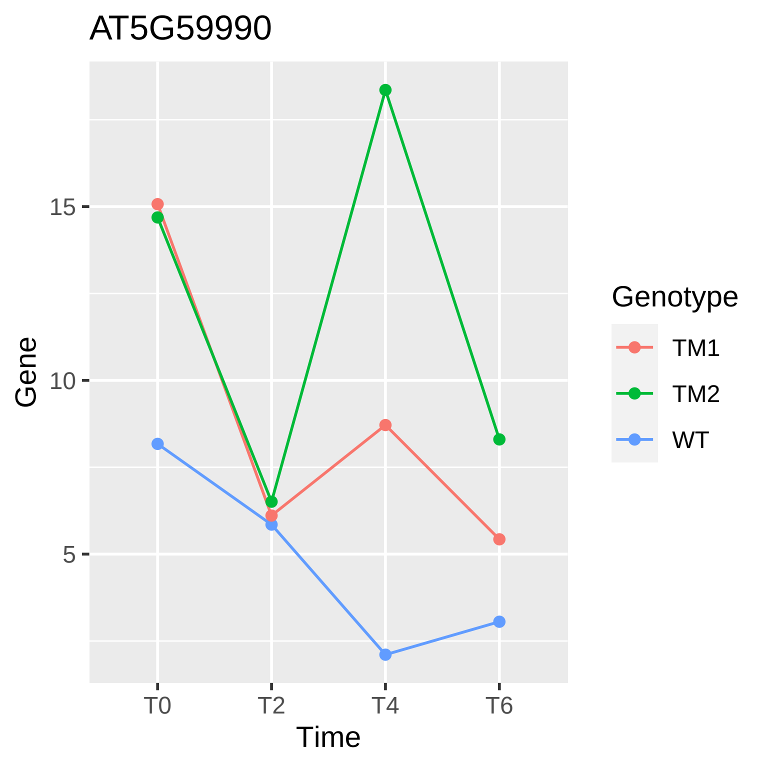
AT5G59990
Gene
15
10
5
T0
T2
T4
T6
Time
Genotype
TM1
TM2
WT

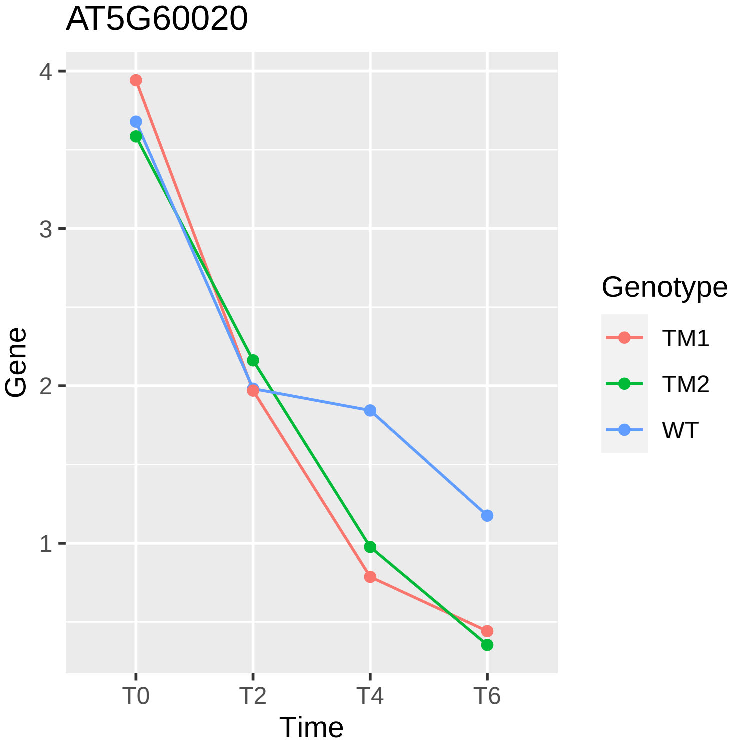
AT5G60020
Gene
4
3
2
1
T0
T2
T4
T6
Time
Genotype
TM1
TM2
WT

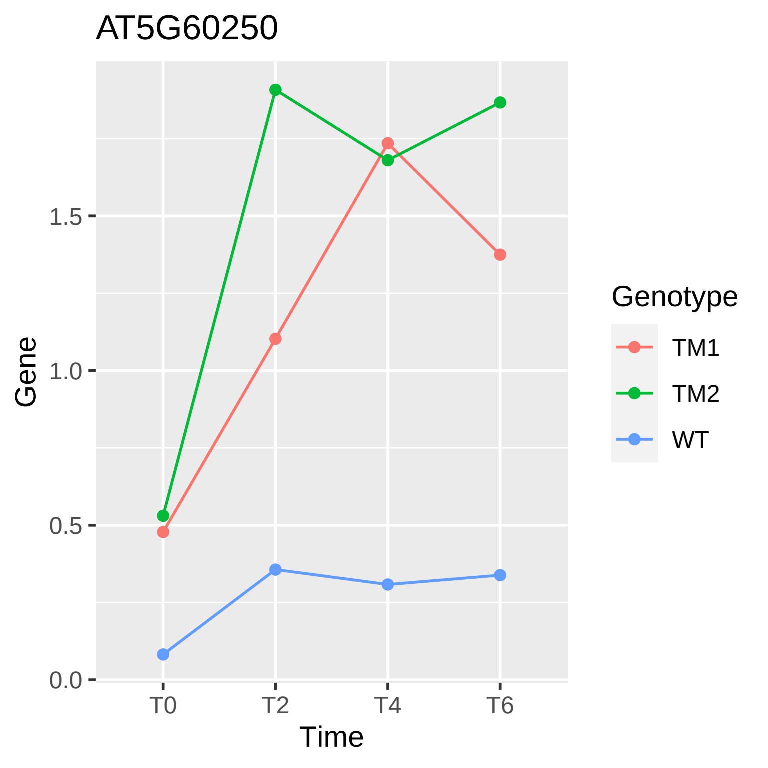
AT5G60250
Gene
1.5
1.0
0.5
0.0
T0
T2
T4
T6
Time
Genotype
TM1
TM2
WT

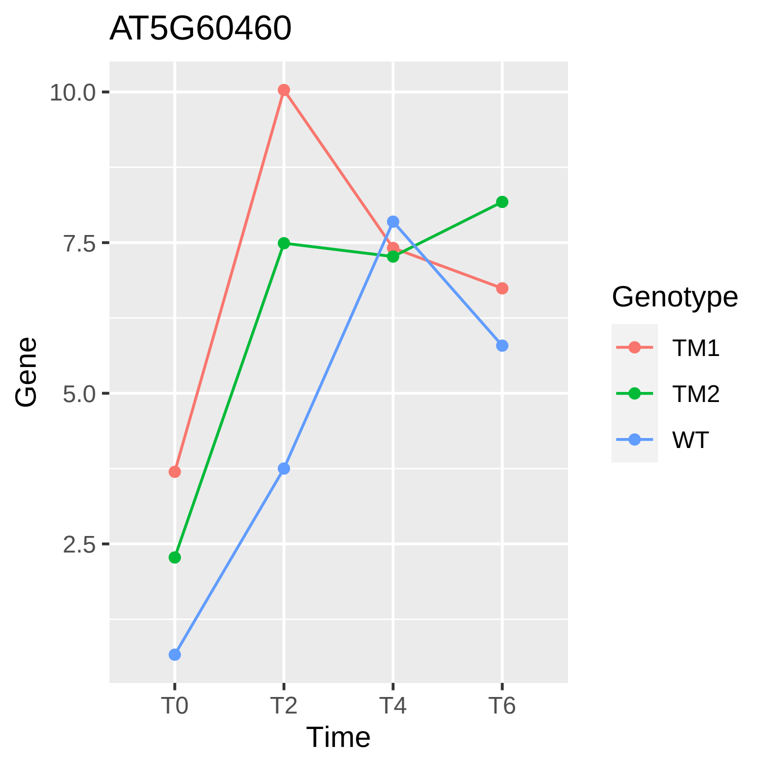
AT5G60460
10.0
7.5
5.0
2.5
Gene
T0
T2
T4
T6
Time
Genotype
TM1
TM2
WT

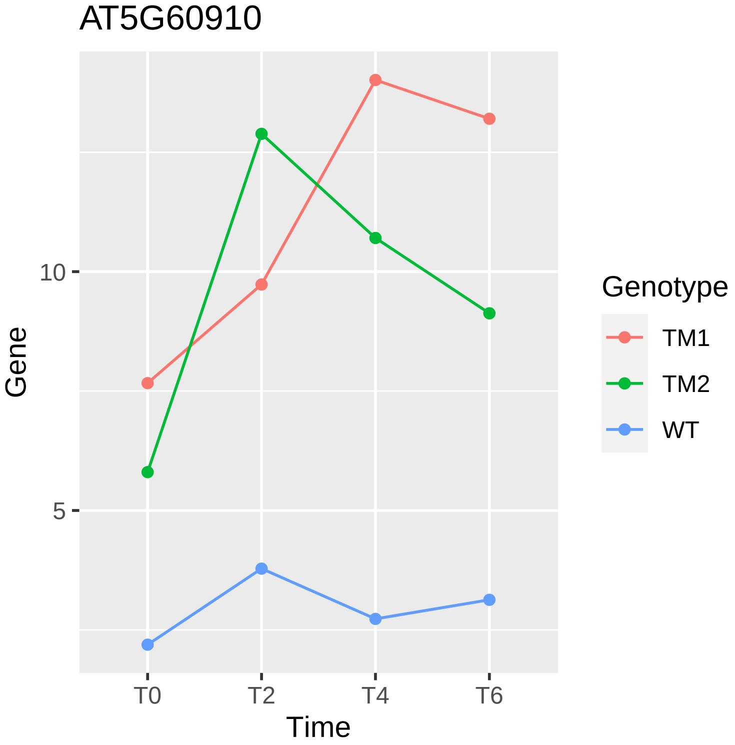
AT5G60910
10
5
Gene
T0
T2
T4
T6
Time
Genotype
TM1
TM2
WT

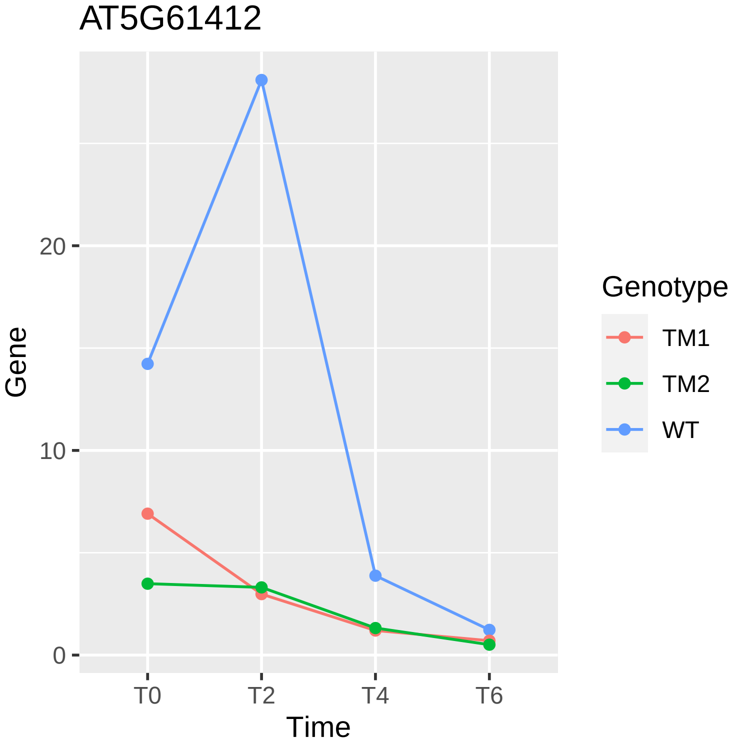
AT5G61412
Gene
20
10
0
T0
T2
T4
T6
Time
Genotype
TM1
TM2
WT

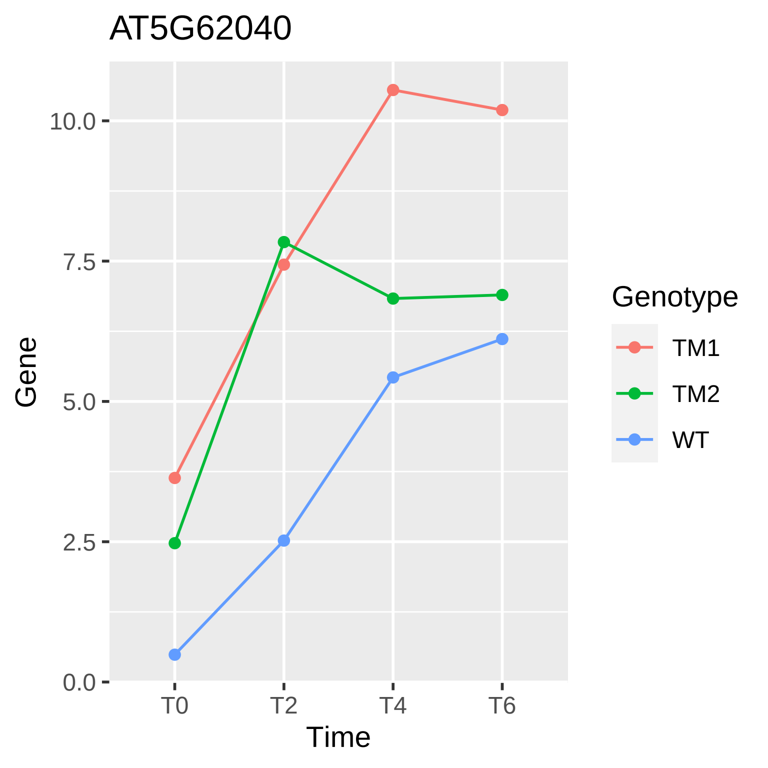
AT5G62040
Gene
10.0
7.5
5.0
2.5
0.0
T0
T2
T4
T6
Time
Genotype
TM1
TM2
WT

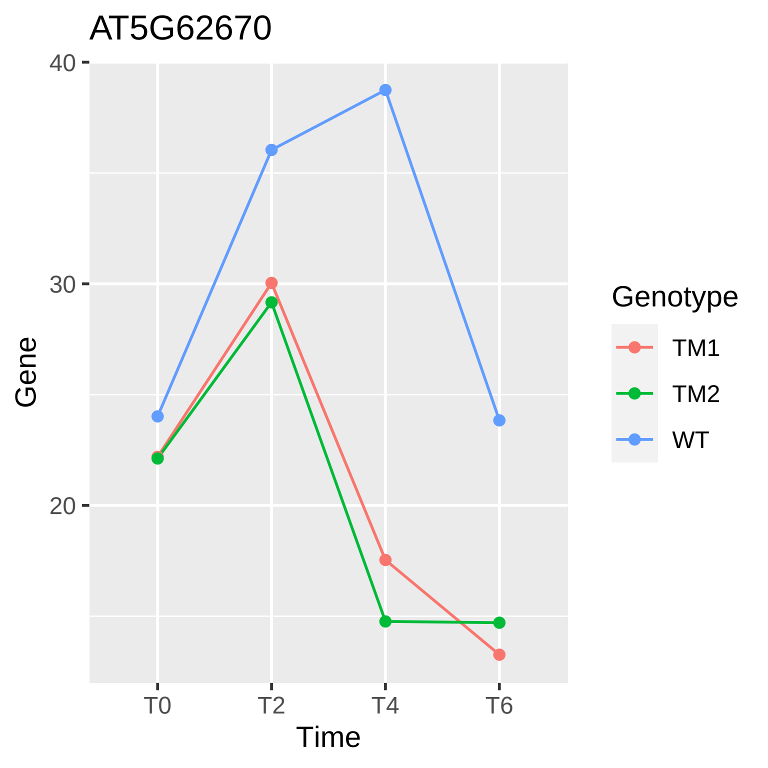

AT5G62670
40
30
20
Gene
T0
T2
T4
T6
Time
Genotype
TM1
TM2
WT

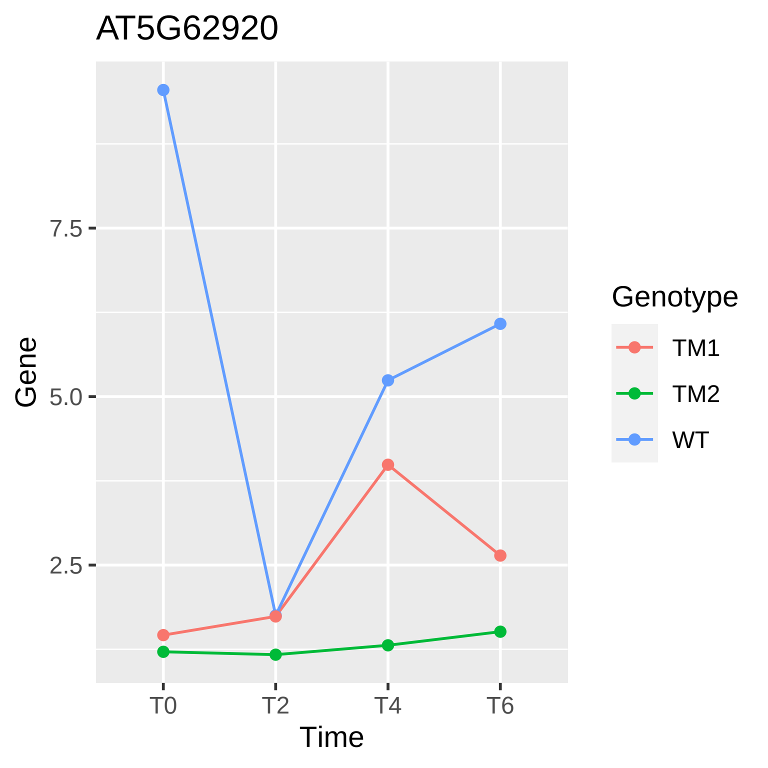

AT5G62920
Gene
7.5
5.0
2.5
T0
T2
T4
T6
Time
Genotype
TM1
TM2
WT

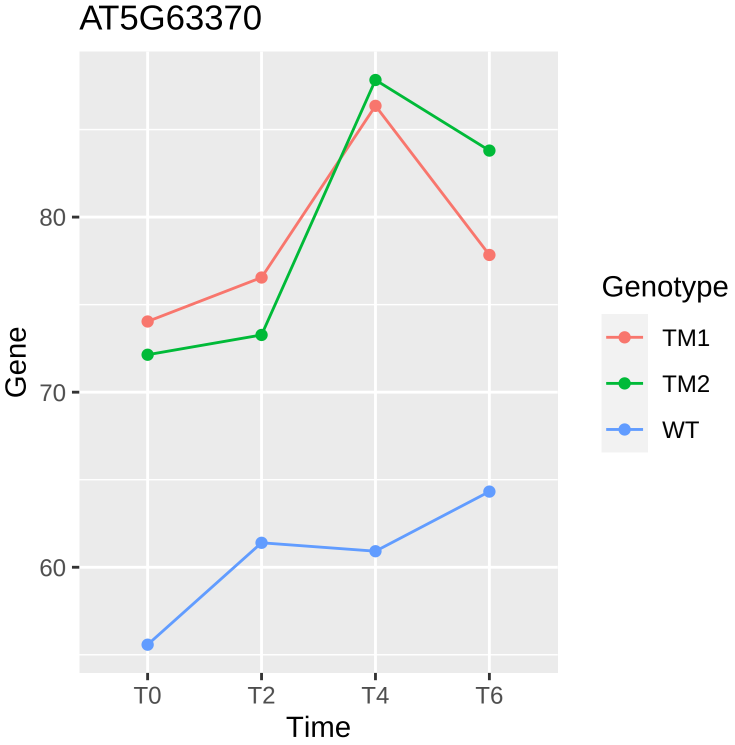
AT5G63370
Gene
80
70
60
T0
T2
T4
T6
Time
Genotype
TM1
TM2
WT

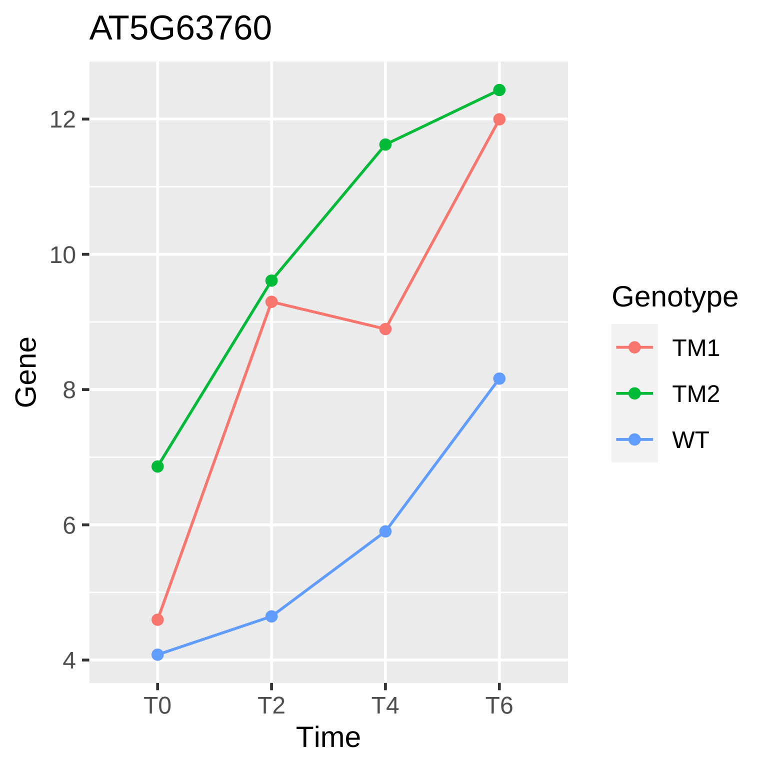

AT5G63760
Gene
12
10
8
6
4
T0
T2
T4
T6
Time
Genotype
TM1
TM2
WT

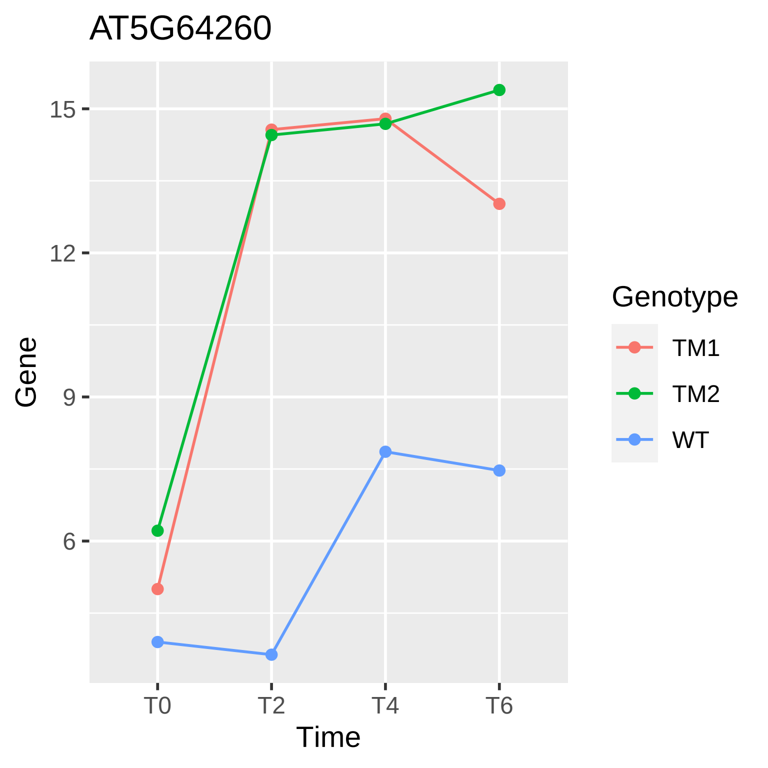
AT5G64260
Gene
15
12
9
6
T0
T2
T4
T6
Time
Genotype
TM1
TM2
WT

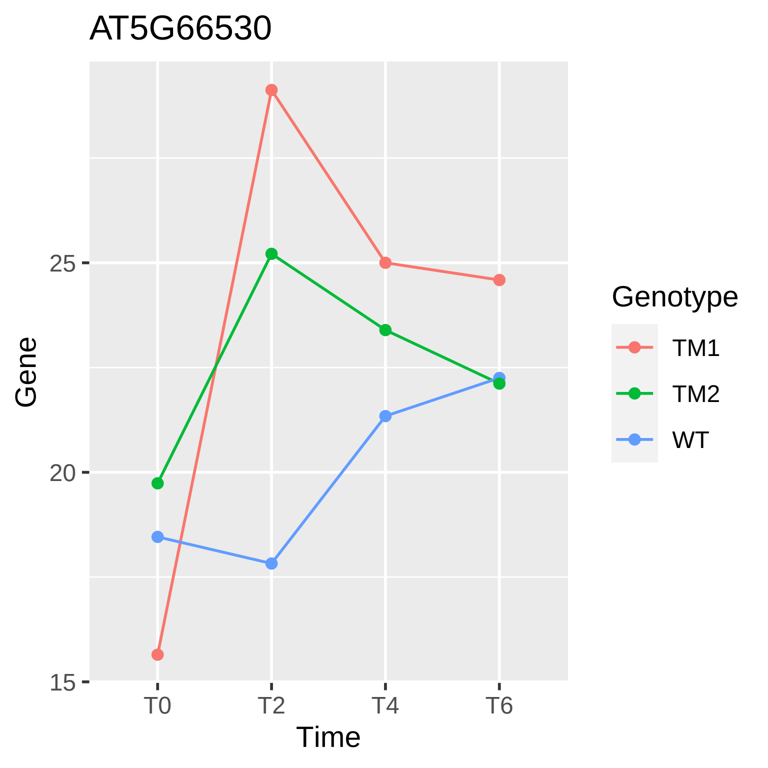
AT5G66530
Gene
25
20
15
T0
T2
T4
T6
Time
Genotype
TM1
TM2
WT

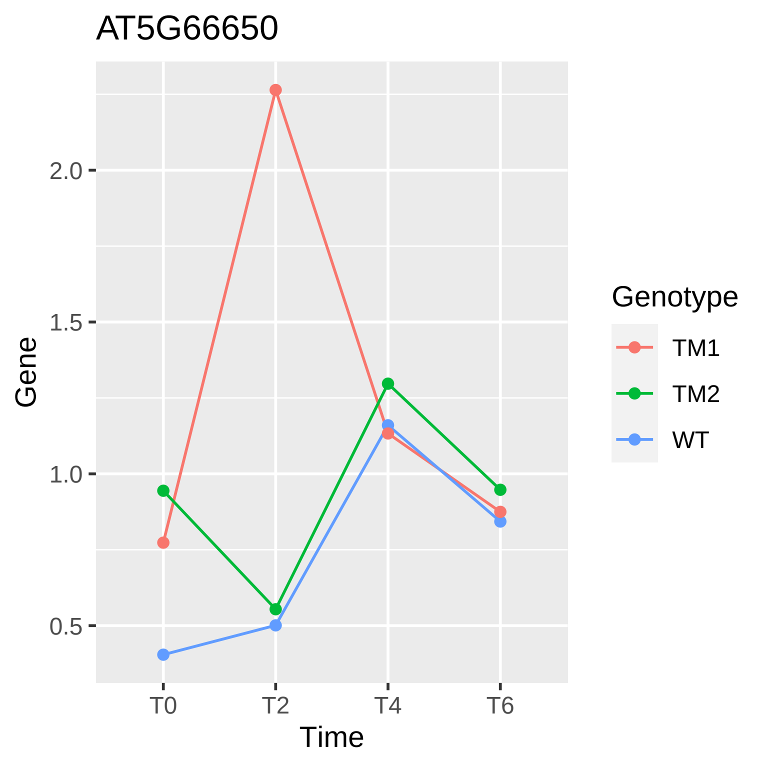
AT5G66650
Gene
2.0
1.5
1.0
0.5
T0
T2
T4
T6
Time
Genotype
TM1
TM2
WT

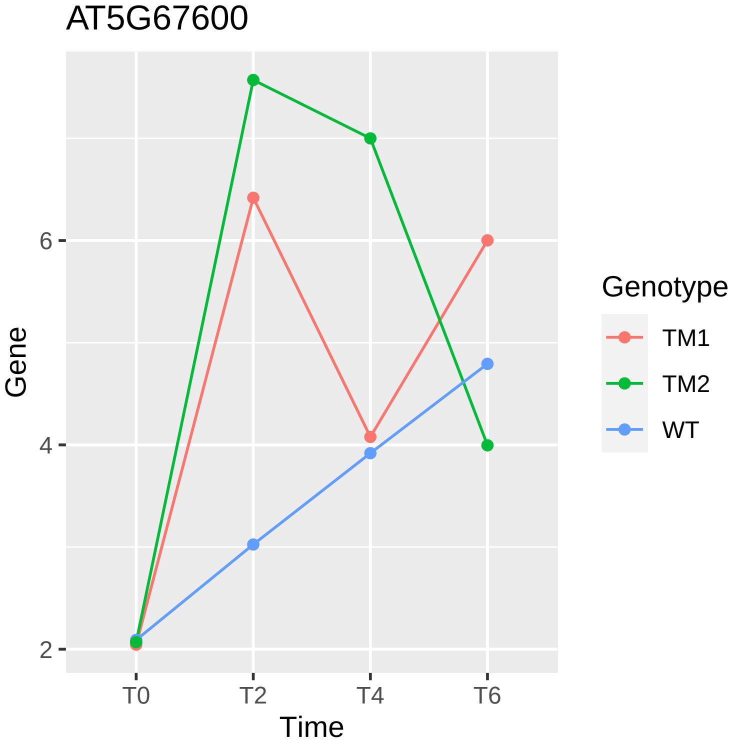
AT5G67600
Gene
2
4
6
T0
T2
T4
T6
Time
Genotype
TM1
TM2
WT

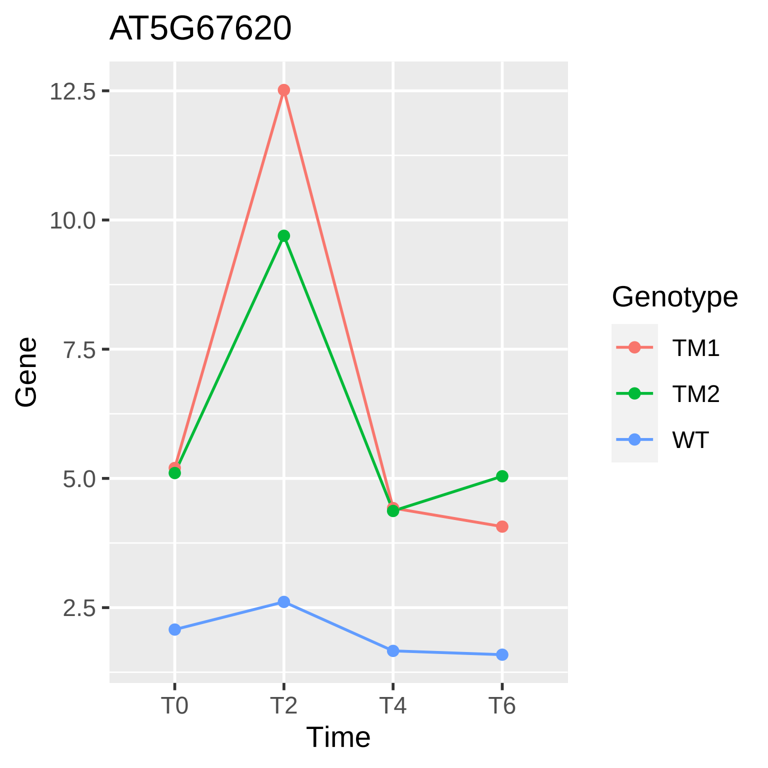
AT5G67620
12.5
10.0
7.5
5.0
2.5
Gene
T0
T2
T4
T6
Time
Genotype
TM1
TM2
WT
