## Supplementals for "Gene Expression Changes Occurring at Bolting Time are Associated with Leaf Senescence in Arabidopsis": Supplemental_Data_File_7.txt

#!/bin/bash
#input file. should be a text file from panther from the JSON option.
#Download the JSON output and paste it to a text file, make that the input here
echo Which file? On Panther, use the JSON output and copy that to a text file to use here! 
read file

#Replace every Result with a ### and Label with a @@@ so i can later select for what is in between those
gsed -e 's/result/###/g' -e 's/label/@@@/g' $file > tempgo1.txt

#select for what is in between the ### and the @@@ which is the at numbers, but excludes the GO TERM
cat -n tempgo1.txt | sed -n '/###/,/@@@/p' > tempgo2.txt

#export the go terms onto a new file
awk '{for(i=1;i<=NF;i++)if($i~/"@@@":/)print $($2)}' tempgo2.txt > tempgo3.txt

#remove everything that is not GO term name
gsed -n 's/:/&\n/;s/.*\n//p' tempgo3.txt > tempgo4.txt
#replace all spaces with _
gsed 's/ /_/g' tempgo4.txt > tempgo5.txt

#remove data to further isolate the AT numbers from tempgo2
cat -n tempgo2.txt | sed -n '/_id":/,/"id":/p' > atnumb.txt
#remove all quotes
cat -n atnumb.txt |tr -d '"' > atnumb15.txt
cat -n atnumb15.txt | sed -n '/: {/,/]},/p' > atnumb1.txt
gsed -e 's/mapped_id/@@/g' atnumb1.txt | sed -n '/@@/,/},/p' > atnumb2.txt
cat atnumb2.txt | cut -f5 -s | tr "}" " " > atnumb3.txt
cat atnumb3.txt | tr -d '\n' | tr -d " \t\n\r" | sed 's/"@@_list":{"@@"://g' > atnumb4.txt
gsed 's/@@_list:{@@://g' atnumb4.txt > atnumb5.txt
#replace brackets with a # so new lines can be added
gsed -e 's/]/###/g' atnumb5.txt  | gsed 's/###/\n/g'  > atnumb6.txt


#merge the at numbers with the go terms and make it into columns
paste tempgo5.txt atnumb6.txt  > goat.txt
cat goat.txt  | column -s, -t > goats.txt
#replace the commas with spaces
gsed 's/,/ /g' goats.txt > spacegoat.txt
#removed unclassified genes
cat spacegoat.txt | cut -f1 -d "{"  > spacegoats.txt
#assign spacegoats to neat columns
column -t spacegoats.txt >colspacegoat.txt

#berlemont wrote this to reformat the page to be fit for R
while read id
do
echo $id
fun=$(echo $id | gsed 's/\ \ */\t/g' | cut  -f 1 | tr -d '"' | gsed 's/_//' )

echo $id | gsed 's/\ \ */\t/g' | cut  -f 2- | tr -d '[' | tr '\t' '\n' | gsed "s/^/$fun\t/g" | tee -a $file.result.txt

done < colspacegoat.txt
#remove excess files
rm atnumb.txt
rm atnumb15.txt
rm atnumb1.txt
rm atnumb2.txt
rm atnumb3.txt
rm atnumb4.txt
rm atnumb5.txt
rm atnumb6.txt
rm tempgo1.txt
rm tempgo2.txt
rm tempgo3.txt
rm tempgo4.txt
rm tempgo5.txt
rm goats.txt
rm spacegoats.txt
rm spacegoat.txt
rm colspacegoat.txt


#Export this to R and merge with the Time/Gene file
