## Supplementals for "Gene Expression Changes Occurring at Bolting Time are Associated with Leaf Senescence in Arabidopsis": Supplemental_Figure_1.docx

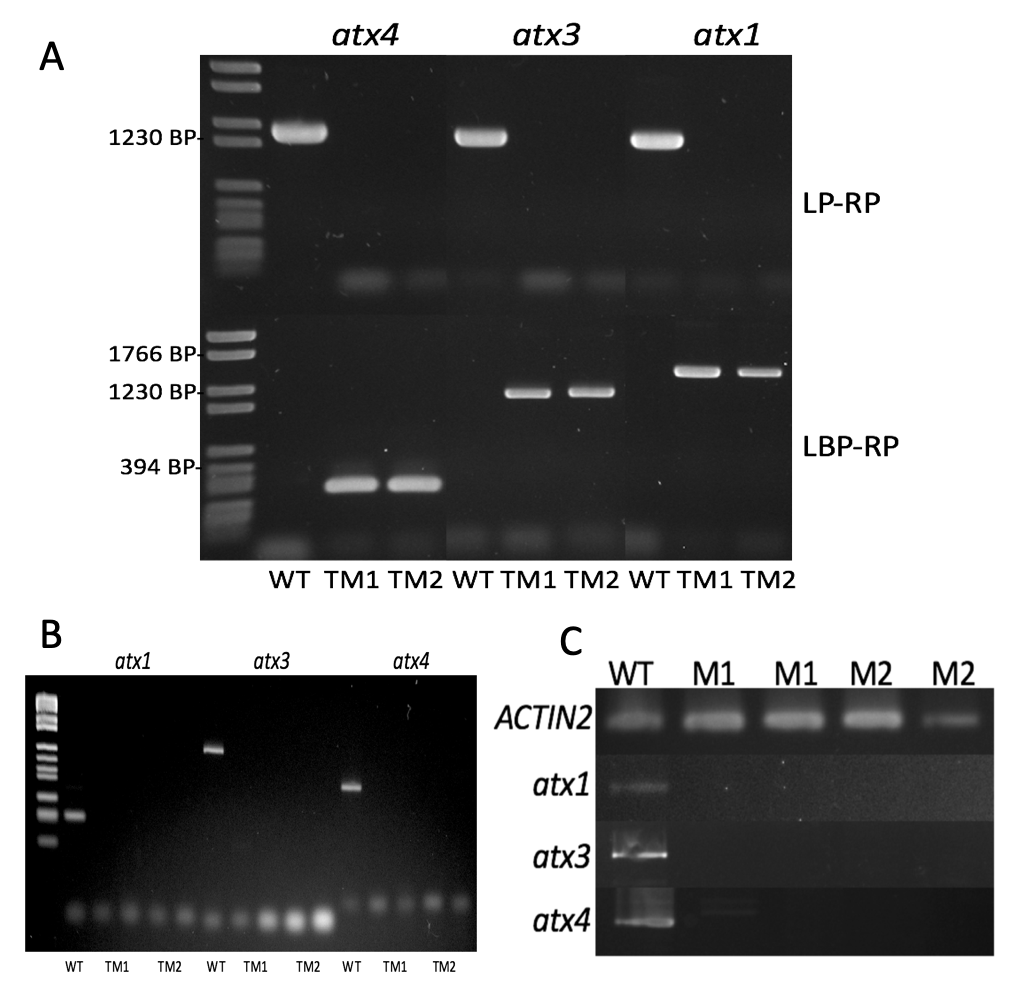
**Figure 1:  *atx* triple mutant genotype analysis.**

A: Genomic DNA was extracted and amplified with PCR using primers that flank the T-DNA insertion site of each allele. LP-RP amplified the wildtype allele while LBP (Left Border Primer) -RP amplified in the presence of a T-DNA insertion. B: This gel shows the product of PCR using cDNA template and exon specific primers that flank the T-DNA insertion of each gene. C: A second replicate was completed for the cDNA analysis on a different generation of plants to include a loading control (ACT2). These gels confirm the lack of full-length mRNA of two homozygous *atx* triple mutants from different F_2_ parents from which seeds were collected and treated as two separate early flowering mutants for further analysis.
